# Planar cell polarity pathway and development of the human visual cortex

**DOI:** 10.1101/404558

**Authors:** Jean Shin, Shaojie Ma, Edith Hofer, Yash Patel, Gennady V. Roshchupkin, André M. Sousa, Xueqiu Jian, Rebecca Gottesman, Thomas H. Mosley, Myriam Fornage, Yasaman Saba, Lukas Pirpamer, Reinhold Schmidt, Helena Schmidt, Amaia Carrion-Castillo, Fabrice Crivello, Bernard Mazoyer, Joshua C. Bis, Shuo Li, Qiong Yang, Michelle Luciano, Sherif Karama, Lindsay Lewis, Mark Bastin, Mathew A. Harris, Joanna M. Wardlaw, Ian E. Deary, Markus Scholz, Markus Loeffler, Veronica Witte, Frauke Beyer, Arno Villringer, Nicola J Armstrong, Karen A. Mather, David Ames, Jiyang Jiang, John B Kwok, Peter R. Schofield, Anbupalam Thalamuthu, Julian N. Trollor, Margaret J. Wright, Henry Brodaty, Wei Wen, Perminder S. Sachdev, Natalie Terzikhan, Tavia E. Evans, Hieab H.H.H. Adams, M. Arfan Ikram, Stefan Frenzel, Sandra van der Auwera-Palitschka, Katharina Wittfeld, Robin Bülow, Hans Jörgen Grabe, Christophe Tzourio, Aniket Mishra, Sophie Maingault, Stephanie Debette, Nathan A. Gillespie, Carol E. Franz, William S. Kremen, Linda Ding, Neda Jahanshad, the ENIGMA Consortium, Nenad Sestan, Zdenka Pausova, Sudha Seshadri, Tomas Paus, for the neuroCHARGE Working Group, Katrina L. Grasby, Neda Jahanshad, Jodie N. Painter, Lucía Colodro-Conde, Janita Bralten, Derrek P. Hibar, Penelope A. Lind, Fabrizio Pizzagalli, Christopher R.K. Ching, Mary Agnes B. McMahon, Natalia Shatokhina, Leo Zsembik, Ingrid Agartz, Saud Alhusaini, Marcio A.A. Almeida, Dag Alnæs, Inge K. Amlien, Micael Andersson, Tyler Ard, Nicola J. Armstrong, Allison Ashley-Koch, Manon Bernard, Rachel M. Brouwer, Elizabeth E.L. Buimer, Robin Bülow, Christian Bürger, Dara M. Cannon, Mallasr Chakravarty, Qiang Chen, Joshua W. Cheung, Baptiste Couvy-Duchesne, Anders M. Dale, Shareefa Dalvie, Tânia K. de Araujo, Greig I. de Zubicaray, Sonja M.C. de Zwarte, Anouk den Braber, Nhat Trung Doan, Katharina Dohm, Stefan Ehrlich, Hannah-Ruth Engelbrecht, Susanne Erk, Chun Chieh Fan, Iryna O. Fedko, Sonya F. Foley, Judith M. Ford, Masaki Fukunaga, Melanie E. Garrett, Tian Ge, Sudheer Giddaluru, Aaron L. Goldman, Nynke A. Groenewold, Dominik Grotegerd, Tiril P. Gurholt, Boris A. Gutman, Narelle K. Hansell, Mathew A. Harris, Marc B. Harrison, Courtney C. Haswell, Michael Hauser, Dirk J. Heslenfeld, David Hoehn, Laurena Holleran, Martine Hoogman, Jouke-Jan Hottenga, Masashi Ikeda, Deborah Janowitz, Iris E. Jansen, Tianye Jia, Christiane Jockwitz, Ryota Kanai, Sherif Karama, Dalia Kasperaviciute, Tobias Kaufmann, Sinead Kelly, Masataka Kikuchi, Marieke Klein, Michael Knapp, Annchen R. Knodt, Bernd Krämer, Thomas M. Lancaster, Phil H. Lee, Tristram A. Lett, Lindsay B. Lewis, Iscia Lopes-Cendes, Michelle Luciano, Fabio Macciardi, Andre F. Marquand, Samuel R. Mathias, Tracy R. Melzer, Yuri Milaneschi, Nazanin Mirza-Schreiber, Jose C.V. Moreira, Thomas W. Mühleisen, Bertram Müller-Myhsok, Pablo Najt, Soichiro Nakahara, Kwangsik Nho, Loes M. Olde Loohuis, Dimitri Papadopoulos Orfanos, John F. Pearson, Toni L. Pitcher, Benno Pütz, Anjanibhargavi Ragothaman, Faisal M. Rashid, Ronny Redlich, Céline S. Reinbold, Jonathan Repple, Geneviève Richard, Brandalyn C. Riedel, Shannon L. Risacher, Cristiane S. Rocha, Nina Roth Mota, Lauren Salminen, Arvin Saremi, Andrew J. Saykin, Fenja Schlag, Lianne Schmaal, Peter R. Schofield, Rodrigo Secolin, Chin Yang Shapland, Li Shen, Jean Shin, Elena Shumskaya, Ida E. Sønderby, Emma Sprooten, Lachlan T. Strike, Katherine E. Tansey, Alexander Teumer, Anbupalam Thalamuthu, Sophia I. Thomopoulos, Diana Tordesillas-Gutiérrez, Jessica A. Turner, Anne Uhlmann, Costanza Ludovica Vallerga, Dennis van der Meer, Marjolein M.J. van Donkelaar, Liza van Eijk, Theo G.M. van Erp, Neeltje E.M. van Haren, Daan van Rooij, Marie-José van Tol, Jan H. Veldink, Ellen Verhoef, Esther Walton, Yunpeng Wang, Joanna M. Wardlaw, Wei Wen, Lars T. Westlye, Christopher D. Whelan, Stephanie H. Witt, Katharina Wittfeld, Christiane Wolf, Thomas Wolfers, Clarissa L. Yasuda, Dario Zaremba, Zuo Zhang, Alyssa H. Zhu, Marcel P. Zwiers, Eric Artiges, Amelia A. Assareh, Rosa Ayesa-Arriola, Aysenil Belger, Christine L. Brandt, Gregory G. Brown, Sven Cichon, Joanne E. Curran, Gareth E. Davies, Franziska Degenhardt, Bruno Dietsche, Srdjan Djurovic, Colin P. Doherty, Ryan Espiritu, Daniel Garijo, Yolanda Gil, Penny A. Gowland, Robert C. Green, Alexander N. Häusler, Walter Heindel, Beng-Choon Ho, Wolfgang U. Hoffmann, Florian Holsboer, Georg Homuth, Norbert Hosten, Clifford R. Jack, MiHyun Jang, Andreas Jansen, Knut Kolskår, Sanne Koops, Axel Krug, Kelvin O. Lim, Jurjen J. Luykx, Daniel H. Mathalon, Karen A. Mather, Venkata S. Mattay, Sarah Matthews, Jaqueline Mayoral Van Son, Sarah C. McEwen, Ingrid Melle, Derek W. Morris, Bryon A. Mueller, Matthias Nauck, Jan E. Nordvik, Markus M. Nöthen, Daniel S. O’Leary, Nils Opel, Marie - Laure Paillère Martinot, G. Bruce Pike, Adrian Preda, Erin B. Quinlan, Varun Ratnakar, Simone Reppermund, Vidar M. Steen, Fábio R. Torres, Dick J. Veltman, James T. Voyvodic, Robert Whelan, Tonya White, Hidenaga Yamamori, Marina K.M. Alvim, David Ames, Tim J. Anderson, Ole A. Andreassen, Alejandro Arias-Vasquez, Mark E. Bastin, Bernhard T. Baune, John Blangero, Dorret I. Boomsma, Henry Brodaty, Han G. Brunner, Randy L. Buckner, Jan K. Buitelaar, Juan R. Bustillo, Wiepke Cahn, Vince Calhoun, Xavier Caseras, Svenja Caspers, Gianpiero L. Cavalleri, Fernando Cendes, Aiden Corvin, Benedicto Crespo-Facorro, John C. Dalrymple-Alford, Udo Dannlowski, Eco J.C. de Geus, Ian J. Deary, Norman Delanty, Chantal Depondt, Sylvane Desrivières, Gary Donohoe, Thomas Espeseth, Guillén Fernández, Simon E. Fisher, Herta Flor, Andreas J. Forstner, Clyde Francks, Barbara Franke, David C. Glahn, Randy L. Gollub, Hans J. Grabe, Oliver Gruber, Asta K. Håberg, Ahmad R. Hariri, Catharina A. Hartman, Ryota Hashimoto, Andreas Heinz, Manon H.J. Hillegers, Pieter J. Hoekstra, Avram J. Holmes, L. Elliot Hong, William D. Hopkins, Hilleke E. Hulshoff Pol, Terry L. Jernigan, Erik G. Jönsson, René S. Kahn, Martin A. Kennedy, Tilo T.J. Kircher, Peter Kochunov, John B.J. Kwok, Stephanie Le Hellard, Nicholas G. Martin, Jean - Luc Martinot, Colm McDonald, Katie L. McMahon, Andreas Meyer-Lindenberg, Rajendra A. Morey, Lars Nyberg, Jaap Oosterlaan, Roel A. Ophoff, Tomas Paus, Zdenka Pausova, Brenda W.J.H. Penninx, Tinca J.C. Polderman, Danielle Posthuma, Marcella Rietschel, Joshua L. Roffman, Laura M. Rowland, Perminder S. Sachdev, Philipp G. Sämann, Gunter Schumann, Kang Sim, Sanjay M. Sisodiya, Jordan W. Smoller, Iris E. Sommer, Beate St Pourcain, Dan J. Stein, Arthur W. Toga, Julian N. Trollor, Nic J.A. Van der Wee, Dennis van’t Ent, Henry Völzke, Henrik Walter, Bernd Weber, Daniel R. Weinberger, Margaret J. Wright, Juan Zhou, Jason L. Stein, Paul M. Thompson, Sarah E. Medland

## Abstract

The radial unit hypothesis provides a framework for global (proliferation) and regional (distribution) expansion of the primate cerebral cortex. Using principal component analysis (PCA), we have identified cortical regions with shared variance in their surface area and cortical thickness, respectively, segmented from magnetic resonance images obtained in 23,800 participants. We then carried out meta-analyses of genome-wide association studies of the first two principal components for each phenotype. For surface area (but not cortical thickness), we have detected strong associations between each of the components and single nucleotide polymorphisms in a number of gene loci. The first (global) component was associated mainly with loci on chromosome 17 (9.5e-32 ≤ *p* ≤ 2.8e-10), including those detected previously as linked with intracranial volume and/or general cognitive function. The second (regional) component captured shared variation in the surface area of the primary and adjacent secondary visual cortices and showed a robust association with polymorphisms in *a* locus on chromosome 14 containing *Disheveled Associated Activator of Morphogenesis 1* (*DAAM1*; *p*=2.4e-34). *DAAM1* is a key component in the planar-cell-polarity signaling pathway. In follow-up studies, we have focused on the latter finding and established that: (1) *DAAM1* is highly expressed between 12^th^ and 22^nd^ post-conception weeks in the human cerebral cortex; (2) genes co-expressed with *DAAM1* in the primary visual cortex are enriched in mitochondria-related pathways; and (3) volume of the lateral geniculate nucleus, which projects to regions of the visual cortex staining for cytochrome oxidase (a mitochondrial enzyme), correlates with the surface area of the visual cortex in major-allele homozygotes but not in carriers of the minor allele. Altogether, we speculate that, in concert with thalamocortical input to cortical subplate, *DAAM1* enables migration of neurons to cytochrome-oxidase rich regions of the visual cortex, and, in turn, facilitates regional expansion of this set of cortical regions during development.

---

The radial unit hypothesis provides a framework for global (proliferation) and regional (distribution) expansion of the primate cerebral cortex^1^. Using principal component analysis (PCA), we have identified cortical regions with shared variance in their surface area and cortical thickness, respectively, segmented from magnetic resonance images obtained in 23,800 participants. We then carried out meta-analyses of genome-wide association studies of the first two principal components for each phenotype. For surface area (but not cortical thickness), we have detected strong associations between each of the components and single nucleotide polymorphisms in a number of gene loci. The first (global) component was associated mainly with loci on chromosome 17 (9.5e-32 ≤ *p* ≤ 2.8e-10), including those detected previously as linked with intracranial volume^2,3^ and/or general cognitive function^4^. The second (regional) component captured shared variation in the surface area of the primary and adjacent secondary visual cortices and showed a robust association with polymorphisms in *a* locus on chromosome 14 containing *Disheveled Associated Activator of Morphogenesis 1* (*DAAM1*; *p*=2.4e-34). *DAAM1* is a key component in the planar-cell-polarity signaling pathway^5,6^. In follow-up studies, we have focused on the latter finding and established that: (1) *DAAM1* is highly expressed between 12^th^ and 22^nd^ post-conception weeks in the human cerebral cortex; (2) genes co-expressed with *DAAM1* in the primary visual cortex are enriched in mitochondria-related pathways; and (3) volume of the lateral geniculate nucleus, which projects to regions of the visual cortex staining for cytochrome oxidase (a mitochondrial enzyme), correlates with the surface area of the visual cortex in major-allele homozygotes but not in carriers of the minor allele. Altogether, we speculate that, in concert with thalamocortical input to cortical subplate, *DAAM1* enables migration of neurons to cytochrome-oxidase rich regions of the visual cortex, and, in turn, facilitates regional expansion of this set of cortical regions during development.

Using magnetic resonance imaging, one can derive a number of metrics informative with regard to development and aging of the human cerebral cortex, including cortical surface area and cortical thickness. The two measures provide insights into different developmental processes, each with a different timeline. <u>Cortical surface area</u> reflects primarily the tangential growth of the cerebral cortex during prenatal development; the phase of *symmetric division of progenitor cells* in the proliferative zones during the first trimester is particularly important for the tangential growth through additions of ontogenetic columns^1^. The subsequent phase of *asymmetric division* continues to increase the number of ontogenetic columns (and thus surface area) but it also begins to contribute to the <u>thickness of cerebral cortex</u> formed by post-mitotic neurons migrating from the proliferative zones to the cortical plate in the inside-out manner^1^. Ionizing radiation of the (monkey) fetus during early gestation reduces surface area (sparing cortical thickness) while the same radiation applied in midgestation affects both the surface area and cortical thickness^7^. While surface area remains stable after early childhood, cortical thickness continues to change, in particular during puberty and aging. Furthermore, both surface area and cortical thickness vary across individuals in global and regional manner.

Here we report findings obtained in 23,800 participants assessed across 19 cohorts from the CHARGE Consortium and the UK Biobank (Table E1 in Extended Data), and replicated in a subsequent release of the UK Biobank participants (n=6,234) as well as *in silico*^*8*^ using region-based summary statistics provided by the ENIGMA Consortium (n=19,512). To identify genetic loci associated with global and regional variations in each cortical phenotype, i.e., the surface area and thickness of the cerebral cortex, we have first carried out principal component analysis of regional values (34 regions segmented by FreeSurfer) in each cohort. For each phenotype, the first (PC1) and second (PC2) components loaded in similar sets of cortical regions across all 13 cohorts (Figures E1 [surface area] and E2 [cortical thickness] in Extended Data). Figure 1 illustrates the loadings for each of the 34 cortical regions in PC1 (Fig. 1A) and PC2 (Fig. 1B) for surface area. Note that PC2 includes only a handful of cortical regions in the medial aspect of the occipital lobe, including the pericalcarine (primary visual) cortex.

**Figure 1.**
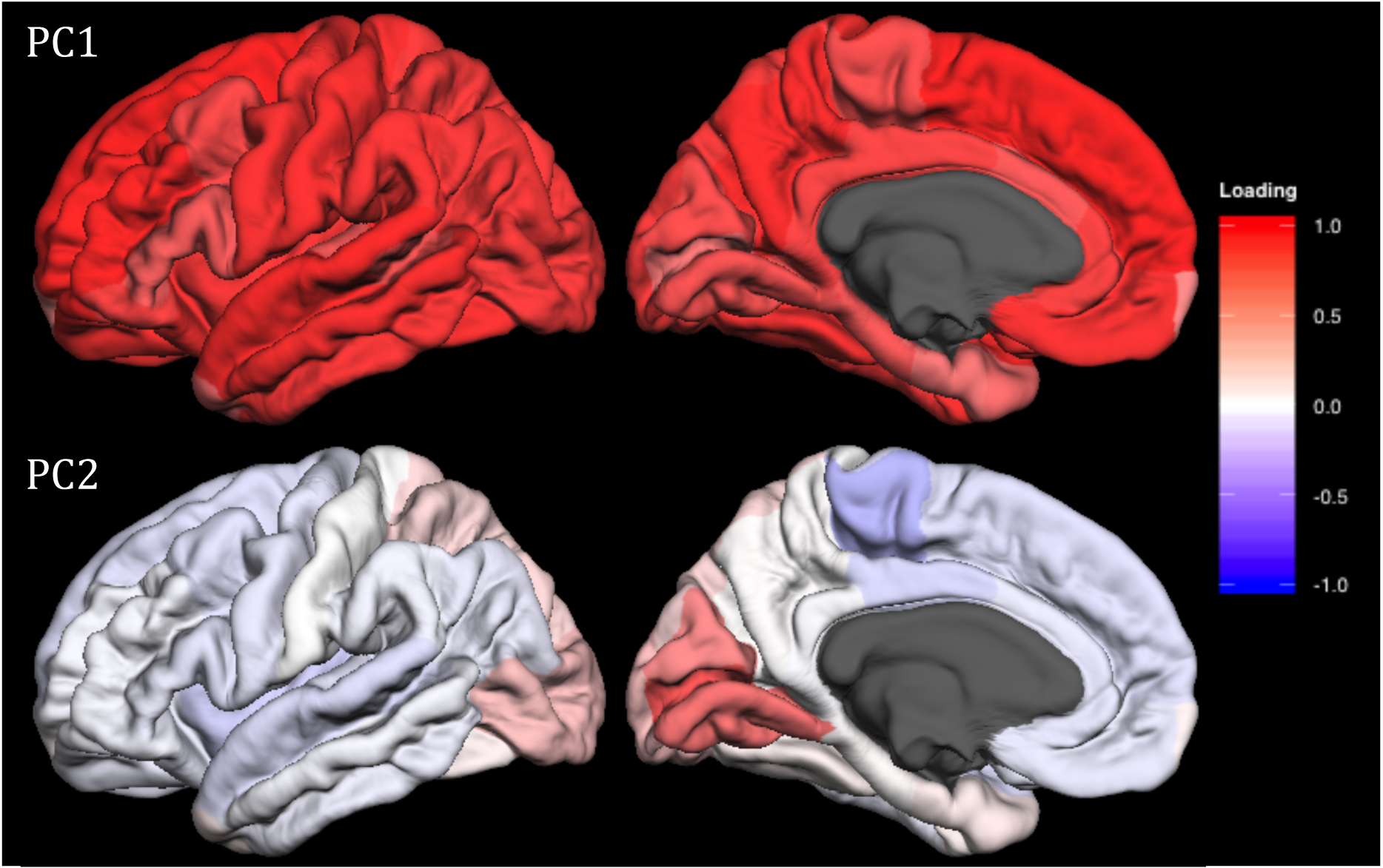
Lateral and medial views of median unrotated principal component (PC) loadings for the surface area of the 34 cortical regions in CHARGE consortium cohorts. Lateral (left column) and medial (right column) views of the median PC loadings are shown for PC1 (top) and PC2 (bottom). Each cohort estimated the surface area of the 34 cortical regions (left and right hemispheres summed), using FreeSurfer and carried out principal component analyses to obtain PC1 and PC2 loadings. Then, for each cortical region, median value of loadings was obtained across the cohorts. The red-to-blue color indicates the positive-to-negative loading values (i.e., correlation between PC scores and raw data) as indicated by the color bar. The median loading values were then used to derive the ‘general’ PC score for each individual and later used as the response variable in the GWAS meta-analyses.

We then executed a genome-wide association analysis (GWAS) in each of the 19 cohorts and, subsequently, meta-analyzed these cohort-based results (Supplementary Information) for each of the four phenotypes, namely PC1 and PC2 of surface area, and PC1 and PC2 of cortical thickness. For surface area, the first (“global”) component was associated mainly with a number of loci on chromosome 17, including those detected previously as linked with intracranial volume^2,3^ [and/or general cognitive function^4^] (Figure E3A and Table E2A in Extended Data). The second (“regional”) component was associated mainly with a locus on chromosome 14 containing *DAAM1* (Figure 2, Figures E3B and E4, and Table E2B in Extended Data). For cortical thickness, meta-GWAS revealed a single locus associated with PC1 and a single locus associated with PC2 (Figure E5). The *<u>DAAM1</u>* locus (top hit: rs73313052) was associated with surface area (but not cortical thickness) of each of the four cortical regions loading on PC2 when examined on a region-by-region basis, as reported in two independent reports from the CHARGE and ENIGMA Consortia (Table E3 in Extended Data).

**Figure 2.**
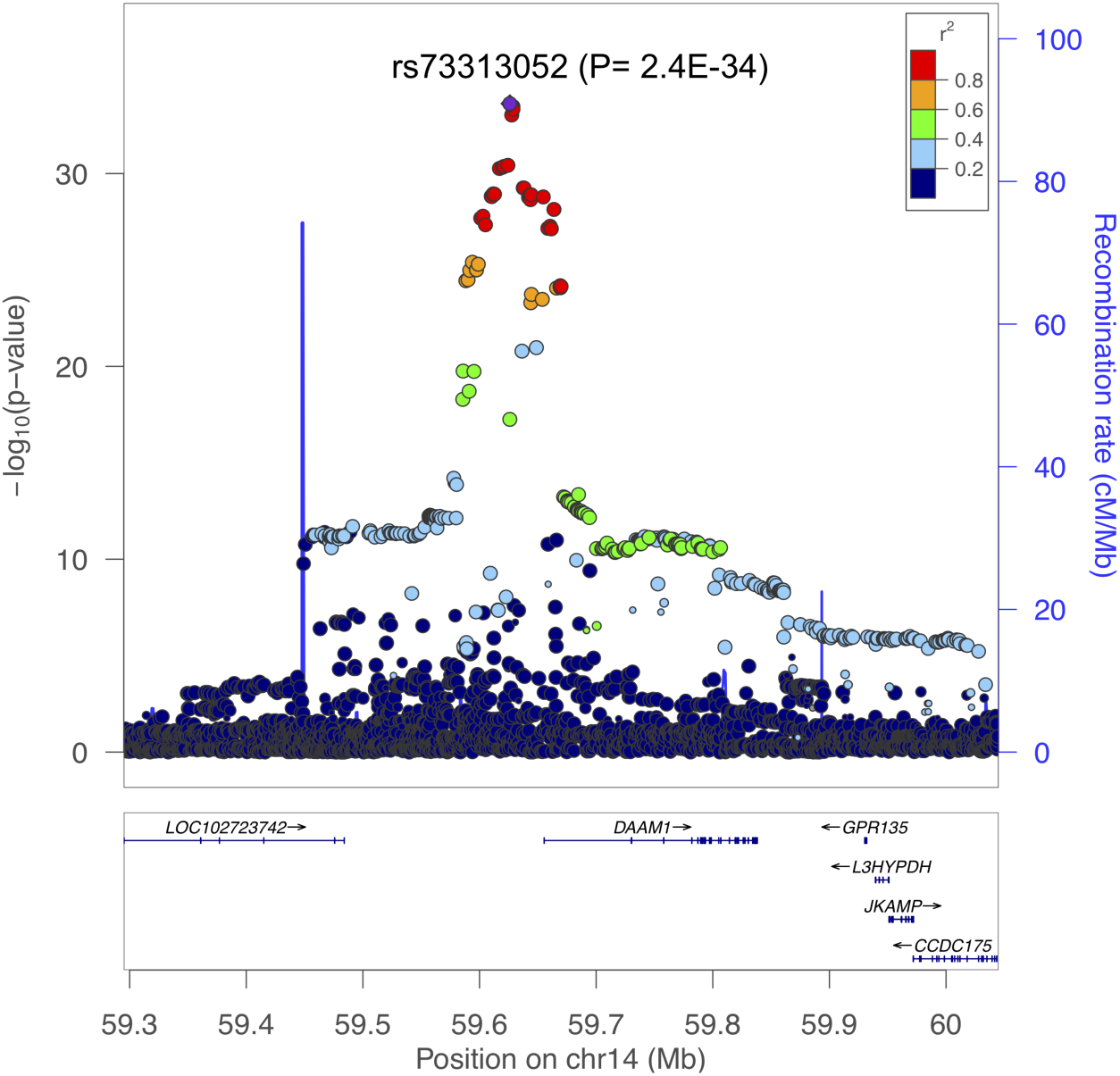
Regional association plot for rs73313052, the top PC2 single nucleotide polymorphism (SNP). Each point indicates a SNP tested in the meta genome-wide association study (GWAS) of surface area PC2 within the shown genomic region. The top PC2 SNP is indicated by the purple diamond. The horizontal axis represents the genomic position on the human chromosome 14 (hg19). The left-vertical axis indicates the –log_10_*P* values obtained from the GWAS meta-analysis (on the left side); and the right-vertical axis, the estimated recombination rate from the HapMap samples. The red-to-blue colors indicate the degree of linkage disequilibrium (LD) between each SNP and rs73313052. The LD was based on the pairwise squared allelic correlation *r*^2^ estimated in the 1000 Genomes European reference panels (Nov 2014 EUR). The plot was created using LocusZoom (http://locuszoom.org/).

*DAAM1* is a key component of the planar-cell-polarity signaling pathway^5,6^; it acts as a bridging factor between Disheveled, Rho-family GTPases and Rho-associated kinases^*9*^, a molecular complex involved in organizing actin cytoskeleton^10^.

In order to gain insights into possible mechanisms by which *DAAM1* contributes to the tangential expansion of the human visual cortex, we have carried out a number of follow-up studies. **First**, we examined *DAAM1* expression in the human brain using the BrainSpan dataset (Table E4 in Extended Data). As shown in Figure 3, *DAAM1* is expressed in the cerebral cortex between ∼80 and ∼150 post-conception days; after birth, its expression is very low. Note that, in monkeys, neurogenesis of the primary visual cortex begins around embryonic day 40 (E40) and ends at E100 (165-day gestation)^1^.

**Figure 3.**
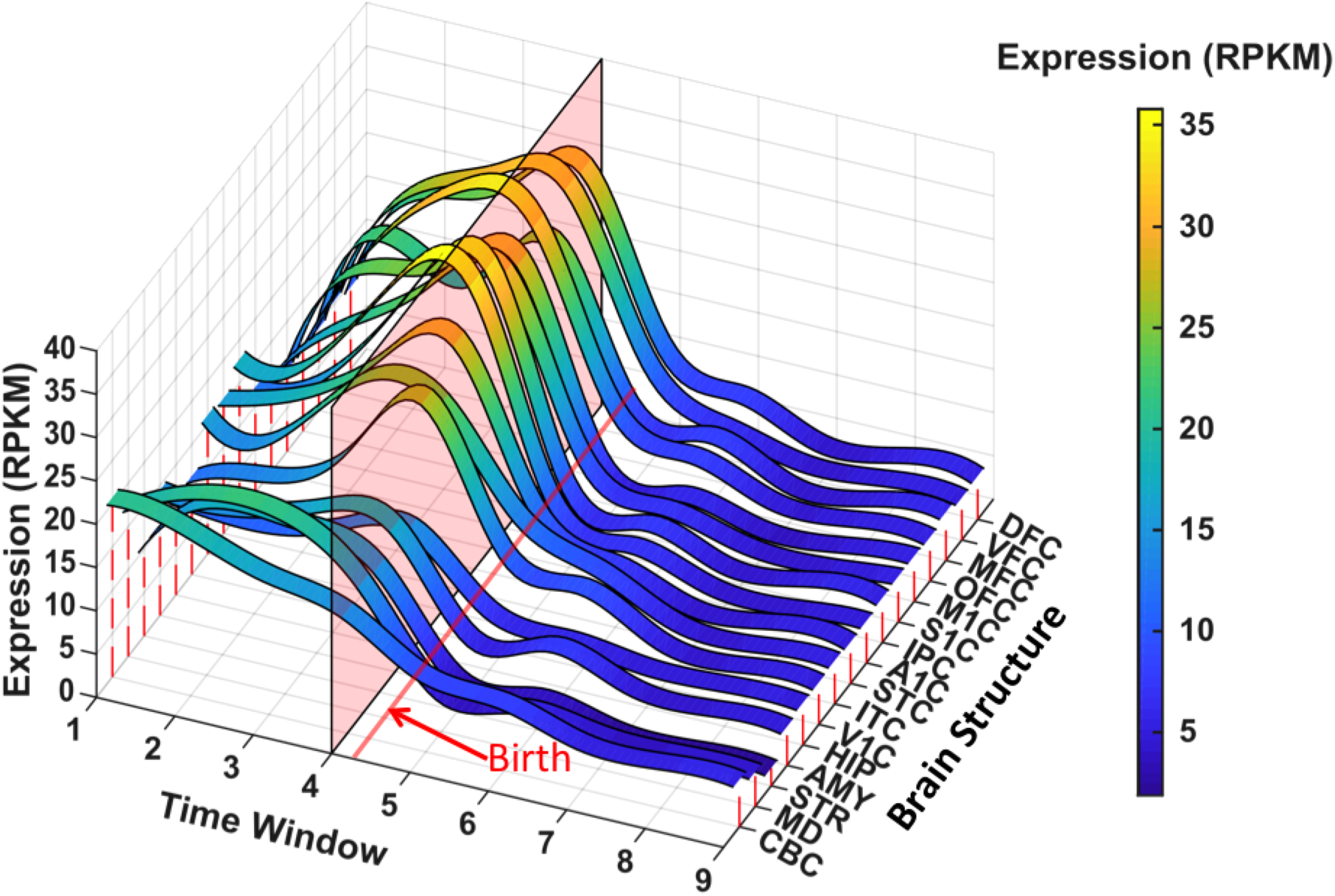
Spatial and temporal expression of *DAAM1* on BrainSpan mRNA-seq data. The mRNA expression levels were measured by RNA sequencing in 607 brain tissues from 18 female and 23 male donors available in BrainSpan database (http://www.brainspan.org/). For a given brain structure, *DAAM1* expression value was averaged and then fitted across differentiation windows using spline function implemented in MATLAB to get smooth *DAAM1* expression dynamics. Each colored band represents the fitted expression levels in RPKM (reads per kilobase per million) of *DAAM1*. Brain structure includes 11 cortical and 5 sub-cortical regions: dorsolateral prefrontal cortex (DFC), ventrolateral prefrontal cortex (VFC), medial frontal cortex (MFC), orbitofrontal cortex (OFC), primary motor cortex (M1C), primary sensory cortex (S1C), inferior parietal cortex (IPC), primary auditory cortex (A1C), superior temporal cortex (STC), inferior temporal cortex (ITC), primary visual cortex (V1C), hippocampus (HIP), amygdala (AMY), striatum (STR), mediodorsal nucleus of thalamus (MD) and cerebellar cortex (CBC). The human brain differentiation was split into 9 windows based on post conception days: 52-69 (Window 1), 70-111 (Window 2), 112-132 (Window 3), 133-167 (Window 4), 168-447 (Window5), 448 - 1299 (Window 6), 1300-4648 (Window 7), 4649-7570 (Window 8), and 7571-14876 (Window 8). The red line indicates the boundary between pre-and postnatal periods. The pink panel indicates the time where *DAAM1* has peak expression in V1C. Blue-to-yellow colors represent low-to-high expression levels of *DAAM1* as indicated in the color bar.

**Second**, we examined co-expression of *DAAM1* across all cortical regions and prenatal time points using the same BrainSpan dataset. As expression of *DAAM1* increases and decreases, so does expression of genes enriched in pathways involving neuron migration and cytoskeleton organization, among others (Table E5A in Extended Data). On the other hand, expression of a large number of genes varies in the direction opposite to that of *DAAM1*, including genes enriched in pathways involved in a number of metabolic processes (Table E5B in Extended Data). To ascertain the pattern of *DAAM1* co-expression *specific* to the primary visual cortex (V1), we have identified genes co-expressed highly (top 1%) in V1 but not in any other cortical region (i.e., not present among top 1% in any of the other eight regions). This analysis yielded striking enrichment for mitochondria-related genes co-expressed strongly in the same direction as *DAAM1* in V1 but not in the other cortical regions (Table E6 in Extended Data). This observation turned our attention to the well-known parcellation of the visual cortex to cytochrome-oxidase rich sub-regions, so-called “blobs” (V1) and “stripes” (V2/V3)^11^. We then examined co-localization of *DAAM1* and a mitochondrial marker ATP5A in the developing (22^nd^ post-conception week) visual cortex (Figure E6); its co-localization is consistent with the co-expression analyses described above. It is known that post-mitotic neurons migrate, along radial glia, from proliferative (ventricular and subventricular) zones to subplate zones, which contain afferents from the thalamic radiation^1^. Activity carried by these afferents from the retina is critical for the development of the primate visual cortex; enucleation of the eyes during fetal development results in the reduction of the surface area (but not thickness) of the monkey primary visual cortex^12,13^, as well as in a reduced size of the lateral geniculate nucleus (LGN)^13^. A tight relationship exists between the volume of LGN and surface area of the primary visual cortex in the human (adult) brain^14^. Furthermore, the koniocellular portion of LGN (which carries signals from short wavelength [blue] cones) appears to project specifically to cytochrome-oxidase rich areas of the visual cortex^15^. Therefore, we hypothesized that DAAM1 contributes to the migration of cytochrome-oxidase positive neurons in response to the LGN inputs in the subplate zones during fetal development. **Third**, to test this hypothesis in our data, we predicted that the expected relationship between the LGN volume (a proxy for retinal inputs during fetal development) and V1 surface area will be present only in *DAAM1* (rs73313052) major-allele homozygotes (GG) but not in the carriers of minor allele (GA or AA). This prediction was confirmed: an interaction between rs73313052 genotype and LGN volume vis-à-vis PC2 magnitude was significant when examined in a cohort with available LGN volumes (GG: r=0.13, p=0.0006, n=694; A carriers: r=-0.06, p=0.37, n=206; interaction: t ratio= −2.47, p=0.014, n=925; Figure E7 in Extended Data).

In summary, we discovered a non-overlapping set of 99 ‘independent’ single nucleotide polymorphisms within 35 genomic loci contributing to the global and regional tangential growth of the human cerebral cortex (Table E7, in Extended Data). On the other hand, our meta-GWAS of cortical thickness, carried out in the same individuals, yielded only two loci. This negative finding is consistent with a very low number of loci associated with both global and regional values of cortical thickness reported in the two reports from the CHARGE^16^ and ENIGMA^17^ Consortia; it may reflect substantial dynamics of cortical thickness during puberty^18^ and aging^19^. For surface area PC1 (PC2), we replicated 695/ 807 (PC1) and 952/1,155 (PC2) GWAS-significant SNPs (Table E2A and E2B, in Extended Data). Through a series of follow-up studies, we formulated a working model by which *DAAM1* regulates tangential expansion of the visual cortex by interacting with LGN inputs, likely at the level of cortical subplate, during mid-gestation. A larger visual cortex is likely to posses more inter-hemispheric connections; *DAAM1* polymorphism is associated with a structure-predicted functional connectivity of the human visual cortex^20^. Overall, these findings illustrate how specification of cortical areas, and their relative growth, might be guided by an interaction between fetal environment and generic developmental mechanisms, such as those constituting planar-cell-polarity signaling pathway.

## Acknowledgements

This research has been conducted using the UK Biobank Resource under Application Number ‘23509’.

## References

1. Rakic P. Specification of cerebral cortical areas. Science. Jul 8 1988;241(4862):170–176.

2. Adams HH, Hibar DP, Chouraki V, et al. Novel genetic loci underlying human intracranial volume identified through genome-wide association. Nat Neurosci. Oct 03 2016.

3. Hibar DP, Stein JL, Renteria ME, et al. Common genetic variants influence human subcortical brain structures. Nature. Apr 9 2015;520(7546):224–229.

4. Trampush JW, Yang ML, Yu J, et al. GWAS meta-analysis reveals novel loci and genetic correlates for general cognitive function: a report from the COGENT consortium. Mol Psychiatry. Mar 2017;22(3):336–345.

5. Tissir F, Goffinet AM. Planar cell polarity signaling in neural development. Curr Opin Neurobiol. Oct 2010;20(5):572–577.

6. Beane WS, Tseng AS, Morokuma J, Lemire JM, Levin M. Inhibition of planar cell polarity extends neural growth during regeneration, homeostasis, and development. Stem cells and development. Aug 10 2012;21(12):2085–2094.

7. Selemon LD, Ceritoglu C, Ratnanather JT, et al. Distinct abnormalities of the primate prefrontal cortex caused by ionizing radiation in early or midgestation. J Comp Neurol. Apr 1 2013;521(5):1040–1053.

8. Nieuwboer HA, Pool R, Dolan CV, Boomsma DI, Nivard MG. GWIS: Genome-Wide Inferred Statistics for Functions of Multiple Phenotypes. Am J Hum Genet. Oct 6 2016;99(4):917–927.

9. Habas R, Kato Y, He X. Wnt/Frizzled activation of Rho regulates vertebrate gastrulation and requires a novel Formin homology protein Daam1. Cell. Dec 28 2001;107(7):843–854.

10. Yang Y, Mlodzik M. Wnt-Frizzled/planar cell polarity signaling: cellular orientation by facing the wind (Wnt). Annual review of cell and developmental biology. 2015;31:623–646.

11. Livingstone MS, Hubel DH. Thalamic inputs to cytochrome oxidase-rich regions in monkey visual cortex. Proc Natl Acad Sci U S A. Oct 1982;79(19):6098–6101.

12. Bourgeois JP, Rakic P. Synaptogenesis in the occipital cortex of macaque monkey devoid of retinal input from early embryonic stages. Eur J Neurosci. May 1996;8(5):942–950.

13. Dehay C, Giroud P, Berland M, Killackey H, Kennedy H. Contribution of thalamic input to the specification of cytoarchitectonic cortical fields in the primate: effects of bilateral enucleation in the fetal monkey on the boundaries, dimensions, and gyrification of striate and extrastriate cortex. J Comp Neurol. Mar 25 1996;367(1):70–89.

14. Andrews TJ, Halpern SD, Purves D. Correlated size variations in human visual cortex, lateral geniculate nucleus, and optic tract. J Neurosci. Apr 15 1997;17(8):2859–2868.

15. Hendry SH, Reid RC. The koniocellular pathway in primate vision. Annu Rev Neurosci. 2000;23:127–153.

16. Hofer Eac. Genetic Determinants of Cortical Structure (Thickness, Surface Area and Volumes) among Disease Free Adults in the CHARGE Consortium. BioRxiv 2018.

17. Grasby Kac. The genetic architecture of the human cerebral cortex. BioRxiv 2018.

18. Walhovd KB, Fjell AM, Giedd J, Dale AM, Brown TT. Through Thick and Thin: a Need to Reconcile Contradictory Results on Trajectories in Human Cortical Development. Cereb Cortex. Feb 01 2017;27(2):1472–1481.

19. Vinke EJ, de Groot M, Venkatraghavan V, et al. Trajectories of imaging markers in brain aging: the Rotterdam Study. Neurobiol Aging. Jul 17 2018;71:32–40.

20. Mollink J, Smith SM, Elliott LT, et al. The spatial correspondence and genetic influence of interhemispheric connectivity with white matter microstructure. Nat Neurosci. Apr 15 2019.

