## Supplemental Tables for "Planar cell polarity pathway and development of the human visual cortex"

**Extended Data Tables**

| | |
|---|---|
| Table E1A | Demograpic information for the cohorts included in the meta-analyses of genome-wide association studies of surface area and cortical thickness principal component scores. |
| Table E1B | Summary of brain imaging measures and acquisition parameters per cohort included in meta analyses of genome-wide association studies of surface area and cortical thickness principal component scores |
| Table E1C | Genotyping, quality control, imputation and GWAS analysis software |
| Table E2A | GWAS meta-analysis results for surface area PC1 for SNPs with P-values< 5E-08 |
| Table E2B | GWAS meta-analysis results for surface area PC2 for SNPs with P-values< 5E-08 |
| Table E3 | Associations between rs73313052 and surface area and thickness in each of the 34 cortical regions estimated from CHARGE- and ENIGMA-meta analyses |
| Table E4 | *DAAM1* mRNA expression levels measured in different brain structures from the donors at different ages |
| Table E5A | Over- or under-represented Gene Ontology biological process terms for the genes positively coexpressed with *DAAM1* across cortical regions in prenatal brains |
| Table E5B | Over- or under-represented Gene Ontology biological process terms for the genes negatively coexpressed with *DAAM1* across cortical regions in prenatal brain |
| Table E6 | Over- or under-represented Gene Ontology biological process terms for the genes positively coexpressed with *DAAM1* exclusively in the primary visual cortex (VIC) |
| Table E7 | Genomic loci with independent significant SNPs identified in meta analyses of GWAS of surface area-PC1 and -PC2, obtained by FUMA |

**Table E1A. Demograpic information for the cohorts included in the meta-analyses of genome-wide association studies of surface area and cortical thickness principal component scores.**

| Cohort | Cohort Abbreviation | Study Design | Ancestry | Total N | Females N (%) | Age Range (years) |
|---|---|---|---|---|---|---|
| **Atherosclerosis Risk in Communities Study** | **ARIC** | Population-based | European | 1,080 | 451 (42%) | 45 - 64 |
| **Austrian Stroke Prevention Family Study** | **ASPSFam** | Family-based | European | 229 | 162 (71%) | 38 - 86 |
| **BIL&GIN** | **BIL&GIN** | Population-based | European | 262 | 127 (48%) | 18 - 57 |
| **Cardiovascular Health Study** | **CHS** | Population-based | European | 567 | 226 (40%) | 73 - 95 |
| **Framingham Heart Study** | **FHS** | Population-based | European | 946 | 404 (43%) | 34 - 85 |
| **LIFE-Adult** | **LIFE-Adult** | Population-based | European | 1,690 | 850 (50%) | 20 - 82 |
| **Lothian Birth Cohort 1936** | **LBC1936** | Population based | European | 568 | 267 (47%) | 70 - 74 |
| **Sydney Memory and Ageing Study** | **MAS** | Population-based | European | 376 | 236 (63%) | 70 - 90 |
| **Older Australian Twin Study** | **OATS** | Population based | European | 280 | 203 (73%) | 65 - 89 |
| **Rotterdam Study 1** | **RS1** | Population-based | European | 1,073 | 459 (43%) | 69 - 96 |
| **Rotterdam Study 2** | **RS2** | Population-based | European | 1,148 | 551 (48%) | 60 - 97 |
| **Rotterdam Study 3** | **RS3** | Population-based | European | 2,551 | 1,142 (45%) | 45 - 89 |
| **Saguenay Youth Study - Adult** | **SYS-Adult** | Family-based | European | 515 | 241 (47%) | 36 - 65 |
| **Saguenay Youth Study - Adolescent** | **SYS-Adolescent** | Family-based | European | 947 | 458 (48%) | 12 - 18 |
| **Study of Health in Pomerania - 2** | **SHIP-2** | Population based | European | 1,073 | 557 (52%) | 30 - 89 |
| **Study of Health in Pomerania - Trend1** | **SHIP-TREND1** | Population based | European | 857 | 482 (56%) | 22 - 81 |
| **Study of Health in Pomerania - Trend2** | **SHIP-TREND2** | Population based | European | 1,076 | 529 (49%) | 21 - 82 |
| **Three Cities-Dijon** | **3CDijon** | Population-based | European | 383 | 225 (58%) | 65 - 83 |
| **UK Biobank** | **UKBB** | Population-based | European | 8,163 | 3,806 (47%) | 48 - 63 |

| Table E1B. Summary of brain imaging measures and acquisition parameters per cohort included in meta analyses of genome-wide association studies of surface area and cortical thickness principal component scores. | | | | |
|---|---|---|---|---|
| **Cohort** | **Scanner(s)** | **Software** | **Citation(s) PMID** | **Acquistion Information** |
| **ARIC** | 1.5 T MRI scanners (General Electric Medical Systems) | FSL FIRST | | Total intracranial volume was measured from T1 weighted spin echo sagittal images, each set consisting of 32 contiguous 5 mm thick interleaved sections with no interslice gap, a field of view of 24 cm, and a matrix of 256 × 192, obtained with the following sequence: scan time, 2.5 minutes; echo time, 14 milliseconds; 2 repetitions; and repetition time, 500 milliseconds. A 3D volumetric spoiled gradient echo sequence with TR = 27 msec, TE = 9 msec, 124 contiguous partitions, 1.6 mm slice thickness, a 22 × 16.5 cm field of view, 192 views, and 45° flip angle was used to measure the volumes of the hippocampus and temporal horn.<br>Interactive image processing steps were performed at the Mayo Clinic by a research associate who had no knowledge of the subjects' personal or medical histories |
| **ASPSFam** | MAGNETOM Trio, A Tim System, Siemens - Erlangen, Germany (3.0T) | Freesurfer v5.1 | | 3D-MPRAGE |
| **BIL&GIN** | 3.0 T Tesla Philips ACHIVA scan ner (Philips Medical Systems, Best, The Netherlands) | FreeSurfer (5.3) | 25840118 | The acquisition protocol included a high resolution 3D T1-weighted sequence (3D-FFE-TFE; TR = 20 ms; TE = 4.6 ms; flip angle = 10°; inversion time = 800 ms; turbo field echo factor = 65; sense factor = 2; matrix size = 256 × 256 × 180 mm3; 1mm3 isotropic voxel size) |
| **CHS** | 1.5 T MRI | FSL FIRST | 21943959; 7847205; 8303738 | MRI scanning was completed at each of the four sites using 1.5 Tesla scanners. The scanning protocol included a 3-D volumetric T1 weighted Spoiled Gradient Recall (SPGR) sequence covering the whole brain (TE/TR = 5/25, flip angle = 40°, NEX = 1, slice thickness = 1.5 mm/0 mm interslice gap), with an in-plane acquisition matrix of 256 × 256 × 124 image elements, 250 × 250 mm field of view and an in-plane voxel size of 0.98 mm3. |
| **FHS** | 1 or 1.5 T Siemens Magnetom | FreeSurfer (5.3) | 15653178 | Subjects were imaged on a Siemens Magnetom(Munich, Germany), using a T2-weighted double spin-echo coronal imaging sequence of 4mm contiguous slices from nasion to occiput with a repetition time (TR) of 2420 ms, echo time (TE) of TE1 20/TE2 90 ms; echo train length 8 ms; field of view (FOV) 22 cm and an acquisition matrix of 182 × 256 interpolated to a 256 × 256 with one excitation. |
| **LBC1936** | GE Signa Horizon HDx 1.5 T | Freesurfer 5.3.0 | 14598854 | fast 3-dimensional spoiled gradient-echo T1-weighted axial acquisition, with 2 excitations; 124 slices (1 mm thick) |
| **LIFE** | 3T Siemens Verio scanner equipped with a 32 channel head coil | FreeSurfer (5.3) | 26973099; 28397392 | High resolution T1-weighted structural images were acquired using an MP-RAGE sequence with 1 mm isotropic voxels, 176 slices, TR=2300 ms, TE=2.98 ms, and inversion time (TI)=900 ms. |
| **MAS** | 3T | Freesurfer 5.3 | 20637138 | The 3D T1-weighted MRIs scans were used for computing the neuroimaging phenotypes and FreeSurfer v5.3 was used in the computation. T1-weighted scans had in-plane resoultion of 1 x 1 mm^2 and slice thickness = 1 mm with no gap in between, yielding 1 × 1 × 1 mm^3 isotropic voxels. |
| **OATS** | 1.5T and 3T | Freesurfer 5.3 | | The 3D T1-weighted MRIs scans were used for computing the neuroimaging phenotypes. The sequence was performed using a similar protocol for the 1.5 Tesla scanners in the three centres with in-plane resolution = 1 × 1 mm, slice thickness = 1.5 mm, slice number = 144. The acquisition parameters for the 3 Tesla Philips scanner in centre 1 in-plane resolution = 1 × 1 mm, slice thickness = 1 mm, slice number = 190, resulting isotropic voxels of 1 × 1 × 1 mm^3. |
| **RS2,RS2,RS3** | 1.5 T MRI unit (GE Healthcare, Milwaukee, USA, software version 11×) | FreeSurfer (4.5) | 22002080 | Structural imaging is performed with T1-weighted (T1w), proton density-weighted (PDw) and fluid-attenuated inversion recovery (FLAIR) sequences. The combination of different MR contrasts provided by these sequences can be used for automated brain tissue and white matter lesion segmentation (see section on processing). For this purpose, the T1w scan is acquired in 3D at high in-plane resolution and with thin slices (voxel size <1 mm3). |
| **SHIP** | Magnetom Avanto, Siemens Medical Systems, Erlangen, Germany, 1.5 Tesla | Freesurfer 5.3 | 20167617 | T1-weighted magnetization prepared rapid acquisition gradient echo (MPRAGE) sequence |
| **SHIP-Trend** | Magnetom Avanto, Siemens Medical Systems, Erlangen, Germany, 1.5 Tesla | Freesurfer 5.3 | 20167617 | T1-weighted magnetization prepared rapid acquisition gradient echo (MPRAGE) sequence |
| **SYS-Adult** | 1.5 T Siemens (Avanto) scanner | FreeSurfer (5.1.0) | 25454417 | T1W, 1-mm, isotropic resolution images acquired with a 3D fast radio frequency (RF)-spoiled gradient-echo scan. |
| **SYS-Adolescent** | 1.0 T Philips | FreeSurfer (5.1.0) | 17469173 | Three-dimensional (3D) radio frequency (RF)-spoiled gradient-echo scan with 140 –160 slices, an isotropic resolution of 1 mm, a repetition time (TR) of 25 ms, an echo time (TE) of 5 ms, and flip angle of 30° |
| **3CDijon** | SIEMENS Magnetom (1.5T) | Freesurfer 5.3.0 | 14598854, 17977521 | 3D IR-FSPGR (T1); 2D FSE |
| **UKBB** | 3.0 T Siemens Skyra running VD13A SP4, with a standard Siemens 32-channel RF receive head coil | FreeSurfer | 27643430 | T1W, acquired in the sagittal plane using a 3D magnetization-prepared rapid gradient-echo sequence at a resolution of 1×1×1 mm, with a 208×256×256 field of view |

| Table E1C. Genotyping, quality control, imputation and GWAS analysis software | | | | | |
|---|---|---|---|---|---|
| Cohort | Genotyping Platform | Sample and variant quality control | Phasing/Imputation software | Reference Panel | Association Software (version) |
| ARIC | Affymetrix SNP Array 6.0 | sample callrate ≥ 95%, MAF > 0.01 & Rsq > 0.3, 0.005 < MAF <= 0.01 & Rsq > 0.8 | SHAPEIT (v1.r532)/IMPUTE2 (v2.2.2) | 1000 Genomes phase 1 version 3 | ProbABEL |
| ASPS | Illumina Human610-Quad BeadChip | Sample call rate 98%, SNP call rate 98%, MAF 0.01, HWE-pvalue 1×10-6, Other sample exclusions: Non-European ancestry, sample failures, sex mismatch, high autosomal heterozygosity, cryptic relatedness | SHAPEIT (v1.r790)/IMPUTE2 (v2.3.1) | 1000 Genomes ALL phase 1 version 3 | Plink |
| BIL&GIN | Illumina InfiniumOmniExpressExome-8v1-4 | MAF 1%, SNP call rate 95%, sample call rate 95%, HWE p-value $10^{-6}$, sample failures, sex mismatch, cryptic relatedness, ancestry outliers (smartPCA) | mach (v1.0.18)/Minimac3 (v2.0.1) | 1000G EUR v3 (http://enigma.ini.usc.edu/wp-content/uploads/2012/07/v3.20101123.ENIGMA2.EUR.20120719.vcf.tgz ) | Plink (v1.90b3w) |
| CHS-EA | Illumina 370 CNV, ITMAT-Broad-CARe (IBC) | For SNPs, the following exclusions were applied : Call rate < 97%, HWE P < 10-5, > 2 duplicate errors or Mendelian inconsistencies (for reference CEPH trios), heterozygote frequency = 0, SNP not found in HapMap. | MaCH + minimac (release stamp 2012-11-16) | 1000 Genomes phase 1 version 3 | R |
| FHS | Affymetrix GeneChip Human Mapping 500K Array Set® and 50K Human Gene Focused Panel® | Sample Level QC: sample call rate ≥ 97%, sample failures, genotyped sex different from recorded sex, discordance based on pedigree analyses; SNP level QC: SNP call rate ≥ 97%, a deviation from Hardy-Weinberg (HWE) ≥ 1x10-6, Mendel errors ≤ 1000, and MAF ≥ 1%, on chromosome 1-22, or X with a physical location on Build 37. | MaCH/minimac | 1000 Genomes phase 1 version 3 | GWAF |
| LBC1936 | Illumina Human 610_Quadv1 | MAF>1%, SNP call rate >0.98, sample call rate >0.95, HWE p-value $10^{-3}$, Non-European ancestry not included | minimac/minimac | 1000G Phase I Version III with All ethnicities (2010-11 data freeze, 2012-03-14 haplotypes) | HASE |
| LIFE-Adult | Affymetrix Axiom CEU1 SNP array | Sample quality filters were dish-QC<0.82, call rate<97%, heterozygosity outliers, sex-mismatch, cryptic relatedness and outliers of principle components analysis (>6SD of first 10 PCs). SNP quality filters were call rate<97%, cluster plot irregularities as recommended by Affymetrix, p-value of exact Hardy-Weinberg equilibrium test <1e-6 and p-value of plate association <1e-7. | SHAPEIT (v2.r790)/ IMPUTE2 (v2.3.2) | 1kG phase 1, version 3 (2012) | PLINK 1.9 |
| MAS | Affymetrix Genome-wide Human SNP Array 6.0 | MAF 1%, HWE p-value $10^{-6}$, SNP call rate 98%, sample call rate 95%, Sex, relatedness and assesment of population stratification | MaCH/minimac | 1000 Genomes phase 1 version 3 | mach2qtl |
| OATS | Illumina Omniexpress | MAF 1%, HWE p-value $10^{-6}$, SNP call rate 98%, sample call rate 95%, Sex, relatedness and assesment of population stratification | MaCH/minimac | 1000 Genomes phase 1 version 3 | merlin |
| RS1 | Illumina 550 (+duo), Illumina 610 quad | SNP call rate 98%, MAF 0.01, HWE p-value $10^{-6}$ | MaCH/minimac | 1000 Genomes phase 1 version 3 | HASE |
| RS2 | Illumina 550 duo | SNP call rate 98%, MAF 0.01, HWE p-value $10^{-6}$ | MaCH/minimac | 1000 Genomes phase 1 version 3 | HASE |
| RS3 | Illumina 610 quad | SNP call rate 98%, MAF 0.01, HWE p-value $10^{-6}$ | MaCH/minimac | 1000 Genomes phase 1 version 3 | HASE |
| SYS | Illumina Human610W-Quad Beadchip (Quad), Illumina HumanOmniExpress BeadChip (Omni) | MAF>1%, call rate >0.95 per chip, HWE p-value <10e-6 | SHAPEIT/IMPUTE2 | 1000 Genomes release March 2012 (EUR reference panel) | GenABEL/ProbABEL |
| 3CDijon | Illumina Human 610 Quad BeadChip® | SNP call rate 95%, sample call rate 98%, HWE p-value $10^{-6}$, MAF 0.01, discordance between genotyped and recorded sex, excess inter/intra heterozygosity, non-European ancestry | MiniMac RELEASE STAMP 2012-11-16 | 1000G (phase-1, version-3) | HASE |
| SHIP | Affymetrix SNP 6.0 | HWE p-value $10^{-4}$, call rate 80%, sample call rate 92%, duplicate samples (by IBS) or reported/genotyped gender mismatch | IMPUTE2 v2.2.2 | ALL 1000 Genomes (phase 1 version 3; March 2012) | QUICKTEST v0.95 |
| SHIP-Trend | Illumina Omni 2.5 | HWE p-value $10^{-4}$, call rate 90%, sample call rate 94%, duplicate samples (by IBS) or reported/genotyped gender mismatch | IMPUTE2 v2.2.2 | ALL 1000 Genomes (phase 1 version 3; March 2012) | QUICKTEST v0.95 |
| UKBB | Applied Biosystems™ UK BiLEVE Axiom™ Array by Affymetrix and Applied Biosystems™ UK Biobank Axiom™ Array | HWE p-value $10^{-6}$, call rate 95%, sample call rate 95%, non-European ancestry, sample failures, sex mismatch, high autosomal heterozygosity, cryptic relatedness | SHPAEIT (v3)/IMPUTE4 | Haplotype Reference Consortium (HRC, version 1.1) | HASE |

**Table E2A. GWAS meta-analysis results for surface area PC1 for SNPs with P-values< 5E-08.**
The table shows the genomic positions, alleles, P-values, nearest genes, and the functions with respect to the nearest genes.

| | | | | | | | | | | | Meta-GWAS | | | UKBB | | | ENIGMA* | | | |
|---|---|---|---|---|---|---|---|---|---|---|---|---|---|---|---|---|---|---|---|---|
| rsID | chr | pos (hg19) | Genomic Locus | Genomic Locus start | Genomic Locus end | A1 | A2 | MAF | Nearest Genes | Function | Beta | SE | P-value | Beta | SE | P-value | Beta | SE | P-value | Replicated** |
| rs58380009 | 1 | 9720496 | 1 | 9720496 | 9721289 | T | C | 0.2197 | PIK3CD | intronic | 0.9904 | 0.1791 | 3.19E-08 | 0.3613 | 0.3344 | 2.80E-01 | 0.0126 | 0.0130 | 3.29E-01 | 0 |
| rs58784124 | 1 | 9720526 | 1 | 9720496 | 9721289 | A | G | 0.2217 | PIK3CD | intronic | -0.9938 | 0.1791 | 2.85E-08 | -0.3588 | 0.3343 | 2.83E-01 | -0.0123 | 0.0130 | 3.43E-01 | 0 |
| rs12088060 | 1 | 9721289 | 1 | 9720496 | 9721289 | A | G | 0.2217 | PIK3CD:RP11-558F | ncRNA_intronic | 0.9924 | 0.1796 | 3.29E-08 | 0.3655 | 0.3342 | 2.74E-01 | 0.0128 | 0.0130 | 3.24E-01 | 0 |
| rs12495978 | 3 | 141707113 | 2 | 141637438 | 141847759 | T | G | 0.1779 | TFDP2 | intronic | -1.0453 | 0.1912 | 4.60E-08 | -1.5674 | 0.3751 | 2.97E-05 | -0.0917 | 0.0139 | 4.52E-11 | 1 |
| rs1511299 | 3 | 141716072 | 2 | 141637438 | 141847759 | T | C | 0.2604 | TFDP2 | intronic | -0.8564 | 0.1557 | 3.80E-08 | -0.8673 | 0.3064 | 4.66E-03 | -0.0506 | 0.0112 | 6.11E-06 | 1 |
| rs34186890 | 3 | 141720712 | 2 | 141637438 | 141847759 | A | G | 0.2614 | TFDP2 | intronic | -0.8576 | 0.1556 | 3.60E-08 | -0.8742 | 0.3061 | 4.30E-03 | -0.0506 | 0.0112 | 6.16E-06 | 1 |
| rs7671110 | 4 | 17874089 | 3 | 17792249 | 18035250 | T | C | 0.1252 | LCORL | intronic | -1.0865 | 0.1914 | 1.37E-08 | -1.1556 | 0.3683 | 1.71E-03 | -0.0389 | 0.0138 | 4.86E-03 | 1 |
| rs7663887 | 4 | 17902920 | 3 | 17792249 | 18035250 | A | C | 0.1252 | LCORL | intronic | -1.0873 | 0.1912 | 1.29E-08 | -1.1351 | 0.3671 | 2.00E-03 | -0.0378 | 0.0138 | 6.15E-03 | 1 |
| rs13138182 | 4 | 17908413 | 3 | 17792249 | 18035250 | A | G | 0.1252 | LCORL | intronic | 1.0899 | 0.1912 | 1.20E-08 | 1.1334 | 0.3671 | 2.03E-03 | 0.0376 | 0.0138 | 6.36E-03 | 1 |
| rs1472852 | 4 | 17910236 | 3 | 17792249 | 18035250 | A | C | 0.1252 | LCORL | intronic | -1.0859 | 0.1911 | 1.33E-08 | -1.1490 | 0.3673 | 1.77E-03 | -0.0373 | 0.0138 | 6.70E-03 | 1 |
| rs6813340 | 4 | 17922866 | 3 | 17792249 | 18035250 | A | G | 0.1252 | LCORL | intronic | 1.0945 | 0.1911 | 1.03E-08 | 1.1242 | 0.3678 | 2.25E-03 | 0.0372 | 0.0137 | 6.75E-03 | 1 |
| rs11938781 | 4 | 17924734 | 3 | 17792249 | 18035250 | T | C | 0.1451 | LCORL | intronic | 1.1313 | 0.1892 | 2.24E-09 | 1.1246 | 0.3647 | 2.05E-03 | 0.0416 | 0.0138 | 2.49E-03 | 1 |
| rs13146142 | 4 | 17931318 | 3 | 17792249 | 18035250 | T | C | 0.1262 | LCORL | intronic | 1.0933 | 0.1907 | 9.81E-09 | 1.1187 | 0.3673 | 2.33E-03 | 0.0359 | 0.0137 | 8.56E-03 | 1 |
| rs73802707 | 4 | 17932319 | 3 | 17792249 | 18035250 | T | C | 0.1193 | LCORL | intronic | -1.1240 | 0.1955 | 8.98E-09 | -1.0215 | 0.3721 | 6.07E-03 | -0.0360 | 0.0143 | 1.16E-02 | 1 |
| rs6841793 | 4 | 17933810 | 3 | 17792249 | 18035250 | C | G | 0.1262 | LCORL | intronic | 1.0820 | 0.1904 | 1.32E-08 | 1.1337 | 0.3671 | 2.02E-03 | 0.0361 | 0.0136 | 8.06E-03 | 1 |
| rs6449345 | 4 | 17934394 | 3 | 17792249 | 18035250 | A | T | 0.1262 | LCORL | intronic | 1.0804 | 0.1904 | 1.39E-08 | 1.1330 | 0.3672 | 2.04E-03 | 0.0362 | 0.0136 | 8.04E-03 | 1 |
| rs6449346 | 4 | 17934461 | 3 | 17792249 | 18035250 | T | C | 0.1262 | LCORL | intronic | 1.0784 | 0.1903 | 1.46E-08 | 1.1343 | 0.3671 | 2.01E-03 | 0.0361 | 0.0136 | 8.05E-03 | 1 |
| rs7663818 | 4 | 17936443 | 3 | 17792249 | 18035250 | T | C | 0.1262 | LCORL | intronic | -1.0620 | 0.1903 | 2.38E-08 | -1.1370 | 0.3671 | 1.96E-03 | -0.0360 | 0.0136 | 8.23E-03 | 1 |
| rs7692995 | 4 | 17936634 | 3 | 17792249 | 18035250 | T | C | 0.1262 | LCORL | intronic | 1.0755 | 0.1903 | 1.59E-08 | 1.1371 | 0.3671 | 1.96E-03 | 0.0360 | 0.0136 | 8.25E-03 | 1 |
| rs34683079 | 4 | 17939302 | 3 | 17792249 | 18035250 | T | C | 0.1272 | LCORL | intronic | 1.0795 | 0.1901 | 1.36E-08 | 1.1363 | 0.3671 | 1.97E-03 | 0.0355 | 0.0136 | 9.14E-03 | 1 |
| rs6840868 | 4 | 17939331 | 3 | 17792249 | 18035250 | T | C | 0.1272 | LCORL | intronic | -1.0794 | 0.1901 | 1.37E-08 | -1.1367 | 0.3671 | 1.97E-03 | -0.0354 | 0.0136 | 9.17E-03 | 1 |
| rs6845118 | 4 | 17939740 | 3 | 17792249 | 18035250 | T | G | 0.1272 | LCORL | intronic | -1.1009 | 0.1911 | 8.33E-09 | -1.1363 | 0.3671 | 1.97E-03 | -0.0362 | 0.0138 | 8.59E-03 | 1 |
| rs12511600 | 4 | 17939921 | 3 | 17792249 | 18035250 | T | C | 0.1272 | LCORL | intronic | -1.0801 | 0.1901 | 1.33E-08 | -1.1364 | 0.3671 | 1.97E-03 | -0.0355 | 0.0136 | 9.14E-03 | 1 |
| rs71603391 | 4 | 17940197 | 3 | 17792249 | 18035250 | A | G | 0.1282 | LCORL | intronic | 1.1222 | 0.1937 | 6.84E-09 | 1.1297 | 0.3670 | 2.09E-03 | 0.0357 | 0.0141 | 1.13E-02 | 1 |
| rs73098845 | 4 | 17940718 | 3 | 17792249 | 18035250 | A | C | 0.1272 | LCORL | intronic | -1.0778 | 0.1901 | 1.44E-08 | -1.1357 | 0.3672 | 1.99E-03 | -0.0354 | 0.0136 | 9.24E-03 | 1 |
| rs73098848 | 4 | 17941027 | 3 | 17792249 | 18035250 | C | G | 0.1272 | LCORL | intronic | -1.0782 | 0.1901 | 1.42E-08 | -1.1359 | 0.3672 | 1.99E-03 | -0.0354 | 0.0136 | 9.28E-03 | 1 |
| rs16896068 | 4 | 17944840 | 3 | 17792249 | 18035250 | A | G | 0.1282 | LCORL | intronic | -1.0815 | 0.19 | 1.25E-08 | -1.1366 | 0.3675 | 1.99E-03 | -0.0353 | 0.0136 | 9.42E-03 | 1 |
| rs16896074 | 4 | 17947873 | 3 | 17792249 | 18035250 | T | C | 0.1272 | LCORL | intronic | 1.0677 | 0.1899 | 1.87E-08 | 1.1663 | 0.3666 | 1.47E-03 | 0.0347 | 0.0136 | 1.05E-02 | 1 |
| rs12503997 | 4 | 17948089 | 3 | 17792249 | 18035250 | A | T | 0.1272 | LCORL | intronic | -1.0726 | 0.1898 | 1.60E-08 | -1.1665 | 0.3665 | 1.46E-03 | -0.0347 | 0.0136 | 1.06E-02 | 1 |
| rs12506752 | 4 | 17948352 | 3 | 17792249 | 18035250 | A | G | 0.1272 | LCORL | intronic | -1.0710 | 0.1898 | 1.68E-08 | -1.1665 | 0.3665 | 1.46E-03 | -0.0346 | 0.0136 | 1.07E-02 | 1 |
| rs11936915 | 4 | 17948497 | 3 | 17792249 | 18035250 | T | C | 0.1272 | LCORL | intronic | 1.0710 | 0.1898 | 1.69E-08 | 1.1665 | 0.3665 | 1.46E-03 | 0.0346 | 0.0136 | 1.07E-02 | 1 |
| rs11936911 | 4 | 17948668 | 3 | 17792249 | 18035250 | A | G | 0.1272 | LCORL | intronic | 1.0727 | 0.1898 | 1.59E-08 | 1.1666 | 0.3665 | 1.46E-03 | 0.0346 | 0.0136 | 1.07E-02 | 1 |
| rs6850259 | 4 | 17949607 | 3 | 17792249 | 18035250 | A | G | 0.1272 | LCORL | intronic | 1.0691 | 0.1899 | 1.80E-08 | 1.1656 | 0.3667 | 1.49E-03 | 0.0346 | 0.0136 | 1.08E-02 | 1 |
| rs34934312 | 4 | 17950172 | 3 | 17792249 | 18035250 | A | T | 0.1272 | LCORL | intronic | -1.0668 | 0.19 | 1.95E-08 | -1.1394 | 0.3676 | 1.95E-03 | -0.0345 | 0.0136 | 1.09E-02 | 1 |
| rs16896101 | 4 | 17950626 | 3 | 17792249 | 18035250 | A | G | 0.1272 | LCORL | intronic | 1.0670 | 0.19 | 1.95E-08 | 1.1394 | 0.3676 | 1.95E-03 | 0.0345 | 0.0136 | 1.10E-02 | 1 |
| rs16896113 | 4 | 17950850 | 3 | 17792249 | 18035250 | T | C | 0.1272 | LCORL | intronic | 1.0721 | 0.1899 | 1.64E-08 | 1.1391 | 0.3676 | 1.95E-03 | 0.0355 | 0.0136 | 8.92E-03 | 1 |
| rs7657678 | 4 | 17951035 | 3 | 17792249 | 18035250 | A | G | 0.1272 | LCORL | intronic | 1.0670 | 0.19 | 1.95E-08 | 1.1392 | 0.3676 | 1.95E-03 | 0.0344 | 0.0136 | 1.13E-02 | 1 |
| rs6843330 | 4 | 17952056 | 3 | 17792249 | 18035250 | A | G | 0.1272 | LCORL | intronic | -1.0671 | 0.19 | 1.94E-08 | -1.1393 | 0.3676 | 1.95E-03 | -0.0342 | 0.0136 | 1.16E-02 | 1 |
| rs6449349 | 4 | 17952609 | 3 | 17792249 | 18035250 | C | G | 0.1272 | LCORL | intronic | -1.0693 | 0.1899 | 1.80E-08 | -1.1645 | 0.3667 | 1.50E-03 | -0.0341 | 0.0136 | 1.20E-02 | 1 |
| rs7675806 | 4 | 17952959 | 3 | 17792249 | 18035250 | T | C | 0.1272 | LCORL | intronic | 1.0704 | 0.1899 | 1.74E-08 | 1.1655 | 0.3666 | 1.48E-03 | 0.0342 | 0.0136 | 1.18E-02 | 1 |
| rs16896128 | 4 | 17953590 | 3 | 17792249 | 18035250 | A | G | 0.1272 | LCORL | intronic | 1.0713 | 0.1899 | 1.68E-08 | 1.1655 | 0.3666 | 1.48E-03 | 0.0341 | 0.0136 | 1.20E-02 | 1 |
| rs7660642 | 4 | 17956017 | 3 | 17792249 | 18035250 | T | C | 0.1272 | LCORL | intronic | 1.0686 | 0.1899 | 1.82E-08 | 1.1578 | 0.3667 | 1.60E-03 | 0.0340 | 0.0136 | 1.21E-02 | 1 |
| rs4642247 | 4 | 17957158 | 3 | 17792249 | 18035250 | A | G | 0.1272 | LCORL | intronic | -1.0674 | 0.19 | 1.94E-08 | -1.1387 | 0.3676 | 1.96E-03 | -0.0338 | 0.0136 | 1.26E-02 | 1 |
| rs4057984 | 4 | 17957200 | 3 | 17792249 | 18035250 | T | C | 0.1272 | LCORL | intronic | 1.0674 | 0.19 | 1.94E-08 | 1.1387 | 0.3676 | 1.96E-03 | 0.0338 | 0.0136 | 1.26E-02 | 1 |
| rs7684221 | 4 | 17957354 | 3 | 17792249 | 18035250 | A | G | 0.1272 | LCORL | intronic | -1.0701 | 0.1902 | 1.85E-08 | -1.1387 | 0.3676 | 1.96E-03 | -0.0340 | 0.0136 | 1.23E-02 | 1 |
| rs7686082 | 4 | 17957576 | 3 | 17792249 | 18035250 | T | C | 0.1272 | LCORL | intronic | -1.0681 | 0.19 | 1.91E-08 | -1.1384 | 0.3675 | 1.96E-03 | -0.0338 | 0.0136 | 1.28E-02 | 1 |
| rs7673321 | 4 | 17957688 | 3 | 17792249 | 18035250 | A | T | 0.1272 | LCORL | intronic | -1.0557 | 0.1902 | 2.84E-08 | -1.0771 | 0.3679 | 3.43E-03 | -0.0335 | 0.0136 | 1.37E-02 | 1 |
| rs7690823 | 4 | 17957698 | 3 | 17792249 | 18035250 | T | C | 0.1272 | LCORL | intronic | -1.0673 | 0.19 | 1.94E-08 | -1.1391 | 0.3676 | 1.95E-03 | -0.0338 | 0.0136 | 1.28E-02 | 1 |
| rs10446630 | 4 | 17958690 | 3 | 17792249 | 18035250 | C | G | 0.1272 | LCORL | intronic | 1.0692 | 0.19 | 1.82E-08 | 1.1402 | 0.3674 | 1.92E-03 | 0.0338 | 0.0136 | 1.28E-02 | 1 |
| rs13151431 | 4 | 17959581 | 3 | 17792249 | 18035250 | A | C | 0.1272 | LCORL | intronic | -1.0672 | 0.1899 | 1.92E-08 | -1.1646 | 0.3667 | 1.50E-03 | -0.0338 | 0.0136 | 1.27E-02 | 1 |
| rs13105471 | 4 | 17959776 | 3 | 17792249 | 18035250 | T | C | 0.1272 | LCORL | intronic | -1.0545 | 0.1894 | 2.60E-08 | -1.1646 | 0.3667 | 1.50E-03 | -0.0328 | 0.0135 | 1.51E-02 | 1 |
| rs11945359 | 4 | 17960008 | 3 | 17792249 | 18035250 | T | C | 0.1272 | LCORL | intronic | 1.0729 | 0.1899 | 1.61E-08 | 1.1646 | 0.3667 | 1.50E-03 | 0.0338 | 0.0136 | 1.27E-02 | 1 |
| rs11930366 | 4 | 17960168 | 3 | 17792249 | 18035250 | A | G | 0.1272 | LCORL | intronic | -1.0675 | 0.1899 | 1.90E-08 | -1.1641 | 0.3667 | 1.51E-03 | -0.0338 | 0.0136 | 1.27E-02 | 1 |
| rs1876219 | 4 | 17961415 | 3 | 17792249 | 18035250 | A | G | 0.1272 | LCORL | intronic | -1.0666 | 0.1899 | 1.96E-08 | -1.1647 | 0.3667 | 1.50E-03 | -0.0338 | 0.0136 | 1.27E-02 | 1 |
| rs7670435 | 4 | 17962260 | 3 | 17792249 | 18035250 | A | G | 0.1272 | LCORL | intronic | -1.0663 | 0.1899 | 1.97E-08 | -1.1646 | 0.3667 | 1.50E-03 | -0.0338 | 0.0136 | 1.27E-02 | 1 |
| rs7695840 | 4 | 17962379 | 3 | 17792249 | 18035250 | A | G | 0.1272 | LCORL | intronic | 1.0604 | 0.1899 | 2.36E-08 | 1.1626 | 0.3667 | 1.53E-03 | 0.0338 | 0.0136 | 1.27E-02 | 1 |
| rs16896155 | 4 | 17962935 | 3 | 17792249 | 18035250 | A | T | 0.1272 | LCORL | intronic | 1.0660 | 0.1899 | 2.00E-08 | 1.1646 | 0.3667 | 1.50E-03 | 0.0338 | 0.0136 | 1.26E-02 | 1 |
| rs7662221 | 4 | 17966468 | 3 | 17792249 | 18035250 | A | G | 0.1272 | LCORL | intronic | -1.0667 | 0.1899 | 1.95E-08 | -1.1650 | 0.3667 | 1.50E-03 | -0.0338 | 0.0136 | 1.26E-02 | 1 |
| rs7697634 | 4 | 17966746 | 3 | 17792249 | 18035250 | T | C | 0.1272 | LCORL | intronic | 1.0650 | 0.1899 | 2.06E-08 | 1.1680 | 0.3667 | 1.45E-03 | 0.0341 | 0.0136 | 1.20E-02 | 1 |
| rs16896164 | 4 | 17967094 | 3 | 17792249 | 18035250 | T | C | 0.1272 | LCORL | intronic | 1.0863 | 0.1907 | 1.22E-08 | 1.1587 | 0.3669 | 1.60E-03 | 0.0341 | 0.0136 | 1.19E-02 | 1 |
| rs34588175 | 4 | 17967114 | 3 | 17792249 | 18035250 | A | G | 0.1272 | LCORL | intronic | -1.0807 | 0.1906 | 1.43E-08 | -1.1280 | 0.3679 | 2.18E-03 | -0.0341 | 0.0136 | 1.19E-02 | 1 |
| rs11933767 | 4 | 17967896 | 3 | 17792249 | 18035250 | C | G | 0.1272 | LCORL | intronic | -1.0571 | 0.19 | 2.64E-08 | -1.1387 | 0.3676 | 1.96E-03 | -0.0339 | 0.0136 | 1.25E-02 | 1 |
| rs34025766 | 4 | 17968811 | 3 | 17792249 | 18035250 | A | T | 0.1272 | LCORL | intronic | -1.0617 | 0.19 | 2.31E-08 | -1.1385 | 0.3674 | 1.95E-03 | -0.0339 | 0.0136 | 1.25E-02 | 1 |
| rs6853216 | 4 | 17970655 | 3 | 17792249 | 18035250 | T | C | 0.1272 | LCORL | intronic | -1.0482 | 0.19 | 3.45E-08 | -1.1317 | 0.3673 | 2.07E-03 | -0.0340 | 0.0136 | 1.21E-02 | 1 |
| rs11933575 | 4 | 17971138 | 3 | 17792249 | 18035250 | A | G | 0.1272 | LCORL | intronic | 1.0567 | 0.1901 | 2.72E-08 | 1.1404 | 0.3673 | 1.91E-03 | 0.0339 | 0.0136 | 1.25E-02 | 1 |
| rs11930105 | 4 | 17972078 | 3 | 17792249 | 18035250 | A | C | 0.1272 | LCORL | intronic | -1.0576 | 0.19 | 2.59E-08 | -1.1307 | 0.3675 | 2.10E-03 | -0.0339 | 0.0136 | 1.24E-02 | 1 |
| rs7661129 | 4 | 17973171 | 3 | 17792249 | 18035250 | C | G | 0.1272 | LCORL | intronic | -1.0605 | 0.19 | 2.40E-08 | -1.1383 | 0.3673 | 1.95E-03 | -0.0339 | 0.0136 | 1.25E-02 | 1 |
| rs11938188 | 4 | 17973698 | 3 | 17792249 | 18035250 | T | C | 0.1272 | LCORL | intronic | 1.0605 | 0.19 | 2.40E-08 | 1.1382 | 0.3673 | 1.95E-03 | 0.0339 | 0.0136 | 1.26E-02 | 1 |
| rs16896182 | 4 | 17973899 | 3 | 17792249 | 18035250 | C | G | 0.1272 | LCORL | intronic | 1.0600 | 0.19 | 2.42E-08 | 1.1316 | 0.3673 | 2.07E-03 | 0.0339 | 0.0136 | 1.25E-02 | 1 |
| rs13144742 | 4 | 17974181 | 3 | 17792249 | 18035250 | A | G | 0.1272 | LCORL | intronic | 1.0585 | 0.19 | 2.53E-08 | 1.1317 | 0.3673 | 2.07E-03 | 0.0348 | 0.0136 | 1.04E-02 | 1 |
| rs12500636 | 4 | 17974883 | 3 | 17792249 | 18035250 | T | G | 0.1272 | LCORL | intronic | 1.0615 | 0.19 | 2.30E-08 | 1.1636 | 0.3665 | 1.50E-03 | 0.0339 | 0.0136 | 1.26E-02 | 1 |
| rs12511456 | 4 | 17975603 | 3 | 17792249 | 18035250 | T | C | 0.1272 | LCORL | intronic | -1.0602 | 0.19 | 2.42E-08 | -1.1382 | 0.3673 | 1.95E-03 | -0.0338 | 0.0136 | 1.27E-02 | 1 |
| rs12501035 | 4 | 17976326 | 3 | 17792249 | 18035250 | A | G | 0.1272 | LCORL | intronic | 1.0586 | 0.19 | 2.52E-08 | 1.1626 | 0.3667 | 1.53E-03 | 0.0338 | 0.0136 | 1.27E-02 | 1 |
| rs13127416 | 4 | 17976779 | 3 | 17792249 | 18035250 | A | G | 0.1272 | LCORL | intronic | 1.0585 | 0.19 | 2.53E-08 | 1.1626 | 0.3667 | 1.53E-03 | 0.0338 | 0.0136 | 1.27E-02 | 1 |
| rs13152352 | 4 | 17976846 | 3 | 17792249 | 18035250 | A | C | 0.1272 | LCORL | intronic | -1.0563 | 0.1901 | 2.73E-08 | -1.1631 | 0.3667 | 1.52E-03 | -0.0335 | 0.0136 | 1.38E-02 | 1 |
| rs6449350 | 4 | 17977535 | 3 | 17792249 | 18035250 | A | C | 0.1272 | LCORL | intronic | 1.0588 | 0.1899 | 2.49E-08 | 1.1637 | 0.3665 | 1.50E-03 | 0.0338 | 0.0136 | 1.28E-02 | 1 |
| rs34156351 | 4 | 17983055 | 3 | 17792249 | 18035250 | A | G | 0.1272 | LCORL | intronic | -1.0543 | 0.19 | 2.89E-08 | -1.1373 | 0.3676 | 1.99E-03 | -0.0336 | 0.0136 | 1.33E-02 | 1 |
| rs6839954 | 4 | 17984129 | 3 | 17792249 | 18035250 | T | G | 0.1272 | LCORL | intronic | 1.0580 | 0.19 | 2.55E-08 | 1.1637 | 0.3665 | 1.51E-03 | 0.0336 | 0.0136 | 1.33E-02 | 1 |
| rs6840334 | 4 | 17984277 | 3 | 17792249 | 18035250 | T | C | 0.1272 | LCORL | intronic | 1.0580 | 0.19 | 2.55E-08 | 1.1637 | 0.3665 | 1.50E-03 | 0.0336 | 0.0136 | 1.34E-02 | 1 |
| rs6835186 | 4 | 17984316 | 3 | 17792249 | 18035250 | A | C | 0.1272 | LCORL | intronic | 1.0579 | 0.19 | 2.56E-08 | 1.1637 | 0.3665 | 1.50E-03 | 0.0336 | 0.0136 | 1.34E-02 | 1 |
| rs6810665 | 4 | 17984462 | 3 | 17792249 | 18035250 | A | C | 0.1272 | LCORL | intronic | -1.0628 | 0.1899 | 2.19E-08 | -1.1637 | 0.3665 | 1.51E-03 | -0.0333 | 0.0136 | 1.44E-02 | 1 |
| rs35338009 | 4 | 17984842 | 3 | 17792249 | 18035250 | T | G | 0.1272 | LCORL | intronic | -1.0528 | 0.191 | 3.55E-08 | -1.1392 | 0.3674 | 1.94E-03 | -0.0330 | 0.0136 | 1.53E-02 | 1 |
| rs35565383 | 4 | 17984874 | 3 | 17792249 | 18035250 | C | G | 0.1272 | LCORL | intronic | 1.0354 | 0.1899 | 4.98E-08 | 1.1383 | 0.3674 | 1.95E-03 | 0.0327 | 0.0136 | 1.61E-02 | 1 |
| rs67113817 | 4 | 17985189 | 3 | 17792249 | 18035250 | T | C | 0.1272 | LCORL | intronic | 1.0571 | 0.19 | 2.62E-08 | 1.1636 | 0.3665 | 1.51E-03 | 0.0336 | 0.0136 | 1.35E-02 | 1 |
| rs35210497 | 4 | 17985374 | 3 | 17792249 | 18035250 | T | C | 0.1272 | LCORL | intronic | 1.0571 | 0.19 | 2.62E-08 | 1.1634 | 0.3665 | 1.51E-03 | 0.0336 | 0.0136 | 1.35E-02 | 1 |
| rs1987068 | 4 | 17986522 | 3 | 17792249 | 18035250 | T | C | 0.1272 | LCORL | intronic | -1.0563 | 0.19 | 2.72E-08 | -1.1382 | 0.3674 | 1.96E-03 | -0.0336 | 0.0136 | 1.35E-02 | 1 |
| rs35317957 | 4 | 17986863 | 3 | 17792249 | 18035250 | T | C | 0.1272 | LCORL | intronic | 1.0563 | 0.19 | 2.72E-08 | 1.1382 | 0.3674 | 1.96E-03 | 0.0335 | 0.0136 | 1.36E-02 | 1 |
| rs7668933 | 4 | 17988141 | 3 | 17792249 | 18035250 | A | G | 0.1272 | LCORL | intronic | 1.0601 | 0.19 | 2.41E-08 | 1.1321 | 0.3673 | 2.06E-03 | 0.0331 | 0.0136 | 1.49E-02 | 1 |
| rs35348660 | 4 | 17988635 | 3 | 17792249 | 18035250 | A | T | 0.1272 | LCORL | intronic | -1.0571 | 0.19 | 2.65E-08 | -1.1627 | 0.3667 | 1.53E-03 | -0.0335 | 0.0136 | 1.38E-02 | 1 |
| rs13132994 | 4 | 17990181 | 3 | 17792249 | 18035250 | A | G | 0.1272 | LCORL | intronic | 1.0558 | 0.19 | 2.77E-08 | 1.1383 | 0.3674 | 1.95E-03 | 0.0335 | 0.0136 | 1.37E-02 | 1 |
| rs67978911 | 4 | 17991423 | 3 | 17792249 | 18035250 | T | C | 0.1272 | LCORL | intronic | -1.0568 | 0.19 | 2.65E-08 | -1.1637 | 0.3665 | 1.50E-03 | -0.0335 | 0.0136 | 1.38E-02 | 1 |
| rs16896210 | 4 | 17992332 | 3 | 17792249 | 18035250 | A | G | 0.1272 | LCORL | intronic | 1.0638 | 0.1919 | 2.95E-08 | 1.1564 | 0.3665 | 1.61E-03 | 0.0325 | 0.0139 | 1.91E-02 | 1 |
| rs16896213 | 4 | 17994661 | 3 | 17792249 | 18035250 | C | G | 0.1272 | LCORL | intronic | 1.0577 | 0.1899 | 2.57E-08 | 1.1630 | 0.3664 | 1.51E-03 | 0.0336 | 0.0136 | 1.34E-02 | 1 |
| rs34358941 | 4 | 17995047 | 3 | 17792249 | 18035250 | T | C | 0.1272 | LCORL | intronic | 1.0580 | 0.19 | 2.55E-08 | 1.1632 | 0.3664 | 1.51E-03 | 0.0336 | 0.0136 | 1.34E-02 | 1 |

| | | | | | | | | | | | Meta-GWAS | | | UKBB | | | ENIGMA* | | | |
|---|---|---|---|---|---|---|---|---|---|---|---|---|---|---|---|---|---|---|---|---|
| rsID | chr | pos (hg19) | Genomic Locus | Genomic Locus start | Genomic Locus end | A1 | A2 | MAF | Nearest Genes | Function | Beta | SE | P-value | Beta | SE | P-value | Beta | SE | P-value | Replicated** |
| rs16896215 | 4 | 17995312 | 3 | 17792249 | 18035250 | T | C | 0.1272 | LCORL | intronic | 1.0574 | 0.19 | 2.60E-08 | 1.1632 | 0.3664 | 1.51E-03 | 0.0336 | 0.0136 | 1.34E-02 | 1 |
| rs6843651 | 4 | 17995500 | 3 | 17792249 | 18035250 | T | C | 0.1272 | LCORL | intronic | 1.0574 | 0.19 | 2.60E-08 | 1.1633 | 0.3664 | 1.51E-03 | 0.0336 | 0.0136 | 1.34E-02 | 1 |
| rs6824748 | 4 | 17997066 | 3 | 17792249 | 18035250 | A | G | 0.1272 | LCORL | intronic | -1.0517 | 0.1901 | 3.15E-08 | -1.1336 | 0.3673 | 2.04E-03 | -0.0337 | 0.0136 | 1.32E-02 | 1 |
| rs12513171 | 4 | 17997541 | 3 | 17792249 | 18035250 | T | C | 0.1292 | LCORL | intronic | 1.0489 | 0.19 | 3.36E-08 | 1.1338 | 0.3672 | 2.02E-03 | 0.0343 | 0.0136 | 1.17E-02 | 1 |
| rs16896236 | 4 | 18003812 | 3 | 17792249 | 18035250 | T | C | 0.1272 | LCORL | intronic | 1.0704 | 0.19 | 1.77E-08 | 1.1769 | 0.3664 | 1.33E-03 | 0.0337 | 0.0136 | 1.32E-02 | 1 |
| rs6821168 | 4 | 18005019 | 3 | 17792249 | 18035250 | A | C | 0.1272 | LCORL | intronic | 1.0561 | 0.1901 | 2.78E-08 | 1.1488 | 0.3675 | 1.78E-03 | 0.0337 | 0.0136 | 1.32E-02 | 1 |
| rs16896245 | 4 | 18005608 | 3 | 17792249 | 18035250 | T | C | 0.1272 | LCORL | intronic | -1.0511 | 0.1902 | 3.25E-08 | -1.1488 | 0.3675 | 1.78E-03 | -0.0338 | 0.0136 | 1.30E-02 | 1 |
| rs35613046 | 4 | 18005966 | 3 | 17792249 | 18035250 | A | G | 0.1272 | LCORL | intronic | 1.0520 | 0.1901 | 3.16E-08 | 1.1487 | 0.3675 | 1.78E-03 | 0.0338 | 0.0136 | 1.30E-02 | 1 |
| rs6828122 | 4 | 18006052 | 3 | 17792249 | 18035250 | A | G | 0.1272 | LCORL | intronic | 1.0503 | 0.1902 | 3.32E-08 | 1.1487 | 0.3675 | 1.78E-03 | 0.0338 | 0.0136 | 1.31E-02 | 1 |
| rs12513324 | 4 | 18006951 | 3 | 17792249 | 18035250 | T | C | 0.1272 | LCORL | intronic | -1.0640 | 0.192 | 2.97E-08 | -1.1388 | 0.3673 | 1.94E-03 | -0.0335 | 0.0140 | 1.66E-02 | 1 |
| rs2085683 | 4 | 18007435 | 3 | 17792249 | 18035250 | C | G | 0.1272 | LCORL | intronic | 1.0525 | 0.1902 | 3.11E-08 | 1.1367 | 0.3676 | 1.99E-03 | 0.0338 | 0.0136 | 1.30E-02 | 1 |
| rs6842114 | 4 | 18008199 | 3 | 17792249 | 18035250 | A | C | 0.1272 | LCORL | intronic | 1.0564 | 0.1901 | 2.74E-08 | 1.1570 | 0.3667 | 1.61E-03 | 0.0339 | 0.0136 | 1.29E-02 | 1 |
| rs6849629 | 4 | 18009488 | 3 | 17792249 | 18035250 | A | G | 0.1272 | LCORL | intronic | 1.0551 | 0.1902 | 2.88E-08 | 1.1378 | 0.3673 | 1.96E-03 | 0.0339 | 0.0136 | 1.28E-02 | 1 |
| rs6811842 | 4 | 18009798 | 3 | 17792249 | 18035250 | A | T | 0.1272 | LCORL | intronic | -1.0584 | 0.19 | 2.56E-08 | -1.1368 | 0.3676 | 1.99E-03 | -0.0342 | 0.0136 | 1.21E-02 | 1 |
| rs16896271 | 4 | 18012454 | 3 | 17792249 | 18035250 | T | C | 0.1272 | LCORL | intronic | 1.0549 | 0.1901 | 2.88E-08 | 1.1368 | 0.3675 | 1.99E-03 | 0.0337 | 0.0137 | 1.38E-02 | 1 |
| rs10516315 | 4 | 18016877 | 3 | 17792249 | 18035250 | A | G | 0.1272 | LCORL | intronic | 1.0595 | 0.19 | 2.47E-08 | 1.1627 | 0.3664 | 1.51E-03 | 0.0337 | 0.0137 | 1.38E-02 | 1 |
| rs6830062 | 4 | 18017730 | 3 | 17792249 | 18035250 | T | C | 0.1272 | LCORL | intronic | 1.0560 | 0.19 | 2.71E-08 | 1.1941 | 0.3660 | 1.11E-03 | 0.0338 | 0.0137 | 1.34E-02 | 1 |
| rs35081654 | 4 | 18017938 | 3 | 17792249 | 18035250 | A | T | 0.1272 | LCORL | intronic | 1.0591 | 0.19 | 2.50E-08 | 1.1629 | 0.3664 | 1.51E-03 | 0.0336 | 0.0137 | 1.41E-02 | 1 |
| rs6831102 | 4 | 18018236 | 3 | 17792249 | 18035250 | T | C | 0.1272 | LCORL | intronic | 1.0585 | 0.1901 | 2.57E-08 | 1.1629 | 0.3664 | 1.51E-03 | 0.0336 | 0.0137 | 1.43E-02 | 1 |
| rs1585334 | 4 | 18018944 | 3 | 17792249 | 18035250 | T | C | 0.1272 | LCORL | intronic | -1.0602 | 0.1901 | 2.46E-08 | -1.1630 | 0.3664 | 1.51E-03 | -0.0337 | 0.0137 | 1.40E-02 | 1 |
| rs1585333 | 4 | 18019025 | 3 | 17792249 | 18035250 | T | C | 0.1272 | LCORL | intronic | 1.0602 | 0.1901 | 2.46E-08 | 1.1629 | 0.3664 | 1.51E-03 | 0.0335 | 0.0137 | 1.45E-02 | 1 |
| rs7698271 | 4 | 18019413 | 3 | 17792249 | 18035250 | T | C | 0.1272 | LCORL | intronic | -1.0604 | 0.1901 | 2.45E-08 | -1.1629 | 0.3664 | 1.51E-03 | -0.0335 | 0.0137 | 1.45E-02 | 1 |
| rs7698644 | 4 | 18019572 | 3 | 17792249 | 18035250 | T | C | 0.1272 | LCORL | intronic | -1.0672 | 0.1901 | 1.98E-08 | -1.1629 | 0.3664 | 1.51E-03 | -0.0335 | 0.0137 | 1.46E-02 | 1 |
| rs7659195 | 4 | 18022692 | 3 | 17792249 | 18035250 | T | C | 0.1272 | LCORL | intronic | 1.0710 | 0.1956 | 4.36E-08 | 1.1629 | 0.3664 | 1.51E-03 | 0.0342 | 0.0144 | 1.73E-02 | 1 |
| rs1380294 | 4 | 18024121 | 3 | 17792249 | 18035250 | T | C | 0.1272 | LCORL | upstream | -1.0630 | 0.1904 | 2.35E-08 | -1.1304 | 0.3673 | 2.09E-03 | -0.0331 | 0.0138 | 1.62E-02 | 1 |
| rs7668417 | 4 | 18025029 | 3 | 17792249 | 18035250 | A | G | 0.1272 | LCORL | intergenic | -1.0636 | 0.1905 | 2.38E-08 | -1.1449 | 0.3673 | 1.83E-03 | -0.0331 | 0.0138 | 1.64E-02 | 1 |
| rs2125654 | 4 | 18025368 | 3 | 17792249 | 18035250 | A | T | 0.1272 | LCORL | intergenic | -1.0644 | 0.1906 | 2.34E-08 | -1.1496 | 0.3668 | 1.73E-03 | -0.0330 | 0.0138 | 1.66E-02 | 1 |
| rs1380293 | 4 | 18027235 | 3 | 17792249 | 18035250 | T | C | 0.1272 | LCORL | intergenic | 1.0646 | 0.1906 | 2.33E-08 | 1.1674 | 0.3657 | 1.42E-03 | 0.0328 | 0.0138 | 1.74E-02 | 1 |
| rs7694606 | 4 | 18028987 | 3 | 17792249 | 18035250 | A | G | 0.1272 | LCORL | intergenic | -1.0548 | 0.1909 | 3.26E-08 | -1.1585 | 0.3669 | 1.60E-03 | -0.0330 | 0.0139 | 1.72E-02 | 1 |
| rs7694806 | 4 | 18029100 | 3 | 17792249 | 18035250 | A | G | 0.1272 | LCORL | intergenic | -1.0582 | 0.1909 | 2.97E-08 | -1.1477 | 0.3669 | 1.77E-03 | -0.0329 | 0.0139 | 1.76E-02 | 1 |
| rs7696532 | 4 | 18029299 | 3 | 17792249 | 18035250 | T | C | 0.1272 | LCORL | intergenic | -1.0595 | 0.1908 | 2.82E-08 | -1.1429 | 0.3666 | 1.83E-03 | -0.0327 | 0.0139 | 1.85E-02 | 1 |
| rs16896312 | 4 | 18030853 | 3 | 17792249 | 18035250 | T | C | 0.1272 | LCORL | intergenic | -1.0581 | 0.1912 | 3.11E-08 | -1.1732 | 0.3660 | 1.36E-03 | -0.0323 | 0.0139 | 2.06E-02 | 1 |
| rs6449353 | 4 | 18033488 | 3 | 17792249 | 18035250 | T | C | 0.1272 | LCORL | intergenic | 1.0675 | 0.1919 | 2.67E-08 | 1.1658 | 0.3655 | 1.43E-03 | 0.0323 | 0.0141 | 2.20E-02 | 1 |
| rs13107325 | 4 | 103188709 | 4 | 103001649 | 103387161 | T | C | 0.07952 | SLC39A8 | exonic | 1.5264 | 0.279 | 4.47E-08 | 0.8030 | 0.5276 | 1.28E-01 | 0.0590 | 0.0208 | 4.62E-03 | 0 |
| rs13109272 | 4 | 103292422 | 4 | 103001649 | 103387161 | T | C | 0.07654 | SLC39A8 | NA | -1.4937 | 0.2679 | 2.46E-08 | -1.1935 | 0.5276 | 2.37E-02 | -0.0546 | 0.0195 | 5.06E-03 | 1 |
| rs2301718 | 4 | 106009763 | 5 | 106009763 | 106009763 | A | G | 0.2286 | RP11-556I14.1 | ncRNA_intronic | 0.9752 | 0.1649 | 3.32E-09 | 0.2256 | 0.3311 | 4.96E-01 | 0.0451 | 0.0118 | 1.32E-04 | 0 |
| rs2764264 | 6 | 108934461 | 6 | 108861264 | 109019323 | T | C | 0.3728 | FOXO3 | intronic | 0.8178 | 0.1481 | 3.34E-08 | 1.2551 | 0.2906 | 1.60E-05 | 0.0408 | 0.0108 | 1.51E-04 | 1 |
| rs2022464 | 6 | 108945370 | 6 | 108861264 | 109019323 | A | C | 0.3648 | FOXO3 | intronic | -0.8338 | 0.1483 | 1.88E-08 | -1.1162 | 0.2923 | 1.36E-04 | -0.0437 | 0.0108 | 5.21E-05 | 1 |
| rs10457180 | 6 | 108965039 | 6 | 108861264 | 109019323 | A | G | 0.3678 | FOXO3 | intronic | 0.8338 | 0.1482 | 1.84E-08 | 1.1101 | 0.2921 | 1.46E-04 | 0.0433 | 0.0108 | 5.94E-05 | 1 |
| rs13217795 | 6 | 108974098 | 6 | 108861264 | 109019323 | T | C | 0.3658 | FOXO3 | intronic | 0.8365 | 0.1484 | 1.73E-08 | 1.0821 | 0.2931 | 2.24E-04 | 0.0430 | 0.0108 | 6.63E-05 | 1 |
| rs4946932 | 6 | 108974746 | 6 | 108861264 | 109019323 | A | C | 0.3668 | FOXO3 | intronic | -0.8408 | 0.1482 | 1.41E-08 | -1.0950 | 0.2924 | 1.83E-04 | -0.0437 | 0.0108 | 5.00E-05 | 1 |
| rs9400239 | 6 | 108977663 | 6 | 108861264 | 109019323 | T | C | 0.3678 | FOXO3 | UTR5 | -0.8456 | 0.1482 | 1.17E-08 | -1.0976 | 0.2925 | 1.76E-04 | -0.0439 | 0.0108 | 4.54E-05 | 1 |
| rs2153960 | 6 | 108988184 | 6 | 108861264 | 109019323 | A | G | 0.3579 | FOXO3 | intronic | 0.8127 | 0.1486 | 4.54E-08 | 0.8810 | 0.2944 | 2.77E-03 | 0.0434 | 0.0108 | 5.58E-05 | 1 |
| rs35396874 | 6 | 108994433 | 6 | 108861264 | 109019323 | T | C | 0.3519 | FOXO3 | intronic | 0.8352 | 0.1495 | 2.34E-08 | 0.8850 | 0.2959 | 2.79E-03 | 0.0421 | 0.0108 | 9.87E-05 | 1 |
| rs9398172 | 6 | 108994826 | 6 | 108861264 | 109019323 | A | G | 0.3519 | FOXO3 | intronic | 0.8303 | 0.1496 | 2.86E-08 | 0.8682 | 0.2963 | 3.40E-03 | 0.0421 | 0.0108 | 9.68E-05 | 1 |
| rs61192764 | 6 | 108995187 | 6 | 108861264 | 109019323 | A | G | 0.1581 | FOXO3 | intronic | 1.1803 | 0.2082 | 1.44E-08 | 1.0491 | 0.4085 | 1.02E-02 | 0.0432 | 0.0150 | 3.99E-03 | 1 |
| rs3800228 | 6 | 108996748 | 6 | 108861264 | 109019323 | T | G | 0.3469 | FOXO3 | intronic | -0.8269 | 0.15 | 3.53E-08 | -0.8599 | 0.2974 | 3.85E-03 | -0.0424 | 0.0108 | 9.24E-05 | 1 |
| rs3800229 | 6 | 108996963 | 6 | 108861264 | 109019323 | T | G | 0.3499 | FOXO3 | intronic | 0.8255 | 0.1496 | 3.45E-08 | 0.8666 | 0.2966 | 3.50E-03 | 0.0427 | 0.0108 | 7.66E-05 | 1 |
| rs9374040 | 6 | 108997435 | 6 | 108861264 | 109019323 | A | G | 0.3499 | FOXO3 | intronic | 0.8241 | 0.1497 | 3.69E-08 | 0.8664 | 0.2967 | 3.51E-03 | 0.0428 | 0.0108 | 7.46E-05 | 1 |
| rs9400240 | 6 | 108997611 | 6 | 108861264 | 109019323 | A | G | 0.3499 | FOXO3 | intronic | 0.8232 | 0.1497 | 3.78E-08 | 0.8671 | 0.2967 | 3.49E-03 | 0.0428 | 0.0108 | 7.56E-05 | 1 |
| rs3800230 | 6 | 108998128 | 6 | 108861264 | 109019323 | T | G | 0.1581 | FOXO3 | intronic | 1.1733 | 0.2084 | 1.79E-08 | 1.0466 | 0.4091 | 1.05E-02 | 0.0433 | 0.0150 | 3.93E-03 | 1 |
| rs1935952 | 6 | 108998905 | 6 | 108861264 | 109019323 | C | G | 0.3509 | FOXO3 | intronic | 0.8331 | 0.1497 | 2.62E-08 | 0.8859 | 0.2967 | 2.84E-03 | 0.0430 | 0.0108 | 6.96E-05 | 1 |
| rs3800232 | 6 | 108998953 | 6 | 108861264 | 109019323 | T | C | 0.1581 | FOXO3 | intronic | -1.1713 | 0.2084 | 1.90E-08 | -1.0413 | 0.4092 | 1.10E-02 | -0.0434 | 0.0150 | 3.85E-03 | 1 |
| rs1935951 | 6 | 108999101 | 6 | 108861264 | 109019323 | A | G | 0.3499 | FOXO3 | intronic | 0.8255 | 0.1497 | 3.51E-08 | 0.8677 | 0.2970 | 3.50E-03 | 0.0429 | 0.0108 | 7.23E-05 | 1 |
| rs1935949 | 6 | 108999287 | 6 | 108861264 | 109019323 | A | G | 0.3499 | FOXO3 | intronic | -0.8263 | 0.1497 | 3.40E-08 | -0.8669 | 0.2970 | 3.53E-03 | -0.0429 | 0.0108 | 7.26E-05 | 1 |
| rs9398173 | 6 | 109000316 | 6 | 108861264 | 109019323 | T | C | 0.3489 | FOXO3 | intronic | -0.8641 | 0.1558 | 2.94E-08 | -0.8602 | 0.2973 | 3.83E-03 | -0.0395 | 0.0113 | 4.64E-04 | 1 |
| rs4946935 | 6 | 109000742 | 6 | 108861264 | 109019323 | A | G | 0.3479 | FOXO3 | intronic | -0.8255 | 0.1502 | 3.86E-08 | -0.8943 | 0.2981 | 2.71E-03 | -0.0429 | 0.0109 | 7.87E-05 | 1 |
| rs4945816 | 6 | 109002042 | 6 | 108861264 | 109019323 | T | C | 0.3499 | FOXO3 | UTR3 | 0.8469 | 0.1504 | 1.79E-08 | 0.8703 | 0.2976 | 3.46E-03 | 0.0441 | 0.0109 | 5.28E-05 | 1 |
| rs4946936 | 6 | 109003321 | 6 | 108861264 | 109019323 | T | C | 0.3499 | FOXO3 | UTR3 | -0.8455 | 0.1503 | 1.87E-08 | -0.8754 | 0.2978 | 3.29E-03 | -0.0436 | 0.0109 | 6.17E-05 | 1 |
| rs9388490 | 6 | 126704795 | 7 | 126648460 | 127369230 | T | C | 0.4414 | CENPW | intergenic | 0.8320 | 0.1386 | 1.95E-09 | 1.5643 | 0.2707 | 7.94E-09 | 0.0545 | 0.0100 | 5.76E-08 | 1 |
| rs11759026 | 6 | 126792095 | 7 | 126648460 | 127369230 | A | G | 0.2545 | MIR588 | intergenic | -1.5906 | 0.1729 | 3.58E-20 | -1.5237 | 0.3193 | 1.87E-06 | -0.0811 | 0.0123 | 4.51E-11 | 1 |
| rs1490384 | 6 | 126851160 | 7 | 126648460 | 127369230 | T | C | 0.493 | MIR588 | intergenic | 0.7445 | 0.1365 | 4.91E-08 | 1.5604 | 0.2669 | 5.25E-09 | 0.0494 | 0.0099 | 5.94E-07 | 1 |
| rs4273712 | 6 | 126964510 | 7 | 126648460 | 127369230 | A | G | 0.2803 | PRELID1P1 | upstream | -1.3614 | 0.157 | 4.34E-18 | -1.3746 | 0.3017 | 5.29E-06 | -0.0807 | 0.0113 | 1.12E-12 | 1 |
| rs1262474 | 6 | 126976071 | 7 | 126648460 | 127369230 | C | G | 0.2416 | PRELID1P1 | intergenic | 0.9648 | 0.1586 | 1.19E-09 | 1.2168 | 0.3060 | 7.06E-05 | 0.0453 | 0.0116 | 9.22E-05 | 1 |
| rs1262476 | 6 | 126986996 | 7 | 126648460 | 127369230 | A | G | 0.2406 | RPS4XP9 | intergenic | -0.9717 | 0.1582 | 8.10E-10 | -1.2103 | 0.3060 | 7.72E-05 | -0.0457 | 0.0116 | 7.90E-05 | 1 |
| rs13212044 | 6 | 127025661 | 7 | 126648460 | 127369230 | T | G | 0.2416 | RPS4XP9 | intergenic | -0.9598 | 0.1581 | 1.28E-09 | -1.2103 | 0.3056 | 7.55E-05 | -0.0470 | 0.0116 | 4.95E-05 | 1 |
| rs10484759 | 6 | 127037042 | 7 | 126648460 | 127369230 | T | C | 0.2316 | RPS4XP9 | intergenic | -0.9684 | 0.1601 | 1.46E-09 | -1.2024 | 0.3106 | 1.09E-04 | -0.0477 | 0.0117 | 4.73E-05 | 1 |
| rs853974 | 6 | 127068983 | 7 | 126648460 | 127369230 | T | C | 0.2753 | RPS4XP9 | intergenic | 1.3524 | 0.1576 | 9.28E-18 | 1.4284 | 0.3043 | 2.74E-06 | 0.0775 | 0.0115 | 1.83E-11 | 1 |
| rs12525327 | 6 | 127107248 | 7 | 126648460 | 127369230 | A | G | 0.2435 | RPS4XP9 | intergenic | 0.9502 | 0.1625 | 4.95E-09 | 1.1350 | 0.3049 | 1.99E-04 | 0.0488 | 0.0122 | 6.74E-05 | 1 |
| rs35848181 | 6 | 127118646 | 7 | 126648460 | 127369230 | A | G | 0.2396 | RPS4XP9 | intergenic | -0.9572 | 0.162 | 3.46E-09 | -1.0865 | 0.3031 | 3.40E-04 | -0.0496 | 0.0122 | 4.93E-05 | 1 |
| rs9375477 | 6 | 127204623 | 7 | 126648460 | 127369230 | A | G | 0.163 | RPS4XP9 | intergenic | -1.2224 | 0.191 | 1.57E-10 | -0.7766 | 0.3722 | 3.70E-02 | -0.0687 | 0.0141 | 1.13E-06 | 1 |
| rs139849708 | 6 | 127369230 | 7 | 126648460 | 127369230 | T | C | 0.02087 | RSPO3 | intergenic | 2.7827 | 0.4746 | 4.53E-09 | 0.3436 | 0.8553 | 6.88E-01 | NA | NA | NA | NA |
| rs149352678 | 7 | 54920906 | 8 | 54909928 | 55015505 | T | C | 0.1203 | SNORA73 | intergenic | 1.3946 | 0.2439 | 1.08E-08 | 2.1007 | 0.4663 | 6.75E-06 | 0.0399 | 0.0173 | 2.08E-02 | 1 |
| rs76928645 | 7 | 54941328 | 8 | 54909928 | 55015505 | T | C | 0.1272 | SNORA73 | intergenic | 1.3466 | 0.2352 | 1.04E-08 | 2.0573 | 0.4440 | 3.66E-06 | 0.0355 | 0.0168 | 3.41E-02 | 1 |
| rs151057105 | 7 | 54944920 | 8 | 54909928 | 55015505 | T | C | 0.1213 | SNORA73 | intergenic | 1.4185 | 0.2442 | 6.32E-09 | 2.1533 | 0.4619 | 3.20E-06 | 0.0347 | 0.0172 | 4.37E-02 | 1 |
| rs74504435 | 7 | 54949256 | 8 | 54909928 | 55015505 | A | G | 0.1213 | SNORA73 | intergenic | -1.3984 | 0.2457 | 1.26E-08 | -2.1608 | 0.4665 | 3.70E-06 | -0.0375 | 0.0173 | 3.04E-02 | 1 |
| rs77126132 | 7 | 54966738 | 8 | 54909928 | 55015505 | A | G | 0.1203 | SNORA73 | intergenic | 1.4570 | 0.2472 | 3.78E-09 | 2.2156 | 0.4656 | 2.00E-06 | 0.0398 | 0.0176 | 2.39E-02 | 1 |
| rs77324770 | 7 | 155772027 | 9 | 155772027 | 155778152 | T | C | 0.05964 | AC021218.2 | intergenic | -1.9547 | 0.3499 | 2.32E-08 | -1.2594 | 0.5246 | 1.64E-02 | NA | NA | NA | NA |
| rs75745313 | 7 | 155778152 | 9 | 155772027 | 155778152 | A | G | 0.05964 | AC021218.2 | intergenic | -1.9833 | 0.3522 | 1.80E-08 | -1.1903 | 0.5218 | 2.26E-02 | NA | NA | NA | NA |
| rs10883848 | 10 | 104973061 | 10 | 104591182 | 105175131 | A | G | 0.34 | ST13P13 | intergenic | 0.8478 | 0.1505 | 1.76E-08 | 0.5639 | 0.2863 | 4.89E-02 | 0.0575 | 0.0109 | 1.53E-07 | 1 |
| rs6584545 | 10 | 104999266 | 10 | 104591182 | 105175131 | A | T | 0.2525 | RPEL1 | intergenic | 0.9854 | 0.1663 | 3.13E-09 | 0.5240 | 0.3136 | 9.48E-02 | 0.0710 | 0.0119 | 2.74E-09 | 0 |
| rs1628768 | 10 | 105012994 | 10 | 104591182 | 105175131 | T | C | 0.2555 | RPEL1 | intergenic | -0.9972 | 0.1662 | 1.99E-09 | -0.5130 | 0.3136 | 1.02E-01 | -0.0723 | 0.0119 | 1.24E-09 | 0 |
| rs2487862 | 10 | 105045477 | 10 | 104591182 | 105175131 | T | C | 0.2535 | INA | intronic | 1.0330 | 0.1746 | 3.32E-09 | 0.5131 | 0.3146 | 1.03E-01 | 0.0758 | 0.0131 | 7.48E-09 | 0 |
| rs2030205 | 10 | 105056473 | 10 | 104591182 | 105175131 | C | G | 0.2545 | PCGF6 | intergenic | 0.9600 | 0.1667 | 8.51E-09 | 0.5014 | 0.3147 | 1.11E-01 | 0.0729 | 0.0120 | 1.38E-09 | 0 |
| rs4420190 | 10 | 105085191 | 10 | 104591182 | 105175131 | T | C | 0.2555 | PCGF6 | intronic | -0.9257 | 0.1665 | 2.71E-08 | -0.5476 | 0.3153 | 8.24E-02 | -0.0712 | 0.0120 | 3.30E-09 | 0 |
| rs72846192 | 10 | 105091459 | 10 | 104591182 | 105175131 | A | C | 0.2545 | PCGF6 | intronic | -0.9323 | 0.1671 | 2.41E-08 | -0.5292 | 0.3153 | 9.34E-02 | -0.0721 | 0.0121 | 2.40E-09 | 0 |
| rs4311992 | 10 | 105120649 | 10 | 104591182 | 105175131 | A | G | 0.2505 | TAF5 | intergenic | 0.9480 | 0.1674 | 1.48E-08 | 0.4899 | 0.3158 | 1.21E-01 | 0.0723 | 0.0122 | 2.94E-09 | 0 |
| rs55948574 | 11 | 17310391 | 11 | 17067849 | 17365209 | T | C | 0.2793 | NUCB2 | intronic | 0.8248 | 0.1502 | 3.96E-08 | 0.0138 | 0.2944 | 9.63E-01 | 0.0031 | 0.0109 | 7.77E-01 | 0 |
| rs3911562 | 11 | 92355990 | 12 | 92317858 | 92558342 | A | T | 0.336 | FAT3 | intronic | 0.7941 | 0.1455 | 4.83E-08 | 0.3092 | 0.2900 | 2.86E-01 | 0.0325 | 0.0103 | 1.69E-03 | 0 |
| rs1791558 | 11 | 92356881 | 12 | 92317858 | 92558342 | A | G | 0.339 | FAT3 | intronic | -0.7920 | 0.1452 | 4.94E-08 | -0.3188 | 0.2897 | 2.71E-01 | -0.0323 | 0.0103 | 1.76E-03 | 0 |
| rs1613721 | 11 | 92357002 | 12 | 92317858 | 92558342 | A | G | 0.339 | FAT3 | intronic | -0.7924 | 0.1452 | 4.85E-08 | -0.3232 | 0.2897 | 2.65E-01 | -0.0323 | 0.0103 | 1.76E-03 | 0 |
| rs56021037 | 11 | 92363215 | 12 | 92317858 | 92558342 | T | C | 0.336 | FAT3 | intronic | -0.7935 | 0.1455 | 4.89E-08 | -0.3066 | 0.2896 | 2.90E-01 | -0.0314 | 0.0104 | 2.41E-03 | 0 |
| rs10830949 | 11 | 92540758 | 12 | 92317858 | 92558342 | T | C | 0.328 | FAT3 | intronic | -0.8014 | 0.1469 | 4.93E-08 | -0.2978 | 0.2919 | 3.08E-01 | -0.0352 | 0.0104 | 7.24E-04 | 0 |
| rs12298541 | 12 | 66306441 | 13 | 66257355 | 66389968 | A | C | 0.341 | HMGA2:AC090673. | intronic | 0.8290 | 0.1477 | 2.00E-08 | 0.7320 | 0.2826 | 9.61E-03 | 0.0612 | 0.0110 | 2.31E-08 | 1 |
| rs343086 | 12 | 66325689 | 13 | 66257355 | 66389968 | T | C | 0.3032 | HMGA2 | intronic | -0.8945 | 0.1501 | 2.53E-09 | -0.7404 | 0.2908 | 1.09E-02 | -0.0533 | 0.0110 | 1.25E-06 | 1 |
| rs1480474 | 12 | 66326943 | 13 | 66257355 | 66389968 | A | G | 0.4563 | HMGA2 | intronic | 0.9000 | 0.1397 | 1.19E-10 | 1.1525 | 0.2698 | 1.97E-05 | 0.0555 | 0.0100 | 3.07E-08 | 1 |

| | | | | | | | | | | | Meta-GWAS | | | UKBB | | | ENIGMA* | | | |
|---|---|---|---|---|---|---|---|---|---|---|---|---|---|---|---|---|---|---|---|---|
| rsID | chr | pos (hg19) | Genomic Locus | Genomic Locus start | Genomic Locus end | A1 | A2 | MAF | Nearest Genes | Function | Beta | SE | P-value | Beta | SE | P-value | Beta | SE | P-value | Replicated** |
| rs10878349 | 12 | 66327632 | 13 | 66257355 | 66389968 | A | G | 0.4563 | HMGA2 | intronic | 1.0999 | 0.1385 | 2.01E-15 | 1.3509 | 0.2701 | 5.88E-07 | 0.0585 | 0.0099 | 3.50E-09 | 1 |
| rs1038196 | 12 | 66343400 | 13 | 66257355 | 66389968 | C | G | 0.4573 | HMGA2 | intronic | -1.0574 | 0.1371 | 1.26E-14 | -1.3437 | 0.2693 | 6.20E-07 | NA | NA | NA | NA |
| rs10784502 | 12 | 66343810 | 13 | 66257355 | 66389968 | T | C | 0.4583 | HMGA2 | intronic | -1.0612 | 0.1372 | 1.02E-14 | -1.3513 | 0.2693 | 5.36E-07 | -0.0571 | 0.0098 | 5.67E-09 | 1 |
| rs1979440 | 12 | 66346624 | 13 | 66257355 | 66389968 | T | C | 0.4433 | HMGA2 | UTR3 | 0.8671 | 0.1402 | 6.20E-10 | 1.0719 | 0.2701 | 7.31E-05 | 0.0504 | 0.0101 | 5.69E-07 | 1 |
| rs7959830 | 12 | 66347368 | 13 | 66257355 | 66389968 | T | G | 0.4513 | HMGA2 | UTR3 | -0.8618 | 0.1391 | 5.80E-10 | -1.1403 | 0.2696 | 2.38E-05 | -0.0534 | 0.0100 | 9.62E-08 | 1 |
| rs1351394 | 12 | 66351826 | 13 | 66257355 | 66389968 | T | C | 0.4503 | HMGA2 | UTR3 | 1.0888 | 0.1384 | 3.63E-15 | 1.3310 | 0.2695 | 8.04E-07 | 0.0570 | 0.0098 | 6.54E-09 | 1 |
| rs7138102 | 12 | 66353891 | 13 | 66257355 | 66389968 | A | G | 0.4354 | HMGA2 | intronic | -0.7934 | 0.1405 | 1.61E-08 | -0.9293 | 0.2722 | 6.44E-04 | -0.0499 | 0.0102 | 9.19E-07 | 1 |
| rs867633 | 12 | 66354911 | 13 | 66257355 | 66389968 | A | G | 0.4354 | HMGA2 | intronic | -0.8061 | 0.1406 | 9.93E-09 | -0.9463 | 0.2722 | 5.11E-04 | -0.0505 | 0.0102 | 6.53E-07 | 1 |
| rs1042725 | 12 | 66358347 | 13 | 66257355 | 66389968 | T | C | 0.4632 | HMGA2 | UTR3 | -1.0669 | 0.1376 | 8.94E-15 | -1.2216 | 0.2694 | 5.89E-06 | -0.0560 | 0.0098 | 1.21E-08 | 1 |
| rs8756 | 12 | 66359752 | 13 | 66257355 | 66389968 | A | C | 0.4473 | HMGA2 | UTR3 | -1.0735 | 0.1373 | 5.30E-15 | -1.5180 | 0.2694 | 1.83E-08 | -0.0557 | 0.0098 | 1.26E-08 | 1 |
| rs7970350 | 12 | 66360164 | 13 | 66257355 | 66389968 | T | C | 0.4632 | HMGA2 | downstream | -1.0625 | 0.1375 | 1.11E-14 | -1.2226 | 0.2695 | 5.82E-06 | -0.0553 | 0.0098 | 1.75E-08 | 1 |
| rs7968902 | 12 | 66363070 | 13 | 66257355 | 66389968 | T | G | 0.4076 | HMGA2 | intergenic | 0.9456 | 0.1397 | 1.30E-11 | 1.1022 | 0.2753 | 6.32E-05 | 0.0547 | 0.0102 | 7.93E-08 | 1 |
| rs61921611 | 12 | 66367726 | 13 | 66257355 | 66389968 | T | C | 0.338 | HMGA2 | intergenic | 0.9190 | 0.1482 | 5.67E-10 | 1.0801 | 0.2936 | 2.37E-04 | 0.0412 | 0.0107 | 1.10E-04 | 1 |
| rs7968682 | 12 | 66371880 | 13 | 66257355 | 66389968 | T | G | 0.4473 | HMGA2 | intergenic | -1.0756 | 0.1379 | 6.32E-15 | -1.5048 | 0.2703 | 2.70E-08 | -0.0559 | 0.0099 | 1.49E-08 | 1 |
| NA | 12 | 66373602 | 13 | 66257355 | 66389968 | T | C | 0.4473 | HMGA2 | intergenic | -1.1921 | 0.2052 | 6.30E-09 | -1.4941 | 0.2709 | 3.60E-08 | -0.0560 | 0.0103 | 5.24E-08 | 1 |
| NA | 12 | 66374247 | 13 | 66257355 | 66389968 | A | G | 0.4473 | HMGA2 | intergenic | 1.3211 | 0.2382 | 2.93E-08 | 1.5142 | 0.2711 | 2.42E-08 | 0.0693 | 0.0115 | 1.88E-09 | 1 |
| rs9669278 | 12 | 66374587 | 13 | 66257355 | 66389968 | T | C | 0.4473 | HMGA2 | intergenic | 1.1151 | 0.1403 | 1.86E-15 | 1.4970 | 0.2708 | 3.38E-08 | 0.0572 | 0.0101 | 1.30E-08 | 1 |
| rs7306710 | 12 | 66376091 | 13 | 66257355 | 66389968 | T | C | 0.4463 | HMGA2 | intergenic | 1.1329 | 0.1405 | 7.56E-16 | 1.5214 | 0.2708 | 2.01E-08 | 0.0573 | 0.0101 | 1.48E-08 | 1 |
| NA | 12 | 66376202 | 13 | 66257355 | 66389968 | C | G | 0.4463 | HMGA2 | intergenic | -1.1938 | 0.2033 | 4.32E-09 | -1.5255 | 0.2717 | 2.06E-08 | NA | NA | NA | NA |
| rs1585897 | 12 | 66383320 | 13 | 66257355 | 66389968 | A | C | 0.4215 | HMGA2 | intergenic | -1.0364 | 0.1405 | 1.60E-13 | -1.2410 | 0.2714 | 4.92E-06 | -0.0471 | 0.0101 | 2.85E-06 | 1 |
| rs7966895 | 12 | 66383843 | 13 | 66257355 | 66389968 | A | G | 0.338 | HMGA2 | intergenic | 0.9027 | 0.1472 | 8.54E-10 | 1.4106 | 0.2844 | 7.23E-07 | 0.0338 | 0.0105 | 1.27E-03 | 1 |
| rs11175990 | 12 | 66386396 | 13 | 66257355 | 66389968 | T | C | 0.2475 | HMGA2 | intergenic | 0.9150 | 0.1668 | 4.15E-08 | 1.3119 | 0.3162 | 3.38E-05 | 0.0409 | 0.0121 | 7.39E-04 | 1 |
| rs10748027 | 12 | 66387621 | 13 | 66257355 | 66389968 | A | T | 0.2952 | HMGA2 | intergenic | 0.8781 | 0.1606 | 4.57E-08 | 0.8900 | 0.2980 | 2.83E-03 | 0.0491 | 0.0127 | 1.08E-04 | 1 |
| rs10400419 | 12 | 66389968 | 13 | 66257355 | 66389968 | T | C | 0.4324 | HMGA2 | intergenic | 0.9381 | 0.14 | 2.09E-11 | 1.3632 | 0.2760 | 8.06E-07 | 0.0455 | 0.0101 | 6.38E-06 | 1 |
| rs35227403 | 12 | 68216239 | 14 | 68216239 | 68225876 | A | T | 0.05368 | RP11-43N5.1 | intergenic | -1.7779 | 0.3182 | 2.31E-08 | -2.2433 | 0.5797 | 1.10E-04 | -0.0760 | 0.0239 | 1.45E-03 | 1 |
| rs28792763 | 15 | 90081905 | 15 | 90081905 | 90208310 | A | G | 0.4632 | RP11-429B14.1 | ncRNA_intronic | 0.8393 | 0.1489 | 1.72E-08 | 0.4019 | 0.2773 | 1.47E-01 | 0.0139 | 0.0110 | 2.08E-01 | 0 |
| rs1596062 | 15 | 90091513 | 15 | 90081905 | 90208310 | A | C | 0.4652 | RP11-429B14.1 | ncRNA_intronic | 0.8061 | 0.1471 | 4.23E-08 | 0.3038 | 0.2729 | 2.66E-01 | 0.0120 | 0.0108 | 2.68E-01 | 0 |
| rs11630039 | 15 | 90123991 | 15 | 90081905 | 90208310 | T | C | 0.4831 | RP11-429B14.1:TIC | ncRNA_intronic | 0.8263 | 0.1383 | 2.30E-09 | 0.3007 | 0.2710 | 2.67E-01 | 0.0085 | 0.0099 | 3.93E-01 | 0 |
| rs893725 | 15 | 90128223 | 15 | 90081905 | 90208310 | A | C | 0.3777 | RP11-429B14.1:TIC | ncRNA_intronic | 0.9023 | 0.14 | 1.15E-10 | 0.5440 | 0.2743 | 4.74E-02 | 0.0116 | 0.0101 | 2.51E-01 | 0 |
| rs11639246 | 15 | 90128834 | 15 | 90081905 | 90208310 | A | G | 0.4851 | RP11-429B14.1:TIC | ncRNA_intronic | 0.8145 | 0.1375 | 3.19E-09 | 0.3128 | 0.2708 | 2.48E-01 | 0.0080 | 0.0099 | 4.18E-01 | 0 |
| rs11629584 | 15 | 90128966 | 15 | 90081905 | 90208310 | T | C | 0.4871 | RP11-429B14.1:TIC | exonic | 0.7978 | 0.1375 | 6.53E-09 | 0.3234 | 0.2705 | 2.32E-01 | 0.0075 | 0.0099 | 4.46E-01 | 0 |
| rs4932136 | 15 | 90132578 | 15 | 90081905 | 90208310 | C | G | 0.4831 | RP11-429B14.1:TIC | ncRNA_intronic | -0.8462 | 0.1393 | 1.24E-09 | -0.3500 | 0.2705 | 1.96E-01 | NA | NA | NA | NA |
| rs35892197 | 15 | 90135172 | 15 | 90081905 | 90208310 | T | C | 0.4821 | RP11-429B14.1:TIC | ncRNA_exonic | 0.8324 | 0.1377 | 1.48E-09 | 0.3459 | 0.2704 | 2.01E-01 | 0.0085 | 0.0098 | 3.91E-01 | 0 |
| rs8032553 | 15 | 90137325 | 15 | 90081905 | 90208310 | A | G | 0.4831 | TICRR | intronic | -0.8395 | 0.1377 | 1.08E-09 | -0.3343 | 0.2708 | 2.17E-01 | -0.0084 | 0.0098 | 3.92E-01 | 0 |
| rs4556765 | 15 | 90139262 | 15 | 90081905 | 90208310 | T | C | 0.4821 | TICRR | intronic | 0.8317 | 0.1376 | 1.51E-09 | 0.3266 | 0.2704 | 2.27E-01 | 0.0076 | 0.0098 | 4.39E-01 | 0 |
| rs11634571 | 15 | 90195712 | 15 | 90081905 | 90208310 | A | G | 0.4881 | KIF7 | intronic | 0.8506 | 0.1415 | 1.85E-09 | 0.2488 | 0.2704 | 3.58E-01 | 0.0102 | 0.0102 | 3.18E-01 | 0 |
| rs11652522 | 17 | 43055579 | 16 | 43055579 | 43055579 | A | C | 0.0835 | CTD-2534I21.9 | intergenic | -1.2996 | 0.2299 | 1.59E-08 | -0.5817 | 0.4502 | 1.96E-01 | -0.0438 | 0.0165 | 7.74E-03 | 0 |
| rs117642368 | 17 | 43399058 | 17 | 43399058 | 44874453 | C | G | 0.1014 | MAP3K14 | intergenic | -1.6616 | 0.2435 | 8.94E-12 | -1.6470 | 0.4427 | 2.01E-04 | -0.0634 | 0.0203 | 1.73E-03 | 1 |
| rs17686238 | 17 | 43417273 | 17 | 43399058 | 44874453 | T | G | 0.1054 | RNA5SP443 | intergenic | -1.9576 | 0.328 | 2.39E-09 | -1.5384 | 0.4397 | 4.70E-04 | -0.0577 | 0.0216 | 7.64E-03 | 1 |
| rs62064594 | 17 | 43460181 | 17 | 43399058 | 44874453 | A | G | 0.1581 | CTB-39G8.2 | intergenic | -2.1586 | 0.2826 | 2.21E-14 | -1.6112 | 0.3883 | 3.39E-05 | -0.0933 | 0.0210 | 8.57E-06 | 1 |
| rs62064595 | 17 | 43460374 | 17 | 43399058 | 44874453 | A | G | 0.16 | ARHGAP27 | intergenic | 2.1324 | 0.2823 | 4.21E-14 | 1.6341 | 0.3859 | 2.33E-05 | 0.0923 | 0.0209 | 1.05E-05 | 1 |
| rs61572747 | 17 | 43460891 | 17 | 43399058 | 44874453 | A | G | 0.2893 | ARHGAP27 | intergenic | 1.7478 | 0.2293 | 2.48E-14 | 1.0532 | 0.3114 | 7.23E-04 | NA | NA | NA | NA |
| rs79724577 | 17 | 43463493 | 17 | 43399058 | 44874453 | A | C | 0.1918 | ARHGAP27 | intergenic | 2.1791 | 0.2466 | 1.00E-18 | 1.9384 | 0.3514 | 3.59E-08 | 0.1150 | 0.0157 | 2.02E-13 | 1 |
| rs4763 | 17 | 43471489 | 17 | 43399058 | 44874453 | A | G | 0.1909 | ARHGAP27 | UTR3 | -2.1628 | 0.2471 | 2.09E-18 | -1.9442 | 0.3500 | 2.89E-08 | -0.1184 | 0.0157 | 4.49E-14 | 1 |
| rs62064597 | 17 | 43473307 | 17 | 43399058 | 44874453 | C | G | 0.1909 | ARHGAP27 | intronic | 2.1355 | 0.2471 | 5.47E-18 | 2.0340 | 0.3503 | 6.70E-09 | 0.1183 | 0.0156 | 3.44E-14 | 1 |
| rs62064598 | 17 | 43474668 | 17 | 43399058 | 44874453 | C | G | 0.1938 | ARHGAP27:CTB-39( | ncRNA_intronic | 2.1222 | 0.2441 | 3.51E-18 | 1.9805 | 0.3504 | 1.65E-08 | 0.1178 | 0.0152 | 7.82E-15 | 1 |
| rs2028078 | 17 | 43475929 | 17 | 43399058 | 44874453 | A | G | 0.1938 | ARHGAP27 | intronic | -2.1143 | 0.2444 | 5.15E-18 | -1.9825 | 0.3503 | 1.59E-08 | -0.1178 | 0.0152 | 7.59E-15 | 1 |
| rs56220387 | 17 | 43476807 | 17 | 43399058 | 44874453 | A | G | 0.1918 | ARHGAP27 | intronic | 2.1123 | 0.2443 | 5.24E-18 | 2.0181 | 0.3509 | 9.30E-09 | 0.1176 | 0.0151 | 8.29E-15 | 1 |
| rs55793500 | 17 | 43479748 | 17 | 43399058 | 44874453 | T | C | 0.1938 | ARHGAP27 | intronic | 2.1184 | 0.244 | 3.91E-18 | 1.9971 | 0.3498 | 1.19E-08 | 0.1179 | 0.0151 | 6.72E-15 | 1 |
| rs62064600 | 17 | 43480701 | 17 | 43399058 | 44874453 | A | G | 0.1938 | ARHGAP27 | intronic | -2.1165 | 0.2439 | 3.98E-18 | -1.9977 | 0.3498 | 1.18E-08 | -0.1181 | 0.0151 | 6.03E-15 | 1 |
| rs56236914 | 17 | 43483551 | 17 | 43399058 | 44874453 | T | C | 0.1938 | ARHGAP27 | intronic | -2.1355 | 0.2437 | 1.89E-18 | -1.9967 | 0.3499 | 1.21E-08 | -0.1182 | 0.0151 | 4.61E-15 | 1 |
| rs62064603 | 17 | 43484496 | 17 | 43399058 | 44874453 | T | C | 0.1938 | ARHGAP27 | intronic | -2.1048 | 0.2431 | 4.81E-18 | -1.9983 | 0.3499 | 1.17E-08 | -0.1183 | 0.0151 | 4.19E-15 | 1 |
| rs73984391 | 17 | 43484598 | 17 | 43399058 | 44874453 | T | G | 0.1938 | ARHGAP27 | intronic | -2.1458 | 0.2453 | 2.17E-18 | -2.0051 | 0.3496 | 1.02E-08 | -0.1177 | 0.0151 | 5.89E-15 | 1 |
| NA | 17 | 43484903 | 17 | 43399058 | 44874453 | T | C | 0.1938 | ARHGAP27 | intronic | 2.3006 | 0.3023 | 2.71E-14 | 2.0937 | 0.3564 | 4.45E-09 | 0.1223 | 0.0161 | 3.11E-14 | 1 |
| rs62064637 | 17 | 43485551 | 17 | 43399058 | 44874453 | T | G | 0.1938 | ARHGAP27 | intronic | 2.1308 | 0.2435 | 2.10E-18 | 2.0054 | 0.3498 | 1.03E-08 | 0.1182 | 0.0151 | 4.13E-15 | 1 |
| rs36078910 | 17 | 43487217 | 17 | 43399058 | 44874453 | A | G | 0.1968 | ARHGAP27 | intronic | -2.1047 | 0.2419 | 3.25E-18 | -2.0332 | 0.3495 | 6.28E-09 | -0.1177 | 0.0150 | 4.30E-15 | 1 |
| rs35389313 | 17 | 43487424 | 17 | 43399058 | 44874453 | T | C | 0.1968 | ARHGAP27 | intronic | 2.1133 | 0.2419 | 2.37E-18 | 2.0346 | 0.3495 | 6.13E-09 | 0.1182 | 0.0150 | 3.29E-15 | 1 |
| rs35327136 | 17 | 43487574 | 17 | 43399058 | 44874453 | A | C | 0.1928 | ARHGAP27 | intronic | -2.1170 | 0.2433 | 3.27E-18 | -2.0809 | 0.3499 | 2.89E-09 | -0.1170 | 0.0150 | 5.58E-15 | 1 |
| rs8071011 | 17 | 43488382 | 17 | 43399058 | 44874453 | T | C | 0.1968 | ARHGAP27 | intronic | -2.1077 | 0.2418 | 2.89E-18 | -2.0378 | 0.3495 | 5.81E-09 | -0.1176 | 0.0150 | 3.95E-15 | 1 |
| rs62064641 | 17 | 43488792 | 17 | 43399058 | 44874453 | T | C | 0.1968 | ARHGAP27 | intronic | 2.1109 | 0.2417 | 2.48E-18 | 2.0301 | 0.3492 | 6.42E-09 | 0.1176 | 0.0150 | 3.72E-15 | 1 |
| rs62064643 | 17 | 43490853 | 17 | 43399058 | 44874453 | A | G | 0.1948 | ARHGAP27 | intronic | -2.1067 | 0.2428 | 4.04E-18 | -2.0747 | 0.3500 | 3.25E-09 | -0.1172 | 0.0149 | 3.56E-15 | 1 |
| rs34063617 | 17 | 43491003 | 17 | 43399058 | 44874453 | A | G | 0.1968 | ARHGAP27 | intronic | -2.1129 | 0.2416 | 2.26E-18 | -2.0406 | 0.3493 | 5.43E-09 | -0.1177 | 0.0149 | 2.61E-15 | 1 |
| rs12940792 | 17 | 43492357 | 17 | 43399058 | 44874453 | T | C | 0.1968 | ARHGAP27 | intronic | -2.1048 | 0.2414 | 2.76E-18 | -2.0422 | 0.3493 | 5.27E-09 | -0.1175 | 0.0149 | 2.51E-15 | 1 |
| rs12947718 | 17 | 43493101 | 17 | 43399058 | 44874453 | A | G | 0.1968 | ARHGAP27 | intronic | -2.0986 | 0.2411 | 3.24E-18 | -2.0426 | 0.3493 | 5.21E-09 | -0.1177 | 0.0148 | 2.11E-15 | 1 |
| rs71373560 | 17 | 43493504 | 17 | 43399058 | 44874453 | T | C | 0.1968 | ARHGAP27 | intronic | -2.1306 | 0.2435 | 2.15E-18 | -2.0429 | 0.3492 | 5.18E-09 | -0.1202 | 0.0152 | 2.28E-15 | 1 |
| rs34915103 | 17 | 43493560 | 17 | 43399058 | 44874453 | C | G | 0.1968 | ARHGAP27 | intronic | 2.1045 | 0.2413 | 2.77E-18 | 2.0430 | 0.3492 | 5.17E-09 | 0.1179 | 0.0148 | 1.58E-15 | 1 |
| rs12942951 | 17 | 43493835 | 17 | 43399058 | 44874453 | T | C | 0.1909 | ARHGAP27 | intronic | 2.1351 | 0.2472 | 5.76E-18 | 2.0433 | 0.3492 | 5.14E-09 | 0.1199 | 0.0157 | 2.09E-14 | 1 |
| rs35884427 | 17 | 43495216 | 17 | 43399058 | 44874453 | A | G | 0.1968 | ARHGAP27 | intronic | -2.1042 | 0.2413 | 2.79E-18 | -2.0430 | 0.3492 | 5.14E-09 | -0.1179 | 0.0148 | 1.48E-15 | 1 |
| NA | 17 | 43495235 | 17 | 43399058 | 44874453 | C | G | 0.1968 | ARHGAP27 | intronic | 2.1056 | 0.2749 | 1.88E-14 | 2.0843 | 0.3509 | 3.00E-09 | 0.1178 | 0.0148 | 1.50E-15 | 1 |
| NA | 17 | 43495420 | 17 | 43399058 | 44874453 | A | T | 0.1948 | ARHGAP27 | intronic | 2.1992 | 0.313 | 2.13E-12 | 2.2590 | 0.3874 | 5.76E-09 | 0.1100 | 0.0188 | 5.12E-09 | 1 |
| rs34286926 | 17 | 43495852 | 17 | 43399058 | 44874453 | A | G | 0.1958 | ARHGAP27 | intronic | 2.0711 | 0.2376 | 2.90E-18 | 2.0733 | 0.3491 | 3.01E-09 | 0.1157 | 0.0145 | 1.51E-15 | 1 |
| rs56378631 | 17 | 43496465 | 17 | 43399058 | 44874453 | T | C | 0.1958 | ARHGAP27 | intronic | 2.0777 | 0.2377 | 2.29E-18 | 2.0745 | 0.3490 | 2.93E-09 | 0.1154 | 0.0145 | 1.52E-15 | 1 |
| rs12946723 | 17 | 43496828 | 17 | 43399058 | 44874453 | T | C | 0.1958 | ARHGAP27 | intronic | -2.0839 | 0.2392 | 2.99E-18 | -2.0748 | 0.3490 | 2.91E-09 | -0.1163 | 0.0146 | 1.35E-15 | 1 |
| rs12952764 | 17 | 43497210 | 17 | 43399058 | 44874453 | A | G | 0.1958 | ARHGAP27 | intronic | -2.0778 | 0.2378 | 2.40E-18 | -2.0752 | 0.3490 | 2.89E-09 | -0.1151 | 0.0144 | 1.61E-15 | 1 |
| rs35420583 | 17 | 43498180 | 17 | 43399058 | 44874453 | T | C | 0.1958 | ARHGAP27 | intronic | -2.1433 | 0.2573 | 7.97E-17 | -2.0759 | 0.3489 | 2.84E-09 | -0.1250 | 0.0163 | 1.54E-14 | 1 |
| rs34792542 | 17 | 43498181 | 17 | 43399058 | 44874453 | A | G | 0.1958 | ARHGAP27 | intronic | 2.1573 | 0.2571 | 4.76E-17 | 2.0893 | 0.3507 | 2.71E-09 | 0.1249 | 0.0163 | 1.54E-14 | 1 |
| rs35519908 | 17 | 43499328 | 17 | 43399058 | 44874453 | T | C | 0.1958 | ARHGAP27 | intronic | 2.0736 | 0.2374 | 2.48E-18 | 2.0736 | 0.3489 | 2.95E-09 | 0.1148 | 0.0144 | 1.54E-15 | 1 |
| rs12942666 | 17 | 43499839 | 17 | 43399058 | 44874453 | A | G | 0.1958 | ARHGAP27 | intronic | 2.0736 | 0.2374 | 2.45E-18 | 2.0773 | 0.3488 | 2.74E-09 | 0.1148 | 0.0144 | 1.52E-15 | 1 |
| rs62064645 | 17 | 43500477 | 17 | 43399058 | 44874453 | A | G | 0.1938 | ARHGAP27 | intronic | -2.0678 | 0.2388 | 4.82E-18 | -2.1035 | 0.3495 | 1.86E-09 | -0.1139 | 0.0143 | 2.00E-15 | 1 |
| rs62064646 | 17 | 43500587 | 17 | 43399058 | 44874453 | A | G | 0.1958 | ARHGAP27 | intronic | -2.0900 | 0.239 | 2.22E-18 | -2.0806 | 0.3485 | 2.51E-09 | -0.1142 | 0.0143 | 1.68E-15 | 1 |
| rs62064647 | 17 | 43500621 | 17 | 43399058 | 44874453 | A | G | 0.1958 | ARHGAP27 | intronic | -2.0720 | 0.2377 | 2.82E-18 | -2.0775 | 0.3488 | 2.73E-09 | -0.1145 | 0.0143 | 1.35E-15 | 1 |
| rs7222389 | 17 | 43501442 | 17 | 43399058 | 44874453 | T | C | 0.1958 | ARHGAP27 | intronic | 2.0669 | 0.237 | 2.73E-18 | 2.0903 | 0.3487 | 2.16E-09 | 0.1143 | 0.0143 | 1.38E-15 | 1 |
| rs55648326 | 17 | 43501591 | 17 | 43399058 | 44874453 | T | C | 0.1958 | ARHGAP27 | intronic | 2.0807 | 0.2383 | 2.53E-18 | 2.0933 | 0.3485 | 1.99E-09 | 0.1139 | 0.0143 | 1.73E-15 | 1 |
| rs56212100 | 17 | 43501940 | 17 | 43399058 | 44874453 | A | G | 0.1968 | ARHGAP27 | intronic | -2.0672 | 0.237 | 2.75E-18 | -2.0902 | 0.3487 | 2.16E-09 | -0.1143 | 0.0143 | 1.39E-15 | 1 |
| NA | 17 | 43502012 | 17 | 43399058 | 44874453 | T | C | 0.214 | ARHGAP27 | intronic | 2.1308 | 0.2734 | 6.57E-15 | NA | NA | NA | 0.1160 | 0.0145 | 1.30E-15 | NA |
| rs62064649 | 17 | 43502111 | 17 | 43399058 | 44874453 | A | G | 0.1958 | ARHGAP27 | intronic | 2.0867 | 0.2371 | 1.37E-18 | 2.0883 | 0.3487 | 2.24E-09 | 0.1141 | 0.0143 | 1.53E-15 | 1 |
| rs35626715 | 17 | 43502241 | 17 | 43399058 | 44874453 | T | C | 0.1958 | ARHGAP27 | intronic | 2.0831 | 0.2376 | 1.82E-18 | 2.0868 | 0.3487 | 2.30E-09 | 0.1145 | 0.0144 | 1.54E-15 | 1 |
| rs62064651 | 17 | 43503000 | 17 | 43399058 | 44874453 | A | G | 0.1938 | ARHGAP27 | intronic | -2.0600 | 0.2382 | 5.21E-18 | -2.1008 | 0.3494 | 1.94E-09 | -0.1138 | 0.0143 | 1.66E-15 | 1 |
| rs76344126 | 17 | 43503284 | 17 | 43399058 | 44874453 | A | G | 0.1938 | ARHGAP27 | intronic | -2.0662 | 0.2389 | 5.27E-18 | -2.0978 | 0.3500 | 2.16E-09 | -0.1137 | 0.0143 | 1.67E-15 | 1 |
| rs7209501 | 17 | 43503294 | 17 | 43399058 | 44874453 | A | C | 0.1948 | ARHGAP27 | intronic | 2.0637 | 0.2372 | 3.32E-18 | 2.0797 | 0.3491 | 2.71E-09 | 0.1139 | 0.0143 | 1.49E-15 | 1 |
| rs12949256 | 17 | 43507297 | 17 | 43399058 | 44874453 | T | C | 0.1938 | ARHGAP27 | exonic | -2.1569 | 0.2441 | 9.79E-19 | -2.1130 | 0.3492 | 1.53E-09 | -0.1133 | 0.0145 | 6.31E-15 | 1 |
| rs7220206 | 17 | 43507403 | 17 | 43399058 | 44874453 | A | G | 0.1938 | ARHGAP27 | exonic | -1.9788 | 0.2367 | 6.32E-17 | -2.1205 | 0.3493 | 1.34E-09 | -0.1127 | 0.0142 | 2.41E-15 | 1 |
| rs7222444 | 17 | 43507649 | 17 | 43399058 | 44874453 | A | G | 0.1938 | ARHGAP27 | UTR5 | 2.0795 | 0.2392 | 3.52E-18 | 2.1140 | 0.3492 | 1.50E-09 | 0.1148 | 0.0143 | 1.21E-15 | 1 |
| rs12939187 | 17 | 43508223 | 17 | 43399058 | 44874453 | T | G | 0.1938 | ARHGAP27 | intronic | -2.0472 | 0.2367 | 5.19E-18 | -2.1171 | 0.3496 | 1.48E-09 | -0.1133 | 0.0142 | 1.43E-15 | 1 |
| rs34018943 | 17 | 43508303 | 17 | 43399058 | 44874453 | A | G | 0.1938 | ARHGAP27 | intronic | 2.0482 | 0.2369 | 5.27E-18 | 2.1144 | 0.3492 | 1.49E-09 | 0.1133 | 0.0142 | 1.35E-15 | 1 |
| rs34104358 | 17 | 43508616 | 17 | 43399058 | 44874453 | T | G | 0.1938 | ARHGAP27 | intronic | -2.0477 | 0.2367 | 5.08E-18 | -2.1161 | 0.3492 | 1.45E-09 | -0.1132 | 0.0142 | 1.43E-15 | 1 |

| | | | | | | | | | | | Meta-GWAS | | | UKBB | | | ENIGMA* | | | |
|---|---|---|---|---|---|---|---|---|---|---|---|---|---|---|---|---|---|---|---|---|
| rsID | chr | pos (hg19) | Genomic Locus | Genomic Locus start | Genomic Locus end | A1 | A2 | MAF | Nearest Genes | Function | Beta | SE | P-value | Beta | SE | P-value | Beta | SE | P-value | Replicated** |
| rs62064652 | 17 | 43509310 | 17 | 43399058 | 44874453 | A | G | 0.1928 | ARHGAP27 | intronic | 2.0797 | 0.244 | 1.56E-17 | 2.1196 | 0.3492 | 1.36E-09 | 0.1185 | 0.0148 | 1.20E-15 | 1 |
| rs62064653 | 17 | 43509316 | 17 | 43399058 | 44874453 | T | C | 0.1928 | ARHGAP27 | intronic | -2.0588 | 0.2385 | 6.09E-18 | -2.1196 | 0.3492 | 1.36E-09 | -0.1147 | 0.0144 | 1.71E-15 | 1 |
| rs56020833 | 17 | 43509778 | 17 | 43399058 | 44874453 | T | C | 0.1938 | ARHGAP27 | intronic | -2.0401 | 0.2387 | 1.27E-17 | -2.1149 | 0.3494 | 1.51E-09 | -0.1137 | 0.0142 | 1.12E-15 | 1 |
| rs55642947 | 17 | 43510187 | 17 | 43399058 | 44874453 | C | G | 0.1938 | ARHGAP27 | UTR5 | -2.0607 | 0.2386 | 5.72E-18 | -2.1169 | 0.3494 | 1.46E-09 | -0.1156 | 0.0140 | 1.91E-16 | 1 |
| rs34465449 | 17 | 43511435 | 17 | 43399058 | 44874453 | A | C | 0.1938 | ARHGAP27 | intronic | -1.9944 | 0.1843 | 2.68E-27 | -2.1249 | 0.3494 | 1.26E-09 | -0.1136 | 0.0142 | 1.04E-15 | 1 |
| rs12946900 | 17 | 43512206 | 17 | 43399058 | 44874453 | A | G | 0.1938 | ARHGAP27 | upstream | 1.9811 | 0.1842 | 5.74E-27 | 2.1347 | 0.3493 | 1.05E-09 | 0.1154 | 0.0140 | 1.77E-16 | 1 |
| rs56168933 | 17 | 43512318 | 17 | 43399058 | 44874453 | A | G | 0.1938 | ARHGAP27 | upstream:downstrea | -2.0304 | 0.1856 | 7.66E-28 | -2.1351 | 0.3493 | 1.04E-09 | -0.1153 | 0.0142 | 5.79E-16 | 1 |
| rs55790407 | 17 | 43512439 | 17 | 43399058 | 44874453 | C | G | 0.1938 | ARHGAP27 | upstream:downstrea | -1.9814 | 0.1843 | 5.73E-27 | -2.1371 | 0.3493 | 1.01E-09 | -0.1153 | 0.0140 | 1.82E-16 | 1 |
| rs11012 | 17 | 43513441 | 17 | 43399058 | 44874453 | T | C | 0.1948 | PLEKHM1 | UTR3 | -1.9828 | 0.184 | 4.48E-27 | -2.1405 | 0.3493 | 9.47E-10 | -0.1157 | 0.0140 | 1.39E-16 | 1 |
| rs9730 | 17 | 43513551 | 17 | 43399058 | 44874453 | C | G | 0.1938 | PLEKHM1 | UTR3 | 1.9820 | 0.1842 | 5.42E-27 | 2.1414 | 0.3493 | 9.32E-10 | 0.1153 | 0.0140 | 1.84E-16 | 1 |
| rs62064654 | 17 | 43513896 | 17 | 43399058 | 44874453 | T | C | 0.1938 | PLEKHM1 | UTR3 | -1.9883 | 0.1852 | 6.69E-27 | -2.1317 | 0.3495 | 1.12E-09 | -0.1155 | 0.0140 | 1.90E-16 | 1 |
| rs62064655 | 17 | 43514954 | 17 | 43399058 | 44874453 | A | G | 0.1938 | PLEKHM1 | UTR3 | -2.0297 | 0.2362 | 8.34E-18 | -2.1406 | 0.3493 | 9.44E-10 | -0.1158 | 0.0140 | 1.59E-16 | 1 |
| NA | 17 | 43515846 | 17 | 43399058 | 44874453 | T | C | 0.1938 | PLEKHM1 | intronic | -2.0105 | 0.267 | 5.05E-14 | -2.1367 | 0.3540 | 1.67E-09 | -0.1158 | 0.0140 | 1.67E-16 | 1 |
| rs36114997 | 17 | 43515885 | 17 | 43399058 | 44874453 | A | G | 0.1938 | PLEKHM1 | intronic | 2.0291 | 0.2365 | 9.49E-18 | 2.1416 | 0.3493 | 9.27E-10 | 0.1158 | 0.0140 | 1.64E-16 | 1 |
| rs568655892 | 17 | 43515927 | 17 | 43399058 | 44874453 | T | C | 0.1948 | PLEKHM1 | intronic | -2.0261 | 0.2361 | 9.37E-18 | -2.1416 | 0.3493 | 9.27E-10 | -0.1156 | 0.0141 | 2.12E-16 | 1 |
| rs17631303 | 17 | 43516402 | 17 | 43399058 | 44874453 | A | G | 0.1938 | PLEKHM1 | intronic | 1.9763 | 0.1848 | 1.11E-26 | 2.1348 | 0.3494 | 1.06E-09 | 0.1168 | 0.0141 | 1.08E-16 | 1 |
| NA | 17 | 43516739 | 17 | 43399058 | 44874453 | A | G | 0.1938 | PLEKHM1 | intronic | -2.0653 | 0.2706 | 2.33E-14 | -2.1534 | 0.3498 | 7.94E-10 | -0.1176 | 0.0142 | 1.52E-16 | 1 |
| rs62064657 | 17 | 43517054 | 17 | 43399058 | 44874453 | A | G | 0.1938 | PLEKHM1 | intronic | -2.0409 | 0.2382 | 1.05E-17 | -2.1375 | 0.3494 | 1.01E-09 | -0.1167 | 0.0141 | 1.25E-16 | 1 |
| rs35489312 | 17 | 43517252 | 17 | 43399058 | 44874453 | T | C | 0.1938 | PLEKHM1 | intronic | 1.9734 | 0.1843 | 9.41E-27 | 2.1452 | 0.3493 | 8.70E-10 | 0.1166 | 0.0141 | 1.30E-16 | 1 |
| NA | 17 | 43519564 | 17 | 43399058 | 44874453 | A | C | 0.1938 | PLEKHM1 | intronic | -2.0102 | 0.2674 | 5.56E-14 | -2.1445 | 0.3498 | 9.33E-10 | -0.1169 | 0.0141 | 1.25E-16 | 1 |
| rs62065374 | 17 | 43520272 | 17 | 43399058 | 44874453 | A | G | 0.1938 | PLEKHM1 | intronic | -2.0427 | 0.2386 | 1.13E-17 | -2.1438 | 0.3495 | 9.09E-10 | -0.1170 | 0.0141 | 1.25E-16 | 1 |
| rs62065376 | 17 | 43521161 | 17 | 43399058 | 44874453 | C | G | 0.1948 | PLEKHM1 | intronic | 2.0413 | 0.2382 | 1.03E-17 | 2.1393 | 0.3495 | 9.84E-10 | 0.1173 | 0.0141 | 1.04E-16 | 1 |
| rs62065377 | 17 | 43521193 | 17 | 43399058 | 44874453 | A | G | 0.1948 | PLEKHM1 | intronic | 2.0992 | 0.2448 | 9.89E-18 | 2.1497 | 0.3493 | 8.04E-10 | 0.1206 | 0.0150 | 1.03E-15 | 1 |
| rs62065378 | 17 | 43522361 | 17 | 43399058 | 44874453 | T | C | 0.1938 | PLEKHM1 | intronic | 1.9910 | 0.1847 | 4.34E-27 | 2.1475 | 0.3494 | 8.42E-10 | 0.1178 | 0.0142 | 1.09E-16 | 1 |
| rs113575082 | 17 | 43524526 | 17 | 43399058 | 44874453 | T | C | 0.1938 | PLEKHM1 | intronic | -2.0675 | 0.2355 | 1.64E-18 | -2.1395 | 0.3495 | 9.88E-10 | -0.1181 | 0.0142 | 8.25E-17 | 1 |
| rs71373572 | 17 | 43525022 | 17 | 43399058 | 44874453 | A | G | 0.1938 | PLEKHM1 | intronic | -2.0574 | 0.2341 | 1.53E-18 | -2.1541 | 0.3494 | 7.45E-10 | -0.1181 | 0.0142 | 9.09E-17 | 1 |
| rs62065379 | 17 | 43525365 | 17 | 43399058 | 44874453 | T | C | 0.1938 | PLEKHM1 | intronic | -2.0818 | 0.2362 | 1.19E-18 | -2.1440 | 0.3496 | 9.12E-10 | -0.1182 | 0.0142 | 8.84E-17 | 1 |
| NA | 17 | 43527025 | 17 | 43399058 | 44874453 | T | C | 0.1938 | PLEKHM1 | intronic | -2.0972 | 0.2714 | 1.11E-14 | -2.1415 | 0.3500 | 1.00E-09 | -0.1182 | 0.0142 | 8.25E-17 | 1 |
| rs112538459 | 17 | 43527323 | 17 | 43399058 | 44874453 | T | C | 0.1938 | PLEKHM1 | intronic | -1.9999 | 0.1846 | 2.36E-27 | -2.1557 | 0.3494 | 7.29E-10 | -0.1181 | 0.0142 | 8.79E-17 | 1 |
| rs2077606 | 17 | 43529293 | 17 | 43399058 | 44874453 | A | G | 0.1938 | PLEKHM1 | intronic | -2.0799 | 0.2373 | 1.88E-18 | -2.1613 | 0.3494 | 6.56E-10 | -0.1186 | 0.0142 | 7.75E-17 | 1 |
| rs2960000 | 17 | 43534353 | 17 | 43399058 | 44874453 | T | C | 0.1978 | PLEKHM1:AC09113 | ncRNA_intronic | 2.1462 | 0.2522 | 1.76E-17 | 2.0753 | 0.3483 | 2.69E-09 | 0.1249 | 0.0148 | 3.60E-17 | 1 |
| rs62065385 | 17 | 43534694 | 17 | 43399058 | 44874453 | T | C | 0.1958 | PLEKHM1:AC09113 | ncRNA_intronic | -2.0975 | 0.2404 | 2.68E-18 | -2.1603 | 0.3495 | 6.79E-10 | -0.1199 | 0.0143 | 5.32E-17 | 1 |
| rs55703888 | 17 | 43536408 | 17 | 43399058 | 44874453 | T | C | 0.1938 | PLEKHM1:AC09113 | ncRNA_intronic | -2.0886 | 0.2396 | 2.81E-18 | -2.1695 | 0.3493 | 5.63E-10 | -0.1198 | 0.0143 | 5.66E-17 | 1 |
| rs56005713 | 17 | 43536743 | 17 | 43399058 | 44874453 | T | C | 0.1938 | PLEKHM1:AC09113 | ncRNA_intronic | -2.0125 | 0.1858 | 2.47E-27 | -2.1698 | 0.3493 | 5.61E-10 | -0.1200 | 0.0143 | 5.01E-17 | 1 |
| rs111423688 | 17 | 43538523 | 17 | 43399058 | 44874453 | A | G | 0.1938 | PLEKHM1:AC09113 | ncRNA_intronic | -2.1113 | 0.242 | 2.64E-18 | -2.1651 | 0.3495 | 6.20E-10 | -0.1205 | 0.0143 | 4.14E-17 | 1 |
| rs62065389 | 17 | 43538807 | 17 | 43399058 | 44874453 | T | G | 0.1938 | PLEKHM1:AC09113 | ncRNA_intronic | 2.1054 | 0.2433 | 5.05E-18 | 2.1650 | 0.3495 | 6.20E-10 | 0.1209 | 0.0143 | 3.46E-17 | 1 |
| NA | 17 | 43538991 | 17 | 43399058 | 44874453 | A | G | 0.1938 | PLEKHM1:AC09113 | ncRNA_intronic | -2.1131 | 0.2783 | 3.13E-14 | -2.1548 | 0.3499 | 7.78E-10 | -0.1216 | 0.0144 | 3.41E-17 | 1 |
| NA | 17 | 43538993 | 17 | 43399058 | 44874453 | C | G | 0.1938 | PLEKHM1:AC09113 | ncRNA_intronic | -2.1278 | 0.2793 | 2.57E-14 | -2.1548 | 0.3499 | 7.78E-10 | -0.1217 | 0.0145 | 4.25E-17 | 1 |
| NA | 17 | 43539035 | 17 | 43399058 | 44874453 | A | G | 0.1938 | PLEKHM1:AC09113 | ncRNA_intronic | -2.1146 | 0.2785 | 3.13E-14 | -2.1549 | 0.3499 | 7.76E-10 | -0.1215 | 0.0144 | 3.39E-17 | 1 |
| NA | 17 | 43539437 | 17 | 43399058 | 44874453 | C | G | 0.1938 | PLEKHM1:AC09113 | ncRNA_intronic | 2.3295 | 0.3204 | 3.56E-13 | 2.1447 | 0.3500 | 9.46E-10 | 0.1345 | 0.0161 | 7.08E-17 | 1 |
| NA | 17 | 43539723 | 17 | 43399058 | 44874453 | T | C | 0.1938 | PLEKHM1:AC09113 | ncRNA_intronic | -2.3142 | 0.3745 | 6.46E-10 | -2.1554 | 0.3499 | 7.68E-10 | NA | NA | NA | NA |
| rs77099723 | 17 | 43539968 | 17 | 43399058 | 44874453 | A | G | 0.1938 | PLEKHM1:AC09113 | ncRNA_intronic | -2.1333 | 0.2501 | 1.45E-17 | -2.1739 | 0.3493 | 5.18E-10 | -0.1234 | 0.0145 | 1.81E-17 | 1 |
| NA | 17 | 43540472 | 17 | 43399058 | 44874453 | A | T | 0.1938 | PLEKHM1:AC09113 | ncRNA_intronic | 2.3531 | 0.3419 | 5.91E-12 | 2.1626 | 0.3531 | 9.61E-10 | 0.1349 | 0.0174 | 8.65E-15 | 1 |
| rs2139890 | 17 | 43541627 | 17 | 43399058 | 44874453 | A | C | 0.1938 | PLEKHM1 | intronic | -2.0469 | 0.1879 | 1.27E-27 | -2.1734 | 0.3493 | 5.24E-10 | -0.1220 | 0.0144 | 2.63E-17 | 1 |
| rs3946526 | 17 | 43541656 | 17 | 43399058 | 44874453 | T | C | 0.1938 | PLEKHM1 | intronic | -2.1552 | 0.2475 | 3.10E-18 | -2.1638 | 0.3495 | 6.34E-10 | -0.1235 | 0.0146 | 2.97E-17 | 1 |
| NA | 17 | 43544206 | 17 | 43399058 | 44874453 | C | G | 0.1958 | PLEKHM1 | intronic | -2.3996 | 0.3678 | 6.87E-11 | -2.1601 | 0.3486 | 6.17E-10 | -0.1463 | 0.0191 | 1.95E-14 | 1 |
| rs55663797 | 17 | 43544379 | 17 | 43399058 | 44874453 | A | G | 0.1958 | PLEKHM1 | intronic | -2.0504 | 0.1882 | 1.26E-27 | -2.1607 | 0.3484 | 5.96E-10 | -0.1220 | 0.0144 | 2.05E-17 | 1 |
| rs1879581 | 17 | 43545893 | 17 | 43399058 | 44874453 | T | C | 0.1958 | PLEKHM1 | exonic | 2.0594 | 0.1893 | 1.48E-27 | 2.1633 | 0.3485 | 5.72E-10 | 0.1186 | 0.0147 | 8.18E-16 | 1 |
| NA | 17 | 43546057 | 17 | 43399058 | 44874453 | T | G | 0.1958 | PLEKHM1 | intronic | -2.2274 | 0.2866 | 7.68E-15 | -2.1539 | 0.3486 | 6.85E-10 | -0.1184 | 0.0147 | 9.68E-16 | 1 |
| rs55671319 | 17 | 43548424 | 17 | 43399058 | 44874453 | A | G | 0.1958 | PLEKHM1 | intronic | 2.0872 | 0.1934 | 3.67E-27 | 2.1564 | 0.3487 | 6.66E-10 | 0.1207 | 0.0153 | 2.84E-15 | 1 |
| rs55652155 | 17 | 43548481 | 17 | 43399058 | 44874453 | A | G | 0.1958 | PLEKHM1 | intronic | 2.2426 | 0.256 | 1.97E-18 | 2.1669 | 0.3486 | 5.42E-10 | 0.1206 | 0.0153 | 2.93E-15 | 1 |
| rs17631676 | 17 | 43549526 | 17 | 43399058 | 44874453 | A | G | 0.1958 | PLEKHM1 | intronic | 2.0748 | 0.1931 | 6.47E-27 | 2.1682 | 0.3486 | 5.33E-10 | 0.1214 | 0.0151 | 7.12E-16 | 1 |
| rs149366495 | 17 | 43549608 | 17 | 43399058 | 44874453 | A | G | 0.1958 | PLEKHM1 | intronic | -2.0695 | 0.1936 | 1.15E-26 | -2.1610 | 0.3488 | 6.17E-10 | -0.1219 | 0.0151 | 7.37E-16 | 1 |
| rs2090847 | 17 | 43550107 | 17 | 43399058 | 44874453 | A | G | 0.1958 | PLEKHM1 | intronic | 2.0868 | 0.1949 | 9.63E-27 | 2.1688 | 0.3487 | 5.29E-10 | 0.1235 | 0.0154 | 1.18E-15 | 1 |
| rs55746869 | 17 | 43551083 | 17 | 43399058 | 44874453 | T | C | 0.1958 | PLEKHM1 | intronic | -2.2435 | 0.2604 | 7.01E-18 | -2.1574 | 0.3490 | 6.78E-10 | -0.1256 | 0.0156 | 6.83E-16 | 1 |
| rs12950965 | 17 | 43551151 | 17 | 43399058 | 44874453 | C | G | 0.1958 | PLEKHM1 | intronic | 2.2886 | 0.2658 | 7.35E-18 | 2.1700 | 0.3487 | 5.21E-10 | 0.1276 | 0.0160 | 1.70E-15 | 1 |
| rs56015792 | 17 | 43551321 | 17 | 43399058 | 44874453 | C | G | 0.1958 | PLEKHM1 | intronic | -2.2303 | 0.2578 | 5.04E-18 | -2.1702 | 0.3487 | 5.19E-10 | -0.1236 | 0.0153 | 7.58E-16 | 1 |
| NA | 17 | 43551523 | 17 | 43399058 | 44874453 | T | C | 0.1958 | PLEKHM1 | intronic | 2.2841 | 0.304 | 5.77E-14 | 2.2021 | 0.3498 | 3.26E-10 | 0.1260 | 0.0157 | 1.01E-15 | 1 |
| NA | 17 | 43551546 | 17 | 43399058 | 44874453 | T | C | 0.1958 | PLEKHM1 | intronic | -2.2921 | 0.3047 | 5.38E-14 | -2.1878 | 0.3500 | 4.36E-10 | -0.1265 | 0.0157 | 8.76E-16 | 1 |
| rs62065404 | 17 | 43551613 | 17 | 43399058 | 44874453 | T | C | 0.1958 | PLEKHM1 | intronic | -2.2357 | 0.2599 | 7.91E-18 | -2.1705 | 0.3487 | 5.17E-10 | -0.1268 | 0.0157 | 8.21E-16 | 1 |
| rs71238846 | 17 | 43552537 | 17 | 43399058 | 44874453 | A | G | 0.1958 | PLEKHM1 | exonic | -2.2373 | 0.2602 | 8.09E-18 | -2.1714 | 0.3488 | 5.11E-10 | -0.1271 | 0.0158 | 8.11E-16 | 1 |
| rs147243132 | 17 | 43552717 | 17 | 43399058 | 44874453 | C | G | 0.1958 | PLEKHM1 | exonic | -2.3170 | 0.2628 | 1.18E-18 | -2.1716 | 0.3488 | 5.09E-10 | -0.1281 | 0.0160 | 1.01E-15 | 1 |
| rs12452273 | 17 | 43552921 | 17 | 43399058 | 44874453 | T | C | 0.1958 | PLEKHM1 | exonic | -2.3051 | 0.2637 | 2.27E-18 | -2.1718 | 0.3488 | 5.08E-10 | -0.1287 | 0.0161 | 1.29E-15 | 1 |
| rs56192752 | 17 | 43555253 | 17 | 43399058 | 44874453 | A | G | 0.1958 | PLEKHM1 | UTR3 | 2.1184 | 0.196 | 3.18E-27 | 2.1736 | 0.3488 | 4.93E-10 | 0.1293 | 0.0161 | 8.69E-16 | 1 |
| rs62065436 | 17 | 43556652 | 17 | 43399058 | 44874453 | A | G | 0.1958 | PLEKHM1 | intronic | -2.3218 | 0.2686 | 5.42E-18 | -2.1666 | 0.3490 | 5.72E-10 | -0.1297 | 0.0162 | 1.09E-15 | 1 |
| rs55925547 | 17 | 43556807 | 17 | 43399058 | 44874453 | T | C | 0.1978 | PLEKHM1 | intronic | 2.3578 | 0.269 | 1.85E-18 | 2.1511 | 0.3480 | 6.77E-10 | 0.1316 | 0.0163 | 6.62E-16 | 1 |
| NA | 17 | 43556862 | 17 | 43399058 | 44874453 | T | C | 0.1958 | PLEKHM1 | intronic | -2.5074 | 0.3345 | 6.61E-14 | -2.1530 | 0.3487 | 7.05E-10 | -0.1341 | 0.0166 | 7.15E-16 | 1 |
| rs55643511 | 17 | 43556982 | 17 | 43399058 | 44874453 | A | G | 0.1958 | PLEKHM1 | intronic | -2.3190 | 0.2685 | 5.79E-18 | -2.1740 | 0.3489 | 4.93E-10 | -0.1313 | 0.0163 | 7.76E-16 | 1 |
| rs62065437 | 17 | 43557612 | 17 | 43399058 | 44874453 | A | G | 0.1958 | PLEKHM1 | intronic | -2.3195 | 0.2686 | 5.92E-18 | -2.1742 | 0.3489 | 4.92E-10 | -0.1314 | 0.0163 | 8.25E-16 | 1 |
| rs62065438 | 17 | 43558092 | 17 | 43399058 | 44874453 | T | C | 0.1958 | PLEKHM1 | intronic | -2.1191 | 0.1993 | 2.10E-26 | -2.1667 | 0.3490 | 5.73E-10 | -0.1323 | 0.0164 | 6.90E-16 | 1 |
| NA | 17 | 43563093 | 17 | 43399058 | 44874453 | A | G | 0.1958 | PLEKHM1 | intronic | 2.5529 | 0.3565 | 7.96E-13 | 2.1622 | 0.3489 | 6.14E-10 | 0.1431 | 0.0176 | 5.13E-16 | 1 |
| rs62065442 | 17 | 43563894 | 17 | 43399058 | 44874453 | T | C | 0.1958 | PLEKHM1 | intronic | 2.3014 | 0.2859 | 8.29E-16 | 2.1761 | 0.3490 | 4.80E-10 | 0.1355 | 0.0169 | 1.25E-15 | 1 |
| rs144733372 | 17 | 43564222 | 17 | 43399058 | 44874453 | T | G | 0.1958 | PLEKHM1 | intronic | 2.2869 | 0.2865 | 1.45E-15 | 2.1747 | 0.3489 | 4.88E-10 | 0.1360 | 0.0170 | 1.13E-15 | 1 |
| rs62065445 | 17 | 43565840 | 17 | 43399058 | 44874453 | A | G | 0.1958 | PLEKHM1 | intronic | -2.3002 | 0.2869 | 1.08E-15 | -2.1791 | 0.3490 | 4.54E-10 | -0.1368 | 0.0171 | 1.13E-15 | 1 |
| rs1879585 | 17 | 43567175 | 17 | 43399058 | 44874453 | T | G | 0.1968 | PLEKHM1 | intronic | 2.3038 | 0.2879 | 1.21E-15 | 2.1589 | 0.3493 | 6.77E-10 | 0.1379 | 0.0172 | 1.21E-15 | 1 |
| NA | 17 | 43568928 | 17 | 43399058 | 44874453 | A | T | 0.1958 | PLEKHM1 | upstream | -2.3919 | 0.3549 | 1.58E-11 | -2.1487 | 0.3490 | 7.93E-10 | -0.1414 | 0.0176 | 1.01E-15 | 1 |
| NA | 17 | 43569083 | 17 | 43399058 | 44874453 | T | C | 0.1988 | PLEKHM1 | upstream | -2.3911 | 0.3513 | 9.99E-12 | -2.1430 | 0.3487 | 8.47E-10 | -0.1403 | 0.0174 | 8.88E-16 | 1 |
| rs113322852 | 17 | 43569245 | 17 | 43399058 | 44874453 | A | T | 0.1988 | PLEKHM1 | intergenic | -2.1015 | 0.2058 | 1.74E-24 | -2.1731 | 0.3494 | 5.30E-10 | -0.1369 | 0.0177 | 1.21E-14 | 1 |
| rs62065448 | 17 | 43569770 | 17 | 43399058 | 44874453 | T | C | 0.1998 | PLEKHM1 | intergenic | -2.3205 | 0.2893 | 1.05E-15 | -2.1505 | 0.3509 | 9.38E-10 | -0.1371 | 0.0179 | 1.72E-14 | 1 |
| rs62065449 | 17 | 43569909 | 17 | 43399058 | 44874453 | T | C | 0.1988 | PLEKHM1 | intergenic | 2.3158 | 0.2886 | 1.02E-15 | 2.1598 | 0.3491 | 6.54E-10 | 0.1374 | 0.0180 | 2.01E-14 | 1 |
| rs1879583 | 17 | 43570893 | 17 | 43399058 | 44874453 | T | C | 0.1988 | PLEKHM1 | intergenic | -2.1765 | 0.3035 | 7.41E-13 | -2.1595 | 0.3491 | 6.57E-10 | -0.1393 | 0.0187 | 9.34E-14 | 1 |
| rs1724390 | 17 | 43663247 | 17 | 43399058 | 44874453 | A | C | 0.2435 | DND1P1 | ncRNA_exonic | -1.8194 | 0.2047 | 6.10E-19 | -2.0648 | 0.3208 | 1.31E-10 | NA | NA | NA | NA |
| rs117368197 | 17 | 43715924 | 17 | 43399058 | 44874453 | A | G | 0.2396 | CRHR1-IT1:CRHR1:I | ncRNA_intronic | 1.8520 | 0.1976 | 7.08E-21 | 2.2034 | 0.3214 | 7.80E-12 | NA | NA | NA | NA |
| rs111273167 | 17 | 43740967 | 17 | 43399058 | 44874453 | T | C | 0.2396 | CRHR1:RP11-105N1 | ncRNA_intronic | -1.7940 | 0.1964 | 6.50E-20 | -2.2033 | 0.3214 | 7.80E-12 | NA | NA | NA | NA |
| rs112155389 | 17 | 43758078 | 17 | 43399058 | 44874453 | A | G | 0.2396 | CRHR1:RP11-105N1 | ncRNA_intronic | -1.8394 | 0.1957 | 5.43E-21 | -2.2034 | 0.3214 | 7.80E-12 | NA | NA | NA | NA |
| rs112431991 | 17 | 43758885 | 17 | 43399058 | 44874453 | A | G | 0.2396 | CRHR1:RP11-105N1 | ncRNA_intronic | 1.8499 | 0.1965 | 4.68E-21 | 2.1833 | 0.3233 | 1.58E-11 | NA | NA | NA | NA |
| rs113661667 | 17 | 43791610 | 17 | 43399058 | 44874453 | T | C | 0.2396 | CRHR1:RP11-105N1 | ncRNA_intronic | 1.7881 | 0.1959 | 7.05E-20 | 2.2118 | 0.3216 | 6.71E-12 | NA | NA | NA | NA |
| rs111415173 | 17 | 43795573 | 17 | 43399058 | 44874453 | T | C | 0.2396 | CRHR1:RP11-105N1 | ncRNA_intronic | -1.7533 | 0.196 | 3.73E-19 | -2.2115 | 0.3216 | 6.75E-12 | NA | NA | NA | NA |
| rs113991678 | 17 | 43795634 | 17 | 43399058 | 44874453 | T | C | 0.2386 | CRHR1:RP11-105N1 | ncRNA_intronic | -1.7536 | 0.1961 | 3.81E-19 | -2.2115 | 0.3216 | 6.75E-12 | NA | NA | NA | NA |
| rs112995313 | 17 | 43795768 | 17 | 43399058 | 44874453 | T | C | 0.2396 | CRHR1:RP11-105N1 | ncRNA_intronic | 1.7436 | 0.1961 | 5.96E-19 | 2.2115 | 0.3216 | 6.75E-12 | NA | NA | NA | NA |
| rs117615688 | 17 | 43798308 | 17 | 43399058 | 44874453 | A | G | 0.07058 | CRHR1:RP11-105N1 | ncRNA_intronic | -1.9423 | 0.3549 | 4.43E-08 | -1.9337 | 0.5391 | 3.37E-04 | NA | NA | NA | NA |
| rs112570965 | 17 | 43801092 | 17 | 43399058 | 44874453 | T | C | 0.2396 | CRHR1:RP11-105N1 | ncRNA_intronic | 1.7962 | 0.1962 | 5.36E-20 | 2.2111 | 0.3216 | 6.80E-12 | NA | NA | NA | NA |
| rs113790915 | 17 | 43810782 | 17 | 43399058 | 44874453 | C | G | 0.2396 | CRHR1:RP11-105N1 | ncRNA_exonic | -1.8211 | 0.1952 | 1.06E-20 | -2.2124 | 0.3216 | 6.59E-12 | NA | NA | NA | NA |
| rs62055869 | 17 | 43830685 | 17 | 43399058 | 44874453 | A | G | 0.2396 | CRHR1:RP11-105N1 | ncRNA_intronic | -1.7872 | 0.1947 | 4.35E-20 | -2.2130 | 0.3216 | 6.49E-12 | NA | NA | NA | NA |
| rs17426174 | 17 | 43830938 | 17 | 43399058 | 44874453 | C | G | 0.2396 | CRHR1:RP11-105N1 | ncRNA_intronic | -1.7870 | 0.1947 | 4.39E-20 | -2.2135 | 0.3216 | 6.42E-12 | NA | NA | NA | NA |
| rs62055885 | 17 | 43838014 | 17 | 43399058 | 44874453 | A | G | 0.2396 | CRHR1:RP11-105N1 | ncRNA_intronic | 2.0658 | 0.2631 | 4.07E-15 | 2.2165 | 0.3217 | 6.15E-12 | 0.1214 | 0.0181 | 1.88E-11 | 1 |

| | | | | | | | | | | | Meta-GWAS | | | UKBB | | | ENIGMA* | | | |
|---|---|---|---|---|---|---|---|---|---|---|---|---|---|---|---|---|---|---|---|---|
| rsID | chr | pos (hg19) | Genomic Locus | Genomic Locus start | Genomic Locus end | A1 | A2 | MAF | Nearest Genes | Function | Beta | SE | P-value | Beta | SE | P-value | Beta | SE | P-value | Replicated** |
| rs28439278 | 17 | 43838071 | 17 | 43399058 | 44874453 | T | C | 0.2396 | CRHR1:RP11-105N1 | ncRNA_intronic | -2.0653 | 0.263 | 4.12E-15 | -2.2147 | 0.3216 | 6.28E-12 | -0.1213 | 0.0181 | 1.88E-11 | 1 |
| rs12150621 | 17 | 43838482 | 17 | 43399058 | 44874453 | T | G | 0.2396 | CRHR1:RP11-105N1 | ncRNA_intronic | -2.0640 | 0.2627 | 3.89E-15 | -2.2146 | 0.3216 | 6.33E-12 | -0.1185 | 0.0178 | 3.06E-11 | 1 |
| rs62055886 | 17 | 43838678 | 17 | 43399058 | 44874453 | A | G | 0.2396 | CRHR1:RP11-105N1 | ncRNA_intronic | -2.0627 | 0.2631 | 4.48E-15 | -2.2080 | 0.3218 | 7.47E-12 | -0.1183 | 0.0178 | 3.13E-11 | 1 |
| rs62055887 | 17 | 43838710 | 17 | 43399058 | 44874453 | T | C | 0.2396 | CRHR1:RP11-105N1 | ncRNA_intronic | 2.0626 | 0.2631 | 4.48E-15 | 2.2080 | 0.3218 | 7.49E-12 | 0.1213 | 0.0176 | 5.73E-12 | 1 |
| rs62055888 | 17 | 43838720 | 17 | 43399058 | 44874453 | T | C | 0.2366 | CRHR1:RP11-105N1 | ncRNA_intronic | -2.0513 | 0.2644 | 8.58E-15 | -2.1850 | 0.3243 | 1.76E-11 | -0.1193 | 0.0176 | 1.16E-11 | 1 |
| rs62055889 | 17 | 43838919 | 17 | 43399058 | 44874453 | A | G | 0.2396 | CRHR1:RP11-105N1 | ncRNA_intronic | -2.0635 | 0.2635 | 4.85E-15 | -2.2147 | 0.3216 | 6.30E-12 | -0.1221 | 0.0174 | 2.09E-12 | 1 |
| rs71375313 | 17 | 43839253 | 17 | 43399058 | 44874453 | A | G | 0.2396 | CRHR1:RP11-105N1 | ncRNA_intronic | -2.0607 | 0.263 | 4.64E-15 | -2.2183 | 0.3219 | 6.06E-12 | -0.1219 | 0.0173 | 2.05E-12 | 1 |
| rs11079718 | 17 | 43839951 | 17 | 43399058 | 44874453 | A | T | 0.2435 | CRHR1:RP11-105N1 | ncRNA_intronic | 2.2439 | 0.2612 | 8.74E-18 | 2.2498 | 0.3222 | 3.20E-12 | 0.1243 | 0.0176 | 1.68E-12 | 1 |
| rs11079719 | 17 | 43840006 | 17 | 43399058 | 44874453 | T | G | 0.2435 | CRHR1:RP11-105N1 | ncRNA_intronic | 2.1870 | 0.2595 | 3.54E-17 | 2.2218 | 0.3239 | 7.56E-12 | 0.1227 | 0.0175 | 2.65E-12 | 1 |
| rs11079720 | 17 | 43840016 | 17 | 43399058 | 44874453 | A | G | 0.2386 | CRHR1:RP11-105N1 | ncRNA_intronic | -2.1530 | 0.2593 | 1.02E-16 | -2.1971 | 0.3243 | 1.35E-11 | -0.1211 | 0.0172 | 1.74E-12 | 1 |
| rs11079721 | 17 | 43840107 | 17 | 43399058 | 44874453 | A | C | 0.2396 | CRHR1:RP11-105N1 | ncRNA_intronic | -2.1517 | 0.2585 | 8.59E-17 | -2.2300 | 0.3219 | 4.73E-12 | -0.1233 | 0.0169 | 2.94E-13 | 1 |
| rs62055890 | 17 | 43840681 | 17 | 43399058 | 44874453 | T | C | 0.2396 | CRHR1:RP11-105N1 | ncRNA_intronic | -2.1538 | 0.2585 | 7.89E-17 | -2.2137 | 0.3217 | 6.47E-12 | -0.1227 | 0.0168 | 3.19E-13 | 1 |
| rs77849344 | 17 | 43840864 | 17 | 43399058 | 44874453 | A | G | 0.2396 | CRHR1:RP11-105N1 | ncRNA_intronic | -2.1510 | 0.2583 | 8.20E-17 | -2.2177 | 0.3217 | 5.99E-12 | -0.1220 | 0.0167 | 3.08E-13 | 1 |
| rs74918686 | 17 | 43840899 | 17 | 43399058 | 44874453 | C | G | 0.2396 | CRHR1:RP11-105N1 | ncRNA_intronic | -2.1511 | 0.2582 | 8.09E-17 | -2.2178 | 0.3217 | 6.00E-12 | -0.1220 | 0.0167 | 3.09E-13 | 1 |
| rs79545140 | 17 | 43840935 | 17 | 43399058 | 44874453 | T | C | 0.2396 | CRHR1:RP11-105N1 | ncRNA_intronic | 2.1506 | 0.2582 | 8.24E-17 | 2.2172 | 0.3217 | 6.08E-12 | 0.1223 | 0.0167 | 2.14E-13 | 1 |
| rs56369036 | 17 | 43841571 | 17 | 43399058 | 44874453 | A | T | 0.2396 | CRHR1:RP11-105N1 | ncRNA_intronic | -2.1454 | 0.2581 | 9.28E-17 | -2.2146 | 0.3217 | 6.37E-12 | -0.1219 | 0.0166 | 2.22E-13 | 1 |
| rs11079723 | 17 | 43841729 | 17 | 43399058 | 44874453 | T | C | 0.2406 | CRHR1:RP11-105N1 | ncRNA_intronic | 2.1456 | 0.258 | 9.18E-17 | 2.2056 | 0.3217 | 7.79E-12 | 0.1219 | 0.0167 | 2.91E-13 | 1 |
| rs11079724 | 17 | 43841912 | 17 | 43399058 | 44874453 | T | C | 0.2396 | CRHR1:RP11-105N1 | ncRNA_intronic | -2.1421 | 0.2578 | 9.52E-17 | -2.2155 | 0.3217 | 6.23E-12 | -0.1219 | 0.0166 | 2.12E-13 | 1 |
| rs55707339 | 17 | 43842462 | 17 | 43399058 | 44874453 | T | C | 0.2406 | CRHR1:RP11-105N1 | ncRNA_intronic | 2.1336 | 0.2578 | 1.26E-16 | 2.1980 | 0.3220 | 9.51E-12 | 0.1217 | 0.0166 | 2.17E-13 | 1 |
| rs62055893 | 17 | 43842494 | 17 | 43399058 | 44874453 | A | G | 0.2396 | CRHR1:RP11-105N1 | ncRNA_intronic | 2.1328 | 0.2577 | 1.28E-16 | 2.2039 | 0.3219 | 8.32E-12 | 0.1216 | 0.0166 | 2.14E-13 | 1 |
| rs75257002 | 17 | 43843395 | 17 | 43399058 | 44874453 | T | C | 0.2406 | CRHR1:RP11-105N1 | ncRNA_intronic | -2.1332 | 0.2577 | 1.26E-16 | -2.2151 | 0.3220 | 6.66E-12 | -0.1211 | 0.0165 | 2.38E-13 | 1 |
| rs62055894 | 17 | 43843943 | 17 | 43399058 | 44874453 | A | G | 0.2396 | CRHR1:RP11-105N1 | ncRNA_intronic | -2.1304 | 0.2576 | 1.35E-16 | -2.2215 | 0.3217 | 5.46E-12 | -0.1211 | 0.0165 | 2.34E-13 | 1 |
| rs62055895 | 17 | 43844044 | 17 | 43399058 | 44874453 | A | G | 0.2396 | CRHR1:RP11-105N1 | ncRNA_intronic | 2.1315 | 0.2576 | 1.28E-16 | 2.2228 | 0.3216 | 5.29E-12 | 0.1211 | 0.0165 | 2.36E-13 | 1 |
| rs62055896 | 17 | 43844201 | 17 | 43399058 | 44874453 | T | C | 0.2396 | CRHR1:RP11-105N1 | ncRNA_intronic | -2.1315 | 0.2577 | 1.31E-16 | -2.2228 | 0.3216 | 5.29E-12 | -0.1209 | 0.0165 | 2.41E-13 | 1 |
| rs55725840 | 17 | 43844486 | 17 | 43399058 | 44874453 | A | T | 0.2396 | CRHR1:RP11-105N1 | ncRNA_intronic | -2.1315 | 0.2575 | 1.27E-16 | -2.2228 | 0.3216 | 5.29E-12 | -0.1208 | 0.0165 | 2.24E-13 | 1 |
| rs56194509 | 17 | 43844559 | 17 | 43399058 | 44874453 | T | G | 0.2396 | CRHR1:RP11-105N1 | ncRNA_intronic | 2.1647 | 0.2605 | 9.56E-17 | 2.2337 | 0.3248 | 6.66E-12 | 0.1218 | 0.0167 | 3.10E-13 | 1 |
| rs55657917 | 17 | 43844560 | 17 | 43399058 | 44874453 | T | G | 0.2396 | CRHR1:RP11-105N1 | ncRNA_intronic | 2.1650 | 0.2609 | 1.05E-16 | 2.2337 | 0.3248 | 6.67E-12 | 0.1221 | 0.0166 | 2.16E-13 | 1 |
| rs56082319 | 17 | 43844798 | 17 | 43399058 | 44874453 | A | G | 0.2406 | CRHR1:RP11-105N1 | ncRNA_intronic | -2.1333 | 0.2573 | 1.11E-16 | -2.2169 | 0.3217 | 6.04E-12 | -0.1200 | 0.0164 | 2.45E-13 | 1 |
| rs56109643 | 17 | 43844859 | 17 | 43399058 | 44874453 | A | G | 0.2396 | CRHR1:RP11-105N1 | ncRNA_intronic | 2.1321 | 0.2572 | 1.15E-16 | 2.2227 | 0.3216 | 5.29E-12 | 0.1227 | 0.0162 | 3.49E-14 | 1 |
| rs62055899 | 17 | 43844977 | 17 | 43399058 | 44874453 | A | T | 0.2396 | CRHR1:RP11-105N1 | ncRNA_intronic | -2.1324 | 0.2572 | 1.14E-16 | -2.2244 | 0.3217 | 5.19E-12 | -0.1225 | 0.0162 | 3.49E-14 | 1 |
| rs62055900 | 17 | 43845002 | 17 | 43399058 | 44874453 | C | G | 0.2396 | CRHR1:RP11-105N1 | ncRNA_intronic | 2.1322 | 0.2572 | 1.14E-16 | 2.2246 | 0.3217 | 5.16E-12 | 0.1225 | 0.0162 | 3.44E-14 | 1 |
| rs62055901 | 17 | 43845041 | 17 | 43399058 | 44874453 | A | G | 0.2396 | CRHR1:RP11-105N1 | ncRNA_intronic | -2.1321 | 0.2572 | 1.14E-16 | -2.2164 | 0.3216 | 6.04E-12 | -0.1225 | 0.0162 | 3.43E-14 | 1 |
| rs58089049 | 17 | 43845480 | 17 | 43399058 | 44874453 | T | C | 0.2883 | CRHR1:RP11-105N1 | ncRNA_intronic | -1.6212 | 0.2353 | 5.62E-12 | -1.6788 | 0.3032 | 3.21E-08 | -0.0926 | 0.0167 | 3.02E-08 | 1 |
| rs62055903 | 17 | 43846668 | 17 | 43399058 | 44874453 | A | C | 0.2396 | CRHR1:RP11-105N1 | ncRNA_intronic | -2.1310 | 0.257 | 1.11E-16 | -2.2227 | 0.3216 | 5.29E-12 | -0.1220 | 0.0161 | 3.47E-14 | 1 |
| rs111374028 | 17 | 43846820 | 17 | 43399058 | 44874453 | A | G | 0.2396 | CRHR1:RP11-105N1 | ncRNA_intronic | -2.1309 | 0.257 | 1.12E-16 | -2.2228 | 0.3216 | 5.28E-12 | -0.1218 | 0.0161 | 3.45E-14 | 1 |
| rs113934115 | 17 | 43847039 | 17 | 43399058 | 44874453 | T | C | 0.2396 | CRHR1:RP11-105N1 | ncRNA_intronic | -2.1645 | 0.2594 | 7.24E-17 | -2.2228 | 0.3216 | 5.28E-12 | -0.1239 | 0.0163 | 2.52E-14 | 1 |
| NA | 17 | 43847095 | 17 | 43399058 | 44874453 | T | C | 0.2396 | CRHR1:RP11-105N1 | ncRNA_intronic | -2.1207 | 0.3085 | 6.20E-12 | -2.2286 | 0.3217 | 4.72E-12 | -0.1216 | 0.0160 | 3.31E-14 | 1 |
| rs62055928 | 17 | 43847374 | 17 | 43399058 | 44874453 | A | G | 0.2396 | CRHR1:RP11-105N1 | ncRNA_intronic | -2.1339 | 0.2571 | 1.03E-16 | -2.2229 | 0.3216 | 5.28E-12 | -0.1210 | 0.0160 | 4.26E-14 | 1 |
| rs56268325 | 17 | 43847741 | 17 | 43399058 | 44874453 | T | C | 0.2396 | CRHR1:RP11-105N1 | ncRNA_intronic | 2.1303 | 0.2569 | 1.12E-16 | 2.2228 | 0.3216 | 5.28E-12 | 0.1211 | 0.0160 | 3.25E-14 | 1 |
| rs56070245 | 17 | 43847868 | 17 | 43399058 | 44874453 | T | C | 0.2396 | CRHR1:RP11-105N1 | ncRNA_intronic | 2.1396 | 0.2522 | 2.18E-17 | 2.2228 | 0.3216 | 5.28E-12 | 0.1211 | 0.0160 | 3.28E-14 | 1 |
| rs56387266 | 17 | 43847912 | 17 | 43399058 | 44874453 | T | C | 0.2396 | CRHR1:RP11-105N1 | ncRNA_intronic | 2.1397 | 0.2522 | 2.16E-17 | 2.2228 | 0.3216 | 5.28E-12 | 0.1211 | 0.0159 | 3.18E-14 | 1 |
| rs34303488 | 17 | 43848181 | 17 | 43399058 | 44874453 | T | C | 0.2396 | CRHR1:RP11-105N1 | ncRNA_intronic | -2.1397 | 0.2522 | 2.15E-17 | -2.2227 | 0.3216 | 5.30E-12 | -0.1209 | 0.0159 | 3.19E-14 | 1 |
| rs62055932 | 17 | 43848412 | 17 | 43399058 | 44874453 | A | G | 0.2396 | CRHR1:RP11-105N1 | ncRNA_intronic | 2.1431 | 0.2517 | 1.66E-17 | 2.2118 | 0.3216 | 6.69E-12 | 0.1208 | 0.0159 | 3.17E-14 | 1 |
| rs62055933 | 17 | 43848461 | 17 | 43399058 | 44874453 | A | G | 0.2396 | CRHR1:RP11-105N1 | ncRNA_intronic | -2.1214 | 0.2543 | 7.36E-17 | -2.2407 | 0.3220 | 3.78E-12 | -0.1211 | 0.0159 | 2.82E-14 | 1 |
| NA | 17 | 43848638 | 17 | 43399058 | 44874453 | A | G | 0.2386 | CRHR1:RP11-105N1 | ncRNA_intronic | -2.1272 | 0.3117 | 8.82E-12 | -2.2823 | 0.3227 | 1.68E-12 | -0.1217 | 0.0161 | 3.91E-14 | 1 |
| rs62055935 | 17 | 43848750 | 17 | 43399058 | 44874453 | T | C | 0.2396 | CRHR1:RP11-105N1 | ncRNA_intronic | 2.1369 | 0.2529 | 2.94E-17 | 2.2681 | 0.3235 | 2.60E-12 | 0.1211 | 0.0159 | 2.93E-14 | 1 |
| rs62055936 | 17 | 43848761 | 17 | 43399058 | 44874453 | A | T | 0.2406 | CRHR1:RP11-105N1 | ncRNA_intronic | -2.1367 | 0.2528 | 2.88E-17 | -2.2671 | 0.3234 | 2.61E-12 | -0.1208 | 0.0159 | 3.34E-14 | 1 |
| rs62055937 | 17 | 43848968 | 17 | 43399058 | 44874453 | T | G | 0.2396 | CRHR1:RP11-105N1 | ncRNA_intronic | -2.1444 | 0.2521 | 1.80E-17 | -2.2494 | 0.3223 | 3.27E-12 | -0.1204 | 0.0158 | 3.03E-14 | 1 |
| rs76294809 | 17 | 43849327 | 17 | 43399058 | 44874453 | A | G | 0.2396 | CRHR1:RP11-105N1 | ncRNA_intronic | -2.1376 | 0.252 | 2.22E-17 | -2.2207 | 0.3216 | 5.52E-12 | -0.1206 | 0.0158 | 2.70E-14 | 1 |
| rs75916678 | 17 | 43849366 | 17 | 43399058 | 44874453 | T | C | 0.2396 | CRHR1:RP11-105N1 | ncRNA_intronic | 2.1384 | 0.252 | 2.17E-17 | 2.2160 | 0.3217 | 6.17E-12 | 0.1205 | 0.0158 | 2.69E-14 | 1 |
| rs79730878 | 17 | 43849415 | 17 | 43399058 | 44874453 | T | C | 0.2455 | CRHR1:RP11-105N1 | ncRNA_intronic | 2.1374 | 0.2553 | 5.59E-17 | 2.1507 | 0.3198 | 1.90E-11 | 0.1189 | 0.0150 | 2.58E-15 | 1 |
| rs62055938 | 17 | 43849656 | 17 | 43399058 | 44874453 | A | C | 0.2396 | CRHR1:RP11-105N1 | ncRNA_intronic | -2.1389 | 0.2518 | 1.97E-17 | -2.2229 | 0.3216 | 5.27E-12 | -0.1200 | 0.0158 | 2.77E-14 | 1 |
| rs62055939 | 17 | 43849787 | 17 | 43399058 | 44874453 | A | T | 0.2396 | CRHR1:RP11-105N1 | ncRNA_intronic | 2.1384 | 0.2517 | 1.99E-17 | 2.2229 | 0.3216 | 5.27E-12 | 0.1201 | 0.0154 | 7.20E-15 | 1 |
| rs62055940 | 17 | 43849896 | 17 | 43399058 | 44874453 | A | G | 0.2396 | CRHR1:RP11-105N1 | ncRNA_intronic | 2.1386 | 0.2517 | 1.94E-17 | 2.2229 | 0.3216 | 5.27E-12 | 0.1201 | 0.0154 | 7.08E-15 | 1 |
| rs76885724 | 17 | 43850519 | 17 | 43399058 | 44874453 | T | C | 0.2396 | CRHR1:RP11-105N1 | ncRNA_intronic | -2.1379 | 0.2516 | 1.96E-17 | -2.2229 | 0.3216 | 5.27E-12 | -0.1201 | 0.0154 | 6.96E-15 | 1 |
| rs60814418 | 17 | 43850645 | 17 | 43399058 | 44874453 | T | C | 0.2435 | CRHR1:RP11-105N1 | ncRNA_intronic | -2.0436 | 0.2493 | 2.47E-16 | -2.2215 | 0.3205 | 4.60E-12 | -0.1197 | 0.0158 | 3.61E-14 | 1 |
| NA | 17 | 43850932 | 17 | 43399058 | 44874453 | T | C | 0.2396 | CRHR1:RP11-105N1 | ncRNA_intronic | -2.1178 | 0.3169 | 2.33E-11 | -2.2290 | 0.3217 | 4.67E-12 | -0.1192 | 0.0163 | 2.66E-13 | 1 |
| rs62055942 | 17 | 43850966 | 17 | 43399058 | 44874453 | A | G | 0.2396 | CRHR1:RP11-105N1 | ncRNA_intronic | 2.1415 | 0.252 | 1.93E-17 | 2.2229 | 0.3216 | 5.28E-12 | 0.1199 | 0.0154 | 6.92E-15 | 1 |
| rs62055943 | 17 | 43851018 | 17 | 43399058 | 44874453 | T | C | 0.2396 | CRHR1:RP11-105N1 | ncRNA_intronic | 2.1398 | 0.251 | 1.52E-17 | 2.2228 | 0.3216 | 5.28E-12 | 0.1197 | 0.0154 | 7.09E-15 | 1 |
| rs62055945 | 17 | 43851851 | 17 | 43399058 | 44874453 | A | G | 0.2396 | CRHR1:RP11-105N1 | ncRNA_intronic | -2.1368 | 0.2514 | 1.92E-17 | -2.2228 | 0.3216 | 5.29E-12 | -0.1169 | 0.0149 | 4.20E-15 | 1 |
| rs62055946 | 17 | 43851971 | 17 | 43399058 | 44874453 | A | C | 0.2396 | CRHR1:RP11-105N1 | ncRNA_intronic | -2.1365 | 0.2514 | 1.93E-17 | -2.2231 | 0.3216 | 5.25E-12 | -0.1166 | 0.0149 | 4.43E-15 | 1 |
| rs62055947 | 17 | 43852605 | 17 | 43399058 | 44874453 | T | C | 0.2396 | CRHR1:RP11-105N1 | ncRNA_intronic | -2.1374 | 0.2519 | 2.15E-17 | -2.2265 | 0.3228 | 5.80E-12 | -0.1165 | 0.0149 | 4.49E-15 | 1 |
| rs62055948 | 17 | 43852621 | 17 | 43399058 | 44874453 | T | C | 0.2396 | CRHR1:RP11-105N1 | ncRNA_intronic | 2.1373 | 0.2518 | 2.10E-17 | 2.2260 | 0.3228 | 5.85E-12 | 0.1164 | 0.0148 | 4.48E-15 | 1 |
| rs113871181 | 17 | 43852733 | 17 | 43399058 | 44874453 | A | G | 0.2396 | CRHR1:RP11-105N1 | ncRNA_intronic | -1.9957 | 0.1853 | 4.74E-27 | -2.2135 | 0.3223 | 7.13E-12 | -0.1163 | 0.0148 | 4.59E-15 | 1 |
| rs111370985 | 17 | 43852742 | 17 | 43399058 | 44874453 | T | C | 0.2396 | CRHR1:RP11-105N1 | ncRNA_intronic | 1.9961 | 0.1853 | 4.63E-27 | 2.2072 | 0.3223 | 8.24E-12 | 0.1163 | 0.0148 | 4.48E-15 | 1 |
| rs55787105 | 17 | 43853109 | 17 | 43399058 | 44874453 | A | G | 0.2396 | CRHR1:RP11-105N1 | ncRNA_intronic | -2.1353 | 0.2513 | 1.92E-17 | -2.2231 | 0.3216 | 5.26E-12 | -0.1164 | 0.0148 | 4.10E-15 | 1 |
| rs62055950 | 17 | 43853133 | 17 | 43399058 | 44874453 | T | C | 0.2396 | CRHR1:RP11-105N1 | ncRNA_intronic | 2.1350 | 0.2512 | 1.93E-17 | 2.2227 | 0.3216 | 5.29E-12 | 0.1162 | 0.0148 | 4.34E-15 | 1 |
| rs56043078 | 17 | 43853235 | 17 | 43399058 | 44874453 | T | G | 0.2396 | CRHR1:RP11-105N1 | ncRNA_intronic | 2.1348 | 0.2512 | 1.90E-17 | 2.2227 | 0.3216 | 5.29E-12 | 0.1161 | 0.0148 | 4.35E-15 | 1 |
| rs77965652 | 17 | 43853457 | 17 | 43399058 | 44874453 | A | G | 0.2396 | CRHR1:RP11-105N1 | ncRNA_intronic | 2.1346 | 0.2512 | 1.91E-17 | 2.2227 | 0.3216 | 5.29E-12 | 0.1160 | 0.0148 | 4.47E-15 | 1 |
| rs55849949 | 17 | 43853526 | 17 | 43399058 | 44874453 | A | G | 0.2396 | CRHR1:RP11-105N1 | ncRNA_intronic | 2.1345 | 0.2511 | 1.91E-17 | 2.2228 | 0.3216 | 5.28E-12 | 0.1161 | 0.0148 | 4.31E-15 | 1 |
| rs56303031 | 17 | 43853922 | 17 | 43399058 | 44874453 | A | G | 0.2396 | CRHR1:RP11-105N1 | ncRNA_intronic | -2.0659 | 0.2792 | 1.37E-13 | -2.2181 | 0.3216 | 5.87E-12 | -0.1160 | 0.0148 | 4.00E-15 | 1 |
| rs55991914 | 17 | 43854267 | 17 | 43399058 | 44874453 | T | C | 0.2396 | CRHR1:RP11-105N1 | ncRNA_intronic | -2.1345 | 0.2511 | 1.91E-17 | -2.2228 | 0.3216 | 5.28E-12 | -0.1158 | 0.0147 | 3.83E-15 | 1 |
| rs62055955 | 17 | 43854340 | 17 | 43399058 | 44874453 | A | G | 0.2406 | CRHR1:RP11-105N1 | ncRNA_intronic | 2.1327 | 0.2511 | 2.01E-17 | 2.2239 | 0.3215 | 5.09E-12 | 0.1156 | 0.0147 | 4.21E-15 | 1 |
| rs62055956 | 17 | 43854449 | 17 | 43399058 | 44874453 | T | C | 0.2406 | CRHR1:RP11-105N1 | ncRNA_intronic | -2.1399 | 0.2511 | 1.58E-17 | -2.2303 | 0.3215 | 4.41E-12 | -0.1156 | 0.0147 | 4.09E-15 | 1 |
| rs12938031 | 17 | 43854502 | 17 | 43399058 | 44874453 | A | G | 0.3887 | CRHR1:RP11-105N1 | ncRNA_intronic | 1.1816 | 0.2108 | 2.08E-08 | 1.6389 | 0.2714 | 1.64E-09 | 0.0458 | 0.0125 | 2.42E-04 | 1 |
| rs62055957 | 17 | 43854536 | 17 | 43399058 | 44874453 | A | C | 0.2406 | CRHR1:RP11-105N1 | ncRNA_intronic | -2.1334 | 0.2511 | 1.95E-17 | -2.2240 | 0.3215 | 5.08E-12 | -0.1155 | 0.0147 | 4.10E-15 | 1 |
| rs4074462 | 17 | 43855228 | 17 | 43399058 | 44874453 | T | G | 0.2406 | CRHR1:RP11-105N1 | ncRNA_intronic | -2.1157 | 0.2501 | 2.71E-17 | -2.2180 | 0.3215 | 5.78E-12 | -0.1153 | 0.0147 | 4.31E-15 | 1 |
| rs574201100 | 17 | 43855603 | 17 | 43399058 | 44874453 | A | G | 0.3926 | CRHR1:RP11-105N1 | ncRNA_intronic | -1.1837 | 0.2105 | 1.87E-08 | -1.5945 | 0.2708 | 4.14E-09 | -0.0487 | 0.0130 | 1.82E-04 | 1 |
| NA | 17 | 43856372 | 17 | 43399058 | 44874453 | A | G | 0.2386 | CRHR1:RP11-105N1 | ncRNA_intronic | -2.1170 | 0.2964 | 9.26E-13 | -2.2455 | 0.3232 | 4.07E-12 | -0.1144 | 0.0141 | 5.16E-16 | 1 |
| rs62057061 | 17 | 43856639 | 17 | 43399058 | 44874453 | C | G | 0.2396 | CRHR1:RP11-105N1 | ncRNA_intronic | 2.1370 | 0.2496 | 1.12E-17 | 2.2126 | 0.3219 | 6.83E-12 | 0.1142 | 0.0141 | 5.02E-16 | 1 |
| rs62057062 | 17 | 43856710 | 17 | 43399058 | 44874453 | A | G | 0.2396 | CRHR1:RP11-105N1 | ncRNA_intronic | -1.9869 | 0.1843 | 4.22E-27 | -2.2227 | 0.3216 | 5.30E-12 | -0.1142 | 0.0141 | 5.02E-16 | 1 |
| rs62057063 | 17 | 43856730 | 17 | 43399058 | 44874453 | T | C | 0.2396 | CRHR1:RP11-105N1 | ncRNA_intronic | -1.9914 | 0.1848 | 4.49E-27 | -2.2323 | 0.3219 | 4.48E-12 | -0.1153 | 0.0142 | 4.94E-16 | 1 |
| rs62057064 | 17 | 43857033 | 17 | 43399058 | 44874453 | T | C | 0.2396 | CRHR1:RP11-105N1 | ncRNA_intronic | -1.9859 | 0.183 | 1.90E-27 | -2.2228 | 0.3216 | 5.30E-12 | -0.1141 | 0.0141 | 5.07E-16 | 1 |
| rs76830096 | 17 | 43857129 | 17 | 43399058 | 44874453 | A | C | 0.2396 | CRHR1:RP11-105N1 | ncRNA_intronic | 1.9820 | 0.183 | 2.40E-27 | 2.2227 | 0.3216 | 5.29E-12 | 0.1140 | 0.0141 | 5.11E-16 | 1 |
| rs62057065 | 17 | 43858187 | 17 | 43399058 | 44874453 | T | G | 0.2396 | CRHR1:RP11-105N1 | ncRNA_intronic | -2.1329 | 0.251 | 1.91E-17 | -2.2128 | 0.3217 | 6.60E-12 | -0.1144 | 0.0141 | 5.42E-16 | 1 |
| NA | 17 | 43858307 | 17 | 43399058 | 44874453 | T | C | 0.2406 | CRHR1:RP11-105N1 | ncRNA_intronic | 2.1086 | 0.3031 | 3.48E-12 | 2.2364 | 0.3231 | 4.92E-12 | 0.1150 | 0.0143 | 7.17E-16 | 1 |
| rs62057067 | 17 | 43858326 | 17 | 43399058 | 44874453 | T | C | 0.2396 | CRHR1:RP11-105N1 | ncRNA_intronic | -1.9871 | 0.1847 | 5.58E-27 | -2.2218 | 0.3216 | 5.39E-12 | -0.1150 | 0.0142 | 5.38E-16 | 1 |
| rs78917495 | 17 | 43858482 | 17 | 43399058 | 44874453 | T | C | 0.2396 | CRHR1:RP11-105N1 | ncRNA_intronic | -1.9812 | 0.1828 | 2.30E-27 | -2.2154 | 0.3216 | 6.22E-12 | -0.1131 | 0.0140 | 5.27E-16 | 1 |
| rs62057068 | 17 | 43858629 | 17 | 43399058 | 44874453 | A | G | 0.2396 | CRHR1:RP11-105N1 | ncRNA_intronic | 1.9810 | 0.1828 | 2.31E-27 | 2.2215 | 0.3216 | 5.42E-12 | 0.1130 | 0.0139 | 5.16E-16 | 1 |
| rs62057069 | 17 | 43859065 | 17 | 43399058 | 44874453 | T | C | 0.2396 | CRHR1:RP11-105N1 | ncRNA_intronic | 1.9806 | 0.1828 | 2.36E-27 | 2.2209 | 0.3216 | 5.47E-12 | 0.1131 | 0.0139 | 4.72E-16 | 1 |
| rs62057070 | 17 | 43859640 | 17 | 43399058 | 44874453 | A | G | 0.2396 | CRHR1:RP11-105N1 | ncRNA_intronic | 1.9806 | 0.1828 | 2.33E-27 | 2.2210 | 0.3216 | 5.46E-12 | 0.1126 | 0.0139 | 4.80E-16 | 1 |
| rs62057071 | 17 | 43859691 | 17 | 43399058 | 44874453 | C | G | 0.2396 | CRHR1:RP11-105N1 | ncRNA_intronic | 1.9804 | 0.1828 | 2.37E-27 | 2.2210 | 0.3216 | 5.45E-12 | 0.1125 | 0.0139 | 5.00E-16 | 1 |
| rs62057073 | 17 | 43861117 | 17 | 43399058 | 44874453 | T | C | 0.2396 | CRHR1:RP11-105N1 | ncRNA_intronic | -1.9866 | 0.1851 | 7.08E-27 | -2.2536 | 0.3221 | 2.91E-12 | -0.1136 | 0.0141 | 9.45E-16 | 1 |
| rs12944712 | 17 | 43871147 | 17 | 43399058 | 44874453 | A | G | 0.4682 | CRHR1:RP11-105N1 | ncRNA_intronic | -0.9449 | 0.1438 | 4.99E-11 | -1.3643 | 0.2672 | 3.41E-07 | -0.0328 | 0.0106 | 2.06E-03 | 1 |
| rs56319902 | 17 | 43871982 | 17 | 43399058 | 44874453 | T | C | 0.2396 | CRHR1:RP11-105N1 | ncRNA_intronic | -2.0011 | 0.1821 | 4.19E-28 | -2.2227 | 0.3220 | 5.62E-12 | -0.1102 | 0.0136 | 4.30E-16 | 1 |
| rs55943044 | 17 | 43872228 | 17 | 43399058 | 44874453 | A | G | 0.2396 | CRHR1:RP11-105N1 | ncRNA_intronic | -1.9906 | 0.1821 | 8.00E-28 | -2.2187 | 0.3216 | 5.74E-12 | -0.1099 | 0.0135 | 4.83E-16 | 1 |

| rsID | chr | pos (hg19) | Genomic Locus | Genomic Locus start | Genomic Locus end | A1 | A2 | MAF | Nearest Genes | Function | Meta-GWAS | | | UKBB | | | ENIGMA* | | | Replicated** |
|---|---|---|---|---|---|---|---|---|---|---|---|---|---|---|---|---|---|---|---|---|
| | | | | | | | | | | | Beta | SE | P-value | Beta | SE | P-value | Beta | SE | P-value | |
| rs80184151 | 17 | 43879308 | 17 | 43399058 | 44874453 | A | G | 0.2396 | CRHR1:RP11-105N1 | ncRNA_intronic | 1.9699 | 0.1808 | 1.21E-27 | 2.2310 | 0.3214 | 4.25E-12 | 0.1089 | 0.0134 | 4.38E-16 | 1 |
| rs17689378 | 17 | 43881790 | 17 | 43399058 | 44874453 | T | C | 0.2396 | CRHR1:RP11-105N1 | ncRNA_intronic | -1.9418 | 0.1802 | 4.34E-27 | -2.2310 | 0.3214 | 4.25E-12 | -0.1086 | 0.0133 | 4.05E-16 | 1 |
| rs62057101 | 17 | 43885291 | 17 | 43399058 | 44874453 | A | G | 0.2396 | CRHR1:RP11-105N1 | ncRNA_intronic | -1.9399 | 0.1801 | 4.77E-27 | -2.2298 | 0.3214 | 4.39E-12 | -0.1087 | 0.0133 | 3.28E-16 | 1 |
| rs62057103 | 17 | 43887480 | 17 | 43399058 | 44874453 | T | C | 0.2396 | CRHR1:RP11-105N1 | ncRNA_intronic | 2.0545 | 0.2396 | 1.00E-17 | 2.2289 | 0.3216 | 4.59E-12 | 0.1093 | 0.0133 | 2.12E-16 | 1 |
| rs55915917 | 17 | 43892784 | 17 | 43399058 | 44874453 | T | G | 0.2256 | CRHR1:RP11-105N1 | ncRNA_intronic | 2.0065 | 0.1846 | 1.66E-27 | 2.2602 | 0.3244 | 3.56E-12 | 0.1135 | 0.0138 | 2.23E-16 | 1 |
| rs55668363 | 17 | 43892788 | 17 | 43399058 | 44874453 | A | G | 0.2247 | CRHR1:RP11-105N1 | ncRNA_intronic | -1.9993 | 0.1848 | 2.79E-27 | -2.2555 | 0.3248 | 4.17E-12 | -0.1129 | 0.0138 | 3.67E-16 | 1 |
| rs17689471 | 17 | 43892973 | 17 | 43399058 | 44874453 | T | C | 0.2396 | CRHR1:RP11-105N1 | ncRNA_intronic | 1.9470 | 0.1799 | 2.73E-27 | 2.2382 | 0.3214 | 3.66E-12 | 0.1092 | 0.0132 | 1.69E-16 | 1 |
| rs117365970 | 17 | 43893259 | 17 | 43399058 | 44874453 | A | G | 0.2396 | CRHR1:RP11-105N1 | ncRNA_intronic | -2.0072 | 0.1836 | 7.92E-28 | -2.2534 | 0.3221 | 2.92E-12 | -0.1144 | 0.0139 | 2.11E-16 | 1 |
| rs117646503 | 17 | 43893260 | 17 | 43399058 | 44874453 | T | C | 0.2396 | CRHR1:RP11-105N1 | ncRNA_intronic | 2.0258 | 0.1831 | 1.94E-28 | 2.2385 | 0.3214 | 3.66E-12 | 0.1150 | 0.0139 | 1.35E-16 | 1 |
| rs17762769 | 17 | 43893403 | 17 | 43399058 | 44874453 | A | G | 0.2396 | CRHR1:RP11-105N1 | ncRNA_intronic | -1.9477 | 0.1799 | 2.59E-27 | -2.2386 | 0.3215 | 3.64E-12 | -0.1093 | 0.0132 | 1.54E-16 | 1 |
| rs8072451 | 17 | 43893716 | 17 | 43399058 | 44874453 | T | C | 0.2396 | CRHR1:RP11-105N1 | ncRNA_intronic | -1.9544 | 0.1799 | 1.69E-27 | -2.2398 | 0.3221 | 3.90E-12 | -0.1093 | 0.0132 | 1.54E-16 | 1 |
| rs8073146 | 17 | 43893751 | 17 | 43399058 | 44874453 | A | G | 0.2396 | CRHR1:RP11-105N1 | ncRNA_intronic | 1.9523 | 0.1799 | 1.95E-27 | 2.2434 | 0.3223 | 3.71E-12 | 0.1093 | 0.0132 | 1.57E-16 | 1 |
| rs28364025 | 17 | 43894102 | 17 | 43399058 | 44874453 | T | C | 0.2396 | CRHR1 | intronic | 1.9486 | 0.1799 | 2.46E-27 | 2.2413 | 0.3216 | 3.50E-12 | 0.1093 | 0.0132 | 1.50E-16 | 1 |
| rs28364023 | 17 | 43894159 | 17 | 43399058 | 44874453 | T | C | 0.2396 | CRHR1 | intronic | -1.9495 | 0.18 | 2.46E-27 | -2.2403 | 0.3216 | 3.58E-12 | -0.1092 | 0.0132 | 1.55E-16 | 1 |
| rs55779147 | 17 | 43894510 | 17 | 43399058 | 44874453 | A | G | 0.2396 | CRHR1 | intronic | -1.9519 | 0.1801 | 2.31E-27 | -2.2414 | 0.3215 | 3.44E-12 | -0.1097 | 0.0133 | 1.48E-16 | 1 |
| rs56357543 | 17 | 43894547 | 17 | 43399058 | 44874453 | T | C | 0.2396 | CRHR1 | intronic | -1.9497 | 0.1802 | 2.69E-27 | -2.2539 | 0.3217 | 2.70E-12 | -0.1097 | 0.0133 | 1.48E-16 | 1 |
| rs56099546 | 17 | 43894609 | 17 | 43399058 | 44874453 | A | G | 0.2396 | CRHR1 | intronic | 1.9498 | 0.1802 | 2.68E-27 | 2.2541 | 0.3217 | 2.69E-12 | 0.1097 | 0.0133 | 1.48E-16 | 1 |
| rs739645 | 17 | 43894990 | 17 | 43399058 | 44874453 | T | G | 0.2396 | CRHR1 | intronic | 1.9532 | 0.1802 | 2.18E-27 | 2.2428 | 0.3215 | 3.35E-12 | 0.1096 | 0.0133 | 1.56E-16 | 1 |
| rs739644 | 17 | 43895008 | 17 | 43399058 | 44874453 | C | G | 0.2396 | CRHR1 | intronic | 1.9519 | 0.1801 | 2.31E-27 | 2.2427 | 0.3215 | 3.35E-12 | 0.1096 | 0.0133 | 1.54E-16 | 1 |
| rs4564621 | 17 | 43895501 | 17 | 43399058 | 44874453 | C | G | 0.2396 | CRHR1 | intronic | -1.9523 | 0.1802 | 2.32E-27 | -2.2443 | 0.3215 | 3.25E-12 | -0.1095 | 0.0133 | 1.59E-16 | 1 |
| rs2316763 | 17 | 43895530 | 17 | 43399058 | 44874453 | T | C | 0.2396 | CRHR1 | intronic | -1.9717 | 0.1801 | 6.89E-28 | -2.2458 | 0.3216 | 3.17E-12 | -0.1100 | 0.0133 | 1.15E-16 | 1 |
| rs2316764 | 17 | 43895602 | 17 | 43399058 | 44874453 | T | G | 0.2396 | CRHR1 | intronic | 1.9426 | 0.18 | 3.63E-27 | 2.2370 | 0.3216 | 3.87E-12 | 0.1090 | 0.0132 | 1.63E-16 | 1 |
| rs4277389 | 17 | 43895653 | 17 | 43399058 | 44874453 | A | G | 0.2396 | CRHR1 | intronic | 1.9455 | 0.1801 | 3.28E-27 | 2.2356 | 0.3216 | 3.96E-12 | 0.1096 | 0.0132 | 1.14E-16 | 1 |
| rs4566211 | 17 | 43895696 | 17 | 43399058 | 44874453 | A | G | 0.2396 | CRHR1 | intronic | -1.9526 | 0.18 | 2.10E-27 | -2.2464 | 0.3216 | 3.13E-12 | -0.1090 | 0.0132 | 1.58E-16 | 1 |
| rs4566212 | 17 | 43895751 | 17 | 43399058 | 44874453 | A | G | 0.2396 | CRHR1 | intronic | -1.9525 | 0.18 | 2.10E-27 | -2.2462 | 0.3215 | 3.12E-12 | -0.1090 | 0.0132 | 1.58E-16 | 1 |
| rs4309444 | 17 | 43895797 | 17 | 43399058 | 44874453 | T | C | 0.2396 | CRHR1 | intronic | 1.9478 | 0.1801 | 2.86E-27 | 2.2372 | 0.3216 | 3.84E-12 | 0.1090 | 0.0132 | 1.58E-16 | 1 |
| rs62057107 | 17 | 43896032 | 17 | 43399058 | 44874453 | T | C | 0.2396 | CRHR1 | intronic | -1.9597 | 0.18 | 1.33E-27 | -2.2355 | 0.3215 | 3.96E-12 | -0.1092 | 0.0132 | 1.61E-16 | 1 |
| rs12150390 | 17 | 43896228 | 17 | 43399058 | 44874453 | T | C | 0.2396 | CRHR1 | intronic | 1.9609 | 0.1799 | 1.15E-27 | 2.2446 | 0.3215 | 3.21E-12 | 0.1091 | 0.0132 | 1.51E-16 | 1 |
| rs17689608 | 17 | 43896528 | 17 | 43399058 | 44874453 | C | G | 0.2396 | CRHR1 | intronic | 1.9585 | 0.1797 | 1.20E-27 | 2.2358 | 0.3215 | 3.93E-12 | 0.1091 | 0.0132 | 1.47E-16 | 1 |
| rs62057108 | 17 | 43896616 | 17 | 43399058 | 44874453 | T | C | 0.2396 | CRHR1 | intronic | 1.9562 | 0.1799 | 1.57E-27 | 2.2359 | 0.3215 | 3.93E-12 | 0.1091 | 0.0132 | 1.51E-16 | 1 |
| rs62057109 | 17 | 43896637 | 17 | 43399058 | 44874453 | T | C | 0.2396 | CRHR1 | intronic | 1.9561 | 0.1799 | 1.58E-27 | 2.2359 | 0.3215 | 3.93E-12 | 0.1091 | 0.0132 | 1.50E-16 | 1 |
| rs78074121 | 17 | 43896690 | 17 | 43399058 | 44874453 | T | C | 0.2396 | CRHR1 | intronic | 1.9563 | 0.1799 | 1.53E-27 | 2.2379 | 0.3216 | 3.77E-12 | 0.1091 | 0.0132 | 1.49E-16 | 1 |
| rs62057110 | 17 | 43896734 | 17 | 43399058 | 44874453 | T | C | 0.2396 | CRHR1 | intronic | 1.9549 | 0.18 | 1.77E-27 | 2.2359 | 0.3215 | 3.92E-12 | 0.1092 | 0.0132 | 1.50E-16 | 1 |
| rs62057111 | 17 | 43897130 | 17 | 43399058 | 44874453 | A | G | 0.2396 | CRHR1 | intronic | 1.9499 | 0.1799 | 2.19E-27 | 2.2420 | 0.3214 | 3.33E-12 | 0.1091 | 0.0132 | 1.40E-16 | 1 |
| rs62057112 | 17 | 43897202 | 17 | 43399058 | 44874453 | A | T | 0.2396 | CRHR1 | intronic | -1.9520 | 0.1798 | 1.92E-27 | -2.2496 | 0.3215 | 2.88E-12 | -0.1092 | 0.0132 | 1.36E-16 | 1 |
| rs78587102 | 17 | 43897246 | 17 | 43399058 | 44874453 | A | G | 0.2396 | CRHR1 | intronic | 1.9543 | 0.1797 | 1.47E-27 | 2.2495 | 0.3215 | 2.89E-12 | 0.1092 | 0.0132 | 1.32E-16 | 1 |
| rs62057113 | 17 | 43897449 | 17 | 43399058 | 44874453 | A | T | 0.2396 | CRHR1 | intronic | 1.9866 | 0.1811 | 5.32E-28 | 2.2363 | 0.3216 | 3.89E-12 | 0.1109 | 0.0133 | 8.92E-17 | 1 |
| rs78506181 | 17 | 43897480 | 17 | 43399058 | 44874453 | A | G | 0.2396 | CRHR1 | intronic | 1.9838 | 0.181 | 6.07E-28 | 2.2454 | 0.3215 | 3.16E-12 | 0.1109 | 0.0133 | 8.90E-17 | 1 |
| rs79600142 | 17 | 43897722 | 17 | 43399058 | 44874453 | T | C | 0.2396 | CRHR1 | intronic | 1.9856 | 0.1811 | 5.54E-28 | 2.2368 | 0.3217 | 3.93E-12 | 0.1109 | 0.0133 | 8.82E-17 | 1 |
| rs111739681 | 17 | 43898459 | 17 | 43399058 | 44874453 | T | C | 0.2396 | CRHR1 | intronic | -1.9508 | 0.1799 | 2.12E-27 | -2.2282 | 0.3217 | 4.78E-12 | -0.1093 | 0.0132 | 1.25E-16 | 1 |
| rs17762882 | 17 | 43898887 | 17 | 43399058 | 44874453 | T | C | 0.2396 | CRHR1 | intronic | 1.9494 | 0.1799 | 2.26E-27 | 2.2357 | 0.3216 | 3.96E-12 | 0.1093 | 0.0132 | 1.23E-16 | 1 |
| rs17689653 | 17 | 43898963 | 17 | 43399058 | 44874453 | A | T | 0.2396 | CRHR1 | intronic | 1.9495 | 0.1799 | 2.23E-27 | 2.2357 | 0.3216 | 3.96E-12 | 0.1093 | 0.0132 | 1.23E-16 | 1 |
| rs17762912 | 17 | 43899161 | 17 | 43399058 | 44874453 | A | C | 0.2396 | CRHR1 | intronic | 1.9501 | 0.1799 | 2.21E-27 | 2.2406 | 0.3217 | 3.62E-12 | 0.1093 | 0.0132 | 1.21E-16 | 1 |
| rs62057114 | 17 | 43899401 | 17 | 43399058 | 44874453 | T | C | 0.2396 | CRHR1 | intronic | -1.9507 | 0.1799 | 2.13E-27 | -2.2380 | 0.3216 | 3.79E-12 | -0.1093 | 0.0132 | 1.20E-16 | 1 |
| rs62057115 | 17 | 43899417 | 17 | 43399058 | 44874453 | C | G | 0.2396 | CRHR1 | intronic | -1.9492 | 0.1799 | 2.30E-27 | -2.2380 | 0.3216 | 3.79E-12 | -0.1093 | 0.0132 | 1.19E-16 | 1 |
| rs78917479 | 17 | 43899611 | 17 | 43399058 | 44874453 | T | C | 0.2396 | CRHR1 | intronic | -1.9492 | 0.1799 | 2.29E-27 | -2.2380 | 0.3216 | 3.78E-12 | -0.1093 | 0.0132 | 1.17E-16 | 1 |
| rs62057116 | 17 | 43899655 | 17 | 43399058 | 44874453 | T | C | 0.2396 | CRHR1 | intronic | 1.9496 | 0.1799 | 2.23E-27 | 2.2382 | 0.3216 | 3.79E-12 | 0.1093 | 0.0132 | 1.19E-16 | 1 |
| rs62057117 | 17 | 43899657 | 17 | 43399058 | 44874453 | C | G | 0.2396 | CRHR1 | intronic | 1.9497 | 0.1799 | 2.22E-27 | 2.2385 | 0.3216 | 3.76E-12 | 0.1093 | 0.0132 | 1.16E-16 | 1 |
| rs62057118 | 17 | 43899727 | 17 | 43399058 | 44874453 | A | G | 0.2396 | CRHR1 | intronic | 1.9498 | 0.1799 | 2.31E-27 | 2.2487 | 0.3217 | 3.05E-12 | 0.1093 | 0.0132 | 1.13E-16 | 1 |
| rs62057119 | 17 | 43899736 | 17 | 43399058 | 44874453 | A | G | 0.2396 | CRHR1 | intronic | -1.9468 | 0.1799 | 2.71E-27 | -2.2475 | 0.3217 | 3.11E-12 | -0.1093 | 0.0132 | 1.12E-16 | 1 |
| rs17762954 | 17 | 43899786 | 17 | 43399058 | 44874453 | T | C | 0.2396 | CRHR1 | intronic | -1.9468 | 0.1799 | 2.71E-27 | -2.2473 | 0.3217 | 3.11E-12 | -0.1093 | 0.0132 | 1.12E-16 | 1 |
| NA | 17 | 43900081 | 17 | 43399058 | 44874453 | T | C | 0.2396 | CRHR1 | intronic | -2.0190 | 0.2787 | 4.35E-13 | -2.2321 | 0.3217 | 4.39E-12 | -0.1093 | 0.0132 | 1.11E-16 | 1 |
| rs55638417 | 17 | 43900434 | 17 | 43399058 | 44874453 | A | G | 0.2396 | CRHR1 | intronic | 2.0678 | 0.2408 | 8.92E-18 | 2.2400 | 0.3216 | 3.61E-12 | 0.1109 | 0.0133 | 6.40E-17 | 1 |
| rs79501144 | 17 | 43900697 | 17 | 43399058 | 44874453 | T | C | 0.2396 | CRHR1 | intronic | -1.9479 | 0.1798 | 2.41E-27 | -2.2381 | 0.3216 | 3.75E-12 | -0.1093 | 0.0132 | 1.11E-16 | 1 |
| NA | 17 | 43900760 | 17 | 43399058 | 44874453 | A | G | 0.2396 | CRHR1 | intronic | -2.0175 | 0.2827 | 9.62E-13 | -2.2535 | 0.3253 | 4.74E-12 | -0.1114 | 0.0135 | 1.29E-16 | 1 |
| rs112137135 | 17 | 43900817 | 17 | 43399058 | 44874453 | A | G | 0.2396 | CRHR1 | intronic | 2.0711 | 0.2396 | 5.35E-18 | 2.2384 | 0.3216 | 3.73E-12 | 0.1102 | 0.0133 | 1.12E-16 | 1 |
| rs62057122 | 17 | 43901001 | 17 | 43399058 | 44874453 | T | C | 0.2396 | CRHR1 | intronic | -1.9504 | 0.1796 | 1.77E-27 | -2.2376 | 0.3216 | 3.82E-12 | -0.1098 | 0.0132 | 8.60E-17 | 1 |
| rs173365 | 17 | 43901074 | 17 | 43399058 | 44874453 | A | G | 0.4433 | CRHR1 | intronic | -0.8810 | 0.1404 | 3.54E-10 | -1.1331 | 0.2736 | 3.50E-05 | -0.0537 | 0.0110 | 1.16E-06 | 1 |
| rs62057123 | 17 | 43901238 | 17 | 43399058 | 44874453 | A | G | 0.2396 | CRHR1 | intronic | 1.9481 | 0.1798 | 2.38E-27 | 2.2381 | 0.3216 | 3.75E-12 | 0.1093 | 0.0132 | 1.08E-16 | 1 |
| rs62057143 | 17 | 43901528 | 17 | 43399058 | 44874453 | T | C | 0.2396 | CRHR1 | intronic | 1.9528 | 0.1798 | 1.74E-27 | 2.2475 | 0.3215 | 3.03E-12 | 0.1094 | 0.0132 | 1.06E-16 | 1 |
| rs62057144 | 17 | 43901558 | 17 | 43399058 | 44874453 | A | G | 0.2396 | CRHR1 | intronic | 1.9579 | 0.1799 | 1.34E-27 | 2.2457 | 0.3215 | 3.15E-12 | 0.1089 | 0.0132 | 1.58E-16 | 1 |
| rs4335809 | 17 | 43902216 | 17 | 43399058 | 44874453 | T | C | 0.2396 | CRHR1 | intronic | -1.9531 | 0.1798 | 1.73E-27 | -2.2473 | 0.3215 | 3.04E-12 | -0.1094 | 0.0132 | 1.05E-16 | 1 |
| rs4341787 | 17 | 43902505 | 17 | 43399058 | 44874453 | T | C | 0.2406 | CRHR1 | intronic | -1.9531 | 0.1798 | 1.73E-27 | -2.2471 | 0.3215 | 3.06E-12 | -0.1094 | 0.0132 | 1.04E-16 | 1 |
| rs4523962 | 17 | 43902522 | 17 | 43399058 | 44874453 | A | G | 0.2396 | CRHR1 | intronic | -1.9531 | 0.1798 | 1.74E-27 | -2.2471 | 0.3215 | 3.06E-12 | -0.1094 | 0.0132 | 1.04E-16 | 1 |
| rs4327090 | 17 | 43902541 | 17 | 43399058 | 44874453 | A | G | 0.2038 | CRHR1 | intronic | 1.9705 | 0.1944 | 3.87E-24 | 2.1389 | 0.3398 | 3.28E-10 | 0.1162 | 0.0144 | 8.19E-16 | 1 |
| rs3885074 | 17 | 43902738 | 17 | 43399058 | 44874453 | A | C | 0.2396 | CRHR1 | intronic | -1.9531 | 0.1798 | 1.74E-27 | -2.2471 | 0.3215 | 3.07E-12 | -0.1094 | 0.0132 | 9.86E-17 | 1 |
| rs3885075 | 17 | 43902799 | 17 | 43399058 | 44874453 | A | G | 0.2396 | CRHR1 | intronic | 1.9530 | 0.1798 | 1.76E-27 | 2.2470 | 0.3215 | 3.07E-12 | 0.1094 | 0.0132 | 9.83E-17 | 1 |
| rs41280116 | 17 | 43902842 | 17 | 43399058 | 44874453 | T | C | 0.2396 | CRHR1 | intronic | -1.9547 | 0.1802 | 1.99E-27 | -2.2471 | 0.3215 | 3.07E-12 | -0.1099 | 0.0132 | 1.04E-16 | 1 |
| rs34283254 | 17 | 43902861 | 17 | 43399058 | 44874453 | A | C | 0.2396 | CRHR1 | exonic | -2.0702 | 0.2398 | 5.95E-18 | -2.2471 | 0.3215 | 3.07E-12 | -0.1099 | 0.0132 | 1.04E-16 | 1 |
| rs1912151 | 17 | 43902944 | 17 | 43399058 | 44874453 | T | C | 0.2396 | CRHR1 | UTR3 | -1.9530 | 0.1798 | 1.76E-27 | -2.2468 | 0.3215 | 3.09E-12 | -0.1095 | 0.0132 | 9.63E-17 | 1 |
| rs1396862 | 17 | 43902997 | 17 | 43399058 | 44874453 | A | G | 0.2396 | CRHR1 | intronic | -1.9535 | 0.1796 | 1.52E-27 | -2.2468 | 0.3215 | 3.09E-12 | -0.1095 | 0.0132 | 9.58E-17 | 1 |
| rs41280118 | 17 | 43903089 | 17 | 43399058 | 44874453 | T | C | 0.2396 | CRHR1 | intronic | -1.9530 | 0.1798 | 1.76E-27 | -2.2469 | 0.3216 | 3.09E-12 | -0.1096 | 0.0132 | 9.32E-17 | 1 |
| rs62057146 | 17 | 43903106 | 17 | 43399058 | 44874453 | A | G | 0.2396 | CRHR1 | intronic | 1.9528 | 0.1798 | 1.79E-27 | 2.2471 | 0.3216 | 3.07E-12 | 0.1096 | 0.0132 | 9.34E-17 | 1 |
| rs62057147 | 17 | 43903298 | 17 | 43399058 | 44874453 | A | G | 0.2396 | CRHR1 | intronic | -1.9505 | 0.18 | 2.32E-27 | -2.2568 | 0.3216 | 2.51E-12 | -0.1095 | 0.0132 | 1.01E-16 | 1 |
| rs17763050 | 17 | 43903336 | 17 | 43399058 | 44874453 | A | G | 0.2396 | CRHR1 | intronic | -1.9567 | 0.1802 | 1.81E-27 | -2.2332 | 0.3223 | 4.66E-12 | -0.1096 | 0.0132 | 9.21E-17 | 1 |
| rs62057148 | 17 | 43903485 | 17 | 43399058 | 44874453 | C | G | 0.2396 | CRHR1 | intronic | -1.9534 | 0.1798 | 1.72E-27 | -2.2466 | 0.3216 | 3.11E-12 | -0.1097 | 0.0132 | 9.14E-17 | 1 |
| rs62057149 | 17 | 43903546 | 17 | 43399058 | 44874453 | A | G | 0.2396 | CRHR1 | intronic | 1.9535 | 0.1798 | 1.72E-27 | 2.2467 | 0.3216 | 3.10E-12 | 0.1097 | 0.0132 | 9.20E-17 | 1 |
| rs62057150 | 17 | 43903548 | 17 | 43399058 | 44874453 | T | C | 0.2396 | CRHR1 | intronic | -1.9535 | 0.1798 | 1.71E-27 | -2.2467 | 0.3216 | 3.10E-12 | -0.1097 | 0.0132 | 9.20E-17 | 1 |
| rs62057151 | 17 | 43903842 | 17 | 43399058 | 44874453 | T | C | 0.2465 | CRHR1 | intronic | -1.9045 | 0.1786 | 1.52E-26 | -2.1634 | 0.3175 | 1.04E-11 | -0.1095 | 0.0132 | 1.09E-16 | 1 |
| rs62057152 | 17 | 43904110 | 17 | 43399058 | 44874453 | C | G | 0.2396 | CRHR1 | intronic | -2.0678 | 0.2391 | 5.16E-18 | -2.2467 | 0.3216 | 3.10E-12 | -0.1098 | 0.0132 | 8.81E-17 | 1 |
| rs17689824 | 17 | 43904397 | 17 | 43399058 | 44874453 | T | C | 0.2396 | CRHR1 | intronic | -1.9582 | 0.1799 | 1.36E-27 | -2.2466 | 0.3216 | 3.11E-12 | -0.1098 | 0.0132 | 8.92E-17 | 1 |
| rs62057153 | 17 | 43904528 | 17 | 43399058 | 44874453 | T | C | 0.2396 | CRHR1 | intronic | 1.9574 | 0.1798 | 1.32E-27 | 2.2466 | 0.3216 | 3.11E-12 | 0.1103 | 0.0132 | 6.21E-17 | 1 |
| rs62057154 | 17 | 43904610 | 17 | 43399058 | 44874453 | T | C | 0.2396 | CRHR1 | intronic | -1.9549 | 0.1799 | 1.61E-27 | -2.2466 | 0.3216 | 3.11E-12 | -0.1098 | 0.0132 | 8.64E-17 | 1 |
| rs62057155 | 17 | 43904673 | 17 | 43399058 | 44874453 | T | C | 0.2396 | CRHR1 | intronic | -1.9549 | 0.1799 | 1.62E-27 | -2.2466 | 0.3216 | 3.11E-12 | -0.1098 | 0.0132 | 8.67E-17 | 1 |
| rs62057156 | 17 | 43904948 | 17 | 43399058 | 44874453 | C | G | 0.2396 | CRHR1 | intronic | -1.9551 | 0.1799 | 1.61E-27 | -2.2465 | 0.3216 | 3.13E-12 | -0.1099 | 0.0132 | 8.53E-17 | 1 |
| rs78872653 | 17 | 43905134 | 17 | 43399058 | 44874453 | A | G | 0.2396 | CRHR1 | intronic | -1.9571 | 0.1802 | 1.82E-27 | -2.2464 | 0.3216 | 3.13E-12 | -0.1103 | 0.0133 | 9.02E-17 | 1 |
| rs62057157 | 17 | 43905313 | 17 | 43399058 | 44874453 | C | G | 0.2396 | CRHR1 | intronic | -1.9553 | 0.1799 | 1.60E-27 | -2.2463 | 0.3216 | 3.14E-12 | -0.1099 | 0.0132 | 8.29E-17 | 1 |
| rs17763086 | 17 | 43905481 | 17 | 43399058 | 44874453 | T | G | 0.2396 | CRHR1 | intronic | 1.9548 | 0.1799 | 1.69E-27 | 2.2467 | 0.3216 | 3.11E-12 | 0.1097 | 0.0132 | 9.74E-17 | 1 |
| rs17425752 | 17 | 43906726 | 17 | 43399058 | 44874453 | A | C | 0.2386 | CRHR1 | intronic | 1.9403 | 0.1804 | 5.63E-27 | 2.2560 | 0.3218 | 2.63E-12 | 0.1094 | 0.0133 | 1.58E-16 | 1 |
| rs17689882 | 17 | 43906828 | 17 | 43399058 | 44874453 | A | G | 0.2386 | CRHR1 | intronic | -1.9325 | 0.1804 | 8.73E-27 | -2.2580 | 0.3217 | 2.47E-12 | -0.1089 | 0.0133 | 2.15E-16 | 1 |
| rs62057158 | 17 | 43907143 | 17 | 43399058 | 44874453 | A | G | 0.2396 | CRHR1 | intronic | -1.9554 | 0.18 | 1.68E-27 | -2.2459 | 0.3216 | 3.18E-12 | -0.1095 | 0.0132 | 1.07E-16 | 1 |
| rs1876831 | 17 | 43907745 | 17 | 43399058 | 44874453 | T | C | 0.2396 | CRHR1 | intronic | -1.9574 | 0.18 | 1.52E-27 | -2.2456 | 0.3216 | 3.21E-12 | -0.1096 | 0.0132 | 1.11E-16 | 1 |
| rs16940665 | 17 | 43907896 | 17 | 43399058 | 44874453 | T | C | 0.2396 | CRHR1 | exonic | 1.9556 | 0.18 | 1.67E-27 | 2.2456 | 0.3216 | 3.21E-12 | 0.1099 | 0.0132 | 8.89E-17 | 1 |
| rs16940668 | 17 | 43907966 | 17 | 43399058 | 44874453 | A | G | 0.2396 | CRHR1 | intronic | -1.9552 | 0.18 | 1.72E-27 | -2.2456 | 0.3216 | 3.21E-12 | -0.1098 | 0.0132 | 8.95E-17 | 1 |
| rs16940671 | 17 | 43908151 | 17 | 43399058 | 44874453 | T | C | 0.2396 | CRHR1 | intronic | -2.0299 | 0.1873 | 2.28E-27 | -2.2497 | 0.3221 | 3.17E-12 | -0.1211 | 0.0147 | 1.94E-16 | 1 |
| rs16940672 | 17 | 43908152 | 17 | 43399058 | 44874453 | T | C | 0.2396 | CRHR1 | intronic | -2.0290 | 0.1873 | 2.35E-27 | -2.2493 | 0.3221 | 3.19E-12 | -0.1211 | 0.0147 | 1.95E-16 | 1 |

| | | | | | | | | | | | Meta-GWAS | | | UKBB | | | ENIGMA* | | | |
|---|---|---|---|---|---|---|---|---|---|---|---|---|---|---|---|---|---|---|---|---|
| rsID | chr | pos (hg19) | Genomic Locus | Genomic Locus start | Genomic Locus end | A1 | A2 | MAF | Nearest Genes | Function | Beta | SE | P-value | Beta | SE | P-value | Beta | SE | P-value | Replicated** |
| rs55763795 | 17 | 43908773 | 17 | 43399058 | 44874453 | C | G | 0.2396 | CRHR1 | intronic | -1.9556 | 0.18 | 1.68E-27 | -2.2458 | 0.3216 | 3.18E-12 | -0.1101 | 0.0132 | 9.21E-17 | 1 |
| rs55865707 | 17 | 43908826 | 17 | 43399058 | 44874453 | T | G | 0.2396 | CRHR1 | intronic | 1.9555 | 0.18 | 1.70E-27 | 2.2459 | 0.3216 | 3.18E-12 | 0.1101 | 0.0132 | 9.27E-17 | 1 |
| rs62054760 | 17 | 43908989 | 17 | 43399058 | 44874453 | T | C | 0.2366 | CRHR1 | intronic | -1.9626 | 0.1805 | 1.54E-27 | -2.2948 | 0.3235 | 1.46E-12 | -0.1102 | 0.0133 | 9.71E-17 | 1 |
| rs62054761 | 17 | 43909008 | 17 | 43399058 | 44874453 | T | C | 0.2366 | CRHR1 | intronic | -1.9631 | 0.1805 | 1.50E-27 | -2.2939 | 0.3235 | 1.49E-12 | -0.1101 | 0.0133 | 1.04E-16 | 1 |
| rs62054762 | 17 | 43909022 | 17 | 43399058 | 44874453 | T | G | 0.2366 | CRHR1 | intronic | -1.9643 | 0.1805 | 1.41E-27 | -2.2942 | 0.3236 | 1.48E-12 | -0.1101 | 0.0133 | 1.03E-16 | 1 |
| rs242951 | 17 | 43909412 | 17 | 43399058 | 44874453 | T | C | 0.4463 | CRHR1 | intronic | 0.8272 | 0.1437 | 8.57E-09 | 1.1656 | 0.2731 | 2.01E-05 | 0.0555 | 0.0112 | 7.78E-07 | 1 |
| rs17689918 | 17 | 43910088 | 17 | 43399058 | 44874453 | A | G | 0.2396 | CRHR1 | intronic | -1.9537 | 0.18 | 1.95E-27 | -2.2486 | 0.3219 | 3.12E-12 | -0.1102 | 0.0133 | 1.12E-16 | 1 |
| rs17763199 | 17 | 43910183 | 17 | 43399058 | 44874453 | A | G | 0.2386 | CRHR1 | intronic | -1.9569 | 0.18 | 1.59E-27 | -2.2471 | 0.3216 | 3.11E-12 | -0.1101 | 0.0133 | 1.11E-16 | 1 |
| rs62054763 | 17 | 43910262 | 17 | 43399058 | 44874453 | C | G | 0.2396 | CRHR1 | intronic | 1.9570 | 0.18 | 1.59E-27 | 2.2458 | 0.3216 | 3.18E-12 | 0.1104 | 0.0133 | 1.05E-16 | 1 |
| rs17689966 | 17 | 43910455 | 17 | 43399058 | 44874453 | A | G | 0.4443 | CRHR1 | intronic | 0.8370 | 0.1444 | 6.76E-09 | 1.1752 | 0.2731 | 1.71E-05 | 0.0550 | 0.0112 | 9.84E-07 | 1 |
| rs16940674 | 17 | 43910507 | 17 | 43399058 | 44874453 | T | C | 0.2396 | CRHR1 | exonic | -1.9567 | 0.1801 | 1.67E-27 | -2.2458 | 0.3216 | 3.18E-12 | -0.1104 | 0.0133 | 1.12E-16 | 1 |
| rs16940676 | 17 | 43911036 | 17 | 43399058 | 44874453 | A | G | 0.2386 | CRHR1 | intronic | -1.9539 | 0.1802 | 2.17E-27 | -2.2459 | 0.3216 | 3.18E-12 | -0.1105 | 0.0133 | 1.12E-16 | 1 |
| rs1876830 | 17 | 43911352 | 17 | 43399058 | 44874453 | T | C | 0.2396 | CRHR1 | intronic | -1.9556 | 0.1802 | 1.96E-27 | -2.2459 | 0.3216 | 3.18E-12 | -0.1106 | 0.0133 | 1.11E-16 | 1 |
| rs41457044 | 17 | 43911424 | 17 | 43399058 | 44874453 | T | C | 0.2396 | CRHR1 | intronic | -1.9556 | 0.1802 | 1.97E-27 | -2.2455 | 0.3216 | 3.21E-12 | -0.1106 | 0.0133 | 1.14E-16 | 1 |
| rs1876829 | 17 | 43911443 | 17 | 43399058 | 44874453 | T | C | 0.2396 | CRHR1 | intronic | 1.9559 | 0.1802 | 1.97E-27 | 2.2459 | 0.3216 | 3.18E-12 | 0.1108 | 0.0134 | 1.11E-16 | 1 |
| rs1876828 | 17 | 43911525 | 17 | 43399058 | 44874453 | T | C | 0.2396 | CRHR1 | intronic | -1.9559 | 0.1802 | 1.98E-27 | -2.2458 | 0.3216 | 3.18E-12 | -0.1108 | 0.0134 | 1.15E-16 | 1 |
| rs1876827 | 17 | 43911832 | 17 | 43399058 | 44874453 | T | C | 0.2396 | CRHR1 | intronic | 1.9560 | 0.1803 | 1.99E-27 | 2.2459 | 0.3216 | 3.18E-12 | 0.1110 | 0.0134 | 1.11E-16 | 1 |
| rs16940677 | 17 | 43911898 | 17 | 43399058 | 44874453 | T | C | 0.2396 | CRHR1 | intronic | -1.9589 | 0.1802 | 1.60E-27 | -2.2452 | 0.3217 | 3.25E-12 | -0.1099 | 0.0135 | 3.50E-16 | 1 |
| rs16940681 | 17 | 43912159 | 17 | 43399058 | 44874453 | C | G | 0.2396 | CRHR1 | exonic | -1.9561 | 0.1804 | 2.15E-27 | -2.2545 | 0.3217 | 2.69E-12 | -0.1100 | 0.0135 | 3.57E-16 | 1 |
| rs28364021 | 17 | 43912282 | 17 | 43399058 | 44874453 | T | C | 0.2396 | CRHR1 | UTR3 | -1.9565 | 0.1805 | 2.25E-27 | -2.2459 | 0.3216 | 3.18E-12 | -0.1101 | 0.0135 | 3.52E-16 | 1 |
| rs2316765 | 17 | 43912454 | 17 | 43399058 | 44874453 | T | C | 0.2396 | CRHR1 | UTR3 | 1.9536 | 0.1811 | 4.04E-27 | 2.2474 | 0.3217 | 3.13E-12 | 0.1103 | 0.0135 | 3.41E-16 | 1 |
| rs878886 | 17 | 43912490 | 17 | 43399058 | 44874453 | C | G | 0.2396 | CRHR1 | UTR3 | 1.9537 | 0.1811 | 4.01E-27 | 2.2476 | 0.3217 | 3.11E-12 | 0.1103 | 0.0135 | 3.51E-16 | 1 |
| rs878887 | 17 | 43912582 | 17 | 43399058 | 44874453 | T | C | 0.2396 | CRHR1 | UTR3 | -1.9532 | 0.1814 | 5.03E-27 | -2.2475 | 0.3217 | 3.11E-12 | -0.1104 | 0.0135 | 3.50E-16 | 1 |
| rs878888 | 17 | 43912635 | 17 | 43399058 | 44874453 | A | G | 0.2396 | CRHR1 | UTR3 | 1.9457 | 0.1815 | 8.02E-27 | 2.2473 | 0.3217 | 3.13E-12 | 0.1102 | 0.0135 | 4.01E-16 | 1 |
| rs4525537 | 17 | 43912723 | 17 | 43399058 | 44874453 | T | C | 0.2396 | CRHR1 | UTR3 | 1.9535 | 0.1814 | 4.97E-27 | 2.2468 | 0.3217 | 3.17E-12 | 0.1105 | 0.0135 | 3.42E-16 | 1 |
| rs4640231 | 17 | 43912786 | 17 | 43399058 | 44874453 | C | G | 0.2396 | CRHR1 | UTR3 | -1.9554 | 0.1816 | 5.00E-27 | -2.2473 | 0.3217 | 3.13E-12 | -0.1109 | 0.0136 | 2.93E-16 | 1 |
| rs4482334 | 17 | 43912830 | 17 | 43399058 | 44874453 | T | C | 0.2396 | CRHR1 | UTR3 | 1.9529 | 0.1829 | 1.34E-26 | 2.2473 | 0.3217 | 3.13E-12 | 0.1107 | 0.0136 | 3.83E-16 | 1 |
| rs56127111 | 17 | 43913315 | 17 | 43399058 | 44874453 | T | C | 0.2396 | CRHR1 | downstream | -1.9530 | 0.1829 | 1.33E-26 | -2.2482 | 0.3217 | 3.05E-12 | -0.1109 | 0.0136 | 3.99E-16 | 1 |
| rs242948 | 17 | 43913544 | 17 | 43399058 | 44874453 | T | G | 0.4453 | CRHR1 | downstream | 0.8361 | 0.1487 | 1.87E-08 | 1.1664 | 0.2735 | 2.03E-05 | 0.0570 | 0.0116 | 9.56E-07 | 1 |
| rs75104593 | 17 | 43913557 | 17 | 43399058 | 44874453 | T | G | 0.2416 | CRHR1 | downstream | 1.7913 | 0.1954 | 4.79E-20 | 2.2629 | 0.3251 | 3.71E-12 | 0.1204 | 0.0155 | 6.83E-15 | 1 |
| rs74998289 | 17 | 43913558 | 17 | 43399058 | 44874453 | T | G | 0.2416 | CRHR1 | downstream | 1.7887 | 0.1953 | 5.23E-20 | 2.2643 | 0.3250 | 3.59E-12 | 0.1192 | 0.0154 | 1.10E-14 | 1 |
| rs10445362 | 17 | 43914554 | 17 | 43399058 | 44874453 | A | C | 0.2396 | CRHR1 | intergenic | -1.9800 | 0.1835 | 3.86E-27 | -2.2496 | 0.3217 | 2.98E-12 | -0.1123 | 0.0138 | 3.83E-16 | 1 |
| rs10445363 | 17 | 43914558 | 17 | 43399058 | 44874453 | A | G | 0.2396 | CRHR1 | intergenic | -1.9513 | 0.1834 | 1.96E-26 | -2.2496 | 0.3217 | 2.98E-12 | -0.1104 | 0.0140 | 3.57E-15 | 1 |
| rs62054802 | 17 | 43914598 | 17 | 43399058 | 44874453 | C | G | 0.2396 | CRHR1 | intergenic | -1.9802 | 0.1836 | 3.92E-27 | -2.2495 | 0.3217 | 2.99E-12 | -0.1124 | 0.0138 | 3.83E-16 | 1 |
| rs62054803 | 17 | 43914728 | 17 | 43399058 | 44874453 | T | G | 0.2396 | CRHR1 | intergenic | -1.9804 | 0.1836 | 3.89E-27 | -2.2493 | 0.3217 | 3.00E-12 | -0.1113 | 0.0141 | 2.33E-15 | 1 |
| rs62054804 | 17 | 43914809 | 17 | 43399058 | 44874453 | T | C | 0.2396 | CRHR1 | intergenic | -1.9817 | 0.1835 | 3.44E-27 | -2.2503 | 0.3217 | 2.94E-12 | -0.1113 | 0.0141 | 2.34E-15 | 1 |
| rs62054805 | 17 | 43915054 | 17 | 43399058 | 44874453 | A | G | 0.2396 | CRHR1 | intergenic | 1.9714 | 0.1844 | 1.09E-26 | 2.2495 | 0.3217 | 2.99E-12 | 0.1155 | 0.0147 | 3.95E-15 | 1 |
| rs62054806 | 17 | 43915312 | 17 | 43399058 | 44874453 | T | C | 0.2396 | CRHR1 | intergenic | -2.1203 | 0.2484 | 1.37E-17 | -2.2495 | 0.3217 | 2.99E-12 | -0.1146 | 0.0145 | 3.06E-15 | 1 |
| rs62054807 | 17 | 43915497 | 17 | 43399058 | 44874453 | T | C | 0.3201 | CRHR1 | intergenic | -1.4929 | 0.2229 | 2.12E-11 | -1.5541 | 0.2950 | 1.43E-07 | -0.0900 | 0.0127 | 1.28E-12 | 1 |
| rs10445364 | 17 | 43916356 | 17 | 43399058 | 44874453 | A | G | 0.2396 | CRHR1 | intergenic | -2.1244 | 0.2489 | 1.40E-17 | -2.2495 | 0.3217 | 2.99E-12 | -0.1160 | 0.0147 | 2.61E-15 | 1 |
| rs10445333 | 17 | 43916509 | 17 | 43399058 | 44874453 | A | G | 0.2396 | CRHR1 | intergenic | 2.1188 | 0.249 | 1.76E-17 | 2.2493 | 0.3217 | 3.01E-12 | 0.1168 | 0.0149 | 5.15E-15 | 1 |
| rs17690176 | 17 | 43916773 | 17 | 43399058 | 44874453 | A | C | 0.2406 | CRHR1 | intergenic | 2.1120 | 0.2491 | 2.30E-17 | 2.2651 | 0.3216 | 2.08E-12 | 0.1169 | 0.0149 | 5.26E-15 | 1 |
| rs78328427 | 17 | 43916932 | 17 | 43399058 | 44874453 | A | G | 0.2396 | CRHR1 | intergenic | -2.1132 | 0.2493 | 2.31E-17 | -2.2554 | 0.3218 | 2.66E-12 | -0.1171 | 0.0150 | 5.60E-15 | 1 |
| rs77692262 | 17 | 43917086 | 17 | 43399058 | 44874453 | A | G | 0.2396 | CRHR1 | intergenic | 2.1132 | 0.2493 | 2.31E-17 | 2.2496 | 0.3217 | 2.99E-12 | 0.1172 | 0.0150 | 5.63E-15 | 1 |
| rs56023973 | 17 | 43917776 | 17 | 43399058 | 44874453 | T | G | 0.2396 | MAPT-AS1 | intergenic | -2.1382 | 0.2513 | 1.79E-17 | -2.2492 | 0.3217 | 3.01E-12 | -0.1174 | 0.0150 | 5.61E-15 | 1 |
| rs17763515 | 17 | 43917818 | 17 | 43399058 | 44874453 | A | G | 0.2396 | MAPT-AS1 | intergenic | -2.1386 | 0.2513 | 1.76E-17 | -2.2491 | 0.3217 | 3.01E-12 | -0.1176 | 0.0151 | 5.57E-15 | 1 |
| rs17763533 | 17 | 43918190 | 17 | 43399058 | 44874453 | T | C | 0.2396 | MAPT-AS1 | intergenic | 2.1431 | 0.2513 | 1.49E-17 | 2.2493 | 0.3217 | 3.00E-12 | 0.1181 | 0.0151 | 5.16E-15 | 1 |
| rs62054809 | 17 | 43918239 | 17 | 43399058 | 44874453 | T | C | 0.2396 | MAPT-AS1 | intergenic | -2.1397 | 0.2514 | 1.73E-17 | -2.2479 | 0.3217 | 3.11E-12 | -0.1180 | 0.0151 | 5.77E-15 | 1 |
| rs112583797 | 17 | 43918418 | 17 | 43399058 | 44874453 | A | G | 0.2396 | MAPT-AS1 | intergenic | 1.9877 | 0.1849 | 6.01E-27 | 2.2494 | 0.3217 | 3.00E-12 | 0.1181 | 0.0151 | 5.93E-15 | 1 |
| rs74922289 | 17 | 43918524 | 17 | 43399058 | 44874453 | A | G | 0.2396 | MAPT-AS1 | intergenic | 2.1383 | 0.2514 | 1.79E-17 | 2.2493 | 0.3217 | 3.00E-12 | 0.1182 | 0.0151 | 5.87E-15 | 1 |
| rs56971664 | 17 | 43918613 | 17 | 43399058 | 44874453 | T | C | 0.2396 | MAPT-AS1 | intergenic | 2.1319 | 0.251 | 2.01E-17 | 2.2508 | 0.3217 | 2.89E-12 | 0.1182 | 0.0151 | 6.10E-15 | 1 |
| rs62054811 | 17 | 43918651 | 17 | 43399058 | 44874453 | C | G | 0.2396 | MAPT-AS1 | intergenic | 2.1389 | 0.2514 | 1.77E-17 | 2.2492 | 0.3217 | 3.02E-12 | 0.1184 | 0.0152 | 5.77E-15 | 1 |
| rs2106785 | 17 | 43919105 | 17 | 43399058 | 44874453 | T | C | 0.2386 | MAPT-AS1 | intergenic | -2.1223 | 0.2587 | 2.35E-16 | -2.2404 | 0.3249 | 5.92E-12 | -0.1215 | 0.0155 | 4.09E-15 | 1 |
| rs56150806 | 17 | 43919301 | 17 | 43399058 | 44874453 | T | C | 0.2396 | MAPT-AS1 | intergenic | 2.1358 | 0.2518 | 2.20E-17 | 2.2494 | 0.3217 | 3.00E-12 | 0.1189 | 0.0152 | 5.64E-15 | 1 |
| rs17690314 | 17 | 43919884 | 17 | 43399058 | 44874453 | T | G | 0.2396 | MAPT-AS1 | intergenic | 2.1359 | 0.2518 | 2.19E-17 | 2.2493 | 0.3217 | 3.00E-12 | 0.1190 | 0.0152 | 5.62E-15 | 1 |
| rs56233672 | 17 | 43920300 | 17 | 43399058 | 44874453 | T | C | 0.2396 | MAPT-AS1 | downstream | 2.0733 | 0.2562 | 5.87E-16 | 2.2492 | 0.3217 | 3.01E-12 | 0.1199 | 0.0153 | 4.93E-15 | 1 |
| rs56354807 | 17 | 43920378 | 17 | 43399058 | 44874453 | T | C | 0.2396 | MAPT-AS1 | downstream | -2.1364 | 0.2518 | 2.18E-17 | -2.2492 | 0.3217 | 3.01E-12 | -0.1191 | 0.0152 | 5.82E-15 | 1 |
| rs55718212 | 17 | 43920411 | 17 | 43399058 | 44874453 | T | C | 0.2396 | MAPT-AS1 | downstream | 2.1362 | 0.2518 | 2.19E-17 | 2.2493 | 0.3217 | 3.01E-12 | 0.1191 | 0.0153 | 5.84E-15 | 1 |
| rs17690326 | 17 | 43920974 | 17 | 43399058 | 44874453 | T | C | 0.2396 | MAPT-AS1 | downstream | 2.1360 | 0.2519 | 2.27E-17 | 2.2492 | 0.3217 | 3.01E-12 | 0.1192 | 0.0153 | 5.90E-15 | 1 |
| rs17763596 | 17 | 43921210 | 17 | 43399058 | 44874453 | T | G | 0.2396 | MAPT-AS1 | ncRNA_exonic | -2.1358 | 0.2519 | 2.29E-17 | -2.2491 | 0.3217 | 3.01E-12 | -0.1193 | 0.0153 | 5.56E-15 | 1 |
| rs62054814 | 17 | 43921572 | 17 | 43399058 | 44874453 | A | T | 0.2396 | MAPT-AS1 | ncRNA_intronic | -2.1362 | 0.2519 | 2.27E-17 | -2.2491 | 0.3217 | 3.01E-12 | -0.1193 | 0.0153 | 5.93E-15 | 1 |
| rs17763634 | 17 | 43921754 | 17 | 43399058 | 44874453 | T | C | 0.2396 | MAPT-AS1 | ncRNA_intronic | 2.1360 | 0.2519 | 2.29E-17 | 2.2402 | 0.3218 | 3.70E-12 | 0.1193 | 0.0153 | 5.94E-15 | 1 |
| rs62621252 | 17 | 43922942 | 17 | 43399058 | 44874453 | T | C | 0.2396 | MAPT-AS1:SPPL2C | exonic | 2.1351 | 0.252 | 2.37E-17 | 2.2216 | 0.3216 | 5.43E-12 | 0.1198 | 0.0153 | 5.92E-15 | 1 |
| rs62054815 | 17 | 43923266 | 17 | 43399058 | 44874453 | A | G | 0.2396 | MAPT-AS1:SPPL2C | exonic | -2.1345 | 0.252 | 2.44E-17 | -2.2505 | 0.3217 | 2.90E-12 | -0.1198 | 0.0154 | 5.93E-15 | 1 |
| rs12185233 | 17 | 43923654 | 17 | 43399058 | 44874453 | C | G | 0.2396 | MAPT-AS1:SPPL2C | exonic | -2.1341 | 0.252 | 2.49E-17 | -2.2518 | 0.3218 | 2.87E-12 | -0.1199 | 0.0154 | 5.92E-15 | 1 |
| rs12185268 | 17 | 43923683 | 17 | 43399058 | 44874453 | A | G | 0.2396 | MAPT-AS1:SPPL2C | exonic | 2.1338 | 0.252 | 2.51E-17 | 2.2404 | 0.3215 | 3.54E-12 | 0.1200 | 0.0154 | 5.86E-15 | 1 |
| rs12185235 | 17 | 43923703 | 17 | 43399058 | 44874453 | T | C | 0.2396 | MAPT-AS1:SPPL2C | exonic | -2.1340 | 0.252 | 2.51E-17 | -2.2647 | 0.3216 | 2.09E-12 | -0.1201 | 0.0154 | 5.59E-15 | 1 |
| rs11079725 | 17 | 43923934 | 17 | 43399058 | 44874453 | T | C | 0.2396 | MAPT-AS1:SPPL2C | exonic | 2.1341 | 0.252 | 2.50E-17 | 2.2473 | 0.3215 | 3.04E-12 | 0.1200 | 0.0154 | 5.90E-15 | 1 |
| rs12373123 | 17 | 43924073 | 17 | 43399058 | 44874453 | T | C | 0.2396 | MAPT-AS1:SPPL2C | exonic | 2.1328 | 0.252 | 2.62E-17 | 2.2475 | 0.3215 | 3.03E-12 | 0.1201 | 0.0154 | 5.84E-15 | 1 |
| rs12373139 | 17 | 43924130 | 17 | 43399058 | 44874453 | A | G | 0.2396 | MAPT-AS1:SPPL2C | exonic | -2.1328 | 0.252 | 2.63E-17 | -2.2477 | 0.3215 | 3.01E-12 | -0.1201 | 0.0154 | 5.88E-15 | 1 |
| rs12373142 | 17 | 43924200 | 17 | 43399058 | 44874453 | C | G | 0.2396 | MAPT-AS1:SPPL2C | exonic | 2.0535 | 0.2564 | 1.15E-15 | 2.2616 | 0.3236 | 3.04E-12 | 0.1202 | 0.0155 | 1.02E-14 | 1 |
| rs12373124 | 17 | 43924219 | 17 | 43399058 | 44874453 | T | C | 0.2396 | MAPT-AS1:SPPL2C | exonic | 2.0531 | 0.2564 | 1.17E-15 | 2.2621 | 0.3237 | 3.06E-12 | 0.1208 | 0.0155 | 7.72E-15 | 1 |
| rs12373140 | 17 | 43924231 | 17 | 43399058 | 44874453 | A | G | 0.2396 | MAPT-AS1:SPPL2C | exonic | -2.0530 | 0.2564 | 1.17E-15 | -2.2619 | 0.3237 | 3.07E-12 | -0.1209 | 0.0155 | 7.71E-15 | 1 |
| rs12373168 | 17 | 43924337 | 17 | 43399058 | 44874453 | A | C | 0.2425 | MAPT-AS1:SPPL2C | ncRNA_intronic | 2.1048 | 0.2501 | 3.85E-17 | 2.2035 | 0.3206 | 6.92E-12 | 0.1216 | 0.0157 | 8.79E-15 | 1 |
| rs17690661 | 17 | 43924521 | 17 | 43399058 | 44874453 | A | G | 0.2396 | MAPT-AS1 | ncRNA_intronic | -2.0333 | 0.2611 | 6.85E-15 | -2.2472 | 0.3215 | 3.04E-12 | -0.1213 | 0.0156 | 7.57E-15 | 1 |
| rs55943825 | 17 | 43924776 | 17 | 43399058 | 44874453 | T | C | 0.2396 | MAPT-AS1 | ncRNA_intronic | -2.0267 | 0.2607 | 7.55E-15 | -2.2474 | 0.3215 | 3.03E-12 | -0.1218 | 0.0156 | 6.05E-15 | 1 |
| rs17690679 | 17 | 43924803 | 17 | 43399058 | 44874453 | A | G | 0.2396 | MAPT-AS1 | ncRNA_intronic | 2.0267 | 0.2607 | 7.55E-15 | 2.2474 | 0.3215 | 3.03E-12 | 0.1220 | 0.0156 | 5.99E-15 | 1 |
| rs17690703 | 17 | 43925297 | 17 | 43399058 | 44874453 | T | C | 0.2763 | MAPT-AS1 | ncRNA_intronic | -1.8035 | 0.2474 | 3.08E-13 | -1.8754 | 0.3059 | 9.27E-10 | -0.1012 | 0.0154 | 4.93E-11 | 1 |
| rs17769490 | 17 | 43925605 | 17 | 43399058 | 44874453 | A | G | 0.2406 | MAPT-AS1 | ncRNA_intronic | -2.0004 | 0.2603 | 1.54E-14 | -2.2321 | 0.3205 | 3.64E-12 | -0.1219 | 0.0157 | 7.04E-15 | 1 |
| rs77063768 | 17 | 43925729 | 17 | 43399058 | 44874453 | A | G | 0.2406 | MAPT-AS1 | ncRNA_intronic | 2.0212 | 0.2604 | 8.31E-15 | 2.2321 | 0.3205 | 3.65E-12 | 0.1218 | 0.0157 | 7.94E-15 | 1 |
| rs62054817 | 17 | 43925966 | 17 | 43399058 | 44874453 | A | G | 0.2406 | MAPT-AS1 | ncRNA_intronic | -2.0225 | 0.2604 | 8.05E-15 | -2.2319 | 0.3205 | 3.66E-12 | -0.1220 | 0.0157 | 8.48E-15 | 1 |
| rs62054818 | 17 | 43926056 | 17 | 43399058 | 44874453 | A | C | 0.2406 | MAPT-AS1 | ncRNA_intronic | 2.0204 | 0.2604 | 8.60E-15 | 2.2315 | 0.3205 | 3.69E-12 | 0.1221 | 0.0157 | 8.53E-15 | 1 |
| rs62054819 | 17 | 43926149 | 17 | 43399058 | 44874453 | A | G | 0.2406 | MAPT-AS1 | ncRNA_intronic | 2.0424 | 0.2611 | 5.22E-15 | 2.2314 | 0.3205 | 3.70E-12 | 0.1220 | 0.0158 | 1.33E-14 | 1 |
| rs62054820 | 17 | 43926755 | 17 | 43399058 | 44874453 | A | G | 0.2406 | MAPT-AS1 | ncRNA_intronic | 2.0209 | 0.2605 | 8.56E-15 | 2.2320 | 0.3205 | 3.67E-12 | 0.1223 | 0.0158 | 8.57E-15 | 1 |
| rs56113217 | 17 | 43926948 | 17 | 43399058 | 44874453 | A | G | 0.2406 | MAPT-AS1 | ncRNA_intronic | 2.0209 | 0.2605 | 8.60E-15 | 2.2196 | 0.3209 | 5.05E-12 | 0.1225 | 0.0158 | 8.39E-15 | 1 |
| rs56269136 | 17 | 43926992 | 17 | 43399058 | 44874453 | T | C | 0.2406 | MAPT-AS1 | ncRNA_intronic | -2.0208 | 0.2605 | 8.65E-15 | -2.2305 | 0.3208 | 3.94E-12 | -0.1226 | 0.0158 | 8.56E-15 | 1 |
| rs56385754 | 17 | 43927255 | 17 | 43399058 | 44874453 | T | G | 0.2406 | MAPT-AS1 | ncRNA_intronic | -2.0207 | 0.2605 | 8.65E-15 | -2.2360 | 0.3205 | 3.35E-12 | -0.1227 | 0.0158 | 8.56E-15 | 1 |
| rs17769552 | 17 | 43927290 | 17 | 43399058 | 44874453 | A | G | 0.2326 | MAPT-AS1 | ncRNA_intronic | -1.7713 | 0.2736 | 9.48E-11 | -2.2569 | 0.3234 | 3.28E-12 | -0.1250 | 0.0161 | 9.78E-15 | 1 |
| rs62054822 | 17 | 43927708 | 17 | 43399058 | 44874453 | A | G | 0.2406 | MAPT-AS1 | ncRNA_intronic | 2.0225 | 0.26 | 7.26E-15 | 2.2241 | 0.3201 | 4.09E-12 | 0.1230 | 0.0159 | 8.84E-15 | 1 |
| rs885639 | 17 | 43928614 | 17 | 43399058 | 44874453 | T | G | 0.2406 | MAPT-AS1 | ncRNA_intronic | 2.0210 | 0.2594 | 6.73E-15 | 2.2414 | 0.3202 | 2.84E-12 | 0.1231 | 0.0159 | 1.03E-14 | 1 |
| rs62054823 | 17 | 43929029 | 17 | 43399058 | 44874453 | A | G | 0.2406 | MAPT-AS1 | ncRNA_intronic | 2.0119 | 0.2607 | 1.20E-14 | 2.2246 | 0.3210 | 4.64E-12 | 0.1240 | 0.0159 | 7.08E-15 | 1 |
| rs62054824 | 17 | 43929992 | 17 | 43399058 | 44874453 | T | C | 0.2396 | MAPT-AS1 | ncRNA_intronic | -2.0265 | 0.2615 | 9.16E-15 | -2.2349 | 0.3214 | 3.94E-12 | -0.1252 | 0.0162 | 1.26E-14 | 1 |
| rs62054825 | 17 | 43930033 | 17 | 43399058 | 44874453 | A | G | 0.2406 | MAPT-AS1 | ncRNA_intronic | -2.0273 | 0.2614 | 8.82E-15 | -2.2339 | 0.3214 | 4.02E-12 | -0.1250 | 0.0162 | 1.40E-14 | 1 |
| rs79589869 | 17 | 43930238 | 17 | 43399058 | 44874453 | A | C | 0.2396 | MAPT-AS1 | ncRNA_intronic | -2.0275 | 0.2615 | 9.04E-15 | -2.2374 | 0.3216 | 3.84E-12 | -0.1253 | 0.0163 | 1.27E-14 | 1 |
| rs2316769 | 17 | 43930798 | 17 | 43399058 | 44874453 | A | G | 0.2396 | MAPT-AS1 | ncRNA_intronic | 1.7964 | 0.2707 | 3.23E-11 | 2.2301 | 0.3214 | 4.34E-12 | 0.1299 | 0.0167 | 7.84E-15 | 1 |
| rs2873268 | 17 | 43930955 | 17 | 43399058 | 44874453 | A | G | 0.2396 | MAPT-AS1 | ncRNA_intronic | -1.7955 | 0.2708 | 3.32E-11 | -2.2447 | 0.3216 | 3.25E-12 | -0.1300 | 0.0167 | 8.36E-15 | 1 |

| | | | | | | | | | | | Meta-GWAS | | | UKBB | | | ENIGMA* | | | |
|---|---|---|---|---|---|---|---|---|---|---|---|---|---|---|---|---|---|---|---|---|
| rsID | chr | pos (hg19) | Genomic Locus | Genomic Locus start | Genomic Locus end | A1 | A2 | MAF | Nearest Genes | Function | Beta | SE | P-value | Beta | SE | P-value | Beta | SE | P-value | Replicated** |
| rs113856644 | 17 | 43932277 | 17 | 43399058 | 44874453 | A | G | 0.2396 | MAPT-AS1 | ncRNA_intronic | -1.7630 | 0.1949 | 1.49E-19 | -2.2260 | 0.3214 | 4.73E-12 | NA | NA | NA | NA |
| rs76563578 | 17 | 43933879 | 17 | 43399058 | 44874453 | C | G | 0.2396 | MAPT-AS1 | ncRNA_intronic | 1.7496 | 0.1941 | 1.97E-19 | 2.2031 | 0.3212 | 7.58E-12 | NA | NA | NA | NA |
| rs111962225 | 17 | 43934016 | 17 | 43399058 | 44874453 | C | G | 0.2396 | MAPT-AS1 | ncRNA_intronic | -1.7490 | 0.1941 | 2.02E-19 | -2.2008 | 0.3212 | 7.99E-12 | NA | NA | NA | NA |
| rs62054835 | 17 | 43934672 | 17 | 43399058 | 44874453 | A | C | 0.2396 | MAPT-AS1 | ncRNA_intronic | 1.7514 | 0.1941 | 1.83E-19 | 2.1999 | 0.3211 | 8.09E-12 | NA | NA | NA | NA |
| rs113414067 | 17 | 43955093 | 17 | 43399058 | 44874453 | T | C | 0.2396 | MAPT-AS1 | ncRNA_intronic | 1.7504 | 0.1939 | 1.76E-19 | 2.2023 | 0.3212 | 7.75E-12 | NA | NA | NA | NA |
| rs62056785 | 17 | 43975263 | 17 | 43399058 | 44874453 | T | C | 0.2396 | MAPT | intronic | 1.7524 | 0.1939 | 1.60E-19 | 2.2029 | 0.3212 | 7.67E-12 | NA | NA | NA | NA |
| rs62056786 | 17 | 43975285 | 17 | 43399058 | 44874453 | T | C | 0.2396 | MAPT | intronic | 1.7469 | 0.1939 | 2.04E-19 | 2.2029 | 0.3212 | 7.67E-12 | NA | NA | NA | NA |
| rs62056789 | 17 | 43975415 | 17 | 43399058 | 44874453 | A | C | 0.2396 | MAPT | intronic | 1.7582 | 0.1945 | 1.56E-19 | 2.2030 | 0.3212 | 7.66E-12 | NA | NA | NA | NA |
| rs62056790 | 17 | 43975417 | 17 | 43399058 | 44874453 | A | G | 0.2396 | MAPT | intronic | -1.7581 | 0.1945 | 1.57E-19 | -2.2030 | 0.3212 | 7.66E-12 | NA | NA | NA | NA |
| rs113589236 | 17 | 43981795 | 17 | 43399058 | 44874453 | A | G | 0.2396 | MAPT | intronic | -1.7176 | 0.1951 | 1.33E-18 | -2.2037 | 0.3212 | 7.56E-12 | NA | NA | NA | NA |
| rs113029914 | 17 | 43981831 | 17 | 43399058 | 44874453 | A | T | 0.2396 | MAPT | intronic | 1.7243 | 0.1952 | 9.99E-19 | 2.2037 | 0.3212 | 7.56E-12 | NA | NA | NA | NA |
| rs112275277 | 17 | 43981958 | 17 | 43399058 | 44874453 | T | C | 0.2396 | MAPT | intronic | -1.7104 | 0.1944 | 1.38E-18 | -2.2030 | 0.3212 | 7.66E-12 | NA | NA | NA | NA |
| rs62056851 | 17 | 43992806 | 17 | 43399058 | 44874453 | A | G | 0.2396 | MAPT | intronic | 1.7104 | 0.1944 | 1.37E-18 | 2.1974 | 0.3214 | 8.83E-12 | NA | NA | NA | NA |
| rs9899833 | 17 | 43992943 | 17 | 43399058 | 44874453 | A | G | 0.3608 | MAPT | intronic | -1.0198 | 0.1746 | 5.18E-09 | -1.7456 | 0.2850 | 9.56E-10 | NA | NA | NA | NA |
| rs111541901 | 17 | 43994358 | 17 | 43399058 | 44874453 | T | C | 0.2396 | MAPT | intronic | -1.7080 | 0.1948 | 1.79E-18 | -2.1815 | 0.3218 | 1.32E-11 | NA | NA | NA | NA |
| rs112647192 | 17 | 43994623 | 17 | 43399058 | 44874453 | A | G | 0.2396 | MAPT | intronic | -1.7092 | 0.1944 | 1.45E-18 | -2.1969 | 0.3214 | 8.94E-12 | NA | NA | NA | NA |
| rs62059005 | 17 | 44004472 | 17 | 43399058 | 44874453 | A | G | 0.2396 | MAPT | intronic | 1.7098 | 0.1944 | 1.45E-18 | 2.2034 | 0.3212 | 7.60E-12 | NA | NA | NA | NA |
| rs113796169 | 17 | 44005254 | 17 | 43399058 | 44874453 | A | T | 0.2406 | MAPT | intronic | -1.7089 | 0.1945 | 1.58E-18 | -2.2013 | 0.3213 | 7.99E-12 | NA | NA | NA | NA |
| rs112454267 | 17 | 44005329 | 17 | 43399058 | 44874453 | A | G | 0.2406 | MAPT | intronic | -1.7154 | 0.1944 | 1.12E-18 | -2.2013 | 0.3213 | 7.99E-12 | NA | NA | NA | NA |
| rs78026984 | 17 | 44025592 | 17 | 43399058 | 44874453 | C | G | 0.2396 | MAPT | intronic | -1.7373 | 0.1956 | 6.53E-19 | -2.2161 | 0.3214 | 5.90E-12 | NA | NA | NA | NA |
| rs113520245 | 17 | 44033132 | 17 | 43399058 | 44874453 | T | C | 0.2396 | MAPT | intronic | -1.7524 | 0.1973 | 6.50E-19 | -2.2325 | 0.3270 | 9.50E-12 | NA | NA | NA | NA |
| rs62063271 | 17 | 44036047 | 17 | 43399058 | 44874453 | A | G | 0.2396 | MAPT | intronic | -1.7575 | 0.1955 | 2.44E-19 | -2.2245 | 0.3215 | 4.99E-12 | NA | NA | NA | NA |
| rs62641967 | 17 | 44047216 | 17 | 43399058 | 44874453 | T | G | 0.2406 | MAPT | intronic | 1.7517 | 0.1953 | 2.94E-19 | 2.2173 | 0.3215 | 5.85E-12 | NA | NA | NA | NA |
| rs117124984 | 17 | 44051588 | 17 | 43399058 | 44874453 | C | G | 0.2406 | MAPT | UTR5 | 1.7903 | 0.1991 | 2.41E-19 | 2.2264 | 0.3269 | 1.06E-11 | NA | NA | NA | NA |
| rs118087478 | 17 | 44051589 | 17 | 43399058 | 44874453 | T | G | 0.2406 | MAPT | UTR5 | 1.7955 | 0.1991 | 1.88E-19 | 2.2265 | 0.3269 | 1.06E-11 | NA | NA | NA | NA |
| rs112385572 | 17 | 44066172 | 17 | 43399058 | 44874453 | A | G | 0.2406 | MAPT | intronic | 1.7487 | 0.1952 | 3.26E-19 | 2.2175 | 0.3214 | 5.72E-12 | NA | NA | NA | NA |
| rs112572874 | 17 | 44072984 | 17 | 43399058 | 44874453 | A | G | 0.2406 | MAPT | intronic | 1.7292 | 0.1949 | 7.27E-19 | 2.1708 | 0.3213 | 1.54E-11 | NA | NA | NA | NA |
| rs62062278 | 17 | 44093860 | 17 | 43399058 | 44874453 | A | G | 0.2406 | MAPT | intronic | 1.7451 | 0.1955 | 4.31E-19 | 2.2178 | 0.3214 | 5.68E-12 | NA | NA | NA | NA |
| rs112578465 | 17 | 44125066 | 17 | 43399058 | 44874453 | T | C | 0.2406 | KANSL1 | intronic | -1.7455 | 0.1953 | 4.04E-19 | -2.2174 | 0.3214 | 5.73E-12 | NA | NA | NA | NA |
| rs112746008 | 17 | 44126650 | 17 | 43399058 | 44874453 | T | C | 0.2406 | KANSL1 | intronic | -1.7655 | 0.1971 | 3.34E-19 | -2.2122 | 0.3214 | 6.45E-12 | NA | NA | NA | NA |
| rs112333322 | 17 | 44126673 | 17 | 43399058 | 44874453 | A | G | 0.2406 | KANSL1 | intronic | 1.7418 | 0.1968 | 8.74E-19 | 2.2483 | 0.3230 | 3.73E-12 | NA | NA | NA | NA |
| rs111327992 | 17 | 44126691 | 17 | 43399058 | 44874453 | A | G | 0.2406 | KANSL1 | intronic | -1.7346 | 0.1953 | 6.47E-19 | -2.1932 | 0.3213 | 9.59E-12 | NA | NA | NA | NA |
| rs113434679 | 17 | 44126765 | 17 | 43399058 | 44874453 | A | C | 0.2028 | KANSL1 | intronic | -1.6822 | 0.2087 | 7.64E-16 | -2.0696 | 0.3387 | 1.05E-09 | NA | NA | NA | NA |
| rs17575507 | 17 | 44134095 | 17 | 43399058 | 44874453 | A | G | 0.2406 | KANSL1 | intronic | 1.7394 | 0.1953 | 5.29E-19 | 2.2117 | 0.3214 | 6.52E-12 | NA | NA | NA | NA |
| rs111372048 | 17 | 44136577 | 17 | 43399058 | 44874453 | A | C | 0.2406 | KANSL1 | intronic | 1.7328 | 0.1953 | 7.26E-19 | 2.2035 | 0.3215 | 7.85E-12 | NA | NA | NA | NA |
| rs112197756 | 17 | 44154105 | 17 | 43399058 | 44874453 | A | G | 0.2406 | KANSL1 | intronic | 1.7352 | 0.1954 | 6.58E-19 | 2.2178 | 0.3214 | 5.68E-12 | NA | NA | NA | NA |
| rs111913701 | 17 | 44159631 | 17 | 43399058 | 44874453 | T | G | 0.2406 | KANSL1 | intronic | 1.7346 | 0.1954 | 6.81E-19 | 2.2177 | 0.3214 | 5.69E-12 | NA | NA | NA | NA |
| rs111519055 | 17 | 44159672 | 17 | 43399058 | 44874453 | A | G | 0.2406 | KANSL1 | intronic | 1.7347 | 0.1954 | 6.80E-19 | 2.2177 | 0.3214 | 5.69E-12 | NA | NA | NA | NA |
| rs113788190 | 17 | 44161302 | 17 | 43399058 | 44874453 | A | G | 0.2406 | KANSL1 | intronic | 1.7346 | 0.1954 | 6.83E-19 | 2.2177 | 0.3214 | 5.70E-12 | NA | NA | NA | NA |
| rs112364920 | 17 | 44161360 | 17 | 43399058 | 44874453 | A | T | 0.2396 | KANSL1 | intronic | 1.7346 | 0.1954 | 6.83E-19 | 2.2177 | 0.3214 | 5.70E-12 | NA | NA | NA | NA |
| rs80028338 | 17 | 44161470 | 17 | 43399058 | 44874453 | A | C | 0.2406 | KANSL1 | intronic | 1.7764 | 0.2035 | 2.59E-18 | 2.3703 | 0.3421 | 4.65E-12 | NA | NA | NA | NA |
| rs111970616 | 17 | 44169581 | 17 | 43399058 | 44874453 | T | C | 0.2406 | KANSL1 | intronic | 1.7816 | 0.1966 | 1.26E-19 | 2.2166 | 0.3214 | 5.85E-12 | NA | NA | NA | NA |
| rs112596352 | 17 | 44170238 | 17 | 43399058 | 44874453 | A | G | 0.2406 | KANSL1 | intronic | -1.7259 | 0.1955 | 1.05E-18 | -2.1992 | 0.3213 | 8.36E-12 | NA | NA | NA | NA |
| rs111676341 | 17 | 44183403 | 17 | 43399058 | 44874453 | A | G | 0.2406 | KANSL1 | intronic | 1.7642 | 0.1965 | 2.80E-19 | 2.1985 | 0.3214 | 8.62E-12 | NA | NA | NA | NA |
| rs112560196 | 17 | 44200078 | 17 | 43399058 | 44874453 | A | T | 0.2406 | KANSL1 | intronic | 1.7445 | 0.1962 | 6.00E-19 | 2.1970 | 0.3215 | 9.01E-12 | NA | NA | NA | NA |
| rs112073200 | 17 | 44201791 | 17 | 43399058 | 44874453 | C | G | 0.2406 | KANSL1 | intronic | -1.7299 | 0.1967 | 1.45E-18 | -2.1887 | 0.3216 | 1.09E-11 | NA | NA | NA | NA |
| rs55669501 | 17 | 44202564 | 17 | 43399058 | 44874453 | A | G | 0.2406 | KANSL1 | intronic | -1.7436 | 0.1965 | 7.02E-19 | -2.1888 | 0.3215 | 1.08E-11 | NA | NA | NA | NA |
| rs55686102 | 17 | 44202608 | 17 | 43399058 | 44874453 | T | C | 0.2406 | KANSL1 | intronic | -1.7436 | 0.1965 | 7.03E-19 | -2.1888 | 0.3215 | 1.09E-11 | NA | NA | NA | NA |
| NA | 17 | 44699851 | 17 | 43399058 | 44874453 | A | G | 0.2197 | NSF | intronic | -2.5768 | 0.3085 | 6.72E-17 | -2.2087 | 0.3374 | 6.35E-11 | -0.1189 | 0.0195 | 1.19E-09 | 1 |
| rs117300236 | 17 | 44753350 | 17 | 43399058 | 44874453 | A | G | 0.2843 | NSF | intronic | -1.9285 | 0.1977 | 1.76E-22 | -1.7289 | 0.3222 | 8.36E-08 | NA | NA | NA | NA |
| rs199461 | 17 | 44762589 | 17 | 43399058 | 44874453 | A | G | 0.2555 | NSF | intronic | -1.7751 | 0.2101 | 2.94E-17 | -1.9600 | 0.3171 | 6.80E-10 | -0.0900 | 0.0129 | 2.91E-12 | 1 |
| rs199460 | 17 | 44764775 | 17 | 43399058 | 44874453 | A | C | 0.2187 | NSF | intronic | 2.0439 | 0.2516 | 4.50E-16 | 2.1775 | 0.3491 | 4.73E-10 | 0.1045 | 0.0150 | 3.72E-12 | 1 |
| rs199441 | 17 | 44773783 | 17 | 43399058 | 44874453 | A | G | 0.2306 | NSF | intronic | -2.0726 | 0.2125 | 1.79E-22 | -2.1373 | 0.3259 | 5.88E-11 | -0.1072 | 0.0130 | 1.30E-16 | 1 |
| rs199437 | 17 | 44786336 | 17 | 43399058 | 44874453 | A | T | 0.2575 | NSF | intronic | -1.6670 | 0.1605 | 2.78E-25 | -1.9276 | 0.3164 | 1.19E-09 | -0.0844 | 0.0120 | 1.74E-12 | 1 |
| rs1378358 | 17 | 44787312 | 17 | 43399058 | 44874453 | T | C | 0.2247 | NSF | intronic | -2.0049 | 0.171 | 9.55E-32 | -2.2033 | 0.3276 | 1.91E-11 | -0.1091 | 0.0130 | 4.12E-17 | 1 |
| rs538628 | 17 | 44787313 | 17 | 43399058 | 44874453 | C | G | 0.2247 | NSF | intronic | -2.0044 | 0.171 | 9.74E-32 | -2.1925 | 0.3276 | 2.38E-11 | -0.1089 | 0.0130 | 4.16E-17 | 1 |
| rs183211 | 17 | 44788310 | 17 | 43399058 | 44874453 | A | G | 0.2584 | NSF | intronic | -1.6535 | 0.16 | 4.77E-25 | -1.9271 | 0.3164 | 1.19E-09 | -0.0840 | 0.0119 | 1.69E-12 | 1 |
| rs199436 | 17 | 44789285 | 17 | 43399058 | 44874453 | A | G | 0.2575 | NSF | intronic | 1.6504 | 0.1599 | 5.47E-25 | 1.9271 | 0.3164 | 1.19E-09 | 0.0839 | 0.0119 | 1.62E-12 | 1 |
| rs169201 | 17 | 44790203 | 17 | 43399058 | 44874453 | A | G | 0.2247 | NSF | intronic | 1.9379 | 0.1672 | 4.66E-31 | 2.2175 | 0.3276 | 1.41E-11 | 0.1050 | 0.0125 | 4.53E-17 | 1 |
| rs199438 | 17 | 44791643 | 17 | 43399058 | 44874453 | A | G | 0.2575 | NSF | intronic | -1.6479 | 0.1597 | 5.83E-25 | -1.9273 | 0.3164 | 1.19E-09 | -0.0839 | 0.0118 | 1.43E-12 | 1 |
| rs17692129 | 17 | 44793283 | 17 | 43399058 | 44874453 | T | C | 0.34 | NSF | intronic | 1.0218 | 0.1596 | 1.53E-10 | 1.1869 | 0.2842 | 3.00E-05 | 0.0573 | 0.0125 | 4.32E-06 | 1 |
| rs199439 | 17 | 44793503 | 17 | 43399058 | 44874453 | A | G | 0.2247 | NSF | intronic | 1.9309 | 0.1671 | 6.72E-31 | 2.1957 | 0.3276 | 2.22E-11 | 0.1048 | 0.0125 | 3.94E-17 | 1 |
| rs142167 | 17 | 44795234 | 17 | 43399058 | 44874453 | A | G | 0.2575 | NSF | intronic | 1.6441 | 0.1596 | 6.84E-25 | 1.9241 | 0.3164 | 1.27E-09 | 0.0837 | 0.0118 | 1.23E-12 | 1 |
| rs199457 | 17 | 44795469 | 17 | 43399058 | 44874453 | T | C | 0.2247 | NSF | intronic | -1.9316 | 0.167 | 6.29E-31 | -2.2140 | 0.3276 | 1.52E-11 | -0.1046 | 0.0124 | 3.84E-17 | 1 |
| rs199456 | 17 | 44797919 | 17 | 43399058 | 44874453 | T | C | 0.2247 | NSF | intronic | -1.9254 | 0.167 | 9.74E-31 | -2.2058 | 0.3277 | 1.84E-11 | -0.1045 | 0.0124 | 3.87E-17 | 1 |
| rs199455 | 17 | 44799216 | 17 | 43399058 | 44874453 | T | C | 0.2614 | NSF:RPS7P11 | ncRNA_exonic | 1.6216 | 0.1593 | 2.40E-24 | 1.8856 | 0.3159 | 2.53E-09 | 0.0831 | 0.0118 | 1.61E-12 | 1 |
| rs7224296 | 17 | 44800046 | 17 | 43399058 | 44874453 | A | G | 0.3022 | NSF | intronic | 1.4177 | 0.1545 | 4.53E-20 | 1.7800 | 0.3021 | 4.02E-09 | 0.0768 | 0.0115 | 2.66E-11 | 1 |
| rs199454 | 17 | 44800110 | 17 | 43399058 | 44874453 | A | G | 0.2594 | NSF | intronic | 1.6287 | 0.1595 | 1.71E-24 | 1.8933 | 0.3158 | 2.15E-09 | 0.0830 | 0.0118 | 1.62E-12 | 1 |
| rs199453 | 17 | 44800946 | 17 | 43399058 | 44874453 | T | C | 0.2584 | NSF | intronic | -1.6439 | 0.1596 | 7.17E-25 | -1.9237 | 0.3166 | 1.30E-09 | -0.0831 | 0.0118 | 1.60E-12 | 1 |
| rs199452 | 17 | 44801340 | 17 | 43399058 | 44874453 | T | C | 0.2565 | NSF | intronic | -1.6403 | 0.1597 | 9.68E-25 | -1.9361 | 0.3169 | 1.05E-09 | -0.0836 | 0.0118 | 1.17E-12 | 1 |
| rs199451 | 17 | 44801784 | 17 | 43399058 | 44874453 | A | G | 0.2227 | NSF | intronic | -1.9281 | 0.1671 | 8.17E-31 | -2.2194 | 0.3277 | 1.37E-11 | -0.1052 | 0.0124 | 2.18E-17 | 1 |
| rs35937770 | 17 | 44808360 | 17 | 43399058 | 44874453 | A | G | 0.339 | NSF | intronic | 0.9826 | 0.1592 | 6.70E-10 | 1.2214 | 0.2858 | 1.96E-05 | 0.0602 | 0.0125 | 1.57E-06 | 1 |
| rs199449 | 17 | 44808902 | 17 | 43399058 | 44874453 | A | G | 0.2555 | NSF | intronic | -1.6343 | 0.1599 | 1.57E-24 | -1.9408 | 0.3168 | 9.56E-10 | -0.0839 | 0.0118 | 1.06E-12 | 1 |
| rs199448 | 17 | 44809001 | 17 | 43399058 | 44874453 | A | G | 0.2227 | NSF | intronic | 1.9246 | 0.1672 | 1.15E-30 | 2.2277 | 0.3279 | 1.19E-11 | 0.1054 | 0.0124 | 2.05E-17 | 1 |
| rs199447 | 17 | 44812188 | 17 | 43399058 | 44874453 | T | C | 0.2227 | NSF | intronic | -1.9220 | 0.1672 | 1.39E-30 | -2.2267 | 0.3279 | 1.22E-11 | -0.1054 | 0.0124 | 2.01E-17 | 1 |
| rs199446 | 17 | 44813169 | 17 | 43399058 | 44874453 | A | G | 0.2575 | NSF | intronic | 1.6239 | 0.1597 | 2.81E-24 | 1.9073 | 0.3160 | 1.68E-09 | 0.0836 | 0.0118 | 1.29E-12 | 1 |
| rs199445 | 17 | 44817408 | 17 | 43399058 | 44874453 | T | C | 0.2227 | NSF | intronic | -1.9133 | 0.1672 | 2.64E-30 | -2.2057 | 0.3279 | 1.88E-11 | -0.1056 | 0.0124 | 1.69E-17 | 1 |
| rs199444 | 17 | 44818276 | 17 | 43399058 | 44874453 | T | C | 0.2555 | NSF | intronic | 1.6317 | 0.1601 | 2.14E-24 | 1.9367 | 0.3168 | 1.03E-09 | 0.0849 | 0.0118 | 6.01E-13 | 1 |
| rs199443 | 17 | 44819565 | 17 | 43399058 | 44874453 | T | C | 0.2227 | NSF | intronic | -1.9144 | 0.1673 | 2.49E-30 | -2.2029 | 0.3279 | 1.99E-11 | -0.1059 | 0.0124 | 1.37E-17 | 1 |
| rs199442 | 17 | 44820122 | 17 | 43399058 | 44874453 | A | G | 0.2555 | NSF | intronic | -1.6277 | 0.1601 | 2.88E-24 | -1.9347 | 0.3168 | 1.07E-09 | -0.0853 | 0.0118 | 4.94E-13 | 1 |
| rs199536 | 17 | 44820425 | 17 | 43399058 | 44874453 | T | C | 0.2575 | NSF | intronic | -1.6194 | 0.1599 | 4.26E-24 | -1.9033 | 0.3160 | 1.80E-09 | -0.0847 | 0.0118 | 6.50E-13 | 1 |
| rs199535 | 17 | 44822662 | 17 | 43399058 | 44874453 | A | G | 0.2227 | NSF | intronic | 1.9174 | 0.1673 | 2.03E-30 | 2.2015 | 0.3278 | 2.03E-11 | 0.1063 | 0.0124 | 9.80E-18 | 1 |
| rs199534 | 17 | 44824213 | 17 | 43399058 | 44874453 | T | G | 0.2227 | NSF | intronic | 1.9156 | 0.1672 | 2.25E-30 | 2.2212 | 0.3278 | 1.35E-11 | 0.1059 | 0.0124 | 1.32E-17 | 1 |
| rs9896243 | 17 | 44826056 | 17 | 43399058 | 44874453 | C | G | 0.2256 | NSF | intronic | 1.9152 | 0.1672 | 2.23E-30 | 2.1692 | 0.3272 | 3.66E-11 | 0.1061 | 0.0124 | 1.20E-17 | 1 |
| rs199533 | 17 | 44828931 | 17 | 43399058 | 44874453 | A | G | 0.2227 | NSF | exonic | -1.9104 | 0.1671 | 2.77E-30 | -2.1675 | 0.3264 | 3.41E-11 | -0.1063 | 0.0124 | 9.98E-18 | 1 |
| rs199531 | 17 | 44830414 | 17 | 43399058 | 44874453 | T | C | 0.2565 | NSF | intronic | 1.6310 | 0.1601 | 2.20E-24 | 1.8917 | 0.3152 | 2.06E-09 | 0.0863 | 0.0118 | 2.94E-13 | 1 |
| rs9912530 | 17 | 44836302 | 17 | 43399058 | 44874453 | T | C | 0.3002 | NSF | intergenic | -1.4129 | 0.1555 | 1.06E-19 | -1.7485 | 0.3013 | 6.80E-09 | -0.0809 | 0.0116 | 3.69E-12 | 1 |
| rs199530 | 17 | 44836653 | 17 | 43399058 | 44874453 | A | G | 0.2604 | NSF | intergenic | 1.6122 | 0.16 | 7.19E-24 | 1.8552 | 0.3148 | 4.00E-09 | 0.0856 | 0.0118 | 4.86E-13 | 1 |
| rs199529 | 17 | 44837217 | 17 | 43399058 | 44874453 | A | C | 0.2425 | NSF | intergenic | 1.8033 | 0.1642 | 4.68E-28 | 2.0280 | 0.3187 | 2.11E-10 | 0.0982 | 0.0122 | 9.45E-16 | 1 |
| rs199528 | 17 | 44843136 | 17 | 43399058 | 44874453 | T | C | 0.2207 | WNT3 | intronic | -1.9060 | 0.1681 | 8.55E-30 | -2.1992 | 0.3263 | 1.72E-11 | -0.1076 | 0.0125 | 8.36E-18 | 1 |
| rs199527 | 17 | 44843667 | 17 | 43399058 | 44874453 | A | G | 0.2575 | WNT3 | intronic | 1.6287 | 0.1616 | 7.02E-24 | 1.8801 | 0.3145 | 2.40E-09 | 0.0871 | 0.0120 | 3.58E-13 | 1 |
| rs199526 | 17 | 44847707 | 17 | 43399058 | 44874453 | C | G | 0.2366 | WNT3 | intronic | -1.7499 | 0.1656 | 4.09E-26 | -2.0385 | 0.3212 | 2.37E-10 | -0.0971 | 0.0123 | 2.85E-15 | 1 |
| rs199525 | 17 | 44847834 | 17 | 43399058 | 44874453 | T | G | 0.2207 | WNT3 | intronic | 1.9041 | 0.168 | 8.89E-30 | 2.2045 | 0.3260 | 1.48E-11 | 0.1074 | 0.0125 | 9.21E-18 | 1 |
| rs3809857 | 17 | 44848314 | 17 | 43399058 | 44874453 | T | G | 0.331 | WNT3 | intronic | 1.0365 | 0.1653 | 3.57E-10 | 1.1860 | 0.2865 | 3.52E-05 | 0.0620 | 0.0130 | 1.95E-06 | 1 |
| rs199524 | 17 | 44848438 | 17 | 43399058 | 44874453 | T | G | 0.2575 | WNT3 | intronic | 1.6151 | 0.1614 | 1.38E-23 | 1.8934 | 0.3151 | 1.97E-09 | 0.0869 | 0.0119 | 3.38E-13 | 1 |
| rs199523 | 17 | 44848517 | 17 | 43399058 | 44874453 | A | C | 0.2575 | WNT3 | intronic | 1.6005 | 0.1617 | 4.21E-23 | 1.8950 | 0.3151 | 1.91E-09 | 0.0864 | 0.0120 | 7.11E-13 | 1 |

| | | | | | | | | | | | Meta-GWAS | | | UKBB | | | ENIGMA* | | | |
|---|---|---|---|---|---|---|---|---|---|---|---|---|---|---|---|---|---|---|---|---|
| rsID | chr | pos (hg19) | Genomic Locus | Genomic Locus start | Genomic Locus end | A1 | A2 | MAF | Nearest Genes | Function | Beta | SE | P-value | Beta | SE | P-value | Beta | SE | P-value | Replicated** |
| rs199521 | 17 | 44853456 | 17 | 43399058 | 44874453 | C | G | 0.2555 | WNT3 | intronic | 1.6389 | 0.1616 | 3.64E-24 | 1.8990 | 0.3152 | 1.80E-09 | 0.0856 | 0.0119 | 6.87E-13 | 1 |
| rs199520 | 17 | 44853872 | 17 | 43399058 | 44874453 | A | G | 0.2575 | WNT3 | intronic | 1.6391 | 0.1616 | 3.50E-24 | 1.9083 | 0.3152 | 1.50E-09 | 0.0856 | 0.0119 | 6.72E-13 | 1 |
| rs199519 | 17 | 44853924 | 17 | 43399058 | 44874453 | A | G | 0.2575 | WNT3 | intronic | 1.6447 | 0.1615 | 2.34E-24 | 1.8957 | 0.3150 | 1.86E-09 | 0.0856 | 0.0119 | 6.90E-13 | 1 |
| rs199518 | 17 | 44854580 | 17 | 43399058 | 44874453 | A | C | 0.2575 | WNT3 | intronic | -1.6550 | 0.1632 | 3.68E-24 | -1.8810 | 0.3158 | 2.73E-09 | -0.0896 | 0.0122 | 2.30E-13 | 1 |
| rs199517 | 17 | 44854587 | 17 | 43399058 | 44874453 | A | G | 0.2575 | WNT3 | intronic | -1.6530 | 0.1631 | 3.87E-24 | -1.8807 | 0.3157 | 2.71E-09 | -0.0891 | 0.0122 | 2.85E-13 | 1 |
| rs199516 | 17 | 44856485 | 17 | 43399058 | 44874453 | T | C | 0.2207 | WNT3 | intronic | 1.9081 | 0.1678 | 5.74E-30 | 2.2211 | 0.3249 | 8.96E-12 | 0.1053 | 0.0125 | 2.83E-17 | 1 |
| rs199515 | 17 | 44856641 | 17 | 43399058 | 44874453 | C | G | 0.2177 | WNT3 | intronic | 1.9060 | 0.1678 | 6.78E-30 | 2.2324 | 0.3249 | 7.00E-12 | 0.1049 | 0.0125 | 3.91E-17 | 1 |
| rs199514 | 17 | 44856881 | 17 | 43399058 | 44874453 | A | G | 0.2187 | WNT3 | intronic | 1.9126 | 0.1677 | 3.99E-30 | 2.2162 | 0.3249 | 9.91E-12 | 0.1052 | 0.0124 | 2.71E-17 | 1 |
| rs199513 | 17 | 44856932 | 17 | 43399058 | 44874453 | A | G | 0.2167 | WNT3 | intronic | -1.9148 | 0.1678 | 3.75E-30 | -2.2163 | 0.3249 | 9.89E-12 | -0.1050 | 0.0125 | 3.62E-17 | 1 |
| rs199512 | 17 | 44857352 | 17 | 43399058 | 44874453 | T | C | 0.2197 | WNT3 | intronic | -1.9163 | 0.1679 | 3.47E-30 | -2.2105 | 0.3250 | 1.12E-11 | -0.1052 | 0.0124 | 2.78E-17 | 1 |
| NA | 17 | 44857929 | 17 | 43399058 | 44874453 | A | C | 0.2197 | WNT3 | intronic | -2.0733 | 0.2367 | 1.94E-18 | -2.2053 | 0.3250 | 1.27E-11 | -0.1062 | 0.0126 | 3.19E-17 | 1 |
| rs199509 | 17 | 44858728 | 17 | 43399058 | 44874453 | A | G | 0.2197 | WNT3 | intronic | 1.9140 | 0.1678 | 4.02E-30 | 2.2229 | 0.3250 | 8.67E-12 | 0.1052 | 0.0124 | 2.78E-17 | 1 |
| rs199508 | 17 | 44858838 | 17 | 43399058 | 44874453 | C | G | 0.2565 | WNT3 | intronic | -1.6831 | 0.1624 | 3.58E-25 | -1.9043 | 0.3153 | 1.64E-09 | -0.0870 | 0.0119 | 3.05E-13 | 1 |
| rs199507 | 17 | 44858855 | 17 | 43399058 | 44874453 | A | G | 0.2197 | WNT3 | intronic | -1.9149 | 0.1679 | 3.90E-30 | -2.2231 | 0.3250 | 8.64E-12 | -0.1052 | 0.0124 | 2.78E-17 | 1 |
| rs199506 | 17 | 44859031 | 17 | 43399058 | 44874453 | A | G | 0.2197 | WNT3 | intronic | -1.9138 | 0.168 | 4.69E-30 | -2.2323 | 0.3250 | 7.13E-12 | -0.1052 | 0.0125 | 3.37E-17 | 1 |
| rs415430 | 17 | 44859144 | 17 | 43399058 | 44874453 | T | C | 0.2167 | WNT3 | intronic | 1.9158 | 0.168 | 4.11E-30 | 2.2239 | 0.3250 | 8.55E-12 | 0.1055 | 0.0125 | 2.83E-17 | 1 |
| rs430685 | 17 | 44859148 | 17 | 43399058 | 44874453 | T | C | 0.2167 | WNT3 | intronic | -1.9159 | 0.168 | 4.07E-30 | -2.2239 | 0.3250 | 8.54E-12 | -0.1055 | 0.0125 | 2.92E-17 | 1 |
| rs199505 | 17 | 44859410 | 17 | 43399058 | 44874453 | A | G | 0.2177 | WNT3 | intronic | -1.9149 | 0.1681 | 4.67E-30 | -2.2317 | 0.3252 | 7.47E-12 | -0.1056 | 0.0125 | 2.78E-17 | 1 |
| rs70602 | 17 | 44859715 | 17 | 43399058 | 44874453 | T | C | 0.2177 | WNT3 | intronic | -1.9146 | 0.1682 | 5.15E-30 | -2.2289 | 0.3254 | 8.07E-12 | -0.1056 | 0.0125 | 2.70E-17 | 1 |
| rs70600 | 17 | 44860021 | 17 | 43399058 | 44874453 | T | C | 0.2177 | WNT3 | intronic | -1.9256 | 0.1683 | 2.60E-30 | -2.2457 | 0.3262 | 6.35E-12 | -0.1057 | 0.0125 | 2.67E-17 | 1 |
| rs199504 | 17 | 44861003 | 17 | 43399058 | 44874453 | T | C | 0.2207 | WNT3 | intronic | 1.9200 | 0.1683 | 3.80E-30 | 2.2057 | 0.3251 | 1.28E-11 | 0.1055 | 0.0125 | 3.35E-17 | 1 |
| NA | 17 | 44862162 | 17 | 43399058 | 44874453 | A | G | 0.2197 | WNT3 | intronic | -2.0964 | 0.2437 | 7.78E-18 | -2.2480 | 0.3267 | 6.51E-12 | -0.1097 | 0.0128 | 9.97E-18 | 1 |
| rs199502 | 17 | 44862347 | 17 | 43399058 | 44874453 | A | G | 0.2326 | WNT3 | intronic | -1.8352 | 0.1686 | 1.40E-27 | -2.1448 | 0.3277 | 6.45E-11 | -0.0949 | 0.0124 | 1.70E-14 | 1 |
| rs199501 | 17 | 44862613 | 17 | 43399058 | 44874453 | A | G | 0.2525 | WNT3 | intronic | -1.7066 | 0.1635 | 1.72E-25 | -2.0012 | 0.3162 | 2.65E-10 | -0.0894 | 0.0121 | 1.27E-13 | 1 |
| rs916888 | 17 | 44863133 | 17 | 43399058 | 44874453 | T | C | 0.2654 | WNT3 | intronic | 1.6437 | 0.1673 | 8.71E-23 | 2.1307 | 0.3099 | 6.83E-12 | 0.1001 | 0.0124 | 6.70E-16 | 1 |
| rs199500 | 17 | 44863413 | 17 | 43399058 | 44874453 | T | C | 0.2734 | WNT3 | intronic | -1.5514 | 0.1684 | 3.22E-20 | -1.9120 | 0.3063 | 4.57E-10 | -0.1018 | 0.0127 | 1.37E-15 | 1 |
| rs2074404 | 17 | 44865439 | 17 | 43399058 | 44874453 | T | G | 0.2664 | WNT3 | intronic | 1.4180 | 0.1669 | 1.96E-17 | 1.8595 | 0.3267 | 1.31E-08 | 0.0843 | 0.0120 | 2.53E-12 | 1 |
| rs199499 | 17 | 44865498 | 17 | 43399058 | 44874453 | T | C | 0.2078 | WNT3 | intronic | -1.8702 | 0.182 | 8.85E-25 | -2.2314 | 0.3559 | 3.87E-10 | -0.1123 | 0.0135 | 7.47E-17 | 1 |
| rs199498 | 17 | 44865603 | 17 | 43399058 | 44874453 | T | C | 0.2247 | WNT3 | intronic | 1.7129 | 0.1774 | 4.63E-22 | 2.0385 | 0.3420 | 2.64E-09 | 0.0981 | 0.0130 | 4.01E-14 | 1 |
| rs199497 | 17 | 44866602 | 17 | 43399058 | 44874453 | T | C | 0.1809 | WNT3 | intronic | 1.3391 | 0.2139 | 3.82E-10 | 1.1787 | 0.3699 | 1.45E-03 | NA | NA | NA | NA |
| rs1563304 | 17 | 44874453 | 17 | 43399058 | 44874453 | T | C | 0.1829 | WNT3 | intronic | -1.5441 | 0.2783 | 2.90E-08 | -1.0230 | 0.3474 | 3.25E-03 | -0.0799 | 0.0191 | 2.78E-05 | 1 |
| rs35895680 | 17 | 47060322 | 18 | 47060322 | 47145848 | A | C | 0.2992 | RP11-501C14.5 | intergenic | 0.8719 | 0.1555 | 2.05E-08 | -0.1880 | 0.2875 | 5.13E-01 | 0.0457 | 0.0113 | 5.41E-05 | 0 |
| rs9909861 | 17 | 47079416 | 18 | 47060322 | 47145848 | A | C | 0.3191 | IGF2BP1 | intronic | 0.8659 | 0.1467 | 3.62E-09 | 0.0230 | 0.2812 | 9.35E-01 | 0.0458 | 0.0107 | 1.82E-05 | 0 |
| rs12945020 | 17 | 47082775 | 18 | 47060322 | 47145848 | A | G | 0.3181 | IGF2BP1:RP11-501( | ncRNA_intronic | 0.8547 | 0.1465 | 5.35E-09 | 0.0490 | 0.2810 | 8.62E-01 | 0.0435 | 0.0106 | 4.30E-05 | 0 |
| rs35051752 | 17 | 47089908 | 18 | 47060322 | 47145848 | A | G | 0.3181 | IGF2BP1:RP11-501( | ncRNA_intronic | 0.8605 | 0.1465 | 4.27E-09 | 0.0475 | 0.2809 | 8.66E-01 | 0.0430 | 0.0106 | 5.42E-05 | 0 |
| rs11079849 | 17 | 47090785 | 18 | 47060322 | 47145848 | T | C | 0.2903 | IGF2BP1:RP11-501( | ncRNA_intronic | 0.9559 | 0.1515 | 2.80E-10 | 0.2316 | 0.2877 | 4.21E-01 | 0.0443 | 0.0111 | 7.03E-05 | 0 |
| rs9906710 | 17 | 47091283 | 18 | 47060322 | 47145848 | A | C | 0.3221 | IGF2BP1 | intronic | 0.8553 | 0.1463 | 5.06E-09 | 0.0461 | 0.2803 | 8.69E-01 | 0.0425 | 0.0106 | 6.18E-05 | 0 |
| rs9906944 | 17 | 47091420 | 18 | 47060322 | 47145848 | T | C | 0.2992 | IGF2BP1 | intronic | 0.9051 | 0.1495 | 1.39E-09 | -0.0506 | 0.2833 | 8.58E-01 | 0.0421 | 0.0110 | 1.23E-04 | 0 |
| rs4794018 | 17 | 47093398 | 18 | 47060322 | 47145848 | T | C | 0.3191 | IGF2BP1 | intronic | -0.8486 | 0.1463 | 6.65E-09 | -0.0365 | 0.2801 | 8.96E-01 | -0.0425 | 0.0106 | 6.12E-05 | 0 |
| rs9902512 | 17 | 47094274 | 18 | 47060322 | 47145848 | C | G | 0.3191 | IGF2BP1 | intronic | -0.8421 | 0.1465 | 8.92E-09 | -0.0367 | 0.2801 | 8.96E-01 | -0.0425 | 0.0106 | 6.26E-05 | 0 |
| rs11079852 | 17 | 47095041 | 18 | 47060322 | 47145848 | A | G | 0.3201 | IGF2BP1 | intronic | 0.8455 | 0.1463 | 7.53E-09 | 0.0382 | 0.2801 | 8.92E-01 | 0.0424 | 0.0106 | 6.22E-05 | 0 |

* Based on genome-wide tests of inferred statistics computed from the roi-specific GWASs of cortical surface area[1] in 19,152 participants of ENIGMA study[2].

** Replicated (=1) if the effect size estimates from both 'replication' association tests have the same direction as that from the meta-GWAS and p-values < 0.05.

1 .Nieuwboer, H. A., Pool, R., Dolan, C. V., Boomsma, D. I. & Nivard, M. G. GWIS: Genome-wide inferred statistics for functions of multiple phenotypes. The American Journal of Human Genetics 99, 917–927 (2016).

2. Grasby, K. L. et al. The genetic architecture of the human cerebral cortex. bioRxiv 399402 (2018). doi:10.1101/399402

**Table E2B. GWAS meta-analysis results for surface area PC2 for SNPs with P-values< 5E-08.**

The table shows the genomic positions, alleles, P values, mapped genes, and the functions with respect to the mapped genes.

| | | | | | | | | | | | Meta-GWAS | | | UKBB | | | ENIGMA* | | | |
|---|---|---|---|---|---|---|---|---|---|---|---|---|---|---|---|---|---|---|---|---|
| rsID | chr | pos (hg19) | Genomic Locus | start | end | A1 | A2 | MAF | Nearest Genes | Function | Beta | SE | P-value | Beta | SE | P-value | Beta | SE | P-value | Replicated** |
| rs1934057 | 1 | 18962095 | 19 | 18956404 | 18992466 | T | C | 0.4811 | PAX7 | intronic | -0.0762 | 0.0137 | 2.78E-08 | -0.1065 | 0.0276 | 1.18E-04 | -0.0662 | 0.0103 | 1.49E-10 | 1 |
| rs9439716 | 1 | 18978263 | 19 | 18956404 | 18992466 | A | C | 0.4384 | PAX7 | intronic | 0.0722 | 0.0131 | 3.96E-08 | 0.0817 | 0.0274 | 2.88E-03 | 0.0595 | 0.0099 | 1.97E-09 | 1 |
| rs10917360 | 1 | 23539010 | 20 | 23473592 | 23543929 | A | G | 0.2495 | HTR1D | intergenic | 0.0829 | 0.0151 | 3.84E-08 | 0.1051 | 0.0312 | 7.62E-04 | 0.0355 | 0.0115 | 2.04E-03 | 1 |
| rs7519093 | 1 | 23541078 | 20 | 23473592 | 23543929 | A | T | 0.2495 | HTR1D | intergenic | 0.0836 | 0.0151 | 3.03E-08 | 0.1075 | 0.0312 | 5.81E-04 | 0.0351 | 0.0115 | 2.29E-03 | 1 |
| rs11801120 | 1 | 23543307 | 20 | 23473592 | 23543929 | T | C | 0.2495 | HTR1D | intergenic | 0.0834 | 0.0151 | 3.37E-08 | 0.1074 | 0.0312 | 5.87E-04 | 0.0354 | 0.0115 | 2.16E-03 | 1 |
| rs7414010 | 1 | 23543929 | 20 | 23473592 | 23543929 | T | C | 0.2495 | HTR1D | intergenic | -0.0832 | 0.0152 | 4.11E-08 | -0.1065 | 0.0312 | 6.49E-04 | -0.0346 | 0.0116 | 2.72E-03 | 1 |
| rs4659441 | 1 | 26752992 | 21 | 26752992 | 26902694 | T | C | 0.2157 | LIN28A | UTR3 | -0.0972 | 0.0163 | 2.43E-09 | -0.0369 | 0.0341 | 2.80E-01 | -0.0499 | 0.0122 | 4.27E-05 | 0 |
| rs9438623 | 1 | 26755919 | 21 | 26752992 | 26902694 | A | G | 0.1531 | LIN28A | UTR3 | -0.105 | 0.0181 | 6.21E-09 | -0.0571 | 0.0385 | 1.38E-01 | -0.0543 | 0.0137 | 7.85E-05 | 0 |
| rs3811462 | 1 | 26758602 | 21 | 26752992 | 26902694 | T | C | 0.2177 | DHDDS | upstream | -0.0953 | 0.0162 | 4.14E-09 | -0.0383 | 0.0341 | 2.61E-01 | -0.0481 | 0.0121 | 7.20E-05 | 0 |
| rs9438550 | 1 | 26765576 | 21 | 26752992 | 26902694 | T | C | 0.2157 | DHDDS | intronic | -0.0949 | 0.0162 | 5.07E-09 | -0.0391 | 0.0341 | 2.52E-01 | -0.0467 | 0.0121 | 1.15E-04 | 0 |
| rs3127011 | 1 | 26782680 | 21 | 26752992 | 26902694 | A | G | 0.1511 | DHDDS | intronic | -0.1034 | 0.0182 | 1.35E-08 | -0.0467 | 0.0390 | 2.31E-01 | -0.0494 | 0.0137 | 3.21E-04 | 0 |
| rs2445639 | 1 | 26785926 | 21 | 26752992 | 26902694 | A | G | 0.1501 | DHDDS | intronic | 0.103 | 0.0182 | 1.64E-08 | 0.0466 | 0.0390 | 2.32E-01 | 0.0487 | 0.0138 | 4.29E-04 | 0 |
| rs2445640 | 1 | 26788319 | 21 | 26752992 | 26902694 | A | G | 0.1501 | DHDDS | intronic | -0.1032 | 0.0182 | 1.53E-08 | -0.0468 | 0.0390 | 2.31E-01 | -0.0492 | 0.0138 | 3.50E-04 | 0 |
| rs2494391 | 1 | 26798478 | 21 | 26752992 | 26902694 | T | C | 0.2157 | HMGN2 | upstream:downst | -0.0955 | 0.0163 | 4.33E-09 | -0.0341 | 0.0341 | 3.18E-01 | -0.0449 | 0.0121 | 2.07E-04 | 0 |
| rs2494399 | 1 | 26804796 | 21 | 26752992 | 26902694 | C | G | 0.2147 | HMGN2 | intergenic | -0.0948 | 0.0163 | 5.74E-09 | -0.0326 | 0.0341 | 3.39E-01 | -0.0457 | 0.0121 | 1.65E-04 | 0 |
| rs2061234 | 1 | 26807312 | 21 | 26752992 | 26902694 | A | G | 0.1501 | HMGN2 | intergenic | 0.1012 | 0.0183 | 3.19E-08 | 0.0466 | 0.0390 | 2.32E-01 | 0.0452 | 0.0139 | 1.16E-03 | 0 |
| rs2445636 | 1 | 26808191 | 21 | 26752992 | 26902694 | A | G | 0.1491 | HMGN2 | intergenic | -0.1042 | 0.0183 | 1.16E-08 | -0.0466 | 0.0390 | 2.32E-01 | -0.0490 | 0.0138 | 4.01E-04 | 0 |
| rs7543995 | 1 | 26814811 | 21 | 26752992 | 26902694 | A | G | 0.1501 | HMGN2 | intergenic | -0.1044 | 0.0183 | 1.17E-08 | -0.0466 | 0.0390 | 2.32E-01 | -0.0496 | 0.0139 | 3.48E-04 | 0 |
| rs80117212 | 1 | 26841794 | 21 | 26752992 | 26902694 | T | C | 0.1531 | DPPA2P2 | intergenic | -0.1013 | 0.0183 | 3.07E-08 | -0.0456 | 0.0390 | 2.42E-01 | -0.0463 | 0.0138 | 7.62E-04 | 0 |
| rs11589953 | 1 | 26851886 | 21 | 26752992 | 26902694 | A | T | 0.2177 | RPS6KA1 | intergenic | 0.091 | 0.0163 | 2.28E-08 | 0.0289 | 0.0341 | 3.97E-01 | 0.0431 | 0.0121 | 3.64E-04 | 0 |
| rs11247963 | 1 | 26855401 | 21 | 26752992 | 26902694 | A | G | 0.2177 | RPS6KA1 | upstream | -0.0909 | 0.0163 | 2.27E-08 | -0.0286 | 0.0341 | 4.02E-01 | -0.0421 | 0.0121 | 4.87E-04 | 0 |
| rs2229714 | 1 | 26900708 | 21 | 26752992 | 26902694 | A | G | 0.1372 | RPS6KA1 | UTR3 | -0.1051 | 0.0189 | 2.75E-08 | -0.0272 | 0.0398 | 4.95E-01 | -0.0432 | 0.0140 | 2.03E-03 | 0 |
| rs12121702 | 1 | 26900805 | 21 | 26752992 | 26902694 | A | G | 0.1342 | RPS6KA1 | UTR3 | -0.1055 | 0.0189 | 2.43E-08 | -0.0265 | 0.0399 | 5.06E-01 | -0.0432 | 0.0140 | 2.09E-03 | 0 |
| rs351370 | 1 | 113054659 | 22 | 113054659 | 113252614 | T | C | 0.4006 | WNT2B | intronic | 0.0773 | 0.0138 | 1.94E-08 | 0.1068 | 0.0277 | 1.19E-04 | 0.0677 | 0.0106 | 1.44E-10 | 1 |
| rs351372 | 1 | 113059220 | 22 | 113054659 | 113252614 | A | T | 0.4702 | WNT2B | intronic | 0.0783 | 0.0133 | 3.45E-09 | 0.1052 | 0.0275 | 1.30E-04 | NA | NA | NA | NA |
| rs910697 | 1 | 113063125 | 22 | 113054659 | 113252614 | A | G | 0.4612 | WNT2B | exonic | 0.0833 | 0.0132 | 3.26E-10 | 0.1165 | 0.0275 | 2.32E-05 | 0.0700 | 0.0101 | 5.06E-12 | 1 |
| rs12728001 | 1 | 113076756 | 22 | 113054659 | 113252614 | T | C | 0.4821 | ST7L | intronic | 0.0804 | 0.0131 | 9.02E-10 | 0.1038 | 0.0274 | 1.52E-04 | 0.0629 | 0.0100 | 3.02E-10 | 1 |
| rs7512948 | 1 | 113081176 | 22 | 113054659 | 113252614 | A | G | 0.4761 | ST7L | intronic | -0.0783 | 0.0131 | 2.20E-09 | -0.0991 | 0.0273 | 2.85E-04 | -0.0597 | 0.0100 | 2.55E-09 | 1 |
| rs10776755 | 1 | 113082974 | 22 | 113054659 | 113252614 | C | G | 0.4751 | ST7L | intronic | -0.0793 | 0.0132 | 1.68E-09 | -0.0968 | 0.0273 | 4.01E-04 | -0.0601 | 0.0100 | 2.12E-09 | 1 |
| rs10745330 | 1 | 113083439 | 22 | 113054659 | 113252614 | T | C | 0.4811 | ST7L | UTR3 | -0.0799 | 0.013 | 8.87E-10 | -0.1038 | 0.0274 | 1.51E-04 | -0.0617 | 0.0099 | 5.39E-10 | 1 |
| rs4838959 | 1 | 113085721 | 22 | 113054659 | 113252614 | A | C | 0.4712 | ST7L | intronic | 0.0752 | 0.0131 | 1.03E-08 | 0.0836 | 0.0273 | 2.23E-03 | 0.0588 | 0.0100 | 4.07E-09 | 1 |
| rs7544663 | 1 | 113089006 | 22 | 113054659 | 113252614 | T | C | 0.4761 | ST7L | intronic | -0.0776 | 0.013 | 2.34E-09 | -0.0963 | 0.0274 | 4.40E-04 | -0.0602 | 0.0099 | 1.23E-09 | 1 |
| rs7513716 | 1 | 113089320 | 22 | 113054659 | 113252614 | A | T | 0.4761 | ST7L | intronic | 0.0775 | 0.013 | 2.41E-09 | 0.0962 | 0.0274 | 4.46E-04 | NA | NA | NA | NA |
| rs10857963 | 1 | 113091487 | 22 | 113054659 | 113252614 | T | C | 0.4761 | ST7L | intronic | -0.0776 | 0.013 | 2.33E-09 | -0.0962 | 0.0274 | 4.46E-04 | -0.0601 | 0.0099 | 1.31E-09 | 1 |
| rs6537743 | 1 | 113092527 | 22 | 113054659 | 113252614 | A | G | 0.4761 | ST7L | intronic | 0.0777 | 0.013 | 2.25E-09 | 0.0961 | 0.0274 | 4.55E-04 | 0.0600 | 0.0099 | 1.38E-09 | 1 |
| rs10776756 | 1 | 113098015 | 22 | 113054659 | 113252614 | A | G | 0.4761 | ST7L | intronic | 0.0818 | 0.0133 | 7.56E-10 | 0.0962 | 0.0274 | 4.45E-04 | 0.0603 | 0.0103 | 5.72E-09 | 1 |
| rs12126277 | 1 | 113106369 | 22 | 113054659 | 113252614 | A | G | 0.4761 | ST7L | intronic | 0.0779 | 0.013 | 2.11E-09 | 0.0970 | 0.0274 | 3.98E-04 | 0.0600 | 0.0099 | 1.45E-09 | 1 |
| rs507213 | 1 | 113109104 | 22 | 113054659 | 113252614 | A | C | 0.4801 | ST7L | intronic | -0.0798 | 0.0133 | 1.89E-09 | -0.0899 | 0.0273 | 9.94E-04 | -0.0552 | 0.0103 | 7.42E-08 | 1 |
| rs11102497 | 1 | 113109274 | 22 | 113054659 | 113252614 | T | C | 0.4761 | ST7L | intronic | 0.0787 | 0.013 | 1.54E-09 | 0.0959 | 0.0274 | 4.61E-04 | 0.0593 | 0.0100 | 2.55E-09 | 1 |
| rs1001494 | 1 | 113110619 | 22 | 113054659 | 113252614 | T | C | 0.4761 | ST7L | intronic | 0.0793 | 0.013 | 1.19E-09 | 0.0981 | 0.0274 | 3.43E-04 | 0.0594 | 0.0100 | 2.49E-09 | 1 |
| rs1032312 | 1 | 113112746 | 22 | 113054659 | 113252614 | A | G | 0.4761 | ST7L | intronic | -0.0781 | 0.013 | 1.92E-09 | -0.0959 | 0.0274 | 4.61E-04 | -0.0596 | 0.0099 | 1.79E-09 | 1 |
| rs10776757 | 1 | 113121673 | 22 | 113054659 | 113252614 | C | G | 0.4801 | ST7L | intronic | -0.0762 | 0.013 | 5.18E-09 | -0.0920 | 0.0273 | 7.47E-04 | NA | NA | NA | NA |
| rs7554345 | 1 | 113122956 | 22 | 113054659 | 113252614 | T | G | 0.4742 | ST7L | intronic | 0.0777 | 0.013 | 2.33E-09 | 0.0969 | 0.0274 | 4.02E-04 | 0.0597 | 0.0099 | 1.70E-09 | 1 |
| rs7554916 | 1 | 113123734 | 22 | 113054659 | 113252614 | A | C | 0.4732 | ST7L | intronic | 0.0778 | 0.013 | 2.28E-09 | 0.0969 | 0.0274 | 4.06E-04 | 0.0596 | 0.0099 | 1.83E-09 | 1 |
| rs663533 | 1 | 113128804 | 22 | 113054659 | 113252614 | C | G | 0.4722 | ST7L | intronic | -0.0742 | 0.0132 | 1.75E-08 | -0.0845 | 0.0273 | 1.99E-03 | -0.0586 | 0.0101 | 6.06E-09 | 1 |
| rs6669096 | 1 | 113132361 | 22 | 113054659 | 113252614 | T | C | 0.4751 | ST7L | intronic | 0.0777 | 0.013 | 2.32E-09 | 0.0959 | 0.0274 | 4.64E-04 | 0.0594 | 0.0099 | 1.99E-09 | 1 |
| rs6666579 | 1 | 113132393 | 22 | 113054659 | 113252614 | A | G | 0.4742 | ST7L | intronic | 0.0776 | 0.013 | 2.40E-09 | 0.0969 | 0.0274 | 4.05E-04 | 0.0594 | 0.0099 | 2.11E-09 | 1 |
| rs6682737 | 1 | 113136229 | 22 | 113054659 | 113252614 | T | C | 0.4751 | ST7L | intronic | 0.0784 | 0.013 | 1.79E-09 | 0.0959 | 0.0274 | 4.66E-04 | 0.0609 | 0.0099 | 8.99E-10 | 1 |
| rs4839254 | 1 | 113140926 | 22 | 113054659 | 113252614 | T | C | 0.4811 | ST7L | intronic | -0.0758 | 0.013 | 5.98E-09 | -0.0904 | 0.0273 | 9.26E-04 | -0.0582 | 0.0100 | 5.50E-09 | 1 |
| rs6691932 | 1 | 113144263 | 22 | 113054659 | 113252614 | T | C | 0.4851 | ST7L | intronic | 0.0766 | 0.013 | 3.98E-09 | 0.0963 | 0.0274 | 4.40E-04 | 0.0591 | 0.0099 | 2.82E-09 | 1 |
| rs11102512 | 1 | 113145394 | 22 | 113054659 | 113252614 | A | G | 0.4742 | ST7L | intronic | 0.0774 | 0.013 | 2.80E-09 | 0.0973 | 0.0274 | 3.85E-04 | 0.0594 | 0.0099 | 2.17E-09 | 1 |
| rs7522288 | 1 | 113151205 | 22 | 113054659 | 113252614 | T | C | 0.4583 | ST7L | intronic | 0.0746 | 0.0132 | 1.42E-08 | 0.0843 | 0.0274 | 2.06E-03 | 0.0584 | 0.0101 | 6.40E-09 | 1 |
| rs7546094 | 1 | 113154400 | 22 | 113054659 | 113252614 | T | C | 0.4871 | ST7L | intronic | -0.0767 | 0.013 | 3.74E-09 | -0.0970 | 0.0274 | 4.02E-04 | -0.0584 | 0.0100 | 4.44E-09 | 1 |
| rs1994604 | 1 | 113156426 | 22 | 113054659 | 113252614 | T | C | 0.4871 | ST7L | intronic | -0.0767 | 0.013 | 3.70E-09 | -0.0970 | 0.0274 | 4.04E-04 | -0.0584 | 0.0100 | 4.43E-09 | 1 |
| rs6700673 | 1 | 113157054 | 22 | 113054659 | 113252614 | T | C | 0.4871 | ST7L | intronic | -0.0766 | 0.013 | 3.88E-09 | -0.0969 | 0.0274 | 4.10E-04 | -0.0584 | 0.0100 | 4.41E-09 | 1 |
| rs9429486 | 1 | 113157372 | 22 | 113054659 | 113252614 | T | C | 0.4692 | ST7L | intronic | 0.0748 | 0.013 | 9.31E-09 | 0.0928 | 0.0273 | 6.72E-04 | 0.0574 | 0.0100 | 9.95E-09 | 1 |
| rs55675817 | 1 | 113166876 | 22 | 113054659 | 113252614 | A | C | 0.4583 | CAPZA1 | intronic | 0.0749 | 0.0132 | 1.25E-08 | 0.0817 | 0.0273 | 2.83E-03 | 0.0578 | 0.0101 | 9.48E-09 | 1 |
| rs7415820 | 1 | 113169449 | 22 | 113054659 | 113252614 | A | G | 0.4871 | CAPZA1 | intronic | 0.077 | 0.013 | 3.15E-09 | 0.0947 | 0.0274 | 5.52E-04 | 0.0577 | 0.0100 | 6.72E-09 | 1 |
| rs12046208 | 1 | 113171589 | 22 | 113054659 | 113252614 | T | C | 0.4682 | CAPZA1 | intronic | 0.0749 | 0.013 | 8.78E-09 | 0.0919 | 0.0273 | 7.71E-04 | 0.0570 | 0.0100 | 1.21E-08 | 1 |
| rs11584092 | 1 | 113176262 | 22 | 113054659 | 113252614 | A | T | 0.4682 | CAPZA1 | intronic | 0.0764 | 0.0131 | 5.78E-09 | 0.0897 | 0.0273 | 1.03E-03 | 0.0587 | 0.0101 | 5.31E-09 | 1 |
| rs72699100 | 1 | 113181401 | 22 | 113054659 | 113252614 | A | G | 0.4583 | CAPZA1 | intronic | -0.0754 | 0.0132 | 1.12E-08 | -0.0842 | 0.0274 | 2.08E-03 | -0.0582 | 0.0101 | 8.59E-09 | 1 |
| rs12042407 | 1 | 113183879 | 22 | 113054659 | 113252614 | A | T | 0.4583 | CAPZA1 | intronic | -0.0766 | 0.0132 | 6.17E-09 | -0.0832 | 0.0273 | 2.34E-03 | -0.0583 | 0.0101 | 8.45E-09 | 1 |
| rs3013440 | 1 | 113195789 | 22 | 113054659 | 113252614 | A | G | 0.4871 | CAPZA1 | intronic | 0.0768 | 0.013 | 3.73E-09 | 0.0948 | 0.0274 | 5.51E-04 | 0.0577 | 0.0100 | 7.99E-09 | 1 |
| rs3103450 | 1 | 113197730 | 22 | 113054659 | 113252614 | A | G | 0.4871 | CAPZA1 | intronic | 0.0769 | 0.013 | 3.47E-09 | 0.0945 | 0.0274 | 5.73E-04 | 0.0578 | 0.0100 | 7.21E-09 | 1 |
| rs7524494 | 1 | 113197800 | 22 | 113054659 | 113252614 | A | G | 0.4871 | CAPZA1 | intronic | -0.0768 | 0.013 | 3.58E-09 | -0.0945 | 0.0274 | 5.73E-04 | -0.0577 | 0.0100 | 7.96E-09 | 1 |
| rs3013439 | 1 | 113200671 | 22 | 113054659 | 113252614 | T | G | 0.4871 | CAPZA1 | intronic | -0.077 | 0.013 | 3.36E-09 | -0.0943 | 0.0274 | 5.82E-04 | -0.0576 | 0.0100 | 8.11E-09 | 1 |
| rs2932536 | 1 | 113202417 | 22 | 113054659 | 113252614 | A | G | 0.4871 | CAPZA1 | intronic | 0.077 | 0.013 | 3.26E-09 | 0.0944 | 0.0274 | 5.74E-04 | 0.0575 | 0.0100 | 8.99E-09 | 1 |
| rs12046329 | 1 | 113207531 | 22 | 113054659 | 113252614 | T | C | 0.4871 | CAPZA1 | intronic | -0.0781 | 0.013 | 2.13E-09 | -0.0951 | 0.0274 | 5.23E-04 | -0.0570 | 0.0100 | 1.22E-08 | 1 |
| rs12046466 | 1 | 113208039 | 22 | 113054659 | 113252614 | T | C | 0.4871 | CAPZA1 | intronic | -0.0785 | 0.013 | 1.64E-09 | -0.0952 | 0.0274 | 5.20E-04 | -0.0578 | 0.0100 | 7.37E-09 | 1 |
| rs6537749 | 1 | 113218503 | 22 | 113054659 | 113252614 | T | C | 0.4573 | MOV10 | intronic | -0.0762 | 0.0132 | 7.59E-09 | -0.0864 | 0.0274 | 1.60E-03 | -0.0577 | 0.0101 | 1.23E-08 | 1 |
| rs2999155 | 1 | 113221658 | 22 | 113054659 | 113252614 | A | G | 0.4881 | MOV10 | intronic | 0.0793 | 0.0131 | 1.22E-09 | 0.0958 | 0.0274 | 4.84E-04 | 0.0569 | 0.0100 | 1.34E-08 | 1 |
| rs2999154 | 1 | 113222456 | 22 | 113054659 | 113252614 | T | G | 0.4881 | MOV10 | intronic | 0.0786 | 0.013 | 1.68E-09 | 0.0972 | 0.0274 | 3.96E-04 | 0.0570 | 0.0100 | 1.33E-08 | 1 |
| rs2999157 | 1 | 113233193 | 22 | 113054659 | 113252614 | A | G | 0.4871 | MOV10 | intronic | 0.076 | 0.0131 | 7.49E-09 | 0.0980 | 0.0274 | 3.55E-04 | 0.0598 | 0.0102 | 3.83E-09 | 1 |
| rs2932532 | 1 | 113234456 | 22 | 113054659 | 113252614 | A | G | 0.4881 | MOV10 | intronic | 0.078 | 0.0131 | 2.54E-09 | 0.0976 | 0.0274 | 3.73E-04 | 0.0570 | 0.0101 | 1.51E-08 | 1 |
| rs902289 | 1 | 113236087 | 22 | 113054659 | 113252614 | T | C | 0.4881 | MOV10 | intronic | 0.0778 | 0.0131 | 2.79E-09 | 0.0973 | 0.0274 | 3.88E-04 | 0.0570 | 0.0101 | 1.50E-08 | 1 |
| rs1110043 | 1 | 113236421 | 22 | 113054659 | 113252614 | A | G | 0.4583 | MOV10:RP11-4 | ncRNA_exonic | 0.0746 | 0.0131 | 1.38E-08 | 0.0857 | 0.0273 | 1.73E-03 | 0.0559 | 0.0102 | 3.91E-08 | 1 |
| rs2999158 | 1 | 113239478 | 22 | 113054659 | 113252614 | T | C | 0.334 | MOV10 | intronic | -0.0931 | 0.0139 | 2.13E-11 | -0.1279 | 0.0291 | 1.09E-05 | -0.0759 | 0.0108 | 1.76E-12 | 1 |
| rs3013431 | 1 | 113242488 | 22 | 113054659 | 113252614 | T | C | 0.4016 | MOV10 | intronic | 0.0803 | 0.0134 | 2.02E-09 | 0.1008 | 0.0281 | 3.40E-04 | 0.0660 | 0.0103 | 1.48E-10 | 1 |
| rs937053 | 1 | 113246229 | 22 | 113054659 | 113252614 | A | T | 0.4543 | RHOC:RP11-42 | intronic | 0.0746 | 0.0132 | 1.50E-08 | 0.0861 | 0.0276 | 1.80E-03 | NA | NA | NA | NA |
| rs2999156 | 1 | 113247127 | 22 | 113054659 | 113252614 | C | G | 0.4543 | RHOC:RP11-42 | intronic | 0.0761 | 0.0132 | 8.08E-09 | 0.0872 | 0.0275 | 1.56E-03 | NA | NA | NA | NA |
| rs954679 | 1 | 113248365 | 22 | 113054659 | 113252614 | A | G | 0.3936 | RHOC:RP11-42 | UTR5 | -0.0838 | 0.0134 | 4.38E-10 | -0.1050 | 0.0283 | 2.10E-04 | -0.0614 | 0.0104 | 3.43E-09 | 1 |
| rs340496 | 2 | 45134871 | 23 | 45130410 | 45175585 | C | G | 0.4254 | RP11-89K21.1 | intergenic | 0.0777 | 0.0132 | 4.57E-09 | 0.0727 | 0.0275 | 8.23E-03 | 0.0227 | 0.0101 | 2.44E-02 | 1 |
| rs340497 | 2 | 45134900 | 23 | 45130410 | 45175585 | C | G | 0.4264 | RP11-89K21.1 | intergenic | 0.078 | 0.0132 | 3.88E-09 | 0.0709 | 0.0275 | 1.00E-02 | 0.0228 | 0.0101 | 2.36E-02 | 1 |
| rs340498 | 2 | 45135885 | 23 | 45130410 | 45175585 | A | C | 0.4334 | RP11-89K21.1 | intergenic | -0.0779 | 0.0133 | 4.48E-09 | -0.0703 | 0.0274 | 1.05E-02 | -0.0232 | 0.0101 | 2.15E-02 | 1 |
| rs564906 | 2 | 45136693 | 23 | 45130410 | 45175585 | C | G | 0.4264 | RP11-89K21.1 | intergenic | 0.0775 | 0.0133 | 5.48E-09 | 0.0687 | 0.0274 | 1.23E-02 | 0.0233 | 0.0101 | 2.12E-02 | 1 |
| rs564207 | 2 | 45136742 | 23 | 45130410 | 45175585 | T | C | 0.4225 | RP11-89K21.1 | intergenic | -0.0793 | 0.0133 | 2.41E-09 | -0.0694 | 0.0275 | 1.15E-02 | -0.0219 | 0.0102 | 3.09E-02 | 1 |
| rs540395 | 2 | 45138444 | 23 | 45130410 | 45175585 | A | C | 0.4165 | RP11-89K21.1 | intergenic | -0.0739 | 0.0134 | 3.80E-08 | -0.0593 | 0.0279 | 3.37E-02 | -0.0285 | 0.0102 | 5.25E-03 | 1 |
| rs540606 | 2 | 45138507 | 23 | 45130410 | 45175585 | A | G | 0.4254 | RP11-89K21.1 | intergenic | -0.0781 | 0.0133 | 4.01E-09 | -0.0708 | 0.0274 | 9.72E-03 | -0.0236 | 0.0101 | 1.97E-02 | 1 |
| rs567047 | 2 | 45139093 | 23 | 45130410 | 45175585 | A | G | 0.4225 | RP11-89K21.1 | intergenic | -0.0785 | 0.0133 | 3.42E-09 | -0.0696 | 0.0274 | 1.11E-02 | -0.0235 | 0.0101 | 2.03E-02 | 1 |

| | | | | | | | | | | | Meta-GWAS | | | UKBB | | | ENIGMA* | | | |
|---|---|---|---|---|---|---|---|---|---|---|---|---|---|---|---|---|---|---|---|---|
| rsID | chr | pos (hg19) | Genomic Locus | start | end | A1 | A2 | MAF | Nearest Genes | Function | Beta | SE | P-value | Beta | SE | P-value | Beta | SE | P-value | Replicated** |
| rs567897 | 2 | 45139153 | 23 | 45130410 | 45175585 | A | G | 0.4185 | RP11-89K21.1 | intergenic | -0.0733 | 0.0134 | 4.86E-08 | -0.0570 | 0.0279 | 4.12E-02 | -0.0274 | 0.0102 | 7.14E-03 | 1 |
| rs473098 | 2 | 45139779 | 23 | 45130410 | 45175585 | T | C | 0.4235 | RP11-89K21.1 | intergenic | 0.0782 | 0.0133 | 3.86E-09 | 0.0686 | 0.0273 | 1.21E-02 | 0.0234 | 0.0101 | 2.08E-02 | 1 |
| rs472140 | 2 | 45139904 | 23 | 45130410 | 45175585 | T | C | 0.4205 | RP11-89K21.1 | intergenic | 0.0792 | 0.0133 | 2.46E-09 | 0.0680 | 0.0274 | 1.30E-02 | 0.0235 | 0.0101 | 2.02E-02 | 1 |
| rs518853 | 2 | 45141218 | 23 | 45130410 | 45175585 | A | G | 0.4205 | RP11-89K21.1 | intergenic | 0.0799 | 0.0133 | 1.87E-09 | 0.0679 | 0.0274 | 1.33E-02 | 0.0236 | 0.0101 | 1.94E-02 | 1 |
| rs570436 | 2 | 45142673 | 23 | 45130410 | 45175585 | T | C | 0.4225 | RP11-89K21.1 | intergenic | 0.0784 | 0.0133 | 3.69E-09 | 0.0751 | 0.0275 | 6.31E-03 | 0.0243 | 0.0101 | 1.62E-02 | 1 |
| rs578584 | 2 | 45143175 | 23 | 45130410 | 45175585 | A | T | 0.4245 | RP11-89K21.1 | intergenic | -0.0772 | 0.0133 | 6.43E-09 | -0.0732 | 0.0275 | 7.83E-03 | -0.0234 | 0.0101 | 2.08E-02 | 1 |
| rs542883 | 2 | 45143382 | 23 | 45130410 | 45175585 | C | G | 0.4215 | RP11-89K21.1 | intergenic | 0.0789 | 0.0133 | 2.99E-09 | 0.0690 | 0.0275 | 1.22E-02 | 0.0248 | 0.0101 | 1.42E-02 | 1 |
| rs490340 | 2 | 45151616 | 23 | 45130410 | 45175585 | T | C | 0.4324 | RP11-89K21.1 | ncRNA_intronic | 0.081 | 0.0133 | 1.27E-09 | 0.0710 | 0.0275 | 9.73E-03 | 0.0256 | 0.0101 | 1.15E-02 | 1 |
| rs528301 | 2 | 45154908 | 23 | 45130410 | 45175585 | A | G | 0.4294 | RP11-89K21.1 | ncRNA_intronic | 0.0844 | 0.0133 | 2.30E-10 | 0.0672 | 0.0274 | 1.43E-02 | 0.0249 | 0.0101 | 1.40E-02 | 1 |
| rs1004787 | 2 | 45159091 | 23 | 45130410 | 45175585 | A | G | 0.4602 | RP11-89K21.1 | ncRNA_intronic | 0.0813 | 0.0132 | 7.56E-10 | 0.0607 | 0.0273 | 2.63E-02 | 0.0218 | 0.0100 | 2.99E-02 | 1 |
| rs895632 | 2 | 45165045 | 23 | 45130410 | 45175585 | A | C | 0.4324 | RP11-89K21.1 | ncRNA_intronic | -0.0811 | 0.0133 | 9.65E-10 | -0.0505 | 0.0274 | 6.50E-02 | -0.0281 | 0.0101 | 5.31E-03 | 0 |
| rs1561227 | 2 | 45165389 | 23 | 45130410 | 45175585 | T | C | 0.4602 | RP11-89K21.1 | ncRNA_intronic | 0.0805 | 0.0132 | 1.08E-09 | 0.0541 | 0.0272 | 4.70E-02 | 0.0265 | 0.0100 | 7.94E-03 | 1 |
| rs83995 | 2 | 45167570 | 23 | 45130410 | 45175585 | T | G | 0.4095 | SIX3-AS1 | ncRNA_exonic | 0.0851 | 0.0133 | 1.49E-10 | 0.0553 | 0.0277 | 4.63E-02 | 0.0285 | 0.0101 | 4.59E-03 | 1 |
| rs741813 | 2 | 45167886 | 23 | 45130410 | 45175585 | A | T | 0.4602 | SIX3-AS1 | ncRNA_exonic | 0.0814 | 0.0132 | 6.93E-10 | 0.0592 | 0.0273 | 2.99E-02 | 0.0269 | 0.0100 | 7.17E-03 | 1 |
| rs338070 | 2 | 45175289 | 23 | 45130410 | 45175585 | C | G | 0.4076 | SIX3 | intergenic | -0.0812 | 0.0133 | 1.12E-09 | -0.0587 | 0.0279 | 3.55E-02 | -0.0288 | 0.0102 | 4.86E-03 | 1 |
| rs173076 | 2 | 45175585 | 23 | 45130410 | 45175585 | A | G | 0.4105 | SIX3 | intergenic | 0.0811 | 0.0133 | 1.13E-09 | 0.0579 | 0.0279 | 3.81E-02 | 0.0295 | 0.0102 | 3.92E-03 | 1 |
| rs4895120 | 3 | 104647183 | 24 | 104646815 | 104828166 | T | C | 0.493 | RP11-281P11.1 | intergenic | -0.1009 | 0.0132 | 1.93E-14 | -0.0965 | 0.0272 | 3.86E-04 | -0.0689 | 0.0101 | 1.04E-11 | 1 |
| rs9826055 | 3 | 104647968 | 24 | 104646815 | 104828166 | A | T | 0.493 | RP11-281P11.1 | intergenic | -0.1009 | 0.0132 | 1.88E-14 | -0.0953 | 0.0272 | 4.54E-04 | NA | NA | NA | NA |
| rs2196494 | 3 | 104650052 | 24 | 104646815 | 104828166 | T | C | 0.4911 | RP11-281P11.1 | intergenic | -0.1015 | 0.0132 | 1.41E-14 | -0.0954 | 0.0271 | 4.40E-04 | -0.0688 | 0.0101 | 1.12E-11 | 1 |
| rs12695100 | 3 | 104650477 | 24 | 104646815 | 104828166 | T | C | 0.4911 | RP11-281P11.1 | intergenic | -0.1008 | 0.0131 | 1.64E-14 | -0.0954 | 0.0271 | 4.42E-04 | -0.0683 | 0.0101 | 1.29E-11 | 1 |
| rs6794834 | 3 | 104651280 | 24 | 104646815 | 104828166 | T | C | 0.4911 | RP11-281P11.1 | intergenic | 0.1009 | 0.0131 | 1.55E-14 | 0.0945 | 0.0271 | 5.00E-04 | 0.0683 | 0.0101 | 1.29E-11 | 1 |
| rs4264756 | 3 | 104651489 | 24 | 104646815 | 104828166 | C | G | 0.493 | RP11-281P11.1 | intergenic | -0.1008 | 0.0131 | 1.56E-14 | -0.0945 | 0.0271 | 5.02E-04 | NA | NA | NA | NA |
| rs1540462 | 3 | 104651985 | 24 | 104646815 | 104828166 | A | G | 0.493 | RP11-281P11.1 | intergenic | -0.1009 | 0.0131 | 1.48E-14 | -0.0956 | 0.0271 | 4.28E-04 | -0.0681 | 0.0101 | 1.39E-11 | 1 |
| rs1910326 | 3 | 104652528 | 24 | 104646815 | 104828166 | T | C | 0.492 | RP11-281P11.1 | intergenic | -0.1009 | 0.0131 | 1.42E-14 | -0.0953 | 0.0271 | 4.44E-04 | -0.0680 | 0.0101 | 1.41E-11 | 1 |
| rs4895122 | 3 | 104652961 | 24 | 104646815 | 104828166 | A | G | 0.1978 | RP11-281P11.1 | intergenic | 0.1008 | 0.0164 | 7.83E-10 | 0.0789 | 0.0345 | 2.21E-02 | 0.0618 | 0.0125 | 7.17E-07 | 1 |
| rs993463 | 3 | 104653545 | 24 | 104646815 | 104828166 | A | T | 0.492 | RP11-281P11.1 | intergenic | -0.1015 | 0.0131 | 9.65E-15 | -0.0953 | 0.0271 | 4.41E-04 | NA | NA | NA | NA |
| rs6780351 | 3 | 104654075 | 24 | 104646815 | 104828166 | A | C | 0.4911 | ALCAM | intergenic | -0.0999 | 0.0131 | 2.66E-14 | -0.0951 | 0.0271 | 4.54E-04 | -0.0682 | 0.0101 | 1.36E-11 | 1 |
| rs12490814 | 3 | 104654515 | 24 | 104646815 | 104828166 | T | C | 0.4891 | ALCAM | intergenic | -0.1012 | 0.0131 | 1.17E-14 | -0.0966 | 0.0271 | 3.71E-04 | -0.0674 | 0.0100 | 1.90E-11 | 1 |
| rs7649239 | 3 | 104654791 | 24 | 104646815 | 104828166 | A | G | 0.4911 | ALCAM | intergenic | 0.1007 | 0.0131 | 1.45E-14 | 0.0953 | 0.0271 | 4.41E-04 | 0.0669 | 0.0100 | 1.85E-11 | 1 |
| rs2043626 | 3 | 104655212 | 24 | 104646815 | 104828166 | T | C | 0.492 | ALCAM | intergenic | 0.1007 | 0.0131 | 1.45E-14 | 0.0954 | 0.0271 | 4.36E-04 | 0.0668 | 0.0099 | 1.78E-11 | 1 |
| rs6808509 | 3 | 104655582 | 24 | 104646815 | 104828166 | A | C | 0.4911 | ALCAM | intergenic | 0.0986 | 0.013 | 3.55E-14 | 0.0951 | 0.0271 | 4.57E-04 | 0.0665 | 0.0099 | 1.73E-11 | 1 |
| rs6773136 | 3 | 104655765 | 24 | 104646815 | 104828166 | T | C | 0.4911 | ALCAM | intergenic | -0.0986 | 0.013 | 3.56E-14 | -0.0950 | 0.0271 | 4.60E-04 | -0.0666 | 0.0099 | 1.68E-11 | 1 |
| rs6808909 | 3 | 104655818 | 24 | 104646815 | 104828166 | A | G | 0.4911 | ALCAM | intergenic | 0.0986 | 0.013 | 3.59E-14 | 0.0951 | 0.0271 | 4.56E-04 | 0.0666 | 0.0099 | 1.71E-11 | 1 |
| rs2118003 | 3 | 104656639 | 24 | 104646815 | 104828166 | A | C | 0.4911 | ALCAM | intergenic | 0.0988 | 0.013 | 3.26E-14 | 0.0952 | 0.0271 | 4.52E-04 | 0.0666 | 0.0099 | 1.63E-11 | 1 |
| rs2118004 | 3 | 104656715 | 24 | 104646815 | 104828166 | T | C | 0.4911 | ALCAM | intergenic | 0.0986 | 0.013 | 3.64E-14 | 0.0953 | 0.0271 | 4.40E-04 | 0.0666 | 0.0099 | 1.70E-11 | 1 |
| rs6766559 | 3 | 104656989 | 24 | 104646815 | 104828166 | C | G | 0.492 | ALCAM | intergenic | 0.0986 | 0.013 | 3.67E-14 | 0.0957 | 0.0271 | 4.20E-04 | NA | NA | NA | NA |
| rs6766724 | 3 | 104657313 | 24 | 104646815 | 104828166 | C | G | 0.4901 | ALCAM | intergenic | -0.0987 | 0.013 | 3.36E-14 | -0.0963 | 0.0271 | 3.87E-04 | NA | NA | NA | NA |
| rs11713472 | 3 | 104657409 | 24 | 104646815 | 104828166 | A | G | 0.4911 | ALCAM | intergenic | -0.098 | 0.013 | 5.54E-14 | -0.0953 | 0.0271 | 4.39E-04 | -0.0662 | 0.0099 | 2.73E-11 | 1 |
| rs6783013 | 3 | 104658328 | 24 | 104646815 | 104828166 | T | C | 0.2813 | ALCAM | intergenic | -0.0895 | 0.0146 | 9.71E-10 | -0.0501 | 0.0303 | 9.81E-02 | -0.0719 | 0.0111 | 9.20E-11 | 0 |
| rs76664757 | 3 | 104660317 | 24 | 104646815 | 104828166 | A | T | 0.1968 | ALCAM | intergenic | -0.1005 | 0.0164 | 8.14E-10 | -0.0794 | 0.0345 | 2.13E-02 | -0.0624 | 0.0124 | 4.90E-07 | 1 |
| rs13077489 | 3 | 104660608 | 24 | 104646815 | 104828166 | C | G | 0.3062 | ALCAM | intergenic | 0.0845 | 0.0155 | 4.58E-08 | 0.0968 | 0.0298 | 1.18E-03 | 0.0582 | 0.0119 | 1.06E-06 | 1 |
| rs9829580 | 3 | 104660892 | 24 | 104646815 | 104828166 | A | G | 0.4901 | ALCAM | intergenic | 0.0987 | 0.013 | 3.39E-14 | 0.0964 | 0.0271 | 3.78E-04 | 0.0666 | 0.0099 | 1.62E-11 | 1 |
| rs9867223 | 3 | 104661131 | 24 | 104646815 | 104828166 | A | T | 0.4901 | ALCAM | intergenic | -0.0987 | 0.013 | 3.35E-14 | -0.0964 | 0.0271 | 3.78E-04 | NA | NA | NA | NA |
| rs9868127 | 3 | 104661750 | 24 | 104646815 | 104828166 | A | G | 0.4901 | ALCAM | intergenic | -0.0989 | 0.013 | 2.93E-14 | -0.0964 | 0.0271 | 3.83E-04 | -0.0667 | 0.0099 | 1.50E-11 | 1 |
| rs13318870 | 3 | 104662433 | 24 | 104646815 | 104828166 | T | C | 0.4901 | ALCAM | intergenic | 0.0987 | 0.013 | 3.22E-14 | 0.0966 | 0.0271 | 3.72E-04 | 0.0666 | 0.0099 | 1.63E-11 | 1 |
| rs77652765 | 3 | 104663554 | 24 | 104646815 | 104828166 | A | G | 0.1978 | ALCAM | intergenic | 0.1002 | 0.0163 | 8.67E-10 | 0.0782 | 0.0345 | 2.33E-02 | 0.0631 | 0.0124 | 3.67E-07 | 1 |
| rs7618943 | 3 | 104664130 | 24 | 104646815 | 104828166 | A | C | 0.4901 | ALCAM | intergenic | 0.0985 | 0.013 | 3.76E-14 | 0.0938 | 0.0271 | 5.52E-04 | 0.0669 | 0.0099 | 1.39E-11 | 1 |
| rs1393740 | 3 | 104664955 | 24 | 104646815 | 104828166 | A | C | 0.4901 | ALCAM | intergenic | 0.0982 | 0.013 | 4.36E-14 | 0.0938 | 0.0271 | 5.46E-04 | 0.0674 | 0.0099 | 9.25E-12 | 1 |
| rs1393741 | 3 | 104664981 | 24 | 104646815 | 104828166 | A | G | 0.4911 | ALCAM | intergenic | 0.0986 | 0.013 | 3.50E-14 | 0.0963 | 0.0271 | 3.88E-04 | 0.0672 | 0.0099 | 1.10E-11 | 1 |
| rs6792637 | 3 | 104666491 | 24 | 104646815 | 104828166 | T | C | 0.4274 | ALCAM | intergenic | -0.0752 | 0.0132 | 1.28E-08 | -0.0528 | 0.0273 | 5.27E-02 | -0.0627 | 0.0100 | 4.48E-10 | 0 |
| rs6782467 | 3 | 104666925 | 24 | 104646815 | 104828166 | T | C | 0.493 | ALCAM | intergenic | 0.099 | 0.013 | 2.63E-14 | 0.0969 | 0.0271 | 3.58E-04 | 0.0667 | 0.0099 | 1.53E-11 | 1 |
| rs6782636 | 3 | 104666928 | 24 | 104646815 | 104828166 | C | G | 0.493 | ALCAM | intergenic | 0.0986 | 0.013 | 3.46E-14 | 0.0971 | 0.0271 | 3.42E-04 | NA | NA | NA | NA |
| rs6806605 | 3 | 104666970 | 24 | 104646815 | 104828166 | A | C | 0.4911 | ALCAM | intergenic | -0.0986 | 0.013 | 3.38E-14 | -0.0948 | 0.0271 | 4.71E-04 | -0.0667 | 0.0099 | 1.47E-11 | 1 |
| rs13066237 | 3 | 104666976 | 24 | 104646815 | 104828166 | T | C | 0.4245 | ALCAM | intergenic | 0.0951 | 0.0132 | 5.04E-13 | 0.1088 | 0.0274 | 7.11E-05 | 0.0512 | 0.0100 | 3.41E-07 | 1 |
| rs12492893 | 3 | 104667061 | 24 | 104646815 | 104828166 | T | G | 0.4274 | ALCAM | intergenic | -0.0752 | 0.0132 | 1.29E-08 | -0.0528 | 0.0273 | 5.25E-02 | -0.0628 | 0.0100 | 4.11E-10 | 0 |
| rs12374063 | 3 | 104667506 | 24 | 104646815 | 104828166 | A | C | 0.4891 | ALCAM | intergenic | 0.099 | 0.013 | 3.21E-14 | 0.0978 | 0.0271 | 3.12E-04 | 0.0669 | 0.0099 | 1.63E-11 | 1 |
| rs12374161 | 3 | 104667530 | 24 | 104646815 | 104828166 | A | T | 0.4891 | ALCAM | intergenic | -0.099 | 0.013 | 3.29E-14 | -0.0957 | 0.0271 | 4.16E-04 | NA | NA | NA | NA |
| rs6807218 | 3 | 104667576 | 24 | 104646815 | 104828166 | A | G | 0.1958 | ALCAM | intergenic | 0.1004 | 0.0164 | 8.48E-10 | 0.0784 | 0.0344 | 2.27E-02 | 0.0629 | 0.0124 | 4.03E-07 | 1 |
| rs12374067 | 3 | 104667829 | 24 | 104646815 | 104828166 | A | G | 0.4901 | ALCAM | intergenic | 0.1008 | 0.0131 | 1.20E-14 | 0.0972 | 0.0271 | 3.36E-04 | 0.0671 | 0.0099 | 1.40E-11 | 1 |
| rs13092127 | 3 | 104667944 | 24 | 104646815 | 104828166 | A | G | 0.4911 | ALCAM | intergenic | -0.1008 | 0.0131 | 1.18E-14 | -0.0972 | 0.0271 | 3.35E-04 | -0.0672 | 0.0099 | 1.35E-11 | 1 |
| rs35266846 | 3 | 104668010 | 24 | 104646815 | 104828166 | T | C | 0.4911 | ALCAM | intergenic | 0.102 | 0.0131 | 5.84E-15 | 0.0974 | 0.0271 | 3.30E-04 | 0.0670 | 0.0099 | 1.47E-11 | 1 |
| rs35569867 | 3 | 104668074 | 24 | 104646815 | 104828166 | T | G | 0.4911 | ALCAM | intergenic | -0.1008 | 0.0131 | 1.23E-14 | -0.0951 | 0.0271 | 4.54E-04 | -0.0671 | 0.0099 | 1.40E-11 | 1 |
| rs9839056 | 3 | 104668172 | 24 | 104646815 | 104828166 | A | G | 0.4911 | ALCAM | intergenic | 0.1008 | 0.0131 | 1.17E-14 | 0.0972 | 0.0271 | 3.35E-04 | 0.0672 | 0.0099 | 1.33E-11 | 1 |
| rs9876402 | 3 | 104668183 | 24 | 104646815 | 104828166 | A | G | 0.4911 | ALCAM | intergenic | -0.1009 | 0.0131 | 1.11E-14 | -0.0973 | 0.0271 | 3.34E-04 | -0.0672 | 0.0099 | 1.32E-11 | 1 |
| rs9839153 | 3 | 104668453 | 24 | 104646815 | 104828166 | C | G | 0.492 | ALCAM | intergenic | -0.1011 | 0.0131 | 1.07E-14 | -0.0973 | 0.0271 | 3.34E-04 | NA | NA | NA | NA |
| rs6789712 | 3 | 104669023 | 24 | 104646815 | 104828166 | T | G | 0.4264 | ALCAM | intergenic | 0.0753 | 0.0132 | 1.24E-08 | 0.0538 | 0.0272 | 4.86E-02 | 0.0633 | 0.0101 | 4.41E-10 | 1 |
| rs1436337 | 3 | 104669238 | 24 | 104646815 | 104828166 | T | C | 0.4911 | ALCAM | intergenic | 0.1013 | 0.0131 | 9.09E-15 | 0.0971 | 0.0271 | 3.44E-04 | 0.0674 | 0.0099 | 1.18E-11 | 1 |
| rs7646958 | 3 | 104669766 | 24 | 104646815 | 104828166 | T | C | 0.492 | ALCAM | intergenic | -0.1011 | 0.0131 | 1.00E-14 | -0.0971 | 0.0271 | 3.42E-04 | -0.0677 | 0.0099 | 9.56E-12 | 1 |
| rs10804436 | 3 | 104670412 | 24 | 104646815 | 104828166 | A | C | 0.4274 | ALCAM | intergenic | -0.0955 | 0.0132 | 3.99E-13 | -0.1091 | 0.0274 | 6.77E-05 | -0.0515 | 0.0101 | 3.85E-07 | 1 |
| rs1816547 | 3 | 104670606 | 24 | 104646815 | 104828166 | T | G | 0.4911 | ALCAM | intergenic | 0.1009 | 0.0131 | 1.09E-14 | 0.0973 | 0.0271 | 3.35E-04 | 0.0670 | 0.0099 | 1.60E-11 | 1 |
| rs6802252 | 3 | 104670860 | 24 | 104646815 | 104828166 | A | T | 0.4911 | ALCAM | intergenic | -0.1007 | 0.0131 | 1.27E-14 | -0.0948 | 0.0271 | 4.71E-04 | NA | NA | NA | NA |
| rs1393742 | 3 | 104672111 | 24 | 104646815 | 104828166 | A | G | 0.4911 | ALCAM | intergenic | -0.1008 | 0.0131 | 1.23E-14 | -0.0974 | 0.0271 | 3.31E-04 | -0.0670 | 0.0099 | 1.63E-11 | 1 |
| rs1393743 | 3 | 104672291 | 24 | 104646815 | 104828166 | A | C | 0.4911 | ALCAM | intergenic | -0.0999 | 0.0131 | 2.17E-14 | -0.0953 | 0.0271 | 4.43E-04 | -0.0668 | 0.0099 | 1.83E-11 | 1 |
| rs9860835 | 3 | 104672414 | 24 | 104646815 | 104828166 | T | G | 0.4911 | ALCAM | intergenic | 0.1006 | 0.0131 | 1.42E-14 | 0.0974 | 0.0271 | 3.34E-04 | 0.0670 | 0.0099 | 1.63E-11 | 1 |
| rs9823410 | 3 | 104672758 | 24 | 104646815 | 104828166 | A | T | 0.4911 | ALCAM | intergenic | -0.1007 | 0.0131 | 1.36E-14 | -0.0972 | 0.0271 | 3.43E-04 | NA | NA | NA | NA |
| rs76521431 | 3 | 104672841 | 24 | 104646815 | 104828166 | T | G | 0.1968 | ALCAM | intergenic | -0.1004 | 0.0163 | 8.11E-10 | -0.0791 | 0.0344 | 2.15E-02 | -0.0628 | 0.0124 | 3.93E-07 | 1 |
| rs1436336 | 3 | 104673566 | 24 | 104646815 | 104828166 | A | T | 0.1968 | ALCAM | intergenic | -0.1001 | 0.0163 | 9.02E-10 | -0.0785 | 0.0344 | 2.25E-02 | -0.0632 | 0.0124 | 3.34E-07 | 1 |
| rs6788676 | 3 | 104674040 | 24 | 104646815 | 104828166 | A | G | 0.3638 | ALCAM | intergenic | 0.1062 | 0.0136 | 5.31E-15 | 0.0884 | 0.0281 | 1.65E-03 | 0.0679 | 0.0103 | 5.26E-11 | 1 |
| rs112113804 | 3 | 104674454 | 24 | 104646815 | 104828166 | A | G | 0.1968 | ALCAM | intergenic | -0.1011 | 0.0164 | 7.17E-10 | -0.0794 | 0.0344 | 2.10E-02 | -0.0631 | 0.0125 | 4.02E-07 | 1 |
| rs79003825 | 3 | 104676223 | 24 | 104646815 | 104828166 | A | C | 0.2137 | ALCAM | intergenic | -0.1105 | 0.0158 | 2.73E-12 | -0.0573 | 0.0333 | 8.54E-02 | -0.0729 | 0.0120 | 1.25E-09 | 0 |
| rs7618202 | 3 | 104676606 | 24 | 104646815 | 104828166 | A | G | 0.4911 | ALCAM | intergenic | -0.1006 | 0.0131 | 1.40E-14 | -0.0979 | 0.0272 | 3.12E-04 | -0.0661 | 0.0100 | 4.00E-11 | 1 |
| rs2062774 | 3 | 104677139 | 24 | 104646815 | 104828166 | T | C | 0.2137 | ALCAM | intergenic | 0.1103 | 0.0158 | 2.90E-12 | 0.0573 | 0.0333 | 8.56E-02 | 0.0726 | 0.0120 | 1.47E-09 | 0 |
| rs9863836 | 3 | 104678133 | 24 | 104646815 | 104828166 | T | C | 0.2137 | ALCAM | intergenic | -0.1111 | 0.0159 | 2.61E-12 | -0.0573 | 0.0336 | 8.84E-02 | -0.0729 | 0.0120 | 1.28E-09 | 0 |
| rs9863841 | 3 | 104678135 | 24 | 104646815 | 104828166 | T | C | 0.2137 | ALCAM | intergenic | -0.111 | 0.0159 | 2.82E-12 | -0.0570 | 0.0336 | 9.00E-02 | -0.0731 | 0.0120 | 1.24E-09 | 0 |
| rs12639074 | 3 | 104681496 | 24 | 104646815 | 104828166 | T | C | 0.4543 | ALCAM | intergenic | 0.0812 | 0.0133 | 9.77E-10 | 0.0962 | 0.0274 | 4.37E-04 | 0.0514 | 0.0102 | 4.97E-07 | 1 |
| rs74546871 | 3 | 104682814 | 24 | 104646815 | 104828166 | T | C | 0.2137 | ALCAM | intergenic | 0.1104 | 0.0158 | 2.64E-12 | 0.0600 | 0.0333 | 7.14E-02 | 0.0737 | 0.0120 | 8.03E-10 | 0 |
| rs11915534 | 3 | 104682872 | 24 | 104646815 | 104828166 | A | C | 0.3648 | ALCAM | intergenic | -0.0835 | 0.0137 | 9.64E-10 | -0.0547 | 0.0282 | 5.25E-02 | -0.0601 | 0.0105 | 1.07E-08 | 0 |
| rs9288795 | 3 | 104683753 | 24 | 104646815 | 104828166 | T | C | 0.4443 | ALCAM | intergenic | -0.1044 | 0.0131 | 1.72E-15 | -0.0923 | 0.0273 | 7.27E-04 | -0.0606 | 0.0101 | 1.76E-09 | 1 |
| rs7625683 | 3 | 104687585 | 24 | 104646815 | 104828166 | T | C | 0.2137 | ALCAM | intergenic | 0.1103 | 0.0158 | 2.83E-12 | 0.0597 | 0.0333 | 7.27E-02 | 0.0741 | 0.0120 | 6.40E-10 | 0 |
| rs13086204 | 3 | 104687952 | 24 | 104646815 | 104828166 | A | T | 0.4911 | ALCAM | intergenic | 0.0999 | 0.0131 | 2.49E-14 | 0.0971 | 0.0272 | 3.59E-04 | NA | NA | NA | NA |
| rs9850952 | 3 | 104688036 | 24 | 104646815 | 104828166 | A | G | 0.2137 | ALCAM | intergenic | 0.1103 | 0.0158 | 2.86E-12 | 0.0598 | 0.0333 | 7.26E-02 | 0.0744 | 0.0120 | 5.42E-10 | 0 |

| | | | | | | | | | | | Meta-GWAS | | | UKBB | | | ENIGMA* | | | |
|---|---|---|---|---|---|---|---|---|---|---|---|---|---|---|---|---|---|---|---|---|
| rsID | chr | pos (hg19) | Genomic Locus | start | end | A1 | A2 | MAF | Nearest Genes | Function | Beta | SE | P-value | Beta | SE | P-value | Beta | SE | P-value | Replicated** |
| rs4894893 | 3 | 104691743 | 24 | 104646815 | 104828166 | T | C | 0.2137 | ALCAM | intergenic | -0.1103 | 0.0158 | 2.69E-12 | -0.0596 | 0.0333 | 7.32E-02 | -0.0744 | 0.0120 | 5.56E-10 | 0 |
| rs4894894 | 3 | 104691806 | 24 | 104646815 | 104828166 | T | C | 0.4443 | ALCAM | intergenic | 0.1039 | 0.0131 | 2.33E-15 | 0.0924 | 0.0273 | 7.19E-04 | 0.0607 | 0.0101 | 1.68E-09 | 1 |
| rs34044746 | 3 | 104692695 | 24 | 104646815 | 104828166 | T | C | 0.4443 | ALCAM | intergenic | -0.1037 | 0.0131 | 2.56E-15 | -0.0927 | 0.0273 | 6.96E-04 | -0.0609 | 0.0101 | 1.52E-09 | 1 |
| rs13319705 | 3 | 104693252 | 24 | 104646815 | 104828166 | T | C | 0.2942 | ALCAM | intergenic | -0.1186 | 0.0141 | 4.70E-17 | -0.0986 | 0.0297 | 8.89E-04 | -0.0655 | 0.0108 | 1.33E-09 | 1 |
| rs9846639 | 3 | 104694609 | 24 | 104646815 | 104828166 | A | G | 0.2137 | ALCAM | intergenic | -0.1099 | 0.0158 | 3.31E-12 | -0.0597 | 0.0333 | 7.29E-02 | -0.0743 | 0.0120 | 5.86E-10 | 0 |
| rs13320037 | 3 | 104694900 | 24 | 104646815 | 104828166 | T | G | 0.2942 | ALCAM | intergenic | 0.1186 | 0.0141 | 4.55E-17 | 0.0987 | 0.0297 | 8.82E-04 | 0.0654 | 0.0108 | 1.38E-09 | 1 |
| rs9827968 | 3 | 104695956 | 24 | 104646815 | 104828166 | A | G | 0.4911 | ALCAM | intergenic | 0.0998 | 0.0131 | 2.79E-14 | 0.0971 | 0.0272 | 3.60E-04 | 0.0675 | 0.0100 | 1.77E-11 | 1 |
| rs79588215 | 3 | 104697042 | 24 | 104646815 | 104828166 | T | C | 0.2137 | ALCAM | intergenic | 0.1104 | 0.0158 | 2.70E-12 | 0.0604 | 0.0333 | 6.94E-02 | 0.0742 | 0.0120 | 5.86E-10 | 0 |
| rs1354281 | 3 | 104697513 | 24 | 104646815 | 104828166 | A | G | 0.4443 | ALCAM | intergenic | -0.1035 | 0.0132 | 3.88E-15 | -0.0926 | 0.0273 | 6.98E-04 | -0.0599 | 0.0101 | 3.24E-09 | 1 |
| rs9854793 | 3 | 104701066 | 24 | 104646815 | 104828166 | A | G | 0.2137 | ALCAM | intergenic | -0.1101 | 0.0158 | 3.00E-12 | -0.0605 | 0.0333 | 6.90E-02 | -0.0741 | 0.0120 | 6.08E-10 | 0 |
| rs9880189 | 3 | 104702294 | 24 | 104646815 | 104828166 | T | C | 0.4443 | ALCAM | intergenic | 0.1037 | 0.0131 | 2.62E-15 | 0.0928 | 0.0273 | 6.87E-04 | 0.0601 | 0.0100 | 1.89E-09 | 1 |
| rs28374795 | 3 | 104702921 | 24 | 104646815 | 104828166 | A | G | 0.2137 | ALCAM | intergenic | 0.112 | 0.0158 | 1.58E-12 | 0.0608 | 0.0333 | 6.77E-02 | 0.0730 | 0.0120 | 1.15E-09 | 0 |
| rs6771251 | 3 | 104704039 | 24 | 104646815 | 104828166 | T | C | 0.4443 | ALCAM | intergenic | -0.1039 | 0.0131 | 2.34E-15 | -0.0937 | 0.0273 | 6.08E-04 | -0.0601 | 0.0100 | 1.87E-09 | 1 |
| rs9846552 | 3 | 104705960 | 24 | 104646815 | 104828166 | T | C | 0.4443 | ALCAM | intergenic | 0.1038 | 0.0131 | 2.35E-15 | 0.0936 | 0.0273 | 6.12E-04 | 0.0600 | 0.0100 | 2.03E-09 | 1 |
| rs1818149 | 3 | 104711110 | 24 | 104646815 | 104828166 | T | C | 0.4423 | ALCAM | intergenic | 0.1026 | 0.0131 | 3.81E-15 | 0.0918 | 0.0273 | 7.82E-04 | 0.0594 | 0.0101 | 3.43E-09 | 1 |
| rs1354280 | 3 | 104711433 | 24 | 104646815 | 104828166 | A | G | 0.4414 | ALCAM | intergenic | 0.1017 | 0.0131 | 6.88E-15 | 0.0908 | 0.0273 | 8.88E-04 | 0.0583 | 0.0101 | 8.13E-09 | 1 |
| rs9854000 | 3 | 104711549 | 24 | 104646815 | 104828166 | T | C | 0.4423 | ALCAM | intergenic | -0.1022 | 0.0131 | 4.95E-15 | -0.0921 | 0.0273 | 7.45E-04 | -0.0594 | 0.0101 | 3.69E-09 | 1 |
| rs9808934 | 3 | 104713251 | 24 | 104646815 | 104828166 | T | G | 0.2157 | ALCAM | intergenic | 0.1098 | 0.0157 | 2.60E-12 | 0.0611 | 0.0332 | 6.63E-02 | 0.0738 | 0.0119 | 6.30E-10 | 0 |
| rs957485 | 3 | 104713687 | 24 | 104646815 | 104828166 | T | C | 0.2137 | ALCAM | intergenic | -0.1095 | 0.0157 | 3.01E-12 | -0.0608 | 0.0332 | 6.75E-02 | -0.0732 | 0.0119 | 7.94E-10 | 0 |
| rs12495603 | 3 | 104713881 | 24 | 104646815 | 104828166 | A | G | 0.492 | ALCAM | intergenic | 0.0995 | 0.0131 | 3.41E-14 | 0.0975 | 0.0272 | 3.42E-04 | 0.0663 | 0.0100 | 3.48E-11 | 1 |
| rs4894822 | 3 | 104714106 | 24 | 104646815 | 104828166 | A | C | 0.4443 | ALCAM | intergenic | -0.0999 | 0.0131 | 2.34E-14 | -0.0921 | 0.0273 | 7.44E-04 | -0.0540 | 0.0102 | 1.08E-07 | 1 |
| rs9869633 | 3 | 104714210 | 24 | 104646815 | 104828166 | A | G | 0.2167 | ALCAM | intergenic | -0.11 | 0.0157 | 2.41E-12 | -0.0611 | 0.0332 | 6.62E-02 | -0.0739 | 0.0119 | 5.63E-10 | 0 |
| rs13327829 | 3 | 104714898 | 24 | 104646815 | 104828166 | A | G | 0.2157 | ALCAM | intergenic | 0.11 | 0.0157 | 2.34E-12 | 0.0611 | 0.0332 | 6.62E-02 | 0.0745 | 0.0119 | 3.95E-10 | 0 |
| rs13324580 | 3 | 104715038 | 24 | 104646815 | 104828166 | T | C | 0.2127 | ALCAM | intergenic | 0.11 | 0.0157 | 2.46E-12 | 0.0601 | 0.0333 | 7.09E-02 | 0.0740 | 0.0119 | 5.44E-10 | 0 |
| rs6769796 | 3 | 104715856 | 24 | 104646815 | 104828166 | A | G | 0.4443 | ALCAM | intergenic | 0.1031 | 0.013 | 2.72E-15 | 0.0935 | 0.0273 | 6.24E-04 | 0.0593 | 0.0101 | 4.03E-09 | 1 |
| rs6769805 | 3 | 104715884 | 24 | 104646815 | 104828166 | A | C | 0.2157 | ALCAM | intergenic | 0.1098 | 0.0157 | 2.63E-12 | 0.0611 | 0.0332 | 6.62E-02 | 0.0748 | 0.0119 | 3.58E-10 | 0 |
| rs59016227 | 3 | 104716337 | 24 | 104646815 | 104828166 | T | G | 0.2157 | ALCAM | intergenic | -0.109 | 0.0157 | 4.29E-12 | -0.0611 | 0.0332 | 6.62E-02 | -0.0745 | 0.0120 | 5.69E-10 | 0 |
| rs58911379 | 3 | 104716969 | 24 | 104646815 | 104828166 | A | T | 0.2167 | ALCAM | intergenic | -0.1096 | 0.0157 | 2.87E-12 | -0.0630 | 0.0332 | 5.79E-02 | -0.0747 | 0.0119 | 3.52E-10 | 0 |
| rs13327531 | 3 | 104718011 | 24 | 104646815 | 104828166 | T | C | 0.4443 | ALCAM | intergenic | 0.1032 | 0.013 | 2.59E-15 | 0.0937 | 0.0273 | 6.11E-04 | 0.0590 | 0.0101 | 4.69E-09 | 1 |
| rs6802022 | 3 | 104718523 | 24 | 104646815 | 104828166 | A | G | 0.2157 | ALCAM | intergenic | -0.11 | 0.0157 | 2.38E-12 | -0.0611 | 0.0332 | 6.63E-02 | -0.0753 | 0.0119 | 2.38E-10 | 0 |
| rs10222610 | 3 | 104719661 | 24 | 104646815 | 104828166 | A | G | 0.2157 | ALCAM | intergenic | -0.11 | 0.0157 | 2.37E-12 | -0.0608 | 0.0332 | 6.76E-02 | -0.0754 | 0.0119 | 2.34E-10 | 0 |
| rs10222621 | 3 | 104719857 | 24 | 104646815 | 104828166 | T | C | 0.2157 | ALCAM | intergenic | -0.11 | 0.0157 | 2.38E-12 | -0.0611 | 0.0332 | 6.62E-02 | -0.0754 | 0.0119 | 2.36E-10 | 0 |
| rs10222454 | 3 | 104719975 | 24 | 104646815 | 104828166 | A | G | 0.2167 | ALCAM | intergenic | 0.1103 | 0.0157 | 2.08E-12 | 0.0631 | 0.0332 | 5.77E-02 | 0.0757 | 0.0119 | 2.01E-10 | 0 |
| rs10222457 | 3 | 104720112 | 24 | 104646815 | 104828166 | A | C | 0.4443 | ALCAM | intergenic | 0.1024 | 0.013 | 4.26E-15 | 0.0934 | 0.0273 | 6.34E-04 | 0.0589 | 0.0101 | 5.04E-09 | 1 |
| rs10222624 | 3 | 104720126 | 24 | 104646815 | 104828166 | A | G | 0.1889 | ALCAM | intergenic | -0.1064 | 0.0169 | 3.28E-10 | -0.0482 | 0.0357 | 1.77E-01 | -0.0769 | 0.0128 | 2.13E-09 | 0 |
| rs72989330 | 3 | 104721389 | 24 | 104646815 | 104828166 | T | G | 0.2157 | ALCAM | intergenic | -0.11 | 0.0157 | 2.38E-12 | -0.0610 | 0.0333 | 6.68E-02 | -0.0754 | 0.0119 | 2.43E-10 | 0 |
| rs72989332 | 3 | 104721814 | 24 | 104646815 | 104828166 | T | C | 0.2157 | ALCAM | intergenic | -0.11 | 0.0157 | 2.36E-12 | -0.0610 | 0.0333 | 6.68E-02 | -0.0754 | 0.0119 | 2.45E-10 | 0 |
| rs11929686 | 3 | 104722114 | 24 | 104646815 | 104828166 | A | G | 0.4443 | ALCAM | intergenic | -0.1024 | 0.013 | 4.04E-15 | -0.0926 | 0.0273 | 7.02E-04 | -0.0587 | 0.0100 | 4.34E-09 | 1 |
| rs6772884 | 3 | 104722492 | 24 | 104646815 | 104828166 | A | C | 0.2157 | ALCAM | intergenic | 0.111 | 0.0157 | 1.79E-12 | 0.0610 | 0.0333 | 6.67E-02 | 0.0749 | 0.0120 | 4.27E-10 | 0 |
| rs6797921 | 3 | 104722799 | 24 | 104646815 | 104828166 | A | G | 0.2157 | ALCAM | intergenic | -0.11 | 0.0157 | 2.46E-12 | -0.0607 | 0.0333 | 6.83E-02 | -0.0751 | 0.0119 | 2.71E-10 | 0 |
| rs1393744 | 3 | 104722954 | 24 | 104646815 | 104828166 | T | C | 0.2157 | ALCAM | intergenic | 0.1101 | 0.0157 | 2.38E-12 | 0.0610 | 0.0333 | 6.69E-02 | 0.0744 | 0.0119 | 3.42E-10 | 0 |
| rs72989334 | 3 | 104723673 | 24 | 104646815 | 104828166 | T | G | 0.2197 | ALCAM | intergenic | -0.1118 | 0.0156 | 8.36E-13 | -0.0603 | 0.0331 | 6.81E-02 | -0.0767 | 0.0118 | 8.45E-11 | 0 |
| rs7647615 | 3 | 104723971 | 24 | 104646815 | 104828166 | C | G | 0.492 | ALCAM | intergenic | 0.0973 | 0.0134 | 4.20E-13 | 0.0978 | 0.0273 | 3.33E-04 | NA | NA | NA | NA |
| rs9839209 | 3 | 104724315 | 24 | 104646815 | 104828166 | T | G | 0.2157 | ALCAM | intergenic | 0.1102 | 0.0157 | 2.23E-12 | 0.0610 | 0.0333 | 6.68E-02 | 0.0744 | 0.0119 | 3.48E-10 | 0 |
| rs971551 | 3 | 104724634 | 24 | 104646815 | 104828166 | T | C | 0.2157 | ALCAM | intergenic | -0.1102 | 0.0157 | 2.26E-12 | -0.0623 | 0.0333 | 6.13E-02 | -0.0743 | 0.0119 | 3.64E-10 | 0 |
| rs971550 | 3 | 104724787 | 24 | 104646815 | 104828166 | A | T | 0.2972 | ALCAM | intergenic | 0.1192 | 0.0141 | 2.54E-17 | 0.1001 | 0.0297 | 7.50E-04 | 0.0675 | 0.0107 | 2.57E-10 | 1 |
| rs4894896 | 3 | 104726660 | 24 | 104646815 | 104828166 | A | G | 0.2117 | ALCAM | intergenic | 0.1098 | 0.0157 | 2.98E-12 | 0.0593 | 0.0333 | 7.48E-02 | 0.0735 | 0.0119 | 5.92E-10 | 0 |
| rs4894897 | 3 | 104726699 | 24 | 104646815 | 104828166 | C | G | 0.1899 | ALCAM | intergenic | -0.1071 | 0.017 | 2.69E-10 | -0.0504 | 0.0357 | 1.58E-01 | -0.0785 | 0.0128 | 8.13E-10 | 0 |
| rs72989338 | 3 | 104727420 | 24 | 104646815 | 104828166 | T | C | 0.1879 | ALCAM | intergenic | -0.1058 | 0.017 | 4.55E-10 | -0.0488 | 0.0357 | 1.72E-01 | -0.0780 | 0.0128 | 1.08E-09 | 0 |
| rs6766074 | 3 | 104727505 | 24 | 104646815 | 104828166 | C | G | 0.1879 | ALCAM | intergenic | -0.1056 | 0.017 | 4.82E-10 | -0.0504 | 0.0357 | 1.58E-01 | -0.0780 | 0.0128 | 1.05E-09 | 0 |
| rs1604640 | 3 | 104728491 | 24 | 104646815 | 104828166 | C | G | 0.1879 | ALCAM | intergenic | 0.1055 | 0.017 | 5.06E-10 | 0.0490 | 0.0357 | 1.70E-01 | 0.0779 | 0.0128 | 1.15E-09 | 0 |
| rs6769723 | 3 | 104728781 | 24 | 104646815 | 104828166 | A | G | 0.1879 | ALCAM | intergenic | -0.1053 | 0.017 | 5.37E-10 | -0.0504 | 0.0357 | 1.58E-01 | -0.0780 | 0.0128 | 1.09E-09 | 0 |
| rs6796574 | 3 | 104728944 | 24 | 104646815 | 104828166 | A | G | 0.1879 | ALCAM | intergenic | 0.1053 | 0.017 | 5.36E-10 | 0.0506 | 0.0357 | 1.56E-01 | 0.0780 | 0.0128 | 1.10E-09 | 0 |
| rs6785288 | 3 | 104729083 | 24 | 104646815 | 104828166 | T | C | 0.1859 | ALCAM | intergenic | 0.1055 | 0.017 | 5.11E-10 | 0.0482 | 0.0357 | 1.78E-01 | 0.0775 | 0.0128 | 1.36E-09 | 0 |
| rs6772559 | 3 | 104729180 | 24 | 104646815 | 104828166 | A | G | 0.4175 | ALCAM | intergenic | -0.0927 | 0.0132 | 2.00E-12 | -0.0824 | 0.0276 | 2.82E-03 | -0.0555 | 0.0101 | 3.83E-08 | 1 |
| rs17386778 | 3 | 104729589 | 24 | 104646815 | 104828166 | A | T | 0.4175 | ALCAM | intergenic | -0.0926 | 0.0132 | 2.14E-12 | -0.0827 | 0.0276 | 2.72E-03 | -0.0554 | 0.0101 | 3.98E-08 | 1 |
| rs9682035 | 3 | 104730056 | 24 | 104646815 | 104828166 | T | G | 0.4175 | ALCAM | intergenic | 0.0925 | 0.0132 | 2.30E-12 | 0.0827 | 0.0276 | 2.72E-03 | 0.0556 | 0.0101 | 3.56E-08 | 1 |
| rs9682661 | 3 | 104730312 | 24 | 104646815 | 104828166 | A | G | 0.4175 | ALCAM | intergenic | 0.0921 | 0.0132 | 3.01E-12 | 0.0827 | 0.0276 | 2.74E-03 | 0.0544 | 0.0101 | 7.88E-08 | 1 |
| rs9288796 | 3 | 104730581 | 24 | 104646815 | 104828166 | A | G | 0.1879 | ALCAM | intergenic | -0.1055 | 0.017 | 5.00E-10 | -0.0489 | 0.0357 | 1.71E-01 | -0.0763 | 0.0129 | 2.95E-09 | 0 |
| rs4894898 | 3 | 104732673 | 24 | 104646815 | 104828166 | A | G | 0.1879 | ALCAM | intergenic | -0.1046 | 0.017 | 7.25E-10 | -0.0495 | 0.0357 | 1.66E-01 | -0.0769 | 0.0128 | 1.73E-09 | 0 |
| rs9838717 | 3 | 104732961 | 24 | 104646815 | 104828166 | T | G | 0.1769 | ALCAM | intergenic | -0.11 | 0.0172 | 1.66E-10 | -0.0485 | 0.0364 | 1.83E-01 | -0.0772 | 0.0129 | 1.95E-09 | 0 |
| rs9858732 | 3 | 104733096 | 24 | 104646815 | 104828166 | A | T | 0.1769 | ALCAM | intergenic | -0.1098 | 0.0172 | 1.75E-10 | -0.0487 | 0.0364 | 1.80E-01 | -0.0767 | 0.0129 | 2.40E-09 | 0 |
| rs72989342 | 3 | 104733118 | 24 | 104646815 | 104828166 | T | G | 0.1789 | ALCAM | intergenic | 0.1098 | 0.0172 | 1.73E-10 | 0.0512 | 0.0363 | 1.58E-01 | 0.0771 | 0.0129 | 2.04E-09 | 0 |
| rs28578901 | 3 | 104733192 | 24 | 104646815 | 104828166 | T | C | 0.1769 | ALCAM | intergenic | 0.1097 | 0.0172 | 1.79E-10 | 0.0476 | 0.0364 | 1.90E-01 | 0.0766 | 0.0129 | 2.60E-09 | 0 |
| rs16850758 | 3 | 104733368 | 24 | 104646815 | 104828166 | T | C | 0.1779 | ALCAM | intergenic | 0.1092 | 0.0172 | 2.16E-10 | 0.0498 | 0.0363 | 1.70E-01 | 0.0765 | 0.0129 | 2.68E-09 | 0 |
| rs16850761 | 3 | 104733455 | 24 | 104646815 | 104828166 | A | G | 0.1769 | ALCAM | intergenic | 0.1087 | 0.0172 | 2.61E-10 | 0.0500 | 0.0363 | 1.69E-01 | 0.0766 | 0.0129 | 2.58E-09 | 0 |
| rs16850762 | 3 | 104733601 | 24 | 104646815 | 104828166 | T | C | 0.1769 | ALCAM | intergenic | 0.1087 | 0.0172 | 2.67E-10 | 0.0512 | 0.0363 | 1.59E-01 | 0.0766 | 0.0129 | 2.58E-09 | 0 |
| rs16850763 | 3 | 104733610 | 24 | 104646815 | 104828166 | A | G | 0.1769 | ALCAM | intergenic | 0.1082 | 0.0172 | 3.27E-10 | 0.0499 | 0.0363 | 1.70E-01 | 0.0765 | 0.0129 | 2.62E-09 | 0 |
| rs28661692 | 3 | 104733669 | 24 | 104646815 | 104828166 | A | G | 0.1769 | ALCAM | intergenic | -0.1088 | 0.0172 | 2.59E-10 | -0.0511 | 0.0363 | 1.59E-01 | -0.0765 | 0.0129 | 2.68E-09 | 0 |
| rs16850766 | 3 | 104733714 | 24 | 104646815 | 104828166 | A | G | 0.1769 | ALCAM | intergenic | -0.1083 | 0.0172 | 3.08E-10 | -0.0499 | 0.0363 | 1.69E-01 | -0.0763 | 0.0129 | 2.86E-09 | 0 |
| rs28564343 | 3 | 104735374 | 24 | 104646815 | 104828166 | T | G | 0.1769 | ALCAM | intergenic | 0.1082 | 0.0172 | 3.16E-10 | 0.0489 | 0.0362 | 1.77E-01 | 0.0764 | 0.0129 | 2.84E-09 | 0 |
| rs28421781 | 3 | 104735446 | 24 | 104646815 | 104828166 | A | G | 0.1769 | ALCAM | intergenic | 0.1093 | 0.0172 | 2.40E-10 | 0.0502 | 0.0362 | 1.65E-01 | 0.0769 | 0.0129 | 2.29E-09 | 0 |
| rs28551708 | 3 | 104735633 | 24 | 104646815 | 104828166 | T | G | 0.1769 | ALCAM | intergenic | -0.1122 | 0.0177 | 2.55E-10 | -0.0492 | 0.0362 | 1.75E-01 | -0.0838 | 0.0135 | 6.07E-10 | 0 |
| rs28594489 | 3 | 104735634 | 24 | 104646815 | 104828166 | T | C | 0.1769 | ALCAM | intergenic | -0.113 | 0.0178 | 2.12E-10 | -0.0492 | 0.0362 | 1.75E-01 | -0.0844 | 0.0136 | 5.46E-10 | 0 |
| rs28423675 | 3 | 104735824 | 24 | 104646815 | 104828166 | A | G | 0.1769 | ALCAM | intergenic | -0.1083 | 0.0172 | 3.06E-10 | -0.0490 | 0.0362 | 1.76E-01 | -0.0768 | 0.0129 | 2.26E-09 | 0 |
| rs7646614 | 3 | 104736349 | 24 | 104646815 | 104828166 | A | G | 0.1769 | ALCAM | intergenic | 0.1085 | 0.0172 | 2.90E-10 | 0.0494 | 0.0363 | 1.73E-01 | 0.0769 | 0.0129 | 2.24E-09 | 0 |
| rs6437554 | 3 | 104736464 | 24 | 104646815 | 104828166 | A | G | 0.1789 | ALCAM | intergenic | 0.1092 | 0.0172 | 2.25E-10 | 0.0504 | 0.0363 | 1.65E-01 | 0.0773 | 0.0129 | 1.78E-09 | 0 |
| rs34360357 | 3 | 104736675 | 24 | 104646815 | 104828166 | T | G | 0.4135 | ALCAM | intergenic | 0.0877 | 0.0132 | 3.21E-11 | 0.0791 | 0.0277 | 4.28E-03 | 0.0516 | 0.0101 | 2.96E-07 | 1 |
| rs17387070 | 3 | 104736772 | 24 | 104646815 | 104828166 | A | G | 0.4135 | ALCAM | intergenic | -0.0878 | 0.0132 | 3.28E-11 | -0.0798 | 0.0277 | 3.93E-03 | -0.0522 | 0.0101 | 2.31E-07 | 1 |
| rs6781572 | 3 | 104736846 | 24 | 104646815 | 104828166 | A | G | 0.1789 | ALCAM | intergenic | -0.109 | 0.0172 | 2.40E-10 | -0.0504 | 0.0363 | 1.65E-01 | -0.0775 | 0.0129 | 1.64E-09 | 0 |
| rs73852547 | 3 | 104737138 | 24 | 104646815 | 104828166 | T | C | 0.1769 | ALCAM | intergenic | -0.1084 | 0.0172 | 3.06E-10 | -0.0491 | 0.0363 | 1.76E-01 | -0.0769 | 0.0129 | 2.24E-09 | 0 |
| rs72989352 | 3 | 104737400 | 24 | 104646815 | 104828166 | A | T | 0.1769 | ALCAM | intergenic | -0.1089 | 0.0172 | 2.54E-10 | -0.0507 | 0.0363 | 1.63E-01 | -0.0769 | 0.0129 | 2.25E-09 | 0 |
| rs76481720 | 3 | 104737632 | 24 | 104646815 | 104828166 | A | G | 0.1769 | ALCAM | intergenic | 0.1086 | 0.0172 | 2.84E-10 | 0.0471 | 0.0363 | 1.95E-01 | 0.0768 | 0.0129 | 2.42E-09 | 0 |
| rs60072080 | 3 | 104737669 | 24 | 104646815 | 104828166 | A | G | 0.1769 | ALCAM | intergenic | -0.1084 | 0.0172 | 3.00E-10 | -0.0494 | 0.0363 | 1.73E-01 | -0.0770 | 0.0129 | 2.22E-09 | 0 |
| rs58205062 | 3 | 104737769 | 24 | 104646815 | 104828166 | T | G | 0.1759 | ALCAM | intergenic | 0.1077 | 0.0172 | 3.87E-10 | 0.0510 | 0.0363 | 1.60E-01 | 0.0764 | 0.0129 | 2.94E-09 | 0 |
| rs28735393 | 3 | 104738215 | 24 | 104646815 | 104828166 | A | G | 0.1769 | ALCAM | intergenic | -0.1084 | 0.0172 | 3.08E-10 | -0.0495 | 0.0363 | 1.73E-01 | -0.0767 | 0.0129 | 2.52E-09 | 0 |
| rs9857628 | 3 | 104738370 | 24 | 104646815 | 104828166 | A | T | 0.1769 | ALCAM | intergenic | -0.1084 | 0.0172 | 3.08E-10 | -0.0496 | 0.0363 | 1.72E-01 | -0.0767 | 0.0129 | 2.52E-09 | 0 |
| rs6437555 | 3 | 104738746 | 24 | 104646815 | 104828166 | A | G | 0.1769 | ALCAM | intergenic | -0.1105 | 0.0173 | 1.59E-10 | -0.0507 | 0.0365 | 1.65E-01 | -0.0764 | 0.0129 | 2.88E-09 | 0 |
| rs6437556 | 3 | 104738751 | 24 | 104646815 | 104828166 | A | G | 0.1769 | ALCAM | intergenic | 0.1111 | 0.0173 | 1.38E-10 | 0.0513 | 0.0365 | 1.60E-01 | 0.0766 | 0.0129 | 2.70E-09 | 0 |
| rs9848945 | 3 | 104740628 | 24 | 104646815 | 104828166 | T | G | 0.173 | ALCAM | intergenic | -0.1084 | 0.0172 | 3.09E-10 | -0.0519 | 0.0364 | 1.54E-01 | -0.0755 | 0.0129 | 4.36E-09 | 0 |
| rs7626601 | 3 | 104742187 | 24 | 104646815 | 104828166 | A | C | 0.173 | ALCAM | intergenic | -0.1083 | 0.0172 | 3.21E-10 | -0.0520 | 0.0364 | 1.54E-01 | -0.0757 | 0.0129 | 4.00E-09 | 0 |

| | | | | | | | | | | | Meta-GWAS | | | UKBB | | | ENIGMA* | | | |
|---|---|---|---|---|---|---|---|---|---|---|---|---|---|---|---|---|---|---|---|---|
| rsID | chr | pos (hg19) | Genomic Locus | start | end | A1 | A2 | MAF | Nearest Genes | Function | Beta | SE | P-value | Beta | SE | P-value | Beta | SE | P-value | Replicated** |
| rs9883825 | 3 | 104745091 | 24 | 104646815 | 104828166 | A | G | 0.4145 | ALCAM | intergenic | -0.0881 | 0.0132 | 2.68E-11 | -0.0770 | 0.0277 | 5.48E-03 | -0.0508 | 0.0101 | 4.73E-07 | 1 |
| rs9288797 | 3 | 104745255 | 24 | 104646815 | 104828166 | A | G | 0.173 | ALCAM | intergenic | -0.1081 | 0.0172 | 3.73E-10 | -0.0520 | 0.0364 | 1.53E-01 | -0.0752 | 0.0129 | 5.50E-09 | 0 |
| rs79873095 | 3 | 104745777 | 24 | 104646815 | 104828166 | T | G | 0.173 | ALCAM | intergenic | -0.1081 | 0.0172 | 3.67E-10 | -0.0530 | 0.0364 | 1.46E-01 | -0.0751 | 0.0129 | 5.83E-09 | 0 |
| rs9825019 | 3 | 104748374 | 24 | 104646815 | 104828166 | A | G | 0.4155 | ALCAM | intergenic | -0.0876 | 0.0132 | 3.58E-11 | -0.0758 | 0.0277 | 6.27E-03 | -0.0493 | 0.0101 | 1.06E-06 | 1 |
| rs7340550 | 3 | 104754015 | 24 | 104646815 | 104828166 | T | G | 0.173 | ALCAM | intergenic | 0.1066 | 0.0173 | 6.79E-10 | 0.0505 | 0.0362 | 1.64E-01 | 0.0751 | 0.0129 | 5.76E-09 | 0 |
| rs13319556 | 3 | 104754543 | 24 | 104646815 | 104828166 | A | T | 0.174 | ALCAM | intergenic | -0.1069 | 0.0173 | 6.20E-10 | -0.0506 | 0.0362 | 1.63E-01 | -0.0743 | 0.0129 | 8.19E-09 | 0 |
| rs13325696 | 3 | 104755481 | 24 | 104646815 | 104828166 | A | G | 0.173 | ALCAM | intergenic | -0.1068 | 0.0173 | 6.33E-10 | -0.0502 | 0.0362 | 1.66E-01 | -0.0744 | 0.0129 | 7.89E-09 | 0 |
| rs9834649 | 3 | 104755849 | 24 | 104646815 | 104828166 | A | C | 0.2913 | ALCAM | intergenic | -0.1119 | 0.0141 | 2.21E-15 | -0.0945 | 0.0298 | 1.52E-03 | -0.0659 | 0.0107 | 7.15E-10 | 1 |
| rs1503079 | 3 | 104759032 | 24 | 104646815 | 104828166 | T | C | 0.4891 | ALCAM | intergenic | 0.0734 | 0.0129 | 1.35E-08 | 0.0679 | 0.0271 | 1.21E-02 | 0.0318 | 0.0099 | 1.30E-03 | 1 |
| rs955251 | 3 | 104759845 | 24 | 104646815 | 104828166 | A | C | 0.4891 | ALCAM | intergenic | -0.0734 | 0.0129 | 1.38E-08 | -0.0668 | 0.0270 | 1.35E-02 | -0.0320 | 0.0099 | 1.20E-03 | 1 |
| rs955250 | 3 | 104759895 | 24 | 104646815 | 104828166 | A | G | 0.4891 | ALCAM | intergenic | -0.0734 | 0.0129 | 1.34E-08 | -0.0673 | 0.0270 | 1.29E-02 | -0.0320 | 0.0099 | 1.19E-03 | 1 |
| rs6800125 | 3 | 104760179 | 24 | 104646815 | 104828166 | A | G | 0.4881 | ALCAM | intergenic | -0.0733 | 0.0129 | 1.41E-08 | -0.0670 | 0.0270 | 1.33E-02 | -0.0320 | 0.0099 | 1.18E-03 | 1 |
| rs1566720 | 3 | 104761408 | 24 | 104646815 | 104828166 | A | C | 0.4891 | ALCAM | intergenic | 0.0734 | 0.0129 | 1.33E-08 | 0.0670 | 0.0270 | 1.32E-02 | 0.0321 | 0.0099 | 1.14E-03 | 1 |
| rs1566719 | 3 | 104761448 | 24 | 104646815 | 104828166 | A | T | 0.4891 | ALCAM | intergenic | -0.0735 | 0.0129 | 1.31E-08 | -0.0669 | 0.0270 | 1.33E-02 | NA | NA | NA | NA |
| rs1566718 | 3 | 104761637 | 24 | 104646815 | 104828166 | A | G | 0.4891 | ALCAM | intergenic | 0.0732 | 0.0129 | 1.54E-08 | 0.0669 | 0.0270 | 1.33E-02 | 0.0315 | 0.0099 | 1.45E-03 | 1 |
| rs66833862 | 3 | 104762515 | 24 | 104646815 | 104828166 | T | C | 0.4901 | ALCAM | intergenic | 0.075 | 0.013 | 8.92E-09 | 0.0663 | 0.0270 | 1.42E-02 | 0.0331 | 0.0100 | 9.92E-04 | 1 |
| rs12633310 | 3 | 104763667 | 24 | 104646815 | 104828166 | A | G | 0.4891 | ALCAM | intergenic | 0.0735 | 0.0129 | 1.30E-08 | 0.0661 | 0.0270 | 1.46E-02 | 0.0323 | 0.0099 | 1.09E-03 | 1 |
| rs56169547 | 3 | 104764040 | 24 | 104646815 | 104828166 | A | G | 0.4891 | ALCAM | intergenic | 0.075 | 0.013 | 7.39E-09 | 0.0663 | 0.0270 | 1.42E-02 | 0.0321 | 0.0099 | 1.25E-03 | 1 |
| rs35563949 | 3 | 104765663 | 24 | 104646815 | 104828166 | A | T | 0.4901 | ALCAM | intergenic | 0.0738 | 0.0129 | 1.13E-08 | 0.0679 | 0.0270 | 1.20E-02 | NA | NA | NA | NA |
| rs4894900 | 3 | 104766656 | 24 | 104646815 | 104828166 | A | G | 0.4901 | ALCAM | intergenic | -0.0736 | 0.0129 | 1.22E-08 | -0.0675 | 0.0270 | 1.24E-02 | -0.0325 | 0.0099 | 1.00E-03 | 1 |
| rs1566716 | 3 | 104767455 | 24 | 104646815 | 104828166 | T | C | 0.4901 | ALCAM | intergenic | -0.0741 | 0.0129 | 1.00E-08 | -0.0677 | 0.0270 | 1.22E-02 | -0.0325 | 0.0099 | 1.02E-03 | 1 |
| rs1566717 | 3 | 104767509 | 24 | 104646815 | 104828166 | A | T | 0.4901 | ALCAM | intergenic | -0.0741 | 0.0129 | 1.00E-08 | -0.0676 | 0.0270 | 1.23E-02 | NA | NA | NA | NA |
| rs953850 | 3 | 104767889 | 24 | 104646815 | 104828166 | A | T | 0.4881 | ALCAM | intergenic | -0.0733 | 0.0129 | 1.45E-08 | -0.0677 | 0.0270 | 1.22E-02 | NA | NA | NA | NA |
| rs13090983 | 3 | 104768133 | 24 | 104646815 | 104828166 | C | G | 0.4881 | ALCAM | intergenic | 0.0733 | 0.0129 | 1.45E-08 | 0.0678 | 0.0270 | 1.21E-02 | NA | NA | NA | NA |
| rs10933801 | 3 | 104769160 | 24 | 104646815 | 104828166 | T | C | 0.4881 | ALCAM | intergenic | 0.0733 | 0.0129 | 1.42E-08 | 0.0677 | 0.0270 | 1.22E-02 | 0.0325 | 0.0099 | 1.01E-03 | 1 |
| rs12633740 | 3 | 104769506 | 24 | 104646815 | 104828166 | A | C | 0.4881 | ALCAM | intergenic | 0.0746 | 0.013 | 9.61E-09 | 0.0678 | 0.0270 | 1.22E-02 | 0.0345 | 0.0100 | 5.39E-04 | 1 |
| rs13059629 | 3 | 104769921 | 24 | 104646815 | 104828166 | A | G | 0.4881 | ALCAM | intergenic | -0.0735 | 0.0129 | 1.29E-08 | -0.0678 | 0.0270 | 1.20E-02 | -0.0324 | 0.0099 | 1.02E-03 | 1 |
| rs13060672 | 3 | 104770357 | 24 | 104646815 | 104828166 | T | C | 0.4881 | ALCAM | intergenic | 0.0736 | 0.0129 | 1.28E-08 | 0.0678 | 0.0270 | 1.20E-02 | 0.0325 | 0.0099 | 1.01E-03 | 1 |
| rs67773865 | 3 | 104770966 | 24 | 104646815 | 104828166 | T | C | 0.4881 | ALCAM | intergenic | 0.0741 | 0.0129 | 1.03E-08 | 0.0679 | 0.0270 | 1.20E-02 | 0.0329 | 0.0099 | 8.75E-04 | 1 |
| rs13066039 | 3 | 104771161 | 24 | 104646815 | 104828166 | A | G | 0.4881 | ALCAM | intergenic | -0.0737 | 0.0129 | 1.22E-08 | -0.0683 | 0.0270 | 1.15E-02 | -0.0325 | 0.0099 | 1.01E-03 | 1 |
| rs13066506 | 3 | 104771184 | 24 | 104646815 | 104828166 | A | G | 0.4881 | ALCAM | intergenic | 0.0738 | 0.0129 | 1.18E-08 | 0.0679 | 0.0270 | 1.20E-02 | 0.0325 | 0.0099 | 1.02E-03 | 1 |
| rs35154832 | 3 | 104771559 | 24 | 104646815 | 104828166 | T | G | 0.4881 | ALCAM | intergenic | -0.0738 | 0.0129 | 1.16E-08 | -0.0688 | 0.0271 | 1.10E-02 | -0.0324 | 0.0099 | 1.03E-03 | 1 |
| rs34044731 | 3 | 104771580 | 24 | 104646815 | 104828166 | A | G | 0.4881 | ALCAM | intergenic | 0.0738 | 0.0129 | 1.14E-08 | 0.0690 | 0.0270 | 1.08E-02 | 0.0325 | 0.0099 | 1.00E-03 | 1 |
| rs1910328 | 3 | 104771826 | 24 | 104646815 | 104828166 | T | C | 0.4881 | ALCAM | intergenic | 0.0733 | 0.0129 | 1.47E-08 | 0.0678 | 0.0270 | 1.21E-02 | 0.0326 | 0.0099 | 9.76E-04 | 1 |
| rs1910329 | 3 | 104771860 | 24 | 104646815 | 104828166 | C | G | 0.4881 | ALCAM | intergenic | 0.0734 | 0.0129 | 1.35E-08 | 0.0679 | 0.0270 | 1.20E-02 | NA | NA | NA | NA |
| rs35507961 | 3 | 104772393 | 24 | 104646815 | 104828166 | T | C | 0.4881 | ALCAM | intergenic | 0.0735 | 0.0129 | 1.33E-08 | 0.0678 | 0.0270 | 1.21E-02 | 0.0324 | 0.0099 | 1.02E-03 | 1 |
| rs35705376 | 3 | 104773479 | 24 | 104646815 | 104828166 | T | C | 0.4881 | ALCAM | intergenic | -0.0735 | 0.0129 | 1.30E-08 | -0.0670 | 0.0270 | 1.31E-02 | -0.0324 | 0.0099 | 1.05E-03 | 1 |
| rs12637494 | 3 | 104773965 | 24 | 104646815 | 104828166 | T | C | 0.4881 | ALCAM | intergenic | 0.0735 | 0.0129 | 1.28E-08 | 0.0670 | 0.0270 | 1.31E-02 | 0.0326 | 0.0099 | 9.71E-04 | 1 |
| rs10933802 | 3 | 104774304 | 24 | 104646815 | 104828166 | A | G | 0.4881 | ALCAM | intergenic | -0.0737 | 0.0129 | 1.19E-08 | -0.0670 | 0.0270 | 1.31E-02 | -0.0328 | 0.0099 | 9.15E-04 | 1 |
| rs10804438 | 3 | 104774460 | 24 | 104646815 | 104828166 | T | C | 0.4881 | ALCAM | intergenic | -0.0736 | 0.0129 | 1.23E-08 | -0.0670 | 0.0270 | 1.31E-02 | -0.0325 | 0.0099 | 1.00E-03 | 1 |
| rs12163622 | 3 | 104774805 | 24 | 104646815 | 104828166 | T | C | 0.4881 | ALCAM | intergenic | 0.0736 | 0.0129 | 1.25E-08 | 0.0667 | 0.0270 | 1.35E-02 | 0.0325 | 0.0099 | 9.88E-04 | 1 |
| rs12163623 | 3 | 104774833 | 24 | 104646815 | 104828166 | T | G | 0.4881 | ALCAM | intergenic | 0.0737 | 0.0129 | 1.22E-08 | 0.0670 | 0.0270 | 1.31E-02 | 0.0325 | 0.0099 | 1.00E-03 | 1 |
| rs10933803 | 3 | 104775181 | 24 | 104646815 | 104828166 | A | G | 0.4881 | ALCAM | intergenic | -0.0737 | 0.0129 | 1.23E-08 | -0.0670 | 0.0270 | 1.31E-02 | -0.0325 | 0.0099 | 1.01E-03 | 1 |
| rs12639601 | 3 | 104775272 | 24 | 104646815 | 104828166 | A | G | 0.4881 | ALCAM | intergenic | -0.0731 | 0.0129 | 1.63E-08 | -0.0670 | 0.0270 | 1.31E-02 | -0.0327 | 0.0099 | 9.58E-04 | 1 |
| rs13064544 | 3 | 104776332 | 24 | 104646815 | 104828166 | A | C | 0.4881 | ALCAM | intergenic | -0.0735 | 0.0129 | 1.31E-08 | -0.0670 | 0.0270 | 1.31E-02 | -0.0324 | 0.0099 | 1.03E-03 | 1 |
| rs57241486 | 3 | 104776636 | 24 | 104646815 | 104828166 | C | G | 0.4881 | ALCAM | intergenic | -0.0737 | 0.0129 | 1.19E-08 | -0.0670 | 0.0270 | 1.31E-02 | NA | NA | NA | NA |
| rs6797909 | 3 | 104777026 | 24 | 104646815 | 104828166 | T | G | 0.4881 | ALCAM | intergenic | 0.0738 | 0.0129 | 1.18E-08 | 0.0670 | 0.0270 | 1.31E-02 | 0.0328 | 0.0099 | 9.16E-04 | 1 |
| rs6798196 | 3 | 104777290 | 24 | 104646815 | 104828166 | C | G | 0.4881 | ALCAM | intergenic | 0.0736 | 0.0129 | 1.27E-08 | 0.0670 | 0.0270 | 1.31E-02 | NA | NA | NA | NA |
| rs6798649 | 3 | 104777711 | 24 | 104646815 | 104828166 | T | G | 0.4881 | ALCAM | intergenic | 0.0735 | 0.0129 | 1.34E-08 | 0.0668 | 0.0270 | 1.35E-02 | 0.0327 | 0.0099 | 9.48E-04 | 1 |
| rs10933804 | 3 | 104777958 | 24 | 104646815 | 104828166 | T | C | 0.4881 | ALCAM | intergenic | -0.0739 | 0.0129 | 1.15E-08 | -0.0658 | 0.0270 | 1.50E-02 | -0.0329 | 0.0099 | 8.93E-04 | 1 |
| rs10933805 | 3 | 104777973 | 24 | 104646815 | 104828166 | C | G | 0.4881 | ALCAM | intergenic | -0.0739 | 0.0129 | 1.15E-08 | -0.0658 | 0.0270 | 1.50E-02 | NA | NA | NA | NA |
| rs11708913 | 3 | 104778145 | 24 | 104646815 | 104828166 | A | T | 0.4881 | ALCAM | intergenic | -0.0739 | 0.0129 | 1.13E-08 | -0.0666 | 0.0270 | 1.37E-02 | NA | NA | NA | NA |
| rs11708967 | 3 | 104778270 | 24 | 104646815 | 104828166 | A | C | 0.4851 | ALCAM | intergenic | -0.0803 | 0.0133 | 1.61E-09 | -0.0667 | 0.0270 | 1.37E-02 | -0.0410 | 0.0104 | 8.08E-05 | 1 |
| rs34967119 | 3 | 104778430 | 24 | 104646815 | 104828166 | A | G | 0.4881 | ALCAM | intergenic | 0.0765 | 0.0133 | 9.36E-09 | 0.0667 | 0.0270 | 1.36E-02 | 0.0342 | 0.0104 | 1.00E-03 | 1 |
| rs2047806 | 3 | 104778810 | 24 | 104646815 | 104828166 | A | C | 0.4881 | ALCAM | intergenic | 0.0736 | 0.0129 | 1.30E-08 | 0.0679 | 0.0270 | 1.20E-02 | 0.0327 | 0.0099 | 9.47E-04 | 1 |
| rs62260620 | 3 | 104779102 | 24 | 104646815 | 104828166 | A | G | 0.4851 | ALCAM | intergenic | -0.0736 | 0.0129 | 1.29E-08 | -0.0687 | 0.0270 | 1.10E-02 | -0.0329 | 0.0099 | 8.73E-04 | 1 |
| rs56146536 | 3 | 104779138 | 24 | 104646815 | 104828166 | T | C | 0.4881 | ALCAM | intergenic | -0.0736 | 0.0129 | 1.29E-08 | -0.0687 | 0.0270 | 1.10E-02 | -0.0326 | 0.0099 | 9.74E-04 | 1 |
| rs1032796 | 3 | 104779561 | 24 | 104646815 | 104828166 | A | G | 0.4881 | ALCAM | intergenic | -0.0736 | 0.0129 | 1.28E-08 | -0.0694 | 0.0270 | 1.02E-02 | -0.0327 | 0.0099 | 9.44E-04 | 1 |
| rs7651259 | 3 | 104779847 | 24 | 104646815 | 104828166 | T | C | 0.4881 | ALCAM | intergenic | 0.0735 | 0.0129 | 1.36E-08 | 0.0699 | 0.0270 | 9.71E-03 | 0.0326 | 0.0099 | 9.70E-04 | 1 |
| rs2399028 | 3 | 104779893 | 24 | 104646815 | 104828166 | A | G | 0.4881 | ALCAM | intergenic | 0.0736 | 0.0129 | 1.26E-08 | 0.0694 | 0.0270 | 1.02E-02 | 0.0327 | 0.0099 | 9.39E-04 | 1 |
| rs2134526 | 3 | 104780244 | 24 | 104646815 | 104828166 | A | G | 0.4881 | ALCAM | intergenic | -0.0736 | 0.0129 | 1.28E-08 | -0.0694 | 0.0270 | 1.02E-02 | -0.0327 | 0.0099 | 9.30E-04 | 1 |
| rs13073998 | 3 | 104780902 | 24 | 104646815 | 104828166 | T | C | 0.4881 | ALCAM | intergenic | -0.0738 | 0.0129 | 1.16E-08 | -0.0694 | 0.0270 | 1.02E-02 | -0.0327 | 0.0099 | 9.37E-04 | 1 |
| rs13095276 | 3 | 104781455 | 24 | 104646815 | 104828166 | T | C | 0.4861 | ALCAM | intergenic | 0.0739 | 0.0129 | 1.11E-08 | 0.0696 | 0.0270 | 9.94E-03 | 0.0329 | 0.0099 | 8.71E-04 | 1 |
| rs9883267 | 3 | 104782005 | 24 | 104646815 | 104828166 | T | C | 0.4165 | ALCAM | intergenic | -0.0841 | 0.0132 | 1.69E-10 | -0.0702 | 0.0275 | 1.07E-02 | -0.0486 | 0.0101 | 1.33E-06 | 1 |
| rs6809713 | 3 | 104782428 | 24 | 104646815 | 104828166 | T | C | 0.4881 | ALCAM | intergenic | -0.0737 | 0.0129 | 1.20E-08 | -0.0694 | 0.0270 | 1.02E-02 | -0.0327 | 0.0099 | 9.17E-04 | 1 |
| rs6800325 | 3 | 104783012 | 24 | 104646815 | 104828166 | A | G | 0.4881 | ALCAM | intergenic | 0.0738 | 0.0129 | 1.16E-08 | 0.0695 | 0.0270 | 1.00E-02 | 0.0326 | 0.0099 | 9.55E-04 | 1 |
| rs6800248 | 3 | 104783055 | 24 | 104646815 | 104828166 | T | C | 0.4881 | ALCAM | intergenic | 0.0737 | 0.0129 | 1.18E-08 | 0.0695 | 0.0270 | 1.00E-02 | 0.0327 | 0.0099 | 9.35E-04 | 1 |
| rs10933806 | 3 | 104783475 | 24 | 104646815 | 104828166 | T | G | 0.4881 | ALCAM | intergenic | -0.0745 | 0.0129 | 8.51E-09 | -0.0695 | 0.0270 | 1.00E-02 | -0.0327 | 0.0099 | 9.26E-04 | 1 |
| rs9814707 | 3 | 104783737 | 24 | 104646815 | 104828166 | A | G | 0.1779 | ALCAM | intergenic | -0.1057 | 0.0172 | 7.23E-10 | -0.0422 | 0.0362 | 2.44E-01 | -0.0725 | 0.0128 | 1.68E-08 | 0 |
| rs11706888 | 3 | 104783773 | 24 | 104646815 | 104828166 | T | C | 0.4881 | ALCAM | intergenic | 0.0736 | 0.0129 | 1.27E-08 | 0.0696 | 0.0270 | 1.00E-02 | 0.0326 | 0.0099 | 9.68E-04 | 1 |
| rs12635716 | 3 | 104784020 | 24 | 104646815 | 104828166 | A | G | 0.4881 | ALCAM | intergenic | -0.0737 | 0.0129 | 1.19E-08 | -0.0695 | 0.0270 | 1.00E-02 | -0.0326 | 0.0099 | 9.53E-04 | 1 |
| rs10933807 | 3 | 104784330 | 24 | 104646815 | 104828166 | T | C | 0.4881 | ALCAM | intergenic | 0.0743 | 0.0129 | 9.14E-09 | 0.0707 | 0.0270 | 8.84E-03 | 0.0326 | 0.0099 | 9.54E-04 | 1 |
| rs6437560 | 3 | 104784922 | 24 | 104646815 | 104828166 | A | G | 0.4881 | ALCAM | intergenic | -0.0738 | 0.0129 | 1.14E-08 | -0.0702 | 0.0270 | 9.35E-03 | -0.0326 | 0.0099 | 9.76E-04 | 1 |
| rs7618073 | 3 | 104785204 | 24 | 104646815 | 104828166 | T | C | 0.4881 | ALCAM | intergenic | -0.0737 | 0.0129 | 1.21E-08 | -0.0695 | 0.0270 | 1.01E-02 | -0.0326 | 0.0099 | 9.68E-04 | 1 |
| rs7650627 | 3 | 104785216 | 24 | 104646815 | 104828166 | A | C | 0.4881 | ALCAM | intergenic | 0.0738 | 0.0129 | 1.12E-08 | 0.0702 | 0.0270 | 9.30E-03 | 0.0326 | 0.0099 | 9.70E-04 | 1 |
| rs7650875 | 3 | 104785261 | 24 | 104646815 | 104828166 | A | G | 0.4881 | ALCAM | intergenic | 0.0737 | 0.0129 | 1.23E-08 | 0.0702 | 0.0270 | 9.33E-03 | 0.0328 | 0.0099 | 9.27E-04 | 1 |
| rs7628576 | 3 | 104785371 | 24 | 104646815 | 104828166 | A | G | 0.4881 | ALCAM | intergenic | -0.0738 | 0.0129 | 1.13E-08 | -0.0702 | 0.0270 | 9.34E-03 | -0.0326 | 0.0099 | 9.80E-04 | 1 |
| rs1104503 | 3 | 104785888 | 24 | 104646815 | 104828166 | T | C | 0.4881 | ALCAM | intergenic | -0.0738 | 0.0129 | 1.13E-08 | -0.0702 | 0.0270 | 9.34E-03 | -0.0325 | 0.0099 | 9.82E-04 | 1 |
| rs1104502 | 3 | 104785935 | 24 | 104646815 | 104828166 | T | C | 0.4871 | ALCAM | intergenic | 0.0738 | 0.0129 | 1.13E-08 | 0.0702 | 0.0270 | 9.33E-03 | 0.0326 | 0.0099 | 9.66E-04 | 1 |
| rs1587707 | 3 | 104786133 | 24 | 104646815 | 104828166 | T | G | 0.4891 | ALCAM | intergenic | -0.0746 | 0.013 | 1.05E-08 | -0.0680 | 0.0270 | 1.20E-02 | -0.0330 | 0.0100 | 9.57E-04 | 1 |
| rs2399027 | 3 | 104786221 | 24 | 104646815 | 104828166 | T | G | 0.4881 | ALCAM | intergenic | 0.0739 | 0.0129 | 1.12E-08 | 0.0702 | 0.0270 | 9.29E-03 | 0.0326 | 0.0099 | 9.74E-04 | 1 |
| rs57951267 | 3 | 104786428 | 24 | 104646815 | 104828166 | A | G | 0.4881 | ALCAM | intergenic | -0.0739 | 0.0129 | 1.12E-08 | -0.0702 | 0.0270 | 9.29E-03 | -0.0326 | 0.0099 | 9.74E-04 | 1 |
| rs7609968 | 3 | 104786544 | 24 | 104646815 | 104828166 | T | G | 0.4881 | ALCAM | intergenic | 0.0739 | 0.0129 | 1.11E-08 | 0.0702 | 0.0270 | 9.37E-03 | 0.0326 | 0.0099 | 9.73E-04 | 1 |
| rs2174113 | 3 | 104786848 | 24 | 104646815 | 104828166 | A | C | 0.4871 | ALCAM | intergenic | -0.0738 | 0.0129 | 1.17E-08 | -0.0675 | 0.0270 | 1.24E-02 | -0.0326 | 0.0099 | 9.65E-04 | 1 |
| rs2134527 | 3 | 104787185 | 24 | 104646815 | 104828166 | A | G | 0.4881 | ALCAM | intergenic | -0.0738 | 0.0129 | 1.17E-08 | -0.0675 | 0.0270 | 1.24E-02 | -0.0326 | 0.0099 | 9.68E-04 | 1 |
| rs2174114 | 3 | 104787255 | 24 | 104646815 | 104828166 | A | C | 0.4881 | ALCAM | intergenic | 0.0737 | 0.0129 | 1.25E-08 | 0.0675 | 0.0270 | 1.24E-02 | 0.0328 | 0.0099 | 9.14E-04 | 1 |
| rs62258876 | 3 | 104787528 | 24 | 104646815 | 104828166 | A | G | 0.4881 | ALCAM | intergenic | -0.0738 | 0.0129 | 1.17E-08 | -0.0675 | 0.0270 | 1.24E-02 | -0.0326 | 0.0099 | 9.68E-04 | 1 |
| rs6437561 | 3 | 104788531 | 24 | 104646815 | 104828166 | A | C | 0.4881 | ALCAM | intergenic | 0.0738 | 0.0129 | 1.18E-08 | 0.0675 | 0.0270 | 1.24E-02 | 0.0326 | 0.0099 | 9.68E-04 | 1 |
| rs2895296 | 3 | 104788762 | 24 | 104646815 | 104828166 | A | G | 0.4881 | ALCAM | intergenic | -0.0742 | 0.0129 | 9.83E-09 | -0.0674 | 0.0270 | 1.26E-02 | -0.0326 | 0.0099 | 9.70E-04 | 1 |
| rs2399026 | 3 | 104789107 | 24 | 104646815 | 104828166 | T | G | 0.4881 | ALCAM | intergenic | 0.0737 | 0.0129 | 1.23E-08 | 0.0676 | 0.0270 | 1.23E-02 | 0.0326 | 0.0099 | 9.71E-04 | 1 |
| rs4522800 | 3 | 104789253 | 24 | 104646815 | 104828166 | A | T | 0.4881 | ALCAM | intergenic | -0.0737 | 0.0129 | 1.23E-08 | -0.0676 | 0.0270 | 1.23E-02 | NA | NA | NA | NA |

| rsID | chr | pos (hg19) | Genomic Locus | start | end | A1 | A2 | MAF | Nearest Genes | Function | Meta-GWAS | | | UKBB | | | ENIGMA* | | | Replicated** |
|---|---|---|---|---|---|---|---|---|---|---|---|---|---|---|---|---|---|---|---|---|
| | | | | | | | | | | | Beta | SE | P-value | Beta | SE | P-value | Beta | SE | P-value | |
| rs4525898 | 3 | 104789294 | 24 | 104646815 | 104828166 | A | C | 0.4881 | ALCAM | intergenic | -0.0737 | 0.0129 | 1.23E-08 | -0.0676 | 0.0270 | 1.23E-02 | -0.0326 | 0.0099 | 9.69E-04 | 1 |
| rs4276216 | 3 | 104789511 | 24 | 104646815 | 104828166 | A | G | 0.4881 | ALCAM | intergenic | 0.0732 | 0.013 | 1.70E-08 | 0.0680 | 0.0270 | 1.19E-02 | 0.0326 | 0.0099 | 1.03E-03 | 1 |
| rs4444734 | 3 | 104789572 | 24 | 104646815 | 104828166 | T | C | 0.4881 | ALCAM | intergenic | 0.0737 | 0.0129 | 1.24E-08 | 0.0680 | 0.0270 | 1.18E-02 | 0.0323 | 0.0099 | 1.06E-03 | 1 |
| rs62258878 | 3 | 104789859 | 24 | 104646815 | 104828166 | A | G | 0.4881 | ALCAM | intergenic | -0.0753 | 0.0131 | 8.75E-09 | -0.0680 | 0.0270 | 1.19E-02 | -0.0335 | 0.0101 | 9.06E-04 | 1 |
| rs17453821 | 3 | 104790120 | 24 | 104646815 | 104828166 | A | G | 0.4881 | ALCAM | intergenic | -0.0737 | 0.0129 | 1.24E-08 | -0.0680 | 0.0270 | 1.19E-02 | -0.0325 | 0.0099 | 1.00E-03 | 1 |
| rs2399025 | 3 | 104790262 | 24 | 104646815 | 104828166 | C | G | 0.4881 | ALCAM | intergenic | 0.0737 | 0.0129 | 1.22E-08 | 0.0680 | 0.0270 | 1.19E-02 | NA | NA | NA | NA |
| rs2895295 | 3 | 104790415 | 24 | 104646815 | 104828166 | T | G | 0.4881 | ALCAM | intergenic | -0.0737 | 0.0129 | 1.24E-08 | -0.0680 | 0.0270 | 1.19E-02 | -0.0326 | 0.0099 | 9.78E-04 | 1 |
| rs12633839 | 3 | 104790781 | 24 | 104646815 | 104828166 | T | C | 0.4891 | ALCAM | intergenic | -0.0744 | 0.0135 | 3.26E-08 | -0.0659 | 0.0273 | 1.57E-02 | -0.0362 | 0.0104 | 5.28E-04 | 1 |
| rs2174117 | 3 | 104790914 | 24 | 104646815 | 104828166 | A | G | 0.4881 | ALCAM | intergenic | 0.0751 | 0.0132 | 1.14E-08 | 0.0669 | 0.0271 | 1.34E-02 | 0.0335 | 0.0102 | 9.75E-04 | 1 |
| rs2174116 | 3 | 104790927 | 24 | 104646815 | 104828166 | C | G | 0.4881 | ALCAM | intergenic | 0.0761 | 0.0131 | 6.88E-09 | 0.0669 | 0.0271 | 1.34E-02 | NA | NA | NA | NA |
| rs2134542 | 3 | 104791106 | 24 | 104646815 | 104828166 | A | G | 0.4881 | ALCAM | intergenic | -0.0737 | 0.0129 | 1.22E-08 | -0.0680 | 0.0270 | 1.19E-02 | -0.0325 | 0.0099 | 9.97E-04 | 1 |
| rs2134541 | 3 | 104791188 | 24 | 104646815 | 104828166 | T | C | 0.4881 | ALCAM | intergenic | -0.074 | 0.013 | 1.14E-08 | -0.0680 | 0.0270 | 1.19E-02 | -0.0329 | 0.0099 | 9.00E-04 | 1 |
| rs12637068 | 3 | 104791391 | 24 | 104646815 | 104828166 | A | G | 0.4881 | ALCAM | intergenic | -0.0737 | 0.0129 | 1.25E-08 | -0.0680 | 0.0270 | 1.19E-02 | -0.0325 | 0.0099 | 1.00E-03 | 1 |
| rs1503082 | 3 | 104791710 | 24 | 104646815 | 104828166 | T | C | 0.4881 | ALCAM | intergenic | -0.0737 | 0.0129 | 1.25E-08 | -0.0680 | 0.0270 | 1.19E-02 | -0.0325 | 0.0099 | 1.00E-03 | 1 |
| rs1503083 | 3 | 104791792 | 24 | 104646815 | 104828166 | A | G | 0.4881 | ALCAM | intergenic | -0.0738 | 0.0129 | 1.18E-08 | -0.0680 | 0.0270 | 1.19E-02 | -0.0325 | 0.0099 | 1.01E-03 | 1 |
| rs1503084 | 3 | 104791812 | 24 | 104646815 | 104828166 | T | G | 0.4881 | ALCAM | intergenic | 0.0741 | 0.0129 | 1.05E-08 | 0.0679 | 0.0270 | 1.20E-02 | 0.0327 | 0.0099 | 9.30E-04 | 1 |
| rs2202246 | 3 | 104792169 | 24 | 104646815 | 104828166 | A | T | 0.4881 | ALCAM | intergenic | -0.0737 | 0.0129 | 1.25E-08 | -0.0680 | 0.0270 | 1.19E-02 | NA | NA | NA | NA |
| rs12632482 | 3 | 104792646 | 24 | 104646815 | 104828166 | A | G | 0.4881 | ALCAM | intergenic | 0.0736 | 0.0129 | 1.26E-08 | 0.0680 | 0.0270 | 1.18E-02 | 0.0325 | 0.0099 | 1.02E-03 | 1 |
| rs12635443 | 3 | 104792828 | 24 | 104646815 | 104828166 | T | G | 0.4881 | ALCAM | intergenic | -0.0736 | 0.0129 | 1.26E-08 | -0.0680 | 0.0270 | 1.18E-02 | -0.0324 | 0.0099 | 1.02E-03 | 1 |
| rs12496781 | 3 | 104793180 | 24 | 104646815 | 104828166 | A | G | 0.4881 | ALCAM | intergenic | -0.0737 | 0.0129 | 1.26E-08 | -0.0680 | 0.0270 | 1.18E-02 | -0.0324 | 0.0099 | 1.04E-03 | 1 |
| rs4894903 | 3 | 104794115 | 24 | 104646815 | 104828166 | A | T | 0.4881 | ALCAM | intergenic | 0.0736 | 0.0129 | 1.29E-08 | 0.0680 | 0.0270 | 1.19E-02 | NA | NA | NA | NA |
| rs7427968 | 3 | 104794254 | 24 | 104646815 | 104828166 | T | C | 0.4881 | ALCAM | intergenic | 0.0735 | 0.013 | 1.40E-08 | 0.0680 | 0.0270 | 1.19E-02 | 0.0326 | 0.0099 | 9.84E-04 | 1 |
| rs1116674 | 3 | 104794380 | 24 | 104646815 | 104828166 | A | G | 0.4881 | ALCAM | intergenic | -0.0736 | 0.0129 | 1.28E-08 | -0.0680 | 0.0270 | 1.19E-02 | -0.0322 | 0.0099 | 1.12E-03 | 1 |
| rs10933808 | 3 | 104795049 | 24 | 104646815 | 104828166 | T | C | 0.4881 | ALCAM | intergenic | -0.0735 | 0.0129 | 1.38E-08 | -0.0680 | 0.0270 | 1.19E-02 | -0.0322 | 0.0099 | 1.13E-03 | 1 |
| rs12632078 | 3 | 104796874 | 24 | 104646815 | 104828166 | T | C | 0.493 | ALCAM | intergenic | 0.0766 | 0.013 | 3.48E-09 | 0.0695 | 0.0271 | 1.02E-02 | 0.0339 | 0.0099 | 6.30E-04 | 1 |
| rs1587709 | 3 | 104796983 | 24 | 104646815 | 104828166 | A | G | 0.4861 | ALCAM | intergenic | -0.0734 | 0.0129 | 1.45E-08 | -0.0682 | 0.0270 | 1.17E-02 | -0.0323 | 0.0099 | 1.07E-03 | 1 |
| rs1587710 | 3 | 104797217 | 24 | 104646815 | 104828166 | T | C | 0.4861 | ALCAM | intergenic | -0.0733 | 0.013 | 1.71E-08 | -0.0682 | 0.0270 | 1.18E-02 | -0.0322 | 0.0099 | 1.14E-03 | 1 |
| rs1587716 | 3 | 104798102 | 24 | 104646815 | 104828166 | T | G | 0.493 | ALCAM | intergenic | 0.0764 | 0.013 | 4.41E-09 | 0.0703 | 0.0270 | 9.36E-03 | 0.0336 | 0.0099 | 7.05E-04 | 1 |
| rs11715847 | 3 | 104798853 | 24 | 104646815 | 104828166 | T | G | 0.4861 | ALCAM | intergenic | -0.073 | 0.013 | 1.97E-08 | -0.0680 | 0.0271 | 1.20E-02 | -0.0321 | 0.0099 | 1.20E-03 | 1 |
| rs79553445 | 3 | 104799105 | 24 | 104646815 | 104828166 | A | C | 0.169 | ALCAM | intergenic | 0.1025 | 0.0176 | 5.15E-09 | 0.0477 | 0.0370 | 1.98E-01 | 0.0771 | 0.0132 | 5.37E-09 | 0 |
| rs9809567 | 3 | 104799941 | 24 | 104646815 | 104828166 | A | C | 0.169 | ALCAM | intergenic | 0.1026 | 0.0176 | 5.06E-09 | 0.0475 | 0.0370 | 2.00E-01 | 0.0774 | 0.0132 | 4.63E-09 | 0 |
| rs10933809 | 3 | 104800350 | 24 | 104646815 | 104828166 | A | G | 0.4881 | ALCAM | intergenic | 0.0735 | 0.013 | 1.57E-08 | 0.0677 | 0.0271 | 1.24E-02 | 0.0319 | 0.0099 | 1.29E-03 | 1 |
| rs6437565 | 3 | 104801746 | 24 | 104646815 | 104828166 | A | G | 0.4881 | ALCAM | intergenic | -0.0732 | 0.013 | 1.81E-08 | -0.0676 | 0.0271 | 1.25E-02 | -0.0318 | 0.0099 | 1.33E-03 | 1 |
| rs1503085 | 3 | 104802096 | 24 | 104646815 | 104828166 | A | G | 0.4881 | ALCAM | intergenic | 0.0731 | 0.013 | 1.90E-08 | 0.0667 | 0.0271 | 1.38E-02 | 0.0318 | 0.0099 | 1.31E-03 | 1 |
| rs2654247 | 3 | 104805201 | 24 | 104646815 | 104828166 | A | G | 0.4602 | ALCAM | intergenic | -0.0934 | 0.0131 | 9.66E-13 | -0.0885 | 0.0272 | 1.13E-03 | -0.0464 | 0.0099 | 3.07E-06 | 1 |
| rs9881967 | 3 | 104821917 | 24 | 104646815 | 104828166 | A | G | 0.163 | ALCAM | intergenic | -0.1099 | 0.0181 | 1.15E-09 | -0.0352 | 0.0380 | 3.55E-01 | -0.0774 | 0.0136 | 1.22E-08 | 0 |
| rs2399023 | 3 | 104823350 | 24 | 104646815 | 104828166 | A | G | 0.3668 | ALCAM | intergenic | 0.0779 | 0.0136 | 9.74E-09 | 0.0584 | 0.0282 | 3.81E-02 | 0.0384 | 0.0103 | 2.04E-04 | 1 |
| rs7615032 | 3 | 104828081 | 24 | 104646815 | 104828166 | A | G | 0.163 | ALCAM | intergenic | -0.1118 | 0.0183 | 9.92E-10 | -0.0413 | 0.0384 | 2.81E-01 | -0.0747 | 0.0138 | 6.38E-08 | 0 |
| rs153152 | 5 | 59842971 | 25 | 59823118 | 60465365 | A | G | 0.4076 | PART1 | ncRNA_exonic | 0.0767 | 0.0134 | 1.12E-08 | 0.0857 | 0.0277 | 1.96E-03 | 0.0221 | 0.0103 | 3.11E-02 | 1 |
| rs248912 | 5 | 59844079 | 25 | 59823118 | 60465365 | T | C | 0.4085 | PART1 | downstream | -0.0771 | 0.0134 | 9.17E-09 | -0.0857 | 0.0277 | 1.95E-03 | -0.0220 | 0.0102 | 3.18E-02 | 1 |
| rs249070 | 5 | 59851527 | 25 | 59823118 | 60465365 | T | C | 0.4085 | PART1 | intergenic | 0.0767 | 0.0133 | 9.06E-09 | 0.0829 | 0.0275 | 2.59E-03 | 0.0207 | 0.0103 | 4.36E-02 | 1 |
| rs633628 | 5 | 59852636 | 25 | 59823118 | 60465365 | T | C | 0.4085 | PART1 | intergenic | -0.0769 | 0.0133 | 7.79E-09 | -0.0829 | 0.0275 | 2.59E-03 | -0.0218 | 0.0102 | 3.24E-02 | 1 |
| rs60148983 | 5 | 59855032 | 25 | 59823118 | 60465365 | C | G | 0.4085 | PART1 | intergenic | -0.0773 | 0.0134 | 8.25E-09 | -0.0828 | 0.0275 | 2.62E-03 | -0.0207 | 0.0103 | 4.54E-02 | 1 |
| rs3958943 | 5 | 59876749 | 25 | 59823118 | 60465365 | T | C | 0.4085 | DEPDC1B | intergenic | -0.0762 | 0.0132 | 8.71E-09 | -0.0843 | 0.0275 | 2.20E-03 | -0.0211 | 0.0101 | 3.66E-02 | 1 |
| rs55768631 | 5 | 59877191 | 25 | 59823118 | 60465365 | A | T | 0.4085 | DEPDC1B | intergenic | 0.0759 | 0.0133 | 1.02E-08 | 0.0836 | 0.0275 | 2.39E-03 | 0.0207 | 0.0101 | 4.03E-02 | 1 |
| rs56166952 | 5 | 59906395 | 25 | 59823118 | 60465365 | T | C | 0.4085 | DEPDC1B | intronic | 0.0764 | 0.0132 | 7.04E-09 | 0.0832 | 0.0274 | 2.44E-03 | 0.0208 | 0.0101 | 3.87E-02 | 1 |
| rs11951503 | 5 | 59918840 | 25 | 59823118 | 60465365 | A | G | 0.4085 | DEPDC1B | intronic | 0.0767 | 0.0132 | 6.35E-09 | 0.0819 | 0.0274 | 2.85E-03 | 0.0207 | 0.0101 | 3.99E-02 | 1 |
| rs10939855 | 5 | 59930271 | 25 | 59823118 | 60465365 | A | G | 0.4215 | DEPDC1B | intronic | -0.0838 | 0.0135 | 5.62E-10 | -0.0829 | 0.0275 | 2.57E-03 | -0.0211 | 0.0105 | 4.47E-02 | 1 |
| rs1379116 | 5 | 59942400 | 25 | 59823118 | 60465365 | A | G | 0.4085 | DEPDC1B | intronic | -0.077 | 0.0132 | 5.47E-09 | -0.0840 | 0.0274 | 2.20E-03 | -0.0218 | 0.0101 | 3.11E-02 | 1 |
| rs72755157 | 5 | 59947386 | 25 | 59823118 | 60465365 | A | C | 0.4095 | DEPDC1B | intronic | 0.0811 | 0.0135 | 1.98E-09 | 0.0849 | 0.0274 | 1.95E-03 | 0.0203 | 0.0104 | 5.20E-02 | 0 |
| rs4504343 | 5 | 59948395 | 25 | 59823118 | 60465365 | C | G | 0.4205 | DEPDC1B | intronic | 0.0764 | 0.0132 | 7.57E-09 | 0.0884 | 0.0274 | 1.27E-03 | NA | NA | NA | NA |
| rs56347749 | 5 | 59951824 | 25 | 59823118 | 60465365 | A | G | 0.4205 | DEPDC1B | intronic | 0.0761 | 0.0132 | 8.78E-09 | 0.0888 | 0.0274 | 1.21E-03 | 0.0219 | 0.0102 | 3.14E-02 | 1 |
| rs2198865 | 5 | 59954057 | 25 | 59823118 | 60465365 | T | C | 0.4095 | DEPDC1B | intronic | -0.0754 | 0.0132 | 1.14E-08 | -0.0858 | 0.0274 | 1.73E-03 | -0.0210 | 0.0101 | 3.84E-02 | 1 |
| rs59155887 | 5 | 59961543 | 25 | 59823118 | 60465365 | A | G | 0.4095 | DEPDC1B | intronic | -0.0756 | 0.0132 | 1.07E-08 | -0.0858 | 0.0274 | 1.73E-03 | -0.0208 | 0.0101 | 4.02E-02 | 1 |
| rs17387940 | 5 | 59967716 | 25 | 59823118 | 60465365 | A | C | 0.4095 | DEPDC1B | intronic | 0.074 | 0.0132 | 1.89E-08 | 0.0860 | 0.0274 | 1.69E-03 | 0.0214 | 0.0100 | 3.23E-02 | 1 |
| rs12517174 | 5 | 59969015 | 25 | 59823118 | 60465365 | T | C | 0.4095 | DEPDC1B | intronic | 0.074 | 0.0132 | 1.86E-08 | 0.0860 | 0.0274 | 1.69E-03 | 0.0214 | 0.0100 | 3.23E-02 | 1 |
| rs56726435 | 5 | 59975064 | 25 | 59823118 | 60465365 | T | C | 0.4095 | DEPDC1B | intronic | 0.0743 | 0.0132 | 1.63E-08 | 0.0853 | 0.0274 | 1.86E-03 | 0.0214 | 0.0100 | 3.26E-02 | 1 |
| rs11741754 | 5 | 59977058 | 25 | 59823118 | 60465365 | A | G | 0.4095 | DEPDC1B | intronic | -0.0746 | 0.0132 | 1.46E-08 | -0.0851 | 0.0274 | 1.89E-03 | -0.0213 | 0.0100 | 3.33E-02 | 1 |
| rs11955398 | 5 | 59981085 | 25 | 59823118 | 60465365 | A | C | 0.4095 | DEPDC1B | intronic | -0.0745 | 0.0132 | 1.51E-08 | -0.0847 | 0.0274 | 2.00E-03 | -0.0213 | 0.0100 | 3.31E-02 | 1 |
| rs10939858 | 5 | 59983481 | 25 | 59823118 | 60465365 | A | T | 0.4095 | DEPDC1B | intronic | -0.0766 | 0.0135 | 1.38E-08 | -0.0849 | 0.0274 | 1.95E-03 | -0.0211 | 0.0105 | 4.46E-02 | 1 |
| rs7704338 | 5 | 59986155 | 25 | 59823118 | 60465365 | T | C | 0.4553 | DEPDC1B | intronic | -0.0748 | 0.013 | 9.28E-09 | -0.0752 | 0.0270 | 5.39E-03 | -0.0218 | 0.0100 | 2.90E-02 | 1 |
| rs7719611 | 5 | 59988547 | 25 | 59823118 | 60465365 | A | G | 0.4553 | DEPDC1B | intronic | -0.0746 | 0.013 | 1.03E-08 | -0.0748 | 0.0270 | 5.68E-03 | -0.0217 | 0.0100 | 2.94E-02 | 1 |
| rs12517207 | 5 | 59993268 | 25 | 59823118 | 60465365 | T | C | 0.4095 | DEPDC1B | intronic | 0.0745 | 0.0132 | 1.56E-08 | 0.0860 | 0.0274 | 1.71E-03 | 0.0211 | 0.0100 | 3.53E-02 | 1 |
| rs716344 | 5 | 60012521 | 25 | 59823118 | 60465365 | C | G | 0.4066 | DEPDC1B | intergenic | -0.0799 | 0.0132 | 1.27E-09 | -0.0753 | 0.0274 | 5.98E-03 | -0.0214 | 0.0101 | 3.38E-02 | 1 |
| rs7709056 | 5 | 60024091 | 25 | 59823118 | 60465365 | A | G | 0.4076 | KRT8P31 | intergenic | -0.0796 | 0.0133 | 2.10E-09 | -0.0736 | 0.0274 | 7.20E-03 | -0.0199 | 0.0103 | 5.23E-02 | 0 |
| rs7381195 | 5 | 60030791 | 25 | 59823118 | 60465365 | A | T | 0.3638 | KRT8P31 | intergenic | 0.0866 | 0.014 | 5.27E-10 | 0.0750 | 0.0279 | 7.13E-03 | 0.0221 | 0.0107 | 4.00E-02 | 1 |
| rs7720233 | 5 | 60046832 | 25 | 59823118 | 60465365 | T | C | 0.4085 | ELOVL7 | downstream | 0.0763 | 0.0131 | 6.37E-09 | 0.0713 | 0.0274 | 9.23E-03 | 0.0232 | 0.0101 | 2.13E-02 | 1 |
| rs10043291 | 5 | 60058858 | 25 | 59823118 | 60465365 | T | C | 0.4095 | ELOVL7 | intronic | 0.0761 | 0.0131 | 6.96E-09 | 0.0716 | 0.0274 | 8.98E-03 | 0.0233 | 0.0100 | 2.03E-02 | 1 |
| rs4527549 | 5 | 60062745 | 25 | 59823118 | 60465365 | T | C | 0.4095 | ELOVL7 | intronic | -0.0768 | 0.0131 | 4.93E-09 | -0.0730 | 0.0274 | 7.79E-03 | -0.0233 | 0.0100 | 2.03E-02 | 1 |
| rs1460958 | 5 | 60063576 | 25 | 59823118 | 60465365 | A | G | 0.4105 | ELOVL7 | intronic | 0.0766 | 0.0131 | 5.45E-09 | 0.0724 | 0.0274 | 8.25E-03 | 0.0231 | 0.0100 | 2.15E-02 | 1 |
| rs1444237 | 5 | 60063953 | 25 | 59823118 | 60465365 | T | C | 0.4085 | ELOVL7 | intronic | 0.0767 | 0.0131 | 5.17E-09 | 0.0715 | 0.0274 | 9.05E-03 | 0.0226 | 0.0100 | 2.44E-02 | 1 |
| rs13158665 | 5 | 60069057 | 25 | 59823118 | 60465365 | T | C | 0.4394 | ELOVL7 | intronic | 0.0779 | 0.0132 | 3.20E-09 | 0.0820 | 0.0271 | 2.47E-03 | 0.0213 | 0.0101 | 3.56E-02 | 1 |
| rs1444240 | 5 | 60069934 | 25 | 59823118 | 60465365 | A | G | 0.4264 | ELOVL7 | intronic | 0.079 | 0.0132 | 2.13E-09 | 0.0753 | 0.0273 | 5.86E-03 | 0.0233 | 0.0102 | 2.18E-02 | 1 |
| rs6874699 | 5 | 60074000 | 25 | 59823118 | 60465365 | T | C | 0.4254 | ELOVL7 | intronic | -0.0788 | 0.0132 | 2.33E-09 | -0.0738 | 0.0273 | 6.91E-03 | -0.0236 | 0.0102 | 2.02E-02 | 1 |
| rs4353980 | 5 | 60075328 | 25 | 59823118 | 60465365 | T | C | 0.4264 | ELOVL7 | intronic | 0.0787 | 0.0132 | 2.42E-09 | 0.0736 | 0.0273 | 7.04E-03 | 0.0236 | 0.0102 | 2.04E-02 | 1 |
| rs10068474 | 5 | 60077912 | 25 | 59823118 | 60465365 | T | G | 0.4254 | ELOVL7 | intronic | -0.0784 | 0.0132 | 2.72E-09 | -0.0737 | 0.0273 | 6.95E-03 | -0.0236 | 0.0102 | 2.06E-02 | 1 |
| rs10062240 | 5 | 60078412 | 25 | 59823118 | 60465365 | T | C | 0.4254 | ELOVL7 | intronic | 0.0785 | 0.0132 | 2.67E-09 | 0.0737 | 0.0273 | 6.95E-03 | 0.0242 | 0.0102 | 1.72E-02 | 1 |
| rs4700391 | 5 | 60082641 | 25 | 59823118 | 60465365 | T | C | 0.4264 | ELOVL7 | intronic | -0.0821 | 0.0138 | 2.89E-09 | -0.0725 | 0.0273 | 7.98E-03 | -0.0186 | 0.0108 | 8.55E-02 | 0 |
| rs10072745 | 5 | 60083275 | 25 | 59823118 | 60465365 | A | C | 0.4642 | ELOVL7 | UTR5 | -0.0732 | 0.0131 | 2.20E-08 | -0.0961 | 0.0271 | 4.03E-04 | -0.0251 | 0.0100 | 1.25E-02 | 1 |
| rs6449500 | 5 | 60083903 | 25 | 59823118 | 60465365 | A | G | 0.4642 | ELOVL7 | intronic | 0.0731 | 0.0131 | 2.29E-08 | 0.0968 | 0.0271 | 3.67E-04 | 0.0252 | 0.0100 | 1.21E-02 | 1 |
| rs1870015 | 5 | 60085763 | 25 | 59823118 | 60465365 | A | G | 0.4642 | ELOVL7 | intronic | -0.0729 | 0.0131 | 2.57E-08 | -0.0968 | 0.0271 | 3.65E-04 | -0.0240 | 0.0101 | 1.71E-02 | 1 |
| rs12519587 | 5 | 60086900 | 25 | 59823118 | 60465365 | T | C | 0.4284 | ELOVL7 | intronic | -0.0771 | 0.0132 | 5.15E-09 | -0.0857 | 0.0273 | 1.70E-03 | -0.0226 | 0.0101 | 2.50E-02 | 1 |
| rs6872863 | 5 | 60088148 | 25 | 59823118 | 60465365 | A | G | 0.4672 | ELOVL7 | intronic | 0.0724 | 0.013 | 2.57E-08 | 0.0968 | 0.0271 | 3.64E-04 | 0.0238 | 0.0099 | 1.62E-02 | 1 |
| rs6873181 | 5 | 60088250 | 25 | 59823118 | 60465365 | T | C | 0.4672 | ELOVL7 | intronic | -0.0724 | 0.013 | 2.55E-08 | -0.0968 | 0.0271 | 3.64E-04 | -0.0238 | 0.0099 | 1.62E-02 | 1 |
| rs6894672 | 5 | 60088255 | 25 | 59823118 | 60465365 | T | C | 0.4672 | ELOVL7 | intronic | 0.0724 | 0.013 | 2.52E-08 | 0.0968 | 0.0271 | 3.64E-04 | 0.0238 | 0.0099 | 1.64E-02 | 1 |
| rs10057395 | 5 | 60088794 | 25 | 59823118 | 60465365 | A | G | 0.4463 | ELOVL7 | intronic | 0.0729 | 0.0131 | 2.61E-08 | 0.0835 | 0.0272 | 2.17E-03 | 0.0266 | 0.0100 | 8.14E-03 | 1 |
| rs921897 | 5 | 60089137 | 25 | 59823118 | 60465365 | A | G | 0.4642 | ELOVL7 | intronic | -0.0734 | 0.013 | 1.67E-08 | -0.0975 | 0.0271 | 3.32E-04 | -0.0237 | 0.0099 | 1.66E-02 | 1 |
| rs10076679 | 5 | 60090951 | 25 | 59823118 | 60465365 | C | G | 0.4642 | ELOVL7 | intronic | 0.0736 | 0.013 | 1.52E-08 | 0.0950 | 0.0271 | 4.68E-04 | NA | NA | NA | NA |
| rs10076742 | 5 | 60091046 | 25 | 59823118 | 60465365 | T | C | 0.4642 | ELOVL7 | intronic | -0.0738 | 0.013 | 1.40E-08 | -0.0946 | 0.0271 | 4.95E-04 | -0.0236 | 0.0099 | 1.69E-02 | 1 |
| rs6890201 | 5 | 60091192 | 25 | 59823118 | 60465365 | C | G | 0.4642 | ELOVL7 | intronic | 0.0738 | 0.013 | 1.40E-08 | 0.0946 | 0.0271 | 4.94E-04 | NA | NA | NA | NA |
| rs6875456 | 5 | 60091330 | 25 | 59823118 | 60465365 | T | C | 0.4642 | ELOVL7 | intronic | 0.0737 | 0.013 | 1.44E-08 | 0.0946 | 0.0271 | 4.94E-04 | 0.0236 | 0.0099 | 1.70E-02 | 1 |

| | | | | | | | | | | | Meta-GWAS | | | UKBB | | | ENIGMA* | | | |
|---|---|---|---|---|---|---|---|---|---|---|---|---|---|---|---|---|---|---|---|---|
| rsID | chr | pos (hg19) | Genomic Locus | start | end | A1 | A2 | MAF | Nearest Genes | Function | Beta | SE | P-value | Beta | SE | P-value | Beta | SE | P-value | Replicated** |
| rs6894750 | 5 | 60091556 | 25 | 59823118 | 60465365 | A | G | 0.4642 | ELOVL7 | intronic | 0.0737 | 0.013 | 1.43E-08 | 0.0946 | 0.0271 | 4.95E-04 | 0.0235 | 0.0099 | 1.79E-02 | 1 |
| rs6876335 | 5 | 60091818 | 25 | 59823118 | 60465365 | T | C | 0.4642 | ELOVL7 | intronic | 0.0739 | 0.013 | 1.29E-08 | 0.0946 | 0.0271 | 4.95E-04 | 0.0235 | 0.0099 | 1.75E-02 | 1 |
| rs6449501 | 5 | 60092687 | 25 | 59823118 | 60465365 | A | G | 0.4642 | ELOVL7 | intronic | -0.0739 | 0.013 | 1.33E-08 | -0.0949 | 0.0271 | 4.75E-04 | -0.0235 | 0.0099 | 1.75E-02 | 1 |
| rs4235481 | 5 | 60093387 | 25 | 59823118 | 60465365 | T | C | 0.4642 | ELOVL7 | intronic | -0.0739 | 0.013 | 1.32E-08 | -0.0951 | 0.0271 | 4.59E-04 | -0.0235 | 0.0099 | 1.74E-02 | 1 |
| rs4235482 | 5 | 60093612 | 25 | 59823118 | 60465365 | A | G | 0.4642 | ELOVL7 | intronic | 0.0738 | 0.013 | 1.37E-08 | 0.0950 | 0.0271 | 4.69E-04 | 0.0235 | 0.0099 | 1.75E-02 | 1 |
| rs57468227 | 5 | 60095004 | 25 | 59823118 | 60465365 | T | C | 0.3678 | ELOVL7 | intronic | 0.0801 | 0.0137 | 5.45E-09 | 0.0841 | 0.0278 | 2.53E-03 | 0.0213 | 0.0106 | 4.36E-02 | 1 |
| rs4482855 | 5 | 60096497 | 25 | 59823118 | 60465365 | T | C | 0.4652 | ELOVL7 | intronic | 0.0726 | 0.013 | 2.45E-08 | 0.0953 | 0.0272 | 4.57E-04 | 0.0236 | 0.0099 | 1.73E-02 | 1 |
| rs4700393 | 5 | 60098267 | 25 | 59823118 | 60465365 | A | G | 0.4602 | ELOVL7 | intronic | 0.0733 | 0.013 | 1.79E-08 | 0.0993 | 0.0272 | 2.56E-04 | 0.0238 | 0.0099 | 1.66E-02 | 1 |
| rs10077663 | 5 | 60103492 | 25 | 59823118 | 60465365 | A | G | 0.4652 | ELOVL7 | intronic | 0.0719 | 0.013 | 3.27E-08 | 0.0955 | 0.0272 | 4.40E-04 | 0.0238 | 0.0099 | 1.64E-02 | 1 |
| rs61160187 | 5 | 60111579 | 25 | 59823118 | 60465365 | A | G | 0.3678 | ELOVL7 | intronic | 0.0797 | 0.0138 | 7.04E-09 | 0.0834 | 0.0278 | 2.75E-03 | 0.0224 | 0.0106 | 3.39E-02 | 1 |
| rs4700394 | 5 | 60117202 | 25 | 59823118 | 60465365 | A | T | 0.4642 | ELOVL7 | intronic | 0.0711 | 0.013 | 4.74E-08 | 0.0906 | 0.0271 | 8.43E-04 | NA | NA | NA | NA |
| rs6893642 | 5 | 60117723 | 25 | 59823118 | 60465365 | T | C | 0.4642 | ELOVL7 | intronic | 0.0714 | 0.013 | 4.24E-08 | 0.0898 | 0.0271 | 9.38E-04 | 0.0260 | 0.0099 | 8.89E-03 | 1 |
| rs1445293 | 5 | 60119737 | 25 | 59823118 | 60465365 | A | G | 0.4642 | ELOVL7 | intronic | 0.0713 | 0.013 | 4.57E-08 | 0.0906 | 0.0271 | 8.49E-04 | 0.0259 | 0.0099 | 9.06E-03 | 1 |
| rs1807017 | 5 | 60127724 | 25 | 59823118 | 60465365 | T | C | 0.4254 | ELOVL7 | intronic | 0.0788 | 0.0132 | 2.44E-09 | 0.0846 | 0.0274 | 2.05E-03 | 0.0261 | 0.0101 | 9.80E-03 | 1 |
| rs62372102 | 5 | 60139881 | 25 | 59823118 | 60465365 | A | G | 0.4553 | ELOVL7 | intronic | 0.0796 | 0.0131 | 1.43E-09 | 0.0792 | 0.0272 | 3.57E-03 | 0.0249 | 0.0101 | 1.35E-02 | 1 |
| rs12654306 | 5 | 60184856 | 25 | 59823118 | 60465365 | A | C | 0.4275 | ERCC8 | intronic | 0.074 | 0.0133 | 2.94E-08 | 0.0872 | 0.0276 | 1.60E-03 | 0.0308 | 0.0103 | 2.67E-03 | 1 |
| rs7722373 | 5 | 60247815 | 25 | 59823118 | 60465365 | A | G | 0.4612 | NDUFAF2 | intronic | -0.079 | 0.0131 | 1.61E-09 | -0.0816 | 0.0272 | 2.67E-03 | -0.0299 | 0.0101 | 3.17E-03 | 1 |
| rs17419290 | 5 | 60307904 | 25 | 59823118 | 60465365 | A | G | 0.4056 | NDUFAF2 | intronic | 0.0731 | 0.0133 | 4.15E-08 | 0.0909 | 0.0275 | 9.29E-04 | 0.0363 | 0.0102 | 3.71E-04 | 1 |
| rs10939879 | 5 | 60315823 | 25 | 59823118 | 60465365 | T | C | 0.4135 | NDUFAF2 | intronic | 0.0727 | 0.0133 | 4.12E-08 | 0.0973 | 0.0274 | 3.80E-04 | 0.0395 | 0.0101 | 9.14E-05 | 1 |
| rs76470478 | 6 | 126613946 | 26 | 126525715 | 127369230 | T | C | 0.02783 | PPP1R14BP5 | intergenic | -0.2206 | 0.0373 | 3.43E-09 | -0.1571 | 0.0726 | 3.05E-02 | -0.2034 | 0.0301 | 1.36E-11 | 1 |
| rs117892760 | 6 | 126666648 | 26 | 126525715 | 127369230 | T | C | 0.02386 | CENPW | intronic | -0.2224 | 0.0391 | 1.27E-08 | -0.2087 | 0.0741 | 4.85E-03 | -0.2376 | 0.0335 | 1.29E-12 | 1 |
| rs1578060 | 6 | 126712247 | 26 | 126525715 | 127369230 | C | G | 0.4533 | CENPW | intergenic | -0.0744 | 0.0131 | 1.37E-08 | -0.0805 | 0.0273 | 3.21E-03 | NA | NA | NA | NA |
| rs1591805 | 6 | 126717064 | 26 | 126525715 | 127369230 | A | G | 0.4911 | CENPW | intergenic | 0.0793 | 0.0132 | 2.01E-09 | 0.0763 | 0.0271 | 4.92E-03 | 0.0532 | 0.0102 | 2.06E-07 | 1 |
| rs9398805 | 6 | 126726295 | 26 | 126525715 | 127369230 | T | C | 0.4543 | CENPW | intergenic | -0.0749 | 0.0131 | 1.04E-08 | -0.0772 | 0.0273 | 4.69E-03 | -0.0550 | 0.0100 | 3.63E-08 | 1 |
| rs4897179 | 6 | 126727930 | 26 | 126525715 | 127369230 | A | G | 0.4543 | CENPW | intergenic | -0.0785 | 0.0133 | 3.29E-09 | -0.0774 | 0.0273 | 4.62E-03 | -0.0592 | 0.0102 | 6.23E-09 | 1 |
| rs4897180 | 6 | 126727950 | 26 | 126525715 | 127369230 | A | T | 0.4543 | CENPW | intergenic | 0.0751 | 0.0132 | 1.19E-08 | 0.0776 | 0.0273 | 4.51E-03 | NA | NA | NA | NA |
| rs75343521 | 6 | 126732511 | 26 | 126525715 | 127369230 | T | C | 0.02684 | CENPW | intergenic | -0.2458 | 0.0373 | 4.25E-11 | -0.1573 | 0.0702 | 2.50E-02 | -0.1964 | 0.0308 | 1.93E-10 | 1 |
| rs9321065 | 6 | 126744087 | 26 | 126525715 | 127369230 | A | G | 0.4543 | MIR588 | intergenic | -0.0754 | 0.0131 | 8.62E-09 | -0.0776 | 0.0273 | 4.49E-03 | -0.0552 | 0.0100 | 3.42E-08 | 1 |
| rs2039735 | 6 | 126745076 | 26 | 126525715 | 127369230 | T | C | 0.4543 | MIR588 | intergenic | 0.0785 | 0.0132 | 3.11E-09 | 0.0779 | 0.0273 | 4.35E-03 | 0.0578 | 0.0102 | 1.27E-08 | 1 |
| rs139214174 | 6 | 126745633 | 26 | 126525715 | 127369230 | A | C | 0.02684 | MIR588 | intergenic | -0.2502 | 0.0375 | 2.38E-11 | -0.1568 | 0.0708 | 2.67E-02 | -0.2022 | 0.0313 | 1.12E-10 | 1 |
| rs1538172 | 6 | 126748493 | 26 | 126525715 | 127369230 | A | G | 0.4543 | MIR588 | intergenic | 0.0757 | 0.0131 | 7.24E-09 | 0.0794 | 0.0273 | 3.65E-03 | 0.0550 | 0.0100 | 3.59E-08 | 1 |
| rs9401881 | 6 | 126751921 | 26 | 126525715 | 127369230 | T | C | 0.4543 | MIR588 | intergenic | -0.0755 | 0.0131 | 7.95E-09 | -0.0774 | 0.0273 | 4.59E-03 | -0.0550 | 0.0100 | 3.53E-08 | 1 |
| rs4565329 | 6 | 126752798 | 26 | 126525715 | 127369230 | T | C | 0.4543 | MIR588 | intergenic | -0.0752 | 0.0131 | 9.15E-09 | -0.0773 | 0.0273 | 4.66E-03 | -0.0550 | 0.0100 | 3.52E-08 | 1 |
| rs1538170 | 6 | 126752874 | 26 | 126525715 | 127369230 | T | C | 0.4563 | MIR588 | intergenic | -0.0755 | 0.0131 | 7.69E-09 | -0.0773 | 0.0273 | 4.66E-03 | -0.0550 | 0.0100 | 3.54E-08 | 1 |
| rs1538171 | 6 | 126752884 | 26 | 126525715 | 127369230 | C | G | 0.4563 | MIR588 | intergenic | 0.0754 | 0.0131 | 8.26E-09 | 0.0773 | 0.0273 | 4.66E-03 | NA | NA | NA | NA |
| rs4897181 | 6 | 126753564 | 26 | 126525715 | 127369230 | T | C | 0.4543 | MIR588 | intergenic | -0.0758 | 0.0131 | 6.96E-09 | -0.0773 | 0.0273 | 4.62E-03 | -0.0550 | 0.0100 | 3.51E-08 | 1 |
| rs9401882 | 6 | 126755055 | 26 | 126525715 | 127369230 | A | G | 0.4543 | MIR588 | intergenic | 0.0759 | 0.0131 | 6.70E-09 | 0.0773 | 0.0273 | 4.65E-03 | 0.0550 | 0.0100 | 3.69E-08 | 1 |
| rs4897182 | 6 | 126755642 | 26 | 126525715 | 127369230 | T | G | 0.4553 | MIR588 | intergenic | 0.076 | 0.0131 | 6.33E-09 | 0.0772 | 0.0273 | 4.70E-03 | 0.0547 | 0.0100 | 4.18E-08 | 1 |
| rs9375439 | 6 | 126758540 | 26 | 126525715 | 127369230 | C | G | 0.4543 | MIR588 | intergenic | 0.076 | 0.0131 | 6.19E-09 | 0.0774 | 0.0273 | 4.58E-03 | NA | NA | NA | NA |
| rs9398808 | 6 | 126758791 | 26 | 126525715 | 127369230 | T | G | 0.4543 | MIR588 | intergenic | -0.0758 | 0.0131 | 6.65E-09 | -0.0774 | 0.0273 | 4.58E-03 | -0.0550 | 0.0100 | 3.66E-08 | 1 |
| rs9385399 | 6 | 126759033 | 26 | 126525715 | 127369230 | T | G | 0.4543 | MIR588 | intergenic | -0.0758 | 0.0131 | 6.89E-09 | -0.0769 | 0.0273 | 4.87E-03 | -0.0550 | 0.0100 | 3.64E-08 | 1 |
| rs1415671 | 6 | 126759587 | 26 | 126525715 | 127369230 | T | G | 0.4543 | MIR588 | intergenic | 0.0759 | 0.0131 | 6.40E-09 | 0.0771 | 0.0273 | 4.73E-03 | 0.0550 | 0.0100 | 3.60E-08 | 1 |
| rs2184968 | 6 | 126760994 | 26 | 126525715 | 127369230 | T | C | 0.4543 | MIR588 | intergenic | 0.0748 | 0.0131 | 1.13E-08 | 0.0778 | 0.0273 | 4.44E-03 | 0.0553 | 0.0100 | 3.18E-08 | 1 |
| rs2152876 | 6 | 126761228 | 26 | 126525715 | 127369230 | A | G | 0.4543 | MIR588 | intergenic | -0.0757 | 0.0131 | 7.17E-09 | -0.0780 | 0.0273 | 4.26E-03 | -0.0550 | 0.0100 | 3.78E-08 | 1 |
| rs9385400 | 6 | 126764190 | 26 | 126525715 | 127369230 | T | G | 0.4543 | MIR588 | intergenic | 0.0758 | 0.0131 | 6.96E-09 | 0.0773 | 0.0273 | 4.62E-03 | 0.0547 | 0.0100 | 4.34E-08 | 1 |
| rs1361107 | 6 | 126767511 | 26 | 126525715 | 127369230 | A | C | 0.4583 | MIR588 | intergenic | 0.0758 | 0.0131 | 7.23E-09 | 0.0764 | 0.0273 | 5.18E-03 | 0.0549 | 0.0100 | 4.14E-08 | 1 |
| rs1361108 | 6 | 126767600 | 26 | 126525715 | 127369230 | T | C | 0.4543 | MIR588 | intergenic | -0.0761 | 0.0131 | 5.93E-09 | -0.0772 | 0.0273 | 4.68E-03 | -0.0549 | 0.0100 | 4.03E-08 | 1 |
| rs1337735 | 6 | 126771024 | 26 | 126525715 | 127369230 | A | T | 0.4543 | MIR588 | intergenic | -0.0753 | 0.0131 | 8.87E-09 | -0.0776 | 0.0273 | 4.48E-03 | NA | NA | NA | NA |
| rs1361109 | 6 | 126771143 | 26 | 126525715 | 127369230 | T | C | 0.4543 | MIR588 | intergenic | -0.0759 | 0.0131 | 6.77E-09 | -0.0776 | 0.0273 | 4.46E-03 | -0.0551 | 0.0100 | 3.54E-08 | 1 |
| rs1572569 | 6 | 126771687 | 26 | 126525715 | 127369230 | A | G | 0.4543 | MIR588 | intergenic | -0.0758 | 0.0131 | 7.11E-09 | -0.0778 | 0.0273 | 4.37E-03 | -0.0549 | 0.0100 | 3.94E-08 | 1 |
| rs2184967 | 6 | 126773487 | 26 | 126525715 | 127369230 | T | C | 0.4533 | MIR588 | intergenic | -0.0749 | 0.0131 | 1.05E-08 | -0.0779 | 0.0273 | 4.33E-03 | -0.0548 | 0.0100 | 4.37E-08 | 1 |
| rs4559102 | 6 | 126773580 | 26 | 126525715 | 127369230 | A | G | 0.4533 | MIR588 | intergenic | -0.0752 | 0.0131 | 9.05E-09 | -0.0784 | 0.0273 | 4.06E-03 | -0.0548 | 0.0100 | 4.30E-08 | 1 |
| rs9388494 | 6 | 126776094 | 26 | 126525715 | 127369230 | T | G | 0.4543 | MIR588 | intergenic | -0.074 | 0.0131 | 1.62E-08 | -0.0780 | 0.0273 | 4.27E-03 | -0.0540 | 0.0100 | 7.39E-08 | 1 |
| rs9375441 | 6 | 126778600 | 26 | 126525715 | 127369230 | A | G | 0.4533 | MIR588 | intergenic | -0.0754 | 0.0131 | 8.36E-09 | -0.0786 | 0.0273 | 3.99E-03 | -0.0547 | 0.0100 | 4.52E-08 | 1 |
| rs4895808 | 6 | 126781434 | 26 | 126525715 | 127369230 | T | C | 0.4543 | MIR588 | intergenic | -0.0752 | 0.0131 | 9.47E-09 | -0.0779 | 0.0273 | 4.30E-03 | -0.0547 | 0.0100 | 4.45E-08 | 1 |
| rs9388495 | 6 | 126782457 | 26 | 126525715 | 127369230 | A | G | 0.4553 | MIR588 | intergenic | -0.0756 | 0.0131 | 7.92E-09 | -0.0784 | 0.0273 | 4.06E-03 | -0.0546 | 0.0100 | 4.97E-08 | 1 |
| rs9388496 | 6 | 126785062 | 26 | 126525715 | 127369230 | A | G | 0.4543 | MIR588 | intergenic | 0.0752 | 0.0131 | 9.24E-09 | 0.0776 | 0.0273 | 4.43E-03 | 0.0543 | 0.0100 | 5.65E-08 | 1 |
| rs1844594 | 6 | 126788107 | 26 | 126525715 | 127369230 | A | G | 0.4553 | MIR588 | intergenic | -0.0755 | 0.0131 | 8.19E-09 | -0.0779 | 0.0273 | 4.30E-03 | -0.0542 | 0.0100 | 6.26E-08 | 1 |
| rs9398809 | 6 | 126789144 | 26 | 126525715 | 127369230 | T | C | 0.4543 | MIR588 | intergenic | -0.0755 | 0.0131 | 8.11E-09 | -0.0779 | 0.0273 | 4.30E-03 | -0.0543 | 0.0100 | 5.55E-08 | 1 |
| rs9385401 | 6 | 126789472 | 26 | 126525715 | 127369230 | T | C | 0.4543 | MIR588 | intergenic | -0.075 | 0.0131 | 1.01E-08 | -0.0779 | 0.0273 | 4.30E-03 | -0.0542 | 0.0100 | 5.85E-08 | 1 |
| rs9401883 | 6 | 126797111 | 26 | 126525715 | 127369230 | A | G | 0.4553 | MIR588 | intergenic | 0.0749 | 0.0131 | 1.05E-08 | 0.0782 | 0.0273 | 4.17E-03 | 0.0543 | 0.0100 | 5.43E-08 | 1 |
| rs1159619 | 6 | 126801144 | 26 | 126525715 | 127369230 | A | C | 0.4543 | MIR588 | intergenic | -0.0744 | 0.0131 | 1.33E-08 | -0.0788 | 0.0273 | 3.98E-03 | -0.0541 | 0.0100 | 6.15E-08 | 1 |
| rs9375442 | 6 | 126803137 | 26 | 126525715 | 127369230 | A | C | 0.4543 | MIR588 | intergenic | 0.076 | 0.0131 | 6.69E-09 | 0.0778 | 0.0273 | 4.35E-03 | 0.0550 | 0.0100 | 3.92E-08 | 1 |
| rs2050644 | 6 | 126805255 | 26 | 126525715 | 127369230 | A | G | 0.4543 | MIR588 | upstream | 0.0747 | 0.0131 | 1.15E-08 | 0.0781 | 0.0273 | 4.23E-03 | 0.0541 | 0.0100 | 5.99E-08 | 1 |
| rs4422634 | 6 | 126806676 | 26 | 126525715 | 127369230 | T | C | 0.4543 | MIR588 | downstream | 0.0751 | 0.0131 | 9.54E-09 | 0.0786 | 0.0273 | 3.95E-03 | 0.0535 | 0.0100 | 8.37E-08 | 1 |
| rs1120786 | 6 | 126808637 | 26 | 126525715 | 127369230 | T | G | 0.4543 | MIR588 | intergenic | 0.0745 | 0.0131 | 1.32E-08 | 0.0786 | 0.0273 | 3.98E-03 | 0.0544 | 0.0100 | 5.77E-08 | 1 |
| rs9372839 | 6 | 126809870 | 26 | 126525715 | 127369230 | A | G | 0.4543 | MIR588 | intergenic | -0.0747 | 0.0131 | 1.14E-08 | -0.0788 | 0.0273 | 3.90E-03 | -0.0545 | 0.0100 | 5.09E-08 | 1 |
| rs2326451 | 6 | 126811928 | 26 | 126525715 | 127369230 | A | T | 0.4533 | MIR588 | intergenic | 0.0747 | 0.0131 | 1.18E-08 | 0.0765 | 0.0273 | 5.14E-03 | NA | NA | NA | NA |
| rs9398810 | 6 | 126815604 | 26 | 126525715 | 127369230 | A | C | 0.4543 | MIR588 | intergenic | -0.0748 | 0.0131 | 1.09E-08 | -0.0784 | 0.0273 | 4.09E-03 | -0.0544 | 0.0100 | 5.21E-08 | 1 |
| rs4418209 | 6 | 126819699 | 26 | 126525715 | 127369230 | T | G | 0.4543 | MIR588 | intergenic | 0.0748 | 0.0131 | 1.08E-08 | 0.0790 | 0.0273 | 3.81E-03 | 0.0544 | 0.0100 | 5.10E-08 | 1 |
| rs9372840 | 6 | 126822635 | 26 | 126525715 | 127369230 | A | C | 0.4543 | MIR588 | intergenic | 0.0755 | 0.0131 | 7.97E-09 | 0.0799 | 0.0273 | 3.41E-03 | 0.0549 | 0.0100 | 4.15E-08 | 1 |
| rs9401885 | 6 | 126825837 | 26 | 126525715 | 127369230 | C | G | 0.4543 | MIR588 | intergenic | 0.075 | 0.0131 | 9.80E-09 | 0.0800 | 0.0273 | 3.36E-03 | NA | NA | NA | NA |
| rs2130603 | 6 | 126826819 | 26 | 126525715 | 127369230 | A | C | 0.4543 | MIR588 | intergenic | 0.075 | 0.0131 | 1.04E-08 | 0.0799 | 0.0273 | 3.40E-03 | 0.0553 | 0.0100 | 3.50E-08 | 1 |
| rs7738135 | 6 | 126829981 | 26 | 126525715 | 127369230 | A | G | 0.4543 | MIR588 | intergenic | -0.0749 | 0.0131 | 1.05E-08 | -0.0800 | 0.0273 | 3.35E-03 | -0.0550 | 0.0100 | 4.29E-08 | 1 |
| rs4053271 | 6 | 126830512 | 26 | 126525715 | 127369230 | T | C | 0.4543 | MIR588 | intergenic | 0.0748 | 0.0131 | 1.12E-08 | 0.0800 | 0.0273 | 3.35E-03 | 0.0551 | 0.0100 | 4.08E-08 | 1 |
| rs1490387 | 6 | 126834083 | 26 | 126525715 | 127369230 | A | C | 0.4543 | MIR588 | intergenic | -0.0751 | 0.0131 | 9.48E-09 | -0.0801 | 0.0273 | 3.35E-03 | -0.0551 | 0.0100 | 3.80E-08 | 1 |
| rs9375446 | 6 | 126835015 | 26 | 126525715 | 127369230 | A | G | 0.4543 | MIR588 | intergenic | -0.075 | 0.0131 | 9.61E-09 | -0.0800 | 0.0273 | 3.35E-03 | -0.0551 | 0.0100 | 3.83E-08 | 1 |
| rs1490388 | 6 | 126835655 | 26 | 126525715 | 127369230 | T | C | 0.4543 | MIR588 | intergenic | -0.075 | 0.0131 | 9.76E-09 | -0.0803 | 0.0273 | 3.27E-03 | -0.0551 | 0.0100 | 3.78E-08 | 1 |
| rs1907067 | 6 | 126837663 | 26 | 126525715 | 127369230 | A | C | 0.4543 | MIR588 | intergenic | 0.0749 | 0.0131 | 1.05E-08 | 0.0806 | 0.0273 | 3.15E-03 | 0.0551 | 0.0100 | 3.74E-08 | 1 |
| rs1602278 | 6 | 126838605 | 26 | 126525715 | 127369230 | A | C | 0.4553 | MIR588 | intergenic | 0.0754 | 0.0131 | 8.34E-09 | 0.0809 | 0.0273 | 3.05E-03 | 0.0552 | 0.0100 | 3.73E-08 | 1 |
| rs1337736 | 6 | 126845380 | 26 | 126525715 | 127369230 | A | G | 0.1998 | MIR588 | intergenic | -0.1244 | 0.0164 | 3.40E-14 | -0.1115 | 0.0328 | 6.78E-04 | -0.0822 | 0.0126 | 6.62E-11 | 1 |
| rs9375447 | 6 | 126874091 | 26 | 126525715 | 127369230 | A | G | 0.4821 | RNU6-200P | intergenic | 0.071 | 0.013 | 4.53E-08 | 0.0813 | 0.0272 | 2.79E-03 | 0.0563 | 0.0100 | 1.54E-08 | 1 |
| rs1490385 | 6 | 126875521 | 26 | 126525715 | 127369230 | T | C | 0.4821 | RNU6-200P | intergenic | -0.0712 | 0.013 | 4.35E-08 | -0.0820 | 0.0272 | 2.56E-03 | -0.0561 | 0.0100 | 2.10E-08 | 1 |
| rs9401890 | 6 | 126881295 | 26 | 126525715 | 127369230 | A | G | 0.4831 | RNU6-200P | intergenic | 0.0718 | 0.013 | 3.72E-08 | 0.0815 | 0.0272 | 2.72E-03 | 0.0565 | 0.0101 | 2.08E-08 | 1 |
| rs9401891 | 6 | 126894135 | 26 | 126525715 | 127369230 | T | G | 0.4811 | RNU6-200P | intergenic | -0.0724 | 0.013 | 2.76E-08 | -0.0827 | 0.0272 | 2.38E-03 | -0.0577 | 0.0100 | 8.20E-09 | 1 |
| rs9401892 | 6 | 126895368 | 26 | 126525715 | 127369230 | A | G | 0.4811 | RNU6-200P | intergenic | -0.0708 | 0.013 | 4.90E-08 | -0.0832 | 0.0272 | 2.22E-03 | -0.0555 | 0.0099 | 2.34E-08 | 1 |
| rs4621655 | 6 | 126896107 | 26 | 126525715 | 127369230 | A | G | 0.4811 | RNU6-200P | intergenic | -0.0712 | 0.013 | 4.21E-08 | -0.0832 | 0.0272 | 2.22E-03 | -0.0556 | 0.0099 | 2.32E-08 | 1 |
| rs4621656 | 6 | 126896168 | 26 | 126525715 | 127369230 | C | G | 0.4811 | RNU6-200P | intergenic | -0.0709 | 0.013 | 4.66E-08 | -0.0832 | 0.0272 | 2.20E-03 | NA | NA | NA | NA |
| rs4621657 | 6 | 126896227 | 26 | 126525715 | 127369230 | A | G | 0.4811 | RNU6-200P | intergenic | -0.0711 | 0.013 | 4.33E-08 | -0.0832 | 0.0272 | 2.22E-03 | -0.0556 | 0.0099 | 2.29E-08 | 1 |
| rs1844593 | 6 | 126897827 | 26 | 126525715 | 127369230 | A | G | 0.4821 | RNU6-200P | intergenic | -0.0713 | 0.013 | 3.96E-08 | -0.0832 | 0.0272 | 2.22E-03 | -0.0553 | 0.0100 | 2.76E-08 | 1 |
| rs3861455 | 6 | 126897969 | 26 | 126525715 | 127369230 | A | C | 0.4811 | RNU6-200P | intergenic | 0.0711 | 0.013 | 4.35E-08 | 0.0832 | 0.0272 | 2.21E-03 | 0.0554 | 0.0099 | 2.52E-08 | 1 |

| | | | | | | | | | | | Meta-GWAS | | | UKBB | | | ENIGMA* | | | |
|---|---|---|---|---|---|---|---|---|---|---|---|---|---|---|---|---|---|---|---|---|
| rsID | chr | pos (hg19) | Genomic Locus | start | end | A1 | A2 | MAF | Nearest Genes | Function | Beta | SE | P-value | Beta | SE | P-value | Beta | SE | P-value | Replicated** |
| rs3861457 | 6 | 126902245 | 26 | 126525715 | 127369230 | T | C | 0.4821 | RNU6-200P | intergenic | 0.0717 | 0.013 | 3.36E-08 | 0.0832 | 0.0272 | 2.22E-03 | 0.0551 | 0.0100 | 3.13E-08 | 1 |
| rs9388501 | 6 | 126903011 | 26 | 126525715 | 127369230 | T | C | 0.4811 | RNU6-200P | intergenic | 0.072 | 0.013 | 3.00E-08 | 0.0832 | 0.0272 | 2.21E-03 | 0.0549 | 0.0100 | 3.38E-08 | 1 |
| rs4427021 | 6 | 126938446 | 26 | 126525715 | 127369230 | T | C | 0.1998 | VIMP1 | intergenic | -0.1219 | 0.0161 | 3.64E-14 | -0.1114 | 0.0327 | 6.48E-04 | -0.0796 | 0.0124 | 1.24E-10 | 1 |
| rs6569466 | 6 | 126961319 | 26 | 126525715 | 127369230 | T | C | 0.2008 | PRELID1P1 | intergenic | -0.1202 | 0.0159 | 3.92E-14 | -0.1149 | 0.0326 | 4.29E-04 | -0.0800 | 0.0122 | 5.65E-11 | 1 |
| rs74580701 | 6 | 127000881 | 26 | 126525715 | 127369230 | A | G | 0.02684 | RPS4XP9 | intergenic | 0.2316 | 0.0382 | 1.30E-09 | 0.1248 | 0.0683 | 6.75E-02 | 0.2288 | 0.0361 | 2.34E-10 | 0 |
| rs9375452 | 6 | 127014862 | 26 | 126525715 | 127369230 | T | C | 0.2028 | RPS4XP9 | intergenic | -0.1227 | 0.0159 | 1.09E-14 | -0.1145 | 0.0326 | 4.41E-04 | -0.0803 | 0.0122 | 4.05E-11 | 1 |
| rs853983 | 6 | 127046153 | 26 | 126525715 | 127369230 | A | G | 0.4901 | RPS4XP9 | intergenic | 0.0712 | 0.013 | 4.46E-08 | 0.0730 | 0.0272 | 7.24E-03 | 0.0590 | 0.0100 | 4.29E-09 | 1 |
| rs1262555 | 6 | 127052816 | 26 | 126525715 | 127369230 | A | G | 0.4911 | RPS4XP9 | intergenic | 0.0742 | 0.0132 | 2.16E-08 | 0.0735 | 0.0272 | 6.91E-03 | 0.0600 | 0.0104 | 7.55E-09 | 1 |
| rs190958130 | 6 | 127052822 | 26 | 126525715 | 127369230 | C | G | 0.02684 | RPS4XP9 | intergenic | 0.2397 | 0.0391 | 8.47E-10 | 0.1569 | 0.0718 | 2.89E-02 | 0.2119 | 0.0368 | 8.66E-09 | 1 |
| rs9401903 | 6 | 127055771 | 26 | 126525715 | 127369230 | T | C | 0.2137 | RPS4XP9 | intergenic | -0.1227 | 0.0156 | 4.31E-15 | -0.0999 | 0.0322 | 1.91E-03 | -0.0821 | 0.0120 | 8.13E-12 | 1 |
| rs79636188 | 6 | 127078181 | 26 | 126525715 | 127369230 | T | C | 0.02684 | RPS4XP9 | intergenic | 0.2358 | 0.0385 | 8.67E-10 | 0.1417 | 0.0697 | 4.21E-02 | 0.2069 | 0.0370 | 2.18E-08 | 1 |
| rs1262551 | 6 | 127078211 | 26 | 126525715 | 127369230 | T | C | 0.4901 | RPS4XP9 | intergenic | 0.0714 | 0.0131 | 4.78E-08 | 0.0728 | 0.0272 | 7.46E-03 | 0.0627 | 0.0102 | 7.41E-10 | 1 |
| rs1262552 | 6 | 127078752 | 26 | 126525715 | 127369230 | T | C | 0.4911 | RPS4XP9 | intergenic | 0.0714 | 0.0131 | 4.83E-08 | 0.0722 | 0.0272 | 7.94E-03 | 0.0628 | 0.0102 | 6.98E-10 | 1 |
| rs1343222 | 6 | 127081190 | 26 | 126525715 | 127369230 | T | C | 0.2147 | RPS4XP9 | intergenic | 0.1214 | 0.0155 | 4.24E-15 | 0.0975 | 0.0321 | 2.43E-03 | 0.0802 | 0.0118 | 9.95E-12 | 1 |
| rs9385403 | 6 | 127083941 | 26 | 126525715 | 127369230 | T | C | 0.2147 | RPS4XP9 | intergenic | -0.1216 | 0.0155 | 3.95E-15 | -0.0973 | 0.0322 | 2.48E-03 | -0.0799 | 0.0118 | 1.15E-11 | 1 |
| rs13212467 | 6 | 127086138 | 26 | 126525715 | 127369230 | A | G | 0.2147 | RPS4XP9 | intergenic | -0.1213 | 0.0155 | 5.42E-15 | -0.0984 | 0.0322 | 2.22E-03 | -0.0802 | 0.0118 | 1.02E-11 | 1 |
| rs72971190 | 6 | 127088303 | 26 | 126525715 | 127369230 | A | G | 0.2157 | RPS4XP9 | intergenic | 0.1174 | 0.0154 | 2.85E-14 | 0.0974 | 0.0323 | 2.57E-03 | 0.0820 | 0.0116 | 1.96E-12 | 1 |
| rs72971192 | 6 | 127088318 | 26 | 126525715 | 127369230 | T | C | 0.2157 | RPS4XP9 | intergenic | 0.118 | 0.0154 | 2.14E-14 | 0.0985 | 0.0323 | 2.32E-03 | 0.0819 | 0.0116 | 1.97E-12 | 1 |
| rs4897189 | 6 | 127088550 | 26 | 126525715 | 127369230 | A | T | 0.2167 | RPS4XP9 | intergenic | 0.1183 | 0.0154 | 1.56E-14 | 0.0993 | 0.0321 | 1.98E-03 | 0.0819 | 0.0116 | 1.99E-12 | 1 |
| rs9385404 | 6 | 127088591 | 26 | 126525715 | 127369230 | A | G | 0.2167 | RPS4XP9 | intergenic | 0.1186 | 0.0154 | 1.33E-14 | 0.0981 | 0.0321 | 2.24E-03 | 0.0820 | 0.0116 | 1.78E-12 | 1 |
| rs2223739 | 6 | 127089401 | 26 | 126525715 | 127369230 | T | C | 0.2157 | RPS4XP9 | intergenic | -0.1189 | 0.0154 | 1.17E-14 | -0.0994 | 0.0321 | 1.97E-03 | -0.0819 | 0.0116 | 1.98E-12 | 1 |
| rs76611955 | 6 | 127090025 | 26 | 126525715 | 127369230 | A | T | 0.02684 | RPS4XP9 | intergenic | 0.2416 | 0.0392 | 6.92E-10 | 0.1307 | 0.0698 | 6.10E-02 | 0.2160 | 0.0369 | 4.92E-09 | 0 |
| rs4895815 | 6 | 127090469 | 26 | 126525715 | 127369230 | A | G | 0.2157 | RPS4XP9 | intergenic | 0.1186 | 0.0154 | 1.33E-14 | 0.0977 | 0.0322 | 2.43E-03 | 0.0817 | 0.0116 | 2.22E-12 | 1 |
| rs17054064 | 6 | 127091354 | 26 | 126525715 | 127369230 | T | C | 0.2157 | RPS4XP9 | intergenic | -0.119 | 0.0154 | 1.09E-14 | -0.0995 | 0.0321 | 1.95E-03 | -0.0815 | 0.0116 | 2.35E-12 | 1 |
| rs4895816 | 6 | 127094587 | 26 | 126525715 | 127369230 | A | C | 0.2157 | RPS4XP9 | intergenic | 0.1191 | 0.0154 | 9.88E-15 | 0.1001 | 0.0321 | 1.85E-03 | 0.0818 | 0.0116 | 1.95E-12 | 1 |
| rs4895817 | 6 | 127094898 | 26 | 126525715 | 127369230 | T | C | 0.2157 | RPS4XP9 | intergenic | 0.1191 | 0.0154 | 1.02E-14 | 0.1001 | 0.0321 | 1.83E-03 | 0.0821 | 0.0116 | 1.61E-12 | 1 |
| rs9388517 | 6 | 127095666 | 26 | 126525715 | 127369230 | A | G | 0.2157 | RPS4XP9 | intergenic | 0.1192 | 0.0154 | 9.85E-15 | 0.0996 | 0.0322 | 1.99E-03 | 0.0822 | 0.0116 | 1.56E-12 | 1 |
| rs9401907 | 6 | 127096181 | 26 | 126525715 | 127369230 | T | C | 0.2167 | RPS4XP9 | intergenic | 0.1189 | 0.0154 | 1.14E-14 | 0.1007 | 0.0322 | 1.78E-03 | 0.0821 | 0.0116 | 1.68E-12 | 1 |
| rs983741 | 6 | 127096527 | 26 | 126525715 | 127369230 | A | C | 0.2157 | RPS4XP9 | intergenic | -0.1198 | 0.0154 | 7.63E-15 | -0.1004 | 0.0323 | 1.85E-03 | -0.0826 | 0.0116 | 1.28E-12 | 1 |
| rs9375460 | 6 | 127149538 | 26 | 126525715 | 127369230 | T | C | 0.2237 | RPS4XP9 | intergenic | -0.1254 | 0.016 | 4.14E-15 | -0.1109 | 0.0324 | 6.14E-04 | -0.0865 | 0.0124 | 3.05E-12 | 1 |
| rs77598707 | 6 | 127176330 | 26 | 126525715 | 127369230 | T | C | 0.02485 | RPS4XP9 | intergenic | -0.2513 | 0.0396 | 2.10E-10 | -0.1423 | 0.0716 | 4.68E-02 | -0.2015 | 0.0359 | 1.98E-08 | 1 |
| rs77636716 | 6 | 127179329 | 26 | 126525715 | 127369230 | T | G | 0.02485 | RPS4XP9 | intergenic | -0.2529 | 0.0396 | 1.70E-10 | -0.1417 | 0.0717 | 4.83E-02 | -0.2089 | 0.0363 | 8.63E-09 | 1 |
| rs185867463 | 6 | 127182732 | 26 | 126525715 | 127369230 | C | G | 0.02485 | RPS4XP9 | intergenic | -0.2496 | 0.0404 | 6.48E-10 | -0.1476 | 0.0717 | 3.96E-02 | -0.2001 | 0.0384 | 1.85E-07 | 1 |
| rs138299249 | 6 | 127183470 | 26 | 126525715 | 127369230 | A | G | 0.1968 | RPS4XP9 | intergenic | 0.0998 | 0.0175 | 1.25E-08 | 0.0934 | 0.0348 | 7.26E-03 | 0.0795 | 0.0142 | 2.23E-08 | 1 |
| rs9375476 | 6 | 127185801 | 26 | 126525715 | 127369230 | A | G | 0.2217 | RPS4XP9 | intergenic | -0.1253 | 0.0164 | 2.32E-14 | -0.1103 | 0.0324 | 6.69E-04 | -0.0865 | 0.0131 | 3.91E-11 | 1 |
| rs79398770 | 6 | 127277488 | 26 | 126525715 | 127369230 | A | G | 0.02584 | RSPO3 | intergenic | 0.2545 | 0.0406 | 3.63E-10 | 0.1522 | 0.0723 | 3.53E-02 | NA | NA | NA | NA |
| rs957960 | 7 | 18877408 | 27 | 18869552 | 18933411 | A | C | 0.3638 | HDAC9 | intronic | 0.0794 | 0.0136 | 5.15E-09 | 0.0989 | 0.0282 | 4.51E-04 | 0.0641 | 0.0104 | 6.29E-10 | 1 |
| rs12536836 | 7 | 18881075 | 27 | 18869552 | 18933411 | T | C | 0.3936 | HDAC9 | intronic | 0.0783 | 0.0135 | 6.31E-09 | 0.1095 | 0.0278 | 8.40E-05 | 0.0633 | 0.0103 | 9.38E-10 | 1 |
| rs6461386 | 7 | 18883690 | 27 | 18869552 | 18933411 | A | G | 0.3658 | HDAC9 | intronic | 0.0843 | 0.0139 | 1.23E-09 | 0.1019 | 0.0287 | 3.89E-04 | 0.0662 | 0.0106 | 4.86E-10 | 1 |
| rs13245206 | 7 | 18891259 | 27 | 18869552 | 18933411 | A | G | 0.3807 | HDAC9 | intronic | 0.0741 | 0.0135 | 3.89E-08 | 0.0914 | 0.0281 | 1.16E-03 | 0.0574 | 0.0103 | 2.38E-08 | 1 |
| rs12699995 | 7 | 18897321 | 27 | 18869552 | 18933411 | T | C | 0.4175 | HDAC9 | intronic | 0.0768 | 0.0134 | 1.03E-08 | 0.1114 | 0.0278 | 6.16E-05 | 0.0627 | 0.0102 | 7.84E-10 | 1 |
| rs12699996 | 7 | 18897323 | 27 | 18869552 | 18933411 | T | G | 0.4195 | HDAC9 | intronic | -0.0797 | 0.0135 | 3.54E-09 | -0.1113 | 0.0278 | 6.27E-05 | -0.0660 | 0.0105 | 2.81E-10 | 1 |
| rs6461387 | 7 | 18898305 | 27 | 18869552 | 18933411 | A | G | 0.3917 | HDAC9 | intronic | 0.0784 | 0.0134 | 5.05E-09 | 0.1107 | 0.0279 | 7.39E-05 | 0.0630 | 0.0102 | 7.21E-10 | 1 |
| rs756854 | 7 | 18903015 | 27 | 18869552 | 18933411 | T | C | 0.3976 | HDAC9 | intronic | 0.0804 | 0.0135 | 2.29E-09 | 0.1080 | 0.0279 | 1.08E-04 | 0.0622 | 0.0103 | 1.35E-09 | 1 |
| rs59987684 | 7 | 18903517 | 27 | 18869552 | 18933411 | A | T | 0.3936 | HDAC9 | intronic | -0.0781 | 0.0136 | 8.69E-09 | -0.1091 | 0.0280 | 9.67E-05 | -0.0581 | 0.0104 | 2.43E-08 | 1 |
| rs55761404 | 7 | 18903580 | 27 | 18869552 | 18933411 | A | G | 0.3917 | HDAC9 | intronic | -0.081 | 0.0136 | 2.47E-09 | -0.1070 | 0.0280 | 1.34E-04 | -0.0587 | 0.0104 | 1.78E-08 | 1 |
| rs34854568 | 7 | 18903818 | 27 | 18869552 | 18933411 | A | G | 0.3827 | HDAC9 | intronic | 0.0857 | 0.0135 | 2.21E-10 | 0.1037 | 0.0279 | 2.02E-04 | 0.0661 | 0.0103 | 1.41E-10 | 1 |
| rs12699998 | 7 | 18904175 | 27 | 18869552 | 18933411 | A | G | 0.3847 | HDAC9 | intronic | -0.0845 | 0.0135 | 4.18E-10 | -0.1090 | 0.0279 | 9.23E-05 | -0.0658 | 0.0103 | 1.67E-10 | 1 |
| rs12699999 | 7 | 18904264 | 27 | 18869552 | 18933411 | A | G | 0.3738 | HDAC9 | intronic | -0.0811 | 0.0136 | 2.38E-09 | -0.1115 | 0.0280 | 7.04E-05 | -0.0627 | 0.0104 | 1.51E-09 | 1 |
| rs12700000 | 7 | 18904337 | 27 | 18869552 | 18933411 | T | C | 0.3231 | HDAC9 | intronic | -0.0792 | 0.014 | 1.38E-08 | -0.0837 | 0.0289 | 3.75E-03 | -0.0571 | 0.0106 | 7.92E-08 | 1 |
| rs12700001 | 7 | 18904400 | 27 | 18869552 | 18933411 | C | G | 0.3917 | HDAC9 | intronic | -0.0871 | 0.0136 | 1.33E-10 | -0.1038 | 0.0278 | 1.90E-04 | -0.0662 | 0.0103 | 1.32E-10 | 1 |
| rs12700002 | 7 | 18905450 | 27 | 18869552 | 18933411 | T | C | 0.4175 | HDAC9 | intronic | -0.0816 | 0.0134 | 1.13E-09 | -0.1085 | 0.0276 | 8.58E-05 | -0.0660 | 0.0102 | 1.10E-10 | 1 |
| rs12700003 | 7 | 18905866 | 27 | 18869552 | 18933411 | T | C | 0.4085 | HDAC9 | intronic | 0.0796 | 0.0134 | 2.67E-09 | 0.1186 | 0.0276 | 1.73E-05 | 0.0652 | 0.0102 | 1.71E-10 | 1 |
| rs6461389 | 7 | 18907030 | 27 | 18869552 | 18933411 | T | C | 0.3579 | HDAC9 | intronic | 0.0759 | 0.0137 | 2.80E-08 | 0.0918 | 0.0283 | 1.17E-03 | 0.0558 | 0.0104 | 8.67E-08 | 1 |
| rs12530943 | 7 | 18907748 | 27 | 18869552 | 18933411 | C | G | 0.3519 | HDAC9 | intronic | -0.075 | 0.0137 | 4.21E-08 | -0.0942 | 0.0283 | 8.90E-04 | -0.0567 | 0.0104 | 5.51E-08 | 1 |
| rs12530969 | 7 | 18907849 | 27 | 18869552 | 18933411 | C | G | 0.3539 | HDAC9 | intronic | -0.0755 | 0.0137 | 3.37E-08 | -0.0916 | 0.0283 | 1.21E-03 | -0.0557 | 0.0104 | 9.42E-08 | 1 |
| rs12536081 | 7 | 18907930 | 27 | 18869552 | 18933411 | A | C | 0.3598 | HDAC9 | intronic | -0.075 | 0.0137 | 3.90E-08 | -0.0945 | 0.0281 | 7.87E-04 | -0.0566 | 0.0104 | 5.60E-08 | 1 |
| rs6973029 | 7 | 18908850 | 27 | 18869552 | 18933411 | T | C | 0.3519 | HDAC9 | intronic | 0.0753 | 0.0137 | 3.80E-08 | 0.0945 | 0.0283 | 8.56E-04 | 0.0634 | 0.0104 | 1.27E-09 | 1 |
| rs6461390 | 7 | 18910232 | 27 | 18869552 | 18933411 | A | G | 0.3857 | HDAC9 | intronic | -0.0817 | 0.0135 | 1.35E-09 | -0.1056 | 0.0278 | 1.48E-04 | -0.0661 | 0.0103 | 1.21E-10 | 1 |
| rs6951144 | 7 | 18911789 | 27 | 18869552 | 18933411 | C | G | 0.3807 | HDAC9 | intronic | -0.0765 | 0.0136 | 1.86E-08 | -0.0952 | 0.0280 | 6.81E-04 | -0.0593 | 0.0104 | 1.05E-08 | 1 |
| rs6951745 | 7 | 18911957 | 27 | 18869552 | 18933411 | A | G | 0.3628 | HDAC9 | intronic | 0.0775 | 0.0136 | 1.25E-08 | 0.0889 | 0.0282 | 1.63E-03 | 0.0599 | 0.0104 | 7.92E-09 | 1 |
| rs6461391 | 7 | 18912777 | 27 | 18869552 | 18933411 | T | C | 0.3827 | HDAC9 | intronic | 0.082 | 0.0135 | 1.39E-09 | 0.1032 | 0.0280 | 2.26E-04 | 0.0667 | 0.0103 | 8.62E-11 | 1 |
| rs6461392 | 7 | 18912802 | 27 | 18869552 | 18933411 | A | G | 0.3628 | HDAC9 | intronic | 0.0775 | 0.0136 | 1.33E-08 | 0.0883 | 0.0283 | 1.78E-03 | 0.0601 | 0.0104 | 7.02E-09 | 1 |
| rs10237149 | 7 | 18915426 | 27 | 18869552 | 18933411 | A | G | 0.3807 | HDAC9 | intronic | 0.0796 | 0.0135 | 4.26E-09 | 0.1086 | 0.0281 | 1.12E-04 | 0.0723 | 0.0104 | 3.27E-12 | 1 |
| rs10237366 | 7 | 18915559 | 27 | 18869552 | 18933411 | C | G | 0.3827 | HDAC9 | intronic | 0.0815 | 0.0136 | 1.95E-09 | 0.1010 | 0.0281 | 3.33E-04 | 0.0711 | 0.0104 | 6.88E-12 | 1 |
| rs10237280 | 7 | 18915666 | 27 | 18869552 | 18933411 | T | C | 0.3847 | HDAC9 | intronic | 0.0814 | 0.0136 | 1.99E-09 | 0.0987 | 0.0281 | 4.54E-04 | 0.0713 | 0.0104 | 6.29E-12 | 1 |
| rs13233322 | 7 | 18915701 | 27 | 18869552 | 18933411 | A | C | 0.3767 | HDAC9 | intronic | 0.0768 | 0.0136 | 1.81E-08 | 0.1061 | 0.0283 | 1.78E-04 | 0.0674 | 0.0104 | 1.07E-10 | 1 |
| rs10486314 | 7 | 18915874 | 27 | 18869552 | 18933411 | A | G | 0.3767 | HDAC9 | intronic | -0.0769 | 0.0136 | 1.72E-08 | -0.1071 | 0.0282 | 1.48E-04 | -0.0669 | 0.0105 | 1.56E-10 | 1 |
| rs10270210 | 7 | 18920210 | 27 | 18869552 | 18933411 | T | C | 0.4354 | HDAC9 | intronic | -0.0745 | 0.0136 | 4.72E-08 | -0.1172 | 0.0279 | 2.64E-05 | -0.0667 | 0.0104 | 1.58E-10 | 1 |
| rs12338443 | 9 | 98147278 | 28 | 98147278 | 98279801 | T | C | 0.3231 | RP11-435O5.7 | intergenic | -0.0788 | 0.0142 | 2.67E-08 | -0.0494 | 0.0296 | 9.47E-02 | -0.0518 | 0.0109 | 2.06E-06 | 0 |
| rs7860361 | 9 | 98147804 | 28 | 98147278 | 98279801 | T | C | 0.3231 | RP11-435O5.7 | intergenic | -0.0803 | 0.0141 | 1.34E-08 | -0.0523 | 0.0296 | 7.70E-02 | -0.0525 | 0.0109 | 1.37E-06 | 0 |
| rs35422011 | 9 | 98149355 | 28 | 98147278 | 98279801 | A | T | 0.3221 | RP11-435O5.7 | intergenic | -0.085 | 0.0154 | 3.13E-08 | -0.0578 | 0.0294 | 4.90E-02 | -0.0528 | 0.0110 | 1.51E-06 | 1 |
| rs34392622 | 9 | 98150349 | 28 | 98147278 | 98279801 | T | C | 0.3221 | RP11-435O5.7 | intergenic | -0.0802 | 0.0142 | 1.50E-08 | -0.0527 | 0.0296 | 7.51E-02 | -0.0523 | 0.0109 | 1.53E-06 | 0 |
| rs10993531 | 9 | 98152242 | 28 | 98147278 | 98279801 | T | C | 0.326 | RP11-435O5.7 | intergenic | 0.0785 | 0.0142 | 3.01E-08 | 0.0422 | 0.0295 | 1.53E-01 | 0.0528 | 0.0109 | 1.25E-06 | 0 |
| rs10993532 | 9 | 98152282 | 28 | 98147278 | 98279801 | T | C | 0.325 | RP11-435O5.7 | intergenic | 0.0787 | 0.0142 | 2.72E-08 | 0.0449 | 0.0295 | 1.28E-01 | 0.0526 | 0.0109 | 1.38E-06 | 0 |
| rs13289815 | 9 | 98160000 | 28 | 98147278 | 98279801 | T | C | 0.3221 | RP11-435O5.7 | intergenic | -0.0805 | 0.0142 | 1.48E-08 | -0.0519 | 0.0296 | 8.00E-02 | -0.0527 | 0.0109 | 1.40E-06 | 0 |
| rs2202303 | 9 | 98163443 | 28 | 98147278 | 98279801 | T | C | 0.3221 | RP11-435O5.7 | intergenic | 0.0804 | 0.0142 | 1.53E-08 | 0.0527 | 0.0297 | 7.54E-02 | 0.0521 | 0.0109 | 1.76E-06 | 0 |
| rs7854702 | 9 | 98164782 | 28 | 98147278 | 98279801 | T | C | 0.3241 | RP11-435O5.7 | intergenic | -0.0802 | 0.0142 | 1.75E-08 | -0.0533 | 0.0296 | 7.20E-02 | -0.0520 | 0.0109 | 1.95E-06 | 0 |
| rs28573122 | 9 | 98190335 | 28 | 98147278 | 98279801 | A | G | 0.3718 | RP11-435O5.2 | intergenic | 0.0785 | 0.0144 | 4.53E-08 | 0.0399 | 0.0292 | 1.73E-01 | 0.0479 | 0.0111 | 1.75E-05 | 0 |
| rs28710957 | 9 | 98192383 | 28 | 98147278 | 98279801 | A | T | 0.337 | RP11-435O5.2 | intergenic | -0.0868 | 0.0154 | 1.82E-08 | -0.0623 | 0.0303 | 3.99E-02 | -0.0547 | 0.0124 | 9.80E-06 | 1 |
| rs9632916 | 9 | 98213728 | 28 | 98147278 | 98279801 | C | G | 0.2227 | PTCH1 | UTR3 | 0.1023 | 0.016 | 1.83E-10 | 0.1035 | 0.0326 | 1.50E-03 | 0.0667 | 0.0122 | 4.04E-08 | 1 |
| rs9632897 | 9 | 98214367 | 28 | 98147278 | 98279801 | A | G | 0.2237 | PTCH1 | UTR3 | 0.1005 | 0.016 | 3.09E-10 | 0.1004 | 0.0323 | 1.88E-03 | 0.0665 | 0.0121 | 4.36E-08 | 1 |
| rs9632898 | 9 | 98214758 | 28 | 98147278 | 98279801 | T | C | 0.2237 | PTCH1 | UTR3 | -0.1 | 0.0159 | 3.56E-10 | -0.0981 | 0.0323 | 2.38E-03 | -0.0666 | 0.0121 | 4.00E-08 | 1 |
| rs2134721 | 9 | 98215306 | 28 | 98147278 | 98279801 | T | C | 0.2237 | PTCH1 | UTR3 | -0.1006 | 0.0159 | 2.68E-10 | -0.0969 | 0.0323 | 2.68E-03 | -0.0665 | 0.0121 | 4.31E-08 | 1 |
| rs17368876 | 9 | 98222665 | 28 | 98147278 | 98279801 | T | C | 0.2237 | PTCH1 | intronic | -0.1015 | 0.0159 | 1.85E-10 | -0.0962 | 0.0321 | 2.76E-03 | -0.0659 | 0.0121 | 4.65E-08 | 1 |
| rs28571635 | 9 | 98225658 | 28 | 98147278 | 98279801 | T | C | 0.2068 | PTCH1 | intronic | -0.1001 | 0.0164 | 1.04E-09 | -0.0886 | 0.0330 | 7.23E-03 | -0.0672 | 0.0125 | 8.00E-08 | 1 |
| rs28394554 | 9 | 98228225 | 28 | 98147278 | 98279801 | T | C | 0.2237 | PTCH1 | intronic | -0.099 | 0.0158 | 3.94E-10 | -0.0974 | 0.0320 | 2.38E-03 | -0.0654 | 0.0120 | 4.75E-08 | 1 |
| rs2274693 | 9 | 98229297 | 28 | 98147278 | 98279801 | A | G | 0.2237 | PTCH1 | intronic | -0.0994 | 0.0158 | 2.96E-10 | -0.0952 | 0.0320 | 2.93E-03 | -0.0645 | 0.0120 | 6.86E-08 | 1 |
| rs28702657 | 9 | 98231908 | 28 | 98147278 | 98279801 | A | G | 0.2237 | PTCH1 | intronic | -0.0991 | 0.0158 | 3.14E-10 | -0.0948 | 0.0320 | 3.05E-03 | -0.0642 | 0.0119 | 7.74E-08 | 1 |
| rs28442014 | 9 | 98234563 | 28 | 98147278 | 98279801 | T | C | 0.2237 | PTCH1 | intronic | -0.0992 | 0.0158 | 3.28E-10 | -0.0971 | 0.0320 | 2.41E-03 | -0.0647 | 0.0119 | 5.88E-08 | 1 |
| rs62558340 | 9 | 98234903 | 28 | 98147278 | 98279801 | T | C | 0.2237 | PTCH1 | intronic | -0.0989 | 0.0158 | 3.55E-10 | -0.0979 | 0.0320 | 2.21E-03 | -0.0645 | 0.0119 | 6.36E-08 | 1 |

| | | | | | | | | | | | Meta-GWAS | | | UKBB | | | ENIGMA* | | | |
|---|---|---|---|---|---|---|---|---|---|---|---|---|---|---|---|---|---|---|---|---|
| rsID | chr | pos (hg19) | Genomic Locus | start | end | A1 | A2 | MAF | Nearest Genes | Function | Beta | SE | P-value | Beta | SE | P-value | Beta | SE | P-value | Replicated** |
| rs2066836 | 9 | 98238358 | 28 | 98147278 | 98279801 | A | G | 0.1948 | PTCH1 | exonic | -0.0925 | 0.0164 | 1.77E-08 | -0.0818 | 0.0332 | 1.39E-02 | -0.0555 | 0.0125 | 8.36E-06 | 1 |
| rs2274694 | 9 | 98242112 | 28 | 98147278 | 98279801 | A | G | 0.2237 | PTCH1 | intronic | 0.0985 | 0.0158 | 4.08E-10 | 0.0957 | 0.0320 | 2.78E-03 | 0.0644 | 0.0119 | 5.87E-08 | 1 |
| rs2297087 | 9 | 98242925 | 28 | 98147278 | 98279801 | A | G | 0.2237 | PTCH1 | intronic | 0.0981 | 0.0158 | 4.95E-10 | 0.0945 | 0.0320 | 3.14E-03 | 0.0646 | 0.0119 | 5.42E-08 | 1 |
| rs28488553 | 9 | 98245855 | 28 | 98147278 | 98279801 | T | C | 0.2247 | PTCH1 | intronic | 0.0989 | 0.0157 | 3.28E-10 | 0.0917 | 0.0320 | 4.18E-03 | 0.0640 | 0.0119 | 7.22E-08 | 1 |
| rs17369383 | 9 | 98245974 | 28 | 98147278 | 98279801 | A | G | 0.2247 | PTCH1 | intronic | 0.099 | 0.0157 | 3.14E-10 | 0.0916 | 0.0320 | 4.22E-03 | 0.0644 | 0.0119 | 5.86E-08 | 1 |
| rs28504341 | 9 | 98249963 | 28 | 98147278 | 98279801 | T | C | 0.2256 | PTCH1 | intronic | -0.0983 | 0.0158 | 5.50E-10 | -0.0969 | 0.0321 | 2.57E-03 | -0.0647 | 0.0119 | 5.60E-08 | 1 |
| rs28411363 | 9 | 98252313 | 28 | 98147278 | 98279801 | T | C | 0.2237 | PTCH1 | intronic | 0.099 | 0.0157 | 3.10E-10 | 0.0905 | 0.0321 | 4.82E-03 | 0.0638 | 0.0119 | 7.91E-08 | 1 |
| rs28831479 | 9 | 98254526 | 28 | 98147278 | 98279801 | A | C | 0.2475 | PTCH1 | intronic | -0.0888 | 0.0152 | 5.07E-09 | -0.0864 | 0.0312 | 5.62E-03 | -0.0573 | 0.0115 | 6.97E-07 | 1 |
| rs28633576 | 9 | 98257305 | 28 | 98147278 | 98279801 | T | C | 0.2008 | PTCH1:RP11-43 | ncRNA_exonic | -0.1026 | 0.0164 | 4.31E-10 | -0.0923 | 0.0334 | 5.72E-03 | -0.0647 | 0.0123 | 1.33E-07 | 1 |
| rs28468571 | 9 | 98262735 | 28 | 98147278 | 98279801 | A | G | 0.2107 | PTCH1 | intronic | 0.0938 | 0.0158 | 3.05E-09 | 0.1045 | 0.0326 | 1.36E-03 | 0.0631 | 0.0119 | 1.13E-07 | 1 |
| rs3758301 | 9 | 98272464 | 28 | 98147278 | 98279801 | T | C | 0.2078 | PTCH1 | intronic | 0.093 | 0.016 | 6.70E-09 | 0.1089 | 0.0328 | 8.98E-04 | 0.0640 | 0.0120 | 9.42E-08 | 1 |
| rs28412122 | 9 | 98273791 | 28 | 98147278 | 98279801 | A | G | 0.2097 | PTCH1 | intronic | -0.0932 | 0.0159 | 5.05E-09 | -0.1059 | 0.0328 | 1.24E-03 | -0.0648 | 0.0120 | 6.27E-08 | 1 |
| rs1877456 | 9 | 98275097 | 28 | 98147278 | 98279801 | C | G | 0.2097 | PTCH1 | intronic | 0.0931 | 0.0159 | 5.14E-09 | 0.1051 | 0.0328 | 1.35E-03 | 0.0646 | 0.0120 | 6.83E-08 | 1 |
| rs28485705 | 9 | 98276105 | 28 | 98147278 | 98279801 | C | G | 0.2058 | PTCH1 | intronic | 0.0962 | 0.0162 | 2.91E-09 | 0.1146 | 0.0330 | 5.16E-04 | 0.0660 | 0.0121 | 4.81E-08 | 1 |
| rs28410513 | 9 | 98276371 | 28 | 98147278 | 98279801 | T | G | 0.2097 | PTCH1:RP11-43 | ncRNA_exonic | -0.0922 | 0.016 | 7.61E-09 | -0.1066 | 0.0329 | 1.18E-03 | -0.0648 | 0.0120 | 6.55E-08 | 1 |
| rs28491365 | 9 | 98276374 | 28 | 98147278 | 98279801 | A | T | 0.2097 | PTCH1:RP11-43 | ncRNA_exonic | -0.0922 | 0.016 | 7.62E-09 | -0.1066 | 0.0329 | 1.18E-03 | -0.0647 | 0.0120 | 6.85E-08 | 1 |
| rs149658356 | 10 | 102245664 | 29 | 102245664 | 102371946 | T | C | 0.0169 | SEC31B | downstream | 0.3262 | 0.0598 | 4.85E-08 | 0.1960 | 0.1043 | 6.02E-02 | NA | NA | NA | NA |
| rs118158982 | 10 | 102324683 | 29 | 102245664 | 102371946 | A | G | 0.01988 | HIF1AN | intergenic | -0.3315 | 0.0584 | 1.38E-08 | -0.1571 | 0.0999 | 1.16E-01 | NA | NA | NA | NA |
| rs78638570 | 10 | 102335663 | 29 | 102245664 | 102371946 | A | G | 0.02584 | HIF1AN | intergenic | -0.3154 | 0.0528 | 2.32E-09 | -0.1413 | 0.0934 | 1.30E-01 | NA | NA | NA | NA |
| rs138740906 | 10 | 102366639 | 29 | 102245664 | 102371946 | T | C | 0.02485 | HIF1AN | intergenic | -0.3267 | 0.0542 | 1.70E-09 | -0.1332 | 0.0926 | 1.50E-01 | NA | NA | NA | NA |
| rs4439488 | 11 | 30804996 | 30 | 30798255 | 31020084 | C | G | 0.2952 | DCDC1 | intergenic | -0.0982 | 0.0151 | 7.71E-11 | -0.0713 | 0.0291 | 1.45E-02 | -0.0716 | 0.0113 | 2.60E-10 | 1 |
| rs614943 | 11 | 30806623 | 30 | 30798255 | 31020084 | T | C | 0.3877 | DCDC1 | intergenic | 0.0771 | 0.0141 | 4.76E-08 | 0.0457 | 0.0278 | 1.00E-01 | 0.0648 | 0.0109 | 2.64E-09 | 0 |
| rs2021807 | 11 | 30806998 | 30 | 30798255 | 31020084 | T | C | 0.3022 | DCDC1 | intergenic | 0.0932 | 0.0149 | 4.42E-10 | 0.0747 | 0.0291 | 1.02E-02 | 0.0703 | 0.0114 | 6.06E-10 | 1 |
| rs7113110 | 11 | 30807171 | 30 | 30798255 | 31020084 | T | C | 0.3022 | DCDC1 | intergenic | -0.0932 | 0.0149 | 4.43E-10 | -0.0748 | 0.0291 | 1.01E-02 | -0.0704 | 0.0114 | 5.71E-10 | 1 |
| rs911268 | 11 | 30809653 | 30 | 30798255 | 31020084 | T | C | 0.3022 | DCDC1 | intergenic | 0.0927 | 0.0149 | 4.81E-10 | 0.0746 | 0.0290 | 1.01E-02 | 0.0699 | 0.0113 | 6.83E-10 | 1 |
| rs927224 | 11 | 30810665 | 30 | 30798255 | 31020084 | T | C | 0.2932 | DCDC1 | intergenic | 0.0956 | 0.015 | 1.74E-10 | 0.0735 | 0.0291 | 1.15E-02 | 0.0704 | 0.0112 | 3.82E-10 | 1 |
| rs7930532 | 11 | 30811294 | 30 | 30798255 | 31020084 | A | G | 0.2952 | DCDC1 | intergenic | -0.0954 | 0.015 | 1.76E-10 | -0.0724 | 0.0291 | 1.27E-02 | -0.0710 | 0.0112 | 2.64E-10 | 1 |
| rs35194615 | 11 | 30813303 | 30 | 30798255 | 31020084 | T | C | 0.2952 | DCDC1 | intergenic | -0.0975 | 0.0149 | 5.46E-11 | -0.0736 | 0.0291 | 1.14E-02 | -0.0708 | 0.0112 | 2.45E-10 | 1 |
| rs7128775 | 11 | 30816001 | 30 | 30798255 | 31020084 | A | T | 0.3887 | DCDC1 | intergenic | 0.077 | 0.014 | 3.50E-08 | 0.0469 | 0.0277 | 9.10E-02 | 0.0632 | 0.0106 | 2.64E-09 | 0 |
| rs10835708 | 11 | 30816258 | 30 | 30798255 | 31020084 | T | C | 0.3926 | DCDC1 | intergenic | 0.0766 | 0.014 | 4.42E-08 | 0.0451 | 0.0278 | 1.04E-01 | 0.0630 | 0.0107 | 3.34E-09 | 0 |
| rs2022130 | 11 | 30819811 | 30 | 30798255 | 31020084 | T | C | 0.2962 | DCDC1 | intergenic | 0.0981 | 0.0148 | 3.74E-11 | 0.0726 | 0.0290 | 1.24E-02 | 0.0700 | 0.0111 | 2.77E-10 | 1 |
| rs507969 | 11 | 30830612 | 30 | 30798255 | 31020084 | A | C | 0.2962 | DCDC1 | intergenic | 0.0974 | 0.0147 | 3.77E-11 | 0.0722 | 0.0290 | 1.28E-02 | 0.0685 | 0.0110 | 4.66E-10 | 1 |
| rs2145795 | 11 | 30831065 | 30 | 30798255 | 31020084 | A | G | 0.3926 | DCDC1 | intergenic | -0.0761 | 0.0138 | 3.70E-08 | -0.0445 | 0.0277 | 1.08E-01 | -0.0606 | 0.0104 | 6.51E-09 | 0 |
| rs7950105 | 11 | 30837058 | 30 | 30798255 | 31020084 | T | C | 0.3956 | DCDC1 | intergenic | -0.0736 | 0.0135 | 4.85E-08 | -0.0473 | 0.0277 | 8.79E-02 | -0.0574 | 0.0102 | 1.88E-08 | 0 |
| rs535845 | 11 | 30839228 | 30 | 30798255 | 31020084 | A | C | 0.2962 | DCDC1 | intergenic | 0.0945 | 0.0142 | 3.19E-11 | 0.0733 | 0.0289 | 1.14E-02 | 0.0649 | 0.0107 | 1.19E-09 | 1 |
| rs532202 | 11 | 30839635 | 30 | 30798255 | 31020084 | T | C | 0.2962 | DCDC1 | intergenic | 0.0943 | 0.0142 | 3.33E-11 | 0.0736 | 0.0289 | 1.10E-02 | 0.0650 | 0.0107 | 1.15E-09 | 1 |
| rs113026 | 11 | 30846842 | 30 | 30798255 | 31020084 | A | G | 0.2962 | DCDC1 | intergenic | 0.094 | 0.0142 | 4.20E-11 | 0.0726 | 0.0289 | 1.20E-02 | 0.0650 | 0.0107 | 1.22E-09 | 1 |
| NA | 11 | 30853147 | 30 | 30798255 | 31020084 | C | G | 0.2962 | DCDC1 | intronic | -0.1057 | 0.0194 | 4.87E-08 | -0.0726 | 0.0288 | 1.17E-02 | -0.0663 | 0.0108 | 9.05E-10 | 1 |
| rs604672 | 11 | 30853226 | 30 | 30798255 | 31020084 | C | G | 0.2962 | DCDC1 | intronic | -0.0934 | 0.0142 | 4.56E-11 | -0.0723 | 0.0288 | 1.21E-02 | -0.0642 | 0.0106 | 1.54E-09 | 1 |
| rs2021806 | 11 | 30853618 | 30 | 30798255 | 31020084 | A | G | 0.2992 | DCDC1 | intronic | -0.0924 | 0.0142 | 6.58E-11 | -0.0730 | 0.0288 | 1.12E-02 | -0.0646 | 0.0106 | 1.19E-09 | 1 |
| rs686901 | 11 | 30854334 | 30 | 30798255 | 31020084 | C | G | 0.2952 | DCDC1 | intronic | 0.093 | 0.0142 | 5.77E-11 | 0.0735 | 0.0288 | 1.08E-02 | 0.0643 | 0.0106 | 1.43E-09 | 1 |
| rs1015541 | 11 | 30854905 | 30 | 30798255 | 31020084 | T | G | 0.2992 | DCDC1 | intronic | -0.092 | 0.0142 | 7.99E-11 | -0.0730 | 0.0288 | 1.12E-02 | -0.0645 | 0.0106 | 1.27E-09 | 1 |
| rs546181 | 11 | 30855237 | 30 | 30798255 | 31020084 | T | G | 0.2972 | DCDC1 | intronic | 0.0928 | 0.0142 | 5.72E-11 | 0.0721 | 0.0288 | 1.23E-02 | 0.0640 | 0.0106 | 1.61E-09 | 1 |
| rs2312068 | 11 | 30855691 | 30 | 30798255 | 31020084 | A | G | 0.2992 | DCDC1 | intronic | 0.0925 | 0.0142 | 6.31E-11 | 0.0730 | 0.0288 | 1.12E-02 | 0.0645 | 0.0106 | 1.25E-09 | 1 |
| rs7127215 | 11 | 30858131 | 30 | 30798255 | 31020084 | A | G | 0.2992 | DCDC1 | intronic | -0.0927 | 0.0142 | 5.86E-11 | -0.0730 | 0.0288 | 1.12E-02 | -0.0644 | 0.0106 | 1.31E-09 | 1 |
| rs1375378 | 11 | 30859389 | 30 | 30798255 | 31020084 | T | C | 0.2992 | DCDC1 | intronic | -0.0926 | 0.0141 | 5.86E-11 | -0.0730 | 0.0288 | 1.12E-02 | -0.0644 | 0.0106 | 1.30E-09 | 1 |
| rs1823307 | 11 | 30859628 | 30 | 30798255 | 31020084 | C | G | 0.2992 | DCDC1 | intronic | 0.0939 | 0.0142 | 3.72E-11 | 0.0748 | 0.0288 | 9.38E-03 | 0.0647 | 0.0107 | 1.38E-09 | 1 |
| rs1823306 | 11 | 30859760 | 30 | 30798255 | 31020084 | A | G | 0.2982 | DCDC1 | intronic | -0.092 | 0.0142 | 8.14E-11 | -0.0730 | 0.0288 | 1.12E-02 | -0.0649 | 0.0106 | 9.57E-10 | 1 |
| rs2122135 | 11 | 30860112 | 30 | 30798255 | 31020084 | T | C | 0.2992 | DCDC1 | intronic | 0.0925 | 0.0141 | 6.13E-11 | 0.0730 | 0.0288 | 1.13E-02 | 0.0642 | 0.0106 | 1.41E-09 | 1 |
| rs10767877 | 11 | 30860580 | 30 | 30798255 | 31020084 | A | G | 0.3002 | DCDC1 | intronic | -0.0939 | 0.0141 | 3.27E-11 | -0.0726 | 0.0288 | 1.16E-02 | -0.0647 | 0.0106 | 1.03E-09 | 1 |
| rs808471 | 11 | 30861472 | 30 | 30798255 | 31020084 | C | G | 0.2972 | DCDC1 | intronic | -0.0945 | 0.0142 | 2.45E-11 | -0.0686 | 0.0288 | 1.71E-02 | -0.0645 | 0.0106 | 1.16E-09 | 1 |
| rs1028643 | 11 | 30861880 | 30 | 30798255 | 31020084 | A | G | 0.3002 | DCDC1 | intronic | 0.094 | 0.0142 | 3.02E-11 | 0.0714 | 0.0288 | 1.31E-02 | 0.0647 | 0.0106 | 1.07E-09 | 1 |
| rs273558 | 11 | 30864319 | 30 | 30798255 | 31020084 | A | G | 0.3012 | DCDC1 | intronic | 0.0938 | 0.0142 | 3.49E-11 | 0.0714 | 0.0288 | 1.31E-02 | 0.0647 | 0.0106 | 1.04E-09 | 1 |
| rs273557 | 11 | 30864854 | 30 | 30798255 | 31020084 | A | G | 0.3002 | DCDC1 | intronic | -0.0943 | 0.0141 | 2.64E-11 | -0.0696 | 0.0288 | 1.56E-02 | -0.0650 | 0.0106 | 8.88E-10 | 1 |
| rs273556 | 11 | 30866279 | 30 | 30798255 | 31020084 | A | C | 0.2962 | DCDC1 | intronic | 0.0925 | 0.0142 | 6.57E-11 | 0.0723 | 0.0288 | 1.21E-02 | 0.0639 | 0.0106 | 1.65E-09 | 1 |
| rs273555 | 11 | 30866864 | 30 | 30798255 | 31020084 | T | C | 0.2962 | DCDC1 | intronic | -0.0924 | 0.0142 | 6.68E-11 | -0.0723 | 0.0288 | 1.21E-02 | -0.0639 | 0.0106 | 1.66E-09 | 1 |
| rs273554 | 11 | 30868366 | 30 | 30798255 | 31020084 | A | G | 0.2982 | DCDC1 | intronic | 0.0936 | 0.0141 | 3.77E-11 | 0.0722 | 0.0288 | 1.20E-02 | 0.0647 | 0.0106 | 1.01E-09 | 1 |
| rs273553 | 11 | 30871743 | 30 | 30798255 | 31020084 | T | C | 0.2972 | DCDC1 | intronic | -0.0912 | 0.0141 | 1.14E-10 | -0.0758 | 0.0288 | 8.42E-03 | -0.0639 | 0.0106 | 1.61E-09 | 1 |
| rs273552 | 11 | 30872241 | 30 | 30798255 | 31020084 | T | C | 0.2962 | DCDC1 | intronic | 0.0922 | 0.0142 | 7.29E-11 | 0.0723 | 0.0288 | 1.20E-02 | 0.0637 | 0.0106 | 1.80E-09 | 1 |
| rs273551 | 11 | 30872579 | 30 | 30798255 | 31020084 | A | C | 0.2972 | DCDC1 | intronic | 0.0914 | 0.0141 | 1.06E-10 | 0.0764 | 0.0288 | 7.97E-03 | 0.0637 | 0.0106 | 1.79E-09 | 1 |
| rs273550 | 11 | 30873650 | 30 | 30798255 | 31020084 | T | C | 0.2972 | DCDC1 | intronic | 0.0928 | 0.0142 | 6.26E-11 | 0.0764 | 0.0288 | 7.95E-03 | 0.0642 | 0.0107 | 2.03E-09 | 1 |
| rs273597 | 11 | 30875093 | 30 | 30798255 | 31020084 | T | C | 0.2982 | DCDC1 | intronic | -0.0925 | 0.0141 | 6.01E-11 | -0.0725 | 0.0288 | 1.17E-02 | -0.0645 | 0.0106 | 1.12E-09 | 1 |
| rs273594 | 11 | 30876113 | 30 | 30798255 | 31020084 | C | G | 0.3012 | DCDC1 | intronic | 0.0901 | 0.0143 | 3.13E-10 | 0.0719 | 0.0288 | 1.24E-02 | 0.0622 | 0.0109 | 1.17E-08 | 1 |
| rs273593 | 11 | 30878012 | 30 | 30798255 | 31020084 | T | C | 0.3012 | DCDC1 | intronic | -0.093 | 0.0141 | 4.70E-11 | -0.0707 | 0.0288 | 1.40E-02 | -0.0653 | 0.0106 | 7.12E-10 | 1 |
| rs273592 | 11 | 30878849 | 30 | 30798255 | 31020084 | T | C | 0.3012 | DCDC1 | UTR3 | 0.093 | 0.0141 | 4.76E-11 | 0.0707 | 0.0288 | 1.40E-02 | 0.0654 | 0.0106 | 6.66E-10 | 1 |
| rs273589 | 11 | 30881109 | 30 | 30798255 | 31020084 | A | G | 0.2982 | DCDC1 | intronic | -0.0944 | 0.0142 | 3.48E-11 | -0.0738 | 0.0288 | 1.05E-02 | -0.0664 | 0.0108 | 7.19E-10 | 1 |
| rs273587 | 11 | 30883876 | 30 | 30798255 | 31020084 | A | T | 0.2922 | DCDC1 | intronic | -0.0929 | 0.0142 | 6.78E-11 | -0.0706 | 0.0291 | 1.52E-02 | -0.0648 | 0.0107 | 1.27E-09 | 1 |
| rs273582 | 11 | 30890857 | 30 | 30798255 | 31020084 | T | C | 0.2982 | DCDC1 | intronic | 0.0926 | 0.0141 | 5.83E-11 | 0.0678 | 0.0287 | 1.83E-02 | 0.0649 | 0.0106 | 9.07E-10 | 1 |
| rs612171 | 11 | 30893425 | 30 | 30798255 | 31020084 | A | G | 0.3012 | DCDC1 | intronic | 0.0894 | 0.0142 | 2.81E-10 | 0.0738 | 0.0287 | 1.03E-02 | 0.0643 | 0.0107 | 1.69E-09 | 1 |
| rs273618 | 11 | 30894383 | 30 | 30798255 | 31020084 | A | G | 0.3052 | DCDC1 | intronic | -0.0909 | 0.0141 | 1.17E-10 | -0.0720 | 0.0287 | 1.22E-02 | -0.0651 | 0.0106 | 8.01E-10 | 1 |
| rs273617 | 11 | 30895612 | 30 | 30798255 | 31020084 | T | C | 0.3012 | DCDC1 | intronic | -0.0904 | 0.0141 | 1.56E-10 | -0.0739 | 0.0287 | 1.01E-02 | -0.0644 | 0.0106 | 1.24E-09 | 1 |
| rs273616 | 11 | 30896283 | 30 | 30798255 | 31020084 | T | C | 0.2992 | DCDC1 | intronic | 0.0903 | 0.0141 | 1.61E-10 | 0.0737 | 0.0287 | 1.03E-02 | 0.0643 | 0.0106 | 1.35E-09 | 1 |
| rs273615 | 11 | 30899836 | 30 | 30798255 | 31020084 | A | C | 0.2992 | DCDC1 | intronic | -0.0911 | 0.0141 | 1.12E-10 | -0.0737 | 0.0287 | 1.03E-02 | -0.0641 | 0.0106 | 1.67E-09 | 1 |
| rs273614 | 11 | 30901982 | 30 | 30798255 | 31020084 | A | G | 0.3022 | DCDC1 | intronic | -0.088 | 0.0145 | 1.21E-09 | -0.0706 | 0.0287 | 1.39E-02 | -0.0649 | 0.0106 | 9.63E-10 | 1 |
| rs273613 | 11 | 30902893 | 30 | 30798255 | 31020084 | T | C | 0.3032 | DCDC1 | intronic | -0.0924 | 0.0141 | 5.67E-11 | -0.0705 | 0.0287 | 1.40E-02 | -0.0651 | 0.0106 | 8.51E-10 | 1 |
| rs273612 | 11 | 30903063 | 30 | 30798255 | 31020084 | A | G | 0.2962 | DCDC1 | intronic | 0.092 | 0.0142 | 8.65E-11 | 0.0732 | 0.0287 | 1.08E-02 | 0.0653 | 0.0107 | 9.18E-10 | 1 |
| rs273579 | 11 | 30908049 | 30 | 30798255 | 31020084 | T | C | 0.2992 | DCDC1 | intronic | -0.092 | 0.0141 | 7.97E-11 | -0.0721 | 0.0287 | 1.20E-02 | -0.0652 | 0.0107 | 9.84E-10 | 1 |
| rs273576 | 11 | 30910036 | 30 | 30798255 | 31020084 | A | G | 0.2992 | DCDC1 | intronic | -0.0922 | 0.0142 | 7.35E-11 | -0.0720 | 0.0287 | 1.21E-02 | -0.0638 | 0.0107 | 2.65E-09 | 1 |
| rs273573 | 11 | 30910991 | 30 | 30798255 | 31020084 | A | C | 0.2992 | DCDC1 | intronic | -0.0921 | 0.0141 | 6.70E-11 | -0.0720 | 0.0287 | 1.21E-02 | -0.0643 | 0.0107 | 1.62E-09 | 1 |
| rs273572 | 11 | 30911742 | 30 | 30798255 | 31020084 | T | C | 0.2992 | DCDC1 | intronic | 0.0922 | 0.0141 | 6.52E-11 | 0.0720 | 0.0287 | 1.22E-02 | 0.0643 | 0.0107 | 1.62E-09 | 1 |
| rs273571 | 11 | 30911936 | 30 | 30798255 | 31020084 | C | G | 0.2992 | DCDC1 | intronic | -0.0922 | 0.0141 | 6.43E-11 | -0.0720 | 0.0287 | 1.22E-02 | -0.0644 | 0.0107 | 1.59E-09 | 1 |
| rs273569 | 11 | 30914035 | 30 | 30798255 | 31020084 | A | G | 0.2992 | DCDC1 | intronic | 0.0923 | 0.0141 | 6.39E-11 | 0.0719 | 0.0287 | 1.23E-02 | 0.0645 | 0.0107 | 1.48E-09 | 1 |
| rs273567 | 11 | 30917214 | 30 | 30798255 | 31020084 | T | C | 0.3052 | DCDC1 | intronic | 0.0938 | 0.0141 | 2.88E-11 | 0.0683 | 0.0287 | 1.74E-02 | 0.0646 | 0.0107 | 1.38E-09 | 1 |
| rs273565 | 11 | 30918425 | 30 | 30798255 | 31020084 | A | G | 0.3022 | DCDC1 | intronic | 0.093 | 0.0141 | 4.44E-11 | 0.0702 | 0.0288 | 1.48E-02 | 0.0640 | 0.0107 | 1.92E-09 | 1 |
| rs273564 | 11 | 30918860 | 30 | 30798255 | 31020084 | A | G | 0.3022 | DCDC1 | intronic | -0.0935 | 0.0142 | 4.12E-11 | -0.0701 | 0.0288 | 1.48E-02 | -0.0640 | 0.0107 | 2.37E-09 | 1 |
| rs273563 | 11 | 30919884 | 30 | 30798255 | 31020084 | A | G | 0.3042 | DCDC1 | intronic | 0.0938 | 0.0141 | 3.10E-11 | 0.0685 | 0.0288 | 1.73E-02 | 0.0643 | 0.0107 | 1.64E-09 | 1 |
| rs273562 | 11 | 30920430 | 30 | 30798255 | 31020084 | A | T | 0.3042 | DCDC1 | intronic | -0.0937 | 0.0141 | 3.20E-11 | -0.0663 | 0.0288 | 2.11E-02 | -0.0646 | 0.0107 | 1.32E-09 | 1 |
| rs273560 | 11 | 30922081 | 30 | 30798255 | 31020084 | T | C | 0.3042 | DCDC1 | intronic | 0.0937 | 0.0141 | 3.23E-11 | 0.0663 | 0.0288 | 2.11E-02 | 0.0646 | 0.0107 | 1.41E-09 | 1 |
| rs273559 | 11 | 30923223 | 30 | 30798255 | 31020084 | T | C | 0.3012 | DCDC1 | intronic | 0.0929 | 0.0141 | 4.67E-11 | 0.0700 | 0.0288 | 1.50E-02 | 0.0640 | 0.0107 | 2.00E-09 | 1 |
| rs395032 | 11 | 30927200 | 30 | 30798255 | 31020084 | T | C | 0.3012 | DCDC1 | intronic | -0.0928 | 0.0141 | 4.97E-11 | -0.0700 | 0.0288 | 1.50E-02 | -0.0637 | 0.0107 | 2.31E-09 | 1 |
| rs175399 | 11 | 30930374 | 30 | 30798255 | 31020084 | A | C | 0.3042 | DCDC1 | intronic | -0.0934 | 0.0141 | 3.76E-11 | -0.0664 | 0.0288 | 2.11E-02 | -0.0643 | 0.0107 | 1.68E-09 | 1 |

| | | | | | | | | | | | Meta-GWAS | | | UKBB | | | ENIGMA* | | | |
|---|---|---|---|---|---|---|---|---|---|---|---|---|---|---|---|---|---|---|---|---|
| rsID | chr | pos (hg19) | Genomic Locus | start | end | A1 | A2 | MAF | Nearest Genes | Function | Beta | SE | P-value | Beta | SE | P-value | Beta | SE | P-value | Replicated** |
| rs175398 | 11 | 30930393 | 30 | 30798255 | 31020084 | T | G | 0.3042 | DCDC1 | intronic | -0.0934 | 0.0141 | 3.80E-11 | -0.0664 | 0.0288 | 2.11E-02 | -0.0642 | 0.0107 | 1.76E-09 | 1 |
| rs273603 | 11 | 30931904 | 30 | 30798255 | 31020084 | T | C | 0.3052 | DCDC1 | intronic | 0.0931 | 0.0141 | 4.24E-11 | 0.0667 | 0.0288 | 2.03E-02 | 0.0643 | 0.0107 | 1.72E-09 | 1 |
| rs273601 | 11 | 30933646 | 30 | 30798255 | 31020084 | T | C | 0.3062 | DCDC1 | intronic | 0.0917 | 0.0141 | 8.38E-11 | 0.0683 | 0.0288 | 1.76E-02 | 0.0641 | 0.0107 | 1.95E-09 | 1 |
| rs172073 | 11 | 30934274 | 30 | 30798255 | 31020084 | T | C | 0.3052 | DCDC1 | intronic | -0.0921 | 0.0141 | 7.07E-11 | -0.0677 | 0.0288 | 1.87E-02 | -0.0641 | 0.0107 | 1.87E-09 | 1 |
| rs273610 | 11 | 30938218 | 30 | 30798255 | 31020084 | A | G | 0.3062 | DCDC1 | intronic | 0.0924 | 0.0141 | 6.12E-11 | 0.0675 | 0.0288 | 1.90E-02 | 0.0642 | 0.0107 | 1.82E-09 | 1 |
| rs273608 | 11 | 30941272 | 30 | 30798255 | 31020084 | A | C | 0.3052 | DCDC1 | intronic | 0.0924 | 0.0141 | 6.30E-11 | 0.0679 | 0.0288 | 1.84E-02 | 0.0642 | 0.0107 | 1.80E-09 | 1 |
| rs163886 | 11 | 30943561 | 30 | 30798255 | 31020084 | A | T | 0.3052 | DCDC1 | intronic | 0.093 | 0.0142 | 5.05E-11 | 0.0677 | 0.0288 | 1.87E-02 | 0.0638 | 0.0107 | 2.31E-09 | 1 |
| rs163885 | 11 | 30945365 | 30 | 30798255 | 31020084 | T | C | 0.3052 | DCDC1 | intronic | 0.0923 | 0.0141 | 6.80E-11 | 0.0673 | 0.0288 | 1.94E-02 | 0.0643 | 0.0107 | 1.81E-09 | 1 |
| rs163883 | 11 | 30947248 | 30 | 30798255 | 31020084 | A | G | 0.3012 | DCDC1 | intronic | -0.091 | 0.0142 | 1.36E-10 | -0.0701 | 0.0288 | 1.51E-02 | -0.0644 | 0.0107 | 1.74E-09 | 1 |
| rs163882 | 11 | 30947681 | 30 | 30798255 | 31020084 | C | G | 0.3012 | DCDC1 | intronic | -0.091 | 0.0142 | 1.37E-10 | -0.0701 | 0.0288 | 1.50E-02 | -0.0644 | 0.0107 | 1.76E-09 | 1 |
| rs163881 | 11 | 30948244 | 30 | 30798255 | 31020084 | T | G | 0.3052 | DCDC1 | intronic | -0.0914 | 0.0141 | 1.04E-10 | -0.0686 | 0.0288 | 1.74E-02 | -0.0642 | 0.0107 | 1.89E-09 | 1 |
| rs163879 | 11 | 30951674 | 30 | 30798255 | 31020084 | T | C | 0.3062 | DCDC1 | intronic | 0.091 | 0.0141 | 1.27E-10 | 0.0677 | 0.0288 | 1.89E-02 | 0.0641 | 0.0107 | 2.00E-09 | 1 |
| rs163877 | 11 | 30953608 | 30 | 30798255 | 31020084 | A | G | 0.2982 | DCDC1 | intronic | -0.09 | 0.0142 | 2.52E-10 | -0.0706 | 0.0289 | 1.48E-02 | -0.0634 | 0.0107 | 3.38E-09 | 1 |
| rs156399 | 11 | 30955947 | 30 | 30798255 | 31020084 | A | G | 0.2962 | DCDC1 | intronic | -0.0904 | 0.0142 | 2.07E-10 | -0.0717 | 0.0289 | 1.33E-02 | -0.0636 | 0.0107 | 3.16E-09 | 1 |
| rs163876 | 11 | 30956077 | 30 | 30798255 | 31020084 | A | T | 0.2982 | DCDC1 | intronic | -0.0905 | 0.0142 | 2.00E-10 | -0.0706 | 0.0289 | 1.47E-02 | -0.0634 | 0.0107 | 3.33E-09 | 1 |
| rs158629 | 11 | 30962964 | 30 | 30798255 | 31020084 | C | G | 0.3072 | DCDC1 | intronic | -0.0963 | 0.017 | 1.56E-08 | -0.0679 | 0.0288 | 1.85E-02 | -0.0642 | 0.0107 | 1.92E-09 | 1 |
| rs163854 | 11 | 30964632 | 30 | 30798255 | 31020084 | T | G | 0.3012 | DCDC1 | intronic | 0.0901 | 0.0142 | 2.12E-10 | 0.0699 | 0.0289 | 1.55E-02 | 0.0640 | 0.0107 | 2.38E-09 | 1 |
| rs273605 | 11 | 30965855 | 30 | 30798255 | 31020084 | A | T | 0.3052 | DCDC1 | UTR3 | -0.0912 | 0.0142 | 1.19E-10 | -0.0691 | 0.0289 | 1.67E-02 | -0.0637 | 0.0107 | 2.62E-09 | 1 |
| rs273604 | 11 | 30966112 | 30 | 30798255 | 31020084 | A | G | 0.3032 | DCDC1 | intronic | 0.0908 | 0.0142 | 1.46E-10 | 0.0700 | 0.0289 | 1.53E-02 | 0.0639 | 0.0107 | 2.39E-09 | 1 |
| rs163874 | 11 | 30970207 | 30 | 30798255 | 31020084 | T | C | 0.3012 | DCDC1 | intronic | 0.0907 | 0.0142 | 1.59E-10 | 0.0698 | 0.0289 | 1.57E-02 | 0.0632 | 0.0107 | 3.35E-09 | 1 |
| rs158505 | 11 | 30981248 | 30 | 30798255 | 31020084 | T | C | 0.3002 | DCDC1 | intronic | -0.0905 | 0.0142 | 1.66E-10 | -0.0702 | 0.0289 | 1.51E-02 | -0.0632 | 0.0107 | 3.42E-09 | 1 |
| rs158582 | 11 | 30995033 | 30 | 30798255 | 31020084 | T | C | 0.2922 | DCDC1 | intronic | -0.0882 | 0.0143 | 6.29E-10 | -0.0738 | 0.0292 | 1.14E-02 | -0.0630 | 0.0107 | 4.49E-09 | 1 |
| rs9667150 | 11 | 31010455 | 30 | 30798255 | 31020084 | A | G | 0.4046 | DCDC1 | intronic | 0.0741 | 0.0132 | 2.18E-08 | 0.0361 | 0.0274 | 1.89E-01 | 0.0507 | 0.0100 | 4.38E-07 | 0 |
| rs290108 | 11 | 31010962 | 30 | 30798255 | 31020084 | T | C | 0.4026 | DCDC1 | intronic | 0.0736 | 0.0132 | 2.75E-08 | 0.0389 | 0.0275 | 1.56E-01 | 0.0502 | 0.0100 | 5.71E-07 | 0 |
| rs290106 | 11 | 31012288 | 30 | 30798255 | 31020084 | T | G | 0.4026 | DCDC1 | intronic | 0.0737 | 0.0133 | 2.68E-08 | 0.0390 | 0.0275 | 1.56E-01 | 0.0512 | 0.0101 | 3.64E-07 | 0 |
| rs3102163 | 11 | 103988675 | 31 | 103985306 | 104041862 | T | C | 0.4105 | PDGFD | intronic | -0.0762 | 0.0139 | 4.01E-08 | -0.0754 | 0.0277 | 6.54E-03 | -0.0524 | 0.0112 | 2.62E-06 | 1 |
| rs1545865 | 11 | 103989190 | 31 | 103985306 | 104041862 | T | C | 0.4105 | PDGFD | intronic | -0.0762 | 0.0139 | 4.27E-08 | -0.0751 | 0.0277 | 6.80E-03 | -0.0538 | 0.0112 | 1.52E-06 | 1 |
| rs361293 | 11 | 103994966 | 31 | 103985306 | 104041862 | A | C | 0.4105 | PDGFD | intronic | -0.082 | 0.0133 | 7.45E-10 | -0.0758 | 0.0277 | 6.26E-03 | -0.0488 | 0.0103 | 2.10E-06 | 1 |
| rs361301 | 11 | 103998180 | 31 | 103985306 | 104041862 | T | C | 0.4105 | PDGFD | intronic | -0.0812 | 0.0133 | 1.07E-09 | -0.0751 | 0.0277 | 6.76E-03 | -0.0485 | 0.0103 | 2.29E-06 | 1 |
| rs361300 | 11 | 103998386 | 31 | 103985306 | 104041862 | A | C | 0.4115 | PDGFD | intronic | 0.0814 | 0.0133 | 9.55E-10 | 0.0749 | 0.0277 | 6.87E-03 | 0.0487 | 0.0103 | 2.10E-06 | 1 |
| rs667801 | 11 | 104003647 | 31 | 103985306 | 104041862 | T | C | 0.4105 | PDGFD | intronic | 0.083 | 0.0133 | 3.93E-10 | 0.0757 | 0.0277 | 6.29E-03 | 0.0486 | 0.0102 | 1.95E-06 | 1 |
| rs551198 | 11 | 104004552 | 31 | 103985306 | 104041862 | A | G | 0.4105 | PDGFD | intronic | 0.083 | 0.0133 | 3.77E-10 | 0.0763 | 0.0277 | 5.96E-03 | 0.0479 | 0.0102 | 2.65E-06 | 1 |
| rs551300 | 11 | 104004596 | 31 | 103985306 | 104041862 | A | G | 0.4115 | PDGFD | intronic | -0.0833 | 0.0133 | 3.40E-10 | -0.0758 | 0.0277 | 6.26E-03 | -0.0484 | 0.0102 | 2.07E-06 | 1 |
| rs361273 | 11 | 104006346 | 31 | 103985306 | 104041862 | A | G | 0.4105 | PDGFD | intronic | -0.0809 | 0.0132 | 8.22E-10 | -0.0763 | 0.0277 | 5.97E-03 | -0.0478 | 0.0102 | 2.48E-06 | 1 |
| rs361270 | 11 | 104008724 | 31 | 103985306 | 104041862 | A | C | 0.4115 | PDGFD | intronic | -0.0809 | 0.0132 | 8.18E-10 | -0.0759 | 0.0277 | 6.16E-03 | -0.0475 | 0.0101 | 2.77E-06 | 1 |
| rs361269 | 11 | 104009590 | 31 | 103985306 | 104041862 | T | C | 0.4115 | PDGFD | intronic | 0.0809 | 0.0132 | 8.13E-10 | 0.0759 | 0.0277 | 6.17E-03 | 0.0475 | 0.0101 | 2.75E-06 | 1 |
| rs361268 | 11 | 104010052 | 31 | 103985306 | 104041862 | T | C | 0.4105 | PDGFD | intronic | 0.0806 | 0.0132 | 9.39E-10 | 0.0764 | 0.0277 | 5.86E-03 | 0.0466 | 0.0101 | 4.20E-06 | 1 |
| rs1681464 | 11 | 104012656 | 31 | 103985306 | 104041862 | A | G | 0.4105 | PDGFD | intronic | -0.0829 | 0.0132 | 3.02E-10 | -0.0751 | 0.0277 | 6.74E-03 | -0.0469 | 0.0101 | 3.49E-06 | 1 |
| rs2170719 | 11 | 104012767 | 31 | 103985306 | 104041862 | A | G | 0.4105 | PDGFD | intronic | 0.0826 | 0.0132 | 3.51E-10 | 0.0751 | 0.0277 | 6.75E-03 | 0.0464 | 0.0101 | 4.55E-06 | 1 |
| rs2760105 | 13 | 80109867 | 32 | 79859456 | 80251200 | A | G | 0.3767 | NDFIP2 | intronic | -0.0744 | 0.0134 | 2.78E-08 | -0.0628 | 0.0276 | 2.30E-02 | -0.0407 | 0.0102 | 6.70E-05 | 1 |
| rs9601242 | 13 | 80135678 | 32 | 79859456 | 80251200 | A | C | 0.4145 | LINC01068 | intergenic | -0.0721 | 0.0132 | 4.28E-08 | -0.0637 | 0.0272 | 1.94E-02 | -0.0444 | 0.0101 | 1.01E-05 | 1 |
| rs73235963 | 13 | 80136940 | 32 | 79859456 | 80251200 | A | C | 0.4145 | LINC01068 | intergenic | -0.0723 | 0.0132 | 3.95E-08 | -0.0636 | 0.0272 | 1.95E-02 | -0.0444 | 0.0101 | 1.00E-05 | 1 |
| rs2484343 | 13 | 80137722 | 32 | 79859456 | 80251200 | A | G | 0.4145 | LINC01068 | intergenic | 0.0724 | 0.0132 | 3.76E-08 | 0.0637 | 0.0272 | 1.94E-02 | 0.0445 | 0.0101 | 9.77E-06 | 1 |
| rs2476173 | 13 | 80139875 | 32 | 79859456 | 80251200 | A | G | 0.4145 | LINC01068 | intergenic | -0.0726 | 0.0132 | 3.65E-08 | -0.0636 | 0.0272 | 1.95E-02 | -0.0446 | 0.0101 | 9.24E-06 | 1 |
| rs1208049 | 13 | 80144355 | 32 | 79859456 | 80251200 | T | G | 0.4145 | LINC01068 | ncRNA_intronic | -0.073 | 0.0132 | 3.18E-08 | -0.0638 | 0.0272 | 1.91E-02 | -0.0439 | 0.0101 | 1.37E-05 | 1 |
| rs2783125 | 13 | 80157890 | 32 | 79859456 | 80251200 | A | G | 0.4105 | LINC01068 | intergenic | 0.0732 | 0.0132 | 2.85E-08 | 0.0699 | 0.0273 | 1.03E-02 | 0.0452 | 0.0101 | 7.67E-06 | 1 |
| rs2783126 | 13 | 80158701 | 32 | 79859456 | 80251200 | T | C | 0.4115 | LINC01068 | intergenic | -0.0732 | 0.0132 | 2.83E-08 | -0.0699 | 0.0273 | 1.04E-02 | -0.0451 | 0.0101 | 8.15E-06 | 1 |
| rs9545142 | 13 | 80162555 | 32 | 79859456 | 80251200 | T | C | 0.4105 | LINC01068 | intergenic | 0.0737 | 0.0132 | 2.39E-08 | 0.0700 | 0.0273 | 1.03E-02 | 0.0454 | 0.0101 | 7.21E-06 | 1 |
| rs2783129 | 13 | 80168720 | 32 | 79859456 | 80251200 | C | G | 0.496 | LINC01068 | intergenic | -0.1062 | 0.0132 | 7.62E-16 | -0.0970 | 0.0271 | 3.55E-04 | NA | NA | NA | NA |
| rs9601248 | 13 | 80169004 | 32 | 79859456 | 80251200 | T | C | 0.4602 | LINC01068 | intergenic | -0.1029 | 0.013 | 2.62E-15 | -0.1002 | 0.0272 | 2.24E-04 | -0.0497 | 0.0098 | 4.09E-07 | 1 |
| rs2783130 | 13 | 80170160 | 32 | 79859456 | 80251200 | A | G | 0.497 | LINC01068 | intergenic | -0.1192 | 0.013 | 4.73E-20 | -0.1064 | 0.0270 | 8.43E-05 | -0.0527 | 0.0099 | 9.17E-08 | 1 |
| rs9545145 | 13 | 80173760 | 32 | 79859456 | 80251200 | A | G | 0.4702 | LINC01068 | intergenic | -0.1046 | 0.0131 | 1.24E-15 | -0.1040 | 0.0270 | 1.20E-04 | -0.0492 | 0.0099 | 6.71E-07 | 1 |
| rs7319427 | 13 | 80174329 | 32 | 79859456 | 80251200 | T | C | 0.4702 | LINC01068 | intergenic | -0.1041 | 0.013 | 1.07E-15 | -0.1040 | 0.0270 | 1.21E-04 | -0.0475 | 0.0098 | 1.28E-06 | 1 |
| rs6563121 | 13 | 80175000 | 32 | 79859456 | 80251200 | T | C | 0.496 | LINC01068 | intergenic | 0.1211 | 0.013 | 1.45E-20 | 0.1076 | 0.0270 | 6.99E-05 | 0.0507 | 0.0099 | 3.17E-07 | 1 |
| rs9545146 | 13 | 80175606 | 32 | 79859456 | 80251200 | T | G | 0.4284 | LINC01068 | intergenic | 0.1034 | 0.0131 | 2.75E-15 | 0.0909 | 0.0270 | 7.80E-04 | 0.0417 | 0.0099 | 2.36E-05 | 1 |
| rs928640 | 13 | 80177483 | 32 | 79859456 | 80251200 | A | G | 0.3867 | LINC01068 | intergenic | -0.0918 | 0.0133 | 4.45E-12 | -0.0951 | 0.0278 | 6.39E-04 | -0.0333 | 0.0100 | 8.85E-04 | 1 |
| rs9530937 | 13 | 80177836 | 32 | 79859456 | 80251200 | A | G | 0.4115 | LINC01068 | intergenic | 0.1053 | 0.0132 | 1.24E-15 | 0.0984 | 0.0275 | 3.52E-04 | 0.0363 | 0.0100 | 2.79E-04 | 1 |
| rs2760111 | 13 | 80178344 | 32 | 79859456 | 80251200 | T | C | 0.327 | LINC01068 | intergenic | -0.1029 | 0.0147 | 2.80E-12 | -0.0987 | 0.0287 | 5.87E-04 | -0.0529 | 0.0114 | 3.71E-06 | 1 |
| rs2146593 | 13 | 80180112 | 32 | 79859456 | 80251200 | T | C | 0.3429 | LINC01068 | intergenic | -0.0937 | 0.0135 | 4.03E-12 | -0.0849 | 0.0281 | 2.58E-03 | -0.0282 | 0.0102 | 5.67E-03 | 1 |
| rs2146592 | 13 | 80180281 | 32 | 79859456 | 80251200 | A | G | 0.3857 | LINC01068 | intergenic | -0.0917 | 0.0133 | 4.82E-12 | -0.0969 | 0.0278 | 4.98E-04 | -0.0334 | 0.0100 | 8.62E-04 | 1 |
| rs9545151 | 13 | 80182535 | 32 | 79859456 | 80251200 | A | T | 0.3429 | LINC01068 | intergenic | 0.0938 | 0.0135 | 3.70E-12 | 0.0851 | 0.0281 | 2.50E-03 | 0.0287 | 0.0102 | 4.96E-03 | 1 |
| rs1932197 | 13 | 80183353 | 32 | 79859456 | 80251200 | C | G | 0.499 | LINC01068 | intergenic | -0.1197 | 0.013 | 3.54E-20 | -0.1063 | 0.0270 | 8.56E-05 | NA | NA | NA | NA |
| rs7990151 | 13 | 80183985 | 32 | 79859456 | 80251200 | A | G | 0.3867 | LINC01068 | intergenic | -0.0919 | 0.0133 | 4.42E-12 | -0.0971 | 0.0278 | 4.86E-04 | -0.0341 | 0.0101 | 6.90E-04 | 1 |
| rs2876741 | 13 | 80186712 | 32 | 79859456 | 80251200 | A | T | 0.3429 | LINC01068 | intergenic | -0.093 | 0.0135 | 5.62E-12 | -0.0848 | 0.0281 | 2.57E-03 | -0.0288 | 0.0102 | 4.91E-03 | 1 |
| NA | 13 | 80188088 | 32 | 79859456 | 80251200 | A | T | 0.498 | LINC01068 | intergenic | 0.1035 | 0.0179 | 6.86E-09 | 0.0975 | 0.0273 | 3.56E-04 | NA | NA | NA | NA |
| rs2093620 | 13 | 80188089 | 32 | 79859456 | 80251200 | A | C | 0.498 | LINC01068 | intergenic | 0.1208 | 0.0132 | 5.22E-20 | 0.1073 | 0.0270 | 7.42E-05 | 0.0551 | 0.0101 | 5.19E-08 | 1 |
| rs9601250 | 13 | 80188702 | 32 | 79859456 | 80251200 | T | C | 0.3648 | LINC01068 | intergenic | 0.1037 | 0.0141 | 1.71E-13 | 0.1124 | 0.0279 | 5.83E-05 | 0.0489 | 0.0109 | 7.46E-06 | 1 |
| rs9601251 | 13 | 80189717 | 32 | 79859456 | 80251200 | T | C | 0.3847 | LINC01068 | intergenic | -0.0904 | 0.0133 | 1.01E-11 | -0.0967 | 0.0278 | 5.16E-04 | -0.0348 | 0.0101 | 5.67E-04 | 1 |
| rs9601252 | 13 | 80189742 | 32 | 79859456 | 80251200 | T | C | 0.3857 | LINC01068 | intergenic | -0.0914 | 0.0133 | 5.58E-12 | -0.0966 | 0.0278 | 5.25E-04 | -0.0341 | 0.0101 | 7.04E-04 | 1 |
| rs4141825 | 13 | 80190848 | 32 | 79859456 | 80251200 | A | G | 0.498 | LINC01068 | intergenic | -0.1191 | 0.013 | 4.89E-20 | -0.1069 | 0.0271 | 7.94E-05 | -0.0523 | 0.0099 | 1.19E-07 | 1 |
| rs9545154 | 13 | 80191817 | 32 | 79859456 | 80251200 | C | G | 0.4729 | LINC01068 | intergenic | -0.1239 | 0.0132 | 4.62E-21 | -0.0954 | 0.0272 | 4.48E-04 | NA | NA | NA | NA |
| rs9545155 | 13 | 80191873 | 32 | 79859456 | 80251200 | T | C | 0.498 | LINC01068 | intergenic | -0.1203 | 0.013 | 2.19E-20 | -0.1067 | 0.0271 | 8.08E-05 | -0.0528 | 0.0099 | 1.01E-07 | 1 |
| rs9545156 | 13 | 80192236 | 32 | 79859456 | 80251200 | A | G | 0.498 | LINC01068 | intergenic | -0.1252 | 0.015 | 7.53E-17 | -0.1026 | 0.0367 | 5.16E-03 | -0.0528 | 0.0103 | 2.88E-07 | 1 |
| rs9545158 | 13 | 80192906 | 32 | 79859456 | 80251200 | A | G | 0.498 | LINC01068 | intergenic | 0.1203 | 0.013 | 2.71E-20 | 0.1066 | 0.0271 | 8.28E-05 | 0.0532 | 0.0099 | 8.34E-08 | 1 |
| rs7997134 | 13 | 80193729 | 32 | 79859456 | 80251200 | A | G | 0.3857 | LINC01068 | intergenic | -0.091 | 0.0133 | 6.97E-12 | -0.0973 | 0.0279 | 4.84E-04 | -0.0340 | 0.0101 | 7.69E-04 | 1 |
| rs2876742 | 13 | 80196829 | 32 | 79859456 | 80251200 | T | C | 0.3678 | LINC01068 | intergenic | -0.0907 | 0.0135 | 1.81E-11 | -0.0701 | 0.0283 | 1.32E-02 | -0.0470 | 0.0103 | 4.83E-06 | 1 |
| rs767778 | 13 | 80197115 | 32 | 79859456 | 80251200 | A | T | 0.3668 | LINC01068 | intergenic | 0.092 | 0.0135 | 9.12E-12 | 0.0706 | 0.0283 | 1.27E-02 | 0.0468 | 0.0103 | 5.18E-06 | 1 |
| rs9318645 | 13 | 80199411 | 32 | 79859456 | 80251200 | A | G | 0.3678 | LINC01068 | intergenic | -0.0922 | 0.0135 | 7.90E-12 | -0.0702 | 0.0283 | 1.31E-02 | -0.0467 | 0.0103 | 5.41E-06 | 1 |
| rs4885647 | 13 | 80199998 | 32 | 79859456 | 80251200 | A | C | 0.3668 | LINC01068 | intergenic | -0.0922 | 0.0135 | 7.96E-12 | -0.0703 | 0.0283 | 1.29E-02 | -0.0468 | 0.0103 | 5.20E-06 | 1 |
| rs9545159 | 13 | 80202674 | 32 | 79859456 | 80251200 | T | C | 0.3668 | LINC01068 | intergenic | -0.0922 | 0.0135 | 7.85E-12 | -0.0695 | 0.0283 | 1.40E-02 | -0.0467 | 0.0103 | 5.45E-06 | 1 |
| rs2329126 | 13 | 80203805 | 32 | 79859456 | 80251200 | A | G | 0.3668 | LINC01068 | intergenic | -0.092 | 0.0135 | 8.28E-12 | -0.0693 | 0.0283 | 1.42E-02 | -0.0466 | 0.0103 | 5.55E-06 | 1 |
| rs9601256 | 13 | 80208050 | 32 | 79859456 | 80251200 | T | C | 0.3668 | LINC01068 | intergenic | 0.0922 | 0.0135 | 7.72E-12 | 0.0696 | 0.0283 | 1.39E-02 | 0.0464 | 0.0103 | 6.19E-06 | 1 |
| rs4885649 | 13 | 80208801 | 32 | 79859456 | 80251200 | A | T | 0.3668 | LINC01068 | intergenic | 0.0921 | 0.0135 | 8.05E-12 | 0.0696 | 0.0283 | 1.38E-02 | 0.0463 | 0.0103 | 6.33E-06 | 1 |
| rs9530940 | 13 | 80209455 | 32 | 79859456 | 80251200 | A | G | 0.3678 | LINC01068 | intergenic | 0.0922 | 0.0135 | 7.35E-12 | 0.0696 | 0.0283 | 1.39E-02 | 0.0463 | 0.0103 | 6.21E-06 | 1 |
| rs3920578 | 13 | 80212443 | 32 | 79859456 | 80251200 | T | G | 0.3678 | LINC01068 | intergenic | 0.0923 | 0.0135 | 7.23E-12 | 0.0697 | 0.0282 | 1.36E-02 | 0.0463 | 0.0102 | 6.17E-06 | 1 |
| rs9530941 | 13 | 80213514 | 32 | 79859456 | 80251200 | T | C | 0.3678 | LINC01068 | intergenic | 0.0923 | 0.0135 | 7.11E-12 | 0.0698 | 0.0282 | 1.36E-02 | 0.0463 | 0.0102 | 6.17E-06 | 1 |
| rs9318647 | 13 | 80216846 | 32 | 79859456 | 80251200 | A | G | 0.3688 | LINC01068 | intergenic | -0.092 | 0.0135 | 7.91E-12 | -0.0705 | 0.0282 | 1.26E-02 | -0.0466 | 0.0102 | 5.28E-06 | 1 |
| rs9565516 | 13 | 80217881 | 32 | 79859456 | 80251200 | A | T | 0.341 | LINC01068 | intergenic | 0.0802 | 0.0138 | 5.41E-09 | 0.0673 | 0.0288 | 1.98E-02 | 0.0446 | 0.0104 | 1.86E-05 | 1 |
| rs12430226 | 13 | 80218239 | 32 | 79859456 | 80251200 | A | G | 0.3688 | LINC01068 | intergenic | 0.0921 | 0.0135 | 7.57E-12 | 0.0705 | 0.0282 | 1.25E-02 | 0.0468 | 0.0102 | 4.87E-06 | 1 |
| rs2181432 | 13 | 80218861 | 32 | 79859456 | 80251200 | T | C | 0.3678 | LINC01068 | intergenic | 0.092 | 0.0135 | 8.08E-12 | 0.0705 | 0.0282 | 1.25E-02 | 0.0467 | 0.0103 | 5.19E-06 | 1 |
| rs9530942 | 13 | 80219035 | 32 | 79859456 | 80251200 | A | T | 0.3678 | LINC01068 | intergenic | 0.092 | 0.0135 | 8.16E-12 | 0.0705 | 0.0282 | 1.25E-02 | 0.0467 | 0.0103 | 5.22E-06 | 1 |

| | | | | | | | | | | | Meta-GWAS | | | UKBB | | | ENIGMA* | | | |
|---|---|---|---|---|---|---|---|---|---|---|---|---|---|---|---|---|---|---|---|---|
| rsID | chr | pos (hg19) | Genomic Locus | start | end | A1 | A2 | MAF | Nearest Genes | Function | Beta | SE | P-value | Beta | SE | P-value | Beta | SE | P-value | Replicated** |
| rs9574450 | 13 | 80219665 | 32 | 79859456 | 80251200 | A | C | 0.3658 | LINC01068 | intergenic | -0.0914 | 0.0135 | 1.12E-11 | -0.0716 | 0.0282 | 1.11E-02 | -0.0467 | 0.0103 | 5.22E-06 | 1 |
| rs7994255 | 13 | 80219883 | 32 | 79859456 | 80251200 | T | G | 0.3658 | LINC01068 | intergenic | -0.0918 | 0.0135 | 9.12E-12 | -0.0734 | 0.0282 | 9.37E-03 | -0.0466 | 0.0103 | 5.47E-06 | 1 |
| rs7994620 | 13 | 80220015 | 32 | 79859456 | 80251200 | A | G | 0.34 | LINC01068 | intergenic | -0.0797 | 0.0137 | 6.21E-09 | -0.0706 | 0.0288 | 1.41E-02 | -0.0443 | 0.0104 | 2.16E-05 | 1 |
| rs4885652 | 13 | 80222402 | 32 | 79859456 | 80251200 | T | C | 0.34 | LINC01068 | intergenic | -0.0798 | 0.0137 | 6.16E-09 | -0.0720 | 0.0288 | 1.23E-02 | -0.0448 | 0.0104 | 1.74E-05 | 1 |
| rs9574452 | 13 | 80223671 | 32 | 79859456 | 80251200 | C | G | 0.34 | LINC01068 | intergenic | 0.0795 | 0.0137 | 7.06E-09 | 0.0721 | 0.0288 | 1.22E-02 | 0.0447 | 0.0104 | 1.83E-05 | 1 |
| rs4885653 | 13 | 80224421 | 32 | 79859456 | 80251200 | T | G | 0.34 | LINC01068 | intergenic | -0.0796 | 0.0137 | 6.62E-09 | -0.0721 | 0.0288 | 1.22E-02 | -0.0446 | 0.0104 | 1.87E-05 | 1 |
| rs9530943 | 13 | 80226450 | 32 | 79859456 | 80251200 | A | G | 0.3688 | LINC01068 | intergenic | -0.0918 | 0.0135 | 9.31E-12 | -0.0733 | 0.0283 | 9.52E-03 | -0.0468 | 0.0103 | 5.05E-06 | 1 |
| rs9545166 | 13 | 80227341 | 32 | 79859456 | 80251200 | A | T | 0.3678 | LINC01068 | intergenic | 0.092 | 0.0135 | 8.42E-12 | 0.0735 | 0.0283 | 9.35E-03 | 0.0467 | 0.0103 | 5.40E-06 | 1 |
| rs2876746 | 13 | 80231558 | 32 | 79859456 | 80251200 | A | G | 0.334 | LINC01068 | intergenic | -0.0781 | 0.0138 | 1.50E-08 | -0.0691 | 0.0289 | 1.67E-02 | -0.0437 | 0.0105 | 2.98E-05 | 1 |
| rs9601258 | 13 | 80232942 | 32 | 79859456 | 80251200 | A | G | 0.341 | LINC01068 | intergenic | 0.0793 | 0.0137 | 7.71E-09 | 0.0732 | 0.0287 | 1.10E-02 | 0.0445 | 0.0104 | 2.05E-05 | 1 |
| rs2876747 | 13 | 80233541 | 32 | 79859456 | 80251200 | T | G | 0.3419 | LINC01068 | intergenic | 0.0798 | 0.0137 | 6.35E-09 | 0.0726 | 0.0288 | 1.16E-02 | 0.0446 | 0.0104 | 1.99E-05 | 1 |
| rs9318650 | 13 | 80236762 | 32 | 79859456 | 80251200 | A | G | 0.3787 | LINC01068 | intergenic | 0.0978 | 0.0134 | 3.43E-13 | 0.0818 | 0.0282 | 3.71E-03 | 0.0444 | 0.0104 | 1.81E-05 | 1 |
| rs9530944 | 13 | 80238477 | 32 | 79859456 | 80251200 | A | C | 0.3678 | LINC01068 | intergenic | 0.092 | 0.0135 | 9.05E-12 | 0.0727 | 0.0283 | 1.01E-02 | 0.0480 | 0.0103 | 3.43E-06 | 1 |
| rs9318651 | 13 | 80238557 | 32 | 79859456 | 80251200 | T | C | 0.3678 | LINC01068 | intergenic | -0.0922 | 0.0135 | 8.24E-12 | -0.0732 | 0.0282 | 9.53E-03 | -0.0477 | 0.0103 | 3.92E-06 | 1 |
| rs2208938 | 13 | 80239283 | 32 | 79859456 | 80251200 | T | C | 0.3817 | LINC01068 | intergenic | 0.0871 | 0.0135 | 1.09E-10 | 0.0701 | 0.0282 | 1.28E-02 | 0.0452 | 0.0103 | 1.24E-05 | 1 |
| rs4885655 | 13 | 80240964 | 32 | 79859456 | 80251200 | C | G | 0.341 | LINC01068 | intergenic | -0.0795 | 0.0138 | 7.62E-09 | -0.0711 | 0.0288 | 1.35E-02 | -0.0460 | 0.0106 | 1.34E-05 | 1 |
| rs9805769 | 13 | 80242330 | 32 | 79859456 | 80251200 | A | C | 0.3678 | LINC01068 | intergenic | 0.092 | 0.0135 | 9.52E-12 | 0.0732 | 0.0283 | 9.59E-03 | 0.0482 | 0.0104 | 3.51E-06 | 1 |
| rs2208936 | 13 | 80242608 | 32 | 79859456 | 80251200 | C | G | 0.3688 | LINC01068 | intergenic | 0.0919 | 0.0135 | 1.02E-11 | 0.0715 | 0.0283 | 1.14E-02 | 0.0481 | 0.0104 | 3.80E-06 | 1 |
| rs9574455 | 13 | 80244736 | 32 | 79859456 | 80251200 | A | G | 0.341 | LINC01068 | intergenic | -0.0793 | 0.0138 | 8.67E-09 | -0.0713 | 0.0288 | 1.32E-02 | -0.0457 | 0.0106 | 1.54E-05 | 1 |
| rs4141826 | 13 | 80247791 | 32 | 79859456 | 80251200 | A | G | 0.3419 | LINC01068 | intergenic | -0.0782 | 0.0139 | 1.76E-08 | -0.0713 | 0.0288 | 1.32E-02 | -0.0462 | 0.0107 | 1.43E-05 | 1 |
| rs6563129 | 13 | 80250475 | 32 | 79859456 | 80251200 | T | C | 0.3459 | LINC01068 | intergenic | -0.0761 | 0.0139 | 4.47E-08 | -0.0715 | 0.0288 | 1.30E-02 | -0.0466 | 0.0107 | 1.41E-05 | 1 |
| rs2208943 | 13 | 80251200 | 32 | 79859456 | 80251200 | T | C | 0.3728 | LINC01068 | intergenic | 0.0894 | 0.0137 | 6.20E-11 | 0.0744 | 0.0283 | 8.53E-03 | 0.0494 | 0.0106 | 3.29E-06 | 1 |
| rs61976308 | 14 | 59449138 | 33 | 59449138 | 59891607 | A | G | 0.0497 | RP11-112J1.2 | ncRNA_intronic | -0.1742 | 0.0273 | 1.69E-10 | -0.1848 | 0.0550 | 7.83E-04 | -0.0433 | 0.0217 | 4.59E-02 | 1 |
| rs80223023 | 14 | 59450602 | 33 | 59449138 | 59891607 | A | C | 0.05169 | RP11-112J1.2 | ncRNA_intronic | 0.1789 | 0.0266 | 1.71E-11 | 0.1726 | 0.0548 | 1.65E-03 | 0.0545 | 0.0206 | 8.10E-03 | 1 |
| rs61976311 | 14 | 59456583 | 33 | 59449138 | 59891607 | A | G | 0.0507 | RP11-112J1.2 | ncRNA_intronic | -0.1811 | 0.0264 | 6.84E-12 | -0.1828 | 0.0548 | 8.48E-04 | -0.0545 | 0.0204 | 7.47E-03 | 1 |
| rs61976312 | 14 | 59457562 | 33 | 59449138 | 59891607 | T | C | 0.05169 | RP11-112J1.2 | ncRNA_intronic | 0.1818 | 0.0264 | 5.42E-12 | 0.1794 | 0.0547 | 1.05E-03 | 0.0550 | 0.0203 | 6.85E-03 | 1 |
| rs61976313 | 14 | 59459140 | 33 | 59449138 | 59891607 | T | C | 0.05268 | RP11-112J1.2 | ncRNA_intronic | 0.1809 | 0.0262 | 5.53E-12 | 0.1815 | 0.0547 | 9.13E-04 | 0.0553 | 0.0203 | 6.39E-03 | 1 |
| rs61976314 | 14 | 59464840 | 33 | 59449138 | 59891607 | C | G | 0.05268 | RP11-112J1.2 | ncRNA_intronic | 0.1812 | 0.0262 | 4.67E-12 | 0.1808 | 0.0547 | 9.56E-04 | 0.0555 | 0.0202 | 5.99E-03 | 1 |
| rs61976316 | 14 | 59465860 | 33 | 59449138 | 59891607 | A | G | 0.05169 | RP11-112J1.2 | ncRNA_intronic | -0.1805 | 0.0262 | 5.81E-12 | -0.1808 | 0.0547 | 9.59E-04 | -0.0544 | 0.0202 | 7.07E-03 | 1 |
| rs79329625 | 14 | 59466003 | 33 | 59449138 | 59891607 | A | G | 0.05169 | RP11-112J1.2 | ncRNA_intronic | -0.1805 | 0.0262 | 5.74E-12 | -0.1807 | 0.0547 | 9.62E-04 | -0.0545 | 0.0202 | 7.00E-03 | 1 |
| rs112851448 | 14 | 59466077 | 33 | 59449138 | 59891607 | T | G | 0.05169 | RP11-112J1.2 | ncRNA_intronic | 0.1801 | 0.0262 | 6.26E-12 | 0.1806 | 0.0547 | 9.71E-04 | 0.0541 | 0.0202 | 7.43E-03 | 1 |
| rs61976317 | 14 | 59466439 | 33 | 59449138 | 59891607 | T | C | 0.05169 | RP11-112J1.2 | ncRNA_intronic | -0.1798 | 0.0262 | 6.77E-12 | -0.1807 | 0.0547 | 9.62E-04 | -0.0546 | 0.0202 | 6.83E-03 | 1 |
| rs61976318 | 14 | 59466465 | 33 | 59449138 | 59891607 | A | G | 0.05169 | RP11-112J1.2 | ncRNA_intronic | -0.1804 | 0.0262 | 5.84E-12 | -0.1807 | 0.0547 | 9.62E-04 | -0.0544 | 0.0202 | 7.10E-03 | 1 |
| rs17833470 | 14 | 59466834 | 33 | 59449138 | 59891607 | A | G | 0.05169 | RP11-112J1.2 | ncRNA_intronic | -0.1794 | 0.0262 | 7.02E-12 | -0.1809 | 0.0547 | 9.54E-04 | -0.0555 | 0.0201 | 5.83E-03 | 1 |
| rs117399967 | 14 | 59467126 | 33 | 59449138 | 59891607 | A | G | 0.05169 | RP11-112J1.2 | ncRNA_intronic | -0.1819 | 0.0262 | 4.01E-12 | -0.1775 | 0.0547 | 1.19E-03 | -0.0541 | 0.0202 | 7.25E-03 | 1 |
| rs112688994 | 14 | 59467454 | 33 | 59449138 | 59891607 | A | C | 0.05268 | RP11-112J1.2 | ncRNA_intronic | -0.1801 | 0.0262 | 5.96E-12 | -0.1807 | 0.0547 | 9.51E-04 | -0.0550 | 0.0202 | 6.43E-03 | 1 |
| rs61976319 | 14 | 59468299 | 33 | 59449138 | 59891607 | T | C | 0.05169 | RP11-112J1.2 | ncRNA_intronic | 0.1809 | 0.0263 | 5.81E-12 | 0.1774 | 0.0548 | 1.20E-03 | 0.0541 | 0.0202 | 7.41E-03 | 1 |
| rs77891125 | 14 | 59470430 | 33 | 59449138 | 59891607 | A | G | 0.05169 | RP11-112J1.2 | ncRNA_intronic | -0.1795 | 0.0261 | 6.64E-12 | -0.1806 | 0.0547 | 9.75E-04 | -0.0545 | 0.0201 | 6.69E-03 | 1 |
| rs61976321 | 14 | 59470576 | 33 | 59449138 | 59891607 | A | T | 0.05169 | RP11-112J1.2 | ncRNA_intronic | 0.1794 | 0.0261 | 6.67E-12 | 0.1806 | 0.0547 | 9.74E-04 | 0.0543 | 0.0201 | 6.98E-03 | 1 |
| rs61976322 | 14 | 59471046 | 33 | 59449138 | 59891607 | T | C | 0.05268 | RP11-112J1.2 | ncRNA_intronic | 0.1796 | 0.0261 | 6.35E-12 | 0.1806 | 0.0547 | 9.74E-04 | 0.0543 | 0.0201 | 6.94E-03 | 1 |
| rs117483338 | 14 | 59471529 | 33 | 59449138 | 59891607 | A | T | 0.05268 | RP11-112J1.2 | ncRNA_intronic | -0.1796 | 0.0265 | 1.21E-11 | -0.1805 | 0.0547 | 9.75E-04 | -0.0549 | 0.0202 | 6.59E-03 | 1 |
| rs17095539 | 14 | 59471853 | 33 | 59449138 | 59891607 | T | C | 0.05268 | RP11-112J1.2 | ncRNA_intronic | -0.1792 | 0.0261 | 6.95E-12 | -0.1789 | 0.0547 | 1.08E-03 | -0.0545 | 0.0201 | 6.72E-03 | 1 |
| rs61976330 | 14 | 59471891 | 33 | 59449138 | 59891607 | A | T | 0.05268 | RP11-112J1.2 | ncRNA_intronic | 0.1802 | 0.0261 | 5.23E-12 | 0.1806 | 0.0547 | 9.73E-04 | 0.0545 | 0.0201 | 6.71E-03 | 1 |
| rs61976331 | 14 | 59471909 | 33 | 59449138 | 59891607 | A | G | 0.0507 | RP11-112J1.2 | ncRNA_intronic | 0.1802 | 0.0261 | 5.27E-12 | 0.1806 | 0.0547 | 9.71E-04 | 0.0542 | 0.0201 | 7.02E-03 | 1 |
| rs79690612 | 14 | 59472282 | 33 | 59449138 | 59891607 | T | C | 0.05169 | RP11-112J1.2 | ncRNA_intronic | -0.1792 | 0.0261 | 6.95E-12 | -0.1806 | 0.0547 | 9.74E-04 | -0.0553 | 0.0201 | 6.00E-03 | 1 |
| rs61976332 | 14 | 59472299 | 33 | 59449138 | 59891607 | A | G | 0.05169 | RP11-112J1.2 | ncRNA_intronic | -0.1792 | 0.0261 | 6.95E-12 | -0.1806 | 0.0547 | 9.74E-04 | -0.0553 | 0.0201 | 6.00E-03 | 1 |
| rs61976333 | 14 | 59472831 | 33 | 59449138 | 59891607 | A | C | 0.05268 | RP11-112J1.2 | ncRNA_intronic | 0.1786 | 0.0261 | 7.95E-12 | 0.1828 | 0.0547 | 8.28E-04 | 0.0541 | 0.0201 | 7.10E-03 | 1 |
| rs113935260 | 14 | 59473028 | 33 | 59449138 | 59891607 | T | C | 0.05268 | RP11-112J1.2 | ncRNA_intronic | 0.1722 | 0.0258 | 2.57E-11 | 0.1805 | 0.0547 | 9.76E-04 | 0.0485 | 0.0198 | 1.42E-02 | 1 |
| rs61976335 | 14 | 59473343 | 33 | 59449138 | 59891607 | T | C | 0.05268 | RP11-112J1.2 | ncRNA_intronic | -0.1792 | 0.0261 | 6.88E-12 | -0.1805 | 0.0547 | 9.76E-04 | -0.0545 | 0.0201 | 6.67E-03 | 1 |
| rs59959620 | 14 | 59473840 | 33 | 59449138 | 59891607 | A | C | 0.05268 | RP11-112J1.2 | ncRNA_intronic | -0.1792 | 0.0261 | 6.84E-12 | -0.1805 | 0.0547 | 9.76E-04 | -0.0543 | 0.0201 | 6.81E-03 | 1 |
| rs61976337 | 14 | 59474677 | 33 | 59449138 | 59891607 | C | G | 0.05268 | RP11-112J1.2 | ncRNA_intronic | -0.1792 | 0.0261 | 6.75E-12 | -0.1805 | 0.0547 | 9.77E-04 | -0.0543 | 0.0201 | 6.84E-03 | 1 |
| rs77981231 | 14 | 59475229 | 33 | 59449138 | 59891607 | A | G | 0.05169 | RP11-112J1.2 | ncRNA_intronic | 0.1792 | 0.0261 | 6.73E-12 | 0.1806 | 0.0547 | 9.75E-04 | 0.0543 | 0.0201 | 6.81E-03 | 1 |
| rs61976340 | 14 | 59479216 | 33 | 59449138 | 59891607 | A | G | 0.05268 | RP11-112J1.2 | ncRNA_intronic | 0.1781 | 0.0261 | 8.90E-12 | 0.1802 | 0.0547 | 9.99E-04 | 0.0542 | 0.0200 | 6.86E-03 | 1 |
| rs61976341 | 14 | 59479479 | 33 | 59449138 | 59891607 | T | C | 0.05268 | RP11-112J1.2 | ncRNA_intronic | -0.1793 | 0.0261 | 6.39E-12 | -0.1805 | 0.0547 | 9.81E-04 | -0.0544 | 0.0200 | 6.70E-03 | 1 |
| rs60607947 | 14 | 59479917 | 33 | 59449138 | 59891607 | A | G | 0.05268 | RP11-112J1.2 | ncRNA_intronic | 0.1794 | 0.0261 | 6.29E-12 | 0.1805 | 0.0547 | 9.80E-04 | 0.0541 | 0.0200 | 6.98E-03 | 1 |
| rs58324275 | 14 | 59479983 | 33 | 59449138 | 59891607 | T | C | 0.05368 | RP11-112J1.2 | ncRNA_intronic | 0.1794 | 0.0261 | 6.16E-12 | 0.1800 | 0.0547 | 1.01E-03 | 0.0535 | 0.0200 | 7.60E-03 | 1 |
| rs58966877 | 14 | 59480028 | 33 | 59449138 | 59891607 | T | C | 0.05268 | RP11-112J1.2 | ncRNA_intronic | -0.1795 | 0.0261 | 6.13E-12 | -0.1805 | 0.0547 | 9.81E-04 | -0.0542 | 0.0200 | 6.81E-03 | 1 |
| rs58601019 | 14 | 59480125 | 33 | 59449138 | 59891607 | A | G | 0.05268 | RP11-112J1.2 | ncRNA_intronic | 0.1794 | 0.0261 | 6.28E-12 | 0.1805 | 0.0547 | 9.81E-04 | 0.0542 | 0.0200 | 6.80E-03 | 1 |
| rs61976345 | 14 | 59480320 | 33 | 59449138 | 59891607 | T | C | 0.05268 | RP11-112J1.2 | ncRNA_intronic | -0.1794 | 0.0261 | 6.28E-12 | -0.1805 | 0.0547 | 9.81E-04 | -0.0542 | 0.0200 | 6.80E-03 | 1 |
| rs28464012 | 14 | 59482537 | 33 | 59449138 | 59891607 | T | C | 0.05268 | RP11-112J1.2 | ncRNA_intronic | -0.1794 | 0.0261 | 6.08E-12 | -0.1811 | 0.0547 | 9.41E-04 | -0.0543 | 0.0200 | 6.71E-03 | 1 |
| rs10146779 | 14 | 59483819 | 33 | 59449138 | 59891607 | C | G | 0.05268 | RP11-112J1.2 | ncRNA_intronic | 0.1794 | 0.0261 | 6.19E-12 | 0.1806 | 0.0547 | 9.76E-04 | 0.0542 | 0.0200 | 6.77E-03 | 1 |
| rs17255080 | 14 | 59484207 | 33 | 59449138 | 59891607 | T | G | 0.05169 | RP11-112J1.2 | downstream | -0.1794 | 0.0261 | 6.23E-12 | -0.1806 | 0.0547 | 9.74E-04 | -0.0543 | 0.0200 | 6.73E-03 | 1 |
| rs10130575 | 14 | 59485903 | 33 | 59449138 | 59891607 | A | G | 0.05169 | RP11-112J1.2 | intergenic | -0.1792 | 0.0261 | 6.81E-12 | -0.1815 | 0.0548 | 9.27E-04 | -0.0535 | 0.0200 | 7.57E-03 | 1 |
| rs61976347 | 14 | 59486765 | 33 | 59449138 | 59891607 | A | G | 0.0507 | RP11-112J1.2 | intergenic | -0.1819 | 0.0263 | 5.01E-12 | -0.1758 | 0.0548 | 1.35E-03 | -0.0535 | 0.0203 | 8.39E-03 | 1 |
| rs10142674 | 14 | 59489070 | 33 | 59449138 | 59891607 | C | G | 0.05169 | RP11-112J1.2 | intergenic | 0.1826 | 0.0263 | 3.76E-12 | 0.1852 | 0.0549 | 7.51E-04 | 0.0538 | 0.0203 | 8.04E-03 | 1 |
| rs10132018 | 14 | 59491212 | 33 | 59449138 | 59891607 | T | C | 0.05268 | RP11-112J1.2 | intergenic | -0.1845 | 0.0262 | 1.98E-12 | -0.1913 | 0.0548 | 4.82E-04 | -0.0584 | 0.0202 | 3.74E-03 | 1 |
| rs111676402 | 14 | 59505196 | 33 | 59449138 | 59891607 | T | C | 0.0507 | RP11-112J1.2 | intergenic | -0.184 | 0.0266 | 4.48E-12 | -0.2001 | 0.0564 | 3.88E-04 | -0.0641 | 0.0206 | 1.83E-03 | 1 |
| rs79741821 | 14 | 59506360 | 33 | 59449138 | 59891607 | A | C | 0.0507 | RP11-112J1.2 | intergenic | -0.1856 | 0.0266 | 3.30E-12 | -0.1985 | 0.0565 | 4.49E-04 | -0.0640 | 0.0206 | 1.87E-03 | 1 |
| rs79924383 | 14 | 59506790 | 33 | 59449138 | 59891607 | A | G | 0.0507 | RP11-112J1.2 | intergenic | -0.1848 | 0.0267 | 4.21E-12 | -0.2027 | 0.0567 | 3.54E-04 | -0.0640 | 0.0206 | 1.88E-03 | 1 |
| rs75951218 | 14 | 59510280 | 33 | 59449138 | 59891607 | A | T | 0.0497 | RP11-112J1.2 | intergenic | 0.1848 | 0.027 | 7.23E-12 | 0.2063 | 0.0567 | 2.77E-04 | 0.0644 | 0.0207 | 1.83E-03 | 1 |
| rs17255157 | 14 | 59515491 | 33 | 59449138 | 59891607 | T | G | 0.04871 | RP11-112J1.2 | intergenic | 0.1845 | 0.0269 | 6.87E-12 | 0.2039 | 0.0567 | 3.24E-04 | 0.0641 | 0.0207 | 1.91E-03 | 1 |
| rs79830598 | 14 | 59517474 | 33 | 59449138 | 59891607 | A | G | 0.0497 | RP11-112J1.2 | intergenic | 0.1857 | 0.0269 | 4.94E-12 | 0.2039 | 0.0567 | 3.25E-04 | 0.0653 | 0.0206 | 1.56E-03 | 1 |
| rs17833596 | 14 | 59522340 | 33 | 59449138 | 59891607 | A | T | 0.0497 | RP11-112J1.2 | intergenic | 0.1878 | 0.027 | 3.28E-12 | 0.2078 | 0.0566 | 2.45E-04 | 0.0656 | 0.0206 | 1.47E-03 | 1 |
| rs17255192 | 14 | 59523309 | 33 | 59449138 | 59891607 | A | G | 0.0497 | RP11-112J1.2 | intergenic | 0.1861 | 0.0269 | 4.43E-12 | 0.2078 | 0.0566 | 2.45E-04 | 0.0657 | 0.0206 | 1.47E-03 | 1 |
| rs17833602 | 14 | 59524433 | 33 | 59449138 | 59891607 | T | C | 0.0497 | RP11-112J1.2 | intergenic | -0.1861 | 0.0269 | 4.47E-12 | -0.2078 | 0.0566 | 2.45E-04 | -0.0656 | 0.0206 | 1.48E-03 | 1 |
| rs77542809 | 14 | 59524583 | 33 | 59449138 | 59891607 | A | G | 0.0497 | RP11-112J1.2 | intergenic | 0.186 | 0.0269 | 4.49E-12 | 0.2078 | 0.0566 | 2.45E-04 | 0.0656 | 0.0206 | 1.48E-03 | 1 |
| rs79187540 | 14 | 59527301 | 33 | 59449138 | 59891607 | C | G | 0.0497 | RP11-112J1.2 | intergenic | 0.1861 | 0.0269 | 4.47E-12 | 0.2078 | 0.0566 | 2.45E-04 | 0.0653 | 0.0206 | 1.57E-03 | 1 |
| rs112313693 | 14 | 59528311 | 33 | 59449138 | 59891607 | A | G | 0.0497 | RP11-112J1.2 | intergenic | 0.1858 | 0.0269 | 4.85E-12 | 0.2076 | 0.0567 | 2.50E-04 | 0.0652 | 0.0206 | 1.59E-03 | 1 |
| rs74749999 | 14 | 59528959 | 33 | 59449138 | 59891607 | A | G | 0.0497 | RP11-112J1.2 | intergenic | -0.186 | 0.0269 | 4.55E-12 | -0.2077 | 0.0566 | 2.46E-04 | -0.0653 | 0.0206 | 1.57E-03 | 1 |
| rs17833614 | 14 | 59531935 | 33 | 59449138 | 59891607 | T | C | 0.0497 | RP11-112J1.2 | intergenic | 0.186 | 0.0269 | 4.61E-12 | 0.2078 | 0.0566 | 2.46E-04 | 0.0652 | 0.0206 | 1.60E-03 | 1 |
| rs75468739 | 14 | 59535773 | 33 | 59449138 | 59891607 | A | G | 0.0497 | RP11-112J1.2 | intergenic | -0.1859 | 0.0269 | 4.67E-12 | -0.2078 | 0.0566 | 2.45E-04 | -0.0651 | 0.0206 | 1.61E-03 | 1 |
| rs17255213 | 14 | 59540008 | 33 | 59449138 | 59891607 | T | C | 0.0497 | RP11-112J1.2 | intergenic | -0.1848 | 0.0269 | 6.40E-12 | -0.2176 | 0.0568 | 1.27E-04 | -0.0652 | 0.0207 | 1.59E-03 | 1 |
| rs17095660 | 14 | 59541935 | 33 | 59449138 | 59891607 | T | G | 0.06163 | RP11-112J1.2 | intergenic | 0.1424 | 0.0245 | 5.92E-09 | 0.1809 | 0.0513 | 4.20E-04 | 0.0559 | 0.0189 | 3.09E-03 | 1 |
| rs17255220 | 14 | 59543877 | 33 | 59449138 | 59891607 | A | G | 0.0497 | CTD-2315A10.2 | intergenic | 0.1849 | 0.0269 | 6.30E-12 | 0.2178 | 0.0568 | 1.26E-04 | 0.0657 | 0.0207 | 1.48E-03 | 1 |
| rs74746058 | 14 | 59544253 | 33 | 59449138 | 59891607 | T | C | 0.0497 | CTD-2315A10.2 | intergenic | -0.1852 | 0.0269 | 5.90E-12 | -0.2178 | 0.0568 | 1.26E-04 | -0.0666 | 0.0207 | 1.28E-03 | 1 |
| rs78091436 | 14 | 59544796 | 33 | 59449138 | 59891607 | A | G | 0.0497 | CTD-2315A10.2 | intergenic | 0.1854 | 0.0269 | 5.74E-12 | 0.2178 | 0.0568 | 1.28E-04 | 0.0665 | 0.0207 | 1.31E-03 | 1 |
| rs17833632 | 14 | 59547738 | 33 | 59449138 | 59891607 | A | G | 0.0497 | CTD-2315A10.2 | intergenic | -0.1857 | 0.0269 | 5.17E-12 | -0.2180 | 0.0568 | 1.25E-04 | -0.0666 | 0.0207 | 1.29E-03 | 1 |
| rs17833651 | 14 | 59548902 | 33 | 59449138 | 59891607 | T | C | 0.0497 | CTD-2315A10.2 | intergenic | -0.1864 | 0.0269 | 4.40E-12 | -0.2179 | 0.0568 | 1.25E-04 | -0.0662 | 0.0207 | 1.38E-03 | 1 |
| rs17255241 | 14 | 59549260 | 33 | 59449138 | 59891607 | T | C | 0.0497 | CTD-2315A10.2 | intergenic | -0.1866 | 0.0269 | 4.18E-12 | -0.2179 | 0.0568 | 1.25E-04 | -0.0663 | 0.0207 | 1.35E-03 | 1 |
| rs75666025 | 14 | 59553059 | 33 | 59449138 | 59891607 | A | G | 0.0497 | CTD-2315A10.2 | intergenic | 0.1892 | 0.0269 | 2.17E-12 | 0.2193 | 0.0569 | 1.16E-04 | 0.0676 | 0.0207 | 1.10E-03 | 1 |
| rs76185024 | 14 | 59556566 | 33 | 59449138 | 59891607 | A | T | 0.04672 | CTD-2315A10.2 | intergenic | -0.1947 | 0.027 | 5.74E-13 | -0.2191 | 0.0573 | 1.33E-04 | -0.0753 | 0.0208 | 2.92E-04 | 1 |

| | | | | | | | | | | | Meta-GWAS | | | UKBB | | | ENIGMA* | | | |
|---|---|---|---|---|---|---|---|---|---|---|---|---|---|---|---|---|---|---|---|---|
| rsID | chr | pos (hg19) | Genomic Locus | start | end | A1 | A2 | MAF | Nearest Genes | Function | Beta | SE | P-value | Beta | SE | P-value | Beta | SE | P-value | Replicated** |
| rs74742608 | 14 | 59556665 | 33 | 59449138 | 59891607 | A | G | 0.04672 | CTD-2315A10.2 | intergenic | -0.1942 | 0.027 | 7.04E-13 | -0.2192 | 0.0573 | 1.32E-04 | -0.0742 | 0.0208 | 3.56E-04 | 1 |
| rs79633637 | 14 | 59556931 | 33 | 59449138 | 59891607 | A | C | 0.04672 | CTD-2315A10.2 | intergenic | 0.1943 | 0.027 | 6.83E-13 | 0.2192 | 0.0573 | 1.32E-04 | 0.0739 | 0.0208 | 3.74E-04 | 1 |
| rs76045330 | 14 | 59557043 | 33 | 59449138 | 59891607 | T | C | 0.04672 | CTD-2315A10.2 | intergenic | 0.1944 | 0.027 | 6.68E-13 | 0.2192 | 0.0573 | 1.32E-04 | 0.0746 | 0.0208 | 3.34E-04 | 1 |
| rs17255248 | 14 | 59557419 | 33 | 59449138 | 59891607 | T | G | 0.04672 | CTD-2315A10.2 | intergenic | 0.1945 | 0.027 | 6.39E-13 | 0.2191 | 0.0573 | 1.32E-04 | 0.0743 | 0.0208 | 3.50E-04 | 1 |
| rs77381622 | 14 | 59557616 | 33 | 59449138 | 59891607 | T | C | 0.04672 | CTD-2315A10.2 | intergenic | -0.1953 | 0.027 | 5.07E-13 | -0.2219 | 0.0572 | 1.07E-04 | -0.0744 | 0.0208 | 3.42E-04 | 1 |
| rs76443390 | 14 | 59557958 | 33 | 59449138 | 59891607 | A | G | 0.04672 | CTD-2315A10.2 | intergenic | 0.1945 | 0.027 | 6.45E-13 | 0.2191 | 0.0573 | 1.32E-04 | 0.0749 | 0.0208 | 3.15E-04 | 1 |
| rs78616162 | 14 | 59558330 | 33 | 59449138 | 59891607 | T | C | 0.04672 | CTD-2315A10.2 | intergenic | 0.1946 | 0.027 | 6.25E-13 | 0.2191 | 0.0573 | 1.32E-04 | 0.0743 | 0.0208 | 3.50E-04 | 1 |
| rs75499021 | 14 | 59558415 | 33 | 59449138 | 59891607 | T | C | 0.04672 | CTD-2315A10.2 | intergenic | 0.1946 | 0.027 | 6.21E-13 | 0.2191 | 0.0573 | 1.32E-04 | 0.0751 | 0.0208 | 3.02E-04 | 1 |
| rs75035809 | 14 | 59558527 | 33 | 59449138 | 59891607 | A | G | 0.04672 | CTD-2315A10.2 | intergenic | -0.1952 | 0.027 | 5.36E-13 | -0.2193 | 0.0572 | 1.28E-04 | -0.0756 | 0.0208 | 2.76E-04 | 1 |
| rs78090044 | 14 | 59558567 | 33 | 59449138 | 59891607 | A | G | 0.04672 | CTD-2315A10.2 | intergenic | 0.1946 | 0.027 | 6.20E-13 | 0.2191 | 0.0573 | 1.32E-04 | 0.0752 | 0.0208 | 2.96E-04 | 1 |
| rs1954003 | 14 | 59558892 | 33 | 59449138 | 59891607 | A | C | 0.04672 | CTD-2315A10.2 | intergenic | -0.1946 | 0.027 | 6.19E-13 | -0.2191 | 0.0573 | 1.32E-04 | -0.0747 | 0.0208 | 3.24E-04 | 1 |
| rs75923883 | 14 | 59559117 | 33 | 59449138 | 59891607 | A | G | 0.04672 | CTD-2315A10.2 | intergenic | 0.1947 | 0.027 | 6.18E-13 | 0.2191 | 0.0573 | 1.32E-04 | 0.0748 | 0.0208 | 3.22E-04 | 1 |
| rs117513981 | 14 | 59559779 | 33 | 59449138 | 59891607 | A | G | 0.04672 | CTD-2315A10.2 | intergenic | 0.1947 | 0.0271 | 6.19E-13 | 0.2200 | 0.0573 | 1.25E-04 | 0.0738 | 0.0208 | 3.83E-04 | 1 |
| rs117804823 | 14 | 59559797 | 33 | 59449138 | 59891607 | T | C | 0.04672 | CTD-2315A10.2 | intergenic | -0.1946 | 0.0271 | 6.24E-13 | -0.2200 | 0.0573 | 1.25E-04 | -0.0754 | 0.0208 | 2.85E-04 | 1 |
| rs75012551 | 14 | 59560013 | 33 | 59449138 | 59891607 | A | G | 0.04672 | CTD-2315A10.2 | intergenic | 0.1946 | 0.0271 | 6.31E-13 | 0.2200 | 0.0573 | 1.25E-04 | 0.0754 | 0.0208 | 2.87E-04 | 1 |
| rs78339200 | 14 | 59560893 | 33 | 59449138 | 59891607 | T | C | 0.04672 | CTD-2315A10.2 | intergenic | 0.1946 | 0.0271 | 6.31E-13 | 0.2200 | 0.0573 | 1.25E-04 | 0.0755 | 0.0208 | 2.80E-04 | 1 |
| rs75514756 | 14 | 59561825 | 33 | 59449138 | 59891607 | T | C | 0.04672 | CTD-2315A10.2 | intergenic | 0.1922 | 0.027 | 1.12E-12 | 0.2200 | 0.0573 | 1.25E-04 | 0.0764 | 0.0207 | 2.23E-04 | 1 |
| rs78035000 | 14 | 59562075 | 33 | 59449138 | 59891607 | T | C | 0.04672 | CTD-2315A10.2 | intergenic | -0.1947 | 0.0271 | 6.25E-13 | -0.2200 | 0.0573 | 1.25E-04 | -0.0772 | 0.0208 | 2.08E-04 | 1 |
| rs117381419 | 14 | 59563176 | 33 | 59449138 | 59891607 | A | G | 0.04672 | CTD-2315A10.2 | intergenic | 0.1949 | 0.0271 | 6.07E-13 | 0.2199 | 0.0573 | 1.26E-04 | 0.0765 | 0.0208 | 2.39E-04 | 1 |
| rs139493105 | 14 | 59563613 | 33 | 59449138 | 59891607 | T | C | 0.04672 | CTD-2315A10.2 | intergenic | -0.1899 | 0.0271 | 2.39E-12 | -0.2199 | 0.0573 | 1.27E-04 | -0.0729 | 0.0211 | 5.68E-04 | 1 |
| rs17833663 | 14 | 59565263 | 33 | 59449138 | 59891607 | T | C | 0.04672 | CTD-2315A10.2 | intergenic | -0.195 | 0.0271 | 6.31E-13 | -0.2151 | 0.0573 | 1.77E-04 | -0.0766 | 0.0208 | 2.33E-04 | 1 |
| rs76048834 | 14 | 59566092 | 33 | 59449138 | 59891607 | A | G | 0.04672 | CTD-2315A10.2 | intergenic | -0.1954 | 0.0271 | 5.87E-13 | -0.2150 | 0.0574 | 1.84E-04 | -0.0764 | 0.0208 | 2.44E-04 | 1 |
| rs76751566 | 14 | 59567311 | 33 | 59449138 | 59891607 | A | G | 0.04771 | CTD-2315A10.2 | intergenic | -0.1943 | 0.0271 | 7.72E-13 | -0.2115 | 0.0573 | 2.25E-04 | -0.0774 | 0.0208 | 2.01E-04 | 1 |
| rs74946438 | 14 | 59570172 | 33 | 59449138 | 59891607 | T | C | 0.04672 | CTD-2315A10.2 | intergenic | -0.1951 | 0.0272 | 7.04E-13 | -0.2137 | 0.0575 | 2.04E-04 | -0.0771 | 0.0208 | 2.16E-04 | 1 |
| rs79999583 | 14 | 59573508 | 33 | 59449138 | 59891607 | A | G | 0.04672 | CTD-2315A10.2 | intergenic | 0.195 | 0.0272 | 7.46E-13 | 0.2139 | 0.0575 | 2.02E-04 | 0.0770 | 0.0208 | 2.19E-04 | 1 |
| rs78570253 | 14 | 59577898 | 33 | 59449138 | 59891607 | T | C | 0.05666 | CTD-2315A10.2 | intergenic | -0.2027 | 0.026 | 6.21E-15 | -0.2214 | 0.0548 | 5.39E-05 | -0.0782 | 0.0200 | 9.03E-05 | 1 |
| rs78601235 | 14 | 59578878 | 33 | 59449138 | 59891607 | A | C | 0.07256 | CTD-2315A10.2 | intergenic | -0.2003 | 0.0259 | 1.07E-14 | -0.2166 | 0.0545 | 7.22E-05 | -0.0756 | 0.0199 | 1.46E-04 | 1 |
| rs145409503 | 14 | 59579984 | 33 | 59449138 | 59891607 | T | C | 0.04672 | CTD-2315A10.2 | intergenic | -0.1955 | 0.0273 | 7.26E-13 | -0.2143 | 0.0575 | 1.98E-04 | -0.0767 | 0.0209 | 2.45E-04 | 1 |
| rs17095711 | 14 | 59580311 | 33 | 59449138 | 59891607 | A | G | 0.05765 | CTD-2315A10.2 | intergenic | -0.1996 | 0.0259 | 1.34E-14 | -0.2165 | 0.0545 | 7.24E-05 | -0.0769 | 0.0199 | 1.15E-04 | 1 |
| rs17255290 | 14 | 59585670 | 33 | 59449138 | 59891607 | A | T | 0.1998 | CTD-2315A10.2 | intergenic | 0.1561 | 0.0175 | 5.12E-19 | 0.1350 | 0.0371 | 2.76E-04 | 0.0511 | 0.0132 | 1.14E-04 | 1 |
| rs4898978 | 14 | 59585932 | 33 | 59449138 | 59891607 | A | C | 0.1859 | CTD-2315A10.2 | intergenic | -0.1669 | 0.018 | 1.75E-20 | -0.1533 | 0.0390 | 8.52E-05 | -0.0567 | 0.0135 | 2.55E-05 | 1 |
| rs10498487 | 14 | 59588323 | 33 | 59449138 | 59891607 | T | C | 0.1491 | CTD-2315A10.2 | intergenic | 0.1965 | 0.019 | 3.60E-25 | 0.1760 | 0.0406 | 1.48E-05 | 0.0913 | 0.0143 | 1.68E-10 | 1 |
| rs72724695 | 14 | 59590174 | 33 | 59449138 | 59891607 | A | G | 0.1491 | CTD-2315A10.2 | intergenic | 0.1969 | 0.019 | 3.20E-25 | 0.1746 | 0.0406 | 1.71E-05 | 0.0903 | 0.0143 | 2.90E-10 | 1 |
| rs17833704 | 14 | 59591082 | 33 | 59449138 | 59891607 | A | G | 0.1889 | CTD-2315A10.2 | intergenic | 0.1608 | 0.0178 | 1.94E-19 | 0.1521 | 0.0383 | 7.33E-05 | 0.0595 | 0.0133 | 7.78E-06 | 1 |
| rs72724697 | 14 | 59591624 | 33 | 59449138 | 59891607 | T | C | 0.1471 | CTD-2315A10.2 | intergenic | 0.2002 | 0.0191 | 1.05E-25 | 0.1709 | 0.0409 | 2.92E-05 | 0.0873 | 0.0145 | 1.57E-09 | 1 |
| rs72724702 | 14 | 59593916 | 33 | 59449138 | 59891607 | A | C | 0.1451 | CTD-2315A10.2 | intergenic | -0.2026 | 0.0192 | 3.98E-26 | -0.1748 | 0.0412 | 2.22E-05 | -0.0874 | 0.0145 | 1.69E-09 | 1 |
| rs55643369 | 14 | 59595245 | 33 | 59449138 | 59891607 | T | C | 0.1859 | CTD-2315A10.2 | intergenic | 0.1671 | 0.018 | 1.82E-20 | 0.1529 | 0.0389 | 8.61E-05 | 0.0569 | 0.0135 | 2.48E-05 | 1 |
| rs17255304 | 14 | 59597337 | 33 | 59449138 | 59891607 | A | G | 0.1461 | CTD-2315A10.2 | intergenic | -0.2008 | 0.0191 | 9.97E-26 | -0.1833 | 0.0410 | 7.94E-06 | -0.0916 | 0.0144 | 2.13E-10 | 1 |
| rs72726305 | 14 | 59597408 | 33 | 59449138 | 59891607 | A | G | 0.1461 | CTD-2315A10.2 | intergenic | -0.2008 | 0.0191 | 9.80E-26 | -0.1832 | 0.0410 | 8.06E-06 | -0.0920 | 0.0144 | 1.82E-10 | 1 |
| rs17255311 | 14 | 59598931 | 33 | 59449138 | 59891607 | T | C | 0.1382 | CTD-2315A10.2 | intergenic | -0.2044 | 0.0194 | 5.05E-26 | -0.1963 | 0.0414 | 2.15E-06 | -0.0903 | 0.0146 | 6.59E-10 | 1 |
| rs77168169 | 14 | 59601064 | 33 | 59449138 | 59891607 | A | C | 0.1342 | CTD-2315A10.2 | intergenic | -0.2193 | 0.0198 | 2.07E-28 | -0.1939 | 0.0424 | 5.04E-06 | -0.0974 | 0.0150 | 7.63E-11 | 1 |
| rs4901898 | 14 | 59602914 | 33 | 59449138 | 59891607 | C | G | 0.1342 | CTD-2315A10.2 | ncRNA_exonic | -0.2199 | 0.0199 | 1.63E-28 | -0.1906 | 0.0424 | 6.95E-06 | -0.0983 | 0.0150 | 5.46E-11 | 1 |
| rs117461235 | 14 | 59605116 | 33 | 59449138 | 59891607 | A | G | 0.1372 | CTD-2315A10.2 | intergenic | -0.2174 | 0.0198 | 4.54E-28 | -0.2056 | 0.0426 | 1.44E-06 | -0.0961 | 0.0148 | 7.56E-11 | 1 |
| rs4258526 | 14 | 59609282 | 33 | 59449138 | 59891607 | T | G | 0.2763 | CTD-2315A10.2 | intergenic | 0.0881 | 0.0142 | 5.34E-10 | 0.0244 | 0.0302 | 4.18E-01 | 0.0373 | 0.0110 | 6.69E-04 | 0 |
| rs78580207 | 14 | 59610076 | 33 | 59449138 | 59891607 | A | T | 0.1342 | CTD-2315A10.2 | intergenic | 0.2214 | 0.0196 | 1.52E-29 | 0.1928 | 0.0420 | 4.62E-06 | 0.1025 | 0.0149 | 5.52E-12 | 1 |
| rs10498489 | 14 | 59610460 | 33 | 59449138 | 59891607 | T | G | 0.1342 | CTD-2315A10.2 | intergenic | 0.2214 | 0.0196 | 1.56E-29 | 0.1928 | 0.0420 | 4.65E-06 | 0.1022 | 0.0149 | 6.51E-12 | 1 |
| rs17255332 | 14 | 59611735 | 33 | 59449138 | 59891607 | C | G | 0.1352 | CTD-2315A10.2 | intergenic | -0.2216 | 0.0196 | 1.14E-29 | -0.1940 | 0.0421 | 4.10E-06 | -0.1030 | 0.0149 | 4.39E-12 | 1 |
| rs78032956 | 14 | 59612317 | 33 | 59449138 | 59891607 | T | C | 0.1332 | CTD-2315A10.2 | intergenic | -0.2221 | 0.0196 | 1.14E-29 | -0.1940 | 0.0421 | 4.19E-06 | -0.1029 | 0.0149 | 4.38E-12 | 1 |
| rs79676525 | 14 | 59612800 | 33 | 59449138 | 59891607 | A | T | 0.1332 | CTD-2315A10.2 | intergenic | 0.2221 | 0.0196 | 1.18E-29 | 0.1943 | 0.0421 | 4.09E-06 | 0.1026 | 0.0149 | 5.09E-12 | 1 |
| rs1438518 | 14 | 59615848 | 33 | 59449138 | 59891607 | A | T | 0.3161 | CTD-2315A10.2 | intergenic | 0.0743 | 0.0136 | 4.25E-08 | 0.0144 | 0.0289 | 6.17E-01 | 0.0321 | 0.0105 | 2.15E-03 | 0 |
| rs11626333 | 14 | 59616097 | 33 | 59449138 | 59891607 | T | C | 0.3161 | CTD-2315A10.2 | intergenic | -0.0742 | 0.0136 | 4.46E-08 | -0.0144 | 0.0289 | 6.18E-01 | -0.0325 | 0.0105 | 1.92E-03 | 0 |
| rs4901902 | 14 | 59617080 | 33 | 59449138 | 59891607 | A | C | 0.1352 | CTD-2315A10.2 | intergenic | 0.2259 | 0.0195 | 5.43E-31 | 0.2000 | 0.0419 | 1.83E-06 | 0.1032 | 0.0148 | 3.18E-12 | 1 |
| rs8005394 | 14 | 59619564 | 33 | 59449138 | 59891607 | C | G | 0.1352 | CTD-2315A10.2 | intergenic | 0.2259 | 0.0195 | 4.75E-31 | 0.2020 | 0.0417 | 1.33E-06 | 0.1030 | 0.0148 | 3.20E-12 | 1 |
| rs17833752 | 14 | 59621080 | 33 | 59449138 | 59891607 | A | G | 0.1352 | CTD-2315A10.2 | intergenic | 0.2264 | 0.0195 | 4.26E-31 | 0.1984 | 0.0419 | 2.22E-06 | 0.1032 | 0.0148 | 3.24E-12 | 1 |
| rs76119478 | 14 | 59622767 | 33 | 59449138 | 59891607 | A | C | 0.04573 | CTD-2315A10.2 | intergenic | 0.176 | 0.0306 | 9.03E-09 | 0.1491 | 0.0638 | 1.93E-02 | 0.1084 | 0.0240 | 6.59E-06 | 1 |
| rs4901904 | 14 | 59624317 | 33 | 59449138 | 59891607 | T | C | 0.1342 | CTD-2315A10.2 | intergenic | -0.2271 | 0.0196 | 3.74E-31 | -0.2002 | 0.0419 | 1.79E-06 | -0.1024 | 0.0148 | 5.33E-12 | 1 |
| rs11158250 | 14 | 59625980 | 33 | 59449138 | 59891607 | A | G | 0.2107 | CTD-2315A10.1 | intergenic | 0.1414 | 0.0164 | 5.57E-18 | 0.1479 | 0.0347 | 2.12E-05 | 0.0738 | 0.0125 | 3.39E-09 | 1 |
| rs73313052 | 14 | 59625997 | 33 | 59449138 | 59891607 | A | G | 0.1302 | CTD-2315A10.1 | intergenic | -0.2388 | 0.0195 | 2.36E-34 | -0.2184 | 0.0421 | 2.19E-07 | -0.1139 | 0.0148 | 1.63E-14 | 1 |
| rs2164950 | 14 | 59627631 | 33 | 59449138 | 59891607 | A | G | 0.1312 | CTD-2315A10.1 | intergenic | -0.2361 | 0.0195 | 9.21E-34 | -0.2096 | 0.0419 | 5.77E-07 | -0.1119 | 0.0148 | 4.70E-14 | 1 |
| rs74826997 | 14 | 59628609 | 33 | 59449138 | 59891607 | T | C | 0.1312 | CTD-2315A10.1 | intergenic | 0.238 | 0.0195 | 3.31E-34 | 0.2138 | 0.0419 | 3.55E-07 | 0.1126 | 0.0148 | 3.34E-14 | 1 |
| rs76341705 | 14 | 59628679 | 33 | 59449138 | 59891607 | A | G | 0.1312 | CTD-2315A10.1 | intergenic | -0.2374 | 0.0195 | 4.59E-34 | -0.2146 | 0.0419 | 3.18E-07 | -0.1123 | 0.0149 | 4.11E-14 | 1 |
| rs2053300 | 14 | 59629611 | 33 | 59449138 | 59891607 | T | G | 0.4583 | CTD-2315A10.1 | intergenic | 0.0737 | 0.0132 | 2.52E-08 | 0.0972 | 0.0275 | 4.19E-04 | 0.0354 | 0.0102 | 4.97E-04 | 1 |
| rs4898980 | 14 | 59633677 | 33 | 59449138 | 59891607 | A | G | 0.02684 | CTD-2315A10.1 | intergenic | -0.27 | 0.0493 | 4.46E-08 | -0.0836 | 0.0994 | 4.00E-01 | -0.1502 | 0.0391 | 1.23E-04 | 0 |
| rs1252914 | 14 | 59636409 | 33 | 59449138 | 59891607 | T | C | 0.3171 | CTD-2315A10.1 | intergenic | 0.1354 | 0.0142 | 1.62E-21 | 0.1124 | 0.0300 | 1.81E-04 | 0.0540 | 0.0108 | 6.13E-07 | 1 |
| rs79814107 | 14 | 59637503 | 33 | 59449138 | 59891607 | A | G | 0.1312 | CTD-2315A10.1 | intergenic | 0.2242 | 0.0197 | 5.57E-30 | 0.2026 | 0.0424 | 1.83E-06 | 0.1020 | 0.0149 | 7.43E-12 | 1 |
| rs113269922 | 14 | 59638174 | 33 | 59449138 | 59891607 | A | G | 0.1312 | CTD-2315A10.1 | intergenic | 0.2242 | 0.0197 | 5.69E-30 | 0.2025 | 0.0424 | 1.86E-06 | 0.1019 | 0.0149 | 7.64E-12 | 1 |
| rs56115079 | 14 | 59642065 | 33 | 59449138 | 59891607 | T | G | 0.1312 | CTD-2315A10.1 | intergenic | 0.2222 | 0.0197 | 1.72E-29 | 0.1993 | 0.0424 | 2.66E-06 | 0.1014 | 0.0149 | 9.42E-12 | 1 |
| rs112195026 | 14 | 59643002 | 33 | 59449138 | 59891607 | A | G | 0.1312 | CTD-2315A10.1 | intergenic | 0.2226 | 0.0197 | 1.34E-29 | 0.2022 | 0.0425 | 1.94E-06 | 0.1010 | 0.0149 | 1.08E-11 | 1 |
| rs76850797 | 14 | 59643746 | 33 | 59449138 | 59891607 | T | C | 0.1312 | CTD-2315A10.1 | intergenic | 0.2216 | 0.0197 | 2.26E-29 | 0.1993 | 0.0424 | 2.64E-06 | 0.1012 | 0.0149 | 9.84E-12 | 1 |
| rs17095819 | 14 | 59643961 | 33 | 59449138 | 59891607 | A | G | 0.165 | CTD-2315A10.1 | intergenic | 0.1804 | 0.0178 | 5.01E-24 | 0.1373 | 0.0382 | 3.25E-04 | 0.0837 | 0.0136 | 8.11E-10 | 1 |
| rs78349529 | 14 | 59644286 | 33 | 59449138 | 59891607 | A | G | 0.1312 | CTD-2315A10.1 | intergenic | 0.2227 | 0.0197 | 1.27E-29 | 0.2024 | 0.0424 | 1.91E-06 | 0.1009 | 0.0149 | 1.10E-11 | 1 |
| rs28673301 | 14 | 59644498 | 33 | 59449138 | 59891607 | T | C | 0.163 | CTD-2315A10.1 | intergenic | 0.1806 | 0.0177 | 1.85E-24 | 0.1354 | 0.0381 | 3.89E-04 | 0.0799 | 0.0135 | 3.01E-09 | 1 |
| rs893516 | 14 | 59648738 | 33 | 59449138 | 59891607 | T | C | 0.3211 | CTD-2315A10.1 | downstream | 0.1354 | 0.0142 | 1.06E-21 | 0.1081 | 0.0300 | 3.12E-04 | 0.0527 | 0.0107 | 9.39E-07 | 1 |
| rs1438521 | 14 | 59653800 | 33 | 59449138 | 59891607 | T | C | 0.167 | DAAM1 | intergenic | 0.1794 | 0.0177 | 3.34E-24 | 0.1264 | 0.0382 | 9.33E-04 | 0.0790 | 0.0134 | 3.86E-09 | 1 |
| rs4901906 | 14 | 59654649 | 33 | 59449138 | 59891607 | T | C | 0.1312 | DAAM1 | upstream | -0.2222 | 0.0197 | 1.63E-29 | -0.2010 | 0.0425 | 2.31E-06 | -0.1000 | 0.0148 | 1.50E-11 | 1 |
| rs79360654 | 14 | 59658600 | 33 | 59449138 | 59891607 | A | C | 0.1292 | DAAM1 | intronic | -0.2156 | 0.0197 | 6.90E-28 | -0.2014 | 0.0421 | 1.72E-06 | -0.0978 | 0.0149 | 4.67E-11 | 1 |
| rs11844752 | 14 | 59658725 | 33 | 59449138 | 59891607 | A | G | 0.4632 | DAAM1 | intronic | 0.088 | 0.0131 | 1.66E-11 | 0.0281 | 0.0276 | 3.09E-01 | 0.0317 | 0.0099 | 1.45E-03 | 0 |
| NA | 14 | 59658860 | 33 | 59449138 | 59891607 | C | G | 0.04573 | DAAM1 | intronic | -0.2745 | 0.0457 | 1.97E-09 | -0.2052 | 0.0723 | 4.54E-03 | -0.1011 | 0.0280 | 3.00E-04 | 1 |
| rs77618329 | 14 | 59660552 | 33 | 59449138 | 59891607 | T | C | 0.1292 | DAAM1 | intronic | -0.2161 | 0.0197 | 5.44E-28 | -0.2014 | 0.0421 | 1.71E-06 | -0.0979 | 0.0149 | 4.49E-11 | 1 |
| rs76256424 | 14 | 59661266 | 33 | 59449138 | 59891607 | C | G | 0.1292 | DAAM1 | intronic | 0.2156 | 0.0197 | 7.13E-28 | 0.2014 | 0.0421 | 1.72E-06 | 0.0978 | 0.0149 | 4.50E-11 | 1 |
| rs76674533 | 14 | 59661566 | 33 | 59449138 | 59891607 | T | C | 0.1292 | DAAM1 | intronic | -0.2156 | 0.0197 | 7.33E-28 | -0.2013 | 0.0421 | 1.73E-06 | -0.0976 | 0.0148 | 4.88E-11 | 1 |
| rs143720575 | 14 | 59663805 | 33 | 59449138 | 59891607 | A | G | 0.1292 | DAAM1 | intronic | -0.2246 | 0.0201 | 7.25E-29 | -0.2013 | 0.0421 | 1.74E-06 | -0.1030 | 0.0155 | 2.81E-11 | 1 |
| rs1252916 | 14 | 59664984 | 33 | 59449138 | 59891607 | T | C | 0.4483 | DAAM1 | intronic | 0.0748 | 0.0135 | 3.02E-08 | 0.0989 | 0.0279 | 3.89E-04 | 0.0226 | 0.0104 | 3.00E-02 | 1 |
| rs8016570 | 14 | 59665934 | 33 | 59449138 | 59891607 | T | C | 0.4702 | DAAM1 | intronic | 0.0888 | 0.0131 | 1.02E-11 | 0.0355 | 0.0275 | 1.97E-01 | 0.0305 | 0.0099 | 2.05E-03 | 0 |
| rs78419480 | 14 | 59666074 | 33 | 59449138 | 59891607 | C | G | 0.1233 | DAAM1 | intronic | 0.2073 | 0.0202 | 8.64E-25 | 0.1929 | 0.0427 | 6.52E-06 | 0.1000 | 0.0152 | 4.21E-11 | 1 |
| rs75255901 | 14 | 59669037 | 33 | 59449138 | 59891607 | T | C | 0.1332 | DAAM1 | intronic | -0.2 | 0.0194 | 7.17E-25 | -0.1803 | 0.0409 | 1.08E-05 | -0.0930 | 0.0146 | 1.90E-10 | 1 |
| rs76787422 | 14 | 59669305 | 33 | 59449138 | 59891607 | T | C | 0.1332 | DAAM1 | intronic | -0.2003 | 0.0194 | 6.56E-25 | -0.1806 | 0.0410 | 1.06E-05 | -0.0929 | 0.0146 | 1.98E-10 | 1 |
| rs17255395 | 14 | 59669394 | 33 | 59449138 | 59891607 | T | C | 0.1322 | DAAM1 | intronic | -0.1998 | 0.0194 | 8.33E-25 | -0.1806 | 0.0410 | 1.05E-05 | -0.0935 | 0.0146 | 1.62E-10 | 1 |
| rs55649771 | 14 | 59669948 | 33 | 59449138 | 59891607 | A | G | 0.1342 | DAAM1 | intronic | -0.2017 | 0.0196 | 7.19E-25 | -0.1766 | 0.0408 | 1.51E-05 | -0.0971 | 0.0148 | 5.57E-11 | 1 |
| rs17255402 | 14 | 59672118 | 33 | 59449138 | 59891607 | A | G | 0.08648 | DAAM1 | intronic | -0.1731 | 0.023 | 5.84E-14 | -0.1782 | 0.0498 | 3.48E-04 | -0.0881 | 0.0170 | 2.14E-07 | 1 |
| rs77652087 | 14 | 59673348 | 33 | 59449138 | 59891607 | A | C | 0.08549 | DAAM1 | intronic | 0.1726 | 0.023 | 6.50E-14 | 0.1809 | 0.0497 | 2.77E-04 | 0.0882 | 0.0170 | 2.06E-07 | 1 |

| | | | | | | | | | | | Meta-GWAS | | | UKBB | | | ENIGMA* | | | |
|---|---|---|---|---|---|---|---|---|---|---|---|---|---|---|---|---|---|---|---|---|
| rsID | chr | pos (hg19) | Genomic Locus | start | end | A1 | A2 | MAF | Nearest Genes | Function | Beta | SE | P-value | Beta | SE | P-value | Beta | SE | P-value | Replicated** |
| rs79530766 | 14 | 59675486 | 33 | 59449138 | 59891607 | A | G | 0.08549 | DAAM1 | intronic | -0.1717 | 0.023 | 9.29E-14 | -0.1781 | 0.0498 | 3.52E-04 | -0.0883 | 0.0170 | 2.00E-07 | 1 |
| rs76957852 | 14 | 59675634 | 33 | 59449138 | 59891607 | T | C | 0.08549 | DAAM1 | intronic | 0.1712 | 0.023 | 1.04E-13 | 0.1780 | 0.0498 | 3.52E-04 | 0.0885 | 0.0170 | 1.88E-07 | 1 |
| rs76558755 | 14 | 59676851 | 33 | 59449138 | 59891607 | T | C | 0.08549 | DAAM1 | intronic | -0.1707 | 0.023 | 1.13E-13 | -0.1785 | 0.0497 | 3.32E-04 | -0.0884 | 0.0170 | 1.95E-07 | 1 |
| rs77216140 | 14 | 59678685 | 33 | 59449138 | 59891607 | T | G | 0.08648 | DAAM1 | intronic | 0.1704 | 0.023 | 1.17E-13 | 0.1784 | 0.0497 | 3.35E-04 | 0.0884 | 0.0170 | 1.90E-07 | 1 |
| rs79009632 | 14 | 59681380 | 33 | 59449138 | 59891607 | T | C | 0.08549 | DAAM1 | intronic | 0.1681 | 0.0229 | 1.92E-13 | 0.1811 | 0.0496 | 2.66E-04 | 0.0874 | 0.0170 | 2.53E-07 | 1 |
| rs75841771 | 14 | 59681450 | 33 | 59449138 | 59891607 | A | T | 0.08549 | DAAM1 | intronic | 0.1681 | 0.0229 | 1.92E-13 | 0.1811 | 0.0496 | 2.65E-04 | 0.0874 | 0.0170 | 2.52E-07 | 1 |
| rs78445564 | 14 | 59683086 | 33 | 59449138 | 59891607 | T | C | 0.04771 | DAAM1 | intronic | 0.2169 | 0.0336 | 1.13E-10 | 0.1581 | 0.0664 | 1.74E-02 | 0.0938 | 0.0264 | 3.92E-04 | 1 |
| rs17833798 | 14 | 59683581 | 33 | 59449138 | 59891607 | A | G | 0.08549 | DAAM1 | intronic | 0.1672 | 0.0229 | 2.61E-13 | 0.1783 | 0.0497 | 3.36E-04 | 0.0875 | 0.0170 | 2.45E-07 | 1 |
| rs112839082 | 14 | 59683921 | 33 | 59449138 | 59891607 | A | G | 0.08549 | DAAM1 | intronic | -0.1671 | 0.0229 | 2.72E-13 | -0.1783 | 0.0497 | 3.37E-04 | -0.0870 | 0.0169 | 2.87E-07 | 1 |
| rs111608226 | 14 | 59684637 | 33 | 59449138 | 59891607 | A | G | 0.08549 | DAAM1 | intronic | -0.1666 | 0.0229 | 3.04E-13 | -0.1783 | 0.0497 | 3.36E-04 | -0.0866 | 0.0169 | 3.18E-07 | 1 |
| rs74874233 | 14 | 59684859 | 33 | 59449138 | 59891607 | A | G | 0.08549 | DAAM1 | intronic | -0.1817 | 0.0241 | 4.45E-14 | -0.1824 | 0.0505 | 3.09E-04 | -0.0851 | 0.0182 | 2.74E-06 | 1 |
| rs77645702 | 14 | 59685139 | 33 | 59449138 | 59891607 | T | G | 0.08648 | DAAM1 | intronic | 0.1663 | 0.0228 | 3.19E-13 | 0.1811 | 0.0496 | 2.65E-04 | 0.0868 | 0.0169 | 2.97E-07 | 1 |
| rs77773396 | 14 | 59686936 | 33 | 59449138 | 59891607 | A | C | 0.08549 | DAAM1 | intronic | -0.1662 | 0.0228 | 3.21E-13 | -0.1785 | 0.0497 | 3.32E-04 | -0.0864 | 0.0169 | 3.29E-07 | 1 |
| rs721069 | 14 | 59687305 | 33 | 59449138 | 59891607 | A | G | 0.08648 | DAAM1 | intronic | 0.1655 | 0.0228 | 3.88E-13 | 0.1798 | 0.0496 | 2.92E-04 | 0.0868 | 0.0169 | 2.93E-07 | 1 |
| rs79059243 | 14 | 59688148 | 33 | 59449138 | 59891607 | A | G | 0.08549 | DAAM1 | intronic | 0.1654 | 0.0228 | 4.03E-13 | 0.1811 | 0.0496 | 2.65E-04 | 0.0865 | 0.0169 | 3.21E-07 | 1 |
| rs113971277 | 14 | 59689547 | 33 | 59449138 | 59891607 | T | C | 0.08648 | DAAM1 | intronic | 0.1653 | 0.0228 | 4.22E-13 | 0.1811 | 0.0496 | 2.64E-04 | 0.0863 | 0.0169 | 3.46E-07 | 1 |
| rs2099637 | 14 | 59689943 | 33 | 59449138 | 59891607 | A | C | 0.08549 | DAAM1 | intronic | -0.1654 | 0.0228 | 4.19E-13 | -0.1811 | 0.0496 | 2.66E-04 | -0.0863 | 0.0169 | 3.38E-07 | 1 |
| rs76786981 | 14 | 59692786 | 33 | 59449138 | 59891607 | A | T | 0.08549 | DAAM1 | intronic | -0.1649 | 0.0228 | 5.22E-13 | -0.1811 | 0.0496 | 2.65E-04 | -0.0856 | 0.0169 | 4.21E-07 | 1 |
| rs17255423 | 14 | 59694225 | 33 | 59449138 | 59891607 | C | G | 0.08648 | DAAM1 | intronic | -0.1639 | 0.0228 | 6.89E-13 | -0.1811 | 0.0496 | 2.65E-04 | -0.0855 | 0.0169 | 4.28E-07 | 1 |
| rs2757117 | 14 | 59694643 | 33 | 59449138 | 59891607 | A | G | 0.2376 | DAAM1 | intronic | -0.0961 | 0.0154 | 3.89E-10 | -0.1283 | 0.0324 | 7.64E-05 | -0.0428 | 0.0116 | 2.16E-04 | 1 |
| rs61986025 | 14 | 59700366 | 33 | 59449138 | 59891607 | T | C | 0.09344 | DAAM1 | intronic | -0.1464 | 0.022 | 2.80E-11 | -0.1676 | 0.0472 | 3.92E-04 | -0.0751 | 0.0164 | 4.42E-06 | 1 |
| rs17095915 | 14 | 59704841 | 33 | 59449138 | 59891607 | C | G | 0.09344 | DAAM1 | intronic | -0.1463 | 0.022 | 2.88E-11 | -0.1676 | 0.0472 | 3.90E-04 | -0.0747 | 0.0163 | 4.90E-06 | 1 |
| rs61986027 | 14 | 59705458 | 33 | 59449138 | 59891607 | A | G | 0.09344 | DAAM1 | intronic | 0.1465 | 0.022 | 2.58E-11 | 0.1702 | 0.0472 | 3.11E-04 | 0.0746 | 0.0163 | 4.97E-06 | 1 |
| rs17095918 | 14 | 59707006 | 33 | 59449138 | 59891607 | T | C | 0.09344 | DAAM1 | intronic | -0.1466 | 0.022 | 2.57E-11 | -0.1682 | 0.0472 | 3.72E-04 | -0.0751 | 0.0163 | 4.31E-06 | 1 |
| rs17095928 | 14 | 59708431 | 33 | 59449138 | 59891607 | C | G | 0.09443 | DAAM1 | intronic | -0.147 | 0.022 | 2.23E-11 | -0.1707 | 0.0472 | 2.98E-04 | -0.0747 | 0.0163 | 4.88E-06 | 1 |
| rs144053467 | 14 | 59709313 | 33 | 59449138 | 59891607 | A | G | 0.09443 | DAAM1 | intronic | 0.149 | 0.0221 | 1.55E-11 | 0.1751 | 0.0477 | 2.46E-04 | 0.0753 | 0.0164 | 4.24E-06 | 1 |
| rs147327353 | 14 | 59709316 | 33 | 59449138 | 59891607 | C | G | 0.09443 | DAAM1 | intronic | 0.1492 | 0.0221 | 1.45E-11 | 0.1751 | 0.0477 | 2.46E-04 | 0.0753 | 0.0164 | 4.21E-06 | 1 |
| rs61986028 | 14 | 59713871 | 33 | 59449138 | 59891607 | T | C | 0.08648 | DAAM1 | intronic | -0.1503 | 0.0226 | 2.98E-11 | -0.1700 | 0.0490 | 5.26E-04 | -0.0770 | 0.0168 | 4.62E-06 | 1 |
| rs17093312 | 14 | 59715923 | 33 | 59449138 | 59891607 | T | G | 0.08648 | DAAM1 | intronic | -0.1493 | 0.0226 | 4.35E-11 | -0.1715 | 0.0490 | 4.71E-04 | -0.0763 | 0.0168 | 5.81E-06 | 1 |
| rs17095939 | 14 | 59716240 | 33 | 59449138 | 59891607 | C | G | 0.08648 | DAAM1 | intronic | 0.1492 | 0.0226 | 4.44E-11 | 0.1716 | 0.0490 | 4.70E-04 | 0.0763 | 0.0168 | 5.78E-06 | 1 |
| rs17095949 | 14 | 59716875 | 33 | 59449138 | 59891607 | A | G | 0.08748 | DAAM1 | intronic | 0.1494 | 0.0226 | 4.24E-11 | 0.1716 | 0.0490 | 4.69E-04 | 0.0763 | 0.0168 | 5.76E-06 | 1 |
| rs17095951 | 14 | 59718628 | 33 | 59449138 | 59891607 | A | G | 0.08748 | DAAM1 | intronic | -0.1496 | 0.0226 | 3.87E-11 | -0.1719 | 0.0490 | 4.59E-04 | -0.0764 | 0.0168 | 5.56E-06 | 1 |
| rs17095960 | 14 | 59723745 | 33 | 59449138 | 59891607 | A | G | 0.08648 | DAAM1 | intronic | -0.1506 | 0.0226 | 2.83E-11 | -0.1736 | 0.0490 | 3.99E-04 | -0.0764 | 0.0168 | 5.46E-06 | 1 |
| rs61986029 | 14 | 59724880 | 33 | 59449138 | 59891607 | C | G | 0.08748 | DAAM1 | intronic | 0.151 | 0.0226 | 2.53E-11 | 0.1738 | 0.0490 | 3.92E-04 | 0.0760 | 0.0168 | 6.23E-06 | 1 |
| rs17095972 | 14 | 59727022 | 33 | 59449138 | 59891607 | A | G | 0.08648 | DAAM1 | intronic | 0.1497 | 0.0226 | 3.48E-11 | 0.1751 | 0.0489 | 3.50E-04 | 0.0764 | 0.0168 | 5.45E-06 | 1 |
| rs61986030 | 14 | 59728098 | 33 | 59449138 | 59891607 | T | C | 0.08648 | DAAM1 | intronic | -0.1507 | 0.0226 | 2.77E-11 | -0.1727 | 0.0490 | 4.32E-04 | -0.0763 | 0.0168 | 5.74E-06 | 1 |
| rs61986031 | 14 | 59728168 | 33 | 59449138 | 59891607 | T | C | 0.08648 | DAAM1 | intronic | -0.1506 | 0.0226 | 2.83E-11 | -0.1725 | 0.0490 | 4.37E-04 | -0.0762 | 0.0168 | 5.87E-06 | 1 |
| rs17095974 | 14 | 59728443 | 33 | 59449138 | 59891607 | T | C | 0.08648 | DAAM1 | intronic | -0.1508 | 0.0226 | 2.72E-11 | -0.1727 | 0.0490 | 4.31E-04 | -0.0762 | 0.0168 | 5.87E-06 | 1 |
| NA | 14 | 59731359 | 33 | 59449138 | 59891607 | T | C | 0.09046 | DAAM1 | intronic | 0.1568 | 0.0286 | 4.32E-08 | 0.1847 | 0.0476 | 1.07E-04 | 0.0784 | 0.0165 | 1.92E-06 | 1 |
| rs17833876 | 14 | 59734171 | 33 | 59449138 | 59891607 | T | C | 0.09245 | DAAM1 | intronic | 0.1497 | 0.022 | 9.84E-12 | 0.1729 | 0.0477 | 2.90E-04 | 0.0786 | 0.0164 | 1.72E-06 | 1 |
| rs17255500 | 14 | 59735696 | 33 | 59449138 | 59891607 | T | C | 0.09245 | DAAM1 | intronic | 0.1493 | 0.022 | 1.12E-11 | 0.1728 | 0.0477 | 2.91E-04 | 0.0787 | 0.0164 | 1.70E-06 | 1 |
| rs17255507 | 14 | 59736657 | 33 | 59449138 | 59891607 | A | G | 0.09245 | DAAM1 | intronic | 0.1492 | 0.022 | 1.15E-11 | 0.1728 | 0.0477 | 2.91E-04 | 0.0787 | 0.0164 | 1.67E-06 | 1 |
| rs17255514 | 14 | 59736709 | 33 | 59449138 | 59891607 | A | T | 0.09245 | DAAM1 | intronic | 0.1492 | 0.022 | 1.13E-11 | 0.1730 | 0.0477 | 2.88E-04 | 0.0787 | 0.0164 | 1.66E-06 | 1 |
| rs61986048 | 14 | 59737176 | 33 | 59449138 | 59891607 | A | G | 0.09245 | DAAM1 | intronic | -0.1492 | 0.022 | 1.16E-11 | -0.1730 | 0.0477 | 2.88E-04 | -0.0787 | 0.0164 | 1.66E-06 | 1 |
| rs17255520 | 14 | 59737475 | 33 | 59449138 | 59891607 | A | C | 0.09245 | DAAM1 | intronic | -0.1494 | 0.022 | 1.10E-11 | -0.1728 | 0.0477 | 2.93E-04 | -0.0790 | 0.0164 | 1.54E-06 | 1 |
| rs17095989 | 14 | 59737859 | 33 | 59449138 | 59891607 | A | G | 0.09245 | DAAM1 | intronic | -0.148 | 0.022 | 1.60E-11 | -0.1678 | 0.0475 | 4.19E-04 | -0.0788 | 0.0164 | 1.60E-06 | 1 |
| rs17093314 | 14 | 59738186 | 33 | 59449138 | 59891607 | A | G | 0.09245 | DAAM1 | intronic | -0.1493 | 0.0219 | 1.01E-11 | -0.1680 | 0.0475 | 4.15E-04 | -0.0793 | 0.0164 | 1.38E-06 | 1 |
| rs17095990 | 14 | 59738654 | 33 | 59449138 | 59891607 | A | G | 0.09145 | DAAM1 | intronic | -0.1482 | 0.022 | 1.55E-11 | -0.1690 | 0.0476 | 3.85E-04 | -0.0785 | 0.0164 | 1.81E-06 | 1 |
| rs61986049 | 14 | 59739068 | 33 | 59449138 | 59891607 | A | G | 0.09245 | DAAM1 | intronic | -0.1493 | 0.022 | 1.09E-11 | -0.1729 | 0.0477 | 2.90E-04 | -0.0786 | 0.0164 | 1.73E-06 | 1 |
| rs17833888 | 14 | 59739970 | 33 | 59449138 | 59891607 | A | G | 0.09245 | DAAM1 | intronic | 0.1493 | 0.022 | 1.14E-11 | 0.1729 | 0.0477 | 2.89E-04 | 0.0785 | 0.0164 | 1.79E-06 | 1 |
| rs79869391 | 14 | 59740164 | 33 | 59449138 | 59891607 | A | G | 0.09245 | DAAM1 | intronic | 0.1492 | 0.022 | 1.15E-11 | 0.1729 | 0.0477 | 2.89E-04 | 0.0785 | 0.0164 | 1.79E-06 | 1 |
| rs17255527 | 14 | 59741542 | 33 | 59449138 | 59891607 | A | T | 0.09245 | DAAM1 | intronic | 0.1511 | 0.022 | 6.76E-12 | 0.1749 | 0.0479 | 2.60E-04 | 0.0774 | 0.0164 | 2.50E-06 | 1 |
| rs111977890 | 14 | 59744282 | 33 | 59449138 | 59891607 | A | G | 0.09245 | DAAM1 | intronic | -0.1505 | 0.022 | 8.81E-12 | -0.1742 | 0.0479 | 2.76E-04 | -0.0776 | 0.0164 | 2.41E-06 | 1 |
| rs61986052 | 14 | 59745427 | 33 | 59449138 | 59891607 | C | G | 0.09145 | DAAM1 | intronic | -0.1509 | 0.022 | 7.51E-12 | -0.1748 | 0.0479 | 2.62E-04 | -0.0780 | 0.0164 | 2.08E-06 | 1 |
| rs17255534 | 14 | 59745532 | 33 | 59449138 | 59891607 | T | C | 0.09245 | DAAM1 | intronic | -0.1509 | 0.022 | 7.52E-12 | -0.1748 | 0.0479 | 2.62E-04 | -0.0776 | 0.0164 | 2.40E-06 | 1 |
| rs2033795 | 14 | 59748893 | 33 | 59449138 | 59891607 | A | G | 0.09245 | DAAM1 | intronic | -0.1506 | 0.022 | 8.31E-12 | -0.1748 | 0.0479 | 2.61E-04 | -0.0774 | 0.0165 | 2.53E-06 | 1 |
| rs4296184 | 14 | 59749007 | 33 | 59449138 | 59891607 | T | G | 0.09145 | DAAM1 | intronic | -0.1505 | 0.022 | 8.49E-12 | -0.1764 | 0.0478 | 2.27E-04 | -0.0775 | 0.0165 | 2.50E-06 | 1 |
| rs61986054 | 14 | 59749187 | 33 | 59449138 | 59891607 | A | T | 0.09245 | DAAM1 | intronic | -0.1505 | 0.022 | 8.58E-12 | -0.1764 | 0.0478 | 2.27E-04 | -0.0773 | 0.0165 | 2.63E-06 | 1 |
| rs61986055 | 14 | 59749226 | 33 | 59449138 | 59891607 | T | C | 0.09245 | DAAM1 | intronic | 0.1506 | 0.022 | 8.20E-12 | 0.1759 | 0.0478 | 2.35E-04 | 0.0772 | 0.0165 | 2.67E-06 | 1 |
| rs61986056 | 14 | 59750695 | 33 | 59449138 | 59891607 | A | G | 0.09245 | DAAM1 | intronic | -0.1504 | 0.022 | 8.89E-12 | -0.1748 | 0.0479 | 2.61E-04 | -0.0774 | 0.0165 | 2.57E-06 | 1 |
| rs78270394 | 14 | 59752088 | 33 | 59449138 | 59891607 | A | G | 0.09245 | DAAM1 | intronic | 0.1507 | 0.022 | 7.98E-12 | 0.1748 | 0.0478 | 2.57E-04 | 0.0770 | 0.0165 | 2.89E-06 | 1 |
| rs75555742 | 14 | 59752121 | 33 | 59449138 | 59891607 | A | G | 0.09245 | DAAM1 | intronic | -0.1495 | 0.022 | 1.03E-11 | -0.1749 | 0.0479 | 2.59E-04 | -0.0739 | 0.0164 | 6.90E-06 | 1 |
| rs61986057 | 14 | 59752957 | 33 | 59449138 | 59891607 | T | C | 0.1133 | DAAM1 | intronic | -0.1203 | 0.02 | 1.90E-09 | -0.1286 | 0.0435 | 3.14E-03 | -0.0515 | 0.0151 | 6.34E-04 | 1 |
| rs61986058 | 14 | 59758165 | 33 | 59449138 | 59891607 | A | C | 0.09245 | DAAM1 | UTR3 | 0.1511 | 0.022 | 6.90E-12 | 0.1755 | 0.0478 | 2.42E-04 | 0.0766 | 0.0165 | 3.23E-06 | 1 |
| rs61986059 | 14 | 59758310 | 33 | 59449138 | 59891607 | T | G | 0.09245 | DAAM1 | intronic | -0.1454 | 0.0258 | 1.82E-08 | -0.1769 | 0.0478 | 2.14E-04 | -0.0767 | 0.0165 | 3.20E-06 | 1 |
| rs74748336 | 14 | 59758919 | 33 | 59449138 | 59891607 | A | G | 0.09245 | DAAM1 | intronic | -0.1503 | 0.022 | 8.90E-12 | -0.1753 | 0.0478 | 2.48E-04 | -0.0766 | 0.0165 | 3.22E-06 | 1 |
| rs78271461 | 14 | 59759114 | 33 | 59449138 | 59891607 | A | C | 0.09245 | DAAM1 | intronic | -0.1509 | 0.022 | 7.43E-12 | -0.1737 | 0.0478 | 2.80E-04 | -0.0766 | 0.0165 | 3.22E-06 | 1 |
| rs17255548 | 14 | 59759818 | 33 | 59449138 | 59891607 | A | G | 0.09245 | DAAM1 | intronic | 0.1503 | 0.022 | 8.91E-12 | 0.1753 | 0.0478 | 2.47E-04 | 0.0767 | 0.0165 | 3.20E-06 | 1 |
| rs10483711 | 14 | 59761126 | 33 | 59449138 | 59891607 | T | C | 0.08748 | DAAM1 | intronic | -0.1523 | 0.0227 | 1.81E-11 | -0.1836 | 0.0491 | 1.87E-04 | -0.0754 | 0.0168 | 7.48E-06 | 1 |
| rs10483713 | 14 | 59762998 | 33 | 59449138 | 59891607 | T | C | 0.09145 | DAAM1 | intronic | 0.1501 | 0.022 | 9.65E-12 | 0.1765 | 0.0477 | 2.20E-04 | 0.0765 | 0.0165 | 3.33E-06 | 1 |
| rs17255583 | 14 | 59763748 | 33 | 59449138 | 59891607 | T | C | 0.09145 | DAAM1 | intronic | -0.1504 | 0.022 | 8.95E-12 | -0.1753 | 0.0478 | 2.46E-04 | -0.0766 | 0.0165 | 3.30E-06 | 1 |
| rs61984460 | 14 | 59764524 | 33 | 59449138 | 59891607 | T | C | 0.09245 | DAAM1 | intronic | 0.1503 | 0.022 | 9.17E-12 | 0.1768 | 0.0478 | 2.15E-04 | 0.0766 | 0.0165 | 3.23E-06 | 1 |
| rs61984461 | 14 | 59764529 | 33 | 59449138 | 59891607 | A | G | 0.09245 | DAAM1 | intronic | 0.1501 | 0.022 | 9.47E-12 | 0.1769 | 0.0478 | 2.14E-04 | 0.0767 | 0.0165 | 3.23E-06 | 1 |
| rs17833930 | 14 | 59764790 | 33 | 59449138 | 59891607 | A | G | 0.09245 | DAAM1 | intronic | -0.1503 | 0.022 | 9.20E-12 | -0.1769 | 0.0478 | 2.13E-04 | -0.0767 | 0.0165 | 3.23E-06 | 1 |
| rs17833942 | 14 | 59765908 | 33 | 59449138 | 59891607 | A | G | 0.09245 | DAAM1 | intronic | 0.1503 | 0.022 | 9.18E-12 | 0.1753 | 0.0478 | 2.46E-04 | 0.0767 | 0.0165 | 3.20E-06 | 1 |
| rs76990548 | 14 | 59768101 | 33 | 59449138 | 59891607 | A | G | 0.08648 | DAAM1 | intronic | -0.1528 | 0.0227 | 1.61E-11 | -0.1836 | 0.0491 | 1.88E-04 | -0.0756 | 0.0168 | 7.19E-06 | 1 |
| rs17096046 | 14 | 59768135 | 33 | 59449138 | 59891607 | T | C | 0.08648 | DAAM1 | intronic | 0.1525 | 0.0227 | 1.72E-11 | 0.1824 | 0.0491 | 2.02E-04 | 0.0756 | 0.0168 | 7.07E-06 | 1 |
| rs17096048 | 14 | 59768381 | 33 | 59449138 | 59891607 | A | G | 0.08648 | DAAM1 | intronic | -0.1527 | 0.0227 | 1.65E-11 | -0.1829 | 0.0490 | 1.93E-04 | -0.0756 | 0.0168 | 7.18E-06 | 1 |
| rs61984462 | 14 | 59769760 | 33 | 59449138 | 59891607 | T | C | 0.08648 | DAAM1 | intronic | -0.1528 | 0.0227 | 1.57E-11 | -0.1843 | 0.0491 | 1.73E-04 | -0.0757 | 0.0168 | 6.94E-06 | 1 |
| rs78212084 | 14 | 59770439 | 33 | 59449138 | 59891607 | A | G | 0.08648 | DAAM1 | intronic | -0.1529 | 0.0227 | 1.57E-11 | -0.1826 | 0.0491 | 2.03E-04 | -0.0758 | 0.0168 | 6.81E-06 | 1 |
| rs61984463 | 14 | 59771730 | 33 | 59449138 | 59891607 | A | G | 0.09145 | DAAM1 | intronic | 0.1482 | 0.022 | 1.67E-11 | 0.1730 | 0.0475 | 2.72E-04 | 0.0772 | 0.0165 | 2.77E-06 | 1 |
| rs75663630 | 14 | 59772068 | 33 | 59449138 | 59891607 | A | C | 0.08648 | DAAM1 | intronic | -0.1525 | 0.0227 | 1.72E-11 | -0.1841 | 0.0490 | 1.75E-04 | -0.0758 | 0.0168 | 6.65E-06 | 1 |
| rs17096053 | 14 | 59772866 | 33 | 59449138 | 59891607 | T | C | 0.08648 | DAAM1 | intronic | 0.1506 | 0.0226 | 2.81E-11 | 0.1814 | 0.0488 | 2.03E-04 | 0.0763 | 0.0168 | 5.91E-06 | 1 |
| rs17096056 | 14 | 59772986 | 33 | 59449138 | 59891607 | T | C | 0.08648 | DAAM1 | intronic | -0.1508 | 0.0226 | 2.66E-11 | -0.1818 | 0.0488 | 1.96E-04 | -0.0763 | 0.0168 | 5.85E-06 | 1 |
| rs61984464 | 14 | 59773180 | 33 | 59449138 | 59891607 | T | C | 0.08648 | DAAM1 | intronic | 0.1511 | 0.0226 | 2.52E-11 | 0.1797 | 0.0488 | 2.35E-04 | 0.0761 | 0.0168 | 6.17E-06 | 1 |
| rs17255610 | 14 | 59775562 | 33 | 59449138 | 59891607 | A | G | 0.09245 | DAAM1 | intronic | -0.1504 | 0.022 | 8.98E-12 | -0.1738 | 0.0478 | 2.78E-04 | -0.0770 | 0.0165 | 2.94E-06 | 1 |
| rs61984489 | 14 | 59781468 | 33 | 59449138 | 59891607 | A | G | 0.09344 | DAAM1 | intronic | 0.1491 | 0.022 | 1.24E-11 | 0.1717 | 0.0475 | 3.04E-04 | 0.0800 | 0.0164 | 1.15E-06 | 1 |
| rs17096066 | 14 | 59782604 | 33 | 59449138 | 59891607 | T | C | 0.08847 | DAAM1 | intronic | -0.1515 | 0.0226 | 2.15E-11 | -0.1789 | 0.0489 | 2.52E-04 | -0.0788 | 0.0168 | 2.75E-06 | 1 |
| rs10483714 | 14 | 59785346 | 33 | 59449138 | 59891607 | A | G | 0.09443 | DAAM1 | intronic | 0.1487 | 0.022 | 1.40E-11 | 0.1703 | 0.0476 | 3.44E-04 | 0.0791 | 0.0164 | 1.49E-06 | 1 |
| rs61984493 | 14 | 59785916 | 33 | 59449138 | 59891607 | C | G | 0.09344 | DAAM1 | intronic | -0.149 | 0.022 | 1.29E-11 | -0.1705 | 0.0476 | 3.41E-04 | -0.0795 | 0.0164 | 1.32E-06 | 1 |
| rs78626699 | 14 | 59786434 | 33 | 59449138 | 59891607 | A | T | 0.09443 | DAAM1 | intronic | -0.1489 | 0.022 | 1.29E-11 | -0.1698 | 0.0475 | 3.55E-04 | -0.0793 | 0.0164 | 1.38E-06 | 1 |

| | | | | | | | | | | | Meta-GWAS | | | UKBB | | | ENIGMA* | | | |
|---|---|---|---|---|---|---|---|---|---|---|---|---|---|---|---|---|---|---|---|---|
| rsID | chr | pos (hg19) | Genomic Locus | start | end | A1 | A2 | MAF | Nearest Genes | Function | Beta | SE | P-value | Beta | SE | P-value | Beta | SE | P-value | Replicated** |
| rs45610932 | 14 | 59787325 | 33 | 59449138 | 59891607 | C | G | 0.09145 | DAAM1 | intronic | -0.1487 | 0.022 | 1.43E-11 | -0.1718 | 0.0476 | 3.10E-04 | -0.0781 | 0.0165 | 2.06E-06 | 1 |
| rs17096074 | 14 | 59789892 | 33 | 59449138 | 59891607 | T | G | 0.08648 | DAAM1 | exonic | -0.1505 | 0.0226 | 2.82E-11 | -0.1807 | 0.0489 | 2.22E-04 | -0.0768 | 0.0168 | 4.91E-06 | 1 |
| rs17096077 | 14 | 59790240 | 33 | 59449138 | 59891607 | A | T | 0.08648 | DAAM1 | intronic | -0.1504 | 0.0226 | 2.92E-11 | -0.1807 | 0.0489 | 2.23E-04 | -0.0767 | 0.0168 | 5.03E-06 | 1 |
| rs61984494 | 14 | 59790624 | 33 | 59449138 | 59891607 | T | C | 0.08648 | DAAM1 | intronic | -0.1502 | 0.0226 | 3.08E-11 | -0.1808 | 0.0489 | 2.22E-04 | -0.0766 | 0.0168 | 5.18E-06 | 1 |
| rs17096084 | 14 | 59791647 | 33 | 59449138 | 59891607 | A | G | 0.08847 | DAAM1 | intronic | 0.1502 | 0.0226 | 3.00E-11 | 0.1788 | 0.0489 | 2.59E-04 | 0.0779 | 0.0168 | 3.48E-06 | 1 |
| rs41285510 | 14 | 59797213 | 33 | 59449138 | 59891607 | A | G | 0.09245 | DAAM1 | intronic | -0.1476 | 0.022 | 1.99E-11 | -0.1700 | 0.0477 | 3.69E-04 | -0.0774 | 0.0165 | 2.57E-06 | 1 |
| rs28927674 | 14 | 59797373 | 33 | 59449138 | 59891607 | A | G | 0.09245 | DAAM1 | exonic | -0.1473 | 0.022 | 2.14E-11 | -0.1700 | 0.0477 | 3.68E-04 | -0.0774 | 0.0165 | 2.55E-06 | 1 |
| rs17096107 | 14 | 59800036 | 33 | 59449138 | 59891607 | C | G | 0.08946 | DAAM1 | intronic | -0.1491 | 0.0226 | 4.14E-11 | -0.1821 | 0.0489 | 1.98E-04 | -0.0769 | 0.0168 | 4.84E-06 | 1 |
| rs61984497 | 14 | 59801714 | 33 | 59449138 | 59891607 | A | G | 0.1203 | DAAM1 | intronic | 0.1215 | 0.0205 | 3.18E-09 | 0.1437 | 0.0438 | 1.04E-03 | 0.0530 | 0.0152 | 4.97E-04 | 1 |
| rs17096117 | 14 | 59805355 | 33 | 59449138 | 59891607 | C | G | 0.08847 | DAAM1 | intronic | 0.1505 | 0.0227 | 3.08E-11 | 0.1775 | 0.0490 | 2.93E-04 | 0.0793 | 0.0169 | 2.61E-06 | 1 |
| rs75341124 | 14 | 59805480 | 33 | 59449138 | 59891607 | T | C | 0.04771 | DAAM1 | intronic | -0.2244 | 0.0363 | 6.44E-10 | -0.1890 | 0.0695 | 6.53E-03 | -0.1143 | 0.0291 | 8.51E-05 | 1 |
| rs17096126 | 14 | 59806629 | 33 | 59449138 | 59891607 | T | G | 0.08847 | DAAM1 | intronic | 0.1512 | 0.0227 | 2.50E-11 | 0.1782 | 0.0490 | 2.74E-04 | 0.0782 | 0.0170 | 4.19E-06 | 1 |
| rs61984499 | 14 | 59815097 | 33 | 59449138 | 59891607 | A | G | 0.08648 | DAAM1 | intronic | 0.1418 | 0.0231 | 8.89E-10 | 0.2174 | 0.0502 | 1.50E-05 | 0.0760 | 0.0171 | 8.45E-06 | 1 |
| rs17834014 | 14 | 59816026 | 33 | 59449138 | 59891607 | A | G | 0.08648 | DAAM1 | intronic | 0.14 | 0.0232 | 1.53E-09 | 0.2044 | 0.0504 | 5.09E-05 | 0.0765 | 0.0171 | 7.47E-06 | 1 |
| rs61984520 | 14 | 59816182 | 33 | 59449138 | 59891607 | C | G | 0.08648 | DAAM1 | intronic | 0.1408 | 0.0232 | 1.28E-09 | 0.1976 | 0.0504 | 9.01E-05 | 0.0765 | 0.0171 | 7.63E-06 | 1 |
| rs61984521 | 14 | 59820189 | 33 | 59449138 | 59891607 | T | C | 0.08549 | DAAM1 | intronic | -0.1395 | 0.0232 | 1.84E-09 | -0.1992 | 0.0505 | 7.97E-05 | -0.0739 | 0.0171 | 1.58E-05 | 1 |
| rs61984524 | 14 | 59824249 | 33 | 59449138 | 59891607 | C | G | 0.08648 | DAAM1 | intronic | 0.1396 | 0.0232 | 1.82E-09 | 0.1975 | 0.0505 | 9.24E-05 | 0.0757 | 0.0171 | 9.55E-06 | 1 |
| rs61984526 | 14 | 59831428 | 33 | 59449138 | 59891607 | T | C | 0.08549 | DAAM1 | intronic | 0.1414 | 0.0233 | 1.27E-09 | 0.1994 | 0.0504 | 7.64E-05 | 0.0735 | 0.0173 | 2.07E-05 | 1 |
| rs45476291 | 14 | 59836655 | 33 | 59449138 | 59891607 | A | G | 0.08648 | DAAM1 | UTR3 | 0.1393 | 0.0233 | 2.10E-09 | 0.1997 | 0.0505 | 7.83E-05 | 0.0739 | 0.0171 | 1.54E-05 | 1 |
| rs61984529 | 14 | 59841236 | 33 | 59449138 | 59891607 | T | G | 0.08549 | DAAM1 | intergenic | -0.138 | 0.0233 | 2.94E-09 | -0.2051 | 0.0506 | 5.09E-05 | -0.0734 | 0.0172 | 1.89E-05 | 1 |
| rs61984530 | 14 | 59844320 | 33 | 59449138 | 59891607 | A | G | 0.08648 | DAAM1 | intergenic | 0.1374 | 0.0233 | 3.54E-09 | 0.1973 | 0.0505 | 9.37E-05 | 0.0739 | 0.0172 | 1.65E-05 | 1 |
| rs75061529 | 14 | 59846735 | 33 | 59449138 | 59891607 | A | C | 0.08549 | DAAM1 | intergenic | -0.1371 | 0.0233 | 4.09E-09 | -0.1971 | 0.0505 | 9.53E-05 | -0.0740 | 0.0172 | 1.66E-05 | 1 |
| rs61984532 | 14 | 59846757 | 33 | 59449138 | 59891607 | A | G | 0.08648 | DAAM1 | intergenic | -0.1369 | 0.0233 | 4.32E-09 | -0.1971 | 0.0505 | 9.53E-05 | -0.0740 | 0.0172 | 1.68E-05 | 1 |
| rs61984533 | 14 | 59847613 | 33 | 59449138 | 59891607 | T | C | 0.08648 | DAAM1 | intergenic | 0.1372 | 0.0233 | 4.09E-09 | 0.1957 | 0.0505 | 1.09E-04 | 0.0742 | 0.0172 | 1.57E-05 | 1 |
| rs61984534 | 14 | 59848499 | 33 | 59449138 | 59891607 | T | C | 0.08648 | DAAM1 | intergenic | -0.1363 | 0.0233 | 5.16E-09 | -0.1988 | 0.0505 | 8.43E-05 | -0.0732 | 0.0172 | 2.04E-05 | 1 |
| rs61984535 | 14 | 59852007 | 33 | 59449138 | 59891607 | T | C | 0.08648 | DAAM1 | intergenic | -0.1408 | 0.0234 | 1.66E-09 | -0.2021 | 0.0506 | 6.51E-05 | -0.0734 | 0.0172 | 1.97E-05 | 1 |
| rs61984536 | 14 | 59852262 | 33 | 59449138 | 59891607 | T | C | 0.08748 | DAAM1 | intergenic | -0.1376 | 0.0234 | 4.01E-09 | -0.2025 | 0.0507 | 6.54E-05 | -0.0738 | 0.0172 | 1.79E-05 | 1 |
| rs61984537 | 14 | 59852500 | 33 | 59449138 | 59891607 | T | C | 0.08648 | AL159140.1 | intergenic | 0.1382 | 0.0233 | 3.21E-09 | 0.2156 | 0.0504 | 1.94E-05 | 0.0747 | 0.0172 | 1.41E-05 | 1 |
| rs78084529 | 14 | 59852629 | 33 | 59449138 | 59891607 | C | G | 0.08748 | AL159140.1 | intergenic | -0.1412 | 0.0237 | 2.56E-09 | -0.2161 | 0.0503 | 1.79E-05 | -0.0765 | 0.0178 | 1.67E-05 | 1 |
| rs79863915 | 14 | 59852630 | 33 | 59449138 | 59891607 | T | G | 0.08748 | AL159140.1 | intergenic | -0.1412 | 0.0237 | 2.56E-09 | -0.2161 | 0.0503 | 1.79E-05 | -0.0765 | 0.0178 | 1.67E-05 | 1 |
| rs61984538 | 14 | 59852686 | 33 | 59449138 | 59891607 | T | C | 0.08847 | AL159140.1 | intergenic | 0.1398 | 0.0233 | 1.93E-09 | 0.2161 | 0.0503 | 1.79E-05 | 0.0746 | 0.0172 | 1.40E-05 | 1 |
| rs61984539 | 14 | 59852891 | 33 | 59449138 | 59891607 | T | C | 0.08748 | AL159140.1 | intergenic | -0.1391 | 0.0233 | 2.36E-09 | -0.2160 | 0.0503 | 1.80E-05 | -0.0737 | 0.0172 | 1.78E-05 | 1 |
| rs7151357 | 14 | 59853140 | 33 | 59449138 | 59891607 | A | C | 0.08748 | AL159140.1 | intergenic | -0.1415 | 0.0235 | 1.72E-09 | -0.2161 | 0.0503 | 1.80E-05 | -0.0747 | 0.0175 | 1.93E-05 | 1 |
| rs73299984 | 14 | 59854408 | 33 | 59449138 | 59891607 | T | C | 0.08748 | AL159140.1 | intergenic | 0.1393 | 0.0233 | 2.27E-09 | 0.2158 | 0.0503 | 1.84E-05 | 0.0739 | 0.0172 | 1.75E-05 | 1 |
| rs61984545 | 14 | 59856978 | 33 | 59449138 | 59891607 | A | G | 0.08648 | AL159140.1 | intergenic | -0.1376 | 0.0234 | 4.35E-09 | -0.1979 | 0.0506 | 9.24E-05 | -0.0736 | 0.0173 | 1.98E-05 | 1 |
| rs80264941 | 14 | 59857852 | 33 | 59449138 | 59891607 | T | C | 0.08648 | AL159140.1 | intergenic | 0.1375 | 0.0234 | 4.19E-09 | 0.2002 | 0.0505 | 7.44E-05 | 0.0754 | 0.0172 | 1.20E-05 | 1 |
| rs112171060 | 14 | 59858269 | 33 | 59449138 | 59891607 | A | G | 0.08648 | AL159140.1 | intergenic | 0.1376 | 0.0234 | 4.13E-09 | 0.1992 | 0.0505 | 8.01E-05 | 0.0758 | 0.0172 | 1.11E-05 | 1 |
| rs61985462 | 14 | 59858837 | 33 | 59449138 | 59891607 | T | C | 0.08648 | AL159140.1 | intergenic | 0.1377 | 0.0234 | 4.06E-09 | 0.1988 | 0.0505 | 8.30E-05 | 0.0755 | 0.0173 | 1.23E-05 | 1 |
| rs113855394 | 14 | 59859472 | 33 | 59449138 | 59891607 | T | C | 0.08648 | AL159140.1 | intergenic | 0.138 | 0.0234 | 3.84E-09 | 0.1971 | 0.0505 | 9.65E-05 | 0.0761 | 0.0173 | 1.05E-05 | 1 |
| rs61985463 | 14 | 59860572 | 33 | 59449138 | 59891607 | T | C | 0.08648 | AL159140.1 | intergenic | -0.1378 | 0.0234 | 4.00E-09 | -0.1990 | 0.0505 | 8.12E-05 | -0.0764 | 0.0173 | 1.00E-05 | 1 |
| rs61985464 | 14 | 59860716 | 33 | 59449138 | 59891607 | T | C | 0.08648 | AL159140.1 | intergenic | 0.1368 | 0.0234 | 5.36E-09 | 0.1989 | 0.0505 | 8.20E-05 | 0.0764 | 0.0173 | 9.81E-06 | 1 |
| rs7500427 | 16 | 52545277 | 34 | 52538040 | 52635000 | A | G | 0.2793 | TOX3 | intronic | -0.0844 | 0.015 | 1.89E-08 | -0.0705 | 0.0314 | 2.49E-02 | -0.0234 | 0.0116 | 4.30E-02 | 1 |
| rs9936081 | 16 | 52549646 | 34 | 52538040 | 52635000 | A | G | 0.2793 | TOX3 | intronic | -0.0854 | 0.015 | 1.20E-08 | -0.0707 | 0.0314 | 2.43E-02 | -0.0244 | 0.0116 | 3.54E-02 | 1 |
| rs1345388 | 16 | 52556293 | 34 | 52538040 | 52635000 | T | C | 0.2793 | TOX3 | intronic | 0.0863 | 0.015 | 8.14E-09 | 0.0719 | 0.0314 | 2.19E-02 | 0.0253 | 0.0116 | 2.84E-02 | 1 |
| rs12918816 | 16 | 52560213 | 34 | 52538040 | 52635000 | A | G | 0.2803 | TOX3 | intronic | -0.0834 | 0.0149 | 2.28E-08 | -0.0730 | 0.0314 | 2.01E-02 | -0.0230 | 0.0115 | 4.60E-02 | 1 |
| rs1362548 | 16 | 52563951 | 34 | 52538040 | 52635000 | C | G | 0.2819 | TOX3 | intronic | -0.0837 | 0.015 | 2.52E-08 | -0.0769 | 0.0313 | 1.39E-02 | -0.0210 | 0.0115 | 6.79E-02 | 0 |
| rs9921569 | 16 | 52572029 | 34 | 52538040 | 52635000 | T | C | 0.2823 | TOX3 | intronic | 0.0837 | 0.0148 | 1.66E-08 | 0.0792 | 0.0313 | 1.14E-02 | 0.0216 | 0.0114 | 5.88E-02 | 0 |
| rs35850695 | 16 | 52574343 | 34 | 52538040 | 52635000 | A | G | 0.2763 | TOX3 | intronic | -0.0824 | 0.015 | 4.24E-08 | -0.0793 | 0.0314 | 1.17E-02 | -0.0212 | 0.0116 | 6.71E-02 | 0 |
| rs4784223 | 16 | 52575907 | 34 | 52538040 | 52635000 | A | G | 0.2823 | TOX3 | intronic | 0.0845 | 0.0148 | 1.23E-08 | 0.0796 | 0.0313 | 1.10E-02 | 0.0215 | 0.0114 | 5.96E-02 | 0 |
| rs3095602 | 16 | 52580500 | 34 | 52538040 | 52635000 | A | G | 0.2903 | TOX3 | intronic | -0.0849 | 0.0147 | 8.14E-09 | -0.0734 | 0.0313 | 1.90E-02 | -0.0213 | 0.0113 | 5.91E-02 | 0 |
| rs12930156 | 16 | 52581424 | 34 | 52538040 | 52635000 | T | C | 0.2913 | TOX3 | intronic | -0.0852 | 0.0147 | 7.28E-09 | -0.0733 | 0.0312 | 1.90E-02 | -0.0208 | 0.0113 | 6.50E-02 | 0 |
| rs3095604 | 16 | 52581979 | 34 | 52538040 | 52635000 | C | G | 0.2903 | TOX3 | upstream | -0.085 | 0.0147 | 7.80E-09 | -0.0736 | 0.0312 | 1.85E-02 | -0.0210 | 0.0113 | 6.22E-02 | 0 |
| rs28463809 | 16 | 52583054 | 34 | 52538040 | 52635000 | T | G | 0.2903 | TOX3 | intergenic | -0.0854 | 0.0147 | 6.52E-09 | -0.0738 | 0.0312 | 1.81E-02 | -0.0212 | 0.0113 | 5.94E-02 | 0 |
| rs4784226 | 16 | 52583143 | 34 | 52538040 | 52635000 | T | C | 0.2495 | TOX3 | intergenic | -0.0878 | 0.0153 | 9.17E-09 | -0.0691 | 0.0318 | 2.99E-02 | -0.0249 | 0.0117 | 3.43E-02 | 1 |
| rs3095606 | 16 | 52584173 | 34 | 52538040 | 52635000 | A | G | 0.2903 | CASC16 | intergenic | 0.0852 | 0.0147 | 7.11E-09 | 0.0740 | 0.0312 | 1.78E-02 | 0.0211 | 0.0113 | 6.03E-02 | 0 |
| rs3095607 | 16 | 52584295 | 34 | 52538040 | 52635000 | T | G | 0.2903 | CASC16 | intergenic | 0.0852 | 0.0147 | 7.02E-09 | 0.0740 | 0.0312 | 1.78E-02 | 0.0213 | 0.0113 | 5.82E-02 | 0 |
| rs3112578 | 16 | 52585440 | 34 | 52538040 | 52635000 | T | C | 0.2555 | CASC16 | downstream | 0.0886 | 0.015 | 3.84E-09 | 0.0713 | 0.0316 | 2.43E-02 | 0.0247 | 0.0115 | 3.22E-02 | 1 |
| rs3803662 | 16 | 52586341 | 34 | 52538040 | 52635000 | A | G | 0.2913 | CASC16 | ncRNA_exonic | -0.0853 | 0.0147 | 6.07E-09 | -0.0741 | 0.0312 | 1.74E-02 | -0.0207 | 0.0112 | 6.52E-02 | 0 |
| rs3803661 | 16 | 52586477 | 34 | 52538040 | 52635000 | A | G | 0.2913 | CASC16 | ncRNA_intronic | -0.0854 | 0.0147 | 5.99E-09 | -0.0741 | 0.0312 | 1.74E-02 | -0.0208 | 0.0112 | 6.36E-02 | 0 |
| rs4784227 | 16 | 52599188 | 34 | 52538040 | 52635000 | T | C | 0.2545 | CASC16 | ncRNA_intronic | -0.0867 | 0.0153 | 1.41E-08 | -0.0636 | 0.0318 | 4.51E-02 | -0.0249 | 0.0118 | 3.49E-02 | 1 |
| rs6097618 | 20 | 52448936 | 35 | 52448936 | 52471667 | C | G | 0.4433 | AC006076.1 | intergenic | 0.0749 | 0.0136 | 3.85E-08 | 0.0529 | 0.0276 | 5.54E-02 | 0.0202 | 0.0104 | 5.28E-02 | 0 |
| rs6126986 | 20 | 52464719 | 35 | 52448936 | 52471667 | T | C | 0.493 | SUMO1P1 | intergenic | -0.0747 | 0.0136 | 4.19E-08 | -0.0307 | 0.0277 | 2.67E-01 | -0.0201 | 0.0106 | 5.83E-02 | 0 |
| rs720387 | 20 | 52465049 | 35 | 52448936 | 52471667 | T | C | 0.494 | SUMO1P1 | intergenic | 0.0748 | 0.0136 | 4.18E-08 | 0.0332 | 0.0276 | 2.29E-01 | 0.0203 | 0.0106 | 5.68E-02 | 0 |
| rs10432728 | 20 | 52466100 | 35 | 52448936 | 52471667 | T | C | 0.492 | SUMO1P1 | intergenic | -0.0785 | 0.0139 | 1.65E-08 | -0.0338 | 0.0276 | 2.20E-01 | -0.0238 | 0.0110 | 3.07E-02 | 0 |
| rs4811476 | 20 | 52468966 | 35 | 52448936 | 52471667 | T | C | 0.496 | SUMO1P1 | intergenic | 0.0801 | 0.0144 | 2.76E-08 | 0.0295 | 0.0274 | 2.82E-01 | 0.0235 | 0.0117 | 4.37E-02 | 0 |

* Based on genome-wide tests of inferred statistics computed from the roi-specific GWASs of cortical surface area[1] in 19,152 participants of ENIGMA study[2].

** Replicated (=1) if the effect size estimates from both 'replication' association tests have the same direction as that from the meta-GWAS and p-values < 0.05.

1 .Nieuwboer, H. A., Pool, R., Dolan, C. V., Boomsma, D. I. & Nivard, M. G. GWIS: Genome-wide inferred statistics for functions of multiple phenotypes. The American Journal of Human Genetics 99, 917–927 (2016).

2. Grasby, K. L. et al. The genetic architecture of the human cerebral cortex. bioRxiv 399402 (2018). doi:10.1101/399402

**Table E3. Associations between rs73313052 and surface area and thickness in each of the 34 cortical regions estimated from CHARGE- and ENIGMA-meta analyses**

| | Surface Area | | | | Thickness | | | |
|---|---|---|---|---|---|---|---|---|
| | CHARGE | | ENIGMA | | CHARGE | | ENIGMA | |
| ROI | z-score | p-value | z-score | p-value | z-score | p-value | z-score | p-value |
| cuneus | -12.072 | 1.48E-33 | -6.681 | 2.37E-11 | -5.535 | 3.11E-08 | 0.853 | 3.94E-01 |
| lateraloccipital | -10.608 | 2.73E-26 | -5.947 | 2.73E-09 | 1.869 | 6.16E-02 | 2.142 | 3.22E-02 |
| pericalcarine | -10.240 | 1.31E-24 | -8.031 | 9.65E-16 | -3.036 | 2.40E-03 | 0.697 | 4.86E-01 |
| precuneus | 10.212 | 1.75E-24 | 5.258 | 1.46E-07 | 1.641 | 1.01E-01 | -4.682 | 2.84E-06 |
| lingual | -9.420 | 4.50E-21 | -6.311 | 2.76E-10 | -0.266 | 7.90E-01 | -0.559 | 5.77E-01 |
| supramarginal | 7.374 | 1.66E-13 | 4.549 | 5.40E-06 | 3.084 | 2.04E-03 | -0.588 | 5.56E-01 |
| inferiortemporal | 4.769 | 1.85E-06 | 3.009 | 2.62E-03 | 2.684 | 7.28E-03 | 0.683 | 4.95E-01 |
| middletemporal | 4.581 | 4.64E-06 | 2.750 | 5.96E-03 | 1.865 | 6.21E-02 | 0.180 | 8.57E-01 |
| superiorfrontal | -4.247 | 2.17E-05 | -1.987 | 4.70E-02 | -1.062 | 2.88E-01 | 2.046 | 4.08E-02 |
| bankssts | 4.095 | 4.22E-05 | 2.342 | 1.92E-02 | 1.212 | 2.26E-01 | 0.783 | 4.34E-01 |
| precentral | -4.093 | 4.26E-05 | -1.370 | 1.71E-01 | -0.157 | 8.75E-01 | -1.132 | 2.58E-01 |
| superiorparietal | 3.999 | 6.35E-05 | 2.078 | 3.77E-02 | 3.959 | 7.53E-05 | -0.087 | 9.30E-01 |
| postcentral | 3.935 | 8.33E-05 | 1.947 | 5.15E-02 | 3.493 | 4.77E-04 | 0.080 | 9.36E-01 |
| inferiorparietal | 3.536 | 4.06E-04 | 1.605 | 1.08E-01 | 1.913 | 5.58E-02 | -1.319 | 1.87E-01 |
| parstriangularis | -3.314 | 9.18E-04 | -3.774 | 1.61E-04 | 1.156 | 2.48E-01 | 0.619 | 5.36E-01 |
| parsopercularis | -2.598 | 9.37E-03 | -1.851 | 6.42E-02 | 2.220 | 2.64E-02 | 0.379 | 7.05E-01 |
| paracentral | -2.281 | 2.26E-02 | -1.768 | 7.70E-02 | 0.248 | 8.04E-01 | -1.036 | 3.00E-01 |
| superiortemporal | 2.275 | 2.29E-02 | 1.613 | 1.07E-01 | 1.684 | 9.21E-02 | -3.430 | 6.03E-04 |
| lateralorbitofrontal | 2.050 | 4.03E-02 | 0.632 | 5.27E-01 | -0.900 | 3.68E-01 | 0.486 | 6.27E-01 |
| fusiform | -1.878 | 6.04E-02 | 1.047 | 2.95E-01 | 1.594 | 1.11E-01 | 1.303 | 1.93E-01 |
| frontalpole | -1.862 | 6.26E-02 | -2.907 | 3.65E-03 | -1.289 | 1.98E-01 | 0.080 | 9.36E-01 |
| temporalpole | 1.553 | 1.20E-01 | 1.475 | 1.40E-01 | 0.831 | 4.06E-01 | -4.767 | 1.87E-06 |
| isthmuscingulate | 1.400 | 1.61E-01 | 0.006 | 9.95E-01 | -0.241 | 8.10E-01 | 1.287 | 1.98E-01 |
| medialorbitofrontal | 1.397 | 1.62E-01 | -0.006 | 9.95E-01 | -0.566 | 5.72E-01 | 0.707 | 4.80E-01 |
| parahippocampal | 1.394 | 1.63E-01 | 1.434 | 1.52E-01 | -2.848 | 4.40E-03 | -1.649 | 9.91E-02 |
| insula | 1.096 | 2.73E-01 | -0.097 | 9.23E-01 | 0.473 | 6.36E-01 | 0.204 | 8.38E-01 |
| parsorbitalis | 0.737 | 4.61E-01 | -0.475 | 6.35E-01 | -0.848 | 3.96E-01 | 2.762 | 5.74E-03 |
| entorhinal | 0.726 | 4.68E-01 | 2.242 | 2.50E-02 | -0.749 | 4.54E-01 | 0.506 | 6.13E-01 |
| transversetemporal | -0.416 | 6.77E-01 | 0.096 | 9.23E-01 | 1.421 | 1.55E-01 | -0.196 | 8.45E-01 |
| caudalanteriorcingulate | -0.368 | 7.13E-01 | -0.209 | 8.35E-01 | 1.608 | 1.08E-01 | 0.673 | 5.01E-01 |
| rostralanteriorcingulate | 0.275 | 7.84E-01 | -0.243 | 8.08E-01 | 0.975 | 3.30E-01 | 2.002 | 4.53E-02 |
| posteriorcingulate | -0.167 | 8.67E-01 | -0.644 | 5.19E-01 | 1.610 | 1.07E-01 | 2.071 | 3.83E-02 |
| rostralmiddlefrontal | -0.123 | 9.03E-01 | -0.863 | 3.88E-01 | 0.189 | 8.50E-01 | 2.016 | 4.38E-02 |
| caudalmiddlefrontal | -0.120 | 9.05E-01 | -0.544 | 5.86E-01 | 1.381 | 1.67E-01 | 1.116 | 2.64E-01 |

**Table E4. *DAAM1* mRNA expression levels measured in different brain structures from the donors at different ages.** The mRNA expression levels were measured by RNA sequencing in 607 brain tissues from 18 female and 23 male donors available in BrainSpan database (http://www.brainspan.org/).

| Period | Braincode | Regioncode | Age | Days | Window | Sex | Hemisphere | pH | Ethnicity | Site | DAAM1 expression (RPKM) |
|---|---|---|---|---|---|---|---|---|---|---|---|
| 2 | HSB112 | OFC | 8PCW | 56 | 1 | M | R | 6.94 | European | USC | 8.327997 |
| 2 | HSB112 | DFC | 8PCW | 56 | 1 | M | R | 6.94 | European | USC | 7.567156 |
| 2 | HSB112 | VFC | 8PCW | 56 | 1 | M | R | 6.94 | European | USC | 10.358724 |
| 2 | HSB112 | MFC | 8PCW | 56 | 1 | M | R | 6.94 | European | USC | 7.741614 |
| 2 | HSB112 | M1CS1C | 8PCW | 56 | 1 | M | R | 6.94 | European | USC | 10.006544 |
| 2 | HSB112 | PC | 8PCW | 56 | 1 | M | R | 6.94 | European | USC | 9.827986 |
| 2 | HSB112 | STC | 8PCW | 56 | 1 | M | R | 6.94 | European | USC | 13.497321 |
| 2 | HSB112 | ITC | 8PCW | 56 | 1 | M | R | 6.94 | European | USC | 14.35899 |
| 2 | HSB112 | OC | 8PCW | 56 | 1 | M | R | 6.94 | European | USC | 9.626848 |
| 2 | HSB112 | HIP | 8PCW | 56 | 1 | M | R | 6.94 | European | USC | 10.136228 |
| 2 | HSB112 | AMY | 8PCW | 56 | 1 | M | R | 6.94 | European | USC | 16.567088 |
| 2 | HSB112 | CGE | 8PCW | 56 | 1 | M | R | 6.94 | European | USC | 10.28541 |
| 2 | HSB112 | LGE | 8PCW | 56 | 1 | M | R | 6.94 | European | USC | 12.209589 |
| 2 | HSB112 | MGE | 8PCW | 56 | 1 | M | R | 6.94 | European | USC | 9.685202 |
| 2 | HSB112 | DTH | 8PCW | 56 | 1 | M | R | 6.94 | European | USC | 15.558806 |
| 2 | HSB112 | URL | 8PCW | 56 | 1 | M | R | 6.94 | European | USC | 18.806048 |
| 2 | HSB148 | OFC | 9PCW | 63 | 1 | M | L | 6.01 | European | YALE | 10.194641 |
| 2 | HSB148 | DFC | 9PCW | 63 | 1 | M | L | 6.01 | European | YALE | 11.475317 |
| 2 | HSB148 | MFC | 9PCW | 63 | 1 | M | L | 6.01 | European | YALE | 11.267463 |
| 2 | HSB148 | M1CS1C | 9PCW | 63 | 1 | M | L | 6.01 | European | YALE | 8.912443 |
| 2 | HSB148 | PC | 9PCW | 63 | 1 | M | L | 6.01 | European | YALE | 14.485067 |
| 2 | HSB148 | TC | 9PCW | 63 | 1 | M | L | 6.01 | European | YALE | 17.739324 |
| 2 | HSB148 | OC | 9PCW | 63 | 1 | M | L | 6.01 | European | YALE | 13.050194 |
| 2 | HSB148 | HIP | 9PCW | 63 | 1 | M | L | 6.01 | European | YALE | 12.627581 |
| 2 | HSB148 | AMY | 9PCW | 63 | 1 | M | L | 6.01 | European | YALE | 12.422518 |
| 2 | HSB148 | CGE | 9PCW | 63 | 1 | M | L | 6.01 | European | YALE | 8.160755 |
| 2 | HSB148 | LGE | 9PCW | 63 | 1 | M | L | 6.01 | European | YALE | 11.736687 |
| 2 | HSB148 | MGE | 9PCW | 63 | 1 | M | L | 6.01 | European | YALE | 10.580124 |
| 2 | HSB148 | DTH | 9PCW | 63 | 1 | M | L | 6.01 | European | YALE | 19.44688 |
| 2 | HSB148 | URL | 9PCW | 63 | 1 | M | L | 6.01 | European | YALE | 22.65548 |
| 3 | HSB153 | OFC | 12PCW | 84 | 2 | F | R | NA | Asian | YALE | 28.259821 |
| 3 | HSB153 | DFC | 12PCW | 84 | 2 | F | R | NA | Asian | YALE | 26.14951 |
| 3 | HSB153 | VFC | 12PCW | 84 | 2 | F | R | NA | Asian | YALE | 31.027307 |
| 3 | HSB153 | MFC | 12PCW | 84 | 2 | F | R | NA | Asian | YALE | 26.079327 |
| 3 | HSB153 | M1C | 12PCW | 84 | 2 | F | R | NA | Asian | YALE | 25.785253 |
| 3 | HSB153 | S1C | 12PCW | 84 | 2 | F | R | NA | Asian | YALE | 19.780301 |
| 3 | HSB153 | IPC | 12PCW | 84 | 2 | F | R | NA | Asian | YALE | 15.089575 |
| 3 | HSB153 | A1C | 12PCW | 84 | 2 | F | R | NA | Asian | YALE | 17.167598 |
| 3 | HSB153 | STC | 12PCW | 84 | 2 | F | R | NA | Asian | YALE | 15.306079 |
| 3 | HSB153 | ITC | 12PCW | 84 | 2 | F | R | NA | Asian | YALE | 16.694582 |
| 3 | HSB153 | V1C | 12PCW | 84 | 2 | F | R | NA | Asian | YALE | 12.509778 |
| 3 | HSB153 | HIP | 12PCW | 84 | 2 | F | R | NA | Asian | YALE | 8.570237 |
| 3 | HSB153 | AMY | 12PCW | 84 | 2 | F | R | NA | Asian | YALE | 10.441828 |
| 3 | HSB153 | STR | 12PCW | 84 | 2 | F | R | NA | Asian | YALE | 19.514787 |
| 3 | HSB153 | MD | 12PCW | 84 | 2 | F | R | NA | Asian | YALE | 17.297369 |
| 3 | HSB150 | OFC | 12PCW | 84 | 2 | F | R | 6.08 | African | YALE | 19.810599 |

| | | | | | | | | | | | |
|---|---|---|---|---|---|---|---|---|---|---|---|
| 3 | HSB150 | DFC | 12PCW | 84 | 2 | F | R | 6.08 | African | YALE | 18.112009 |
| 3 | HSB150 | VFC | 12PCW | 84 | 2 | F | R | 6.08 | African | YALE | 24.501661 |
| 3 | HSB150 | MFC | 12PCW | 84 | 2 | F | R | 6.08 | African | YALE | 15.748532 |
| 3 | HSB150 | M1C | 12PCW | 84 | 2 | F | R | 6.08 | African | YALE | 27.057511 |
| 3 | HSB150 | S1C | 12PCW | 84 | 2 | F | R | 6.08 | African | YALE | 18.075817 |
| 3 | HSB150 | IPC | 12PCW | 84 | 2 | F | R | 6.08 | African | YALE | 18.702728 |
| 3 | HSB150 | A1C | 12PCW | 84 | 2 | F | R | 6.08 | African | YALE | 19.474021 |
| 3 | HSB150 | STC | 12PCW | 84 | 2 | F | R | 6.08 | African | YALE | 18.380044 |
| 3 | HSB150 | ITC | 12PCW | 84 | 2 | F | R | 6.08 | African | YALE | 18.589513 |
| 3 | HSB150 | V1C | 12PCW | 84 | 2 | F | R | 6.08 | African | YALE | 16.716658 |
| 3 | HSB150 | HIP | 12PCW | 84 | 2 | F | R | 6.08 | African | YALE | 10.376959 |
| 3 | HSB150 | AMY | 12PCW | 84 | 2 | F | R | 6.08 | African | YALE | 17.156571 |
| 3 | HSB150 | STR | 12PCW | 84 | 2 | F | R | 6.08 | African | YALE | 14.019839 |
| 3 | HSB150 | MD | 12PCW | 84 | 2 | F | R | 6.08 | African | YALE | 25.303198 |
| 3 | HSB150 | CBC | 12PCW | 84 | 2 | F | R | 6.08 | African | YALE | 23.304143 |
| 3 | HSB113 | OFC | 12PCW | 84 | 2 | F | R | 6.92 | Mix | USC | 26.364573 |
| 3 | HSB113 | DFC | 12PCW | 84 | 2 | F | R | 6.92 | Mix | USC | 23.179041 |
| 3 | HSB113 | VFC | 12PCW | 84 | 2 | F | R | 6.92 | Mix | USC | 21.296247 |
| 3 | HSB113 | MFC | 12PCW | 84 | 2 | F | R | 6.92 | Mix | USC | 25.175533 |
| 3 | HSB113 | M1C | 12PCW | 84 | 2 | F | R | 6.92 | Mix | USC | 20.850598 |
| 3 | HSB113 | S1C | 12PCW | 84 | 2 | F | R | 6.92 | Mix | USC | 16.996746 |
| 3 | HSB113 | IPC | 12PCW | 84 | 2 | F | R | 6.92 | Mix | USC | 13.884302 |
| 3 | HSB113 | A1C | 12PCW | 84 | 2 | F | R | 6.92 | Mix | USC | 21.050523 |
| 3 | HSB113 | ITC | 12PCW | 84 | 2 | F | R | 6.92 | Mix | USC | 14.768944 |
| 3 | HSB113 | V1C | 12PCW | 84 | 2 | F | R | 6.92 | Mix | USC | 12.698091 |
| 3 | HSB113 | HIP | 12PCW | 84 | 2 | F | R | 6.92 | Mix | USC | 10.144653 |
| 3 | HSB113 | AMY | 12PCW | 84 | 2 | F | R | 6.92 | Mix | USC | 15.565239 |
| 3 | HSB113 | STR | 12PCW | 84 | 2 | F | R | 6.92 | Mix | USC | 18.791417 |
| 3 | HSB113 | MD | 12PCW | 84 | 2 | F | R | 6.92 | Mix | USC | 17.695488 |
| 3 | HSB113 | CBC | 12PCW | 84 | 2 | F | R | 6.92 | Mix | USC | 16.102524 |
| 4 | HSB103 | OFC | 13PCW | 91 | 2 | M | R | NA | European | USC | 16.834071 |
| 4 | HSB103 | DFC | 13PCW | 91 | 2 | M | R | NA | European | USC | 17.618864 |
| 4 | HSB103 | VFC | 13PCW | 91 | 2 | M | R | NA | European | USC | 19.028234 |
| 4 | HSB103 | MFC | 13PCW | 91 | 2 | M | R | NA | European | USC | 12.156976 |
| 4 | HSB103 | M1C | 13PCW | 91 | 2 | M | R | NA | European | USC | 18.974216 |
| 4 | HSB103 | S1C | 13PCW | 91 | 2 | M | R | NA | European | USC | 16.488502 |
| 4 | HSB103 | IPC | 13PCW | 91 | 2 | M | R | NA | European | USC | 13.298987 |
| 4 | HSB103 | A1C | 13PCW | 91 | 2 | M | R | NA | European | USC | 17.449427 |
| 4 | HSB103 | ITC | 13PCW | 91 | 2 | M | R | NA | European | USC | 13.884302 |
| 4 | HSB103 | V1C | 13PCW | 91 | 2 | M | R | NA | European | USC | 12.935681 |
| 4 | HSB103 | HIP | 13PCW | 91 | 2 | M | R | NA | European | USC | 10.12817 |
| 4 | HSB103 | AMY | 13PCW | 91 | 2 | M | R | NA | European | USC | 12.560658 |
| 4 | HSB103 | STR | 13PCW | 91 | 2 | M | R | NA | European | USC | 15.1874 |
| 4 | HSB103 | CBC | 13PCW | 91 | 2 | M | R | NA | European | USC | 18.003576 |
| 4 | HSB149 | OFC | 13PCW | 91 | 2 | F | L | 6.39 | European | YALE | 32.522342 |
| 4 | HSB149 | DFC | 13PCW | 91 | 2 | F | L | 6.39 | European | YALE | 29.704147 |
| 4 | HSB149 | VFC | 13PCW | 91 | 2 | F | L | 6.39 | European | YALE | 31.164895 |
| 4 | HSB149 | MFC | 13PCW | 91 | 2 | F | L | 6.39 | European | YALE | 29.169501 |
| 4 | HSB149 | M1C | 13PCW | 91 | 2 | F | L | 6.39 | European | YALE | 29.689315 |
| 4 | HSB149 | S1C | 13PCW | 91 | 2 | F | L | 6.39 | European | YALE | 26.665326 |
| 4 | HSB149 | IPC | 13PCW | 91 | 2 | F | L | 6.39 | European | YALE | 20.752563 |

| | | | | | | | | | | | |
|---|---|---|---|---|---|---|---|---|---|---|---|
| 4 | HSB149 | A1C | 13PCW | 91 | 2 | F | L | 6.39 | European | YALE | 28.259821 |
| 4 | HSB149 | STC | 13PCW | 91 | 2 | F | L | 6.39 | European | YALE | 20.541546 |
| 4 | HSB149 | ITC | 13PCW | 91 | 2 | F | L | 6.39 | European | YALE | 17.956624 |
| 4 | HSB149 | V1C | 13PCW | 91 | 2 | F | L | 6.39 | European | YALE | 15.065999 |
| 4 | HSB149 | HIP | 13PCW | 91 | 2 | F | L | 6.39 | European | YALE | 12.499033 |
| 4 | HSB149 | AMY | 13PCW | 91 | 2 | F | L | 6.39 | European | YALE | 17.5139 |
| 4 | HSB149 | STR | 13PCW | 91 | 2 | F | L | 6.39 | European | YALE | 17.526137 |
| 4 | HSB149 | MD | 13PCW | 91 | 2 | F | L | 6.39 | European | YALE | 24.244192 |
| 4 | HSB149 | CBC | 13PCW | 91 | 2 | F | L | 6.39 | European | YALE | 19.63736 |
| 4 | HSB114 | OFC | 13PCW | 91 | 2 | M | L | NA | European | YALE | 23.871032 |
| 4 | HSB114 | DFC | 13PCW | 91 | 2 | M | L | NA | European | YALE | 21.651434 |
| 4 | HSB114 | VFC | 13PCW | 91 | 2 | M | L | NA | European | YALE | 24.398319 |
| 4 | HSB114 | MFC | 13PCW | 91 | 2 | M | L | NA | European | YALE | 23.929095 |
| 4 | HSB114 | M1C | 13PCW | 91 | 2 | M | L | NA | European | YALE | 18.329112 |
| 4 | HSB114 | S1C | 13PCW | 91 | 2 | M | L | NA | European | YALE | 18.862614 |
| 4 | HSB114 | IPC | 13PCW | 91 | 2 | M | L | NA | European | YALE | 16.037024 |
| 4 | HSB114 | A1C | 13PCW | 91 | 2 | M | L | NA | European | YALE | 17.658724 |
| 4 | HSB114 | STC | 13PCW | 91 | 2 | M | L | NA | European | YALE | 19.255977 |
| 4 | HSB114 | ITC | 13PCW | 91 | 2 | M | L | NA | European | YALE | 15.964999 |
| 4 | HSB114 | V1C | 13PCW | 91 | 2 | M | L | NA | European | YALE | 12.087327 |
| 4 | HSB114 | HIP | 13PCW | 91 | 2 | M | L | NA | European | YALE | 11.719409 |
| 4 | HSB114 | AMY | 13PCW | 91 | 2 | M | L | NA | European | YALE | 13.579365 |
| 4 | HSB114 | STR | 13PCW | 91 | 2 | M | L | NA | European | YALE | 21.500307 |
| 5 | HSB178 | OFC | 16PCW | 112 | 3 | M | L | 6.84 | European | YALE | 34.293141 |
| 5 | HSB178 | DFC | 16PCW | 112 | 3 | M | L | 6.84 | European | YALE | 29.266833 |
| 5 | HSB178 | VFC | 16PCW | 112 | 3 | M | L | 6.84 | European | YALE | 36.002427 |
| 5 | HSB178 | MFC | 16PCW | 112 | 3 | M | L | 6.84 | European | YALE | 24.172027 |
| 5 | HSB178 | M1C | 16PCW | 112 | 3 | M | L | 6.84 | European | YALE | 25.719117 |
| 5 | HSB178 | S1C | 16PCW | 112 | 3 | M | L | 6.84 | European | YALE | 25.435101 |
| 5 | HSB178 | IPC | 16PCW | 112 | 3 | M | L | 6.84 | European | YALE | 22.405292 |
| 5 | HSB178 | A1C | 16PCW | 112 | 3 | M | L | 6.84 | European | YALE | 29.070584 |
| 5 | HSB178 | STC | 16PCW | 112 | 3 | M | L | 6.84 | European | YALE | 24.553953 |
| 5 | HSB178 | ITC | 16PCW | 112 | 3 | M | L | 6.84 | European | YALE | 23.266972 |
| 5 | HSB178 | V1C | 16PCW | 112 | 3 | M | L | 6.84 | European | YALE | 11.322251 |
| 5 | HSB178 | HIP | 16PCW | 112 | 3 | M | L | 6.84 | European | YALE | 15.334903 |
| 5 | HSB178 | AMY | 16PCW | 112 | 3 | M | L | 6.84 | European | YALE | 13.562817 |
| 5 | HSB178 | STR | 16PCW | 112 | 3 | M | L | 6.84 | European | YALE | 23.096238 |
| 5 | HSB178 | MD | 16PCW | 112 | 3 | M | L | 6.84 | European | YALE | 25.546032 |
| 5 | HSB178 | CBC | 16PCW | 112 | 3 | M | L | 6.84 | European | YALE | 14.829206 |
| 5 | HSB154 | DFC | 16PCW | 112 | 3 | M | R | 6.44 | Mix | USC | 26.153699 |
| 5 | HSB154 | VFC | 16PCW | 112 | 3 | M | R | 6.44 | Mix | USC | 31.427562 |
| 5 | HSB154 | MFC | 16PCW | 112 | 3 | M | R | 6.44 | Mix | USC | 17.352897 |
| 5 | HSB154 | MSC | 16PCW | 112 | 3 | M | R | 6.44 | Mix | USC | 26.929878 |
| 5 | HSB154 | IPC | 16PCW | 112 | 3 | M | R | 6.44 | Mix | USC | 29.715362 |
| 5 | HSB154 | A1C | 16PCW | 112 | 3 | M | R | 6.44 | Mix | USC | 34.311121 |
| 5 | HSB154 | STC | 16PCW | 112 | 3 | M | R | 6.44 | Mix | USC | 30.359123 |
| 5 | HSB154 | ITC | 16PCW | 112 | 3 | M | R | 6.44 | Mix | USC | 26.207946 |
| 5 | HSB154 | V1C | 16PCW | 112 | 3 | M | R | 6.44 | Mix | USC | 23.335906 |
| 5 | HSB154 | HIP | 16PCW | 112 | 3 | M | R | 6.44 | Mix | USC | 11.439061 |
| 5 | HSB154 | AMY | 16PCW | 112 | 3 | M | R | 6.44 | Mix | USC | 19.855869 |
| 5 | HSB154 | STR | 16PCW | 112 | 3 | M | R | 6.44 | Mix | USC | 16.71495 |

| | | | | | | | | | | | |
|---|---|---|---|---|---|---|---|---|---|---|---|
| 5 | HSB154 | MD | 16PCW | 112 | 3 | M | R | 6.44 | Mix | USC | 17.211447 |
| 5 | HSB96 | DFC | 16PCW | 112 | 3 | M | R | NA | Mexican | YALE | 20.143533 |
| 5 | HSB96 | VFC | 16PCW | 112 | 3 | M | R | NA | Mexican | YALE | 24.628536 |
| 5 | HSB96 | MFC | 16PCW | 112 | 3 | M | R | NA | Mexican | YALE | 13.317262 |
| 5 | HSB96 | MSC | 16PCW | 112 | 3 | M | R | NA | Mexican | YALE | 25.672903 |
| 5 | HSB96 | IPC | 16PCW | 112 | 3 | M | R | NA | Mexican | YALE | 29.586584 |
| 5 | HSB96 | A1C | 16PCW | 112 | 3 | M | R | NA | Mexican | YALE | 30.899769 |
| 5 | HSB96 | STC | 16PCW | 112 | 3 | M | R | NA | Mexican | YALE | 18.265117 |
| 5 | HSB96 | V1C | 16PCW | 112 | 3 | M | R | NA | Mexican | YALE | 20.760636 |
| 5 | HSB96 | HIP | 16PCW | 112 | 3 | M | R | NA | Mexican | YALE | 9.35578 |
| 5 | HSB96 | STR | 16PCW | 112 | 3 | M | R | NA | Mexican | YALE | 16.188317 |
| 5 | HSB96 | MD | 16PCW | 112 | 3 | M | R | NA | Mexican | YALE | 20.596399 |
| 5 | HSB97 | OFC | 17PCW | 119 | 3 | F | L | NA | European | USC | 30.876259 |
| 5 | HSB97 | DFC | 17PCW | 119 | 3 | F | L | NA | European | USC | 26.973489 |
| 5 | HSB97 | VFC | 17PCW | 119 | 3 | F | L | NA | European | USC | 26.253739 |
| 5 | HSB97 | MFC | 17PCW | 119 | 3 | F | L | NA | European | USC | 23.067623 |
| 5 | HSB97 | MSC | 17PCW | 119 | 3 | F | L | NA | European | USC | 25.606731 |
| 5 | HSB97 | IPC | 17PCW | 119 | 3 | F | L | NA | European | USC | 29.433934 |
| 5 | HSB97 | A1C | 17PCW | 119 | 3 | F | L | NA | European | USC | 30.702751 |
| 5 | HSB97 | STC | 17PCW | 119 | 3 | F | L | NA | European | USC | 25.423641 |
| 5 | HSB97 | V1C | 17PCW | 119 | 3 | F | L | NA | European | USC | 19.265463 |
| 5 | HSB97 | HIP | 17PCW | 119 | 3 | F | L | NA | European | USC | 13.141321 |
| 5 | HSB97 | AMY | 17PCW | 119 | 3 | F | L | NA | European | USC | 17.400362 |
| 5 | HSB97 | STR | 17PCW | 119 | 3 | F | L | NA | European | USC | 14.216526 |
| 5 | HSB97 | MD | 17PCW | 119 | 3 | F | L | NA | European | USC | 20.625169 |
| 5 | HSB97 | CBC | 17PCW | 119 | 3 | F | L | NA | European | USC | 17.591509 |
| 6 | HSB98 | DFC | 19PCW | 133 | 4 | F | R | NA | Mexican | USC | 29.203837 |
| 6 | HSB98 | VFC | 19PCW | 133 | 4 | F | R | NA | Mexican | USC | 25.808878 |
| 6 | HSB98 | MFC | 19PCW | 133 | 4 | F | R | NA | Mexican | USC | 19.556002 |
| 6 | HSB98 | MSC | 19PCW | 133 | 4 | F | R | NA | Mexican | USC | 35.789455 |
| 6 | HSB98 | IPC | 19PCW | 133 | 4 | F | R | NA | Mexican | USC | 41.227347 |
| 6 | HSB98 | A1C | 19PCW | 133 | 4 | F | R | NA | Mexican | USC | 33.515347 |
| 6 | HSB98 | STC | 19PCW | 133 | 4 | F | R | NA | Mexican | USC | 32.759005 |
| 6 | HSB98 | V1C | 19PCW | 133 | 4 | F | R | NA | Mexican | USC | 36.805893 |
| 6 | HSB98 | HIP | 19PCW | 133 | 4 | F | R | NA | Mexican | USC | 14.518952 |
| 6 | HSB98 | STR | 19PCW | 133 | 4 | F | R | NA | Mexican | USC | 17.585147 |
| 6 | HSB98 | MD | 19PCW | 133 | 4 | F | R | NA | Mexican | USC | 24.538151 |
| 6 | HSB107 | OFC | 21PCW | 147 | 4 | F | L | 6.61 | European | USC | 26.395334 |
| 6 | HSB107 | DFC | 21PCW | 147 | 4 | F | L | 6.61 | European | USC | 27.656453 |
| 6 | HSB107 | VFC | 21PCW | 147 | 4 | F | L | 6.61 | European | USC | 27.834155 |
| 6 | HSB107 | MFC | 21PCW | 147 | 4 | F | L | 6.61 | European | USC | 24.702827 |
| 6 | HSB107 | M1C | 21PCW | 147 | 4 | F | L | 6.61 | European | USC | 27.866996 |
| 6 | HSB107 | S1C | 21PCW | 147 | 4 | F | L | 6.61 | European | USC | 27.642222 |
| 6 | HSB107 | IPC | 21PCW | 147 | 4 | F | L | 6.61 | European | USC | 31.847267 |
| 6 | HSB107 | A1C | 21PCW | 147 | 4 | F | L | 6.61 | European | USC | 33.796603 |
| 6 | HSB107 | STC | 21PCW | 147 | 4 | F | L | 6.61 | European | USC | 25.461259 |
| 6 | HSB107 | ITC | 21PCW | 147 | 4 | F | L | 6.61 | European | USC | 22.496952 |
| 6 | HSB107 | V1C | 21PCW | 147 | 4 | F | L | 6.61 | European | USC | 18.55174 |
| 6 | HSB107 | HIP | 21PCW | 147 | 4 | F | L | 6.61 | European | USC | 14.920385 |
| 6 | HSB107 | AMY | 21PCW | 147 | 4 | F | L | 6.61 | European | USC | 16.500266 |
| 6 | HSB107 | STR | 21PCW | 147 | 4 | F | L | 6.61 | European | USC | 14.221259 |

| | | | | | | | | | | | |
|---|---|---|---|---|---|---|---|---|---|---|---|
| 6 | HSB107 | MD | 21PCW | 147 | 4 | F | L | 6.61 | European | USC | 14.122427 |
| 6 | HSB107 | CBC | 21PCW | 147 | 4 | F | L | 6.61 | European | USC | 15.680462 |
| 6 | HSB92 | OFC | 21PCW | 147 | 4 | M | R | 6.65 | African | USC | 33.377866 |
| 6 | HSB92 | DFC | 21PCW | 147 | 4 | M | R | 6.65 | African | USC | 27.900993 |
| 6 | HSB92 | VFC | 21PCW | 147 | 4 | M | R | 6.65 | African | USC | 34.335944 |
| 6 | HSB92 | MFC | 21PCW | 147 | 4 | M | R | 6.65 | African | USC | 32.181025 |
| 6 | HSB92 | M1C | 21PCW | 147 | 4 | M | R | 6.65 | African | USC | 27.818696 |
| 6 | HSB92 | S1C | 21PCW | 147 | 4 | M | R | 6.65 | African | USC | 25.904151 |
| 6 | HSB92 | IPC | 21PCW | 147 | 4 | M | R | 6.65 | African | USC | 26.453438 |
| 6 | HSB92 | STC | 21PCW | 147 | 4 | M | R | 6.65 | African | USC | 28.595321 |
| 6 | HSB92 | ITC | 21PCW | 147 | 4 | M | R | 6.65 | African | USC | 18.202019 |
| 6 | HSB92 | V1C | 21PCW | 147 | 4 | M | R | 6.65 | African | USC | 32.22296 |
| 6 | HSB92 | HIP | 21PCW | 147 | 4 | M | R | 6.65 | African | USC | 15.9742 |
| 6 | HSB92 | AMY | 21PCW | 147 | 4 | M | R | 6.65 | African | USC | 21.685982 |
| 6 | HSB92 | STR | 21PCW | 147 | 4 | M | R | 6.65 | African | USC | 13.617009 |
| 6 | HSB92 | MD | 21PCW | 147 | 4 | M | R | 6.65 | African | USC | 7.915952 |
| 6 | HSB92 | CBC | 21PCW | 147 | 4 | M | R | 6.65 | African | USC | 13.939831 |
| 6 | HSB159 | OFC | 22PCW | 154 | 4 | M | L | 6.58 | European | YALE | 32.522342 |
| 6 | HSB159 | DFC | 22PCW | 154 | 4 | M | L | 6.58 | European | YALE | 35.17769 |
| 6 | HSB159 | VFC | 22PCW | 154 | 4 | M | L | 6.58 | European | YALE | 33.448268 |
| 6 | HSB159 | MFC | 22PCW | 154 | 4 | M | L | 6.58 | European | YALE | 28.472076 |
| 6 | HSB159 | M1C | 22PCW | 154 | 4 | M | L | 6.58 | European | YALE | 23.324971 |
| 6 | HSB159 | S1C | 22PCW | 154 | 4 | M | L | 6.58 | European | YALE | 29.559149 |
| 6 | HSB159 | IPC | 22PCW | 154 | 4 | M | L | 6.58 | European | YALE | 32.157059 |
| 6 | HSB159 | A1C | 22PCW | 154 | 4 | M | L | 6.58 | European | YALE | 32.785345 |
| 6 | HSB159 | STC | 22PCW | 154 | 4 | M | L | 6.58 | European | YALE | 28.792447 |
| 6 | HSB159 | ITC | 22PCW | 154 | 4 | M | L | 6.58 | European | YALE | 31.624659 |
| 6 | HSB159 | V1C | 22PCW | 154 | 4 | M | L | 6.58 | European | YALE | 20.182605 |
| 6 | HSB159 | HIP | 22PCW | 154 | 4 | M | L | 6.58 | European | YALE | 12.534884 |
| 6 | HSB159 | AMY | 22PCW | 154 | 4 | M | L | 6.58 | European | YALE | 12.761413 |
| 6 | HSB159 | STR | 22PCW | 154 | 4 | M | L | 6.58 | European | YALE | 13.103593 |
| 6 | HSB159 | MD | 22PCW | 154 | 4 | M | L | 6.58 | European | YALE | 14.131577 |
| 6 | HSB159 | CBC | 22PCW | 154 | 4 | M | L | 6.58 | European | YALE | 12.892898 |
| 7 | HSB155 | OFC | 35PCW | 245 | 5 | F | R | 6.67 | European | YALE | 13.301226 |
| 7 | HSB155 | DFC | 35PCW | 245 | 5 | F | R | 6.67 | European | YALE | 9.378335 |
| 7 | HSB155 | VFC | 35PCW | 245 | 5 | F | R | 6.67 | European | YALE | 18.082105 |
| 7 | HSB155 | MFC | 35PCW | 245 | 5 | F | R | 6.67 | European | YALE | 3.648535 |
| 7 | HSB155 | M1C | 35PCW | 245 | 5 | F | R | 6.67 | European | YALE | 6.574262 |
| 7 | HSB155 | S1C | 35PCW | 245 | 5 | F | R | 6.67 | European | YALE | 12.638567 |
| 7 | HSB155 | IPC | 35PCW | 245 | 5 | F | R | 6.67 | European | YALE | 4.082129 |
| 7 | HSB155 | ITC | 35PCW | 245 | 5 | F | R | 6.67 | European | YALE | 15.156004 |
| 7 | HSB155 | HIP | 35PCW | 245 | 5 | F | R | 6.67 | European | YALE | 5.893915 |
| 7 | HSB155 | AMY | 35PCW | 245 | 5 | F | R | 6.67 | European | YALE | 4.007573 |
| 7 | HSB155 | STR | 35PCW | 245 | 5 | F | R | 6.67 | European | YALE | 5.517204 |
| 7 | HSB155 | MD | 35PCW | 245 | 5 | F | R | 6.67 | European | YALE | 4.664362 |
| 7 | HSB155 | CBC | 35PCW | 245 | 5 | F | R | 6.67 | European | YALE | 7.931364 |
| 7 | HSB194 | OFC | 37PCW | 259 | 5 | M | L | 6.12 | European | YALE | 18.020502 |
| 7 | HSB194 | DFC | 37PCW | 259 | 5 | M | L | 6.12 | European | YALE | 18.669269 |
| 7 | HSB194 | VFC | 37PCW | 259 | 5 | M | L | 6.12 | European | YALE | 16.372127 |
| 7 | HSB194 | MFC | 37PCW | 259 | 5 | M | L | 6.12 | European | YALE | 14.909251 |
| 7 | HSB194 | M1C | 37PCW | 259 | 5 | M | L | 6.12 | European | YALE | 10.205915 |

| | | | | | | | | | | | |
|---|---|---|---|---|---|---|---|---|---|---|---|
| 7 | HSB194 | S1C | 37PCW | 259 | 5 | M | L | 6.12 | European | YALE | 12.590515 |
| 7 | HSB194 | IPC | 37PCW | 259 | 5 | M | L | 6.12 | European | YALE | 18.194611 |
| 7 | HSB194 | A1C | 37PCW | 259 | 5 | M | L | 6.12 | European | YALE | 16.689455 |
| 7 | HSB194 | STC | 37PCW | 259 | 5 | M | L | 6.12 | European | YALE | 19.951419 |
| 7 | HSB194 | ITC | 37PCW | 259 | 5 | M | L | 6.12 | European | YALE | 20.355636 |
| 7 | HSB194 | V1C | 37PCW | 259 | 5 | M | L | 6.12 | European | YALE | 17.413436 |
| 7 | HSB194 | HIP | 37PCW | 259 | 5 | M | L | 6.12 | European | YALE | 5.907369 |
| 7 | HSB194 | AMY | 37PCW | 259 | 5 | M | L | 6.12 | European | YALE | 6.126825 |
| 7 | HSB194 | STR | 37PCW | 259 | 5 | M | L | 6.12 | European | YALE | 3.656252 |
| 7 | HSB194 | MD | 37PCW | 259 | 5 | M | L | 6.12 | European | YALE | 3.34586 |
| 7 | HSB194 | CBC | 37PCW | 259 | 5 | M | L | 6.12 | European | YALE | 9.662803 |
| 8 | HSB121 | OFC | 4M | 386 | 5 | M | R | 6.26 | European | YALE | 5.631735 |
| 8 | HSB121 | DFC | 4M | 386 | 5 | M | R | 6.26 | European | YALE | 6.654639 |
| 8 | HSB121 | VFC | 4M | 386 | 5 | M | R | 6.26 | European | YALE | 6.11308 |
| 8 | HSB121 | MFC | 4M | 386 | 5 | M | R | 6.26 | European | YALE | 5.034661 |
| 8 | HSB121 | M1C | 4M | 386 | 5 | M | R | 6.26 | European | YALE | 4.599102 |
| 8 | HSB121 | S1C | 4M | 386 | 5 | M | R | 6.26 | European | YALE | 6.308283 |
| 8 | HSB121 | IPC | 4M | 386 | 5 | M | R | 6.26 | European | YALE | 6.41227 |
| 8 | HSB121 | A1C | 4M | 386 | 5 | M | R | 6.26 | European | YALE | 4.933568 |
| 8 | HSB121 | STC | 4M | 386 | 5 | M | R | 6.26 | European | YALE | 6.384937 |
| 8 | HSB121 | ITC | 4M | 386 | 5 | M | R | 6.26 | European | YALE | 6.830393 |
| 8 | HSB121 | V1C | 4M | 386 | 5 | M | R | 6.26 | European | YALE | 6.263897 |
| 8 | HSB121 | HIP | 4M | 386 | 5 | M | R | 6.26 | European | YALE | 9.670572 |
| 8 | HSB121 | AMY | 4M | 386 | 5 | M | R | 6.26 | European | YALE | 5.268768 |
| 8 | HSB121 | STR | 4M | 386 | 5 | M | R | 6.26 | European | YALE | 3.687353 |
| 8 | HSB121 | MD | 4M | 386 | 5 | M | R | 6.26 | European | YALE | 2.833661 |
| 8 | HSB132 | OFC | 4M | 386 | 5 | M | L | 6.6 | European | USC | 8.970919 |
| 8 | HSB132 | DFC | 4M | 386 | 5 | M | L | 6.6 | European | USC | 10.38789 |
| 8 | HSB132 | VFC | 4M | 386 | 5 | M | L | 6.6 | European | USC | 10.062818 |
| 8 | HSB132 | MFC | 4M | 386 | 5 | M | L | 6.6 | European | USC | 9.876871 |
| 8 | HSB132 | M1C | 4M | 386 | 5 | M | L | 6.6 | European | USC | 9.085435 |
| 8 | HSB132 | S1C | 4M | 386 | 5 | M | L | 6.6 | European | USC | 8.925972 |
| 8 | HSB132 | IPC | 4M | 386 | 5 | M | L | 6.6 | European | USC | 8.423549 |
| 8 | HSB132 | A1C | 4M | 386 | 5 | M | L | 6.6 | European | USC | 8.937675 |
| 8 | HSB132 | STC | 4M | 386 | 5 | M | L | 6.6 | European | USC | 9.539147 |
| 8 | HSB132 | ITC | 4M | 386 | 5 | M | L | 6.6 | European | USC | 9.680093 |
| 8 | HSB132 | V1C | 4M | 386 | 5 | M | L | 6.6 | European | USC | 9.658148 |
| 8 | HSB132 | HIP | 4M | 386 | 5 | M | L | 6.6 | European | USC | 6.968261 |
| 8 | HSB132 | AMY | 4M | 386 | 5 | M | L | 6.6 | European | USC | 6.825136 |
| 8 | HSB132 | STR | 4M | 386 | 5 | M | L | 6.6 | European | USC | 3.013502 |
| 8 | HSB132 | MD | 4M | 386 | 5 | M | L | 6.6 | European | USC | 6.005718 |
| 8 | HSB132 | CBC | 4M | 386 | 5 | M | L | 6.6 | European | USC | 8.559966 |
| 8 | HSB139 | OFC | 4M | 386 | 5 | M | L | 6.7 | African | YALE | 10.376959 |
| 8 | HSB139 | DFC | 4M | 386 | 5 | M | L | 6.7 | African | YALE | 12.114945 |
| 8 | HSB139 | VFC | 4M | 386 | 5 | M | L | 6.7 | African | YALE | 10.191869 |
| 8 | HSB139 | MFC | 4M | 386 | 5 | M | L | 6.7 | African | YALE | 12.227983 |
| 8 | HSB139 | M1C | 4M | 386 | 5 | M | L | 6.7 | African | YALE | 8.50957 |
| 8 | HSB139 | S1C | 4M | 386 | 5 | M | L | 6.7 | African | YALE | 11.573531 |
| 8 | HSB139 | IPC | 4M | 386 | 5 | M | L | 6.7 | African | YALE | 10.278468 |
| 8 | HSB139 | A1C | 4M | 386 | 5 | M | L | 6.7 | African | YALE | 11.204799 |
| 8 | HSB139 | STC | 4M | 386 | 5 | M | L | 6.7 | African | YALE | 10.565526 |

| | | | | | | | | | | | |
|---|---|---|---|---|---|---|---|---|---|---|---|
| 8 | HSB139 | ITC | 4M | 386 | 5 | M | L | 6.7 | African | YALE | 13.042506 |
| 8 | HSB139 | V1C | 4M | 386 | 5 | M | L | 6.7 | African | YALE | 12.524239 |
| 8 | HSB139 | HIP | 4M | 386 | 5 | M | L | 6.7 | African | YALE | 6.813121 |
| 8 | HSB139 | AMY | 4M | 386 | 5 | M | L | 6.7 | African | YALE | 7.631385 |
| 8 | HSB139 | STR | 4M | 386 | 5 | M | L | 6.7 | African | YALE | 3.469339 |
| 8 | HSB139 | MD | 4M | 386 | 5 | M | L | 6.7 | African | YALE | 5.916255 |
| 8 | HSB139 | CBC | 4M | 386 | 5 | M | L | 6.7 | African | YALE | 9.707269 |
| 9 | HSB131 | OFC | 6M | 446 | 6 | F | L | 6.26 | European | USC | 8.966365 |
| 9 | HSB131 | DFC | 6M | 446 | 6 | F | L | 6.26 | European | USC | 11.106435 |
| 9 | HSB131 | VFC | 6M | 446 | 6 | F | L | 6.26 | European | USC | 8.421427 |
| 9 | HSB131 | MFC | 6M | 446 | 6 | F | L | 6.26 | European | USC | 9.245956 |
| 9 | HSB131 | M1C | 6M | 446 | 6 | F | L | 6.26 | European | USC | 7.249009 |
| 9 | HSB131 | S1C | 6M | 446 | 6 | F | L | 6.26 | European | USC | 10.833447 |
| 9 | HSB131 | IPC | 6M | 446 | 6 | F | L | 6.26 | European | USC | 10.396562 |
| 9 | HSB131 | A1C | 6M | 446 | 6 | F | L | 6.26 | European | USC | 7.688422 |
| 9 | HSB131 | STC | 6M | 446 | 6 | F | L | 6.26 | European | USC | 11.984958 |
| 9 | HSB131 | ITC | 6M | 446 | 6 | F | L | 6.26 | European | USC | 16.446519 |
| 9 | HSB131 | V1C | 6M | 446 | 6 | F | L | 6.26 | European | USC | 18.309513 |
| 9 | HSB131 | HIP | 6M | 446 | 6 | F | L | 6.26 | European | USC | 7.443317 |
| 9 | HSB131 | AMY | 6M | 446 | 6 | F | L | 6.26 | European | USC | 10.167935 |
| 9 | HSB131 | STR | 6M | 446 | 6 | F | L | 6.26 | European | USC | 7.105242 |
| 9 | HSB131 | MD | 6M | 446 | 6 | F | L | 6.26 | European | USC | 6.385896 |
| 9 | HSB131 | CBC | 6M | 446 | 6 | F | L | 6.26 | European | USC | 7.237984 |
| 9 | HSB171 | OFC | 10M | 566 | 6 | M | L | 5.96 | African | USC | 7.366706 |
| 9 | HSB171 | DFC | 10M | 566 | 6 | M | L | 5.96 | African | USC | 7.694501 |
| 9 | HSB171 | VFC | 10M | 566 | 6 | M | L | 5.96 | African | USC | 7.845031 |
| 9 | HSB171 | MFC | 10M | 566 | 6 | M | L | 5.96 | African | USC | 8.297435 |
| 9 | HSB171 | S1C | 10M | 566 | 6 | M | L | 5.96 | African | USC | 7.135037 |
| 9 | HSB171 | IPC | 10M | 566 | 6 | M | L | 5.96 | African | USC | 7.094436 |
| 9 | HSB171 | STC | 10M | 566 | 6 | M | L | 5.96 | African | USC | 7.311708 |
| 9 | HSB171 | ITC | 10M | 566 | 6 | M | L | 5.96 | African | USC | 8.332271 |
| 9 | HSB171 | V1C | 10M | 566 | 6 | M | L | 5.96 | African | USC | 7.865179 |
| 9 | HSB171 | MD | 10M | 566 | 6 | M | L | 5.96 | African | USC | 6.525674 |
| 9 | HSB171 | CBC | 10M | 566 | 6 | M | L | 5.96 | African | USC | 10.028196 |
| 9 | HSB122 | OFC | 1Y | 631 | 6 | F | R | 6.68 | European | YALE | 7.333525 |
| 9 | HSB122 | DFC | 1Y | 631 | 6 | F | R | 6.68 | European | YALE | 8.230892 |
| 9 | HSB122 | VFC | 1Y | 631 | 6 | F | R | 6.68 | European | YALE | 7.8836 |
| 9 | HSB122 | MFC | 1Y | 631 | 6 | F | R | 6.68 | European | YALE | 7.767694 |
| 9 | HSB122 | M1C | 1Y | 631 | 6 | F | R | 6.68 | European | YALE | 8.012522 |
| 9 | HSB122 | S1C | 1Y | 631 | 6 | F | R | 6.68 | European | YALE | 8.269261 |
| 9 | HSB122 | IPC | 1Y | 631 | 6 | F | R | 6.68 | European | YALE | 8.474032 |
| 9 | HSB122 | A1C | 1Y | 631 | 6 | F | R | 6.68 | European | YALE | 8.436835 |
| 9 | HSB122 | STC | 1Y | 631 | 6 | F | R | 6.68 | European | YALE | 8.290631 |
| 9 | HSB122 | ITC | 1Y | 631 | 6 | F | R | 6.68 | European | YALE | 8.789406 |
| 9 | HSB122 | V1C | 1Y | 631 | 6 | F | R | 6.68 | European | YALE | 8.856047 |
| 9 | HSB122 | HIP | 1Y | 631 | 6 | F | R | 6.68 | European | YALE | 6.828605 |
| 9 | HSB122 | AMY | 1Y | 631 | 6 | F | R | 6.68 | European | YALE | 6.887542 |
| 9 | HSB122 | STR | 1Y | 631 | 6 | F | R | 6.68 | European | YALE | 2.771957 |
| 9 | HSB122 | MD | 1Y | 631 | 6 | F | R | 6.68 | European | YALE | 5.952039 |
| 9 | HSB122 | CBC | 1Y | 631 | 6 | F | R | 6.68 | European | YALE | 10.15297 |
| 10 | HSB143 | OFC | 2Y | 996 | 6 | F | R | 6.3 | European | USC | 8.35185 |

| | | | | | | | | | | | |
|---|---|---|---|---|---|---|---|---|---|---|---|
| 10 | HSB143 | DFC | 2Y | 996 | 6 | F | R | 6.3 | European | USC | 4.638739 |
| 10 | HSB143 | VFC | 2Y | 996 | 6 | F | R | 6.3 | European | USC | 4.805498 |
| 10 | HSB143 | MFC | 2Y | 996 | 6 | F | R | 6.3 | European | USC | 6.90234 |
| 10 | HSB143 | M1C | 2Y | 996 | 6 | F | R | 6.3 | European | USC | 4.205331 |
| 10 | HSB143 | S1C | 2Y | 996 | 6 | F | R | 6.3 | European | USC | 3.457371 |
| 10 | HSB143 | IPC | 2Y | 996 | 6 | F | R | 6.3 | European | USC | 5.939985 |
| 10 | HSB143 | A1C | 2Y | 996 | 6 | F | R | 6.3 | European | USC | 7.942027 |
| 10 | HSB143 | STC | 2Y | 996 | 6 | F | R | 6.3 | European | USC | 4.371088 |
| 10 | HSB143 | ITC | 2Y | 996 | 6 | F | R | 6.3 | European | USC | 7.849039 |
| 10 | HSB143 | V1C | 2Y | 996 | 6 | F | R | 6.3 | European | USC | 8.172615 |
| 10 | HSB143 | HIP | 2Y | 996 | 6 | F | R | 6.3 | European | USC | 5.349472 |
| 10 | HSB143 | AMY | 2Y | 996 | 6 | F | R | 6.3 | European | USC | 8.810844 |
| 10 | HSB143 | STR | 2Y | 996 | 6 | F | R | 6.3 | European | USC | 3.448714 |
| 10 | HSB143 | MD | 2Y | 996 | 6 | F | R | 6.3 | European | USC | 3.122318 |
| 10 | HSB143 | CBC | 2Y | 996 | 6 | F | R | 6.3 | European | USC | 6.344071 |
| 10 | HSB173 | VFC | 3Y | 1361 | 6 | F | L | 6.03 | European | YALE | 4.188513 |
| 10 | HSB173 | M1C | 3Y | 1361 | 6 | F | L | 6.03 | European | YALE | 4.486824 |
| 10 | HSB173 | S1C | 3Y | 1361 | 6 | F | L | 6.03 | European | YALE | 4.33492 |
| 10 | HSB173 | IPC | 3Y | 1361 | 6 | F | L | 6.03 | European | YALE | 4.786953 |
| 10 | HSB173 | A1C | 3Y | 1361 | 6 | F | L | 6.03 | European | YALE | 4.535053 |
| 10 | HSB173 | STC | 3Y | 1361 | 6 | F | L | 6.03 | European | YALE | 4.94242 |
| 10 | HSB173 | ITC | 3Y | 1361 | 6 | F | L | 6.03 | European | YALE | 4.315792 |
| 10 | HSB173 | V1C | 3Y | 1361 | 6 | F | L | 6.03 | European | YALE | 4.046294 |
| 10 | HSB173 | HIP | 3Y | 1361 | 6 | F | L | 6.03 | European | YALE | 3.584834 |
| 10 | HSB173 | AMY | 3Y | 1361 | 6 | F | L | 6.03 | European | YALE | 4.205783 |
| 10 | HSB173 | STR | 3Y | 1361 | 6 | F | L | 6.03 | European | YALE | 2.249351 |
| 10 | HSB173 | MD | 3Y | 1361 | 6 | F | L | 6.03 | European | YALE | 3.320275 |
| 10 | HSB173 | CBC | 3Y | 1361 | 6 | F | L | 6.03 | European | YALE | 6.469318 |
| 10 | HSB172 | OFC | 3Y | 1361 | 7 | M | L | 5.69 | Mexican | USC | 4.718603 |
| 10 | HSB172 | DFC | 3Y | 1361 | 7 | M | L | 5.69 | Mexican | USC | 3.821081 |
| 10 | HSB172 | VFC | 3Y | 1361 | 7 | M | L | 5.69 | Mexican | USC | 4.035348 |
| 10 | HSB172 | MFC | 3Y | 1361 | 7 | M | L | 5.69 | Mexican | USC | 3.965499 |
| 10 | HSB172 | M1C | 3Y | 1361 | 7 | M | L | 5.69 | Mexican | USC | 3.449601 |
| 10 | HSB172 | S1C | 3Y | 1361 | 7 | M | L | 5.69 | Mexican | USC | 4.009949 |
| 10 | HSB172 | IPC | 3Y | 1361 | 7 | M | L | 5.69 | Mexican | USC | 3.658882 |
| 10 | HSB172 | A1C | 3Y | 1361 | 7 | M | L | 5.69 | Mexican | USC | 3.980016 |
| 10 | HSB172 | STC | 3Y | 1361 | 7 | M | L | 5.69 | Mexican | USC | 3.541569 |
| 10 | HSB172 | ITC | 3Y | 1361 | 7 | M | L | 5.69 | Mexican | USC | 4.110632 |
| 10 | HSB172 | V1C | 3Y | 1361 | 7 | M | L | 5.69 | Mexican | USC | 4.459434 |
| 10 | HSB172 | HIP | 3Y | 1361 | 7 | M | L | 5.69 | Mexican | USC | 2.964551 |
| 10 | HSB172 | AMY | 3Y | 1361 | 7 | M | L | 5.69 | Mexican | USC | 4.218646 |
| 10 | HSB172 | CBC | 3Y | 1361 | 7 | M | L | 5.69 | Mexican | USC | 6.968261 |
| 10 | HSB118 | OFC | 4Y | 1726 | 7 | M | R | 6.52 | African | USC | 4.60759 |
| 10 | HSB118 | DFC | 4Y | 1726 | 7 | M | R | 6.52 | African | USC | 4.502624 |
| 10 | HSB118 | VFC | 4Y | 1726 | 7 | M | R | 6.52 | African | USC | 5.396023 |
| 10 | HSB118 | MFC | 4Y | 1726 | 7 | M | R | 6.52 | African | USC | 4.856853 |
| 10 | HSB118 | M1C | 4Y | 1726 | 7 | M | R | 6.52 | African | USC | 4.526518 |
| 10 | HSB118 | S1C | 4Y | 1726 | 7 | M | R | 6.52 | African | USC | 5.457837 |
| 10 | HSB118 | IPC | 4Y | 1726 | 7 | M | R | 6.52 | African | USC | 6.184516 |
| 10 | HSB118 | A1C | 4Y | 1726 | 7 | M | R | 6.52 | African | USC | 5.818379 |
| 10 | HSB118 | STC | 4Y | 1726 | 7 | M | R | 6.52 | African | USC | 5.516166 |

| | | | | | | | | | | | |
|---|---|---|---|---|---|---|---|---|---|---|---|
| 10 | HSB118 | ITC | 4Y | 1726 | 7 | M | R | 6.52 | African | USC | 6.425146 |
| 10 | HSB118 | V1C | 4Y | 1726 | 7 | M | R | 6.52 | African | USC | 6.943632 |
| 10 | HSB118 | HIP | 4Y | 1726 | 7 | M | R | 6.52 | African | USC | 5.06958 |
| 10 | HSB118 | AMY | 4Y | 1726 | 7 | M | R | 6.52 | African | USC | 5.396023 |
| 10 | HSB118 | STR | 4Y | 1726 | 7 | M | R | 6.52 | African | USC | 2.060486 |
| 10 | HSB118 | MD | 4Y | 1726 | 7 | M | R | 6.52 | African | USC | 4.018741 |
| 10 | HSB118 | CBC | 4Y | 1726 | 7 | M | R | 6.52 | African | USC | 4.222756 |
| 11 | HSB141 | OFC | 8Y | 3186 | 7 | M | L | 6.54 | African | YALE | 4.646533 |
| 11 | HSB141 | DFC | 8Y | 3186 | 7 | M | L | 6.54 | African | YALE | 4.884902 |
| 11 | HSB141 | VFC | 8Y | 3186 | 7 | M | L | 6.54 | African | YALE | 4.411362 |
| 11 | HSB141 | MFC | 8Y | 3186 | 7 | M | L | 6.54 | African | YALE | 4.727629 |
| 11 | HSB141 | M1C | 8Y | 3186 | 7 | M | L | 6.54 | African | YALE | 4.822153 |
| 11 | HSB141 | S1C | 8Y | 3186 | 7 | M | L | 6.54 | African | YALE | 4.614405 |
| 11 | HSB141 | IPC | 8Y | 3186 | 7 | M | L | 6.54 | African | YALE | 4.682118 |
| 11 | HSB141 | A1C | 8Y | 3186 | 7 | M | L | 6.54 | African | YALE | 3.64756 |
| 11 | HSB141 | STC | 8Y | 3186 | 7 | M | L | 6.54 | African | YALE | 4.039016 |
| 11 | HSB141 | ITC | 8Y | 3186 | 7 | M | L | 6.54 | African | YALE | 4.861479 |
| 11 | HSB141 | V1C | 8Y | 3186 | 7 | M | L | 6.54 | African | YALE | 3.941058 |
| 11 | HSB141 | HIP | 8Y | 3186 | 7 | M | L | 6.54 | African | YALE | 5.846468 |
| 11 | HSB141 | AMY | 8Y | 3186 | 7 | M | L | 6.54 | African | YALE | 4.416985 |
| 11 | HSB141 | STR | 8Y | 3186 | 7 | M | L | 6.54 | African | YALE | 2.414302 |
| 11 | HSB141 | MD | 8Y | 3186 | 7 | M | L | 6.54 | African | YALE | 4.319251 |
| 11 | HSB141 | CBC | 8Y | 3186 | 7 | M | L | 6.54 | African | YALE | 5.027036 |
| 11 | HSB174 | OFC | 8Y | 3186 | 7 | M | L | 6.25 | African | USC | 6.070479 |
| 11 | HSB174 | DFC | 8Y | 3186 | 7 | M | L | 6.25 | African | USC | 3.813523 |
| 11 | HSB174 | VFC | 8Y | 3186 | 7 | M | L | 6.25 | African | USC | 3.637306 |
| 11 | HSB174 | MFC | 8Y | 3186 | 7 | M | L | 6.25 | African | USC | 3.925076 |
| 11 | HSB174 | IPC | 8Y | 3186 | 7 | M | L | 6.25 | African | USC | 3.725046 |
| 11 | HSB174 | A1C | 8Y | 3186 | 7 | M | L | 6.25 | African | USC | 4.327021 |
| 11 | HSB174 | STC | 8Y | 3186 | 7 | M | L | 6.25 | African | USC | 4.283345 |
| 11 | HSB174 | ITC | 8Y | 3186 | 7 | M | L | 6.25 | African | USC | 4.506992 |
| 11 | HSB174 | V1C | 8Y | 3186 | 7 | M | L | 6.25 | African | USC | 6.297588 |
| 11 | HSB174 | HIP | 8Y | 3186 | 7 | M | L | 6.25 | African | USC | 4.747031 |
| 11 | HSB174 | AMY | 8Y | 3186 | 7 | M | L | 6.25 | African | USC | 4.853335 |
| 11 | HSB174 | CBC | 8Y | 3186 | 7 | M | L | 6.25 | African | USC | 5.812737 |
| 11 | HSB175 | OFC | 11Y | 4281 | 7 | F | L | 6.7 | African | YALE | 4.53389 |
| 11 | HSB175 | DFC | 11Y | 4281 | 7 | F | L | 6.7 | African | YALE | 5.300704 |
| 11 | HSB175 | VFC | 11Y | 4281 | 7 | F | L | 6.7 | African | YALE | 5.001644 |
| 11 | HSB175 | MFC | 11Y | 4281 | 7 | F | L | 6.7 | African | YALE | 5.213292 |
| 11 | HSB175 | M1C | 11Y | 4281 | 7 | F | L | 6.7 | African | YALE | 4.224362 |
| 11 | HSB175 | S1C | 11Y | 4281 | 7 | F | L | 6.7 | African | YALE | 5.082134 |
| 11 | HSB175 | IPC | 11Y | 4281 | 7 | F | L | 6.7 | African | YALE | 4.663143 |
| 11 | HSB175 | A1C | 11Y | 4281 | 7 | F | L | 6.7 | African | YALE | 5.457867 |
| 11 | HSB175 | STC | 11Y | 4281 | 7 | F | L | 6.7 | African | YALE | 4.483474 |
| 11 | HSB175 | ITC | 11Y | 4281 | 7 | F | L | 6.7 | African | YALE | 5.361197 |
| 11 | HSB175 | V1C | 11Y | 4281 | 7 | F | L | 6.7 | African | YALE | 4.597975 |
| 11 | HSB175 | HIP | 11Y | 4281 | 7 | F | L | 6.7 | African | YALE | 5.126031 |
| 11 | HSB175 | AMY | 11Y | 4281 | 7 | F | L | 6.7 | African | YALE | 5.640279 |
| 11 | HSB175 | CBC | 11Y | 4281 | 7 | F | L | 6.7 | African | YALE | 5.440078 |
| 12 | HSB124 | OFC | 13Y | 5011 | 8 | F | R | 6.34 | African | YALE | 4.970213 |
| 12 | HSB124 | DFC | 13Y | 5011 | 8 | F | R | 6.34 | African | YALE | 5.037169 |

| | | | | | | | | | | | |
|---|---|---|---|---|---|---|---|---|---|---|---|
| 12 | HSB124 | VFC | 13Y | 5011 | 8 | F | R | 6.34 | African | YALE | 4.477614 |
| 12 | HSB124 | MFC | 13Y | 5011 | 8 | F | R | 6.34 | African | YALE | 4.745781 |
| 12 | HSB124 | M1C | 13Y | 5011 | 8 | F | R | 6.34 | African | YALE | 5.290018 |
| 12 | HSB124 | S1C | 13Y | 5011 | 8 | F | R | 6.34 | African | YALE | 5.307395 |
| 12 | HSB124 | IPC | 13Y | 5011 | 8 | F | R | 6.34 | African | YALE | 4.672492 |
| 12 | HSB124 | A1C | 13Y | 5011 | 8 | F | R | 6.34 | African | YALE | 4.213409 |
| 12 | HSB124 | STC | 13Y | 5011 | 8 | F | R | 6.34 | African | YALE | 5.26332 |
| 12 | HSB124 | ITC | 13Y | 5011 | 8 | F | R | 6.34 | African | YALE | 4.721562 |
| 12 | HSB124 | V1C | 13Y | 5011 | 8 | F | R | 6.34 | African | YALE | 3.437448 |
| 12 | HSB124 | HIP | 13Y | 5011 | 8 | F | R | 6.34 | African | YALE | 4.828312 |
| 12 | HSB124 | AMY | 13Y | 5011 | 8 | F | R | 6.34 | African | YALE | 4.2829 |
| 12 | HSB124 | STR | 13Y | 5011 | 8 | F | R | 6.34 | African | YALE | 2.290331 |
| 12 | HSB124 | MD | 13Y | 5011 | 8 | F | R | 6.34 | African | YALE | 3.172126 |
| 12 | HSB124 | CBC | 13Y | 5011 | 8 | F | R | 6.34 | African | YALE | 5.501997 |
| 12 | HSB119 | OFC | 15Y | 5741 | 8 | M | L | 6.93 | African | USC | 5.228314 |
| 12 | HSB119 | DFC | 15Y | 5741 | 8 | M | L | 6.93 | African | USC | 5.46712 |
| 12 | HSB119 | VFC | 15Y | 5741 | 8 | M | L | 6.93 | African | USC | 5.372705 |
| 12 | HSB119 | MFC | 15Y | 5741 | 8 | M | L | 6.93 | African | USC | 4.590966 |
| 12 | HSB119 | M1C | 15Y | 5741 | 8 | M | L | 6.93 | African | USC | 4.536367 |
| 12 | HSB119 | S1C | 15Y | 5741 | 8 | M | L | 6.93 | African | USC | 4.720857 |
| 12 | HSB119 | IPC | 15Y | 5741 | 8 | M | L | 6.93 | African | USC | 5.027184 |
| 12 | HSB119 | A1C | 15Y | 5741 | 8 | M | L | 6.93 | African | USC | 4.814781 |
| 12 | HSB119 | STC | 15Y | 5741 | 8 | M | L | 6.93 | African | USC | 5.079346 |
| 12 | HSB119 | ITC | 15Y | 5741 | 8 | M | L | 6.93 | African | USC | 5.220743 |
| 12 | HSB119 | V1C | 15Y | 5741 | 8 | M | L | 6.93 | African | USC | 5.130977 |
| 12 | HSB119 | AMY | 15Y | 5741 | 8 | M | L | 6.93 | African | USC | 4.598707 |
| 12 | HSB119 | STR | 15Y | 5741 | 8 | M | L | 6.93 | African | USC | 1.908759 |
| 12 | HSB119 | MD | 15Y | 5741 | 8 | M | L | 6.93 | African | USC | 4.804353 |
| 12 | HSB119 | CBC | 15Y | 5741 | 8 | M | L | 6.93 | African | USC | 5.953129 |
| 12 | HSB105 | OFC | 18Y | 6836 | 8 | M | L | 6.21 | European | USC | 4.922878 |
| 12 | HSB105 | DFC | 18Y | 6836 | 8 | M | L | 6.21 | European | USC | 5.275066 |
| 12 | HSB105 | VFC | 18Y | 6836 | 8 | M | L | 6.21 | European | USC | 5.309739 |
| 12 | HSB105 | MFC | 18Y | 6836 | 8 | M | L | 6.21 | European | USC | 4.799758 |
| 12 | HSB105 | M1C | 18Y | 6836 | 8 | M | L | 6.21 | European | USC | 3.867083 |
| 12 | HSB105 | S1C | 18Y | 6836 | 8 | M | L | 6.21 | European | USC | 3.19863 |
| 12 | HSB105 | IPC | 18Y | 6836 | 8 | M | L | 6.21 | European | USC | 5.881258 |
| 12 | HSB105 | A1C | 18Y | 6836 | 8 | M | L | 6.21 | European | USC | 4.036356 |
| 12 | HSB105 | STC | 18Y | 6836 | 8 | M | L | 6.21 | European | USC | 4.658847 |
| 12 | HSB105 | ITC | 18Y | 6836 | 8 | M | L | 6.21 | European | USC | 4.783648 |
| 12 | HSB105 | V1C | 18Y | 6836 | 8 | M | L | 6.21 | European | USC | 5.268818 |
| 12 | HSB105 | HIP | 18Y | 6836 | 8 | M | L | 6.21 | European | USC | 4.782492 |
| 12 | HSB105 | CBC | 18Y | 6836 | 8 | M | L | 6.21 | European | USC | 4.796316 |
| 12 | HSB127 | OFC | 19Y | 7201 | 8 | F | L | 5.91 | European | YALE | 4.892355 |
| 12 | HSB127 | DFC | 19Y | 7201 | 8 | F | L | 5.91 | European | YALE | 5.146623 |
| 12 | HSB127 | VFC | 19Y | 7201 | 8 | F | L | 5.91 | European | YALE | 4.739779 |
| 12 | HSB127 | MFC | 19Y | 7201 | 8 | F | L | 5.91 | European | YALE | 5.660126 |
| 12 | HSB127 | M1C | 19Y | 7201 | 8 | F | L | 5.91 | European | YALE | 4.010697 |
| 12 | HSB127 | S1C | 19Y | 7201 | 8 | F | L | 5.91 | European | YALE | 4.37193 |
| 12 | HSB127 | IPC | 19Y | 7201 | 8 | F | L | 5.91 | European | YALE | 5.089881 |
| 12 | HSB127 | A1C | 19Y | 7201 | 8 | F | L | 5.91 | European | YALE | 4.132435 |
| 12 | HSB127 | STC | 19Y | 7201 | 8 | F | L | 5.91 | European | YALE | 4.903591 |

| | | | | | | | | | | | |
|---|---|---|---|---|---|---|---|---|---|---|---|
| 12 | HSB127 | ITC | 19Y | 7201 | 8 | F | L | 5.91 | European | YALE | 4.10677 |
| 12 | HSB127 | V1C | 19Y | 7201 | 8 | F | L | 5.91 | European | YALE | 5.100166 |
| 12 | HSB127 | HIP | 19Y | 7201 | 8 | F | R | 5.91 | European | YALE | 5.123428 |
| 12 | HSB127 | AMY | 19Y | 7201 | 8 | F | R | 5.91 | European | YALE | 4.626231 |
| 12 | HSB127 | STR | 19Y | 7201 | 8 | F | L | 5.91 | European | YALE | 2.183946 |
| 12 | HSB127 | MD | 19Y | 7201 | 8 | F | L | 5.91 | European | YALE | 4.233124 |
| 12 | HSB127 | CBC | 19Y | 7201 | 8 | F | L | 5.91 | European | YALE | 5.369406 |
| 13 | HSB130 | OFC | 21Y | 7931 | 9 | F | L | 6.81 | European | YALE | 5.968639 |
| 13 | HSB130 | DFC | 21Y | 7931 | 9 | F | L | 6.81 | European | YALE | 5.670118 |
| 13 | HSB130 | VFC | 21Y | 7931 | 9 | F | L | 6.81 | European | YALE | 5.36531 |
| 13 | HSB130 | MFC | 21Y | 7931 | 9 | F | L | 6.81 | European | YALE | 5.171412 |
| 13 | HSB130 | M1C | 21Y | 7931 | 9 | F | L | 6.81 | European | YALE | 5.661556 |
| 13 | HSB130 | S1C | 21Y | 7931 | 9 | F | L | 6.81 | European | YALE | 5.514421 |
| 13 | HSB130 | IPC | 21Y | 7931 | 9 | F | L | 6.81 | European | YALE | 4.93984 |
| 13 | HSB130 | A1C | 21Y | 7931 | 9 | F | L | 6.81 | European | YALE | 5.611973 |
| 13 | HSB130 | STC | 21Y | 7931 | 9 | F | L | 6.81 | European | YALE | 5.035931 |
| 13 | HSB130 | ITC | 21Y | 7931 | 9 | F | L | 6.81 | European | YALE | 5.704396 |
| 13 | HSB130 | V1C | 21Y | 7931 | 9 | F | L | 6.81 | European | YALE | 5.315433 |
| 13 | HSB130 | HIP | 21Y | 7931 | 9 | F | L | 6.81 | European | YALE | 5.479896 |
| 13 | HSB130 | AMY | 21Y | 7931 | 9 | F | L | 6.81 | European | YALE | 4.607365 |
| 13 | HSB130 | STR | 21Y | 7931 | 9 | F | L | 6.81 | European | YALE | 1.773284 |
| 13 | HSB130 | MD | 21Y | 7931 | 9 | F | L | 6.81 | European | YALE | 4.409037 |
| 13 | HSB130 | CBC | 21Y | 7931 | 9 | F | L | 6.81 | European | YALE | 6.402651 |
| 13 | HSB136 | OFC | 23Y | 8661 | 9 | M | R | 6.36 | African | USC | 4.008988 |
| 13 | HSB136 | DFC | 23Y | 8661 | 9 | M | R | 6.36 | African | USC | 4.820583 |
| 13 | HSB136 | VFC | 23Y | 8661 | 9 | M | R | 6.36 | African | USC | 4.181974 |
| 13 | HSB136 | MFC | 23Y | 8661 | 9 | M | R | 6.36 | African | USC | 4.304169 |
| 13 | HSB136 | M1C | 23Y | 8661 | 9 | M | R | 6.36 | African | USC | 4.650999 |
| 13 | HSB136 | S1C | 23Y | 8661 | 9 | M | R | 6.36 | African | USC | 4.149765 |
| 13 | HSB136 | IPC | 23Y | 8661 | 9 | M | R | 6.36 | African | USC | 4.390121 |
| 13 | HSB136 | A1C | 23Y | 8661 | 9 | M | R | 6.36 | African | USC | 3.875727 |
| 13 | HSB136 | STC | 23Y | 8661 | 9 | M | R | 6.36 | African | USC | 3.644584 |
| 13 | HSB136 | ITC | 23Y | 8661 | 9 | M | R | 6.36 | African | USC | 4.384766 |
| 13 | HSB136 | V1C | 23Y | 8661 | 9 | M | R | 6.36 | African | USC | 3.307311 |
| 13 | HSB136 | HIP | 23Y | 8661 | 9 | M | R | 6.36 | African | USC | 3.894527 |
| 13 | HSB136 | AMY | 23Y | 8661 | 9 | M | R | 6.36 | African | USC | 3.926063 |
| 13 | HSB136 | STR | 23Y | 8661 | 9 | M | R | 6.36 | African | USC | 1.912328 |
| 13 | HSB136 | MD | 23Y | 8661 | 9 | M | R | 6.36 | African | USC | 2.720664 |
| 13 | HSB136 | CBC | 23Y | 8661 | 9 | M | R | 6.36 | African | USC | 5.410375 |
| 13 | HSB126 | OFC | 30Y | 11216 | 9 | F | R | 6.92 | European | YALE | 4.928515 |
| 13 | HSB126 | DFC | 30Y | 11216 | 9 | F | R | 6.92 | European | YALE | 4.759084 |
| 13 | HSB126 | VFC | 30Y | 11216 | 9 | F | R | 6.92 | European | YALE | 3.853939 |
| 13 | HSB126 | MFC | 30Y | 11216 | 9 | F | R | 6.92 | European | YALE | 5.238299 |
| 13 | HSB126 | M1C | 30Y | 11216 | 9 | F | R | 6.92 | European | YALE | 4.976462 |
| 13 | HSB126 | S1C | 30Y | 11216 | 9 | F | R | 6.92 | European | YALE | 4.878776 |
| 13 | HSB126 | IPC | 30Y | 11216 | 9 | F | R | 6.92 | European | YALE | 4.984017 |
| 13 | HSB126 | A1C | 30Y | 11216 | 9 | F | R | 6.92 | European | YALE | 3.696279 |
| 13 | HSB126 | STC | 30Y | 11216 | 9 | F | R | 6.92 | European | YALE | 5.193616 |
| 13 | HSB126 | ITC | 30Y | 11216 | 9 | F | R | 6.92 | European | YALE | 4.850426 |
| 13 | HSB126 | V1C | 30Y | 11216 | 9 | F | R | 6.92 | European | YALE | 4.613238 |
| 13 | HSB126 | HIP | 30Y | 11216 | 9 | F | R | 6.92 | European | YALE | 4.253025 |

| | | | | | | | | | | | |
|---|---|---|---|---|---|---|---|---|---|---|---|
| 13 | HSB126 | AMY | 30Y | 11216 | 9 | F | R | 6.92 | European | YALE | 4.682118 |
| 13 | HSB126 | STR | 30Y | 11216 | 9 | F | R | 6.92 | European | YALE | 1.904782 |
| 13 | HSB126 | MD | 30Y | 11216 | 9 | F | R | 6.92 | European | YALE | 4.342721 |
| 13 | HSB126 | CBC | 30Y | 11216 | 9 | F | R | 6.92 | European | YALE | 6.637674 |
| 13 | HSB145 | OFC | 36Y | 13406 | 9 | M | R | NA | European | YALE | 5.735972 |
| 13 | HSB145 | DFC | 36Y | 13406 | 9 | M | R | NA | European | YALE | 6.176367 |
| 13 | HSB145 | VFC | 36Y | 13406 | 9 | M | R | NA | European | YALE | 5.756201 |
| 13 | HSB145 | MFC | 36Y | 13406 | 9 | M | R | NA | European | YALE | 6.556032 |
| 13 | HSB145 | M1C | 36Y | 13406 | 9 | M | R | NA | European | YALE | 5.871608 |
| 13 | HSB145 | S1C | 36Y | 13406 | 9 | M | R | NA | European | YALE | 5.931124 |
| 13 | HSB145 | IPC | 36Y | 13406 | 9 | M | R | NA | European | YALE | 6.422097 |
| 13 | HSB145 | A1C | 36Y | 13406 | 9 | M | R | NA | European | YALE | 5.510272 |
| 13 | HSB145 | STC | 36Y | 13406 | 9 | M | R | NA | European | YALE | 5.114432 |
| 13 | HSB145 | ITC | 36Y | 13406 | 9 | M | R | NA | European | YALE | 4.838004 |
| 13 | HSB145 | V1C | 36Y | 13406 | 9 | M | R | NA | European | YALE | 5.492304 |
| 13 | HSB145 | HIP | 36Y | 13406 | 9 | M | R | NA | European | YALE | 4.695329 |
| 13 | HSB145 | AMY | 36Y | 13406 | 9 | M | R | NA | European | YALE | 5.171412 |
| 13 | HSB145 | STR | 36Y | 13406 | 9 | M | R | NA | European | YALE | 2.374714 |
| 13 | HSB145 | MD | 36Y | 13406 | 9 | M | R | NA | European | YALE | 3.97728 |
| 13 | HSB145 | CBC | 36Y | 13406 | 9 | M | R | NA | European | YALE | 5.381481 |
| 13 | HSB123 | OFC | 37Y | 13771 | 9 | M | R | 6.37 | African | YALE | 4.672492 |
| 13 | HSB123 | DFC | 37Y | 13771 | 9 | M | R | 6.37 | African | YALE | 4.617935 |
| 13 | HSB123 | VFC | 37Y | 13771 | 9 | M | R | 6.37 | African | YALE | 5.091138 |
| 13 | HSB123 | MFC | 37Y | 13771 | 9 | M | R | 6.37 | African | YALE | 6.703634 |
| 13 | HSB123 | M1C | 37Y | 13771 | 9 | M | R | 6.37 | African | YALE | 5.518592 |
| 13 | HSB123 | S1C | 37Y | 13771 | 9 | M | R | 6.37 | African | YALE | 5.83177 |
| 13 | HSB123 | IPC | 37Y | 13771 | 9 | M | R | 6.37 | African | YALE | 4.966378 |
| 13 | HSB123 | A1C | 37Y | 13771 | 9 | M | R | 6.37 | African | YALE | 4.973938 |
| 13 | HSB123 | STC | 37Y | 13771 | 9 | M | R | 6.37 | African | YALE | 4.763998 |
| 13 | HSB123 | ITC | 37Y | 13771 | 9 | M | R | 6.37 | African | YALE | 5.574039 |
| 13 | HSB123 | V1C | 37Y | 13771 | 9 | M | R | 6.37 | African | YALE | 4.710795 |
| 13 | HSB123 | HIP | 37Y | 13771 | 9 | M | R | 6.37 | African | YALE | 4.791771 |
| 13 | HSB123 | AMY | 37Y | 13771 | 9 | M | R | 6.37 | African | YALE | 5.180601 |
| 13 | HSB123 | STR | 37Y | 13771 | 9 | M | R | 6.37 | African | YALE | 2.198619 |
| 13 | HSB123 | MD | 37Y | 13771 | 9 | M | R | 6.37 | African | YALE | 4.556999 |
| 13 | HSB123 | CBC | 37Y | 13771 | 9 | M | R | 6.37 | African | YALE | 4.309228 |
| 13 | HSB135 | OFC | 40Y | 14866 | 9 | F | R | 6.82 | African | USC | 5.028387 |
| 13 | HSB135 | DFC | 40Y | 14866 | 9 | F | R | 6.82 | African | USC | 4.099561 |
| 13 | HSB135 | VFC | 40Y | 14866 | 9 | F | R | 6.82 | African | USC | 5.033161 |
| 13 | HSB135 | M1C | 40Y | 14866 | 9 | F | R | 6.82 | African | USC | 4.751593 |
| 13 | HSB135 | S1C | 40Y | 14866 | 9 | F | R | 6.82 | African | USC | 5.253475 |
| 13 | HSB135 | IPC | 40Y | 14866 | 9 | F | R | 6.82 | African | USC | 5.188229 |
| 13 | HSB135 | A1C | 40Y | 14866 | 9 | F | R | 6.82 | African | USC | 5.468498 |
| 13 | HSB135 | STC | 40Y | 14866 | 9 | F | R | 6.82 | African | USC | 4.944258 |
| 13 | HSB135 | ITC | 40Y | 14866 | 9 | F | R | 6.82 | African | USC | 4.719734 |
| 13 | HSB135 | V1C | 40Y | 14866 | 9 | F | R | 6.82 | African | USC | 5.619968 |
| 13 | HSB135 | HIP | 40Y | 14866 | 9 | F | R | 6.82 | African | USC | 6.057431 |
| 13 | HSB135 | AMY | 40Y | 14866 | 9 | F | R | 6.82 | African | USC | 3.360649 |
| 13 | HSB135 | STR | 40Y | 14866 | 9 | F | R | 6.82 | African | USC | 2.082202 |
| 13 | HSB135 | MD | 40Y | 14866 | 9 | F | R | 6.82 | African | USC | 4.011913 |
| 13 | HSB135 | CBC | 40Y | 14866 | 9 | F | R | 6.82 | African | USC | 4.59759 |

**Table E5A. Over- or under-represented Gene Ontology biological process terms for the genes positively coexpressed with *DAAM1* across cortical regions in prenatal brains.** The gene list analysis was carried out using the analysis tool in the PANTHER classification system (PMID: 23868073), with GO annotation data (PMID: 27899567). The GO terms were organized *hierarchically*, where the most post specific terms, indicated by bolded texts, were listed before the more general terms. The background gene set contains all 21,042 protein-coding genes available in PANTHER database (May 21, 2018 version). The test set contains the 339 genes whose expression levels were positively correlated with DAAM1 expression in all cortical regions in the prenatal brains. We used r ≥0.56 as the cut-off for positive-correlation coefficient, which was taken to be the 99th percentile of the empirical distribution of the correlation coefficients between DAAM1 expression levels and the expression levels of all the other 60,153 coding and non-coding genes in neo-cortical regions of prenatal brains available in the BrainSpan database (http://www.brainspan.org/). The Pearson correlation coefficients were obtained using the vector of mRNA expression levels of each gene across all prenatal stages. For the regions that were not well parcellated during 8 – 10 post-conception weeks, their data were either merged (M1C and S1C) or used to represent the expression values in sub-regions (parietal cortex [PC] and temporal cortex [TC]).

| GO BP complete | N REF (21,042) | N Set (589) | expected | Fold Enrichment | over(+) or under(-) represenation | raw P | FDR |
|---|---|---|---|---|---|---|---|
| **membrane fission** | **14** | **4** | **0.23** | **17.73** | **+** | **1.59E-04** | **3.15E-02** |
| cellular component organization | 5584 | 135 | 89.96 | 1.5 | + | 1.64E-07 | 2.56E-04 |
| cellular process | 15478 | 280 | 249.36 | 1.12 | + | 1.10E-04 | 2.54E-02 |
| cellular component organization or biogenesis | 5773 | 138 | 93.01 | 1.48 | + | 2.21E-07 | 2.30E-04 |
| **clathrin-dependent endocytosis** | **25** | **6** | **0.4** | **14.9** | **+** | **8.01E-06** | **3.48E-03** |
| localization | 5608 | 123 | 90.35 | 1.36 | + | 1.20E-04 | 2.61E-02 |
| **positive regulation of dendritic spine morphogenesis** | **17** | **4** | **0.27** | **14.6** | **+** | **3.00E-04** | **4.51E-02** |
| regulation of dendritic spine morphogenesis | 37 | 7 | 0.6 | 11.74 | + | 5.48E-06 | 2.52E-03 |
| regulation of dendrite morphogenesis | 86 | 9 | 1.39 | 6.5 | + | 2.02E-05 | 7.35E-03 |
| regulation of cell morphogenesis involved in differentiation | 292 | 18 | 4.7 | 3.83 | + | 2.56E-06 | 1.60E-03 |
| regulation of cell morphogenesis | 488 | 25 | 7.86 | 3.18 | + | 7.74E-07 | 6.05E-04 |
| regulation of cellular component organization | 2434 | 73 | 39.21 | 1.86 | + | 2.26E-07 | 2.21E-04 |
| biological regulation | 12409 | 234 | 199.92 | 1.17 | + | 1.82E-04 | 3.52E-02 |
| regulation of cell development | 881 | 37 | 14.19 | 2.61 | + | 2.06E-07 | 2.30E-04 |
| regulation of dendrite development | 140 | 14 | 2.26 | 6.21 | + | 1.73E-07 | 2.26E-04 |
| regulation of neuron projection development | 466 | 31 | 7.51 | 4.13 | + | 9.76E-11 | 1.53E-06 |
| regulation of neuron differentiation | 623 | 34 | 10.04 | 3.39 | + | 1.60E-09 | 6.25E-06 |
| regulation of neurogenesis | 765 | 34 | 12.32 | 2.76 | + | 1.91E-07 | 2.30E-04 |
| regulation of nervous system development | 853 | 35 | 13.74 | 2.55 | + | 7.55E-07 | 6.22E-04 |
| generation of neurons | 1480 | 54 | 23.84 | 2.26 | + | 2.91E-08 | 5.69E-05 |
| neurogenesis | 1577 | 54 | 25.41 | 2.13 | + | 2.43E-07 | 2.23E-04 |
| nervous system development | 2298 | 76 | 37.02 | 2.05 | + | 1.64E-09 | 5.14E-06 |
| regulation of plasma membrane bounded cell projection organization | 620 | 35 | 9.99 | 3.5 | + | 3.88E-10 | 3.03E-06 |
| regulation of cell projection organization | 629 | 35 | 10.13 | 3.45 | + | 5.57E-10 | 2.90E-06 |
| regulation of postsynapse organization | 58 | 7 | 0.93 | 7.49 | + | 7.49E-05 | 1.92E-02 |
| regulation of dendritic spine development | 68 | 9 | 1.1 | 8.22 | + | 3.56E-06 | 2.06E-03 |
| positive regulation of dendritic spine development | 40 | 6 | 0.64 | 9.31 | + | 8.34E-05 | 2.07E-02 |
| positive regulation of dendrite development | 69 | 7 | 1.11 | 6.3 | + | 2.03E-04 | 3.77E-02 |
| positive regulation of neuron projection development | 262 | 20 | 4.22 | 4.74 | + | 2.80E-08 | 6.25E-05 |
| positive regulation of neuron differentiation | 348 | 22 | 5.61 | 3.92 | + | 1.30E-07 | 2.26E-04 |
| positive regulation of neurogenesis | 438 | 22 | 7.06 | 3.12 | + | 4.87E-06 | 2.46E-03 |
| positive regulation of cell development | 507 | 26 | 8.17 | 3.18 | + | 4.54E-07 | 3.95E-04 |
| positive regulation of nervous system development | 501 | 23 | 8.07 | 2.85 | + | 1.20E-05 | 4.69E-03 |
| positive regulation of cell projection organization | 354 | 24 | 5.7 | 4.21 | + | 9.80E-09 | 2.55E-05 |
| positive regulation of cellular component organization | 1179 | 45 | 18.99 | 2.37 | + | 1.68E-07 | 2.39E-04 |
| positive regulation of cell morphogenesis involved in differentiation | 144 | 11 | 2.32 | 4.74 | + | 3.84E-05 | 1.18E-02 |
| **negative regulation of microtubule depolymerization** | **26** | **5** | **0.42** | **11.94** | **+** | **1.18E-04** | **2.59E-02** |
| negative regulation of cellular component organization | 666 | 30 | 10.73 | 2.8 | + | 7.99E-07 | 5.95E-04 |
| negative regulation of microtubule polymerization or depolymerization | 38 | 6 | 0.61 | 9.8 | + | 6.46E-05 | 1.71E-02 |
| negative regulation of organelle organization | 356 | 17 | 5.74 | 2.96 | + | 1.05E-04 | 2.50E-02 |
| regulation of cytoskeleton organization | 526 | 23 | 8.47 | 2.71 | + | 2.50E-05 | 8.33E-03 |
| regulation of microtubule polymerization or depolymerization | 79 | 8 | 1.27 | 6.29 | + | 7.24E-05 | 1.89E-02 |
| regulation of microtubule cytoskeleton organization | 186 | 13 | 3 | 4.34 | + | 1.87E-05 | 7.12E-03 |
| regulation of microtubule-based process | 216 | 15 | 3.48 | 4.31 | + | 4.61E-06 | 2.40E-03 |
| negative regulation of protein complex disassembly | 69 | 7 | 1.11 | 6.3 | + | 2.03E-04 | 3.82E-02 |
| regulation of protein complex disassembly | 109 | 9 | 1.76 | 5.13 | + | 1.12E-04 | 2.54E-02 |
| regulation of microtubule depolymerization | 30 | 6 | 0.48 | 12.41 | + | 1.98E-05 | 7.38E-03 |
| **cerebral cortex radially oriented cell migration** | **30** | **5** | **0.48** | **10.35** | **+** | **2.14E-04** | **3.89E-02** |
| movement of cell or subcellular component | 1510 | 44 | 24.33 | 1.81 | + | 1.87E-04 | 3.57E-02 |
| **regulation of microtubule polymerization** | **48** | **7** | **0.77** | **9.05** | **+** | **2.50E-05** | **8.50E-03** |
| regulation of protein polymerization | 223 | 13 | 3.59 | 3.62 | + | 1.08E-04 | 2.53E-02 |
| **axonal transport** | **48** | **6** | **0.77** | **7.76** | **+** | **2.07E-04** | **3.80E-02** |
| transport along microtubule | 140 | 10 | 2.26 | 4.43 | + | 1.45E-04 | 2.98E-02 |
| microtubule-based transport | 140 | 10 | 2.26 | 4.43 | + | 1.45E-04 | 3.02E-02 |
| microtubule-based process | 698 | 29 | 11.25 | 2.58 | + | 5.60E-06 | 2.50E-03 |
| cytoskeleton-dependent intracellular transport | 161 | 12 | 2.59 | 4.63 | + | 2.16E-05 | 7.67E-03 |
| establishment of localization in cell | 1779 | 49 | 28.66 | 1.71 | + | 2.63E-04 | 4.20E-02 |
| **vesicle cytoskeletal trafficking** | **50** | **6** | **0.81** | **7.45** | **+** | **2.53E-04** | **4.17E-02** |
| **organelle transport along microtubule** | **71** | **7** | **1.14** | **6.12** | **+** | **2.38E-04** | **4.10E-02** |
| **dendrite development** | **99** | **9** | **1.59** | **5.64** | **+** | **5.62E-05** | **1.54E-02** |
| neuron projection development | 653 | 26 | 10.52 | 2.47 | + | 5.08E-05 | 1.45E-02 |
| plasma membrane bounded cell projection organization | 1097 | 40 | 17.67 | 2.26 | + | 2.22E-06 | 1.58E-03 |
| cell projection organization | 1134 | 41 | 18.27 | 2.24 | + | 2.77E-06 | 1.67E-03 |
| neuron development | 798 | 32 | 12.86 | 2.49 | + | 4.99E-06 | 2.44E-03 |
| neuron differentiation | 988 | 35 | 15.92 | 2.2 | + | 2.58E-05 | 8.24E-03 |

| GO BP complete | N REF (21,042) | N Set (589) | expected | Fold Enrichment | over(+) or under(-) represenation | raw P | FDR |
|---|---|---|---|---|---|---|---|
| **neuron migration** | **118** | **10** | **1.9** | **5.26** | **+** | **3.82E-05** | **1.19E-02** |
| **regulation of axon extension** | **96** | **8** | **1.55** | **5.17** | **+** | **2.53E-04** | **4.21E-02** |
| **negative regulation of neuron projection development** | **148** | **12** | **2.38** | **5.03** | **+** | **9.83E-06** | **4.15E-03** |
| negative regulation of neuron differentiation | 220 | 13 | 3.54 | 3.67 | + | 9.53E-05 | 2.33E-02 |
| negative regulation of neurogenesis | 274 | 14 | 4.41 | 3.17 | + | 2.21E-04 | 3.97E-02 |
| negative regulation of cell projection organization | 175 | 13 | 2.82 | 4.61 | + | 1.02E-05 | 4.19E-03 |
| **protein-containing complex localization** | **246** | **13** | **3.96** | **3.28** | **+** | **2.70E-04** | **4.27E-02** |
| **axonogenesis** | **358** | **17** | **5.77** | **2.95** | **+** | **1.13E-04** | **2.51E-02** |
| axon development | 389 | 18 | 6.27 | 2.87 | + | 9.72E-05 | 2.34E-02 |
| cell part morphogenesis | 500 | 20 | 8.06 | 2.48 | + | 2.63E-04 | 4.24E-02 |
| cell morphogenesis involved in neuron differentiation | 419 | 19 | 6.75 | 2.81 | + | 8.06E-05 | 2.03E-02 |
| **microtubule cytoskeleton organization** | **486** | **22** | **7.83** | **2.81** | **+** | **2.28E-05** | **7.93E-03** |
| cytoskeleton organization | 1111 | 36 | 17.9 | 2.01 | + | 1.25E-04 | 2.64E-02 |
| organelle organization | 3387 | 89 | 54.57 | 1.63 | + | 2.42E-06 | 1.65E-03 |
| **endomembrane system organization** | **380** | **17** | **6.12** | **2.78** | **+** | **2.21E-04** | **3.93E-02** |
| **chemical synaptic transmission** | **456** | **19** | **7.35** | **2.59** | **+** | **2.29E-04** | **4.02E-02** |
| anterograde trans-synaptic signaling | 456 | 19 | 7.35 | 2.59 | + | 2.29E-04 | 3.98E-02 |
| trans-synaptic signaling | 464 | 19 | 7.48 | 2.54 | + | 2.82E-04 | 4.33E-02 |
| synaptic signaling | 465 | 19 | 7.49 | 2.54 | + | 2.90E-04 | 4.40E-02 |
| **cell projection assembly** | **460** | **19** | **7.41** | **2.56** | **+** | **2.54E-04** | **4.14E-02** |
| cellular component assembly | 2504 | 67 | 40.34 | 1.66 | + | 4.56E-05 | 1.34E-02 |
| cellular component biogenesis | 2728 | 71 | 43.95 | 1.62 | + | 6.08E-05 | 1.64E-02 |
| **regulation of cellular localization** | **802** | **32** | **12.92** | **2.48** | **+** | **5.29E-06** | **2.51E-03** |
| regulation of localization | 2607 | 69 | 42 | 1.64 | + | 4.33E-05 | 1.30E-02 |
| **ubiquitin-dependent protein catabolic process** | **502** | **20** | **8.09** | **2.47** | **+** | **2.76E-04** | **4.32E-02** |
| modification-dependent protein catabolic process | 526 | 23 | 8.47 | 2.71 | + | 2.50E-05 | 8.16E-03 |
| proteolysis involved in cellular protein catabolic process | 584 | 23 | 9.41 | 2.44 | + | 1.68E-04 | 3.28E-02 |
| cellular protein catabolic process | 613 | 24 | 9.88 | 2.43 | + | 1.22E-04 | 2.60E-02 |
| cellular protein metabolic process | 3700 | 90 | 59.61 | 1.51 | + | 5.34E-05 | 1.49E-02 |
| cellular macromolecule metabolic process | 6569 | 138 | 105.83 | 1.3 | + | 2.47E-04 | 4.16E-02 |
| modification-dependent macromolecule catabolic process | 536 | 24 | 8.64 | 2.78 | + | 1.15E-05 | 4.61E-03 |
| **protein ubiquitination** | **654** | **24** | **10.54** | **2.28** | **+** | **2.44E-04** | **4.14E-02** |
| cellular protein modification process | 3065 | 82 | 49.38 | 1.66 | + | 4.16E-06 | 2.33E-03 |
| protein modification process | 3065 | 82 | 49.38 | 1.66 | + | 4.16E-06 | 2.25E-03 |
| macromolecule modification | 3281 | 87 | 52.86 | 1.65 | + | 2.56E-06 | 1.67E-03 |
| **regulation of transport** | **1788** | **52** | **28.81** | **1.81** | **+** | **5.07E-05** | **1.47E-02** |
| **phosphate-containing compound metabolic process** | **2069** | **56** | **33.33** | **1.68** | **+** | **1.53E-04** | **3.07E-02** |
| phosphorus metabolic process | 2165 | 58 | 34.88 | 1.66 | + | 1.45E-04 | 2.95E-02 |
| **immune response** | **1825** | **12** | **29.4** | **0.41** | **-** | **2.78E-04** | **4.31E-02** |
| **detection of stimulus involved in sensory perception** | **531** | **0** | **8.55** | **< 0.01** | **-** | **3.12E-04** | **4.64E-02** |

**Table E5B. Over- or under-represented Gene Ontology biological process terms for the genes negatively coexpressed with DAAM1 across cortical regions in prenatal brains.**

The gene list analysis was carried out using the analysis tool in the PANTHER classification system (PMID: 23868073), with GO annotation data (PMID: 27899567). The GO terms were organized *hierarchically*, where the most post specific terms, indicated by bolded texts, were listed before the more general terms. The background gene set contains all 21,042 protein-coding genes available in PANTHER database (May 21, 2018 version). The test set contains the 583 genes whose expression levels were negatively correlated with DAAM1 expression in all cortical regions in the prenatal brains. We used r≤ -0.55 as the cut-off for the negative-correlation coefficient, which was taken to be the 1st percentile of the empirical distribution of the correlation coefficients between DAAM1 expression levels and the expression levels of all the other 60,153 coding and non-coding genes in neo-cortical regions of prenatal brains available in the BrainSpan database (http://www.brainspan.org/). The Pearson correlation coefficients were obtained using the vector of mRNA expression levels of each gene across all prenatal stages. For the regions that were not well parcellated during 8 – 10 post-conception weeks, their data were either merged (M1C and S1C) or used to represent the expression values in sub-regions (parietal cortex [PC] and temporal cortex [TC]).

| GO BP complete | N REF (21,042) | N Set (589) | expected | Fold Enrichment | over(+) or under(-) represenation | raw P | FDR |
|---|---|---|---|---|---|---|---|
| **macropinocytosis** | **7** | **4** | **0.18** | **21.74** | **+** | **1.22E-04** | **2.41E-02** |
| pinocytosis | 11 | 5 | 0.29 | 17.3 | + | 3.74E-05 | 1.06E-02 |
| transport | 4418 | 156 | 116.11 | 1.34 | + | 7.35E-05 | 1.74E-02 |
| establishment of localization | 4533 | 157 | 119.13 | 1.32 | + | 2.00E-04 | 3.32E-02 |
| vesicle-mediated transport | 1914 | 81 | 50.3 | 1.61 | + | 2.84E-05 | 8.72E-03 |
| **carnitine shuttle** | **8** | **4** | **0.21** | **19.03** | **+** | **1.79E-04** | **3.07E-02** |
| fatty acid transmembrane transport | 9 | 4 | 0.24 | 16.91 | + | 2.53E-04 | 4.00E-02 |
| **glycine metabolic process** | **17** | **6** | **0.45** | **13.43** | **+** | **1.91E-05** | **6.36E-03** |
| drug metabolic process | 669 | 47 | 17.58 | 2.67 | + | 4.04E-09 | 6.32E-06 |
| metabolic process | 9985 | 313 | 262.41 | 1.19 | + | 2.29E-05 | 7.29E-03 |
| serine family amino acid metabolic process | 42 | 9 | 1.1 | 8.15 | + | 5.18E-06 | 2.25E-03 |
| alpha-amino acid metabolic process | 218 | 25 | 5.73 | 4.36 | + | 4.12E-09 | 5.85E-06 |
| cellular amino acid metabolic process | 320 | 31 | 8.41 | 3.69 | + | 2.44E-09 | 4.23E-06 |
| organonitrogen compound metabolic process | 5512 | 190 | 144.86 | 1.31 | + | 3.10E-05 | 9.33E-03 |
| organic substance metabolic process | 9521 | 294 | 250.22 | 1.17 | + | 2.33E-04 | 3.72E-02 |
| carboxylic acid metabolic process | 907 | 68 | 23.84 | 2.85 | + | 5.73E-14 | 2.99E-10 |
| oxoacid metabolic process | 1001 | 72 | 26.31 | 2.74 | + | 6.61E-14 | 2.58E-10 |
| organic acid metabolic process | 1018 | 73 | 26.75 | 2.73 | + | 4.99E-14 | 3.90E-10 |
| small molecule metabolic process | 1836 | 106 | 48.25 | 2.2 | + | 4.79E-14 | 7.50E-10 |
| **serine family amino acid biosynthetic process** | **17** | **5** | **0.45** | **11.19** | **+** | **1.98E-04** | **3.34E-02** |
| alpha-amino acid biosynthetic process | 65 | 9 | 1.71 | 5.27 | + | 1.12E-04 | 2.30E-02 |
| cellular amino acid biosynthetic process | 83 | 10 | 2.18 | 4.58 | + | 1.34E-04 | 2.53E-02 |
| organonitrogen compound biosynthetic process | 1411 | 63 | 37.08 | 1.7 | + | 7.34E-05 | 1.77E-02 |
| carboxylic acid biosynthetic process | 335 | 23 | 8.8 | 2.61 | + | 5.60E-05 | 1.44E-02 |
| organic acid biosynthetic process | 336 | 23 | 8.83 | 2.6 | + | 5.84E-05 | 1.47E-02 |
| small molecule biosynthetic process | 600 | 38 | 15.77 | 2.41 | + | 2.37E-06 | 1.43E-03 |
| **negative regulation of striated muscle cell apoptotic process** | **25** | **6** | **0.66** | **9.13** | **+** | **1.17E-04** | **2.35E-02** |
| negative regulation of muscle cell apoptotic process | 36 | 7 | 0.95 | 7.4 | + | 1.01E-04 | 2.16E-02 |
| **cellular modified amino acid biosynthetic process** | **40** | **7** | **1.05** | **6.66** | **+** | **1.80E-04** | **3.06E-02** |
| **pigment biosynthetic process** | **51** | **8** | **1.34** | **5.97** | **+** | **1.24E-04** | **2.39E-02** |
| pigment metabolic process | 66 | 9 | 1.73 | 5.19 | + | 1.24E-04 | 2.37E-02 |
| **fatty acid beta-oxidation** | **58** | **8** | **1.52** | **5.25** | **+** | **2.74E-04** | **4.16E-02** |
| fatty acid catabolic process | 94 | 11 | 2.47 | 4.45 | + | 7.98E-05 | 1.76E-02 |
| fatty acid metabolic process | 320 | 22 | 8.41 | 2.62 | + | 7.93E-05 | 1.77E-02 |
| monocarboxylic acid metabolic process | 503 | 33 | 13.22 | 2.5 | + | 4.34E-06 | 2.12E-03 |
| lipid metabolic process | 1238 | 57 | 32.54 | 1.75 | + | 8.12E-05 | 1.76E-02 |
| monocarboxylic acid catabolic process | 116 | 12 | 3.05 | 3.94 | + | 1.13E-04 | 2.30E-02 |
| carboxylic acid catabolic process | 258 | 30 | 6.78 | 4.42 | + | 8.39E-11 | 1.64E-07 |
| organic acid catabolic process | 258 | 30 | 6.78 | 4.42 | + | 8.39E-11 | 1.88E-07 |
| cellular catabolic process | 1788 | 76 | 46.99 | 1.62 | + | 5.87E-05 | 1.46E-02 |
| catabolic process | 2016 | 83 | 52.98 | 1.57 | + | 5.89E-05 | 1.44E-02 |
| small molecule catabolic process | 419 | 42 | 11.01 | 3.81 | + | 1.19E-12 | 3.72E-09 |
| organic substance catabolic process | 1726 | 74 | 45.36 | 1.63 | + | 4.60E-05 | 1.28E-02 |
| oxidation-reduction process | 956 | 55 | 25.12 | 2.19 | + | 1.85E-07 | 1.93E-04 |
| **protein targeting to peroxisome** | **68** | **9** | **1.79** | **5.04** | **+** | **1.52E-04** | **2.80E-02** |
| peroxisomal transport | 69 | 9 | 1.81 | 4.96 | + | 1.68E-04 | 2.92E-02 |
| establishment of protein localization to peroxisome | 68 | 9 | 1.79 | 5.04 | + | 1.52E-04 | 2.77E-02 |
| protein localization to peroxisome | 68 | 9 | 1.79 | 5.04 | + | 1.52E-04 | 2.74E-02 |
| peroxisome organization | 82 | 10 | 2.16 | 4.64 | + | 1.23E-04 | 2.40E-02 |
| **drug catabolic process** | **148** | **16** | **3.89** | **4.11** | **+** | **5.16E-06** | **2.30E-03** |
| **regulation of type I interferon production** | **111** | **12** | **2.92** | **4.11** | **+** | **7.70E-05** | **1.77E-02** |
| **alpha-amino acid catabolic process** | **102** | **11** | **2.68** | **4.1** | **+** | **1.55E-04** | **2.72E-02** |
| cellular amino acid catabolic process | 119 | 16 | 3.13 | 5.12 | + | 3.83E-07 | 3.74E-04 |
| **cellular modified amino acid metabolic process** | **183** | **18** | **4.81** | **3.74** | **+** | **4.57E-06** | **2.16E-03** |
| **positive regulation of I-kappaB kinase/NF-kappaB signaling** | **174** | **16** | **4.57** | **3.5** | **+** | **3.28E-05** | **9.50E-03** |
| positive regulation of biological process | 5779 | 206 | 151.88 | 1.36 | + | 8.33E-07 | 6.86E-04 |
| positive regulation of cellular process | 5110 | 186 | 134.29 | 1.39 | + | 1.09E-06 | 8.08E-04 |
| regulation of I-kappaB kinase/NF-kappaB signaling | 218 | 19 | 5.73 | 3.32 | + | 1.24E-05 | 4.52E-03 |
| **protein tetramerization** | **132** | **12** | **3.47** | **3.46** | **+** | **3.42E-04** | **4.95E-02** |
| protein complex oligomerization | 491 | 30 | 12.9 | 2.32 | + | 5.23E-05 | 1.39E-02 |
| **hexose metabolic process** | **159** | **14** | **4.18** | **3.35** | **+** | **1.54E-04** | **2.73E-02** |

| GO BP complete | N REF (21,042) | N Set (589) | expected | Fold Enrichment | over(+) or under(-) represenation | raw P | FDR |
|---|---|---|---|---|---|---|---|
| monosaccharide metabolic process | 202 | 17 | 5.31 | 3.2 | + | 5.29E-05 | 1.38E-02 |
| **coenzyme biosynthetic process** | **173** | **15** | **4.55** | **3.3** | **+** | **1.06E-04** | **2.21E-02** |
| cofactor biosynthetic process | 226 | 23 | 5.94 | 3.87 | + | 1.27E-07 | 1.42E-04 |
| cofactor metabolic process | 496 | 43 | 13.04 | 3.3 | + | 5.02E-11 | 1.31E-07 |
| coenzyme metabolic process | 311 | 26 | 8.17 | 3.18 | + | 6.76E-07 | 5.87E-04 |
| **protein localization to plasma membrane** | **168** | **14** | **4.42** | **3.17** | **+** | **2.61E-04** | **4.03E-02** |
| **positive regulation of cellular catabolic process** | **342** | **26** | **8.99** | **2.89** | **+** | **3.47E-06** | **1.81E-03** |
| positive regulation of catabolic process | 409 | 26 | 10.75 | 2.42 | + | 7.41E-05 | 1.73E-02 |
| regulation of catabolic process | 865 | 43 | 22.73 | 1.89 | + | 1.40E-04 | 2.60E-02 |
| positive regulation of metabolic process | 3339 | 121 | 87.75 | 1.38 | + | 2.60E-04 | 4.06E-02 |
| positive regulation of cellular metabolic process | 3110 | 116 | 81.73 | 1.42 | + | 1.06E-04 | 2.23E-02 |
| regulation of cellular catabolic process | 755 | 40 | 19.84 | 2.02 | + | 4.95E-05 | 1.33E-02 |
| **sulfur compound metabolic process** | **357** | **26** | **9.38** | **2.77** | **+** | **7.11E-06** | **2.71E-03** |
| **neutrophil degranulation** | **483** | **33** | **12.69** | **2.6** | **+** | **1.58E-06** | **1.03E-03** |
| neutrophil mediated immunity | 496 | 33 | 13.04 | 2.53 | + | 3.62E-06 | 1.83E-03 |
| myeloid leukocyte mediated immunity | 517 | 33 | 13.59 | 2.43 | + | 6.68E-06 | 2.61E-03 |
| immune effector process | 1056 | 54 | 27.75 | 1.95 | + | 5.76E-06 | 2.44E-03 |
| neutrophil activation involved in immune response | 487 | 33 | 12.8 | 2.58 | + | 3.04E-06 | 1.64E-03 |
| myeloid cell activation involved in immune response | 516 | 34 | 13.56 | 2.51 | + | 2.86E-06 | 1.59E-03 |
| leukocyte activation involved in immune response | 609 | 36 | 16 | 2.25 | + | 1.87E-05 | 6.37E-03 |
| cell activation | 1033 | 51 | 27.15 | 1.88 | + | 2.82E-05 | 8.81E-03 |
| cell activation involved in immune response | 613 | 36 | 16.11 | 2.23 | + | 1.98E-05 | 6.44E-03 |
| response to stimulus | 8443 | 265 | 221.89 | 1.19 | + | 2.62E-04 | 4.02E-02 |
| myeloid leukocyte activation | 567 | 35 | 14.9 | 2.35 | + | 8.70E-06 | 3.24E-03 |
| neutrophil activation | 495 | 34 | 13.01 | 2.61 | + | 9.80E-07 | 7.66E-04 |
| granulocyte activation | 499 | 34 | 13.11 | 2.59 | + | 1.16E-06 | 8.25E-04 |
| leukocyte degranulation | 505 | 33 | 13.27 | 2.49 | + | 4.60E-06 | 2.11E-03 |
| regulated exocytosis | 695 | 43 | 18.27 | 2.35 | + | 5.97E-07 | 5.49E-04 |
| exocytosis | 782 | 43 | 20.55 | 2.09 | + | 1.33E-05 | 4.73E-03 |
| secretion by cell | 977 | 46 | 25.68 | 1.79 | + | 2.26E-04 | 3.64E-02 |
| secretion | 1090 | 50 | 28.65 | 1.75 | + | 2.20E-04 | 3.58E-02 |
| **regulation of body fluid levels** | **490** | **28** | **12.88** | **2.17** | **+** | **2.03E-04** | **3.34E-02** |
| **carbohydrate derivative metabolic process** | **1049** | **51** | **27.57** | **1.85** | **+** | **4.76E-05** | **1.31E-02** |
| **regulation of locomotion** | **905** | **43** | **23.78** | **1.81** | **+** | **3.03E-04** | **4.42E-02** |
| **response to oxygen-containing compound** | **1484** | **63** | **39** | **1.62** | **+** | **2.97E-04** | **4.38E-02** |
| **response to organic substance** | **2877** | **107** | **75.61** | **1.42** | **+** | **2.79E-04** | **4.19E-02** |
| **Unclassified** | **3188** | **41** | **83.78** | **0.49** | **-** | **6.20E-08** | **7.46E-05** |
| **G-protein coupled receptor signaling pathway** | **1307** | **15** | **34.35** | **0.44** | **-** | **2.92E-04** | **4.35E-02** |
| **chromosome organization** | **1034** | **9** | **27.17** | **0.33** | **-** | **7.77E-05** | **1.76E-02** |
| **detection of chemical stimulus involved in sensory perception o** | **429** | **0** | **11.27** | **< 0.01** | **-** | **3.18E-05** | **9.37E-03** |
| detection of chemical stimulus involved in sensory perception | 477 | 0 | 12.54 | < 0.01 | - | 6.03E-06 | 2.42E-03 |
| sensory perception of chemical stimulus | 535 | 0 | 14.06 | < 0.01 | - | 1.24E-06 | 8.40E-04 |
| sensory perception | 951 | 9 | 24.99 | 0.36 | - | 3.48E-04 | 5.00E-02 |
| detection of chemical stimulus | 512 | 0 | 13.46 | < 0.01 | - | 2.78E-06 | 1.61E-03 |
| detection of stimulus | 683 | 2 | 17.95 | 0.11 | - | 5.88E-06 | 2.42E-03 |
| detection of stimulus involved in sensory perception | 531 | 0 | 13.96 | < 0.01 | - | 1.94E-06 | 1.21E-03 |
| sensory perception of smell | 458 | 0 | 12.04 | < 0.01 | - | 1.43E-05 | 4.96E-03 |

**Table E6. Over- or under-represented Gene Ontology biological process terms for the genes positively coexpressed with *DAAM1* exclusively in the primary visual cortex (VIC).** The gene list analysis was carried out using the analysis tool in the PANTHER classification system (PMID: 23868073), with GO annotation data (PMID: 27899567). The GO terms were organized *hierarchially*, where the most post specific terms, indicated by bolded texts, were listed before the more general terms. The background gene set contains all 21,042 genes available in PANTHER database. The test set contains the 409 protein-coding genes whose expression levels were positively correlated (r>=0.56) with DAAM1 expression levels in the primary visual cortex area (V1C), but not in the other cortical regions in the prenatal brains. We used r ≥0.56 as the cut-off for positive-correlation coefficient, which was taken to be the 99th percentile of the empirical distribution of the correlation coefficients between DAAM1 expression levels and the expression levels of all the other 60,153 coding and non-coding genes in neo-cortical regions of prenatal brains available in the BrainSpan database (http://www.brainspan.org/). The Pearson correlation coefficients were obtained using the vector of mRNA expression levels of each gene across all prenatal stages. For the regions that were not well parcellated during 8 – 10 post-conception weeks, their data were either merged (M1C and S1C) or used to represent the expression values in sub-regions (parietal cortex [PC] and temporal cortex [TC]).

| GO BP complete | N REF (21,042) | N Set (589) | expected | Fold Enrichment | over(+) or under(-) represenation | raw P | FDR |
|---|---|---|---|---|---|---|---|
| **cristae formation** | **31** | **9** | **0.87** | **10.37** | + | **9.96E-07** | **6.18E-04** |
| inner mitochondrial membrane organization | 41 | 9 | 1.15 | 7.84 | + | 7.16E-06 | 2.58E-03 |
| mitochondrial membrane organization | 129 | 16 | 3.61 | 4.43 | + | 2.25E-06 | 1.09E-03 |
| mitochondrion organization | 437 | 39 | 12.23 | 3.19 | + | 1.11E-09 | 1.72E-05 |
| organelle organization | 3182 | 128 | 89.07 | 1.44 | + | 2.73E-05 | 6.95E-03 |
| cellular component organization | 5273 | 196 | 147.6 | 1.33 | + | 1.15E-05 | 3.79E-03 |
| cellular process | 15084 | 467 | 422.23 | 1.11 | + | 3.43E-05 | 8.05E-03 |
| cellular component organization or biogenesis | 5498 | 207 | 153.9 | 1.35 | + | 1.98E-06 | 1.02E-03 |
| membrane organization | 836 | 46 | 23.4 | 1.97 | + | 2.88E-05 | 7.21E-03 |
| **mitochondrial ATP synthesis coupled proton transport** | **21** | **6** | **0.59** | **10.21** | + | **7.22E-05** | **1.51E-02** |
| mitochondrial transport | 243 | 20 | 6.8 | 2.94 | + | 3.83E-05 | 8.61E-03 |
| transport | 4324 | 159 | 121.04 | 1.31 | + | 2.44E-04 | 4.02E-02 |
| ATP synthesis coupled proton transport | 29 | 8 | 0.81 | 9.86 | + | 5.54E-06 | 2.15E-03 |
| ATP biosynthetic process | 41 | 9 | 1.15 | 7.84 | + | 7.16E-06 | 2.64E-03 |
| purine ribonucleotide biosynthetic process | 121 | 13 | 3.39 | 3.84 | + | 7.73E-05 | 1.58E-02 |
| ribonucleotide biosynthetic process | 134 | 15 | 3.75 | 4 | + | 1.43E-05 | 4.42E-03 |
| nucleotide biosynthetic process | 203 | 18 | 5.68 | 3.17 | + | 3.79E-05 | 8.64E-03 |
| nucleoside phosphate biosynthetic process | 205 | 18 | 5.74 | 3.14 | + | 4.27E-05 | 9.32E-03 |
| nucleoside phosphate metabolic process | 525 | 39 | 14.7 | 2.65 | + | 1.14E-07 | 1.26E-04 |
| organophosphate metabolic process | 990 | 52 | 27.71 | 1.88 | + | 3.31E-05 | 7.90E-03 |
| phosphorus metabolic process | 2199 | 100 | 61.55 | 1.62 | + | 2.45E-06 | 1.12E-03 |
| cellular metabolic process | 9069 | 306 | 253.86 | 1.21 | + | 2.39E-05 | 6.39E-03 |
| metabolic process | 9969 | 330 | 279.05 | 1.18 | + | 4.00E-05 | 8.86E-03 |
| phosphate-containing compound metabolic process | 2107 | 99 | 58.98 | 1.68 | + | 5.53E-07 | 3.73E-04 |
| nucleobase-containing small molecule metabolic process | 609 | 42 | 17.05 | 2.46 | + | 2.58E-07 | 2.35E-04 |
| primary metabolic process | 9162 | 303 | 256.46 | 1.18 | + | 1.48E-04 | 2.64E-02 |
| nucleotide metabolic process | 519 | 39 | 14.53 | 2.68 | + | 8.63E-08 | 1.03E-04 |
| ribose phosphate biosynthetic process | 138 | 15 | 3.86 | 3.88 | + | 1.96E-05 | 5.52E-03 |
| carbohydrate derivative metabolic process | 1094 | 54 | 30.62 | 1.76 | + | 1.13E-04 | 2.17E-02 |
| ribose phosphate metabolic process | 397 | 32 | 11.11 | 2.88 | + | 3.04E-07 | 2.48E-04 |
| ribonucleotide metabolic process | 381 | 31 | 10.66 | 2.91 | + | 3.86E-07 | 2.99E-04 |
| purine nucleotide biosynthetic process | 128 | 14 | 3.58 | 3.91 | + | 3.46E-05 | 8.01E-03 |
| purine-containing compound biosynthetic process | 141 | 14 | 3.95 | 3.55 | + | 9.05E-05 | 1.80E-02 |
| organonitrogen compound biosynthetic process | 1438 | 69 | 40.25 | 1.71 | + | 2.41E-05 | 6.33E-03 |
| organonitrogen compound metabolic process | 5534 | 222 | 154.91 | 1.43 | + | 2.81E-09 | 1.45E-05 |
| purine-containing compound metabolic process | 422 | 30 | 11.81 | 2.54 | + | 1.09E-05 | 3.67E-03 |
| purine nucleotide metabolic process | 385 | 30 | 10.78 | 2.78 | + | 1.37E-06 | 7.56E-04 |
| purine ribonucleotide metabolic process | 366 | 29 | 10.24 | 2.83 | + | 1.50E-06 | 8.03E-04 |
| purine ribonucleoside triphosphate biosynthetic process | 53 | 11 | 1.48 | 7.41 | + | 1.12E-06 | 6.66E-04 |
| purine ribonucleoside triphosphate metabolic process | 235 | 27 | 6.58 | 4.1 | + | 3.55E-09 | 1.38E-05 |
| ribonucleoside triphosphate metabolic process | 241 | 28 | 6.75 | 4.15 | + | 1.45E-09 | 1.12E-05 |
| nucleoside triphosphate metabolic process | 262 | 28 | 7.33 | 3.82 | + | 7.78E-09 | 1.51E-05 |
| purine nucleoside triphosphate metabolic process | 242 | 27 | 6.77 | 3.99 | + | 6.29E-09 | 1.39E-05 |
| purine nucleoside triphosphate biosynthetic process | 54 | 11 | 1.51 | 7.28 | + | 1.31E-06 | 7.54E-04 |
| nucleoside triphosphate biosynthetic process | 70 | 12 | 1.96 | 6.12 | + | 2.17E-06 | 1.08E-03 |
| ribonucleoside triphosphate biosynthetic process | 59 | 12 | 1.65 | 7.27 | + | 4.40E-07 | 3.25E-04 |
| ATP metabolic process | 204 | 25 | 5.71 | 4.38 | + | 4.17E-09 | 1.29E-05 |
| purine ribonucleoside monophosphate metabolic process | 246 | 26 | 6.89 | 3.78 | + | 3.28E-08 | 5.08E-05 |
| purine nucleoside monophosphate metabolic process | 247 | 26 | 6.91 | 3.76 | + | 3.53E-08 | 4.98E-05 |
| nucleoside monophosphate metabolic process | 274 | 28 | 7.67 | 3.65 | + | 1.89E-08 | 3.25E-05 |
| ribonucleoside monophosphate metabolic process | 259 | 28 | 7.25 | 3.86 | + | 6.19E-09 | 1.60E-05 |
| drug metabolic process | 615 | 36 | 17.21 | 2.09 | + | 6.68E-05 | 1.42E-02 |
| purine ribonucleoside monophosphate biosynthetic process | 64 | 10 | 1.79 | 5.58 | + | 3.11E-05 | 7.66E-03 |
| purine nucleoside monophosphate biosynthetic process | 64 | 10 | 1.79 | 5.58 | + | 3.11E-05 | 7.54E-03 |
| nucleoside monophosphate biosynthetic process | 87 | 12 | 2.44 | 4.93 | + | 1.59E-05 | 4.83E-03 |
| ribonucleoside monophosphate biosynthetic process | 77 | 12 | 2.16 | 5.57 | + | 5.22E-06 | 2.08E-03 |
| energy coupled proton transport, down electrochemical gradient | 29 | 8 | 0.81 | 9.86 | + | 5.54E-06 | 2.10E-03 |
| hydrogen ion transmembrane transport | 119 | 15 | 3.33 | 4.5 | + | 3.88E-06 | 1.58E-03 |
| transmembrane transport | 1225 | 64 | 34.29 | 1.87 | + | 3.41E-06 | 1.43E-03 |
| proton transport | 151 | 16 | 4.23 | 3.79 | + | 1.40E-05 | 4.42E-03 |
| monovalent inorganic cation transport | 420 | 26 | 11.76 | 2.21 | + | 3.03E-04 | 4.66E-02 |
| **protein peptidyl-prolyl isomerization** | **44** | **9** | **1.23** | **7.31** | + | **1.18E-05** | **3.81E-03** |
| peptidyl-proline modification | 59 | 9 | 1.65 | 5.45 | + | 9.12E-05 | 1.79E-02 |
| cellular protein modification process | 3164 | 123 | 88.57 | 1.39 | + | 1.87E-04 | 3.25E-02 |

| GO BP complete | N REF (21,042) | N Set (589) | expected | Fold Enrichment | over(+) or under(-) represenation | raw P | FDR |
|---|---|---|---|---|---|---|---|
| protein modification process | 3164 | 123 | 88.57 | 1.39 | + | 1.87E-04 | 3.22E-02 |
| protein metabolic process | 4466 | 174 | 125.01 | 1.39 | + | 3.32E-06 | 1.43E-03 |
| macromolecule modification | 3451 | 131 | 96.6 | 1.36 | + | 3.09E-04 | 4.65E-02 |
| cellular protein metabolic process | 3752 | 156 | 105.02 | 1.49 | + | 2.89E-07 | 2.49E-04 |
| **response to amine** | **40** | **7** | **1.12** | **6.25** | + | **2.62E-04** | **4.27E-02** |
| **mitochondrial electron transport, NADH to ubiquinone** | **49** | **8** | **1.37** | **5.83** | + | **1.47E-04** | **2.65E-02** |
| respiratory electron transport chain | 112 | 17 | 3.14 | 5.42 | + | 8.30E-08 | 1.07E-04 |
| cellular respiration | 167 | 19 | 4.67 | 4.06 | + | 8.48E-07 | 5.48E-04 |
| energy derivation by oxidation of organic compounds | 240 | 21 | 6.72 | 3.13 | + | 1.06E-05 | 3.64E-03 |
| generation of precursor metabolites and energy | 376 | 27 | 10.52 | 2.57 | + | 1.82E-05 | 5.32E-03 |
| electron transport chain | 182 | 19 | 5.09 | 3.73 | + | 2.73E-06 | 1.21E-03 |
| mitochondrial ATP synthesis coupled electron transport | 91 | 15 | 2.55 | 5.89 | + | 1.85E-07 | 1.92E-04 |
| ATP synthesis coupled electron transport | 92 | 15 | 2.58 | 5.82 | + | 2.10E-07 | 2.04E-04 |
| oxidative phosphorylation | 100 | 15 | 2.8 | 5.36 | + | 5.47E-07 | 3.86E-04 |
| phosphorylation | 1289 | 64 | 36.08 | 1.77 | + | 1.94E-05 | 5.57E-03 |
| **SCF-dependent proteasomal ubiquitin-dependent protein catabolic process** | **73** | **11** | **2.04** | **5.38** | + | **1.71E-05** | **5.09E-03** |
| proteasome-mediated ubiquitin-dependent protein catabolic process | 294 | 23 | 8.23 | 2.79 | + | 2.18E-05 | 5.93E-03 |
| proteasomal protein catabolic process | 319 | 24 | 8.93 | 2.69 | + | 2.63E-05 | 6.78E-03 |
| cellular protein catabolic process | 579 | 33 | 16.21 | 2.04 | + | 2.07E-04 | 3.52E-02 |
| ubiquitin-dependent protein catabolic process | 492 | 29 | 13.77 | 2.11 | + | 3.04E-04 | 4.63E-02 |
| modification-dependent protein catabolic process | 497 | 30 | 13.91 | 2.16 | + | 1.82E-04 | 3.21E-02 |
| modification-dependent macromolecule catabolic process | 506 | 31 | 14.16 | 2.19 | + | 1.19E-04 | 2.25E-02 |
| **mitochondrial respiratory chain complex assembly** | **96** | **13** | **2.69** | **4.84** | + | **8.48E-06** | **2.99E-03** |
| cellular protein complex assembly | 418 | 31 | 11.7 | 2.65 | + | 2.39E-06 | 1.12E-03 |
| protein complex subunit organization | 1321 | 61 | 36.98 | 1.65 | + | 2.14E-04 | 3.61E-02 |
| **positive regulation of ubiquitin-protein ligase activity involved in regulation of mitotic cell cycle transition** | **77** | **10** | **2.16** | **4.64** | + | **1.26E-04** | **2.33E-02** |
| positive regulation of ubiquitin protein ligase activity | 85 | 10 | 2.38 | 4.2 | + | 2.63E-04 | 4.25E-02 |
| regulation of ubiquitin-protein transferase activity | 125 | 13 | 3.5 | 3.72 | + | 1.04E-04 | 2.03E-02 |
| positive regulation of ubiquitin-protein transferase activity | 104 | 12 | 2.91 | 4.12 | + | 7.78E-05 | 1.57E-02 |
| **NIK/NF-kappaB signaling** | **86** | **10** | **2.41** | **4.15** | + | **2.87E-04** | **4.53E-02** |
| **negative regulation of ubiquitin-protein transferase activity** | **86** | **10** | **2.41** | **4.15** | + | **2.87E-04** | **4.49E-02** |
| **mitochondrial translational elongation** | **86** | **10** | **2.41** | **4.15** | + | **2.87E-04** | **4.44E-02** |
| translational elongation | 121 | 12 | 3.39 | 3.54 | + | 2.86E-04 | 4.57E-02 |
| **Wnt signaling pathway, planar cell polarity pathway** | **101** | **11** | **2.83** | **3.89** | + | **2.43E-04** | **4.05E-02** |
| **post-translational protein modification** | **440** | **28** | **12.32** | **2.27** | + | **1.25E-04** | **2.33E-02** |
| **detection of chemical stimulus involved in sensory perception of smell** | **429** | **1** | **12.01** | **0.08** | - | **1.38E-04** | **2.52E-02** |
| detection of chemical stimulus involved in sensory perception | 472 | 1 | 13.21 | 0.08 | - | 4.49E-05 | 9.67E-03 |
| sensory perception of chemical stimulus | 533 | 2 | 14.92 | 0.13 | - | 7.33E-05 | 1.52E-02 |
| detection of chemical stimulus | 507 | 1 | 14.19 | 0.07 | - | 2.17E-05 | 6.00E-03 |

**Table E7.Genomic loci with independent significant SNPs identified in meta analyses of GWAS of surface area-PC1 and -PC2, obtained by FUMA[1]**

| PC domension | Ind. SNP No | Genomic Locus | Genomic Locus start | Genomic Locus end | chr | pos | uniqID | rsID | p | nSNPs | nGWASSNPs |
|---|---|---|---|---|---|---|---|---|---|---|---|
| 1 | 1 | 1 | 9720496 | 9721289 | 1 | 9720526 | 1:9720526:A:G | rs58784124 | 2.854E-08 | 5 | 5 |
| 1 | 2 | 2 | 141637438 | 141847759 | 3 | 141720712 | 3:141720712:A:G | rs34186890 | 3.596E-08 | 115 | 95 |
| 1 | 3 | 3 | 17792249 | 18035250 | 4 | 17924734 | 4:17924734:C:T | rs11938781 | 2.244E-09 | 253 | 215 |
| 1 | 4 | 4 | 103001649 | 103387161 | 4 | 103188709 | 4:103188709:C:T | rs13107325 | 4.473E-08 | 5 | 5 |
| 1 | 5 | 4 | 103001649 | 103387161 | 4 | 103292422 | 4:103292422:C:T | rs13109272 | 2.458E-08 | 19 | 14 |
| 1 | 6 | 5 | 106009763 | 106009763 | 4 | 106009763 | 4:106009763:A:G | rs2301718 | 3.319E-09 | 1 | 1 |
| 1 | 7 | 6 | 108861264 | 109019323 | 6 | 108977663 | 6:108977663:C:T | rs9400239 | 1.172E-08 | 65 | 60 |
| 1 | 8 | 6 | 108861264 | 109019323 | 6 | 108995187 | 6:108995187:A:G | rs61192764 | 1.442E-08 | 12 | 9 |
| 1 | 9 | 7 | 126648460 | 127369230 | 6 | 126704795 | 6:126704795:C:T | rs9388490 | 1.953E-09 | 246 | 202 |
| 1 | 10 | 7 | 126648460 | 127369230 | 6 | 126792095 | 6:126792095:A:G | rs11759026 | 3.584E-20 | 4 | 4 |
| 1 | 11 | 7 | 126648460 | 127369230 | 6 | 126986996 | 6:126986996:A:G | rs1262476 | 8.103E-10 | 22 | 19 |
| 1 | 12 | 7 | 126648460 | 127369230 | 6 | 127204623 | 6:127204623:A:G | rs9375477 | 1.567E-10 | 3 | 2 |
| 1 | 13 | 7 | 126648460 | 127369230 | 6 | 127369230 | 6:127369230:C:T | rs139849708 | 4.531E-09 | 13 | 13 |
| 1 | 14 | 8 | 54909928 | 55015505 | 7 | 54966738 | 7:54966738:A:G | rs77126132 | 3.776E-09 | 24 | 24 |
| 1 | 15 | 9 | 155772027 | 155778152 | 7 | 155778152 | 7:155778152:A:G | rs75745313 | 1.797E-08 | 2 | 2 |
| 1 | 16 | 10 | 104591182 | 105175131 | 10 | 105012994 | 10:105012994:C:T | rs1628768 | 1.993E-09 | 45 | 38 |
| 1 | 17 | 11 | 17067849 | 17365209 | 11 | 17310391 | 11:17310391:C:T | rs55948574 | 3.959E-08 | 74 | 59 |
| 1 | 18 | 12 | 92317858 | 92558342 | 11 | 92355990 | 11:92355990:A:T | rs3911562 | 4.832E-08 | 85 | 67 |
| 1 | 19 | 13 | 66257355 | 66389968 | 12 | 66306441 | 12:66306441:A:C | rs12298541 | 1.995E-08 | 2 | 1 |
| 1 | 20 | 13 | 66257355 | 66389968 | 12 | 66325689 | 12:66325689:C:T | rs343086 | 2.534E-09 | 1 | 1 |
| 1 | 21 | 13 | 66257355 | 66389968 | 12 | 66367726 | 12:66367726:C:T | rs61921611 | 5.671E-10 | 5 | 3 |
| 1 | 22 | 13 | 66257355 | 66389968 | 12 | 66376091 | 12:66376091:C:T | rs7306710 | 7.556E-16 | 23 | 21 |
| 1 | 23 | 13 | 66257355 | 66389968 | 12 | 66386396 | 12:66386396:C:T | rs11175990 | 4.152E-08 | 6 | 6 |
| 1 | 24 | 13 | 66257355 | 66389968 | 12 | 66387621 | 12:66387621:A:T | rs10748027 | 4.565E-08 | 1 | 1 |
| 1 | 25 | 13 | 66257355 | 66389968 | 12 | 66389968 | 12:66389968:C:T | rs10400419 | 2.086E-11 | 4 | 3 |
| 1 | 26 | 14 | 68216239 | 68225876 | 12 | 68216239 | 12:68216239:A:T | rs35227403 | 2.306E-08 | 2 | 2 |
| 1 | 27 | 15 | 90081905 | 90208310 | 15 | 90081905 | 15:90081905:A:G | rs28792763 | 1.721E-08 | 58 | 53 |
| 1 | 28 | 15 | 90081905 | 90208310 | 15 | 90128223 | 15:90128223:A:C | rs893725 | 1.153E-10 | 144 | 122 |
| 1 | 29 | 16 | 43055579 | 43055579 | 17 | 43055579 | 17:43055579:A:C | rs11652522 | 1.585E-08 | 1 | 1 |
| 1 | 30 | 17 | 43399058 | 44874453 | 17 | 43399058 | 17:43399058:C:G | rs117642368 | 8.939E-12 | 2 | 2 |
| 1 | 31 | 17 | 43399058 | 44874453 | 17 | 43460891 | 17:43460891:A:G | rs61572747 | 2.479E-14 | 1 | 1 |
| 1 | 32 | 17 | 43399058 | 44874453 | 17 | 43512318 | 17:43512318:A:G | rs56168933 | 7.662E-28 | 3241 | 475 |
| 1 | 33 | 17 | 43399058 | 44874453 | 17 | 43798308 | 17:43798308:A:G | rs117615688 | 4.427E-08 | 2 | 1 |
| 1 | 34 | 17 | 43399058 | 44874453 | 17 | 43871147 | 17:43871147:A:G | rs12944712 | 4.993E-11 | 20 | 20 |
| 1 | 35 | 17 | 43399058 | 44874453 | 17 | 43901074 | 17:43901074:A:G | rs173365 | 3.538E-10 | 8 | 8 |
| 1 | 36 | 17 | 43399058 | 44874453 | 17 | 43915497 | 17:43915497:C:T | rs62054807 | 2.118E-11 | 2935 | 336 |
| 1 | 37 | 17 | 43399058 | 44874453 | 17 | 43992943 | 17:43992943:A:G | rs9899833 | 5.179E-09 | 9 | 1 |
| 1 | 38 | 17 | 43399058 | 44874453 | 17 | 44787312 | 17:44787312:C:T | rs1378358 | 9.548E-32 | 3102 | 409 |
| 1 | 39 | 17 | 43399058 | 44874453 | 17 | 44793283 | 17:44793283:C:T | rs17692129 | 1.529E-10 | 4 | 3 |
| 1 | 40 | 17 | 43399058 | 44874453 | 17 | 44865439 | 17:44865439:G:T | rs2074404 | 1.957E-17 | 1840 | 241 |
| 1 | 41 | 17 | 43399058 | 44874453 | 17 | 44866602 | 17:44866602:C:T | rs199497 | 3.822E-10 | 2 | 2 |
| 1 | 42 | 18 | 47060322 | 47145848 | 17 | 47090785 | 17:47090785:C:T | rs11079849 | 2.803E-10 | 36 | 30 |
| 2 | 43 | 19 | 18956404 | 18992466 | 1 | 18962095 | 1:18962095:C:T | rs1934057 | 2.776E-08 | 26 | 23 |
| 2 | 44 | 20 | 23473592 | 23543929 | 1 | 23541078 | 1:23541078:A:T | rs7519093 | 3.028E-08 | 6 | 5 |
| 2 | 45 | 21 | 26752992 | 26902694 | 1 | 26752992 | 1:26752992:C:T | rs4659441 | 2.43E-09 | 25 | 22 |
| 2 | 46 | 21 | 26752992 | 26902694 | 1 | 26900805 | 1:26900805:A:G | rs12121702 | 2.428E-08 | 28 | 25 |
| 2 | 47 | 22 | 113054659 | 113252614 | 1 | 113054659 | 1:113054659:C:T | rs351370 | 1.936E-08 | 1 | 1 |
| 2 | 48 | 22 | 113054659 | 113252614 | 1 | 113063125 | 1:113063125:A:G | rs910697 | 3.26E-10 | 70 | 57 |
| 2 | 49 | 22 | 113054659 | 113252614 | 1 | 113239478 | 1:113239478:C:T | rs2999158 | 2.132E-11 | 4 | 4 |
| 2 | 50 | 22 | 113054659 | 113252614 | 1 | 113248365 | 1:113248365:A:G | rs954679 | 4.384E-10 | 5 | 5 |
| 2 | 51 | 23 | 45130410 | 45175585 | 2 | 45138444 | 2:45138444:A:C | rs540395 | 3.796E-08 | 10 | 10 |
| 2 | 52 | 23 | 45130410 | 45175585 | 2 | 45167570 | 2:45167570:G:T | rs83995 | 1.492E-10 | 24 | 24 |

| PC domension | Ind. SNP No | Genomic Locus | Genomic Locus start | Genomic Locus end | chr | pos | uniqID | rsID | p | nSNPs | nGWASSNPs |
|---|---|---|---|---|---|---|---|---|---|---|---|
| 2 | 53 | 24 | 104646815 | 104828166 | 3 | 104658328 | 3:104658328:C:T | rs6783013 | 9.705E-10 | 52 | 43 |
| 2 | 54 | 24 | 104646815 | 104828166 | 3 | 104660608 | 3:104660608:C:G | rs13077489 | 4.577E-08 | 2 | 2 |
| 2 | 55 | 24 | 104646815 | 104828166 | 3 | 104669023 | 3:104669023:G:T | rs6789712 | 1.236E-08 | 72 | 67 |
| 2 | 56 | 24 | 104646815 | 104828166 | 3 | 104683753 | 3:104683753:C:T | rs9288795 | 1.715E-15 | 224 | 208 |
| 2 | 57 | 24 | 104646815 | 104828166 | 3 | 104724787 | 3:104724787:A:T | rs971550 | 2.537E-17 | 49 | 42 |
| 2 | 58 | 24 | 104646815 | 104828166 | 3 | 104738751 | 3:104738751:A:G | rs6437556 | 1.381E-10 | 103 | 93 |
| 2 | 59 | 24 | 104646815 | 104828166 | 3 | 104823350 | 3:104823350:A:G | rs2399023 | 9.739E-09 | 67 | 63 |
| 2 | 60 | 25 | 59823118 | 60465365 | 5 | 60030791 | 5:60030791:A:T | rs7381195 | 5.274E-10 | 62 | 57 |
| 2 | 61 | 25 | 59823118 | 60465365 | 5 | 60069057 | 5:60069057:C:T | rs13158665 | 3.201E-09 | 39 | 34 |
| 2 | 62 | 25 | 59823118 | 60465365 | 5 | 60139881 | 5:60139881:A:G | rs62372102 | 1.433E-09 | 178 | 148 |
| 2 | 63 | 26 | 126525715 | 127369230 | 6 | 126613946 | 6:126613946:C:T | rs76470478 | 3.427E-09 | 4 | 1 |
| 2 | 64 | 26 | 126525715 | 127369230 | 6 | 126717064 | 6:126717064:A:G | rs1591805 | 2.012E-09 | 240 | 200 |
| 2 | 65 | 26 | 126525715 | 127369230 | 6 | 126745633 | 6:126745633:A:C | rs139214174 | 2.38E-11 | 14 | 13 |
| 2 | 66 | 26 | 126525715 | 127369230 | 6 | 127083941 | 6:127083941:C:T | rs9385403 | 3.946E-15 | 24 | 23 |
| 2 | 67 | 27 | 18869552 | 18933411 | 7 | 18904400 | 7:18904400:C:G | rs12700001 | 1.329E-10 | 49 | 47 |
| 2 | 68 | 28 | 98147278 | 98279801 | 9 | 98147804 | 9:98147804:C:T | rs7860361 | 1.34E-08 | 19 | 17 |
| 2 | 69 | 28 | 98147278 | 98279801 | 9 | 98192383 | 9:98192383:A:T | rs28710957 | 1.823E-08 | 3 | 3 |
| 2 | 70 | 28 | 98147278 | 98279801 | 9 | 98213728 | 9:98213728:C:G | rs9632916 | 1.825E-10 | 36 | 29 |
| 2 | 71 | 29 | 102245664 | 102371946 | 10 | 102245664 | 10:102245664:C:T | rs149658356 | 4.852E-08 | 2 | 2 |
| 2 | 72 | 29 | 102245664 | 102371946 | 10 | 102366639 | 10:102366639:C:T | rs138740906 | 1.702E-09 | 4 | 3 |
| 2 | 73 | 30 | 30798255 | 31020084 | 11 | 30861472 | 11:30861472:C:G | rs808471 | 2.446E-11 | 160 | 139 |
| 2 | 74 | 30 | 30798255 | 31020084 | 11 | 31010455 | 11:31010455:A:G | rs9667150 | 2.175E-08 | 134 | 118 |
| 2 | 75 | 31 | 103985306 | 104041862 | 11 | 104012656 | 11:104012656:A:G | rs1681464 | 3.021E-10 | 31 | 26 |
| 2 | 76 | 32 | 79859456 | 80251200 | 13 | 80162555 | 13:80162555:C:T | rs9545142 | 2.388E-08 | 109 | 85 |
| 2 | 77 | 32 | 79859456 | 80251200 | 13 | 80182535 | 13:80182535:A:T | rs9545151 | 3.696E-12 | 13 | 12 |
| 2 | 78 | 32 | 79859456 | 80251200 | 13 | 80188702 | 13:80188702:C:T | rs9601250 | 1.705E-13 | 2 | 2 |
| 2 | 79 | 32 | 79859456 | 80251200 | 13 | 80191817 | 13:80191817:C:G | rs9545154 | 4.616E-21 | 26 | 23 |
| 2 | 79 | 32 | 79859456 | 80251200 | 13 | 80191817 | 13:80191817:C:G | rs9545154 | 4.616E-21 | 26 | 23 |
| 2 | 80 | 32 | 79859456 | 80251200 | 13 | 80236762 | 13:80236762:A:G | rs9318650 | 3.425E-13 | 44 | 40 |
| 2 | 81 | 33 | 59449138 | 59891607 | 14 | 59466003 | 14:59466003:A:G | rs79329625 | 5.735E-12 | 104 | 96 |
| 2 | 82 | 33 | 59449138 | 59891607 | 14 | 59577898 | 14:59577898:C:T | rs78570253 | 6.214E-15 | 96 | 88 |
| 2 | 83 | 33 | 59449138 | 59891607 | 14 | 59585932 | 14:59585932:A:C | rs4898978 | 1.746E-20 | 14 | 13 |
| 2 | 84 | 33 | 59449138 | 59891607 | 14 | 59609282 | 14:59609282:G:T | rs4258526 | 5.34E-10 | 3 | 3 |
| 2 | 85 | 33 | 59449138 | 59891607 | 14 | 59622767 | 14:59622767:A:C | rs76119478 | 9.026E-09 | 2 | 2 |
| 2 | 86 | 33 | 59449138 | 59891607 | 14 | 59625980 | 14:59625980:A:G | rs11158250 | 5.574E-18 | 2 | 1 |
| 2 | 87 | 33 | 59449138 | 59891607 | 14 | 59625997 | 14:59625997:A:G | rs73313052 | 2.363E-34 | 50 | 43 |
| 2 | 88 | 33 | 59449138 | 59891607 | 14 | 59629611 | 14:59629611:G:T | rs2053300 | 2.516E-08 | 2 | 2 |
| 2 | 89 | 33 | 59449138 | 59891607 | 14 | 59633677 | 14:59633677:A:G | rs4898980 | 4.463E-08 | 2 | 2 |
| 2 | 90 | 33 | 59449138 | 59891607 | 14 | 59648738 | 14:59648738:C:T | rs893516 | 1.064E-21 | 3 | 2 |
| 2 | 91 | 33 | 59449138 | 59891607 | 14 | 59664984 | 14:59664984:C:T | rs1252916 | 3.016E-08 | 2 | 2 |
| 2 | 92 | 33 | 59449138 | 59891607 | 14 | 59665934 | 14:59665934:C:T | rs8016570 | 1.022E-11 | 15 | 12 |
| 2 | 93 | 33 | 59449138 | 59891607 | 14 | 59683086 | 14:59683086:C:T | rs78445564 | 1.129E-10 | 4 | 4 |
| 2 | 94 | 33 | 59449138 | 59891607 | 14 | 59684859 | 14:59684859:A:G | rs74874233 | 4.449E-14 | 126 | 116 |
| 2 | 95 | 33 | 59449138 | 59891607 | 14 | 59694643 | 14:59694643:A:G | rs2757117 | 3.892E-10 | 1 | 1 |
| 2 | 96 | 33 | 59449138 | 59891607 | 14 | 59801714 | 14:59801714:A:G | rs61984497 | 3.18E-09 | 89 | 83 |
| 2 | 97 | 33 | 59449138 | 59891607 | 14 | 59815097 | 14:59815097:A:G | rs61984499 | 8.887E-10 | 150 | 131 |
| 2 | 98 | 34 | 52538040 | 52635000 | 16 | 52585440 | 16:52585440:C:T | rs3112578 | 3.841E-09 | 29 | 25 |
| 2 | 99 | 35 | 52448936 | 52471667 | 20 | 52466100 | 20:52466100:C:T | rs10432728 | 1.648E-08 | 29 | 19 |

[1]K. Watanabe, E. Taskesen, A. van Bochoven and D. Posthuma. Functional mapping and annotation of genetic associations with FUMA. Nat. Commun. 8:1826. (2017)
