## Supplemental Figures for "Planar cell polarity pathway and development of the human visual cortex"

Supplementary Figures

Figure E1. Loadings for the first two leading principal components (PC) of surface area measurements in 34 cortical regions: PC1 (panel A) and PC2 (panel B).

Figure E2. Loadings for the first two leading principal components (PC) of thickness measurements in 34 cortical regions: PC1 (panel A) and PC2 (panel B).

Figure E3. Manhattan plots for the meta-analyses of GWAS for surface area PC1 (panel A) and PC2 (panel B) scores in CHARGE and UKBB cohorts.

Figure E4. Forest plot of the top PC2 SNP rs73313052.

Figure E5. Manhattan plots for the meta-analyses of GWAS thickness PC1 (panel A) and PC2 (panel B) scores in CHARGE and UKBB cohorts.

Figure E6. *DAAM1* co-localizes with mitochondrial marker ATP5A in the primary visual cortical plate.

Figure E7. Associations between lateral geniculate nucleus (LGN) volume and surface area PC2 scores in 694 SYS major-homozygotes (GG) (panel A) and in the 206 carriers of minor allele of rs73313052 (GA or AA) (panel B).

Figure E8. Quantile-quantile plots of cortical surface area and thickness PC1 and PC2 genome-wide association study meta-analyses.

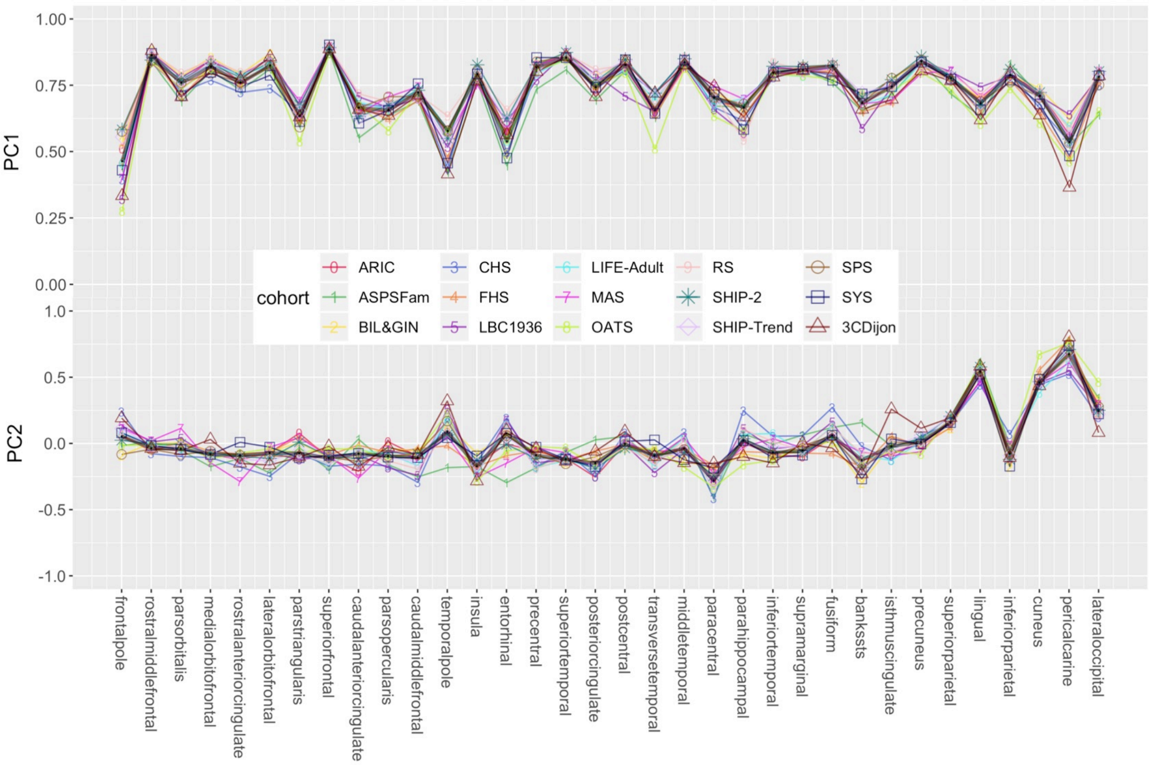


**Figure E1. Loadings for the first two leading principal components (PC) of surface area measurements in 34 cortical regions: PC1 (panel A) and PC2 (panel B).** Each cohort estimated the surface area of the 34 cortical regions (left and right hemispheres summed), using FreeSurfer and carried out principal component analyses to obtain PC loading values for the first two leading PCs, which are indicated by colored lines: 0-red (ARIC), 1-green (ASPSFam), 2-yellow (BIL&GIN), 3-blue (CHS), 4-orange (FHS), 5-purple (LBC1936), 6-cyan (LIFE-Adult), 7-magenta (MAS), 8-lime-green (OATS), 9-light pink (RS), dark green star (SHIP-2), light purple diamond (SHIP-Trend), brown circle (SPS), indigo rectangle (SYS), and dark red triangle (3CDijon) . Black thicker lines indicate the median loading values across the following cohorts: ARIC, ASPSFam, BIL&GIN, CHS, FHS, LIFE-Adult, RS, SPS and SYS. The median loading values were then used to derive the ‘general’ PC score for each individual and later used as the response variable in the meta-GWAS analyses. Cohort abbreviations are presented in Table E1A.

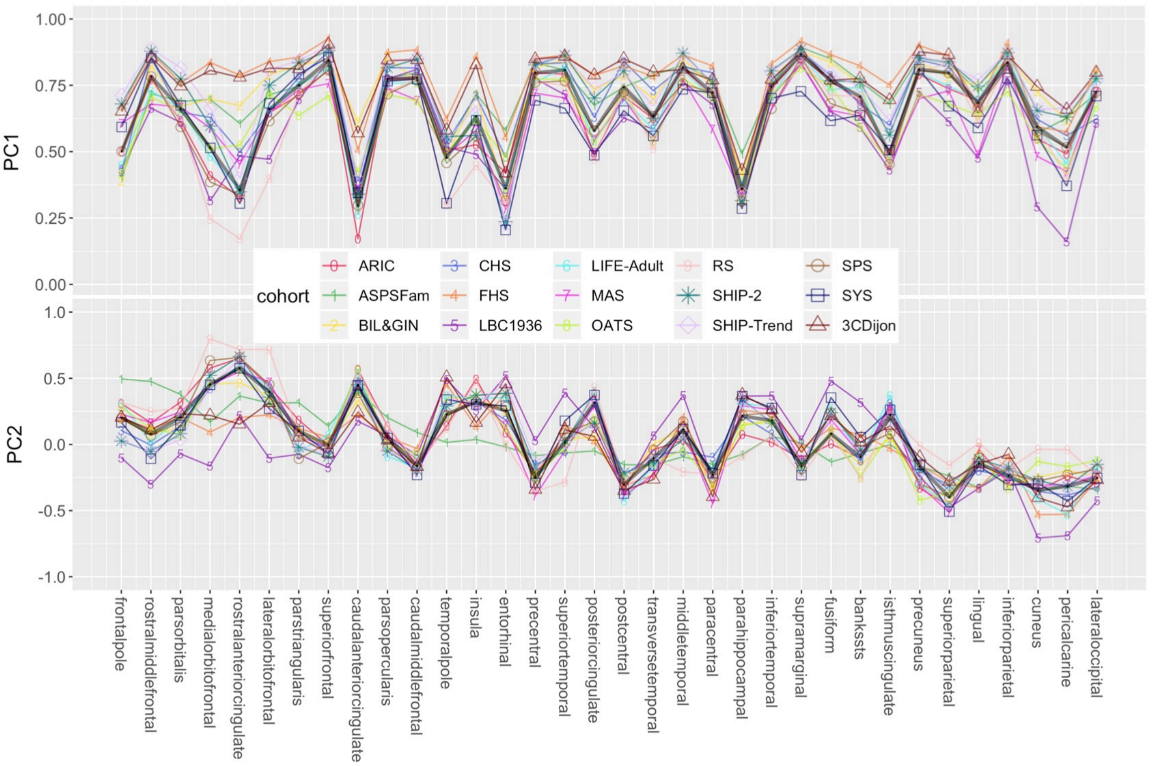


**Figure E2. Loadings for the first two leading principal components (PC) of thickness measurements in 34 cortical regions: PC1 (panel A) and PC2 (panel B).** Each cohort estimated the thickness of the 34 cortical regions (left and right hemispheres averaged), using FreeSurfer and carried out principal component analyses to obtain PC loading values for the first two leading PCs, which are indicated by colored lines: 0-red (ARIC), 1-green (ASPSFam), 2-yellow (BIL&GIN), 3-blue (CHS), 4-orange (FHS), 5-purple (LBC1936), 6-cyan (LIFE-Adult), 7-magenta (MAS), 8-lime-green (OATS), 9-light pink (RS), dark green star (SHIP-2), light purple diamond (SHIP-Trend), brown circle (SPS), indigo rectangle (SYS), and dark red triangle (3CDijon) . Black thicker lines indicate the median loading values across the following cohorts: ARIC, ASPSFam, BIL&GIN, CHS, FHS, LIFE-Adult, RS, SPS and SYS. The median loading values were then used to derive the 'general' PC score for each individual and later used as the response variable in the meta-GWAS analyses. Cohort abbreviations are presented in Table E1A.

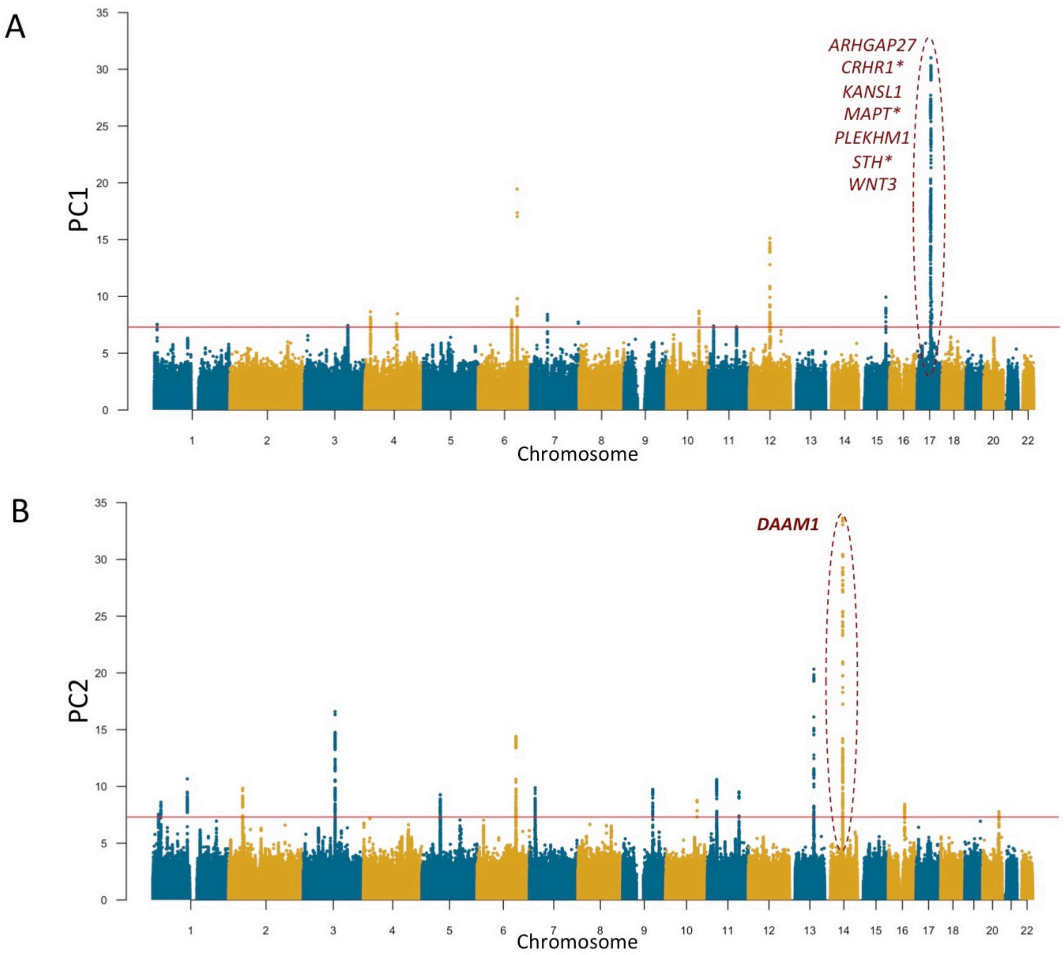


**Figure E3. Manhattan plots for the meta-analyses of GWAS for surface area PC1 (panel A) and PC2 (panel B) scores in CHARGE and UKBB cohorts.** The GWAS meta-analyses were carried out as follows. Using the median PC loadings across the CHARGE consortium cohorts (Figure E1), each cohort derived the 'general' PC scores to be used as phenotypes in the genome-wide association tests in order to ensure 'homogeneity' in phenotype derivation. All the association tests were adjusted for age, sex and other cohort-specific confounding variables, such as study site and/or family structure. The cohort-specific GWAS results were then examined for quality control with Easy QC software[1], and meta-analyzed with METAL[2] using fixed effects models. The vertical axis indicates the $-\log_{10} P$ values of the SNPs in the meta-analyses. The red dotted line indicates the genome-wide significance level of $5.0\times10^{-8}$. The most significant loci are highlighted with red-dashed ovals. The labelled genes within the PC1-GWAS-identified locus on chromosome 17 have been shown to be associated with intracranial volume and/or general cognitive function previously[3]. The *DAAM1* gene within the PC2-GWAS-identified locus on chromosome 14 contains multiple SNPs in moderate LD ($r^2 > 0.4$) with the top SNP rs73313052 ($p=2.4\times10^{-34}$).

[1]Winkler, T. W. *et al.* Quality control and conduct of genome-wide association meta-analyses. *Nat Protoc* 9, 1192–1212 (2014).
[2]Willer, C. J., Li, Y. & Abecasis, G. R. METAL: fast and efficient meta-analysis of genomewide association scans. *Bioinformatics* 26, 2190–2191 (2010).
[3]Trampush, J. W. *et al.* GWAS meta-analysis reveals novel loci and genetic correlates for general cognitive function: a report from the COGENT consortium. *Molecular Psychiatry* 22, 336–345 (2017).

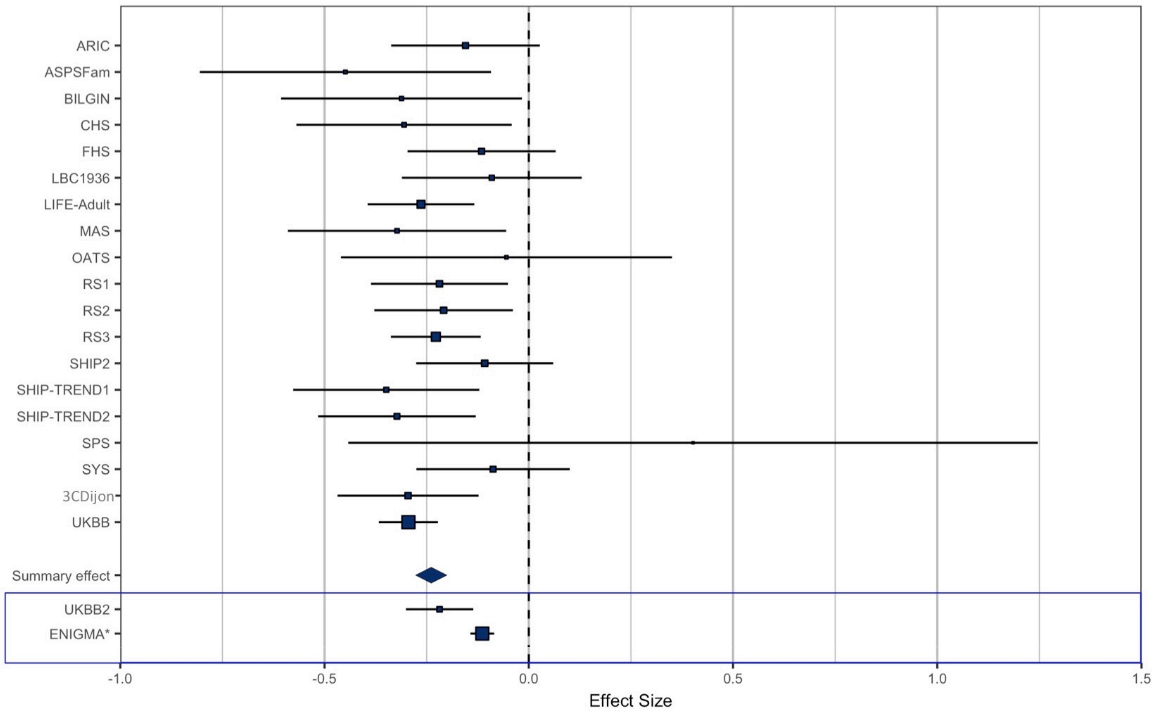


**Figure E4. Forest plot of the top PC2 SNP rs73313052.** Forest plots show the effect at each of the contributing cohort to the meta-analysis. The cohort-specific effect size estimates are indicated by the rectangles, and the meta-analysis effect size estimate, by a diamond; and their 95% confidence intervals (CI) are shown by the horizontal lines. The size of each point and width of CI lines are proportional to the sample size. The last lines indicate the effect size estimates and the 95% CIs obtained based on the 6,234 (UKBB2) individuals in UK Biobank cohort, who were not included in the meta-analysis, and 19,512 individuals included in the EINGMA meta-GWASs of surface area for 34 cortical regions. For ENIGMA analyses, the estimates and CIs were computed by using a method of genome-wide inferred study[1] with the GWAS summary statistics, means and covariances of cortical surface area for each of the 34 regions.

1.Nieuwboer, H. A., Pool, R., Dolan, C. V., Boomsma, D. I. & Nivard, M. G. GWIS: Genome-wide inferred statistics for functions of multiple phenotypes. *The American Journal of Human Genetics* 99, 917–927 (2016).

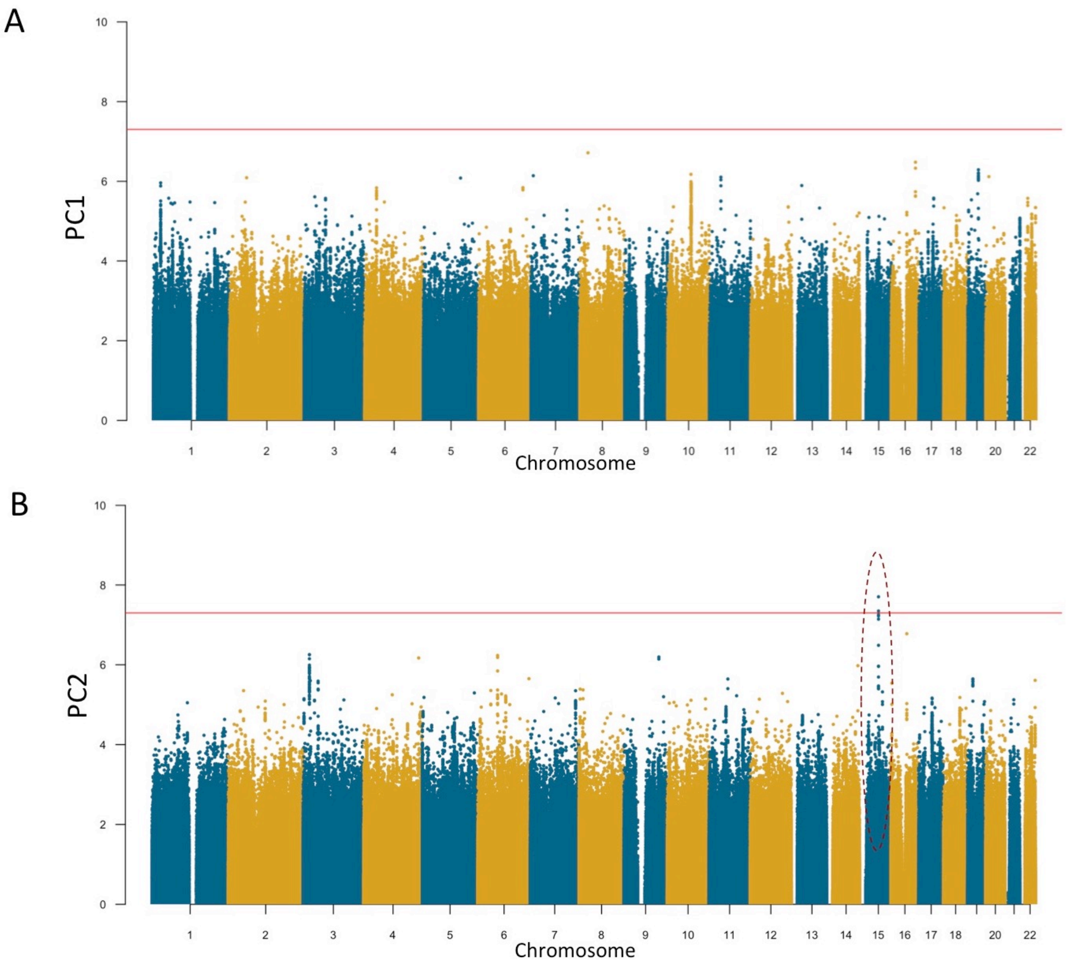


**Figure E5. Manhattan plots for the meta-analyses of GWAS thickness PC1 (panel A) and PC2 (panel B) scores in CHARGE and UKBB cohorts.** The GWAS meta-analyses were carried out as follows. Using the median PC loadings across the CHARGE consortium cohorts (Figure E2), each cohort derived the 'general' PC scores to be used as phenotypes in the genome-wide association tests in order to ensure 'homogeneity' in phenotype derivation. All the association tests were adjusted for age, sex and other cohort-specific confounding variables, such as study site and/or family structure. The cohort-specific GWAS results were then examined for quality-control with Easy QC software[1], and meta-analyzed with METAL[2] using fixed effects models. The vertical axis indicates the $-\log_{10} P$ values of the SNPs in the meta-analyses. The red dotted line indicates the genome-wide significance level of $5.0\times10^{-8}$. The most significant loci are highlighted with black-dashed eclipses. For PC2, there was no protein coding genes mapped to the GWAS-significant locus (rs12438460; $p=2.0\times10^{-8}$).

[1]Winkler, T. W. et al. Quality control and conduct of genome-wide association meta-analyses. *Nat. Protoc.* 9, 1192–1212 (2014).
[2]Willer, C. J., Li, Y. & Abecasis, G. R. METAL: fast and efficient meta-analysis of genomewide association scans. *Bioinformatics* 26, 2190–2191 (2010).

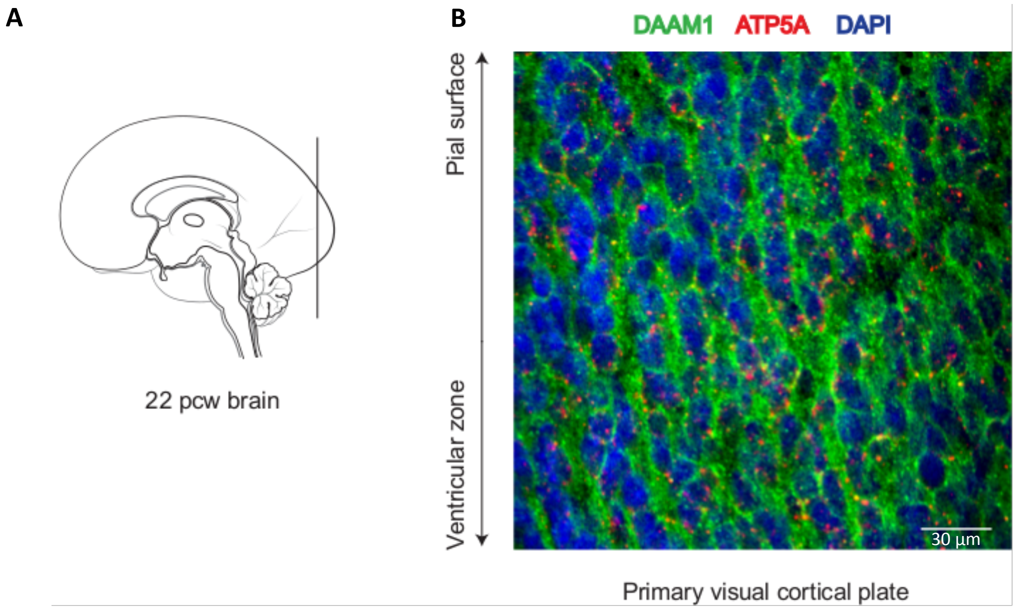


**Figure E6. *DAAM1* co-localizes with mitochondrial marker ATP5A in the primary visual cortical plate**. **A.** Schematic representation of the medial view of a midfetal (22 post-conception weeks) human brain; vertical line indicates the level at which coronal sectioning was performed for immunofluorescent experiments. **B.** Co-localization of DAAM1 and ATP5A in the human midfetal primary visual cortical plate. DAAM1 (green) can be observed in the cytoplasm of cells, whereas ATP5A (red) is localized in mitochondria, which are enriched around the nucleus (labeled with DAPI in blue) of cells. Scale bar represents 30 µm.

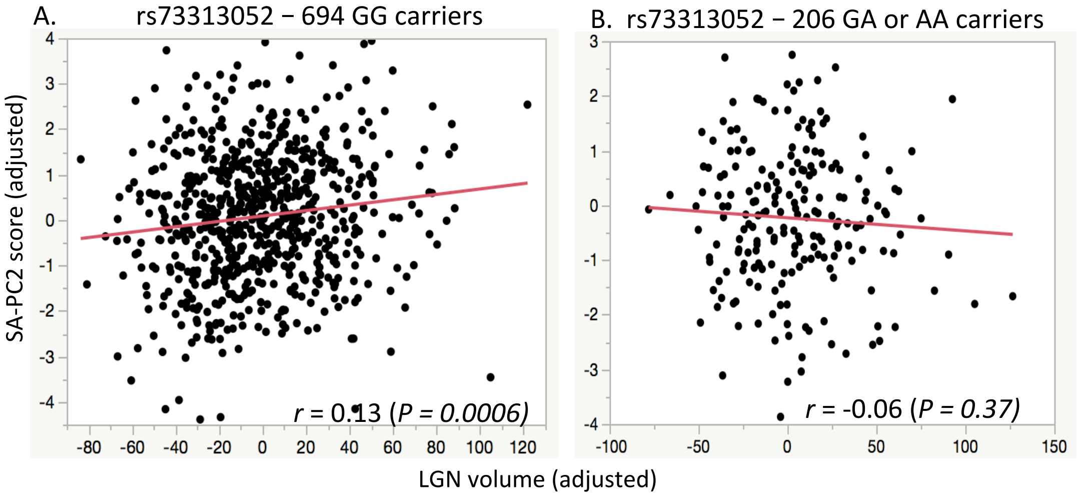


**Figure E7. Associations between lateral geniculate nucleus (LGN) volume and surface area PC2 scores in 694 SYS major-homozygotes (GG) (panel A) and in the 206 carriers of minor allele of rs73313052 (GA or AA) (panel B).** The horizontal and vertical axes indicate, respectively, LGN volume and surface area PC2 score, both adjusted for intracranial volume. The correlation coefficients between LGN and PC2 scores were 0.13 (*P*=0.005) and -0.06 (*P*=0.37), respectively, in GG-carriers and in minor-allele carriers, and t-test indicates that rs73313052-by-LGN interaction effect on PC2-scores is significant (t=-1.97; *P* <0.05) in 925 pooled individuals. The LGN volume was estimated as follows. First, the Jülich Histological atlas was used to obtain a cytoarchitectonic probabilistic map of the lateral geniculate nucleus (LGN) within MNI space[1]; the LGN mask was defined as voxels with a probability of P > 50%. Second, the thresholded LGN mask was non-linearly registered to a group template space (SYS808) using Advanced Normalization Tools (ANTs)[2]. The SYS808 template was created through a series of non-linear registrations of 808 T1-weighted images from the Saguenay Youth Study (SYS)[3]. Prior to registering LGN masks in SYS808 space to native subject space, individual T1-weighted scans are preprocessed by correcting for inhomogeneities using N3 (nonparametric non-uniformity normalization)[4] and skull stripping (Brain Extraction Tool)[5]. Third, the LGN mask from SYS808 template space was warped into individual space using diffeomorphic warping in ANTs[2], and mask volume was extracted using FSL toolkit[6]. ANTs diffeomorphic registration protocol was adapted from a previously reported methodology for extracting LGN Jacobian determinants[7]. Left LGN volumes were extracted for 925 participants from the SYS dataset (448 males, 494 females, average age 15.1 years)[3].

References

1. Bürgel, U. *et al.* White matter fiber tracts of the human brain: three-dimensional mapping at microscopic resolution, topography and intersubject variability. *Neuroimage* **29,** 1092–1105 (2006).

2. Avants, B. B., Tustison, N. & Song, G. Advanced normalization tools (ANTS). *Insight J* **2,** 1–35 (2009).

3. Paus, T. *et al.* Saguenay Youth Study: A multi-generational approach to studying virtual trajectories of the brain and cardio-metabolic health. *Dev. Cogn. Neurosci.* **11,** 129–144 (2015).
4. Sled, J. G., Zijdenbos, A. P. & Evans, A. C. A nonparametric method for automatic correction of intensity nonuniformity in MRI data. *IEEE Trans. Med. Imaging* **17,** 87–97 (1998).
5. Smith, S. M. Fast robust automated brain extraction. *Hum. Brain Mapp.* **17,** 143–155 (2002).
6. Jenkinson, M., Beckmann, C. F., Behrens, T. E., Woolrich, M. W. & Smith, S. M. Fsl. *Neuroimage* **62,** 782–790 (2012).
7. Aguirre, G. K. *et al.* Patterns of individual variation in visual pathway structure and function in the sighted and blind. *PloS One* **11,** e0164677 (2016).

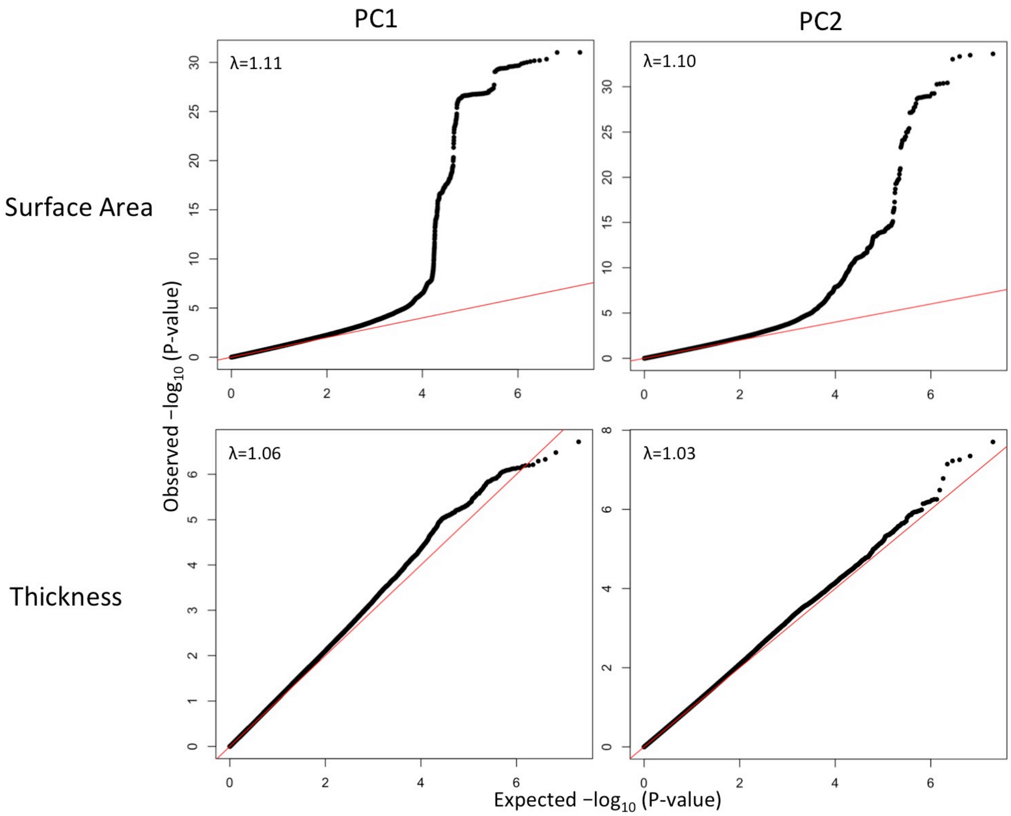


**Figure E8. Quantile-quantile plots of cortical surface area and thickness PC1 and PC2 genome-wide association study meta-analyses.** Observed -log10 transformed P-values of associations with cortical surface and thickness PC1 and PC2 are plotted against the expected values for all SNPs tested in each of the GWAS meta-analyses. Results were inspected for inflation due to confounders such as population stratification or sample overlap, by calculating the inflation factor (λ) as well as the LD score intercept using LD score regression[1]. An inflation factor $\lambda > 1$ suggests an inflation of the association statistics, which can be due to both spurious and genuine effects. It is likely to increase with sample size and degree of polygenicity of the phenotype, as the distribution of effect sizes begins to differ substantially from a null distribution when more variants have true associations. An LD score intercept > 1 suggests that there is spurious association, but an intercept <1.10 is generally considered to suggest that the signal is mostly due to genuine association effects. LD score regression intercepts were 1.05 (se=0.0079), 1.03 (se=0.0076), 1.03 (se=0.0065) and 1.02 (se= 0.007), for PC1-SA, PC2-SA, PC1-TH and PC2-TH, respectively.

1.Bulik-Sullivan, B. K. *et al.* LD Score regression distinguishes confounding from polygenicity in genome-wide association studies. *Nat. Genet.* 47, 291–295 (2015).
